# Phenotypic plasticity evolved for climate variability constrains performance under climate warming

**DOI:** 10.64898/2026.03.15.711905

**Authors:** Alayna Mead, Michelle Zavala-Paez, Joie R. Beasley-Bennett, Andrew C. Bleich, Ian P. Clancy-Mallue, Dylan G. Fischer, Julie M. Golightly, Kristina M. Hufford, Lee A. Kalcsits, Sara K. Klopf, Jesse R. Lasky, Jared M. LeBoldus, David B. Lowry, Nora Mitchell, Emily V. Moran, Jason P. Sexton, Kelsey L. Søndreli, Matthew C. Fitzpatrick, Jason Holliday, Stephen Keller, Jill A. Hamilton

**Affiliations:** Pennsylvania State University; Oregon State University; Michigan State University; Salisbury University; The Evergreen State College; University of Wyoming; Washington State University; Virginia Tech; University of Wisconsin - Eau Claire; University of California Merced; University of Maryland Center for Environmental Science; University of Vermont

**Keywords:** climate variability, ecophysiology, forest trees, genotype x environment, hybrid zone, phenology, phenotypic plasticity, provenance trial, stomata

## Abstract

Phenotypic plasticity allows plants to rapidly respond to changing environments without the need for evolutionary change or migration. While selection can create variation in plasticity across natural populations, these responses are not adaptive in all environments. To predict whether plasticity will be adaptive requires evaluation of its fitness effects across a range of environments, including novel ones. Here, we test how traits and their plasticity vary for genotypes collected across a natural hybrid zone between two tree species with contrasting climatic niches. Fast-growing *Populus trichocarpa* inhabits maritime environments with relatively warm and stable temperatures, while *P. balsamifera* inhabits continental environments with cold winters and large temperature variance throughout the year. We planted 44 clonally replicated genotypes into thirteen common gardens and measured vegetative phenology, leaf morphology, stomata morphology and conductance, and photochemistry. Overall, genotypes from colder, more continental environments exhibited higher plasticity. *P. balsamifera* ancestry was associated with increased plasticity in timing of fall phenology, stomatal conductance, and leaf mass per unit area. We assessed the effects of trait plasticity on fitness estimated as yearly growth across common gardens and found that the plasticity-fitness relationship was often garden-specific, indicating that the planting environment did not consistently mediate plasticity-fitness relationships. When the effects of trait plasticity on growth varied by garden temperature, higher plasticity generally had neutral or negative associations with growth in warmer environments. These results suggest that elevated plasticity evolved in a *P. balsamifera* genomic background as part of a climate generalist strategy to seasonal temperature variability, but that there is a trade-off between plasticity and growth in warmer environments. Consequently, less-plastic but warm-adapted *P. trichocarpa* genotypes are likely to have a fitness advantage under warming climates. These results demonstrate that plasticity may sometimes be maladaptive and will not be universally beneficial in a warming world.

## Introduction

For long-lived, sessile species, the rapid rate of climate change may outpace their ability to migrate or adapt to changing conditions (Aitken et al. 2008). However, phenotypic plasticity may ensure that individuals can adjust phenotypes in response to environmental change, limiting reliance on genetic or demographic responses (Nicotra et al. 2010). While phenotypic plasticity allows an individual to cope with changing environmental cues, plasticity itself can display heritable variation and evolve in response to selection (Scheiner 1993; Ghalambor et al. 2015; De Kort et al. 2020; Leung et al. 2020; Becker et al. 2022). Indeed, natural populations often show extensive variation in plasticity associated with adaptation to their native environments (Schlichting 1986; Ghalambor et al. 2007; Exposito-Alonso et al. 2018; De Kort et al. 2020; Cooper et al. 2022; Eisenring et al. 2022; Martínez-Sancho et al. 2025; Ramírez-Valiente et al. 2025). Theory predicts that phenotypic plasticity is selected for in variable environments where the fitness benefits exceed the costs of sensing and responding to environmental change, and where predictable environmental changes allow organisms to accurately anticipate changes and respond quickly (Scheiner 1993; DeWitt et al. 1998; Stotz et al. 2021; Volk et al. 2022). As a result, it has been hypothesized that more plastic genotypes will have an advantage under increasingly variable and unpredictable environments (Anderson et al. 2012b; Vázquez et al. 2017; De Kort et al. 2020; Jacob et al. 2024). However, it remains difficult to predict when or if the benefits of plasticity will outweigh its costs. Where plasticity has evolved as an adaptive response to local environments, climate change may generate environments outside the historic range of variation, leading to maladaptive plastic responses (Bonamour et al. 2019; Cooper et al. 2019; Schneider 2022). To predict whether phenotypic plasticity will be beneficial under future environments, its fitness effects must be evaluated across a range of historic and novel environmental conditions.

The fitness benefits of plasticity can be highly environmentally dependent (Murren et al. 2015). Within a fitness landscape, adaptive plasticity describes how genotypes can track optimal environment-phenotype combinations by adjusting trait expression (**Figure 1a**) (Richards et al. 2006; Chevin et al. 2010). However, costs and limits of plasticity can constrain reaction norms, preventing a single genotype from occupying fitness peaks in every environment. One form of constraint could arise from relationships between trait plasticity and trait values, as increased plasticity may limit a genotype’s ability to express optimal traits in certain environments (DeWitt et al. 1998). For example, genotypes from regions that experience predictable environmental variation may evolve a highly plastic, “generalist” strategy having high fitness in a wide range of environments, but with a trade-off of lower overall maximum fitness (Genotype C in **Figure 1**) (Richards et al. 2006; Napier et al. 2023). Conversely, genotypes that experience less environmental variation may specialize to a particular climate, expressing optimal trait values for a limited environmental range, but exhibit decreased fitness in environments outside of that narrow range (Genotype A in **Figure 1**). If such trade-offs in flexibility and efficiency exist, it is important to understand which strategy, highly plastic generalists or less plastic specialists, will be favored under increased climate variability combined with warming. Diverse genotypes spanning a continuum of plasticity and planted into multiple environments can be used to determine where phenotypic plasticity versus phenotypic stability are beneficial.

**Figure 1.**
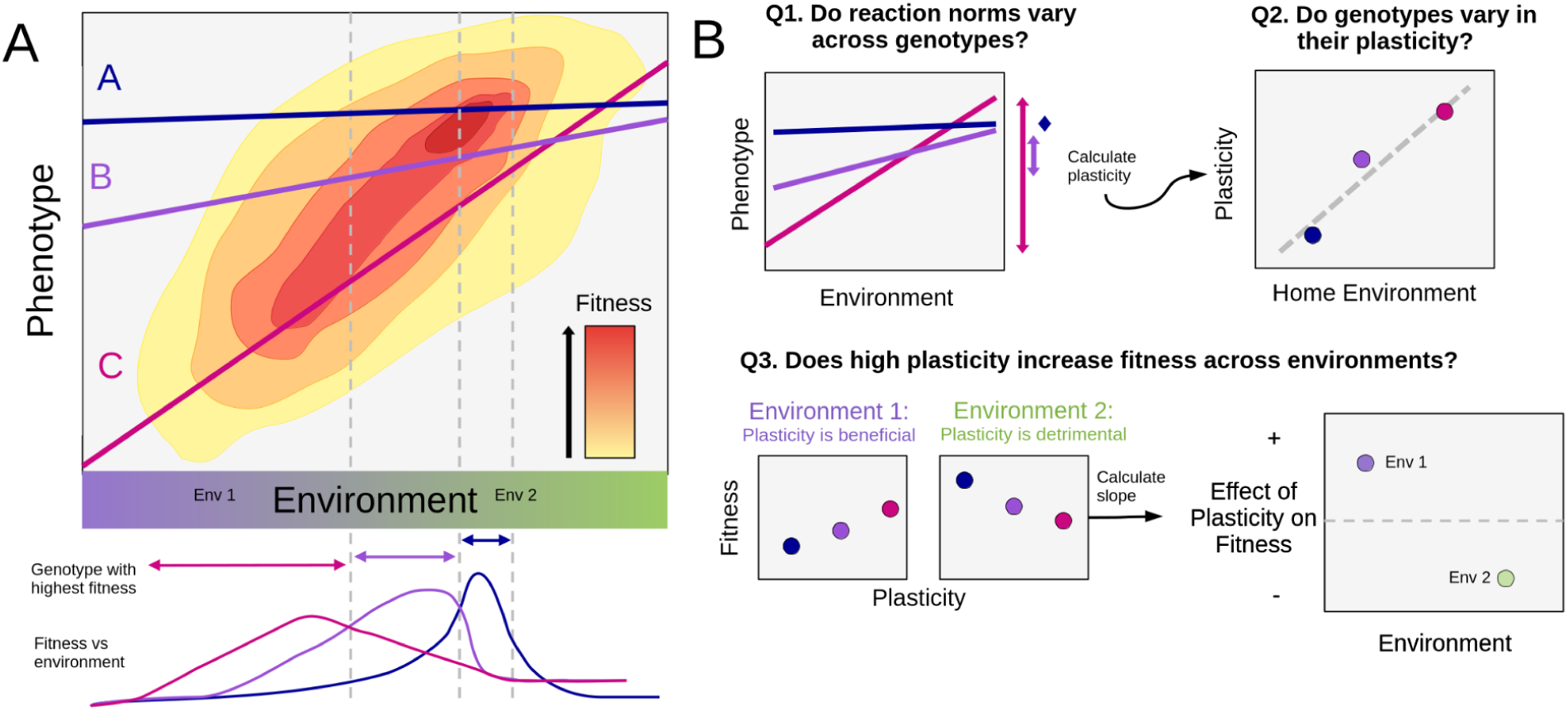
**A)** Hypothetical fitness landscape of trait values across an environmental gradient, in which the optimal trait value is correlated with the environment. Higher fitness is represented by darker red colors. Lines show reaction norms for three genotypes having varying plasticity, which causes variation in their relative fitness across environments. Genotype A (blue) exhibits a “specialist” strategy, expressing the same phenotype across all environments, but only outcompetes other genotypes in a narrow portion of the environmental gradient. Genotype C (pink) exhibits a “generalist” strategy, having high plasticity allowing it to track optimum fitness across a wide range of environments. Genotype B (purple) has intermediate plasticity and an intermediate environmental range. Grey dashed lines indicate environmental thresholds where the genotype having the highest fitness changes. **B)** describes the main questions of this study using the three hypothetical genotypes. First, we ask whether genotypes exhibit variation in their reaction norms. Second, we use the reaction norms to ask whether plasticity varies across genotypes as a function of source climatic or genetic factors. Third, we test whether plasticity is beneficial across different environments. In environment 1, genotype C has the highest fitness, so plasticity is beneficial. In environment 2, genotype A has an advantage, so the higher plasticity of genotypes B and C is detrimental. The relationship between plasticity and fitness in each environment is summarized by the slope and represented on the next plot, where it is plotted with the environmental variable to identify the environments where plasticity may be beneficial versus detrimental.

Where species with distinct climatic niches meet and hybridize, natural hybrid zones can provide dynamic arenas for testing how phenotypic plasticity translates to fitness across environmental gradients. If two species have evolved varying degrees of trait plasticity as adaptations to different levels of environmental variability, hybrids and backcrosses may exhibit a continuum of plasticity that spans, and potentially extends beyond, the range observed in parental species. Some studies have found that hybrids have greater plasticity (Marchal et al. 2019; Cooper and Shaffer 2021; Liu et al. 2021; Schwartz et al. 2024) or reduced vulnerability to climate change (Brauer et al. 2023; Hord et al. 2025), possibly as an outcome of heterosis in F1s or transgressive segregation in advanced-generation hybrids (Hamilton and Miller 2016). Under warming and increasingly variable climates, a higher optimum growth temperature coupled with increased plasticity to tolerate climatic extremes may be needed. Hybridization between generalists adapted to environmental uncertainty but colder average temperatures and specialists adapted to warmer environments could integrate new allelic combinations that give rise to genotypes uniquely suited to a warm-but-variable future climate. However, if there are inherent trade-offs between plasticity and productive growth in warm environments, hybridization and introgression may be unable to produce optimal phenotypes for the future environment and, instead, the warm-adapted species or genotype might replace the other.

Broadly, adaptive plasticity in response to climate change in plants may involve the ability to take advantage of extended growing seasons and accumulate growth under warmer temperatures, while avoiding stresses associated with heat, water limitation, and increased environmental variability. However, because traits vary in their degree of plasticity, their responsiveness to environmental cues, and potential trade-offs, the overall fitness effects of phenotypic plasticity likely vary across trait classes (Bradshaw 1965; DeWitt et al. 1998; Valladares et al. 2006). Here, we focus on four classes of traits that influence fitness across environments: phenology, leaf morphology, stomata morphology and conductance, and photochemistry (Nicotra et al. 2010; Anderson et al. 2012b). Phenological plasticity is common in plants (Anderson et al. 2012a; Vitasse et al. 2013; Cooper et al. 2019; Piao et al. 2019) and may be adaptive when it allows plants to take advantage of an extended growing season with earlier budburst and later budset, while avoiding frost damage during seasonal transitions (Cooper et al. 2019). However, increasing climate variability may result in unreliable environmental cues that trigger maladaptive responses (Bonamour et al. 2019; Chamberlain et al. 2019) which could be avoided by less environmentally-sensitive genotypes. Multiple traits impacting photosynthesis and the rate of carbon fixation respond to the environment. Leaf morphology can determine light capture, carbon diffusion, and water loss, and changes in photosynthetic machinery can alter photosynthetic capacity and temperature optima (Daas et al. 2008; Gunderson et al. 2010; Baird et al. 2017; Zhu et al. 2018; dos Santos and Ferreira 2020). Leaf morphological plasticity can also facilitate rapid environmental responses, such as rapid stomatal closure and opening (Drake et al. 2013). While variation in the responses of these traits to the environment is generally well-characterized, their phenotypic plasticity is often measured in only a handful of environments, and rarely across environmental extremes. Quantifying plasticity across continuous climate gradients is particularly informative for identifying the environments where higher plasticity is advantageous and predicting which responses will remain beneficial under future climates (Arnold et al. 2019).

We used clonally propagated *Populus* genotypes sampled from natural hybrid zones spanning a range of species ancestry proportions and environments to test how genetic variation, home climate, and planting environment interact to determine trait expression, ultimately testing the benefits and limitations of plasticity under changing environments. *Populus* genotypes can be clonally propagated, allowing environmental effects to be isolated from genetic effects. Here we focus on *Populus trichocarpa* Torr & A. Gray and *P. balsamifera* L., which inhabit contrasting environments and hybridize where their ranges come into contact, resulting in advanced-generation hybrids and backcrosses (Suarez-Gonzalez et al. 2016, 2018; Bolte et al. 2024; Mead et al. 2026). *Populus trichocarpa* occurs along the coast of western North America in warmer, wetter, maritime environments that experience less seasonal temperature fluctuation, while *P. balsamifera’s* range extends across the boreal regions of North America, encompassing colder, drier, continental climates (**Figure 2, Figure S1, S2**) (Suarez-Gonzalez et al. 2018; Bolte et al. 2024). In common gardens, *P. trichocarpa* exhibits faster aboveground growth in warm environments, but reduced growth and survival in cold environments when compared to *P. balsamifera* (Mead et al. 2026). Between the two species and among populations within species, functional traits display phenotypic clines consistent with adaptation to local climate variables (Gornall and Guy 2007; Soolanayakanahally et al. 2009, 2013; McKown et al. 2014a,b, 2018; Oubida et al. 2015; Fetter and Keller 2023). Continuous variation in species ancestry alongside substantial phenotypic variation allows us to test the effects of genetic variation, genomic ancestry, and home climate on traits and their plasticity.

**Figure 2.**
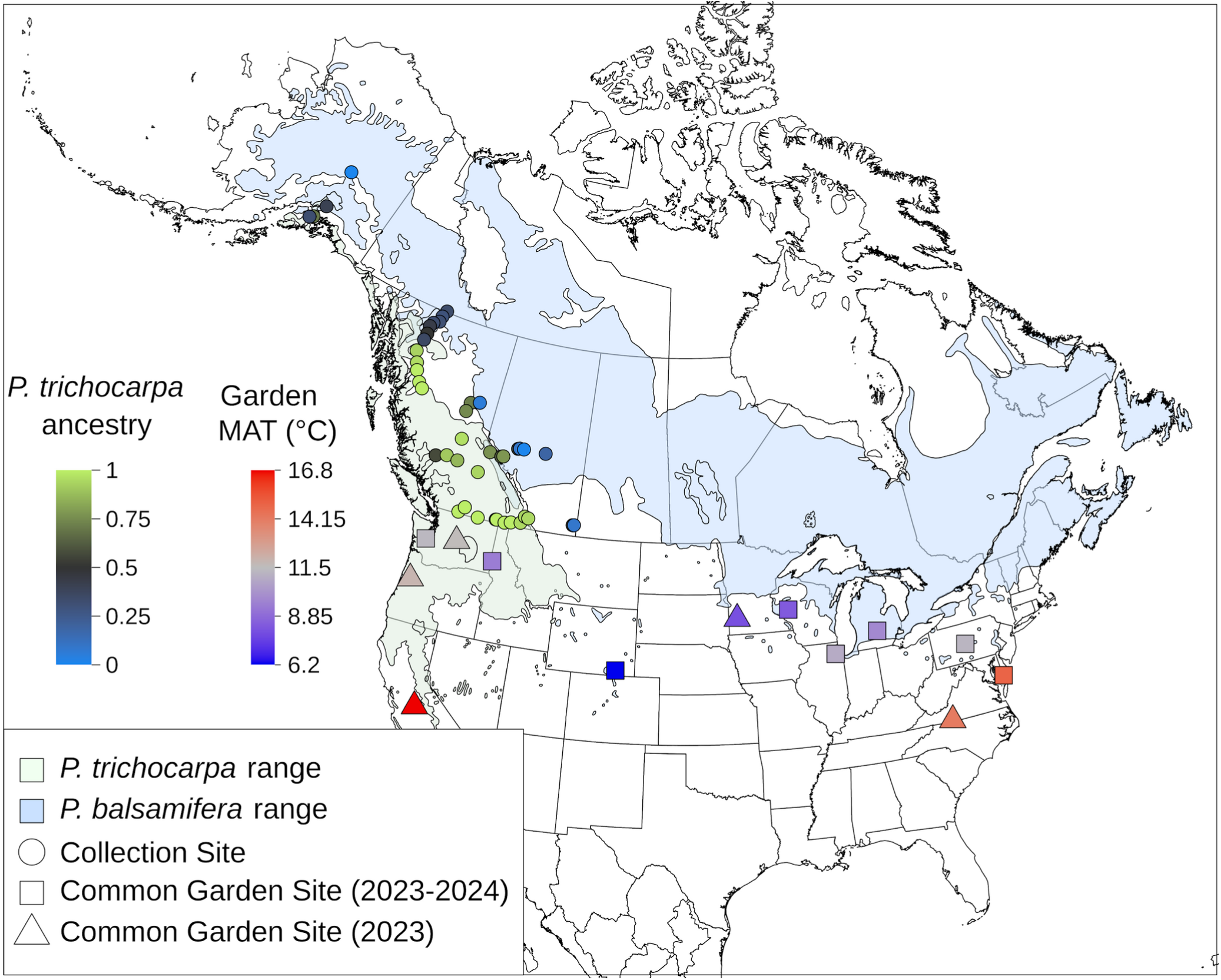
Location of sampled genotypes (circles) and 13 common garden sites (squares if measured in both years, triangles if measured only in 2023). Color of garden sites indicates their mean annual temperature (MAT) in 2023. Color of collection sites indicates species ancestry, determined using genomic data (Bolte et al. 2024; Mead et al. 2026). The light green and blue shading indicates the species ranges of *P. trichocarpa* and *P. balsamifera*, respectively (Little 1971).

Our experiment included 44 hybrid and parental poplar genotypes planted across a continuum of thirteen common garden environments, including novel warmer environments, to address the following questions: **1. Do trait responses to varying environments differ across genotypes based on their home climate, species ancestry, and degree of hybridization?** Genotypes may evolve variation in reaction norms (genotype × environment interactions, G×E) in response to differential selection across the hybrid zone. If we observe G×E effects on traits, selection from future environments could act on standing variation present in existing populations. **2. To what extent do genetic variation and home climate predict plasticity?** Plasticity could vary as a result of neutral genetic structure, environmental selection, or both. If we observe higher plasticity in genotypes originating from more variable or extreme climates, selection may be acting on plasticity itself. If hybrid genotypes have greater plasticity, this could suggest that novel genomic architectures contribute to plasticity. Additionally, if plasticity is associated with species ancestry, this would suggest divergent evolutionary pressures between the species. **3. Under which environments are trait values and their plasticity adaptive?** Plasticity can result in trade-offs with other phenotypic traits, having beneficial, deleterious, or neutral effects depending on the trait and environment. We focus on evaluating whether increased plasticity is beneficial in warming climates. Despite its potential importance, the fitness effects of plasticity are rarely tested in novel climates. By examining how plastic trait responses affect fitness across environments, we can predict whether natural populations may harbor beneficial variation in plasticity, enabling them to rapidly respond to climate change.

## Methods

### Common Gardens

Genotypes were collected as dormant vegetative cuttings from mature, wild trees growing across the natural hybrid zone between *P. trichocarpa* and *P. balsamifera* in western North America (**Figure 2**) between October 2019 and March 2020. Cuttings were clonally propagated at Virginia Tech (Critz, VA, USA), and planted into common gardens across the United States spanning a range of environments (**Figure S1**) in the fall of 2020, as described previously (Mead et al. 2026). Thirteen gardens survived the following two years of growth and are used here for all analyses (**Figure 2, Table S1**). Gardens in the driest environments (SWMN and UCM) were irrigated to ensure survival. All remaining trees in the UCM garden died in late spring 2023 due to irrigation failure during a heatwave, so only spring phenology traits were measured. Each garden included two blocks, with each genotype replicated in each block. In total, the experiment includes 44 genotypes × 13 gardens × 2 blocks/garden × 1 genotype/block.

### Phenotypic trait measurements

Here, we focus on phenotypic traits measured in 2023 and 2024, when trees were 3 and 4 years old, respectively. A list of traits and their abbreviations are provided in **Table 1**. Because phenotyping capacity varied by garden, not all traits were measured in all gardens; a list of traits measured in each garden is given in **Table S1**. All traits included here were measured during the summer of 2023, and an additional year of phenology and growth measures were recorded in 2024. Height was measured each year in the spring before budburst, and in fall following budset. Each year’s growth increment was calculated as the difference between the two height measurements. Survival was recorded at the end of each growing season.

**Table 1.** Phenotypic traits measured, their abbreviations, units, and which years they were measured.

| Trait | Abbreviation | Units | Year(s) measured |
| --- | --- | --- | --- |
| *Phenology* |  |  |  |
| Bud Flush | Stage 2 | Day of year | 2023-2024 |
| Leaf Emergence | Stage 3 | Day of year | 2023-2024 |
| Bud Flush cGDD | Stage 2 cGDD | Cumulative growing-degree days | 2023-2024 |
| Leaf Emergence cGDD | Stage 3 cGDD | Cumulative growing-degree days | 2023-2024 |
| First Budset | Stage 6 | Day of year | 2023-2024 |
| Lammas Growth | Stage 7 | Day of year | 2023-2024 |
| Final Budset | Stage 8 | Day of year | 2023-2024 |
| Growing Season Days |  | Days | 2023-2024 |
| *Leaf Morphology* |  |  |  |
| Leaf Area |  | cm<sup>2</sup> | 2023 |
| Leaf Mass |  | g | 2023 |
| Leaf Mass per Area | LMA | g/m<sup>2</sup> | 2023 |
| Leaf Thickness |  | mm (average of 5 leaves) | 2023 |
| *Stomata morphology and conductance* |  |  |  |
| Abaxial (lower) pore length | P<sub>L</sub> | μm | 2023 |
| Adaxial (upper) pore length | P<sub>U</sub> | μm | 2023 |
| Abaxial (lower) density | D<sub>L</sub> | mm<sup>-2</sup> | 2023 |
| Adaxial (upper) density | D<sub>U</sub> | mm<sup>-2</sup> | 2023 |
| Adaxial (upper) stomata presence |  | presence/absence | 2023 |
| Stomata ratio | SR | unitless | 2023 |
| Stomatal conductance | g<sub>sw</sub> | mol m<sup>-2</sup> s<sup>-1</sup> | 2023 |
| *Photochemistry and conductance* |  |  |  |
| Electron transport rate | ETR | mol m<sup>-2</sup> s<sup>-1</sup> | 2023 |
| Maximum fluorescence in light | Fm' | unitless | 2023 |
| Minimum fluorescence in light | Fs | unitless | 2023 |
| One-sided boundary layer conductance | g<sub>bw</sub> | mol m<sup>-2</sup> s<sup>-1</sup> | 2023 |
| Quantum efficiency in light | ΦPS2 | unitless | 2023 |

### Phenology

Phenological stages were measured in 2023 and 2024 using methods modified from Soolanayakanahally et al. (2013). We defined eight phenology stages: Stage 0 (dormant bud), Stage 1 (swollen bud with green tips), Stage 2 (bud flush), Stage 3 (leaf emergence), Stage 4 (active growth of terminal bud on main stem), Stage 5 (growth cessation, terminal bud beginning to form), Stage 6 (first time terminal bud is observed), Stage 7 (lammas growth, or regrowth from a terminal bud), and Stage 8 (last time bud is set). Dates were recorded for stages 2, 3, 6, 7, and 8. Gardens were monitored twice a week during spring and fall, recording the date that each individual reached a new stage. Because spring phenology is primarily driven by the accumulated heat sum (Soolanayakanahally et al. 2013), we calculated the number of growing-degree days at which stages 2 and 3 occurred. Cumulative growing degree days were calculated by extracting daily minimum and maximum temperatures from PRISM data (PRISM Group 2014), averaging minimum and maximum values for each day starting with January 1, and summing averages above 0°C. Fall phenology timing in *Populus* is thought to be controlled by the combined effect of photoperiod and temperature (Soolanayakanahally et al. 2013; McKown et al. 2018). We found that the daylength at which budset and lammas growth events occurred varied among clonal genotypes growing in different gardens, suggesting that temperature and photoperiod act together to determine fall phenology timing, so here we focus on the effect of garden climate to predict climate change responses.

### Leaf morphology traits

We measured a range of aboveground phenotypic traits across the thirteen gardens during the summer of 2023. Leaf area, mass, and thickness were measured on the same day within each garden, when trees reached phenological stage 4 (active growth). For each individual tree, the first fully expanded neoformed leaf on the dominant shoot was collected to measure leaf mass and area. If the dominant shoot was not producing leaves, we selected a leaf at a similar stage from the largest nondominant stem. Leaves were placed on ice and shipped overnight in a cooler with an ice pack to the Schatz Center for Tree Molecular Genetics in State College, PA. Following arrival, leaves were scanned using an Epson Perfection V39 scanner then placed in a drying oven for at least 24 h until constant mass was reached. The area of each leaf was calculated from leaf scans using ImageJ (Schneider et al. 2012). Leaf mass per area (LMA) was calculated as the dry mass divided by area. Leaf thickness was measured in the field using digital calipers for five fully expanded, sun exposed neoformed leaves, one of which was the same leaf collected for LMA measurements.

Stomatal conductance and photochemistry were measured using a LI-COR 600 Porometer/Fluorometer (LI-COR, inc., Lincoln, NE). For each individual tree, we averaged three replicate measures taken from the center of the first fully expanded leaf (avoiding the midrib and major veins) between 8 AM and 11 AM, before stomatal closure. LI-COR measurements were taken within the same week for each garden, and under similar weather conditions when measurements occurred on multiple days. Measurements across gardens were taken between May 1 and July 12 to facilitate equipment sharing among sites. Following conductance measures, the same leaf was sampled for stomatal morphology measurements. Leaves collected in the field were kept on ice before taking stomatal impressions the same day. We took stomatal impressions by applying New-Skin liquid bandage (Advantice Health, LLC. Bridgewater, NJ) on the upper (adaxial) and lower (abaxial) leaf surfaces, removing the dried impression using double-sided tape, and mounting it to a microscope slide. Stomatal size and density were measured using the methods of Zavala-Paez et al. (2026); see Supplementary Methods for details.

### Climate variables

Climate data was extracted from ClimateNA rasters with 1 km resolution (Wang et al. 2016; AdaptWest Project 2022) using terra 1.8-21 (Hijmans 2025) in R version 4.4.2 (R Core Team 2024) for genotype provenances (locations of origin) and common garden environments, as described by Mead et al. (2026). Historic climate averages for the 30 year period between 1961-1990 were extracted for the genotype collection sites (hereafter ‘home climate’), representing the environments that have likely exerted selective pressures on the mature trees. For each garden, we extracted the climate associated with years of data collection (2023-2024). To visualize how garden sites compare to predicted future climate conditions of natural populations, we used predictions from ClimateNA for the period of 2041-2070 under shared socioeconomic pathway (SSP) 2-4.5, estimated from an ensemble containing 13 general circulation models (GCMs) from the CMIP6 database. To visualize multivariate climatic variation across home and test sites, a PCA of historic and future home climates alongside garden climates over the years of measurement was created using the rda function in the R package vegan (Oksanen et al. 2024) with no conditioning or constraining variables.

Phenotypic variation should arise from the integration of environmental cues experienced in the planting environment and the genotype’s home climate. Previous results have found that winter temperatures and continentality are important factors affecting local adaptation in this hybrid zone (Bolte et al. 2024; Mead et al. 2026); however, plastic traits may be shaped by climate experienced throughout the year, including the growing season. To account for climate as an integrated measure of environmental conditions across seasons, we focused on mean annual temperature (MAT) and mean annual precipitation (MAP), because they are correlated with other temperature and precipitation variables, including seasonal variables, and had the highest loadings on PC1 and PC2 of the climate PCA (**Figure S1**). We also included continentality (temperature difference between mean warmest month and mean coldest month, TD) to test the effects of climatic variability on plasticity, although it is weakly negatively correlated with both MAT and MAP.

### Genetic variables

We used genomic data to evaluate how genetic structure shaped trait responses across environments. Whole-genome sequencing data was used to calculate species ancestry and quantify genetic structure using a principal component analysis (PC) of 334,657 SNPs, as described in previous studies (Bolte et al. 2024; Mead et al. 2026). To evaluate the effects of genomic structure on traits and their plasticity, we used the first three principal component (PC) axes. Genetic PC1 represents species ancestry, with higher values indicating greater *P. balsamifera* ancestry. Genetic PC2 separates *P. balsamifera* and two *P. trichocarpa* lineages from coastal and interior parts of the range. Genetic PC3 separates genotypes from two northern transects and three southern transects, with higher values indicating a northern origin.

### Plasticity Metric

Plasticity was calculated for each genotype using the relative distance plasticity index (RDPI) as described in Valladares et al. (2006), which captures variation in phenotypic values across multiple environments, with values ranging from 0 (no plasticity) to 1 (maximal plasticity). This method calculates the relative distance *(d)* between each pair of individual trees of the same genotype planted in two environments, env1 and env2, as the absolute phenotypic difference between the two measurements divided by their sum:

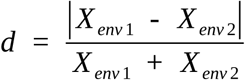

The relative distance is averaged across all pairs of individuals in different environments to calculate the RDPI by genotype:

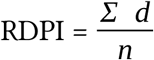

The two genotype replicates in the same garden were treated as individual measurements, with their phenotypes compared to all other individuals except the same-garden individual. Traits which were not normally distributed were log-transformed prior to calculating plasticity indices to meet the assumptions of normality (Valladares et al. 2006): leaf mass, leaf area, adaxial stomata density, and stomata ratio.

### Statistical Analyses

Multiple statistical models were developed to test our three main questions and conducted using R version 4.5.2 (R Core Team 2024). A full list of statistical models and the R functions used is given in **Table S2.**

### Q1. Do responses to varying environments differ across genotypes based on their home climate, species ancestry, and degree of hybridization?

To quantify the variation explained by genotype (G), environment (E), and genotype × environment interactions (G×E), we used a linear mixed-effect model (function lmer in the package lme4, v1.1-37, Bates et al. 2015) with random effects of genotype, garden, genotype × garden interaction, and block nested within garden, and calculated the variance explained by each factor. For stage 7 presence and adaxial stomata presence, which were measured as binary (presence/absence) variables, we used the glmer function from the lme4 package with a binomial link function.

For models 1-3, *p*-values were adjusted for multiple testing across traits and predictors using the Benjamini-Hochberg correction (Benjamini and Hochberg 1995) with a false discovery rate of 0.05. Across all models, trees are represented as individual *i* from genotype *j* planted in garden *k,* B_l(k)_ represents the random intercept of block *l* nested within garden *k*, and ɛ represents the residual error term.

**Model 1** predicts trait Y for the individual *i*, where *u* represents the overall mean of the model, G_j_ represents random intercept of genotype, E_k_ represents random intercept of garden, and G_j_E_k_ represents their interaction.

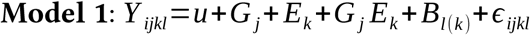

To determine which specific climatic and genomic variables have the strongest effect on trait variation across environments, we used a more complex linear mixed-effects model using lmer (**Model 2**). We tested GxE effects on trait variation in two ways: (1) interactions between genomic PCs and garden climate were used to assess whether responses varied with genetic structure and species ancestry, and (2) interactions between home and garden climates were used to test whether phenotypic responses depended on home climate, consistent with climate-mediated selection on trait values. These two tests are not mutually exclusive and may act together, because climate is correlated with genetic structure (**Figure S2**), but we included both to test which factors have the strongest effect on phenotype. To reduce model complexity, we excluded TD as a climate variable because it is correlated with both genetic PC1 and MAT (**Figure S3**).

This test is represented as **Model 2**, which predicts trait Y for individual *i*, where MAT_k_ and MAP_k_ represent garden climate, MAT_j_ and MAP_j_ represent the genotype’s home climate, and PCs 1, 2, and 3 represent genetic PC scores for genotype. B_l(k)_ represents the random effect of block nested within garden, and G_j_ represents the random effect of genotype.

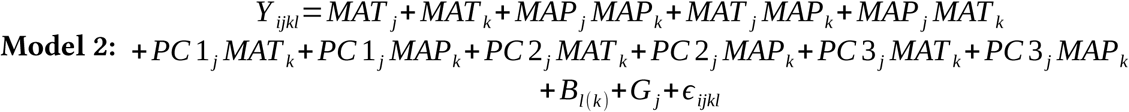

For presence/absence variables (phenological stage 7 presence and adaxial stomata presence), we attempted to fit a model using glmer with a binomial link function as with Model 1, but these models failed to converge, suggesting limited power to predict these variables.

### Q2. To what extent do genetic variation and home climate predict plasticity?

To identify which factors explain variation in trait plasticity across genotypes, we tested whether trait plasticity varied as a function of genetic structure (represented by genetic PCs) or home climate using a linear regression with the function lm (**Model 3**). We used mean annual climate variables represented by MAP and MAT and continentality (TD) as a representation of seasonal temperature variation that may select on plasticity. Because genetic PCs and climate variables covary (**Figure S2**), we tested the effect of each genetic or climatic predictor on plasticity separately to identify those with the largest effects. In **Model 3**, Y_j_ represents trait plasticity for genotype *j* and X_j_ represents the climatic or genetic predictor.

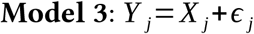

We also tested whether home climate predicts plasticity independently of species ancestry, which would suggest that selection from climate acts similarly across both species. **Model 4** is modified from Model 3, with an additional effect of Pt_j_ representing proportion of *P. trichocarpa* ancestry.

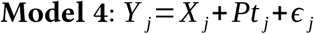

### Q3. Under which environments are trait values and their plasticity adaptive?

To quantify the fitness effects of plasticity as measured by growth increment, we tested whether traits and their plasticity predicted growth increment during the year of measurement. Height is commonly used as a fitness proxy in forest trees representing early-life performance because it confers a competitive benefit in environments experienced by seedlings, and crown height corresponds to reproductive output at maturity (Rehfeldt et al. 1999; Savolainen et al. 2007; Aitken and Bemmels 2016; Younginger et al. 2017; Browne et al. 2019). A linear mixed-effects model (function lmer from the package lme4) was used to test whether trait variation predicted growth increment across environments (**Model 5**). A significant effect of the trait alone would indicate that it influences growth regardless of garden environment, while a significant garden climate × trait interaction would indicate that the relationship between the trait and its effects on growth varied by garden climate. Garden climate was defined using MAT, MAP and TD, with separate models run for each variable. An additional term to control for the species ancestry of each genotype was included, because growth is known to vary between these two species (Mead et al. 2026) and we wanted to isolate effects of the phenotype on growth. This test is represented by Model 5, where Y_ijkl_ represents the yearly growth increment for individual *i*, T_i_ represents its phenotypic trait value, E_k_ represents the garden environment (either MAT, MAP, or TD), T*_i_*E*_k_* represents their interaction, Pt_j_ represents the proportion of *P. trichocarpa* ancestry, B_l(k)_ represents the random effect of block, and G_j_ represents the random effect of genotype.

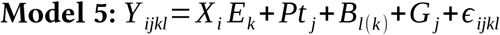

We then tested whether trait plasticity (RDPI) predicted fitness across different garden environments using a similar model, here testing the fitness effects of trait plasticity measured across gardens instead of trait values in each garden. This test is represented by Model 6, where T_j_ represents trait plasticity by genotype.

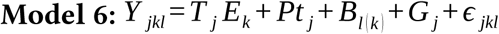

If either trait or trait × climate effects were significant for models 5 and 6, we used a post-hoc test to determine whether the trait or its plasticity predicted growth increment in each individual garden (**Figure 1B**), including a fixed effect of species ancestry and random intercepts for block and genotype. For each trait, p-values for post-hoc tests of individual gardens were adjusted using the Benjamini-Hochberg correction.

## Results

### Q1. Do responses to varying environments differ across genotypes based on their home climate, species ancestry, and degree of hybridization?

On average, phenological traits were best explained by garden environment (E), but there was also some variation due to genotype (G) and G×E effects (**Figure 3**). Among phenological traits, date of lammas growth in 2023 had the largest G×E effect (37%), indicating that genotypes varied in the timing of re-flushing after initial budset across environments. Both leaf and stomata morphology exhibited a relatively higher proportion of unexplained variance compared to phenology, with genotype effects accounting for the majority of explainable variance. In general, traits related to adaxial stomata had a large genotype effect (10-37%), while abaxial stomata traits had a larger environmental effect (27-30%). Variation explained by G×E was particularly high for LMA (25%), indicating genotypic variation in the reaction norm. Factors explaining variance in growth increment differed between the two years: growth increment in 2023 was environmentally dependent, with some impact of G and GxE effects, while growth increment in 2024 had less explainable variance. Ecophysiological traits generally had smaller G×E effects and were explained mostly by the environment. However, stomatal conductance had more variation explained by G and G×E than by the environment alone. Overall, traits with a large GxE effect (9-25% variance explained) included LMA, stomatal conductance, date of lammas growth, and growth increment in 2023, suggesting these traits have genetic variation in plasticity which could be targets for selection when beneficial.

**Figure 3.**
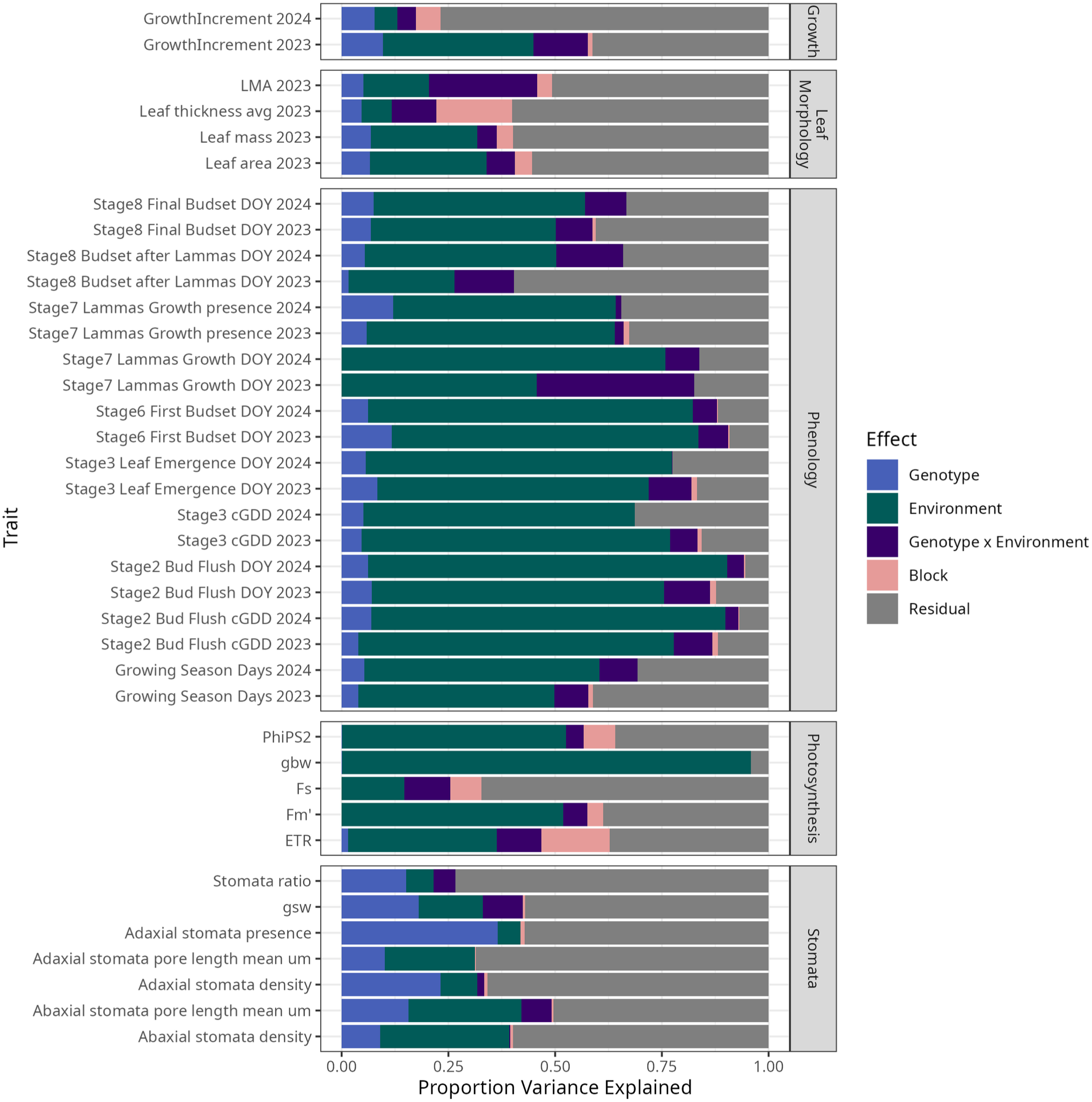
The proportion of phenotypic variation explained by genotype, environment, genotype × environment interactions, and block (nested within garden) varies across trait classes. Most traits were measured in 2023, except phenology and growth, which were also measured in 2024. To calculate variance explained by each factor, we used a linear mixed-effect model with random effects of genotype, garden, and genotype × garden interaction, and block nested within garden.

To identify specific genetic and climate factors contributing to phenotypic variance, we tested the effects of garden climate and home climate (MAT and MAP), species ancestry proportions and intraspecific variation as measured by three genetic PCs, and the interaction of garden climate with genetics or home climate (**Figure 4A**). Of the genetic PCs, species ancestry (PC1) generally had more significant G×E interaction effects. However, intraspecific genetic variation also contributed to G×E, with significant interactions with PC2 (explaining genetic structure within *P. trichocarpa*) and PC3 (separating northern and southern regions). Both home and garden temperatures played an important role in phenology timing. Spring phenology stages occurred earlier in warmer gardens, in genotypes from colder environments, and in genotypes with higher *P. trichocarpa* ancestry, as found previously (Soolanayakanahally et al. 2013). Consistent with variance partitioning (**Figure 3**), garden temperature had the largest effect (β=0.55-0.57) on spring phenology timing (**Figure 4A**), with an increase of 1°C advancing the date of budburst by 4.6 days in 2023. The date of first budset in 2023 had the largest GxE effects (|β|=0.06-0.10), with multiple significant interactions of garden climate with home climate or genetic structure.

**Figure 4.**
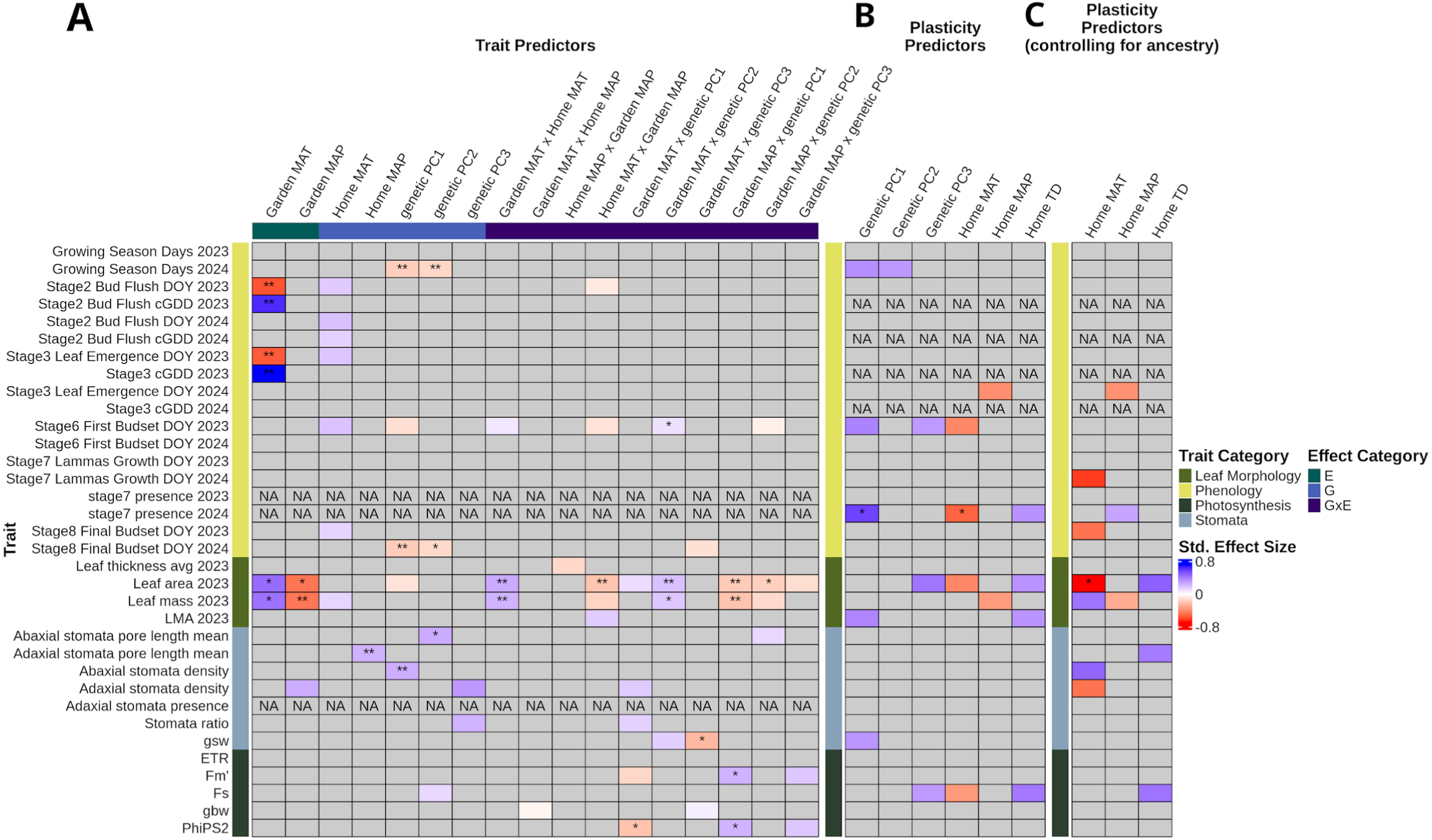
Heatmap showing significant effects of models testing for effects of **A)** garden climate, home climate, genetic PCs, and their interaction on trait values across gardens, **B)** individual genetic and home climate factors on trait plasticity, and **C)** home climate on trait plasticity while controlling for species ancestry. Colored cells indicate a significant effect, and stars indicate a significant effect after p-value correction using the Benjamini-Hochberg method within each panel. Stars indicate p-values as follows: * p≤0.05, ** p≤0.01, ***p≤0.001. Significant G×E effects in panel A suggest there is genotypic variation in phenotypic plasticity for that trait. Significant effects of genetic structure or home climate in B indicate associations with higher (blue) or lower (red) phenotypic plasticity. Significant effects of home climate in C indicate associations with climate that are independent of species ancestry, suggesting the climate exerts selection on plasticity across the hybrid zone of both species.

Leaf morphology, stomata, and photochemistry traits had varying contributions from G, E, and G×E. Leaf mass and area were higher in warmer, drier gardens, although much of this variation was driven by the garden at WSU, which had the largest leaves (**Figure 4A**). Genotypes with higher *P. trichocarpa* ancestry and from warmer environments generally had larger leaves, with degree of genotypic variance differing across gardens (**Figure 4A**). These results suggest that genetic variation in plasticity shapes variation in leaf mass, leaf area, and timing of first budset. Stomatal traits usually had significant genetic effects, especially associated with species ancestry, with *P. balsamifera* genotypes having higher abaxial density (β=0.25) and generally lacking adaxial stomata in colder gardens (**Figure 4**), similar to previous results (Fetter and Keller 2023; Zavala-Paez et al. 2026). Abaxial stomata size and density were more strongly shaped by the environment than adaxial traits, which varied between the two species. Photosynthesis and conductance traits primarily showed significant G×E effects, particularly maximum fluorescence in light (Fm’), stomatal conductance (gsw), and quantum efficiency in light (ΦPS2). *P. trichocarpa* genotypes near the coast and genotypes from northern regions (genetic PC2 and PC3) had higher stomatal conductance. Species ancestry affected Fm’ and ΦPS2, with the direction varying across gardens, suggesting contrasting responses to garden climate in the two species.

### Q2. Does home climate and/or genetic structure predict trait plasticity?

We found greater plasticity in genotypes with higher *P. balsamifera* ancestry and from northern, colder, and more continental regions. In traits for which species ancestry significantly explained plasticity, genotypes with greater *P. balsamifera* ancestry were more plastic (positive effect of genetic PC1) for growing season days in 2024, date of first budset in 2023, presence of lammas growth in 2024, LMA, and stomatal conductance (β=0.32-0.57) (**Figures 4B and 5**). Northern genotypes (represented by genetic PC3) and those from colder regions had higher plasticity in date of first budset in 2023, leaf area, and minimum fluorescence in light (F_s_) (β=0.31-0.42) (**Figure 4B and S4**). Genotypes from more continental regions had higher plasticity in the presence of lammas growth, leaf area, LMA, and F_s_ (β=0.33-0.41) (**Figure 4B and S4**).

**Figure 5.**
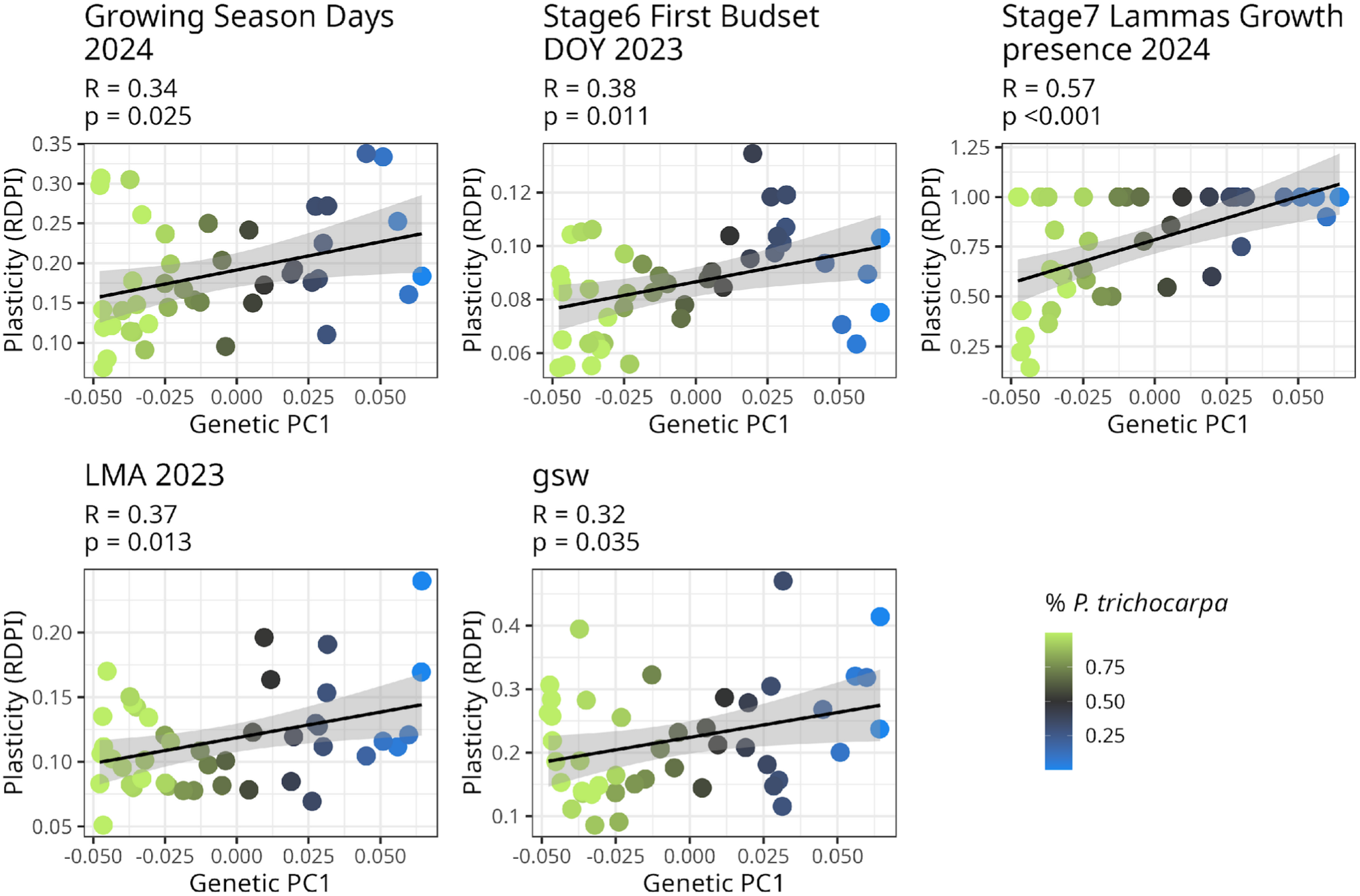
Higher *P. balsamifera* ancestry is associated with higher plasticity in the length of the growing season, date of first budset, occurrence of lammas growth, leaf mass per area, and stomatal conductance (gsw). Plasticity is measured as the relative distance plasticity index (RDPI). Standardized effect sizes, equivalent to Pearson correlations (R) and p-values (p) are shown for each regression model. Points represent genotypes and are colored by the proportion of *P. trichocarpa* ancestry. Significant relationships with genetic PCs 2 and 3, and with home climate, are shown in Figure S4.

We found that higher plasticity in genotypes from colder and more continental environments was not only explained by species ancestry; instead, the environment may be selecting for plasticity across the hybrid zone. Species ancestry was associated with climate variables, with *P. balsamifera* occurring in colder, drier, and more continental regions (**Figure S3**). We tested whether home climate explained plasticity independently from species ancestry. When accounting for species ancestry, the effect of home climate on plasticity was maintained or strengthened for most traits (**Figure 4C**). Genotypes from colder environments had higher plasticity in timing of lammas growth and final budset, leaf area, and adaxial stomata density, but lower plasticity in leaf mass and lower stomata density. Genotypes from drier environments were more plastic in the timing of leaf emergence and accumulation of leaf mass but had lower plasticity in whether lammas growth occurred. Genotypes from more continental climates had higher plasticity in leaf area, adaxial stomata pore length, and F_s_.

### Q3. Under which environments are trait values and their plasticity adaptive?

To test the potential fitness effects of trait variation and its plasticity, and whether those effects varied by garden climate, we tested whether traits predicted growth increment. A significant trait effect on growth increment suggests a fitness effect regardless of environment, while a significant trait × climate interaction suggests that fitness effects of the trait vary by environment. With regard to trait values, earlier bud burst in 2023, thicker leaves, higher leaf mass and area, smaller adaxial stomata, higher adaxial stomatal density, and later date of first budset in 2024 were associated with higher growth increment regardless of garden environment or species ancestry (**Figure S5**). The effects on growth varied by garden MAT (significant trait × climate effect) for date of first budset in 2024, stomatal conductance, ΦPS2, F_s_, LMA, date of last budset in 2024, and growing season days in 2024 (**Figure S5**). A subset of these traits also had growth effects which varied by garden TD: gsw, LMA, and date of last budset in 2024 (**Figure S6**). Fitness effects varied by garden MAP for many of the same traits: date of first and last budset in 2023, stomatal conductance, gbw, ΦPS2, F_s_, LMA, leaf mass, leaf area, and both adaxial and abaxial stomata pore length (**Figure S7**). While multiple traits had significant trait × climate effects on growth, post hoc tests of individual gardens generally only showed significant fitness effects in select gardens.

The effects of trait plasticity on growth varied widely depending on the trait and environment. Greater trait plasticity was associated with higher growth increments independent of planting site climate for SD_abaxial_, and independent of MAP for leaf thickness. However, for most traits with a significant effect of plasticity on growth increment, the strength and/or direction (negative or positive) of the effect varied by garden MAT (**Figure S8**). Only two traits had relationships of plasticity with growth that varied by garden TD (2024 bud flush and leaf emergence, **Figure S9**), and three traits showed relationships of plasticity with growth that varied by garden MAP (gbw, leaf area, and lammas growth presence in 2024, **Figure S10**). Plasticity in earlier spring phenology was associated with higher growth in less continental environments (**Figure S9**). For some traits, plasticity had opposite effects on growth increment in different environments. For example, plasticity in growth increment and leaf thickness generally had positive fitness effects in warmer gardens, while plasticity in fall phenology had positive fitness effects in colder gardens (**Figure S8**). The effects of traits and their plasticity on growth sometimes showed opposite patterns with regard to the environment (**Figure 6, S11**). For example, a constitutively later date of first budset corresponded to increased growth in warmer environments, while greater plasticity in this trait corresponded to higher growth in colder environments. This pattern suggests that trade-offs may exist between the expression of optimal trait values and their plasticity.

**Figure 6.**
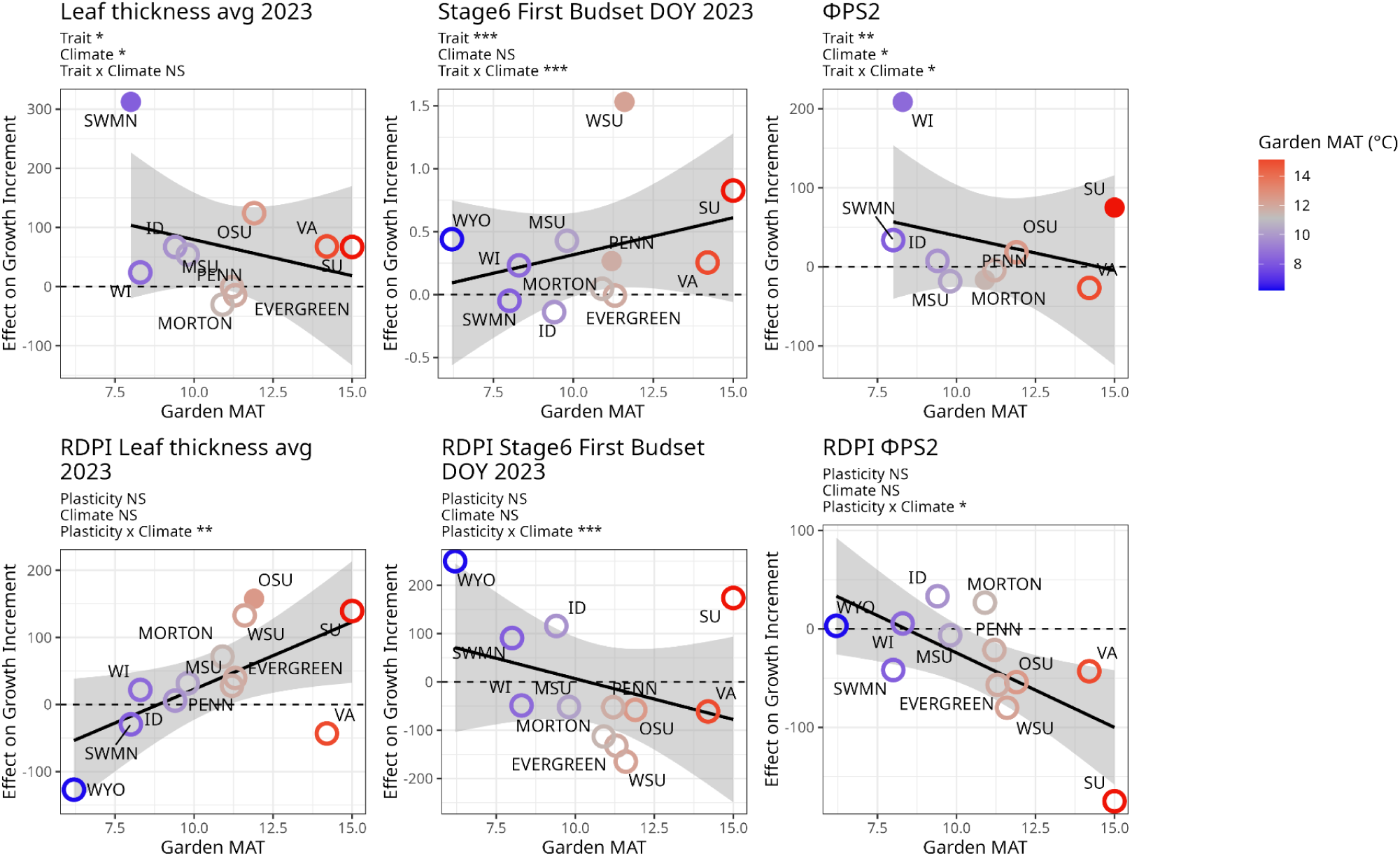
Effects of traits and their plasticity on yearly growth increment. A positive effect indicates that the trait or its plasticity is associated with increased height, while negative effects indicate a deleterious effect for that garden. Filled circles indicate garden sites with a significant relationship after a post-hoc test. Points are colored with the MAT (°C) of the garden and trendline indicates fit and standard error for the effect across garden MAT. Plots are shown for the subset of traits with a significant trait × climate effect and plasticity × climate effect, which suggests that the effect of trait or its plasticity on growth differs by garden MAT. For all traits and climate variables, see Figure S5-S10. Stars indicate p-values for each effect as follows: * p≤0.05, ** p≤0.01, ***p≤0.001.

## Discussion

Using *Populus* genotypes from a natural hybrid zone, we tested how plasticity varies with ancestry and home climate and whether increased plasticity has fitness trade-offs in warming environments. Plasticity appears to reflect historical selection, with genotypes having greater *P. balsamifera* ancestry and originating from colder, more continental climates showing higher plasticity across multiple traits, consistent with adaptation to greater climatic variability. However, for most traits, plasticity had negative or neutral associations with growth in warm environments, suggesting that plasticity is unlikely to offset the advantage of faster-growing, warm-adapted *P. trichocarpa* genotypes under future climates. Overall, our results identify a potential evolutionary tradeoff in which plasticity favored in historically variable climates may come with a cost of reduced growth in warmer environments, limiting its adaptive value under future climates.

### Genetic variation in trait plasticity is shaped by home climate

Several traits across categories, including stomatal conductance, LMA, and lammas growth timing had relatively high variance (9-37%) explained by genotype × environment interactions, suggesting these traits contain genetic variation in plasticity (**Figure 3)** which could be heritable (De Kort et al. 2020). Leaf mass, leaf area, and first budset timing in 2023 showed several significant G×E effects when temperature and precipitation were considered (**Figure 4A**), suggesting that plasticity in these traits could be driven by natural selection associated with climate gradients. Traits strongly influenced by environmental effects, like spring phenology timing, will likely respond to future environments (Anderson et al. 2012a; Piao et al. 2019) but may have relatively limited genetic variation for natural selection to act upon (average variance explained by G×E = 7%). Traits with strong genotypic effects, like stomatal morphology, may be more likely to experience selection on the trait itself than on its plasticity. Broadly, plasticity in leaf allocation traits, stomatal conductance, and fall phenology timing could be targets for natural selection if they influence fitness in future climates.

### Plasticity is associated with P. balsamifera ancestry and cold, continental home environments

Trait plasticity varied with species ancestry and home climate (**Figure 4B**), suggesting that selection has influenced plasticity at interspecific and intraspecific levels across the hybrid zone. Broadly, we found that genotypes with higher *P. balsamifera* ancestry and from colder and more continental environments were more plastic, suggesting that plasticity has evolved as an adaptation to greater seasonal variability. Previous work asking whether slower-growing, resource-conservative species have higher plasticity have observed conflicting results. On one hand, species in more variable environments may evolve increased plasticity as an adaptive strategy to take advantage of relatively rare beneficial conditions while being more resource-conservative in harsh periods (Atkin et al. 2006; Richards et al. 2006; Soolanayakanahally et al. 2009; Solé-Medina et al. 2022). Alternatively, the slower growth rates and higher tissue construction costs of resource-conservative taxa may constrain plastic responses (De Kort et al. 2020; Stotz et al. 2022; Ramírez-Valiente et al. 2025). Here, we found evidence for higher plasticity in *P. balsamifera*, the species from colder and more seasonally variable environments. Genotypes with greater ancestry from *P. balsamifera* had lower growth rates (Mead et al. 2026) and higher plasticity in several traits, including fall phenology timing, LMA, and stomatal conductance (**Figure 4B**), allowing them to take advantage of beneficial conditions (Soolanayakanahally et al. 2009). Plasticity associated with tissue allocation could also allow genotypes to respond to stressors by re-allocating carbon to stress and defense responses rather than growth (Eisenring et al. 2022). The drier climates of *P. balsamifera* may have selected for plasticity in traits affecting evapotranspiration, including LMA and stomatal conductance (**Figure S1 and S2**), allowing plants to avoid water loss in dry periods while increasing carbon fixation when water is available (Chaves et al. 2016). Plasticity in stomatal conductance could also enable greater short-term control of evapotranspiration and may be partly explained by higher stomatal density on the abaxial leaf surface (**Figure 4A**). Overall, these results suggest that the two species have evolved contrasting strategies, with *P. balsamifera* having higher plasticity across multiple traits as part of a “generalist” strategy (similar to genotype C in **Figure 1**), enabling it to maintain fitness in the variable environments it inhabits. However, this strategy may come with a trade-off of reduced growth potential in warmer, less variable environments, where *P. trichocarpa* has higher growth rates (genotype A in **Figure 1**).

We also found that plasticity was associated with the home climate independently of species ancestry, consistent with intraspecific local adaptation (**Figure 4C**). Our findings suggest that increased plasticity in phenology, leaf allocation, stomatal conductance and morphology, and photochemistry could facilitate adaptation to colder and/or more variable environments. Leaf allocation, stomatal conductance, and photosynthetic allocation can all contribute to increased carbon assimilation rates, which may compensate for shorter growing seasons in high-latitude and more continental populations, as found previously in *Populus* (Gornall and Guy 2007; Soolanayakanahally et al. 2009; Keller et al. 2011; McKown et al. 2014a). Here we show evidence for selection on trait plasticity in addition to the traits themselves, which could allow genotypes from more variable environments to take advantage of beneficial growing conditions and alter leaf morphology within a growing season as new leaves develop (McKown et al. 2014a; Lawrence et al. 2021). Previous work in single-species studies has found evidence that *Populus* genotypes from low-resource environments, such as those with short growing seasons or low water availability, may alter their phenotypes to take advantage of high-resource environments (Soolanayakanahally et al. 2009; Oubida et al. 2015; Liu and El-Kassaby 2019) Northern *P. balsamifera* genotypes, which typically have lower growth rates, may actually outcompete southern genotypes in high-resource environments (Soolanayakanahally et al. 2009). Similarly, *P. trichocarpa* genotypes from drier areas had lower water use efficiency than other genotypes in a common garden with high moisture availability, suggesting an ability to take advantage of a relatively less stressful environment (Oubida et al. 2015). Overall, our results are consistent with local adaptation of trait plasticity to different climate regimes, with colder and more variable environments selecting for higher plasticity.

Relative to parental genotypes, genotypes of hybrid ancestry had intermediate plasticity, contrasting with findings that hybrids can exhibit increased plasticity (Marchal et al. 2019; Liu et al. 2021). One possible explanation for this inconsistency is that prior studies were conducted in F1 hybrids, where heterosis is high and thought to be the underlying cause of plasticity (Marchal et al. 2019; Liu et al. 2021). In contrast, our hybrids are advanced-generation backcrosses that do not exhibit heterosis (Mead et al. 2026). This suggests that transgressive segregation has not produced increased phenotypic plasticity compared to the parental species, that natural selection favors intermediate plasticity in hybrids, or both. Given that poplar hybrids typically inhabit intermediate environments, we hypothesize that selection is likely to favor intermediate levels of plasticity, balancing its benefits and costs in environments where both temperature variability and averages are moderate.

### Associations of plasticity with growth suggest neutral or maladaptive effects in warm environments

It has been hypothesized that higher plasticity will be beneficial under future environments, enabling plants to respond to greater climate extremes and variability (Anderson et al. 2012b; Vázquez et al. 2017; De Kort et al. 2020; Jacob et al. 2024). Yet the limited number of previous studies testing this hypothesis in novel environments have found both positive and negative effects of plasticity (Murren et al. 2015; Liu and El-Kassaby 2019; Walter et al. 2023). Here, we find that benefits from increased plasticity are far from universal. The effects of plasticity on growth were highly variable across traits and environments, exhibiting negative, neutral, and beneficial effects. While across-garden effects on growth were not always significant in individual gardens, the direction of the effect across garden climates allows us to describe general patterns where plasticity may be beneficial. Here, we focus on the effect of varying plasticity in warming environments; for further discussion of each trait class, see Supplementary Discussion. The plasticity of budset timing, presence and timing of lammas growth, ΦPS2, and F_s_ had negative or neutral effects on growth in warmer environments. In contrast, plasticity in traits related to aboveground biomass accumulation (growth increment and leaf thickness) had a positive effect on growth in warmer gardens (**Figure 6, Figure S8**) which could confer a competitive advantage under warming climates (Liu and El-Kassaby 2019), but this variation was largely independent of a genotype’s ancestry or home climate.

Budset timing represents a trait where higher plasticity may be detrimental under warming climates by constraining the ability to extend the growing season. Budset timing is likely controlled by both photoperiod and temperature in poplar (Soolanayakanahally et al. 2013; McKown et al. 2018), enabling trees to respond to yearly variation in the onset of freezing temperatures. Having a later budset in 2023 was generally associated with higher growth, particularly in warmer gardens (**Figure 6, Figure S3**), likely because aboveground growth accumulation continued later into the year. However, increased plasticity in fall phenology timing had the opposite effect, generally being associated with lower growth in warmer gardens and higher growth in colder gardens. This could reflect a trade-off between the trait and its plasticity: genotypes that set bud later are less plastic (**Figure S9**), which is beneficial in a warm environment but could indicate an inability to set bud earlier in environments where fall frosts are a greater risk. More-plastic genotypes set bud earlier overall, which is likely beneficial in cold environments but reduces yearly growth in warm environments. Higher plasticity in fall phenology was associated with *P. balsamifera* ancestry (**Figure 5**), so the benefit of increased plasticity in colder environments could be a species-specific strategy that evolved to survive shorter growing seasons and higher frost risk in fall. Similarly, Cooper et al. (2019) observed that earlier budset in a cold environment was associated with higher survival for *P. fremontii* populations, but interestingly, later budset was associated with increased plasticity, not decreased as we find here (Cooper et al. 2022). The correlation between budset timing and its plasticity could vary among species; *P. fremontii* occupies warmer and drier habitats than the species studied here, which may select for different trait-plasticity relationships.

Though genotypes from continental environments had higher plasticity, it was not sufficient to increase their aboveground growth rates in warmer gardens, and in a natural environment they would likely be outcompeted by less plastic but faster-growing genotypes (Mead et al. 2026). This pattern suggests trade-offs associated with increased plasticity, which may arise if the plasticity of a particular trait constrains the expression of optimal trait combinations in warm environments (DeWitt et al. 1998). Inherent trade-offs could arise if the loci underlying trait expression also determines their plasticity (Laitinen and Nikoloski 2019; Li et al. 2019; Mu et al. 2022; Jin et al. 2023; Napier et al. 2023). In some cases, traits and plasticity were correlated with each other (**Figure S11**). Identifying the genomic basis of plasticity will be necessary to determine whether this pattern arises from correlated selection pressures or non-independence of loci controlling traits and their plasticity due to pleiotropy or linkage (Scheiner 1993; Laitinen and Nikoloski 2019; Jin et al. 2023). Importantly, relationships between plasticity and growth do not necessarily result from direct effects of plasticity on fitness (Via 1993; Auld et al. 2009). If there are trade-offs between traits and their plasticity, we cannot disentangle which factor selection is acting on.

The relationship between trait plasticity and growth varied widely among gardens, with positive or negative relationships typically being specific to certain gardens rather than shared among gardens with similar climates. In a meta-analysis of plant phenotypic plasticity, Stotz et al. (2021) found that plastic responses were more often associated with non-climatic factors, like light and nutrient availability, than climatic factors. Our common garden sites have variation in many unmeasured non-climatic factors, including soil characteristics, nutrient availability, and herbivore pressure. Also, annual climate averages are likely correlated with unmeasured factors and may not be the main drivers of selection. Plasticity in the traits we measured was likely shaped by all these factors in combination with climatic variables, resulting in garden-specific effects on plant fitness. Additionally, these results are likely shaped by our study system. In northwestern North America, climatic variability is correlated with colder environments (**Figure S2**), causing phenotypic variation to experience a trade-off between plasticity and growth in warm climates. However, in other systems, warmer environments may have higher variability, selecting for genotypes that are likely well-adapted to future environments that combine higher variability and temperature (Oliver and Palumbi 2011). These findings illustrate that quantifying phenotypic plasticity across novel environments can identify trade-offs between traits and their plasticity, which will shape plant responses to climate change.

## Conclusions

Phenotypic plasticity is often assumed to be an adaptive strategy for coping with environmental variation, but here we found that genotypes with higher plasticity most often had similar or decreased growth compared to those with lower plasticity. Genotypes with higher *P. balsamifera* ancestry and from colder, seasonably variable environments were more plastic, suggesting selection favors greater plasticity for genotypes and species originating from more variable environments. However, increased plasticity generally did not correspond to higher growth rate in warmer environments, suggesting a potential trade-off wherein higher plasticity may come at a cost to fitness in warm environments. Together, our results suggest that warming climates will favor increased growth from *P. trichocarpa* genotypes over higher plasticity from *P. balsamifera* genotypes. Hybrids had plasticity intermediate to the two parental species, which could result if introgression does not produce increased plasticity, or if environments in the hybrid zone select for intermediate plasticity. Future work investigating the genetic basis of traits and their plasticity in hybrids could determine whether selection on standing genetic variation could combine loci underlying the plasticity of *P. balsamifera* and the high growth in warm environments exhibited by *P. trichocarpa*, or if the strategies of the two species result from inherent trade-offs among many phenotypic traits and their plasticity (Zavala-Paez et al.; in prep).

## Supporting information

Analysis Rmarkdown Files

## Acknowledgments

This research was supported by NSF PGR 1856450, the Schatz Center for Tree Molecular Genetics, National Institute of Food and Agriculture Award #PEN04809 to JAH, NIH award R35GM138300 to JRL, and the Life and Environmental Sciences Dept. at UCM. AM was supported by NSF PRFP #2209410. NM was supported by University of Wisconsin– Eau Claire Office of Research and Sponsored Programs through the Biology Research Scholars Program and Summer Research Experience for Undergraduate awards. This material is based upon work supported by the Great Lakes Bioenergy Research Center, U.S. Department of Energy, Office of Science, Biological and Environmental Research Program under Award Number DE-SC0018409.

We would like to thank everyone involved in sampling, garden establishment and maintenance, and data collection, including Marc Aguirre, Shane Baumgart, Debbie Bird, Joseph Braasch, Schaefer Buchanan, Emma Collins, Kevin Cowden, Lionel Di Santo, Katherine Egeler, Marcia Fairbanks, Chase Fillion, Alice Fischer, Abby Ferson-Mitchell, Shelby Flint, Linnea Fraser, Anna Freigen, Carter Guffey, Carri LeRoy, David Hainlen, Carolyn Hanson, Alex Hass, Christian Hernandez, Daria Hutchinson, Erika Jones, Marika Kisgen, Ben Levitt, Eli Levitt, James Levitt, Jessica Lindstrom, Abigail Marshall, Luke McCormack, Pam Mead, Albert Medel, Katie Nelson, George Newcombe, Kyle Peer, Amanda Penn, Tommy Phannareth, Jennifer Pollard, Steven Quick, Michelle Reid, Brendon Reidy, Christy Rollinson, Emma Smidt, Seth Townsend, Guinevere Unterbrink, Paul Warnick, Rebekah Wells, Baxter Worthing, Dailyn Wold, Dean Wu, Kate Volk, Lindsey Zakopal, Thomas Zambiasi, and Qian Zhang. We also thank Alden Stone, who worked on stomatal size and density measurements.

## Author Contributions

JAH, JH, SRK, and MCF designed the study and contributed to data analysis and interpretation. AM performed analyses and lead writing of the manuscript with feedback from JAH. MZ-P contributed to analysis design and interpretation. AM, JRBB, ACB, IPC-M, DGF, JMG, KMH, LAK, SKF, JRL, JML, DBL, NM, EVM, JPS, KLS, and MZ-P contributed to fieldwork and data collection and provided input on garden-specific conditions for data analysis and interpretation. All authors contributed to writing the manuscript.

## Data Availability

Data from this study are included as supplementary material. Analysis scripts are available at https://github.com/alaynamead/popup_poplar_plasticity and will be archived in Zenodo upon manuscript acceptance.

