## Supplementary Material for "Phenotypic plasticity evolved for climate variability constrains performance under climate warming"

**Contents**

Supplementary Methods
Supplementary Discussion

Figure S1. Climate PCA for historic home environments and gardens in 2023 and 2024
Figure S2. Climatic variation by ancestry
Figure S3. Correlation between LeafNet and manual stomata counts
Figure S4. Significant relationships between trait plasticity and genetic PCs or home climate
Figure S5. Fitness effects of traits across garden MAT
Figure S6. Fitness effects of traits across garden TD
Figure S7. Fitness effects of traits across garden MAP
Figure S8. Fitness effects of trait plasticity across garden MAT
Figure S9. Fitness effects of trait plasticity across garden TD
Figure S10. Fitness effects of trait plasticity across garden MAP
Figure S11. Correlations between traits and their plasticity across garden temperatures

Table S1. List of gardens and years/traits measured.
Table S2. List of statistical models.

### Supplementary Methods

*Automated counting of stomata with LeafNet*

Stomatal impressions for abaxial and adaxial leaf surfaces were imaged using the CellSens software on an Olympus BX53 microscope at 40x magnification, taking one image of each leaf surface for each individual while avoiding the midrib and major veins. Adaxial stomatal occurrence was recorded as presence/absence of stomata on the adaxial leaf surface. Pore length was measured for up to five stomata on each leaf surface. If more than five stomata were present, we divided the image into 5 equal regions and measured one stomate per region. The "measure" tool in ImageJ was calibrated using the image's scale bar and used to measure the pore length of each stomate at its widest point. Pore lengths for each leaf surface were averaged for each individual. To automate measurements of stomatal density, we used LeafNet (Li et al. 2022), a deep convolutional network tool which identifies stomata. We used the LeafNet StomaNetUniversal model with a threshold of 40 and an image resolution of 4.12 pixels/μm. To exclude likely artifacts outside of the observed size and shape of stomata in these species, the size range of stomata was set to 20-1500 $\mu m^2$, and the maximum length-to-width ratio to 5. The PeeledDenoiser method was used with a denoise level of 50. To calculate density, we divided the stomata count by the image area evaluated by LeafNet, which excludes a 50-pixel region on all sides, for a total image area of 0.08 $mm^2$. To test the accuracy of LeafNet stomatal counts, we hand-counted a subset of 10 adaxial and 10 abaxial stomatal impressions per garden site (110 total) and correlated the two measurements (**Figure S2).** The two methods produced similar results: the Pearson correlation was 0.88 ($P$ < 2.2e-16) for adaxial stomata and 0.74 ($P$ < 2.2e-16) for abaxial stomata.

### Supplementary Discussion

*Effects of traits and their plasticity on growth*

Leaf allocation traits, like mass, area, and thickness, were positively associated with growth, regardless of garden environment, likely because they reflect aboveground biomass accumulation (**Figure S3**). However, plasticity for growth increment and leaf thickness generally had a positive effect on growth in warmer gardens (**Figure 6, Figure S3**). Increased plasticity in aboveground biomass accumulation likely leads to a competitive advantage in warmer environments.

The effect of plasticity in stomatal morphology on growth was generally weak, but more strongly associated with garden precipitation than temperature (**Figure S8**), consistent with previous studies showing that stomatal trait variation is influenced more by water limitation than thermal tolerance (Liu et al. 2017, 2023; Zavala-Paez et al. 2026). Both adaxial and abaxial pore length had varying effects on growth increment across garden precipitation. Larger pores were associated with decreased growth in dry environments, and smaller pores were neutral or beneficial in wet environments (**Figure S6**), possibly because larger pores increased water loss by increasing time for stomatal closure (Franks et al. 2009). However, plasticity in pore size did not correlate with varying fitness across environments, and the variation in pore length explained by G×E was low relative to the genotype effect (**Figure 3**), suggesting limited ability of selection to act on plasticity.

Some photochemistry traits had fitness effects that varied by environment, but they were generally weak, perhaps because they had limited variance explained by G×E (**Figure 3**) and were measured at different times over the growing season. ΦPS2 was positively associated with growth, more strongly in colder gardens, suggesting increased photosynthetic capacity had a larger benefit in cold gardens with a shorter growing season (Gornall and Guy 2007; Soolanayakanahally et al.

2009; Keller et al. 2011; McKown et al. 2014). However, increased plasticity was associated with lower growth in warmer gardens and had a neutral effect in colder gardens (**Figure 6**), suggesting that the trait can be broadly adaptive but that its plasticity is detrimental in warmer environments. ΦPS2 is correlated with $CO_2$ assimilation rates, assuming similar conditions and stomatal conductance (Murchie and Lawson 2013). We controlled for shading and leaf development, and air temperature was similar within each garden, but stomatal closure could vary with sampling time and among individuals, so these results must be interpreted with caution.

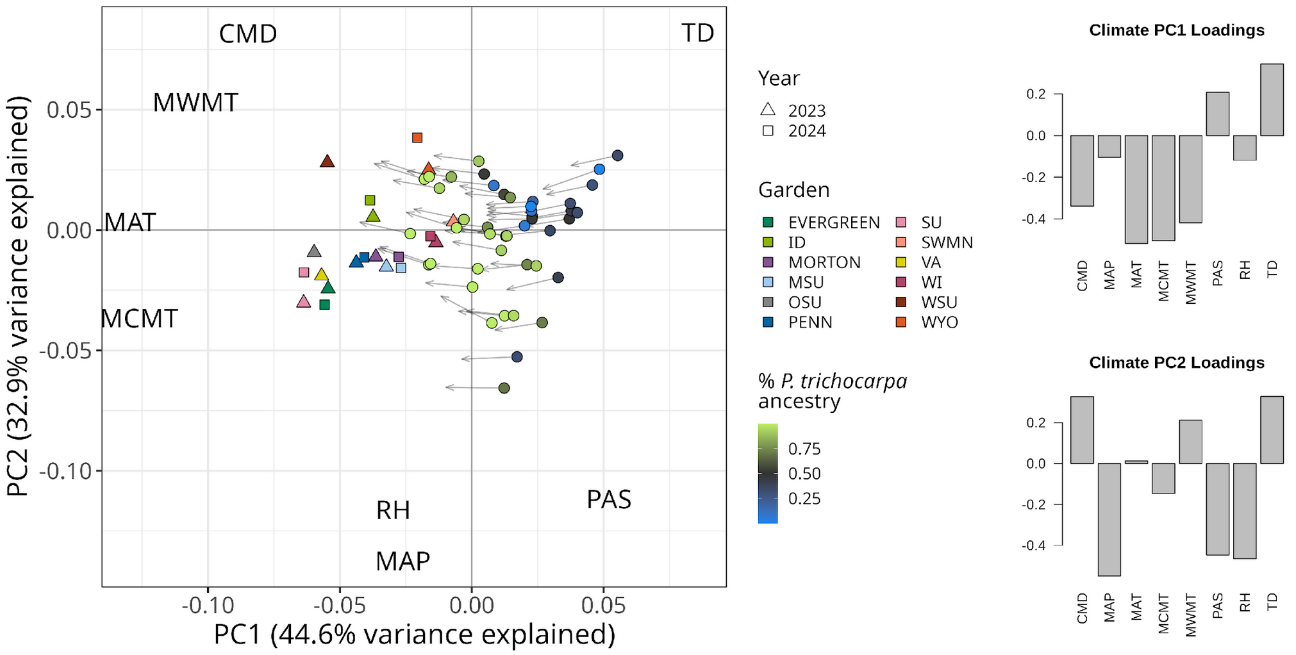


**Figure S1**. Climate PCA for 2023 and 2024 and loadings of each climate variable on PCs 1 and 2. Circles indicate climate of origin for the sampled genotypes for the period between 1961-1990 and are colored by species ancestry. Triangles and squares represent the climate at each common garden site for 2023 and 2024, respectively. Text indicates the loadings of climate variables on each axis, with abbreviations as follows: CMD, climatic moisture deficit; MAP, mean annual precipitation, MAT, mean annual temperature; MCMT, mean coldest month temperature; MWMT, mean warmest month temperature, PAS, precipitation as snow; RH, relative humidity; TD, temperature difference, or continentality. Loadings of climate variables have been downscaled to allow visibility of sites.

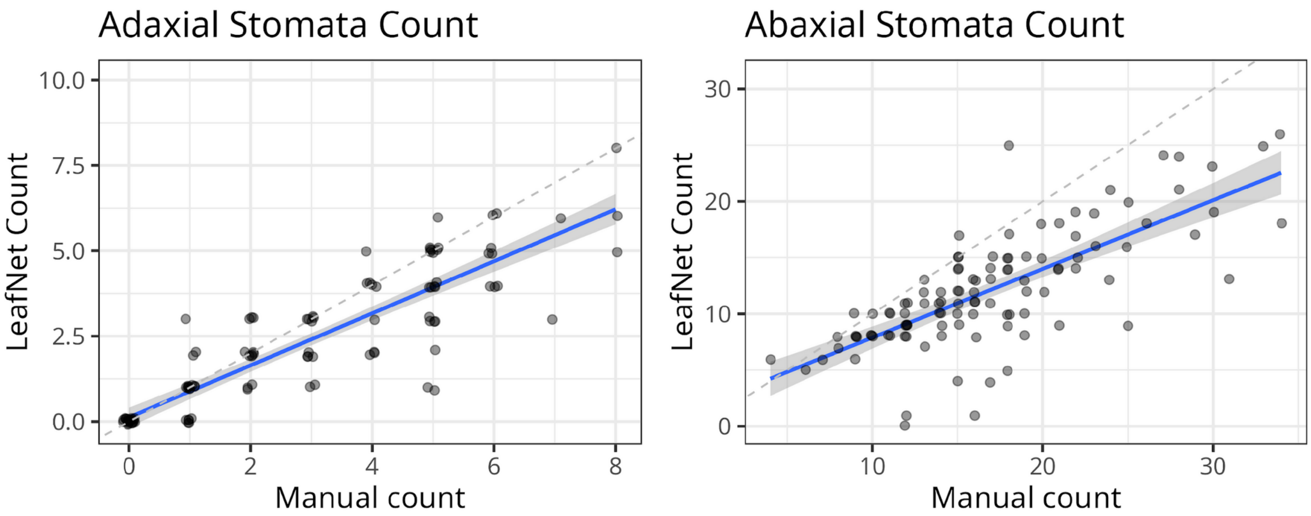


**Figure S2**. Accuracy of stomata counts estimated using LeafNet compared to manual counts for a subset of 110 adaxial and abaxial images (10 per garden). The Pearson correlation was 0.88 ($P$ < 2.2e-16) for adaxial (upper) stomata and 0.74 for abaxial (lower) stomata ($P$ < 2.2e-16). Blue line indicates best fit.

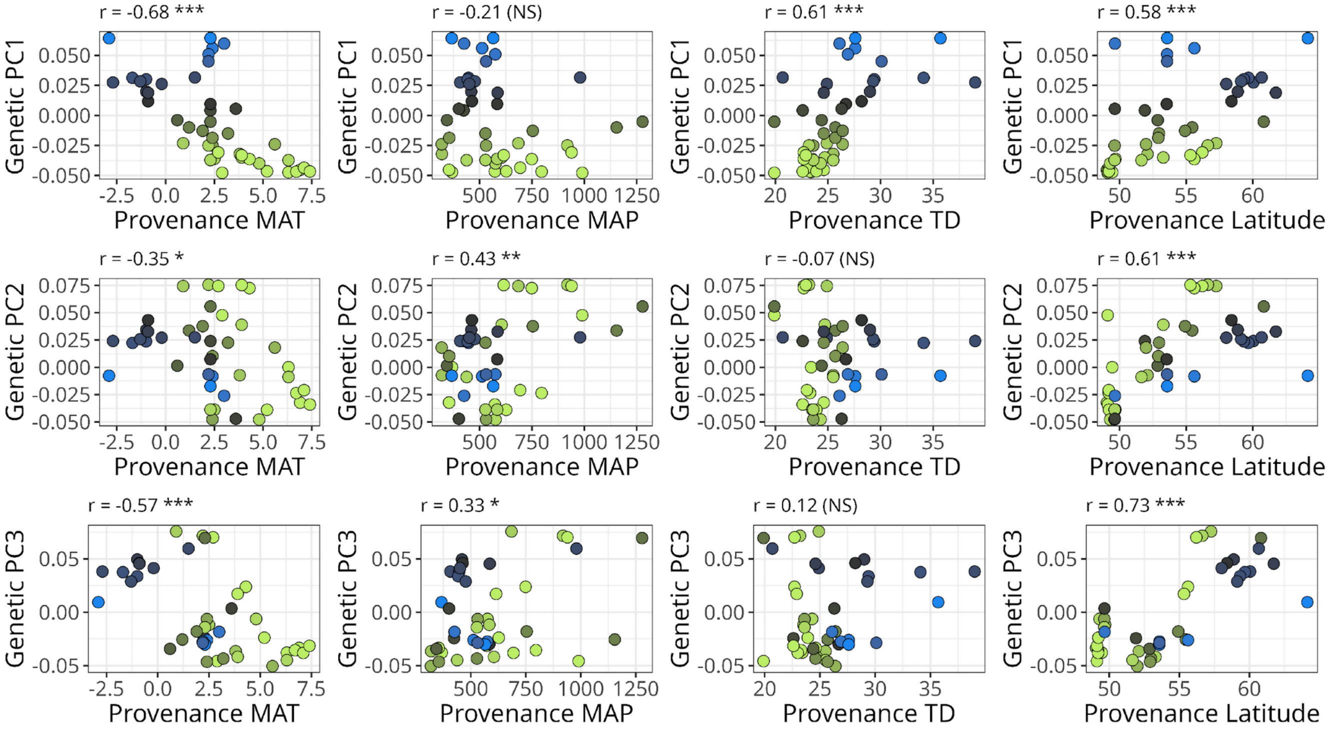


**Figure S3**. Variation in climate across axes of genomic variation. Values for Pearson's correlation (r) between the two variables, and their significance, are given in plot titles. P-values are shown using stars as follows: * indicates p≤0.05, ** indicates p≤0.01, *** indicates p≤0.001, and NS indicates not significant.

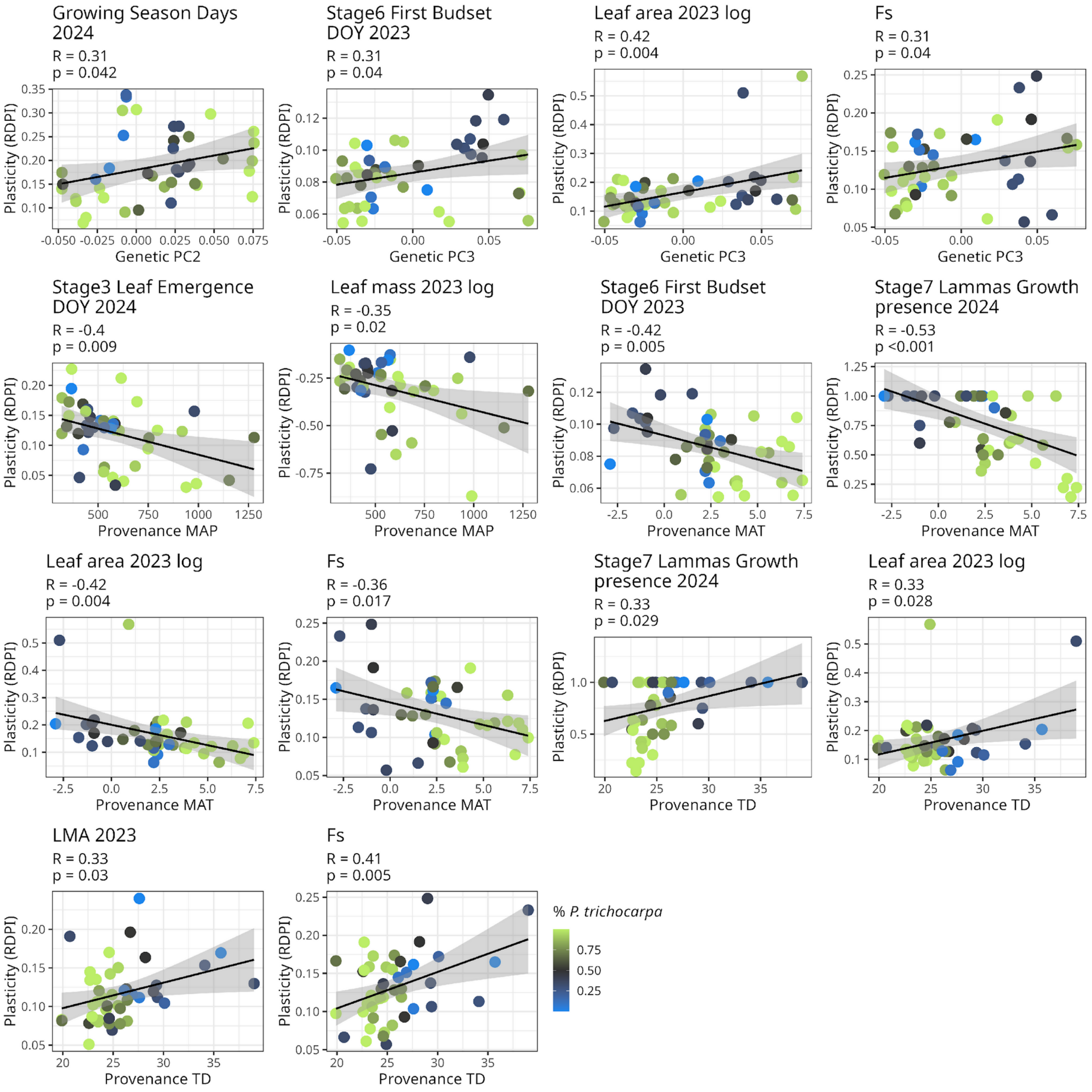


**Figure S4.** Significant relationships of trait plasticity with genetic PCs and climate of origin, as in Figure 5. Standardized effect sizes, equivalent to Pearson correlations (R) and p-values (p) are shown for each regression model. Points represent genotypes and are colored by the proportion of *P. trichocarpa* ancestry.

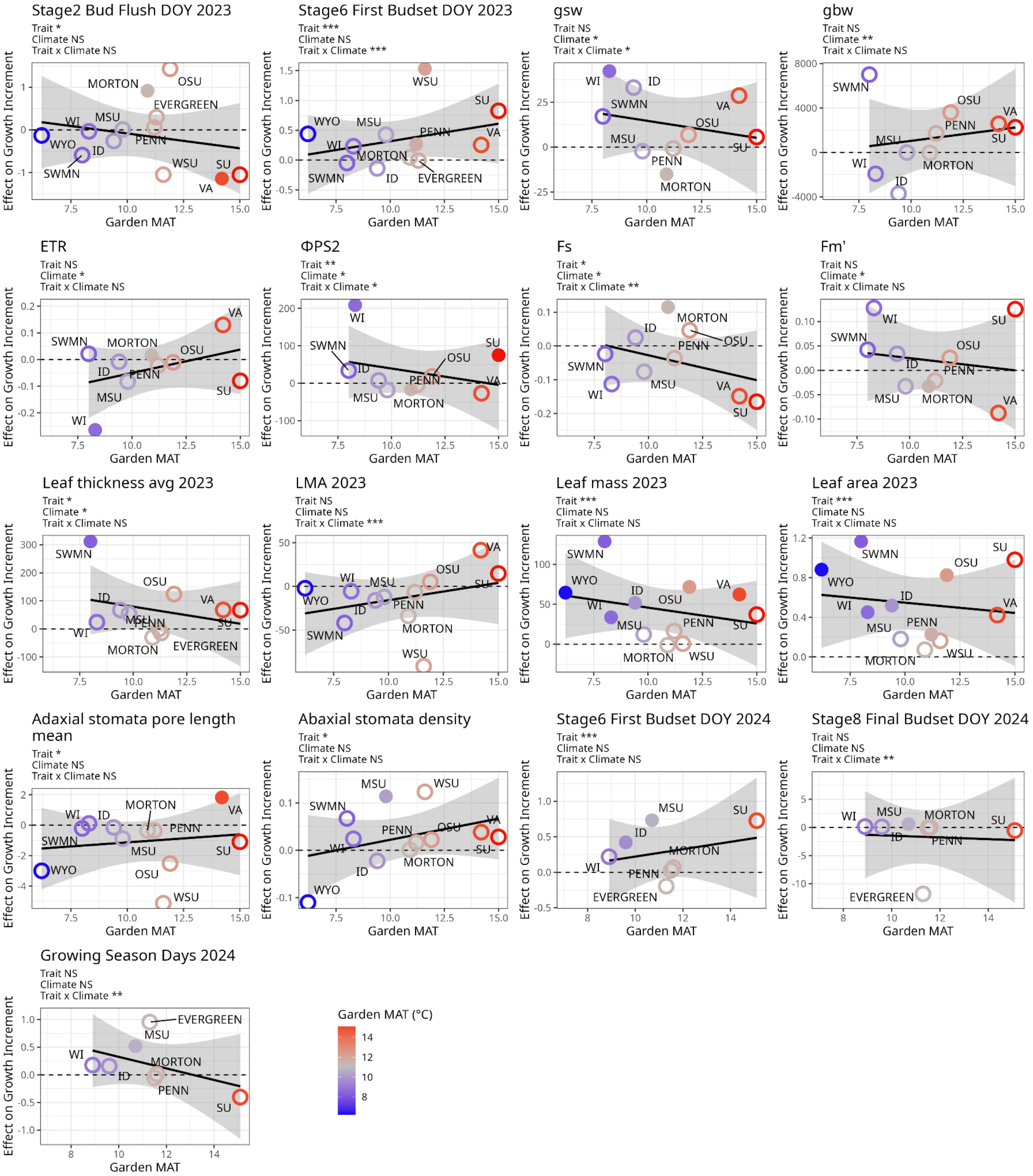


**Figure S5**. Effects of traits on yearly growth increment by garden temperature, as in Figure 5, with all traits that significantly predicted growth increment alone or differently across garden MAT values. A positive effect indicates that the trait is associated with increased height, while negative effects indicate a deleterious effect for that garden. Filled circles indicate garden sites where the relationship was significant in a posthoc test. Points are colored with the MAT of the garden, and trendline indicates fit and standard error for the effect across garden MAT. Stars indicate p-values for each effect as follows: * $p \leq 0.05$, ** $p \leq 0.01$, ***$p \leq 0.001$, and NS not significant.

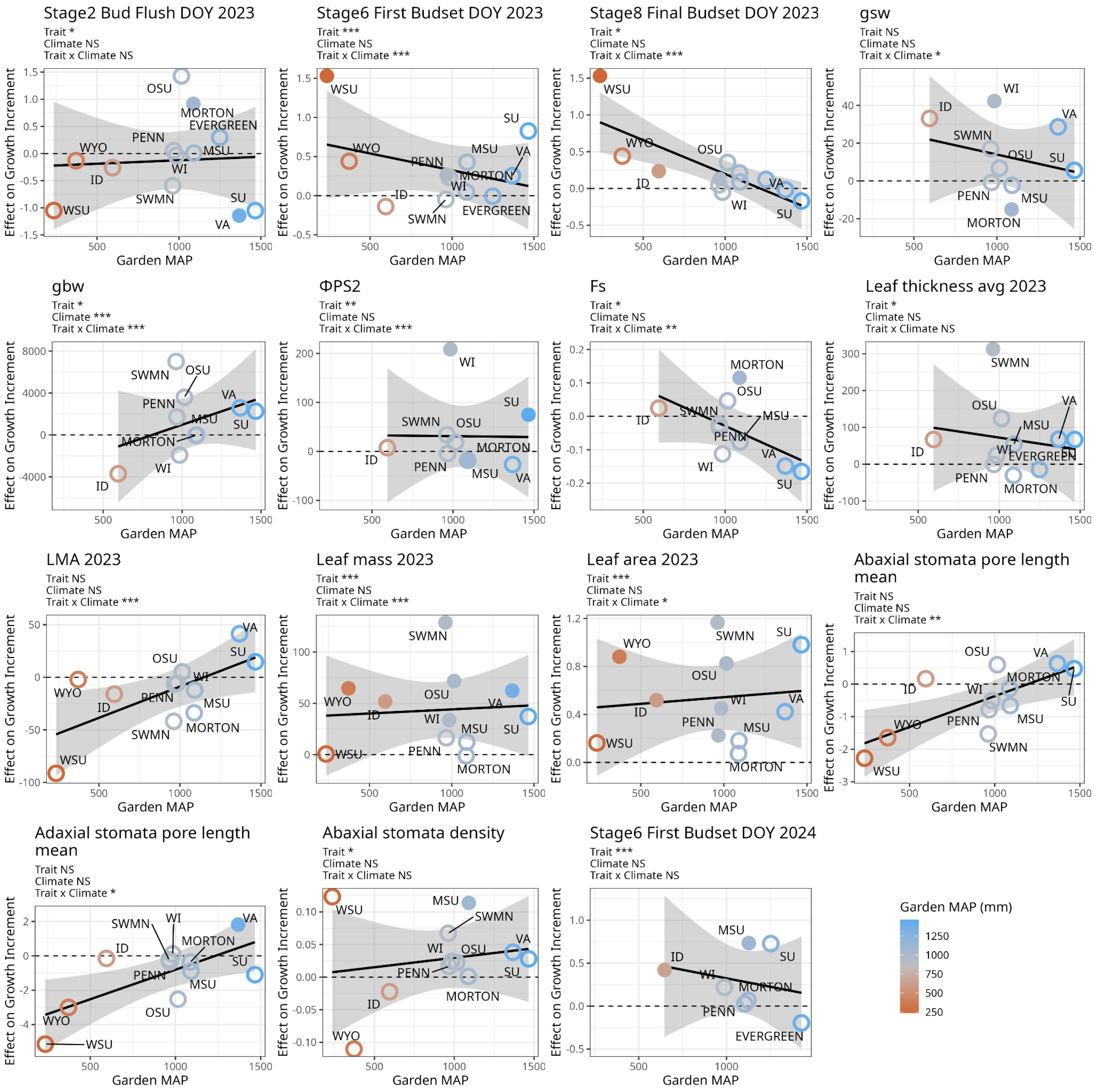


**Figure S6**. Effects of traits on yearly growth increment by garden precipitation, for all traits that significantly predicted growth increment alone or differently across garden MAP values. A positive effect indicates that the trait is associated with increased height, while negative effects indicate a deleterious effect for that garden. Filled circles indicate garden sites where the relationship was significant in a posthoc test. Points are colored with the MAP of the garden and trendline indicates fit and standard error for the effect across garden MAP. Stars indicate p-values for each effect as follows: * p≤0.05, ** p≤0.01, ***p≤0.001, and NS not significant.

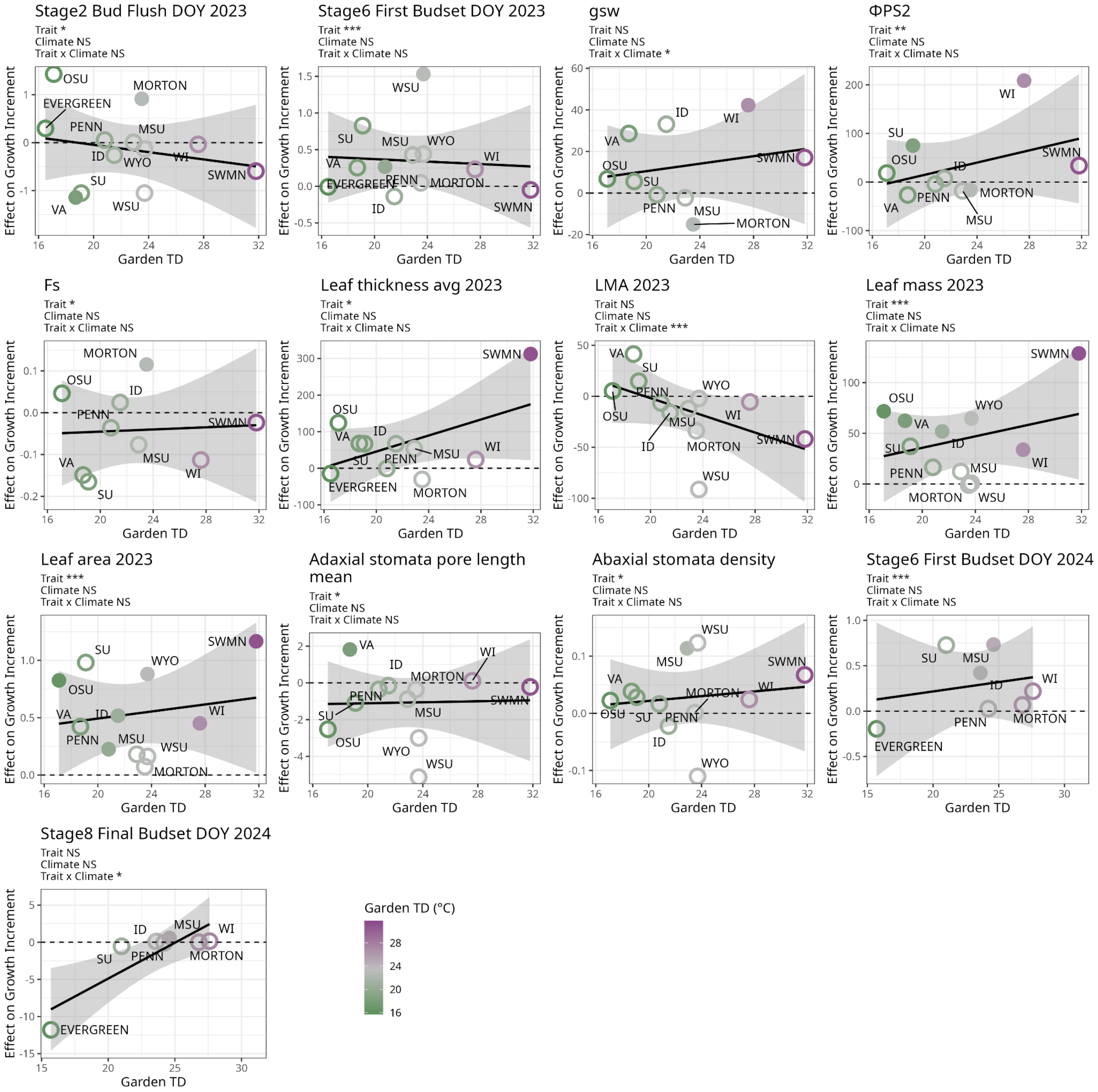


**Figure S7**. Effects of traits on yearly growth increment by garden continentality (TD), for all traits that significantly predicted growth increment alone or differently across garden TD values. A positive effect indicates that the trait is associated with increased height, while negative effects indicate a deleterious effect for that garden. Filled circles indicate garden sites where the relationship was significant in a posthoc test. Points are colored with the MAP of the garden and trendline indicates fit and standard error for the effect across garden TD. Stars indicate p-values for each effect as follows: * p≤0.05, ** p≤0.01, ***p≤0.001, and NS not significant.

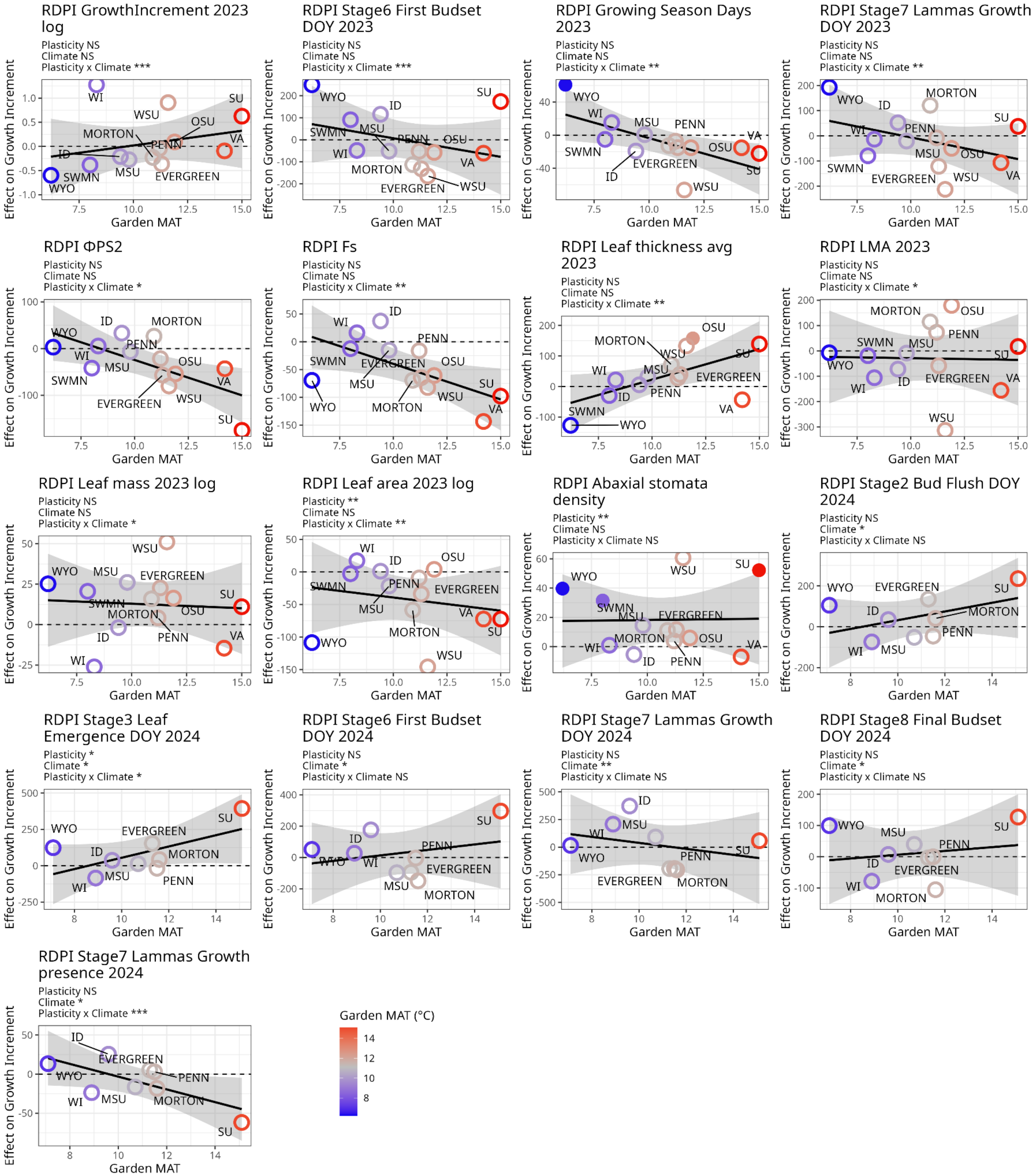


**Figure S8.** Effects of trait plasticity on yearly growth increment by garden temperature, as in Figure 5, with all traits that significantly predicted growth increment alone or differently across garden MAT values. A positive effect indicates that increased plasticity in the trait is associated with increased height, while negative effects indicate a deleterious effect of plasticity for that garden. Filled circles indicate garden sites where the relationship was significant in a posthoc test. Points are colored with the MAT of the garden and trendline indicates fit and standard error for the effect across garden MAT. Stars indicate p-values for each effect as follows: * $p \leq 0.05$, ** $p \leq 0.01$, ***$p \leq 0.001$, and NS not significant.

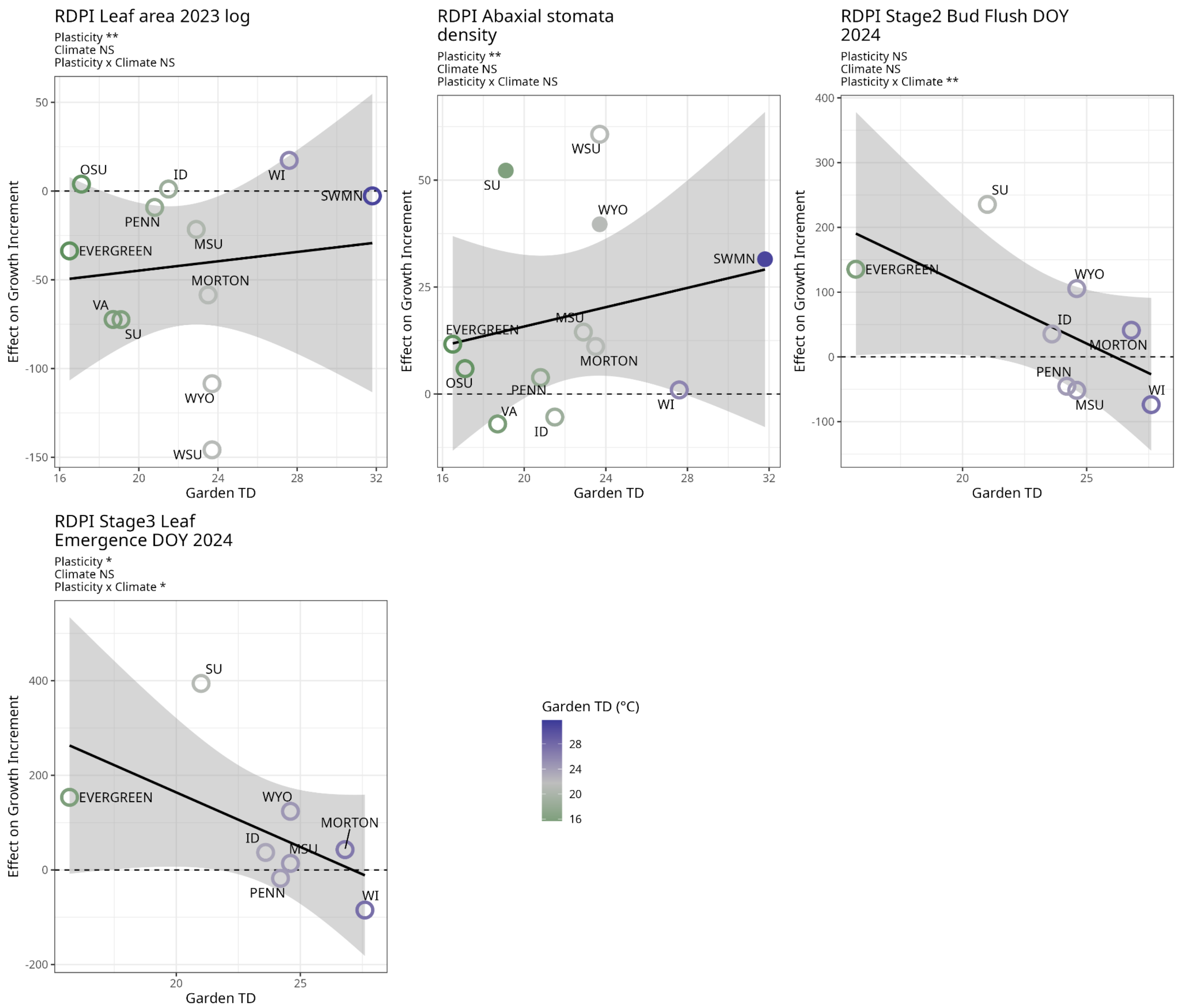


**Figure S10**. Effects of trait plasticity on yearly growth increment by garden continentality (TD), with all traits that significantly predicted growth increment alone or differently across garden TD values. A positive effect indicates that increased plasticity in the trait is associated with increased height, while negative effects indicate a deleterious effect of plasticity for that garden. Filled circles indicate garden sites where the relationship was significant in a posthoc test. Points are colored with the MAP of the garden and trendline indicates fit and standard error for the effect across garden MAP. Stars indicate p-values for each effect as follows: * p≤0.05, ** p≤0.01, ***p≤0.001, and NS not significant.

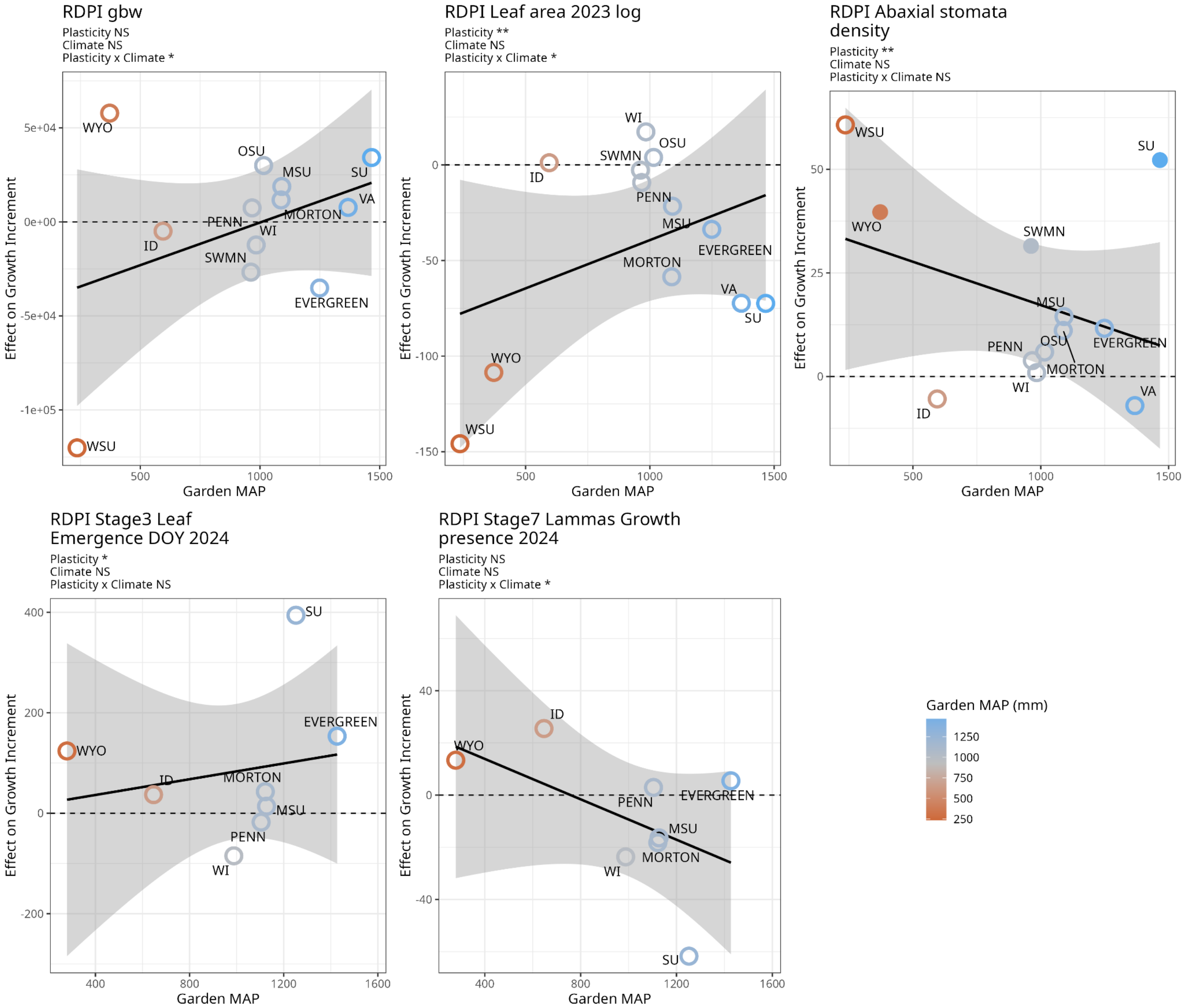


**Figure S9**. Effects of trait plasticity on yearly growth increment by garden precipitation, with all traits that significantly predicted growth increment alone or differently across garden MAP values. A positive effect indicates that increased plasticity in the trait is associated with increased height, while negative effects indicate a deleterious effect of plasticity for that garden. Filled circles indicate garden sites where the relationship was significant in a posthoc test. Points are colored with the MAP of the garden and trendline indicates fit and standard error for the effect across garden MAP. Stars indicate p-values for each effect as follows: * p≤0.05, ** p≤0.01, ***p≤0.001, and NS not significant.

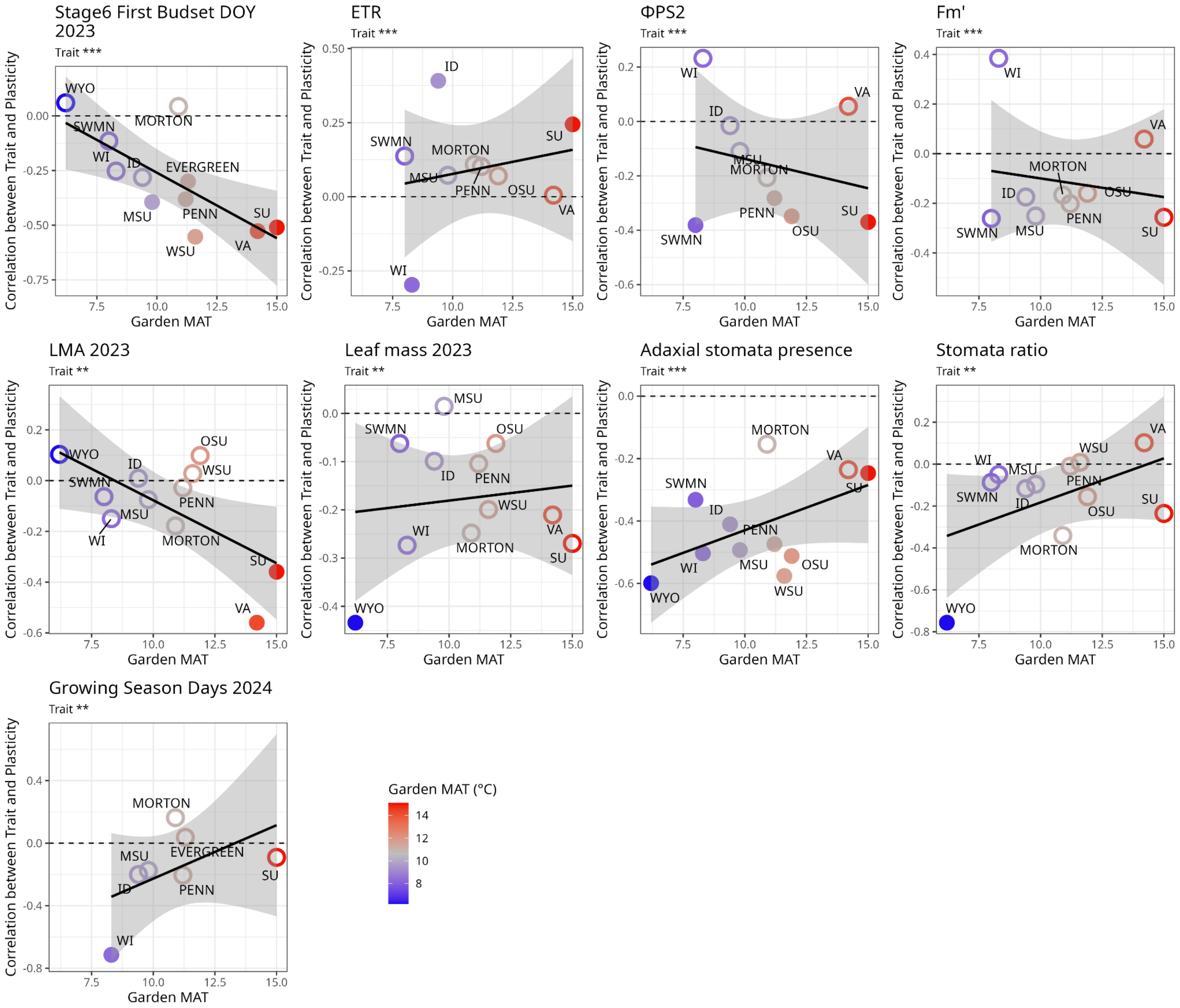


**Figure S11.** Correlations between traits and their plasticity across garden temperatures. A positive effect indicates a positive correlation (the higher values of the trait are associated with increased plasticity). Filled circles indicate garden sites where the relationship was significant in a posthoc test. Points are colored with the MAT of the garden and trendline indicates fit and standard error for the effect across garden MAT. Stars indicate p-values for each effect as follows: * p≤0.05, ** p≤0.01, ***p≤0.001.

**Table S1.** List of gardens and traits measured at each.

| Common Garden Site | Abbreviation | City | State | Spring Phenology 2023 | Fall Phenology 2023 | Leaf thickness 2023 | LMA | Stomata morphology 2023 | LI-COR 2023 | Growth Increment 2023 | Spring Phenology 2024 | Fall Phenology 2024 | Growth Increment 2024 |
|---|---|---|---|---|---|---|---|---|---|---|---|---|---|
| Evergreen State | EVERGREEN | Olympia | WA | x | x | x | | | | x | x | x | x |
| Michigan State University | MSU | East Lansing | MI | x | x | x | x | x | x | x | x | x | x |
| Morton Arboretum | MORTON | Lisle | IL | x | x | x | x | x | x | x | x | x | x |
| Oregon State University | OSU | Corvallis | OR | x | x | x | x | x | x | x | | | |
| Pennsylvania State University | PENN | State College | PA | x | x | x | x | x | x | x | x | x | x |
| Salisbury University Arboretum | SU | Salisbury | MD | x | x | x | x | x | x | x | x | x | x |
| Southwest Minnesota State University | SWMN | Marshall | MN | x | x | x | x | x | x | x | | | |
| University California, Merced | UCM | Merced | CA | x | | | | | | | | | |
| University of Idaho | ID | Moscow | ID | x | x | x | x | x | x | x | x | x | x |
| University of Wisconsin - Eau Claire | WI | Eau Claire | WI | x | x | x | x | x | x | x | x | x | x |
| University of Wyoming | WYO | Laramie | WY | x | Budset only, no lammas | | x | x | | x | x | x | x |
| Virginia Tech | VA | Critz | VA | x | x | x | x | x | x | x | | | |
| Washington State - Wenatchee | WSU | Wenatchee | WA | x | Budset only, no lammas | | x | x | | x | | | |

**Table S2.** Statistical models used in analyses. PCs refer to genomic PCs, Pt refers to *P. trichocarpa* ancestry proportion, MAT is mean annual temperature, MAP is mean annual precipitation, and TD is temperature difference, or continentality.

| Model | Question | R Function | R Equation |
|---|---|---|---|
| *Q1. Do responses to varying environments differ across genotypes based on their climate of origin, species ancestry, and degree of hybridization?* | | | |
| 1 | How much variance in traits is explained by genotype, environment, and genotype × environment effects? | lmer for continuous variables, glmer for binary variables | Trait ~ 1 + (1\|genotype) + (1\|garden/block) + (1\|genotype:garden) |
| 2 | Which environmental, genetic, and G×E effects explain variance in phenotypic traits? | lmer for continuous variables, not run for binary variables | Trait ~ garden_MAT*home_MAT + garden_MAT*home_MAP + garden_MAP*home_MAP + garden_MAP*home_MAT + pc1*garden_MAT + pc2*garden_MAT + pc3*garden_MAT + pc1*garden_MAP + pc2*garden_MAP + pc3*garden_MAP + (1 \| genotype) + (1 \| garden/block) |
| *Q2. To what extent do genetic variation and climate of origin predict plasticity?* | | | |
| 3 | Which measures of genetic variation and home climate predict variation in trait plasticity? (Run separately for each predictor: genetic PCs1-3 and home MAT, MAP, and TD) | lm | Trait plasticity ~ predictor |
| 4 | Which measures of home climate predict variation in trait plasticity independently of species ancestry? (Run separately for each predictor: home MAT, MAP, and TD) | lm | Trait plasticity ~ predictor + Pt |
| *Q3. Under which environments are trait values and their plasticity adaptive?* | | | |
| 5 | Do trait variation, garden climate, and their interaction predict growth increment independently of species ancestry? (Run separately for MAT and MAP as two garden climates) | lmer for continuous variables, not run for binary variables | growth increment ~ trait * garden climate + Pt + (1 \| MiniCG_Site/block) + (1 \| Genotype) |
| 6 | Do trait plasticity variation, garden climate, and their interaction predict growth increment independently of species ancestry? (Run separately for MAT and MAP as two garden climates) | lmer | growth increment ~ trait plasticity * garden climate + Pt + (1 \| MiniCG_Site/block) + (1 \| Genotype) |
