## Supplementary material for "Phenotypic plasticity evolved for climate variability constrains performance under climate warming": Analysis Rmarkdown Files: GxE_models_all_traits.html

Trait GxE Models


### Trait GxE Models

###### Alayna Mead

#### 2026-03-12

- 1 Setup
  - 1.1 Functions for plotting
- 2 Test for GxE effects on traits
  - 2.1 2023 data
    - 2.1.1 Setup
    - 2.1.2 Models
    - 2.1.3 Print model tables -
      2023
    - 2.1.4 Test daylength effects on fall
      phenology
  - 2.2 2024 data
    - 2.2.1 Setup
    - 2.2.2 Models
    - 2.2.3 Print model tables -
      2024
- 3 Summary heatmap

Tests for GxE effects on phenotypic traits. Unlike variance
partitioning script, here we test the effects of specific climate and
genetic variables.

Environment (E): garden MAT and MAP (for year of collection)

Genotype (G): Home MAT and MAP, genetic PCs 1-3

Genotype x Environment interaction (GxE): all possible combinations
of G and E variables above

### 1 Setup

```
library(ggplot2)
library(lme4)
```

```
## Loading required package: Matrix
```

```
library(sjPlot) # model plots
```

```
## Learn more about sjPlot with 'browseVignettes("sjPlot")'.
```

```
library(circlize) # colorramp2
```

```
## ========================================
## circlize version 0.4.16
## CRAN page: https://cran.r-project.org/package=circlize
## Github page: https://github.com/jokergoo/circlize
## Documentation: https://jokergoo.github.io/circlize_book/book/
## 
## If you use it in published research, please cite:
## Gu, Z. circlize implements and enhances circular visualization
##   in R. Bioinformatics 2014.
## 
## This message can be suppressed by:
##   suppressPackageStartupMessages(library(circlize))
## ========================================
```

```
library(ComplexHeatmap)
```

```
## Loading required package: grid
```

```
## ========================================
## ComplexHeatmap version 2.24.1
## Bioconductor page: http://bioconductor.org/packages/ComplexHeatmap/
## Github page: https://github.com/jokergoo/ComplexHeatmap
## Documentation: http://jokergoo.github.io/ComplexHeatmap-reference
## 
## If you use it in published research, please cite either one:
## - Gu, Z. Complex Heatmap Visualization. iMeta 2022.
## - Gu, Z. Complex heatmaps reveal patterns and correlations in multidimensional 
##     genomic data. Bioinformatics 2016.
## 
## 
## The new InteractiveComplexHeatmap package can directly export static 
## complex heatmaps into an interactive Shiny app with zero effort. Have a try!
## 
## This message can be suppressed by:
##   suppressPackageStartupMessages(library(ComplexHeatmap))
## ========================================
```

```
library(effectsize) # standardize_parameters to standardize model coefficients
library(car) # Anova function
```

```
## Loading required package: carData
```

```
library(knitr) # kable() for tables


# load data
load('data/clean/mini_garden_phenotypic_and_climate_data_2021-2024.Rdata')


# set negative heights to NA
dat$GrowthIncrement_2024[dat$GrowthIncrement_2024 < 0] <- NA
dat$GrowthIncrement_2023[dat$GrowthIncrement_2023 < 0] <- NA
dat$GrowthIncrement_2022[dat$GrowthIncrement_2022 < 0] <- NA
dat$GrowthIncrement_2021[dat$GrowthIncrement_2021 < 0] <- NA

# remove NA genotypes and those without genetic info
dat <- dat[! is.na(dat$Genotype),]
dat <- dat[! is.na(dat$k2_tricho),]


# print session info, including package versions
sessionInfo()
```

```
## R version 4.5.2 (2025-10-31)
## Platform: x86_64-pc-linux-gnu
## Running under: Arch Linux
## 
## Matrix products: default
## BLAS:   /usr/lib/libblas.so.3.12.0 
## LAPACK: /usr/lib/liblapack.so.3.12.0  LAPACK version 3.12.0
## 
## locale:
##  [1] LC_CTYPE=en_US.UTF-8       LC_NUMERIC=C              
##  [3] LC_TIME=en_US.UTF-8        LC_COLLATE=en_US.UTF-8    
##  [5] LC_MONETARY=en_US.UTF-8    LC_MESSAGES=en_US.UTF-8   
##  [7] LC_PAPER=en_US.UTF-8       LC_NAME=C                 
##  [9] LC_ADDRESS=C               LC_TELEPHONE=C            
## [11] LC_MEASUREMENT=en_US.UTF-8 LC_IDENTIFICATION=C       
## 
## time zone: US/Eastern
## tzcode source: system (glibc)
## 
## attached base packages:
## [1] grid      stats     graphics  grDevices datasets  utils     methods  
## [8] base     
## 
## other attached packages:
##  [1] knitr_1.50            car_3.1-3             carData_3.0-5        
##  [4] effectsize_1.0.1      ComplexHeatmap_2.24.1 circlize_0.4.16      
##  [7] sjPlot_2.8.17         lme4_1.1-37           Matrix_1.7-4         
## [10] ggplot2_3.5.2        
## 
## loaded via a namespace (and not attached):
##  [1] sjlabelled_1.2.0    tidyselect_1.2.1    dplyr_1.1.4        
##  [4] fastmap_1.2.0       bayestestR_0.17.0   sjstats_0.19.0     
##  [7] digest_0.6.37       estimability_1.5.1  lifecycle_1.0.4    
## [10] cluster_2.1.8.1     magrittr_2.0.3      compiler_4.5.2     
## [13] rlang_1.1.6         sass_0.4.10         tools_4.5.2        
## [16] yaml_2.3.10         RColorBrewer_1.1-3  abind_1.4-8        
## [19] withr_3.0.2         purrr_1.0.4         BiocGenerics_0.54.0
## [22] datawizard_1.2.0    stats4_4.5.2        xtable_1.8-4       
## [25] colorspace_2.1-1    emmeans_1.11.1      scales_1.3.0       
## [28] iterators_1.0.14    MASS_7.3-65         insight_1.4.2      
## [31] cli_3.6.5           mvtnorm_1.3-3       rmarkdown_2.29     
## [34] crayon_1.5.3        reformulas_0.4.0    generics_0.1.4     
## [37] rstudioapi_0.17.1   performance_0.15.1  rjson_0.2.23       
## [40] parameters_0.28.2   minqa_1.2.8         cachem_1.1.0       
## [43] splines_4.5.2       parallel_4.5.2      matrixStats_1.5.0  
## [46] vctrs_0.6.5         boot_1.3-32         jsonlite_2.0.0     
## [49] IRanges_2.42.0      GetoptLong_1.0.5    S4Vectors_0.46.0   
## [52] Formula_1.2-5       clue_0.3-66         foreach_1.5.2      
## [55] tidyr_1.3.1         jquerylib_0.1.4     glue_1.8.0         
## [58] nloptr_2.2.1        codetools_0.2-20    shape_1.4.6.1      
## [61] gtable_0.3.6        ggeffects_2.2.1     munsell_0.5.1      
## [64] tibble_3.2.1        pillar_1.10.2       htmltools_0.5.8.1  
## [67] R6_2.6.1            Rdpack_2.6.4        doParallel_1.0.17  
## [70] evaluate_1.0.3      lattice_0.22-7      rbibutils_2.3      
## [73] png_0.1-8           renv_0.17.3         bslib_0.9.0        
## [76] Rcpp_1.0.14         coda_0.19-4.1       nlme_3.1-168       
## [79] xfun_0.52           sjmisc_2.8.10       pkgconfig_2.0.3    
## [82] GlobalOptions_0.1.2
```

```
# markdown settings
knitr::opts_chunk$set(fig.width = 10, fig.height = 8)
```

```
# ggplot settings
theme_set(theme_bw(base_size = 16))
```

#### 1.1 Functions for plotting

```
# function to plot GxE interaction 
# plot genetics/climate vs trait, separate panel for each garden
plot_int <- function(trait, genetics, env, dat){
  
  dat$garden_sort <- reorder(dat$garden, dat[,env])
  
    p <- ggplot(data = dat, aes_string(x = genetics, y = trait, color = 'k2_tricho'))+
    geom_point(na.rm = TRUE) +
    scale_color_gradient2(high = "darkolivegreen2", mid = "grey20", low = "dodgerblue2", midpoint = 0.5, name = 'P. trichocarpa\nancestry', guide = 'none') +
    geom_smooth(method = lm, col = 'black', na.rm = TRUE)+
    facet_wrap(~garden_sort, drop = T) +
    ggtitle(paste(genetics, ' ~ ', trait,  '\n', 'gardens sorted by ', env, sep = ''))
  
  plot(p)
  
}


# plot variation in trait across gardens, sorting them by garden climate
# boxplot by garden
# use when the only variable is garden climate

plot_env <- function(trait, env, dat){
  
  dat$garden_sort <- reorder(dat$garden, dat[,env])
  #print(dat$garden_sort)
  
      p <- ggplot(data = dat, aes_string(x = 'garden_sort', y = trait))+
    geom_boxplot()+
    geom_jitter(na.rm = TRUE, aes(color = k2_tricho)) +
    scale_color_gradient2(high = "darkolivegreen2", mid = "grey20", low = "dodgerblue2", midpoint = 0.5, name = 'P. trichocarpa\nancestry', guide = 'none') +
    ggtitle(paste('garden ~ ', trait,  '\n', 'gardens sorted by ', env, sep = '')) +
      theme(axis.text.x = element_text(angle = 90, vjust = 0.5, hjust=1))
    
  plot(p)
  
}
```

### 2 Test for GxE effects on traits

Run each year separately, because we want to use the garden climate
for the year of collection

#### 2.1 2023 data

##### 2.1.1 Setup

```
###############################################

# all vars measured in 2023
vars <- c("DOY_Stage2_2023", "Stage2_cGDD_2023", "DOY_Stage3_2023", "Stage3_cGDD_2023",  "DOY_Stage6_2023", "daylength_Stage6_2023", "DOY_Stage7_2023", "daylength_Stage7_2023", "DOY_last_budset_2023", "daylength_last_budset_2023", "stage7_presence_2023", "growing_season_days_2023", 
             "leaf_thickness_avg_mm_2023", "leaf_area_cm2_2023", "leaf_mass_g_2023", "LMA_g_m2_2023", "lower_stomata_pore_length_mean_um", "upper_stomata_pore_length_mean_um", "lower_stomata_density_mm2", "upper_stomata_density_mm2", "stomata_ratio", "upper_stomata_presence", 
            "licor_gsw", "licor_gbw", "licor_ETR", "licor_Fs", "licor_Fm.", "licor_PhiPS2")

###############################################

# setup variables and put into a dataframe

# set climate and phenotype variables
garden_MAT <- dat$garden_MAT_2023
garden_MAT_2 <- dat$garden_MAT_2023^2
home_MAT <- dat$provenance_MAT
home_MAT_2 <- dat$provenance_MAT^2
garden_MAP <- dat$garden_MAP_2023
garden_MAP_2 <- dat$garden_MAP_2023^2
home_MAP <- dat$provenance_MAP
home_MAP_2 <- dat$provenance_MAP^2
home_lat <- dat$provenance_latitude
garden_lat <- dat$garden_Latitude

# random effects
block <- as.character(interaction(dat$MiniCG_Site, dat$block, drop = T))
#block <- as.character(dat$block)
Pt <- dat$Pt
k2_tricho <- dat$k2_tricho
genotype <- as.character(dat$Genotype)
garden <- as.character(dat$MiniCG_Site)
genetic_PC1 <- dat$genetic_PC1
genetic_PC2 <- dat$genetic_PC2
genetic_PC3 <- dat$genetic_PC3

# put in df
df <- data.frame(dat[,vars], garden_MAT, garden_MAT_2, home_MAT, home_MAT_2, garden_MAP, garden_MAP_2, home_MAP, home_MAP_2, home_lat, garden_lat, Pt, k2_tricho, genotype, garden, block, genetic_PC1, genetic_PC2, genetic_PC3)
rownames(df) <- dat$Unique_ID
str(df)
```

```
## 'data.frame':    1510 obs. of  46 variables:
##  $ DOY_Stage2_2023                  : num  97 101 93 101 93 93 101 101 93 97 ...
##  $ Stage2_cGDD_2023                 : num  496 531 472 531 472 ...
##  $ DOY_Stage3_2023                  : num  109 111 109 109 109 109 117 114 101 109 ...
##  $ Stage3_cGDD_2023                 : num  582 590 582 582 582 ...
##  $ DOY_Stage6_2023                  : num  181 163 181 163 162 NA 187 163 162 163 ...
##  $ daylength_Stage6_2023            : num  15.9 15.8 15.9 15.8 15.8 ...
##  $ DOY_Stage7_2023                  : num  NA 201 187 NA 194 194 NA NA NA NA ...
##  $ daylength_Stage7_2023            : num  NA 15.4 15.8 NA 15.6 ...
##  $ DOY_last_budset_2023             : num  181 215 201 163 240 NA 225 163 162 163 ...
##  $ daylength_last_budset_2023       : num  15.9 14.9 15.4 15.8 13.6 ...
##  $ stage7_presence_2023             : num  0 1 1 0 1 1 0 0 0 0 ...
##  $ growing_season_days_2023         : num  84 114 108 62 147 NA 124 62 69 66 ...
##  $ leaf_thickness_avg_mm_2023       : num  0.212 0.25 0.224 0.234 0.232 0.314 0.302 0.226 0.198 0.212 ...
##  $ leaf_area_cm2_2023               : num  NA NA NA NA NA NA NA NA NA NA ...
##  $ leaf_mass_g_2023                 : num  NA NA NA NA NA NA NA NA NA NA ...
##  $ LMA_g_m2_2023                    : num  NA NA NA NA NA NA NA NA NA NA ...
##  $ lower_stomata_pore_length_mean_um: num  NA NA NA NA NA NA NA NA NA NA ...
##  $ upper_stomata_pore_length_mean_um: num  NA NA NA NA NA NA NA NA NA NA ...
##  $ lower_stomata_density_mm2        : num  NA NA NA NA NA NA NA NA NA NA ...
##  $ upper_stomata_density_mm2        : num  NA NA NA NA NA NA NA NA NA NA ...
##  $ stomata_ratio                    : num  NA NA NA NA NA NA NA NA NA NA ...
##  $ upper_stomata_presence           : num  NA NA NA NA NA NA NA NA NA NA ...
##  $ licor_gsw                        : num  NA NA NA NA NA NA NA NA NA NA ...
##  $ licor_gbw                        : num  NA NA NA NA NA NA NA NA NA NA ...
##  $ licor_ETR                        : num  NA NA NA NA NA NA NA NA NA NA ...
##  $ licor_Fs                         : num  NA NA NA NA NA NA NA NA NA NA ...
##  $ licor_Fm.                        : num  NA NA NA NA NA NA NA NA NA NA ...
##  $ licor_PhiPS2                     : num  NA NA NA NA NA NA NA NA NA NA ...
##  $ garden_MAT                       : num  11.3 11.3 11.3 11.3 11.3 11.3 11.3 11.3 11.3 11.3 ...
##  $ garden_MAT_2                     : num  128 128 128 128 128 ...
##  $ home_MAT                         : num  2.3 2.3 3.8 3.8 5.6 5.6 3.9 3.9 1.2 1.2 ...
##  $ home_MAT_2                       : num  5.29 5.29 14.44 14.44 31.36 ...
##  $ garden_MAP                       : num  1249 1249 1249 1249 1249 ...
##  $ garden_MAP_2                     : num  1560001 1560001 1560001 1560001 1560001 ...
##  $ home_MAP                         : int  424 424 320 320 319 319 605 605 1154 1154 ...
##  $ home_MAP_2                       : num  179776 179776 102400 102400 101761 ...
##  $ home_lat                         : num  51.9 51.9 52.1 52.1 52 ...
##  $ garden_lat                       : num  47.1 47.1 47.1 47.1 47.1 ...
##  $ Pt                               : num  0.571 0.571 0.908 0.908 0.864 ...
##  $ k2_tricho                        : num  0.573 0.573 0.903 0.903 0.859 ...
##  $ genotype                         : chr  "206" "206" "210" "210" ...
##  $ garden                           : chr  "EVERGREEN" "EVERGREEN" "EVERGREEN" "EVERGREEN" ...
##  $ block                            : chr  "EVERGREEN.1" "EVERGREEN.2" "EVERGREEN.1" "EVERGREEN.2" ...
##  $ genetic_PC1                      : num  0.00423 0.00423 -0.03207 -0.03207 -0.02393 ...
##  $ genetic_PC2                      : num  0.02407 0.02407 -0.00722 -0.00722 0.01811 ...
##  $ genetic_PC3                      : num  -0.0243 -0.0243 -0.0365 -0.0365 -0.0505 ...
```

##### 2.1.2 Models

```
# run full model for all variables
vars <- c("DOY_Stage2_2023", "Stage2_cGDD_2023", "DOY_Stage3_2023", "Stage3_cGDD_2023",  "DOY_Stage6_2023", "DOY_Stage7_2023", "DOY_last_budset_2023", "stage7_presence_2023", "growing_season_days_2023", 
             "leaf_thickness_avg_mm_2023", "leaf_area_cm2_2023", "leaf_mass_g_2023", "LMA_g_m2_2023", "lower_stomata_pore_length_mean_um", "upper_stomata_pore_length_mean_um", "lower_stomata_density_mm2", "upper_stomata_density_mm2", "stomata_ratio", "upper_stomata_presence", 
            "licor_gsw", "licor_gbw", "licor_ETR", "licor_Fs", "licor_Fm.", "licor_PhiPS2")


dput(vars)
```

```
## c("DOY_Stage2_2023", "Stage2_cGDD_2023", "DOY_Stage3_2023", "Stage3_cGDD_2023", 
## "DOY_Stage6_2023", "DOY_Stage7_2023", "DOY_last_budset_2023", 
## "stage7_presence_2023", "growing_season_days_2023", "leaf_thickness_avg_mm_2023", 
## "leaf_area_cm2_2023", "leaf_mass_g_2023", "LMA_g_m2_2023", "lower_stomata_pore_length_mean_um", 
## "upper_stomata_pore_length_mean_um", "lower_stomata_density_mm2", 
## "upper_stomata_density_mm2", "stomata_ratio", "upper_stomata_presence", 
## "licor_gsw", "licor_gbw", "licor_ETR", "licor_Fs", "licor_Fm.", 
## "licor_PhiPS2")
```

```
# the binary variables won't converge
# Error: (maxstephalfit) PIRLS step-halvings failed to reduce deviance in pwrssUpdate

vars <- c("DOY_Stage2_2023", "Stage2_cGDD_2023", "DOY_Stage3_2023", "Stage3_cGDD_2023",  "DOY_Stage6_2023", "daylength_Stage6_2023", "DOY_Stage7_2023", "daylength_Stage7_2023", "DOY_last_budset_2023", "daylength_last_budset_2023",  "growing_season_days_2023", 
             "leaf_thickness_avg_mm_2023", "leaf_area_cm2_2023", "leaf_mass_g_2023", "LMA_g_m2_2023", "lower_stomata_pore_length_mean_um", "upper_stomata_pore_length_mean_um", "lower_stomata_density_mm2", "upper_stomata_density_mm2", "stomata_ratio", 
            "licor_gsw", "licor_gbw", "licor_ETR", "licor_Fs", "licor_Fm.", "licor_PhiPS2")

# binary variables
vars_bin <- c( "stage7_presence_2023", "upper_stomata_presence")

mods.23 <- list()

for(n in 1:length(vars)){
  
  var <- vars[n]
  
  print(paste('Model for:', var))
  
  if(var %in% vars_bin){

    # these models failed to converge - don't run here
    # mod <- glm(paste0(var, '~ garden_MAT*home_MAT + garden_MAT*home_MAP +   garden_MAP*home_MAP + garden_MAP*home_MAT + genetic_PC1*garden_MAT + genetic_PC2*garden_MAT + genetic_PC3*garden_MAT +  genetic_PC1*garden_MAP + genetic_PC2*garden_MAP + genetic_PC3*garden_MAP + (1 | genotype) + (1 | garden/block) '),
    #             data = df,
    #             family = binomial()
    # )


  } else {
    #  full model with just linear interaction effects
    mod <- lmer(paste0(var, '~ garden_MAT*home_MAT + garden_MAT*home_MAP +   garden_MAP*home_MAP + garden_MAP*home_MAT + genetic_PC1*garden_MAT + genetic_PC2*garden_MAT + genetic_PC3*garden_MAT +  genetic_PC1*garden_MAP + genetic_PC2*garden_MAP + genetic_PC3*garden_MAP + (1 | genotype) + (1 | garden/block)'),
                data = df
    )
  }
    summary(mod)
    mod.info <- car::Anova(mod)
    
    print(mod.info)
    
    # save to list
    mods.23[[n]] <- mod
    
    # plot model effects
    p <- plot_model(mod, type = 'std', vline.color = "black", show.values = T, title = var)
    plot(p)
    
    # plots for significant factors
    pvals <- mod.info$`Pr(>Chisq)`
    if(min(pvals) <= 0.05){
      
      # get significant factors
      sig <- rownames(mod.info)[pvals <= 0.05]
      
      # plot for each significant factor
      for(f in 1:length(sig)){
        
        y <- var # y axis is always the trait
        
        # if it is an interaction effect, we need to include both variables in the plot
        if(grepl(':', sig[f])){
          
          # get two variables in interaction
          interaction_vars <- strsplit(sig[f], ':')[[1]]
          
          # env will start with 'garden' because it's always garden climate
          env <- grep('garden', interaction_vars, value = T)
          # the one without garden is the genetic effect, either PCs or home clim
          x <- grep('garden', interaction_vars, value = T, invert = T)
          
          p2 <- plot_int(trait = y, genetics = x, env = env, dat = df)
          #plot(p2)
          
          # if just a single variable, still plot gardens separately, but sort by MAT by default
        } else{
          
          # if single variable is garden climate, use that and plot boxplots
          if(grepl('garden', sig[f])){
            env <- sig[f]
            plot_env(trait = y, env = env, dat = df)
            
          # if single variable is a genotype characteristic, facet by garden sorted by MAT  
          } else {
            env <- 'garden_MAT'
          x <- sig[f]
           p2 <- plot_int(trait = y, genetics = x, env = env, dat = df)
          #plot(p2)
          }
        }
      }
    }
}
```

```
## [1] "Model for: DOY_Stage2_2023"
```

```
## Warning: Some predictor variables are on very different scales: consider
## rescaling
```

```
## Analysis of Deviance Table (Type II Wald chisquare tests)
## 
## Response: DOY_Stage2_2023
##                          Chisq Df Pr(>Chisq)    
## garden_MAT             15.6181  1  7.751e-05 ***
## home_MAT                4.5723  1    0.03249 *  
## home_MAP                1.7930  1    0.18056    
## garden_MAP              1.3166  1    0.25119    
## genetic_PC1             0.0002  1    0.98972    
## genetic_PC2             0.0564  1    0.81220    
## genetic_PC3             0.0008  1    0.97805    
## garden_MAT:home_MAT     2.7267  1    0.09868 .  
## garden_MAT:home_MAP     0.1488  1    0.69964    
## home_MAP:garden_MAP     1.6976  1    0.19260    
## home_MAT:garden_MAP     4.0913  1    0.04311 *  
## garden_MAT:genetic_PC1  0.5627  1    0.45316    
## garden_MAT:genetic_PC2  0.1474  1    0.70099    
## garden_MAT:genetic_PC3  0.0001  1    0.99163    
## garden_MAP:genetic_PC1  0.1705  1    0.67970    
## garden_MAP:genetic_PC2  0.0170  1    0.89622    
## garden_MAP:genetic_PC3  0.4068  1    0.52358    
## ---
## Signif. codes:  0 '***' 0.001 '**' 0.01 '*' 0.05 '.' 0.1 ' ' 1
```

```
## Warning: `aes_string()` was deprecated in ggplot2 3.0.0.
## ℹ Please use tidy evaluation idioms with `aes()`.
## ℹ See also `vignette("ggplot2-in-packages")` for more information.
## This warning is displayed once every 8 hours.
## Call `lifecycle::last_lifecycle_warnings()` to see where this warning was
## generated.
```

```
## Warning: Removed 744 rows containing non-finite outside the scale range
## (`stat_boxplot()`).
```

```
## `geom_smooth()` using formula = 'y ~ x'
```

```
## `geom_smooth()` using formula = 'y ~ x'
```

```
## [1] "Model for: Stage2_cGDD_2023"
```

```
## Warning: Some predictor variables are on very different scales: consider
## rescaling
```

```
## Analysis of Deviance Table (Type II Wald chisquare tests)
## 
## Response: Stage2_cGDD_2023
##                          Chisq Df Pr(>Chisq)    
## garden_MAT             15.0285  1  0.0001059 ***
## home_MAT                3.2836  1  0.0699733 .  
## home_MAP                1.7113  1  0.1908112    
## garden_MAP              0.0013  1  0.9706956    
## genetic_PC1             0.0263  1  0.8711757    
## genetic_PC2             0.0129  1  0.9094893    
## genetic_PC3             0.0198  1  0.8879893    
## garden_MAT:home_MAT     1.7531  1  0.1854829    
## garden_MAT:home_MAP     0.0061  1  0.9379771    
## home_MAP:garden_MAP     0.8093  1  0.3683306    
## home_MAT:garden_MAP     0.4989  1  0.4799904    
## garden_MAT:genetic_PC1  0.7233  1  0.3950741    
## garden_MAT:genetic_PC2  0.4496  1  0.5025452    
## garden_MAT:genetic_PC3  0.0086  1  0.9262152    
## garden_MAP:genetic_PC1  0.0408  1  0.8399617    
## garden_MAP:genetic_PC2  0.0717  1  0.7888604    
## garden_MAP:genetic_PC3  0.1041  1  0.7470203    
## ---
## Signif. codes:  0 '***' 0.001 '**' 0.01 '*' 0.05 '.' 0.1 ' ' 1
```

```
## Warning: Removed 744 rows containing non-finite outside the scale range
## (`stat_boxplot()`).
```

```
## [1] "Model for: DOY_Stage3_2023"
```

```
## Warning: Some predictor variables are on very different scales: consider
## rescaling
```

```
## Analysis of Deviance Table (Type II Wald chisquare tests)
## 
## Response: DOY_Stage3_2023
##                          Chisq Df Pr(>Chisq)    
## garden_MAT             18.8581  1  1.408e-05 ***
## home_MAT                5.2180  1    0.02235 *  
## home_MAP                0.9917  1    0.31932    
## garden_MAP              2.2113  1    0.13701    
## genetic_PC1             0.0335  1    0.85486    
## genetic_PC2             0.0551  1    0.81439    
## genetic_PC3             0.0490  1    0.82480    
## garden_MAT:home_MAT     0.4129  1    0.52049    
## garden_MAT:home_MAP     0.0290  1    0.86470    
## home_MAP:garden_MAP     0.1838  1    0.66813    
## home_MAT:garden_MAP     1.8251  1    0.17671    
## garden_MAT:genetic_PC1  0.8493  1    0.35674    
## garden_MAT:genetic_PC2  1.3648  1    0.24271    
## garden_MAT:genetic_PC3  0.0477  1    0.82704    
## garden_MAP:genetic_PC1  0.3450  1    0.55693    
## garden_MAP:genetic_PC2  0.1608  1    0.68847    
## garden_MAP:genetic_PC3  0.3950  1    0.52967    
## ---
## Signif. codes:  0 '***' 0.001 '**' 0.01 '*' 0.05 '.' 0.1 ' ' 1
```

```
## Warning: Removed 653 rows containing non-finite outside the scale range
## (`stat_boxplot()`).
```

```
## `geom_smooth()` using formula = 'y ~ x'
```

```
## [1] "Model for: Stage3_cGDD_2023"
```

```
## Warning: Some predictor variables are on very different scales: consider
## rescaling
```

```
## Analysis of Deviance Table (Type II Wald chisquare tests)
## 
## Response: Stage3_cGDD_2023
##                          Chisq Df Pr(>Chisq)    
## garden_MAT             21.6355  1  3.297e-06 ***
## home_MAT                3.7743  1    0.05205 .  
## home_MAP                0.9571  1    0.32793    
## garden_MAP              0.0006  1    0.98008    
## genetic_PC1             0.0358  1    0.84996    
## genetic_PC2             0.0820  1    0.77465    
## genetic_PC3             0.0573  1    0.81081    
## garden_MAT:home_MAT     1.7189  1    0.18983    
## garden_MAT:home_MAP     0.0429  1    0.83598    
## home_MAP:garden_MAP     0.1188  1    0.73036    
## home_MAT:garden_MAP     1.1817  1    0.27701    
## garden_MAT:genetic_PC1  0.2407  1    0.62368    
## garden_MAT:genetic_PC2  1.3926  1    0.23797    
## garden_MAT:genetic_PC3  0.2296  1    0.63182    
## garden_MAP:genetic_PC1  0.1970  1    0.65712    
## garden_MAP:genetic_PC2  0.3629  1    0.54692    
## garden_MAP:genetic_PC3  0.2479  1    0.61857    
## ---
## Signif. codes:  0 '***' 0.001 '**' 0.01 '*' 0.05 '.' 0.1 ' ' 1
```

```
## Warning: Removed 653 rows containing non-finite outside the scale range
## (`stat_boxplot()`).
```

```
## [1] "Model for: DOY_Stage6_2023"
```

```
## Warning: Some predictor variables are on very different scales: consider
## rescaling
```

```
## Analysis of Deviance Table (Type II Wald chisquare tests)
## 
## Response: DOY_Stage6_2023
##                          Chisq Df Pr(>Chisq)    
## garden_MAT              2.1695  1  0.1407721    
## home_MAT                8.4413  1  0.0036678 ** 
## home_MAP                1.1274  1  0.2883398    
## garden_MAP              0.2076  1  0.6486269    
## genetic_PC1             5.1107  1  0.0237789 *  
## genetic_PC2             1.2923  1  0.2556286    
## genetic_PC3             1.2103  1  0.2712727    
## garden_MAT:home_MAT     4.2218  1  0.0399084 *  
## garden_MAT:home_MAP     0.9939  1  0.3187890    
## home_MAP:garden_MAP     3.4926  1  0.0616440 .  
## home_MAT:garden_MAP     8.7803  1  0.0030449 ** 
## garden_MAT:genetic_PC1  2.4587  1  0.1168755    
## garden_MAT:genetic_PC2 11.7425  1  0.0006109 ***
## garden_MAT:genetic_PC3  1.3184  1  0.2508847    
## garden_MAP:genetic_PC1  0.1464  1  0.7020116    
## garden_MAP:genetic_PC2  4.3821  1  0.0363177 *  
## garden_MAP:genetic_PC3  1.0643  1  0.3022436    
## ---
## Signif. codes:  0 '***' 0.001 '**' 0.01 '*' 0.05 '.' 0.1 ' ' 1
```

```
## `geom_smooth()` using formula = 'y ~ x'
```

```
## `geom_smooth()` using formula = 'y ~ x'
```

```
## `geom_smooth()` using formula = 'y ~ x'
```

```
## `geom_smooth()` using formula = 'y ~ x'
```

```
## `geom_smooth()` using formula = 'y ~ x'
```

```
## `geom_smooth()` using formula = 'y ~ x'
```

```
## [1] "Model for: daylength_Stage6_2023"
```

```
## Warning: Some predictor variables are on very different scales: consider
## rescaling
```

```
## Analysis of Deviance Table (Type II Wald chisquare tests)
## 
## Response: daylength_Stage6_2023
##                         Chisq Df Pr(>Chisq)   
## garden_MAT             0.0086  1   0.926126   
## home_MAT               2.6900  1   0.100980   
## home_MAP               0.4396  1   0.507303   
## garden_MAP             0.5046  1   0.477468   
## genetic_PC1            0.1157  1   0.733738   
## genetic_PC2            1.9907  1   0.158266   
## genetic_PC3            0.2023  1   0.652837   
## garden_MAT:home_MAT    7.0321  1   0.008006 **
## garden_MAT:home_MAP    0.0911  1   0.762840   
## home_MAP:garden_MAP    0.7040  1   0.401443   
## home_MAT:garden_MAP    1.5594  1   0.211760   
## garden_MAT:genetic_PC1 1.3125  1   0.251951   
## garden_MAT:genetic_PC2 5.4127  1   0.019990 * 
## garden_MAT:genetic_PC3 0.1186  1   0.730517   
## garden_MAP:genetic_PC1 0.1006  1   0.751120   
## garden_MAP:genetic_PC2 0.4712  1   0.492420   
## garden_MAP:genetic_PC3 0.0993  1   0.752714   
## ---
## Signif. codes:  0 '***' 0.001 '**' 0.01 '*' 0.05 '.' 0.1 ' ' 1
```

```
## `geom_smooth()` using formula = 'y ~ x'
```

```
## `geom_smooth()` using formula = 'y ~ x'
```

```
## [1] "Model for: DOY_Stage7_2023"
```

```
## Warning: Some predictor variables are on very different scales: consider
## rescaling
```

```
## boundary (singular) fit: see help('isSingular')
```

```
## Analysis of Deviance Table (Type II Wald chisquare tests)
## 
## Response: DOY_Stage7_2023
##                         Chisq Df Pr(>Chisq)  
## garden_MAT             0.1187  1    0.73048  
## home_MAT               0.3197  1    0.57177  
## home_MAP               0.6849  1    0.40791  
## garden_MAP             1.3629  1    0.24304  
## genetic_PC1            0.5305  1    0.46640  
## genetic_PC2            1.2173  1    0.26990  
## genetic_PC3            2.4352  1    0.11864  
## garden_MAT:home_MAT    0.7153  1    0.39769  
## garden_MAT:home_MAP    1.6015  1    0.20569  
## home_MAP:garden_MAP    1.4438  1    0.22953  
## home_MAT:garden_MAP    3.7608  1    0.05247 .
## garden_MAT:genetic_PC1 0.0223  1    0.88126  
## garden_MAT:genetic_PC2 1.2813  1    0.25766  
## garden_MAT:genetic_PC3 0.0981  1    0.75411  
## garden_MAP:genetic_PC1 1.1238  1    0.28910  
## garden_MAP:genetic_PC2 0.2163  1    0.64189  
## garden_MAP:genetic_PC3 0.0797  1    0.77768  
## ---
## Signif. codes:  0 '***' 0.001 '**' 0.01 '*' 0.05 '.' 0.1 ' ' 1
```

```
## boundary (singular) fit: see help('isSingular')
```

```
## [1] "Model for: daylength_Stage7_2023"
```

```
## Warning: Some predictor variables are on very different scales: consider
## rescaling
```

```
## boundary (singular) fit: see help('isSingular')
```

```
## Analysis of Deviance Table (Type II Wald chisquare tests)
## 
## Response: daylength_Stage7_2023
##                         Chisq Df Pr(>Chisq)  
## garden_MAT             1.3213  1    0.25035  
## home_MAT               0.0896  1    0.76464  
## home_MAP               0.0010  1    0.97476  
## garden_MAP             1.5021  1    0.22034  
## genetic_PC1            0.3934  1    0.53054  
## genetic_PC2            0.2722  1    0.60183  
## genetic_PC3            1.4469  1    0.22903  
## garden_MAT:home_MAT    0.9852  1    0.32093  
## garden_MAT:home_MAP    0.9120  1    0.33957  
## home_MAP:garden_MAP    0.8455  1    0.35781  
## home_MAT:garden_MAP    3.3163  1    0.06860 .
## garden_MAT:genetic_PC1 0.2749  1    0.60004  
## garden_MAT:genetic_PC2 0.8171  1    0.36602  
## garden_MAT:genetic_PC3 0.0552  1    0.81429  
## garden_MAP:genetic_PC1 3.7801  1    0.05186 .
## garden_MAP:genetic_PC2 0.0638  1    0.80051  
## garden_MAP:genetic_PC3 0.0001  1    0.99264  
## ---
## Signif. codes:  0 '***' 0.001 '**' 0.01 '*' 0.05 '.' 0.1 ' ' 1
```

```
## boundary (singular) fit: see help('isSingular')
```

```
## [1] "Model for: DOY_last_budset_2023"
```

```
## Warning: Some predictor variables are on very different scales: consider
## rescaling
```

```
## Analysis of Deviance Table (Type II Wald chisquare tests)
## 
## Response: DOY_last_budset_2023
##                         Chisq Df Pr(>Chisq)  
## garden_MAT             0.1227  1    0.72613  
## home_MAT               5.1111  1    0.02377 *
## home_MAP               1.8245  1    0.17678  
## garden_MAP             1.0005  1    0.31718  
## genetic_PC1            2.8377  1    0.09208 .
## genetic_PC2            0.9576  1    0.32780  
## genetic_PC3            0.7347  1    0.39138  
## garden_MAT:home_MAT    1.4718  1    0.22506  
## garden_MAT:home_MAP    0.2342  1    0.62844  
## home_MAP:garden_MAP    0.0008  1    0.97716  
## home_MAT:garden_MAP    2.0026  1    0.15703  
## garden_MAT:genetic_PC1 0.4015  1    0.52634  
## garden_MAT:genetic_PC2 1.1941  1    0.27450  
## garden_MAT:genetic_PC3 0.8570  1    0.35457  
## garden_MAP:genetic_PC1 0.6583  1    0.41717  
## garden_MAP:genetic_PC2 0.5846  1    0.44451  
## garden_MAP:genetic_PC3 0.3782  1    0.53856  
## ---
## Signif. codes:  0 '***' 0.001 '**' 0.01 '*' 0.05 '.' 0.1 ' ' 1
```

```
## `geom_smooth()` using formula = 'y ~ x'
```

```
## [1] "Model for: daylength_last_budset_2023"
```

```
## Warning: Some predictor variables are on very different scales: consider
## rescaling
```

```
## Analysis of Deviance Table (Type II Wald chisquare tests)
## 
## Response: daylength_last_budset_2023
##                         Chisq Df Pr(>Chisq)  
## garden_MAT             0.1618  1    0.68749  
## home_MAT               2.8446  1    0.09168 .
## home_MAP               1.6851  1    0.19425  
## garden_MAP             0.5440  1    0.46080  
## genetic_PC1            0.7412  1    0.38928  
## genetic_PC2            2.4795  1    0.11534  
## genetic_PC3            0.0039  1    0.94996  
## garden_MAT:home_MAT    0.0611  1    0.80475  
## garden_MAT:home_MAP    0.5124  1    0.47409  
## home_MAP:garden_MAP    0.0599  1    0.80666  
## home_MAT:garden_MAP    0.1247  1    0.72395  
## garden_MAT:genetic_PC1 0.0040  1    0.94948  
## garden_MAT:genetic_PC2 0.9783  1    0.32261  
## garden_MAT:genetic_PC3 3.1850  1    0.07432 .
## garden_MAP:genetic_PC1 0.0294  1    0.86389  
## garden_MAP:genetic_PC2 0.0697  1    0.79182  
## garden_MAP:genetic_PC3 0.6442  1    0.42220  
## ---
## Signif. codes:  0 '***' 0.001 '**' 0.01 '*' 0.05 '.' 0.1 ' ' 1
```

```
## [1] "Model for: growing_season_days_2023"
```

```
## Warning: Some predictor variables are on very different scales: consider
## rescaling
```

```
## Analysis of Deviance Table (Type II Wald chisquare tests)
## 
## Response: growing_season_days_2023
##                         Chisq Df Pr(>Chisq)  
## garden_MAT             0.7852  1    0.37556  
## home_MAT               0.7764  1    0.37823  
## home_MAP               3.5677  1    0.05891 .
## garden_MAP             0.7147  1    0.39788  
## genetic_PC1            1.4144  1    0.23433  
## genetic_PC2            2.2045  1    0.13761  
## genetic_PC3            0.3925  1    0.53097  
## garden_MAT:home_MAT    2.3254  1    0.12728  
## garden_MAT:home_MAP    0.3501  1    0.55403  
## home_MAP:garden_MAP    0.0558  1    0.81326  
## home_MAT:garden_MAP    2.4136  1    0.12029  
## garden_MAT:genetic_PC1 0.3031  1    0.58193  
## garden_MAT:genetic_PC2 0.5332  1    0.46526  
## garden_MAT:genetic_PC3 2.1125  1    0.14610  
## garden_MAP:genetic_PC1 0.0871  1    0.76785  
## garden_MAP:genetic_PC2 0.0464  1    0.82948  
## garden_MAP:genetic_PC3 2.4448  1    0.11792  
## ---
## Signif. codes:  0 '***' 0.001 '**' 0.01 '*' 0.05 '.' 0.1 ' ' 1
```

```
## [1] "Model for: leaf_thickness_avg_mm_2023"
```

```
## Warning: Some predictor variables are on very different scales: consider
## rescaling
```

```
## Analysis of Deviance Table (Type II Wald chisquare tests)
## 
## Response: leaf_thickness_avg_mm_2023
##                         Chisq Df Pr(>Chisq)  
## garden_MAT             0.5211  1    0.47037  
## home_MAT               0.9432  1    0.33147  
## home_MAP               0.4633  1    0.49608  
## garden_MAP             0.0850  1    0.77059  
## genetic_PC1            0.0007  1    0.97918  
## genetic_PC2            0.4203  1    0.51679  
## genetic_PC3            0.2037  1    0.65171  
## garden_MAT:home_MAT    0.0300  1    0.86259  
## garden_MAT:home_MAP    2.3063  1    0.12885  
## home_MAP:garden_MAP    4.2376  1    0.03954 *
## home_MAT:garden_MAP    0.6888  1    0.40656  
## garden_MAT:genetic_PC1 2.0299  1    0.15423  
## garden_MAT:genetic_PC2 0.0999  1    0.75199  
## garden_MAT:genetic_PC3 0.1400  1    0.70829  
## garden_MAP:genetic_PC1 0.1938  1    0.65976  
## garden_MAP:genetic_PC2 0.2629  1    0.60815  
## garden_MAP:genetic_PC3 0.5728  1    0.44915  
## ---
## Signif. codes:  0 '***' 0.001 '**' 0.01 '*' 0.05 '.' 0.1 ' ' 1
```

```
## `geom_smooth()` using formula = 'y ~ x'
```

```
## [1] "Model for: leaf_area_cm2_2023"
```

```
## Warning: Some predictor variables are on very different scales: consider
## rescaling
```

```
## Analysis of Deviance Table (Type II Wald chisquare tests)
## 
## Response: leaf_area_cm2_2023
##                          Chisq Df Pr(>Chisq)    
## garden_MAT             10.8221  1  0.0010030 ** 
## home_MAT                2.6525  1  0.1033898    
## home_MAP                0.9744  1  0.3235834    
## garden_MAP             11.4428  1  0.0007177 ***
## genetic_PC1             3.8607  1  0.0494286 *  
## genetic_PC2             1.8014  1  0.1795381    
## genetic_PC3             0.1461  1  0.7022888    
## garden_MAT:home_MAT    19.3282  1  1.101e-05 ***
## garden_MAT:home_MAP     0.0488  1  0.8251245    
## home_MAP:garden_MAP     1.7882  1  0.1811478    
## home_MAT:garden_MAP    13.2209  1  0.0002768 ***
## garden_MAT:genetic_PC1  4.2131  1  0.0401139 *  
## garden_MAT:genetic_PC2 15.4241  1  8.588e-05 ***
## garden_MAT:genetic_PC3  3.5559  1  0.0593343 .  
## garden_MAP:genetic_PC1 13.9318  1  0.0001896 ***
## garden_MAP:genetic_PC2 11.2809  1  0.0007831 ***
## garden_MAP:genetic_PC3  6.5454  1  0.0105155 *  
## ---
## Signif. codes:  0 '***' 0.001 '**' 0.01 '*' 0.05 '.' 0.1 ' ' 1
```

```
## Warning: Removed 807 rows containing non-finite outside the scale range
## (`stat_boxplot()`).
```

```
## Warning: Removed 807 rows containing non-finite outside the scale range
## (`stat_boxplot()`).
```

```
## `geom_smooth()` using formula = 'y ~ x'
```

```
## `geom_smooth()` using formula = 'y ~ x'
```

```
## `geom_smooth()` using formula = 'y ~ x'
```

```
## `geom_smooth()` using formula = 'y ~ x'
```

```
## `geom_smooth()` using formula = 'y ~ x'
```

```
## `geom_smooth()` using formula = 'y ~ x'
```

```
## `geom_smooth()` using formula = 'y ~ x'
```

```
## `geom_smooth()` using formula = 'y ~ x'
```

```
## [1] "Model for: leaf_mass_g_2023"
```

```
## Warning: Some predictor variables are on very different scales: consider
## rescaling
```

```
## Analysis of Deviance Table (Type II Wald chisquare tests)
## 
## Response: leaf_mass_g_2023
##                          Chisq Df Pr(>Chisq)    
## garden_MAT             12.3720  1  0.0004358 ***
## home_MAT                3.9393  1  0.0471714 *  
## home_MAP                1.6481  1  0.1992121    
## garden_MAP             13.9465  1  0.0001881 ***
## genetic_PC1             2.7377  1  0.0980057 .  
## genetic_PC2             1.6954  1  0.1928941    
## genetic_PC3             0.6669  1  0.4141330    
## garden_MAT:home_MAT    14.8019  1  0.0001194 ***
## garden_MAT:home_MAP     0.5265  1  0.4680885    
## home_MAP:garden_MAP     3.2492  1  0.0714574 .  
## home_MAT:garden_MAP     6.9971  1  0.0081641 ** 
## garden_MAT:genetic_PC1  2.6083  1  0.1063031    
## garden_MAT:genetic_PC2 11.9494  1  0.0005467 ***
## garden_MAT:genetic_PC3  2.5013  1  0.1137523    
## garden_MAP:genetic_PC1 17.4396  1  2.966e-05 ***
## garden_MAP:genetic_PC2  7.2469  1  0.0071025 ** 
## garden_MAP:genetic_PC3  3.3505  1  0.0671855 .  
## ---
## Signif. codes:  0 '***' 0.001 '**' 0.01 '*' 0.05 '.' 0.1 ' ' 1
```

```
## Warning: Removed 801 rows containing non-finite outside the scale range
## (`stat_boxplot()`).
```

```
## `geom_smooth()` using formula = 'y ~ x'
```

```
## Warning: Removed 801 rows containing non-finite outside the scale range
## (`stat_boxplot()`).
```

```
## `geom_smooth()` using formula = 'y ~ x'
```

```
## `geom_smooth()` using formula = 'y ~ x'
```

```
## `geom_smooth()` using formula = 'y ~ x'
```

```
## `geom_smooth()` using formula = 'y ~ x'
```

```
## `geom_smooth()` using formula = 'y ~ x'
```

```
## [1] "Model for: LMA_g_m2_2023"
```

```
## Warning: Some predictor variables are on very different scales: consider
## rescaling
```

```
## Analysis of Deviance Table (Type II Wald chisquare tests)
## 
## Response: LMA_g_m2_2023
##                         Chisq Df Pr(>Chisq)  
## garden_MAT             0.0437  1    0.83433  
## home_MAT               2.0519  1    0.15201  
## home_MAP               0.1133  1    0.73638  
## garden_MAP             1.4999  1    0.22068  
## genetic_PC1            0.1723  1    0.67810  
## genetic_PC2            0.0291  1    0.86446  
## genetic_PC3            0.3508  1    0.55366  
## garden_MAT:home_MAT    0.2800  1    0.59671  
## garden_MAT:home_MAP    0.0927  1    0.76083  
## home_MAP:garden_MAP    0.1151  1    0.73440  
## home_MAT:garden_MAP    5.3299  1    0.02096 *
## garden_MAT:genetic_PC1 3.6124  1    0.05735 .
## garden_MAT:genetic_PC2 0.7030  1    0.40179  
## garden_MAT:genetic_PC3 0.5154  1    0.47282  
## garden_MAP:genetic_PC1 0.6726  1    0.41214  
## garden_MAP:genetic_PC2 0.3685  1    0.54385  
## garden_MAP:genetic_PC3 1.2406  1    0.26535  
## ---
## Signif. codes:  0 '***' 0.001 '**' 0.01 '*' 0.05 '.' 0.1 ' ' 1
```

```
## `geom_smooth()` using formula = 'y ~ x'
```

```
## [1] "Model for: lower_stomata_pore_length_mean_um"
```

```
## Warning: Some predictor variables are on very different scales: consider
## rescaling
```

```
## Analysis of Deviance Table (Type II Wald chisquare tests)
## 
## Response: lower_stomata_pore_length_mean_um
##                          Chisq Df Pr(>Chisq)    
## garden_MAT              0.4288  1  0.5125922    
## home_MAT                0.1390  1  0.7093118    
## home_MAP                1.2498  1  0.2635864    
## garden_MAP              0.4377  1  0.5082344    
## genetic_PC1             1.6832  1  0.1944999    
## genetic_PC2            11.9153  1  0.0005568 ***
## genetic_PC3             0.8272  1  0.3630699    
## garden_MAT:home_MAT     0.5976  1  0.4395015    
## garden_MAT:home_MAP     0.0409  1  0.8397944    
## home_MAP:garden_MAP     0.2655  1  0.6063960    
## home_MAT:garden_MAP     1.9383  1  0.1638511    
## garden_MAT:genetic_PC1  0.1591  1  0.6900014    
## garden_MAT:genetic_PC2  1.7627  1  0.1842864    
## garden_MAT:genetic_PC3  0.0717  1  0.7888247    
## garden_MAP:genetic_PC1  0.3873  1  0.5337128    
## garden_MAP:genetic_PC2  5.2805  1  0.0215659 *  
## garden_MAP:genetic_PC3  0.9859  1  0.3207444    
## ---
## Signif. codes:  0 '***' 0.001 '**' 0.01 '*' 0.05 '.' 0.1 ' ' 1
```

```
## `geom_smooth()` using formula = 'y ~ x'
```

```
## `geom_smooth()` using formula = 'y ~ x'
```

```
## [1] "Model for: upper_stomata_pore_length_mean_um"
```

```
## Warning: Some predictor variables are on very different scales: consider
## rescaling
```

```
## Analysis of Deviance Table (Type II Wald chisquare tests)
## 
## Response: upper_stomata_pore_length_mean_um
##                          Chisq Df Pr(>Chisq)    
## garden_MAT              0.3203  1  0.5714425    
## home_MAT                1.8620  1  0.1723973    
## home_MAP               13.5863  1  0.0002278 ***
## garden_MAP              0.0000  1  0.9977443    
## genetic_PC1             0.4861  1  0.4856909    
## genetic_PC2             2.8775  1  0.0898250 .  
## genetic_PC3             0.3420  1  0.5586681    
## garden_MAT:home_MAT     0.2738  1  0.6007716    
## garden_MAT:home_MAP     0.0125  1  0.9108846    
## home_MAP:garden_MAP     0.0020  1  0.9642787    
## home_MAT:garden_MAP     0.0817  1  0.7750444    
## garden_MAT:genetic_PC1  0.4681  1  0.4938502    
## garden_MAT:genetic_PC2  0.2796  1  0.5969924    
## garden_MAT:genetic_PC3  0.0559  1  0.8130797    
## garden_MAP:genetic_PC1  1.1731  1  0.2787626    
## garden_MAP:genetic_PC2  0.0669  1  0.7958716    
## garden_MAP:genetic_PC3  0.0295  1  0.8635558    
## ---
## Signif. codes:  0 '***' 0.001 '**' 0.01 '*' 0.05 '.' 0.1 ' ' 1
```

```
## `geom_smooth()` using formula = 'y ~ x'
```

```
## [1] "Model for: lower_stomata_density_mm2"
```

```
## Warning: Some predictor variables are on very different scales: consider
## rescaling
```

```
## Analysis of Deviance Table (Type II Wald chisquare tests)
## 
## Response: lower_stomata_density_mm2
##                          Chisq Df Pr(>Chisq)    
## garden_MAT              0.4527  1  0.5010557    
## home_MAT                1.7287  1  0.1885727    
## home_MAP                1.6535  1  0.1984849    
## garden_MAP              0.2734  1  0.6010727    
## genetic_PC1            13.8817  1  0.0001947 ***
## genetic_PC2             2.5829  1  0.1080259    
## genetic_PC3             0.0720  1  0.7884447    
## garden_MAT:home_MAT     0.6201  1  0.4309985    
## garden_MAT:home_MAP     0.0927  1  0.7608010    
## home_MAP:garden_MAP     0.0060  1  0.9381343    
## home_MAT:garden_MAP     0.1933  1  0.6602119    
## garden_MAT:genetic_PC1  1.7246  1  0.1891007    
## garden_MAT:genetic_PC2  0.0060  1  0.9380801    
## garden_MAT:genetic_PC3  0.5940  1  0.4408781    
## garden_MAP:genetic_PC1  0.0377  1  0.8459923    
## garden_MAP:genetic_PC2  2.1071  1  0.1466201    
## garden_MAP:genetic_PC3  0.0079  1  0.9293697    
## ---
## Signif. codes:  0 '***' 0.001 '**' 0.01 '*' 0.05 '.' 0.1 ' ' 1
```

```
## `geom_smooth()` using formula = 'y ~ x'
```

```
## [1] "Model for: upper_stomata_density_mm2"
```

```
## Warning: Some predictor variables are on very different scales: consider
## rescaling
```

```
## Analysis of Deviance Table (Type II Wald chisquare tests)
## 
## Response: upper_stomata_density_mm2
##                         Chisq Df Pr(>Chisq)   
## garden_MAT             0.5679  1   0.451085   
## home_MAT               3.7573  1   0.052578 . 
## home_MAP               0.1363  1   0.711954   
## garden_MAP             7.0339  1   0.007998 **
## genetic_PC1            0.5579  1   0.455112   
## genetic_PC2            0.4983  1   0.480261   
## genetic_PC3            8.8203  1   0.002979 **
## garden_MAT:home_MAT    1.0338  1   0.309262   
## garden_MAT:home_MAP    0.2073  1   0.648884   
## home_MAP:garden_MAP    0.0203  1   0.886786   
## home_MAT:garden_MAP    0.0229  1   0.879623   
## garden_MAT:genetic_PC1 8.0926  1   0.004445 **
## garden_MAT:genetic_PC2 0.2103  1   0.646534   
## garden_MAT:genetic_PC3 0.0046  1   0.945909   
## garden_MAP:genetic_PC1 0.7463  1   0.387648   
## garden_MAP:genetic_PC2 0.3297  1   0.565810   
## garden_MAP:genetic_PC3 1.0703  1   0.300868   
## ---
## Signif. codes:  0 '***' 0.001 '**' 0.01 '*' 0.05 '.' 0.1 ' ' 1
```

```
## Warning: Removed 811 rows containing non-finite outside the scale range
## (`stat_boxplot()`).
```

```
## `geom_smooth()` using formula = 'y ~ x'
```

```
## `geom_smooth()` using formula = 'y ~ x'
```

```
## [1] "Model for: stomata_ratio"
```

```
## Warning: Some predictor variables are on very different scales: consider
## rescaling
```

```
## boundary (singular) fit: see help('isSingular')
```

```
## Analysis of Deviance Table (Type II Wald chisquare tests)
## 
## Response: stomata_ratio
##                         Chisq Df Pr(>Chisq)  
## garden_MAT             1.1039  1    0.29341  
## home_MAT               2.6177  1    0.10567  
## home_MAP               0.0000  1    0.99798  
## garden_MAP             2.8943  1    0.08889 .
## genetic_PC1            0.5231  1    0.46953  
## genetic_PC2            0.0557  1    0.81345  
## genetic_PC3            5.7485  1    0.01650 *
## garden_MAT:home_MAT    0.2339  1    0.62862  
## garden_MAT:home_MAP    0.0689  1    0.79291  
## home_MAP:garden_MAP    0.0345  1    0.85273  
## home_MAT:garden_MAP    0.3670  1    0.54462  
## garden_MAT:genetic_PC1 5.4042  1    0.02009 *
## garden_MAT:genetic_PC2 0.2928  1    0.58842  
## garden_MAT:genetic_PC3 0.0024  1    0.96056  
## garden_MAP:genetic_PC1 0.0891  1    0.76533  
## garden_MAP:genetic_PC2 0.4838  1    0.48673  
## garden_MAP:genetic_PC3 0.8818  1    0.34772  
## ---
## Signif. codes:  0 '***' 0.001 '**' 0.01 '*' 0.05 '.' 0.1 ' ' 1
```

```
## boundary (singular) fit: see help('isSingular')
```

```
## `geom_smooth()` using formula = 'y ~ x'
```

```
## `geom_smooth()` using formula = 'y ~ x'
```

```
## [1] "Model for: licor_gsw"
```

```
## Warning: Some predictor variables are on very different scales: consider
## rescaling
```

```
## Analysis of Deviance Table (Type II Wald chisquare tests)
## 
## Response: licor_gsw
##                         Chisq Df Pr(>Chisq)   
## garden_MAT             0.0432  1   0.835313   
## home_MAT               0.6337  1   0.425991   
## home_MAP               3.5240  1   0.060487 . 
## garden_MAP             0.0021  1   0.963827   
## genetic_PC1            2.6989  1   0.100415   
## genetic_PC2            0.8707  1   0.350756   
## genetic_PC3            0.0350  1   0.851539   
## garden_MAT:home_MAT    0.2629  1   0.608124   
## garden_MAT:home_MAP    1.5733  1   0.209723   
## home_MAP:garden_MAP    0.0325  1   0.856972   
## home_MAT:garden_MAP    0.0882  1   0.766501   
## garden_MAT:genetic_PC1 0.1046  1   0.746387   
## garden_MAT:genetic_PC2 3.9007  1   0.048265 * 
## garden_MAT:genetic_PC3 9.2012  1   0.002419 **
## garden_MAP:genetic_PC1 0.6916  1   0.405613   
## garden_MAP:genetic_PC2 1.5166  1   0.218132   
## garden_MAP:genetic_PC3 1.5875  1   0.207689   
## ---
## Signif. codes:  0 '***' 0.001 '**' 0.01 '*' 0.05 '.' 0.1 ' ' 1
```

```
## `geom_smooth()` using formula = 'y ~ x'
```

```
## `geom_smooth()` using formula = 'y ~ x'
```

```
## [1] "Model for: licor_gbw"
```

```
## Warning: Some predictor variables are on very different scales: consider
## rescaling
```

```
## Analysis of Deviance Table (Type II Wald chisquare tests)
## 
## Response: licor_gbw
##                         Chisq Df Pr(>Chisq)  
## garden_MAT             0.1597  1    0.68943  
## home_MAT               0.7010  1    0.40246  
## home_MAP               0.1142  1    0.73547  
## garden_MAP             0.5341  1    0.46488  
## genetic_PC1            0.1586  1    0.69042  
## genetic_PC2            0.0033  1    0.95423  
## genetic_PC3            0.8678  1    0.35156  
## garden_MAT:home_MAT    0.8073  1    0.36891  
## garden_MAT:home_MAP    4.6013  1    0.03195 *
## home_MAP:garden_MAP    1.8159  1    0.17780  
## home_MAT:garden_MAP    0.0005  1    0.98141  
## garden_MAT:genetic_PC1 0.0193  1    0.88960  
## garden_MAT:genetic_PC2 1.3308  1    0.24866  
## garden_MAT:genetic_PC3 5.7560  1    0.01643 *
## garden_MAP:genetic_PC1 0.0739  1    0.78569  
## garden_MAP:genetic_PC2 1.2046  1    0.27241  
## garden_MAP:genetic_PC3 2.1685  1    0.14086  
## ---
## Signif. codes:  0 '***' 0.001 '**' 0.01 '*' 0.05 '.' 0.1 ' ' 1
```

```
## `geom_smooth()` using formula = 'y ~ x'
```

```
## `geom_smooth()` using formula = 'y ~ x'
```

```
## [1] "Model for: licor_ETR"
```

```
## Warning: Some predictor variables are on very different scales: consider
## rescaling
```

```
## Analysis of Deviance Table (Type II Wald chisquare tests)
## 
## Response: licor_ETR
##                         Chisq Df Pr(>Chisq)  
## garden_MAT             0.2634  1    0.60781  
## home_MAT               0.3165  1    0.57370  
## home_MAP               0.0161  1    0.89907  
## garden_MAP             0.7963  1    0.37220  
## genetic_PC1            0.0225  1    0.88066  
## genetic_PC2            0.2423  1    0.62253  
## genetic_PC3            0.0000  1    0.99906  
## garden_MAT:home_MAT    0.0135  1    0.90733  
## garden_MAT:home_MAP    2.8489  1    0.09144 .
## home_MAP:garden_MAP    0.5850  1    0.44435  
## home_MAT:garden_MAP    0.0917  1    0.76208  
## garden_MAT:genetic_PC1 1.1909  1    0.27515  
## garden_MAT:genetic_PC2 1.1288  1    0.28803  
## garden_MAT:genetic_PC3 0.0239  1    0.87712  
## garden_MAP:genetic_PC1 1.7053  1    0.19159  
## garden_MAP:genetic_PC2 0.2309  1    0.63083  
## garden_MAP:genetic_PC3 0.2915  1    0.58925  
## ---
## Signif. codes:  0 '***' 0.001 '**' 0.01 '*' 0.05 '.' 0.1 ' ' 1
```

```
## [1] "Model for: licor_Fs"
```

```
## Warning: Some predictor variables are on very different scales: consider
## rescaling
```

```
## boundary (singular) fit: see help('isSingular')
```

```
## Analysis of Deviance Table (Type II Wald chisquare tests)
## 
## Response: licor_Fs
##                         Chisq Df Pr(>Chisq)  
## garden_MAT             0.3607  1    0.54813  
## home_MAT               0.2762  1    0.59921  
## home_MAP               3.6511  1    0.05603 .
## garden_MAP             0.2574  1    0.61192  
## genetic_PC1            0.4082  1    0.52290  
## genetic_PC2            5.4178  1    0.01993 *
## genetic_PC3            0.2733  1    0.60113  
## garden_MAT:home_MAT    0.0073  1    0.93179  
## garden_MAT:home_MAP    2.2321  1    0.13517  
## home_MAP:garden_MAP    1.4155  1    0.23414  
## home_MAT:garden_MAP    0.8446  1    0.35808  
## garden_MAT:genetic_PC1 0.0755  1    0.78356  
## garden_MAT:genetic_PC2 0.0617  1    0.80387  
## garden_MAT:genetic_PC3 0.1023  1    0.74908  
## garden_MAP:genetic_PC1 3.3305  1    0.06801 .
## garden_MAP:genetic_PC2 0.2365  1    0.62672  
## garden_MAP:genetic_PC3 1.3227  1    0.25011  
## ---
## Signif. codes:  0 '***' 0.001 '**' 0.01 '*' 0.05 '.' 0.1 ' ' 1
```

```
## boundary (singular) fit: see help('isSingular')
```

```
## `geom_smooth()` using formula = 'y ~ x'
```

```
## [1] "Model for: licor_Fm."
```

```
## Warning: Some predictor variables are on very different scales: consider
## rescaling
```

```
## Analysis of Deviance Table (Type II Wald chisquare tests)
## 
## Response: licor_Fm.
##                          Chisq Df Pr(>Chisq)    
## garden_MAT              1.9678  1  0.1606805    
## home_MAT                0.2991  1  0.5844753    
## home_MAP                2.2327  1  0.1351185    
## garden_MAP              2.6340  1  0.1046007    
## genetic_PC1             0.5472  1  0.4594774    
## genetic_PC2             0.9919  1  0.3192824    
## genetic_PC3             0.3955  1  0.5294180    
## garden_MAT:home_MAT     0.0010  1  0.9748299    
## garden_MAT:home_MAP     0.4108  1  0.5215847    
## home_MAP:garden_MAP     0.2815  1  0.5956990    
## home_MAT:garden_MAP     2.6303  1  0.1048443    
## garden_MAT:genetic_PC1  4.6595  1  0.0308824 *  
## garden_MAT:genetic_PC2  0.0949  1  0.7580151    
## garden_MAT:genetic_PC3  1.1839  1  0.2765694    
## garden_MAP:genetic_PC1 11.6397  1  0.0006456 ***
## garden_MAP:genetic_PC2  0.4944  1  0.4819786    
## garden_MAP:genetic_PC3  5.3512  1  0.0207080 *  
## ---
## Signif. codes:  0 '***' 0.001 '**' 0.01 '*' 0.05 '.' 0.1 ' ' 1
```

```
## `geom_smooth()` using formula = 'y ~ x'
```

```
## `geom_smooth()` using formula = 'y ~ x'
```

```
## `geom_smooth()` using formula = 'y ~ x'
```

```
## [1] "Model for: licor_PhiPS2"
```

```
## Warning: Some predictor variables are on very different scales: consider
## rescaling
```

```
## Analysis of Deviance Table (Type II Wald chisquare tests)
## 
## Response: licor_PhiPS2
##                          Chisq Df Pr(>Chisq)    
## garden_MAT              2.0489  1  0.1523138    
## home_MAT                0.3918  1  0.5313483    
## home_MAP                0.0818  1  0.7749224    
## garden_MAP              2.7844  1  0.0951883 .  
## genetic_PC1             0.3708  1  0.5425663    
## genetic_PC2             0.3205  1  0.5713240    
## genetic_PC3             0.5753  1  0.4481702    
## garden_MAT:home_MAT     0.7540  1  0.3852146    
## garden_MAT:home_MAP     0.0034  1  0.9532937    
## home_MAP:garden_MAP     0.0000  1  0.9997927    
## home_MAT:garden_MAP     3.5830  1  0.0583729 .  
## garden_MAT:genetic_PC1 11.8535  1  0.0005755 ***
## garden_MAT:genetic_PC2  0.1065  1  0.7441113    
## garden_MAT:genetic_PC3  2.7325  1  0.0983264 .  
## garden_MAP:genetic_PC1 12.6797  1  0.0003697 ***
## garden_MAP:genetic_PC2  0.3094  1  0.5780569    
## garden_MAP:genetic_PC3  5.9499  1  0.0147181 *  
## ---
## Signif. codes:  0 '***' 0.001 '**' 0.01 '*' 0.05 '.' 0.1 ' ' 1
```

```
## `geom_smooth()` using formula = 'y ~ x'
```

```
## `geom_smooth()` using formula = 'y ~ x'
```

```
## `geom_smooth()` using formula = 'y ~ x'
```

```
# plot all model effects together
# there are really too many variables for this to be useful
#plot_models(mods.23,std.est = T, m.labels = vars, p.shape = T, spacing = 1, vline.color = 'black', colors = cols25())

names(mods.23) <- vars

# setup dfs for effect sizes and p-values of each factor
# use last full model run in loop to get effect names - models in mods object have some effects reduced
rows <- names(fixef(mod))[-1] # names of fixed effects, minus intercept
# reorder row so garden clim is first
rows <- rows[c(1,4,2,3,5:17)]

colms <- vars

# effect size
df.eff.23 <- as.data.frame(matrix(ncol = length(colms), nrow = length(rows)))
colnames(df.eff.23) <- colms
rownames(df.eff.23) <- rows
# standardized effect size
df.eff.std.23 <- as.data.frame(matrix(ncol = length(colms), nrow = length(rows)))
colnames(df.eff.std.23) <- colms
rownames(df.eff.std.23) <- rows
# p values
df.pval.23 <- as.data.frame(matrix(ncol = length(colms), nrow = length(rows)))
colnames(df.pval.23) <- colms
rownames(df.pval.23) <- rows
# p values
df.pval.all.23 <- as.data.frame(matrix(ncol = length(colms), nrow = length(rows)))
colnames(df.pval.all.23) <- colms
rownames(df.pval.all.23) <- rows


# get p-values and effect sizes for each variable

for(n in 1:length(vars)){
  
  var <- vars[[n]]
  mod <- mods.23[[n]]
  # model info
  mod.info <- car::Anova(mod)
  # standardize coefficients
  eff.std <- effectsize::standardize_parameters(mod)
  
  # fixed effects, minus intercept
  #fes <- names(fixef(mod)[-1])
  # reorder so garden climates are first
  fes <- rownames(mod.info)
  
  # get their p-values and effect sizes for each fixed effect
    for(f in 1:length(fes)){
      
      fe <- fes[[f]]
      
      # check for significance
      pval <- mod.info[fe, 'Pr(>Chisq)']
      
      if(pval <= 0.05){
        df.eff.23[fe, var] <- fixef(mods.23[[var]])[fe]
        df.eff.std.23[fe,var] <- eff.std[eff.std$Parameter == fe, 'Std_Coefficient']
        df.pval.23[fe, var] <- mod.info[fe, 'Pr(>Chisq)']
      }
  }
}
```

```
## boundary (singular) fit: see help('isSingular')
## boundary (singular) fit: see help('isSingular')
## boundary (singular) fit: see help('isSingular')
## boundary (singular) fit: see help('isSingular')
```

```
# get all pvalues so they can be adjusted
for(n in 1:length(vars)){
  
  var <- vars[[n]]
  mod <- mods.23[[n]]
  #mod.info <- summary(mod)
  mod.info <- car::Anova(mod)

  # loop through retained variables and get their p-values and effect sizes
    for(f in 1:nrow(df.pval.all.23)){
      
      fe <- rownames(df.pval.all.23)[f]
      # get pval 
         df.pval.all.23[fe, var] <- mod.info[fe, 'Pr(>Chisq)']
         
  }
}

# adjust pvals
padj <- p.adjust(unlist(df.pval.all.23), method = 'BH')
df.pval.adj.23 <- as.data.frame(matrix(padj, ncol = ncol(df.pval.all.23)))
colnames(df.pval.adj.23) <- colnames(df.pval.all.23)
rownames(df.pval.adj.23) <- rownames(df.pval.all.23)

hist(unlist(df.pval.all.23))
```

```
hist(unlist(df.pval.adj.23))
```

```
plot(unlist(df.pval.all.23), unlist(df.pval.adj.23))
```

```
plot(unlist(df.eff.std.23), unlist(df.pval.all.23))
```

```
##################
# color scale based on all effect sizes
max_col <- max(unlist(df.eff.std.23), na.rm = T)
min_col <- min(unlist(df.eff.std.23), na.rm = T)

# make scale same at both ends
max_col <- max(abs(max_col), abs(min_col))
min_col <- min(abs(max_col), abs(min_col))*-1
max_ceil <- ceiling(max_col*10)/10 # take ceiling rounded to one decimal place - used for placing ticks on legend

colf <- colorRamp2(c('red', 'white', 'blue'), breaks = c(min_col, 0, max_col))

# raw pvals
# cell_text <- df.pval
# cell_text[is.na(df.pval)] <- ''
# cell_text[df.pval > 0.05] <- ''
# cell_text[df.pval <= 0.05 & df.pval > 0.01] <- '*'
# cell_text[df.pval <= 0.01 & df.pval > 0.001] <- '**'
# cell_text[df.pval<= 0.001] <- '***'
# cell_text <- t(cell_text)

# adjusted pvalues
cell_text <- df.pval.adj.23
cell_text[is.na(df.pval.adj.23)] <- ''
cell_text[df.pval.adj.23 > 0.05] <- ''
cell_text[df.pval.adj.23 <= 0.05 & df.pval.adj.23 > 0.01] <- '*'
cell_text[df.pval.adj.23 <= 0.01 & df.pval.adj.23 > 0.001] <- '**'
cell_text[df.pval.adj.23<= 0.001] <- '***'
cell_text <- t(cell_text)


# heatmap

hm23 <- Heatmap(as.matrix(t(df.eff.std.23)),
        cluster_rows = F,
        cluster_columns = F,
        rect_gp = gpar(col = "black", lwd = 1),
        col = colf,
        na_col = 'grey80',
        row_names_side = 'left',
        column_names_side = 'top',
        column_names_rot = 90,
        column_names_centered = T,
        column_title = 'Predictors',
        column_title_gp = gpar(fontface = 'bold'),
        row_title_gp = gpar(fontface = 'bold'),
        row_title = 'Trait',
        #left_annotation = ha_row,
        heatmap_legend_param = list(title = 'Std. Effect Size', at = c(max_ceil*-1, 0, max_ceil)),
        cell_fun = function(j, i, x, y, width, height, fill) { # add text to each grid
                      grid.text(cell_text[i, j], x, y, gp = gpar(cex = 1.5))}
        )


ComplexHeatmap::draw(hm23, padding = unit(c(2, 30, 2, 15), "mm"))
```

##### 2.1.3 Print model tables - 2023

```
kable(df.eff.23, label = 'Effects for 2023 traits')
```

|  | DOY\_Stage2\_2023 | Stage2\_cGDD\_2023 | DOY\_Stage3\_2023 | Stage3\_cGDD\_2023 | DOY\_Stage6\_2023 | daylength\_Stage6\_2023 | DOY\_Stage7\_2023 | daylength\_Stage7\_2023 | DOY\_last\_budset\_2023 | daylength\_last\_budset\_2023 | growing\_season\_days\_2023 | leaf\_thickness\_avg\_mm\_2023 | leaf\_area\_cm2\_2023 | leaf\_mass\_g\_2023 | LMA\_g\_m2\_2023 | lower\_stomata\_pore\_length\_mean\_um | upper\_stomata\_pore\_length\_mean\_um | lower\_stomata\_density\_mm2 | upper\_stomata\_density\_mm2 | stomata\_ratio | licor\_gsw | licor\_gbw | licor\_ETR | licor\_Fs | licor\_Fm. | licor\_PhiPS2 |
| --- | --- | --- | --- | --- | --- | --- | --- | --- | --- | --- | --- | --- | --- | --- | --- | --- | --- | --- | --- | --- | --- | --- | --- | --- | --- | --- |
| garden\_MAT | -4.6126230 | 54.85756 | -3.882361 | 64.51615 | NA | NA | NA | NA | NA | NA | NA | NA | 0.9642260 | 0.0063582 | NA | NA | NA | NA | NA | NA | NA | NA | NA | NA | NA | NA |
| garden\_MAP | NA | NA | NA | NA | NA | NA | NA | NA | NA | NA | NA | NA | -0.0043970 | -0.0000504 | NA | NA | NA | NA | 0.0156516 | NA | NA | NA | NA | NA | NA | NA |
| home\_MAT | 0.7838523 | NA | 1.323514 | NA | 1.7789570 | NA | NA | NA | 0.6382378 | NA | NA | NA | NA | -0.0396989 | NA | NA | NA | NA | NA | NA | NA | NA | NA | NA | NA | NA |
| home\_MAP | NA | NA | NA | NA | NA | NA | NA | NA | NA | NA | NA | NA | NA | NA | NA | NA | 0.0036169 | NA | NA | NA | NA | NA | NA | NA | NA | NA |
| genetic\_PC1 | NA | NA | NA | NA | 72.4042234 | NA | NA | NA | NA | NA | NA | NA | -47.3533704 | NA | NA | NA | NA | 1025.125 | NA | NA | NA | NA | NA | NA | NA | NA |
| genetic\_PC2 | NA | NA | NA | NA | NA | NA | NA | NA | NA | NA | NA | NA | NA | NA | NA | 25.1772050 | NA | NA | NA | NA | NA | NA | NA | 321.2292 | NA | NA |
| genetic\_PC3 | NA | NA | NA | NA | NA | NA | NA | NA | NA | NA | NA | NA | NA | NA | NA | NA | NA | NA | 102.8045860 | 0.6144987 | NA | NA | NA | NA | NA | NA |
| garden\_MAT:home\_MAT | NA | NA | NA | NA | 0.3621410 | 0.0151878 | NA | NA | NA | NA | NA | NA | 0.7947548 | 0.0074748 | NA | NA | NA | NA | NA | NA | NA | NA | NA | NA | NA | NA |
| garden\_MAT:home\_MAP | NA | NA | NA | NA | NA | NA | NA | NA | NA | NA | NA | NA | NA | NA | NA | NA | NA | NA | NA | NA | NA | -0.0000002 | NA | NA | NA | NA |
| home\_MAP:garden\_MAP | NA | NA | NA | NA | NA | NA | NA | NA | NA | NA | NA | -1e-07 | NA | NA | NA | NA | NA | NA | NA | NA | NA | NA | NA | NA | NA | NA |
| home\_MAT:garden\_MAP | -0.0015509 | NA | NA | NA | -0.0035351 | NA | NA | NA | NA | NA | NA | NA | -0.0042351 | -0.0000331 | 3.68e-05 | NA | NA | NA | NA | NA | NA | NA | NA | NA | NA | NA |
| garden\_MAT:genetic\_PC1 | NA | NA | NA | NA | NA | NA | NA | NA | NA | NA | NA | NA | 23.1054744 | NA | NA | NA | NA | NA | 47.3279704 | 0.3767374 | NA | NA | NA | NA | -167.896335 | -0.4607744 |
| garden\_MAT:genetic\_PC2 | NA | NA | NA | NA | 32.3971876 | -0.7149057 | NA | NA | NA | NA | NA | NA | 39.3847947 | 0.3714492 | NA | NA | NA | NA | NA | NA | 0.2186144 | NA | NA | NA | NA | NA |
| garden\_MAT:genetic\_PC3 | NA | NA | NA | NA | NA | NA | NA | NA | NA | NA | NA | NA | NA | NA | NA | NA | NA | NA | NA | NA | -0.3550011 | 0.0012980 | NA | NA | NA | NA |
| garden\_MAP:genetic\_PC1 | NA | NA | NA | NA | NA | NA | NA | NA | NA | NA | NA | NA | -0.2797732 | -0.0033563 | NA | NA | NA | NA | NA | NA | NA | NA | NA | NA | 2.574194 | 0.0046245 |
| garden\_MAP:genetic\_PC2 | NA | NA | NA | NA | -0.1334996 | NA | NA | NA | NA | NA | NA | NA | -0.2329023 | -0.0020070 | NA | 0.0339568 | NA | NA | NA | NA | NA | NA | NA | NA | NA | NA |
| garden\_MAP:genetic\_PC3 | NA | NA | NA | NA | NA | NA | NA | NA | NA | NA | NA | NA | -0.1554209 | NA | NA | NA | NA | NA | NA | NA | NA | NA | NA | NA | 1.633892 | 0.0029662 |

```
kable(df.eff.std.23, label = 'Standardized effect sizes for 2023 traits')
```

|  | DOY\_Stage2\_2023 | Stage2\_cGDD\_2023 | DOY\_Stage3\_2023 | Stage3\_cGDD\_2023 | DOY\_Stage6\_2023 | daylength\_Stage6\_2023 | DOY\_Stage7\_2023 | daylength\_Stage7\_2023 | DOY\_last\_budset\_2023 | daylength\_last\_budset\_2023 | growing\_season\_days\_2023 | leaf\_thickness\_avg\_mm\_2023 | leaf\_area\_cm2\_2023 | leaf\_mass\_g\_2023 | LMA\_g\_m2\_2023 | lower\_stomata\_pore\_length\_mean\_um | upper\_stomata\_pore\_length\_mean\_um | lower\_stomata\_density\_mm2 | upper\_stomata\_density\_mm2 | stomata\_ratio | licor\_gsw | licor\_gbw | licor\_ETR | licor\_Fs | licor\_Fm. | licor\_PhiPS2 |
| --- | --- | --- | --- | --- | --- | --- | --- | --- | --- | --- | --- | --- | --- | --- | --- | --- | --- | --- | --- | --- | --- | --- | --- | --- | --- | --- |
| garden\_MAT | -0.5692812 | 0.6404178 | -0.5512488 | 0.7093145 | NA | NA | NA | NA | NA | NA | NA | NA | 0.4509394 | 0.4342514 | NA | NA | NA | NA | NA | NA | NA | NA | NA | NA | NA | NA |
| garden\_MAP | NA | NA | NA | NA | NA | NA | NA | NA | NA | NA | NA | NA | -0.4740569 | -0.4714874 | NA | NA | NA | NA | 0.2535938 | NA | NA | NA | NA | NA | NA | NA |
| home\_MAT | 0.1589187 | NA | 0.1714929 | NA | 0.1827911 | NA | NA | NA | 0.1312286 | NA | NA | NA | NA | 0.1274059 | NA | NA | NA | NA | NA | NA | NA | NA | NA | NA | NA | NA |
| home\_MAP | NA | NA | NA | NA | NA | NA | NA | NA | NA | NA | NA | NA | NA | NA | NA | NA | 0.2385671 | NA | NA | NA | NA | NA | NA | NA | NA | NA |
| genetic\_PC1 | NA | NA | NA | NA | -0.1177713 | NA | NA | NA | NA | NA | NA | NA | -0.0994744 | NA | NA | NA | NA | 0.2464168 | NA | NA | NA | NA | NA | NA | NA | NA |
| genetic\_PC2 | NA | NA | NA | NA | NA | NA | NA | NA | NA | NA | NA | NA | NA | NA | NA | 0.2559665 | NA | NA | NA | NA | NA | NA | NA | 0.1165161 | NA | NA |
| genetic\_PC3 | NA | NA | NA | NA | NA | NA | NA | NA | NA | NA | NA | NA | NA | NA | NA | NA | NA | NA | 0.3133116 | 0.2283170 | NA | NA | NA | NA | NA | NA |
| garden\_MAT:home\_MAT | NA | NA | NA | NA | 0.0724660 | 0.1173724 | NA | NA | NA | NA | NA | NA | 0.2570973 | 0.2286461 | NA | NA | NA | NA | NA | NA | NA | NA | NA | NA | NA | NA |
| garden\_MAT:home\_MAP | NA | NA | NA | NA | NA | NA | NA | NA | NA | NA | NA | NA | NA | NA | NA | NA | NA | NA | NA | NA | NA | -0.0378542 | NA | NA | NA | NA |
| home\_MAP:garden\_MAP | NA | NA | NA | NA | NA | NA | NA | NA | NA | NA | NA | -0.1417504 | NA | NA | NA | NA | NA | NA | NA | NA | NA | NA | NA | NA | NA | NA |
| home\_MAT:garden\_MAP | -0.0776557 | NA | NA | NA | -0.1048938 | NA | NA | NA | NA | NA | NA | NA | -0.2089471 | -0.1543864 | 0.153977 | NA | NA | NA | NA | NA | NA | NA | NA | NA | NA | NA |
| garden\_MAT:genetic\_PC1 | NA | NA | NA | NA | NA | NA | NA | NA | NA | NA | NA | NA | 0.1018652 | NA | NA | NA | NA | NA | 0.1560733 | 0.1375364 | NA | NA | NA | NA | -0.1443972 | -0.2170526 |
| garden\_MAT:genetic\_PC2 | NA | NA | NA | NA | 0.0918215 | -0.0782529 | NA | NA | NA | NA | NA | NA | 0.1866634 | 0.1670218 | NA | NA | NA | NA | NA | NA | 0.1442343 | NA | NA | NA | NA | NA |
| garden\_MAT:genetic\_PC3 | NA | NA | NA | NA | NA | NA | NA | NA | NA | NA | NA | NA | NA | NA | NA | NA | NA | NA | NA | NA | -0.2523379 | 0.0513812 | NA | NA | NA | NA |
| garden\_MAP:genetic\_PC1 | NA | NA | NA | NA | NA | NA | NA | NA | NA | NA | NA | NA | -0.1881162 | -0.2136380 | NA | NA | NA | NA | NA | NA | NA | NA | NA | NA | 0.2323322 | 0.2286072 |
| garden\_MAP:genetic\_PC2 | NA | NA | NA | NA | -0.0561060 | NA | NA | NA | NA | NA | NA | NA | -0.1683502 | -0.1374202 | NA | 0.1212302 | NA | NA | NA | NA | NA | NA | NA | NA | NA | NA |
| garden\_MAP:genetic\_PC3 | NA | NA | NA | NA | NA | NA | NA | NA | NA | NA | NA | NA | -0.1196392 | NA | NA | NA | NA | NA | NA | NA | NA | NA | NA | NA | 0.1697261 | 0.1687656 |

```
kable(df.pval.all.23, label = 'p-values for 2023 traits')
```

|  | DOY\_Stage2\_2023 | Stage2\_cGDD\_2023 | DOY\_Stage3\_2023 | Stage3\_cGDD\_2023 | DOY\_Stage6\_2023 | daylength\_Stage6\_2023 | DOY\_Stage7\_2023 | daylength\_Stage7\_2023 | DOY\_last\_budset\_2023 | daylength\_last\_budset\_2023 | growing\_season\_days\_2023 | leaf\_thickness\_avg\_mm\_2023 | leaf\_area\_cm2\_2023 | leaf\_mass\_g\_2023 | LMA\_g\_m2\_2023 | lower\_stomata\_pore\_length\_mean\_um | upper\_stomata\_pore\_length\_mean\_um | lower\_stomata\_density\_mm2 | upper\_stomata\_density\_mm2 | stomata\_ratio | licor\_gsw | licor\_gbw | licor\_ETR | licor\_Fs | licor\_Fm. | licor\_PhiPS2 |
| --- | --- | --- | --- | --- | --- | --- | --- | --- | --- | --- | --- | --- | --- | --- | --- | --- | --- | --- | --- | --- | --- | --- | --- | --- | --- | --- |
| garden\_MAT | 0.0000775 | 0.0001059 | 0.0000141 | 0.0000033 | 0.1407721 | 0.9261265 | 0.7304786 | 0.2503515 | 0.7261349 | 0.6874904 | 0.3755623 | 0.4703679 | 0.0010030 | 0.0004358 | 0.8343274 | 0.5125922 | 0.5714425 | 0.5010557 | 0.4510849 | 0.2934076 | 0.8353127 | 0.6894326 | 0.6078102 | 0.5481321 | 0.1606805 | 0.1523138 |
| garden\_MAP | 0.2511946 | 0.9706956 | 0.1370051 | 0.9800796 | 0.6486269 | 0.4774682 | 0.2430430 | 0.2203434 | 0.3171802 | 0.4607963 | 0.3978803 | 0.7705946 | 0.0007177 | 0.0001881 | 0.2206840 | 0.5082344 | 0.9977443 | 0.6010727 | 0.0079983 | 0.0888912 | 0.9638268 | 0.4648820 | 0.3722037 | 0.6119156 | 0.1046007 | 0.0951883 |
| home\_MAT | 0.0324927 | 0.0699733 | 0.0223543 | 0.0520472 | 0.0036678 | 0.1009795 | 0.5717746 | 0.7646376 | 0.0237728 | 0.0916839 | 0.3782346 | 0.3314654 | 0.1033898 | 0.0471714 | 0.1520148 | 0.7093118 | 0.1723973 | 0.1885727 | 0.0525784 | 0.1056749 | 0.4259908 | 0.4024611 | 0.5737049 | 0.5992059 | 0.5844753 | 0.5313483 |
| home\_MAP | 0.1805627 | 0.1908112 | 0.3193248 | 0.3279296 | 0.2883398 | 0.5073028 | 0.4079138 | 0.9747582 | 0.1767804 | 0.1942472 | 0.0589147 | 0.4960763 | 0.3235834 | 0.1992121 | 0.7363813 | 0.2635864 | 0.0002278 | 0.1984849 | 0.7119541 | 0.9979798 | 0.0604865 | 0.7354651 | 0.8990659 | 0.0560321 | 0.1351185 | 0.7749224 |
| genetic\_PC1 | 0.9897216 | 0.8711757 | 0.8548635 | 0.8499611 | 0.0237789 | 0.7337380 | 0.4664028 | 0.5305391 | 0.0920773 | 0.3892841 | 0.2343268 | 0.9791824 | 0.0494286 | 0.0980057 | 0.6781025 | 0.1944999 | 0.4856909 | 0.0001947 | 0.4551122 | 0.4695307 | 0.1004147 | 0.6904235 | 0.8806564 | 0.5228987 | 0.4594774 | 0.5425663 |
| genetic\_PC2 | 0.8121986 | 0.9094893 | 0.8143862 | 0.7746501 | 0.2556286 | 0.1582659 | 0.2698964 | 0.6018312 | 0.3277970 | 0.1153368 | 0.1376101 | 0.5167921 | 0.1795381 | 0.1928941 | 0.8644610 | 0.0005568 | 0.0898250 | 0.1080259 | 0.4802614 | 0.8134461 | 0.3507555 | 0.9542284 | 0.6225280 | 0.0199324 | 0.3192824 | 0.5713240 |
| genetic\_PC3 | 0.9780453 | 0.8879893 | 0.8247952 | 0.8108060 | 0.2712727 | 0.6528366 | 0.1186400 | 0.2290284 | 0.3913766 | 0.9499643 | 0.5309674 | 0.6517121 | 0.7022888 | 0.4141330 | 0.5536566 | 0.3630699 | 0.5586681 | 0.7884447 | 0.0029789 | 0.0165029 | 0.8515386 | 0.3515635 | 0.9990579 | 0.6011321 | 0.5294180 | 0.4481702 |
| garden\_MAT:home\_MAT | 0.0986820 | 0.1854829 | 0.5204900 | 0.1898293 | 0.0399084 | 0.0080062 | 0.3976884 | 0.3209301 | 0.2250557 | 0.8047496 | 0.1272779 | 0.8625904 | 0.0000110 | 0.0001194 | 0.5967103 | 0.4395015 | 0.6007716 | 0.4309985 | 0.3092616 | 0.6286206 | 0.6081236 | 0.3689139 | 0.9073342 | 0.9317860 | 0.9748299 | 0.3852146 |
| garden\_MAT:home\_MAP | 0.6996375 | 0.9379771 | 0.8646973 | 0.8359842 | 0.3187890 | 0.7628396 | 0.2056881 | 0.3395743 | 0.6284440 | 0.4740913 | 0.5540348 | 0.1288514 | 0.8251245 | 0.4680885 | 0.7608296 | 0.8397944 | 0.9108846 | 0.7608010 | 0.6488844 | 0.7929090 | 0.2097232 | 0.0319481 | 0.0914392 | 0.1351682 | 0.5215847 | 0.9532937 |
| home\_MAP:garden\_MAP | 0.1926007 | 0.3683306 | 0.6681309 | 0.7303648 | 0.0616440 | 0.4014430 | 0.2295313 | 0.3578145 | 0.9771604 | 0.8066648 | 0.8132638 | 0.0395384 | 0.1811478 | 0.0714574 | 0.7344046 | 0.6063960 | 0.9642787 | 0.9381343 | 0.8867856 | 0.8527310 | 0.8569721 | 0.1778019 | 0.4443524 | 0.2341389 | 0.5956990 | 0.9997927 |
| home\_MAT:garden\_MAP | 0.0431057 | 0.4799904 | 0.1767075 | 0.2770088 | 0.0030449 | 0.2117601 | 0.0524686 | 0.0685957 | 0.1570344 | 0.7239472 | 0.1202892 | 0.4065639 | 0.0002768 | 0.0081641 | 0.0209621 | 0.1638511 | 0.7750444 | 0.6602119 | 0.8796234 | 0.5446176 | 0.7665010 | 0.9814078 | 0.7620828 | 0.3580754 | 0.1048443 | 0.0583729 |
| garden\_MAT:genetic\_PC1 | 0.4531596 | 0.3950741 | 0.3567424 | 0.6236784 | 0.1168755 | 0.2519505 | 0.8812623 | 0.6000414 | 0.5263399 | 0.9494770 | 0.5819326 | 0.1542273 | 0.0401139 | 0.1063031 | 0.0573493 | 0.6900014 | 0.4938502 | 0.1891007 | 0.0044445 | 0.0200880 | 0.7463870 | 0.8895968 | 0.2751465 | 0.7835570 | 0.0308824 | 0.0005755 |
| garden\_MAT:genetic\_PC2 | 0.7009945 | 0.5025452 | 0.2427116 | 0.2379730 | 0.0006109 | 0.0199904 | 0.2576627 | 0.3660177 | 0.2745037 | 0.3226119 | 0.4652648 | 0.7519947 | 0.0000859 | 0.0005467 | 0.4017900 | 0.1842864 | 0.5969924 | 0.9380801 | 0.6465339 | 0.5884215 | 0.0482650 | 0.2486609 | 0.2880315 | 0.8038684 | 0.7580151 | 0.7441113 |
| garden\_MAT:genetic\_PC3 | 0.9916250 | 0.9262152 | 0.8270443 | 0.6318180 | 0.2508847 | 0.7305174 | 0.7541125 | 0.8142912 | 0.3545670 | 0.0743178 | 0.1460968 | 0.7082880 | 0.0593343 | 0.1137523 | 0.4728155 | 0.7888247 | 0.8130797 | 0.4408781 | 0.9459091 | 0.9605601 | 0.0024186 | 0.0164328 | 0.8771202 | 0.7490815 | 0.2765694 | 0.0983264 |
| garden\_MAP:genetic\_PC1 | 0.6797036 | 0.8399617 | 0.5569320 | 0.6571247 | 0.7020116 | 0.7511205 | 0.2891034 | 0.0518642 | 0.4171673 | 0.8638906 | 0.7678477 | 0.6597563 | 0.0001896 | 0.0000297 | 0.4121387 | 0.5337128 | 0.2787626 | 0.8459923 | 0.3876476 | 0.7653314 | 0.4056127 | 0.7856917 | 0.1915927 | 0.0680063 | 0.0006456 | 0.0003697 |
| garden\_MAP:genetic\_PC2 | 0.8962207 | 0.7888604 | 0.6884661 | 0.5469207 | 0.0363177 | 0.4924197 | 0.6418876 | 0.8005137 | 0.4445127 | 0.7918189 | 0.8294821 | 0.6081484 | 0.0007831 | 0.0071025 | 0.5438457 | 0.0215659 | 0.7958716 | 0.1466201 | 0.5658099 | 0.4867265 | 0.2181320 | 0.2724098 | 0.6308288 | 0.6267209 | 0.4819786 | 0.5780569 |
| garden\_MAP:genetic\_PC3 | 0.5235845 | 0.7470203 | 0.5296667 | 0.6185723 | 0.3022436 | 0.7527143 | 0.7776779 | 0.9926389 | 0.5385606 | 0.4221966 | 0.1179170 | 0.4491537 | 0.0105155 | 0.0671855 | 0.2653511 | 0.3207444 | 0.8635558 | 0.9293697 | 0.3008676 | 0.3477202 | 0.2076894 | 0.1408627 | 0.5892490 | 0.2501075 | 0.0207080 | 0.0147181 |

```
kable(df.pval.adj.23, label = 'Adjusted p-values for 2023 traits')
```

|  | DOY\_Stage2\_2023 | Stage2\_cGDD\_2023 | DOY\_Stage3\_2023 | Stage3\_cGDD\_2023 | DOY\_Stage6\_2023 | daylength\_Stage6\_2023 | DOY\_Stage7\_2023 | daylength\_Stage7\_2023 | DOY\_last\_budset\_2023 | daylength\_last\_budset\_2023 | growing\_season\_days\_2023 | leaf\_thickness\_avg\_mm\_2023 | leaf\_area\_cm2\_2023 | leaf\_mass\_g\_2023 | LMA\_g\_m2\_2023 | lower\_stomata\_pore\_length\_mean\_um | upper\_stomata\_pore\_length\_mean\_um | lower\_stomata\_density\_mm2 | upper\_stomata\_density\_mm2 | stomata\_ratio | licor\_gsw | licor\_gbw | licor\_ETR | licor\_Fs | licor\_Fm. | licor\_PhiPS2 |
| --- | --- | --- | --- | --- | --- | --- | --- | --- | --- | --- | --- | --- | --- | --- | --- | --- | --- | --- | --- | --- | --- | --- | --- | --- | --- | --- |
| garden\_MAT | 0.0063268 | 0.0065977 | 0.0020747 | 0.0014572 | 0.5986666 | 0.9888578 | 0.9497591 | 0.7231307 | 0.9497591 | 0.9477242 | 0.8383765 | 0.8839608 | 0.0192746 | 0.0128422 | 0.9568635 | 0.8963968 | 0.9081135 | 0.8956653 | 0.8839608 | 0.7628598 | 0.9568635 | 0.9477242 | 0.9081135 | 0.9006483 | 0.6341140 | 0.6233582 |
| garden\_MAP | 0.7231307 | 0.9972006 | 0.5963105 | 0.9972006 | 0.9278257 | 0.8839608 | 0.7231307 | 0.6967308 | 0.7901870 | 0.8839608 | 0.8504618 | 0.9497591 | 0.0151059 | 0.0078227 | 0.6967308 | 0.8963968 | 0.9997927 | 0.9081135 | 0.1127670 | 0.5217715 | 0.9972006 | 0.8839608 | 0.8350966 | 0.9106622 | 0.5279324 | 0.5279324 |
| home\_MAT | 0.2992035 | 0.4356086 | 0.2297810 | 0.3873276 | 0.0600440 | 0.5279324 | 0.9081135 | 0.9497591 | 0.2335615 | 0.5217715 | 0.8400990 | 0.7962376 | 0.5279324 | 0.3861064 | 0.6233582 | 0.9497591 | 0.6562517 | 0.6562517 | 0.3873276 | 0.5279324 | 0.8676864 | 0.8504618 | 0.9081135 | 0.9081135 | 0.9081135 | 0.8963968 |
| home\_MAP | 0.6562517 | 0.6562517 | 0.7901870 | 0.7920486 | 0.7561167 | 0.8963968 | 0.8504618 | 0.9972006 | 0.6562517 | 0.6562517 | 0.4034730 | 0.8913240 | 0.7901870 | 0.6620431 | 0.9497591 | 0.7420479 | 0.0083923 | 0.6620431 | 0.9497591 | 0.9997927 | 0.4050766 | 0.9497591 | 0.9716066 | 0.4034730 | 0.5963105 | 0.9497591 |
| genetic\_PC1 | 0.9997927 | 0.9626491 | 0.9578852 | 0.9578852 | 0.2335615 | 0.9497591 | 0.8839608 | 0.8963968 | 0.5217715 | 0.8504618 | 0.7142929 | 0.9972006 | 0.3873276 | 0.5279324 | 0.9477242 | 0.6562517 | 0.8853214 | 0.0078227 | 0.8839608 | 0.8839608 | 0.5279324 | 0.9477242 | 0.9641533 | 0.8963968 | 0.8839608 | 0.9006483 |
| genetic\_PC2 | 0.9497591 | 0.9772112 | 0.9497591 | 0.9497591 | 0.7289540 | 0.6302118 | 0.7420479 | 0.9081135 | 0.7920486 | 0.5519882 | 0.5963105 | 0.8963968 | 0.6562517 | 0.6562517 | 0.9578852 | 0.0141326 | 0.5217715 | 0.5305271 | 0.8839608 | 0.9497591 | 0.8243195 | 0.9947381 | 0.9156182 | 0.2259822 | 0.7901870 | 0.9081135 |
| genetic\_PC3 | 0.9972006 | 0.9660977 | 0.9568635 | 0.9497591 | 0.7420479 | 0.9278257 | 0.5519882 | 0.7094603 | 0.8504618 | 0.9947381 | 0.8963968 | 0.9278257 | 0.9497591 | 0.8553589 | 0.9036287 | 0.8314865 | 0.9045102 | 0.9497591 | 0.0517640 | 0.2026187 | 0.9578852 | 0.8243195 | 0.9997927 | 0.9081135 | 0.8963968 | 0.8839608 |
| garden\_MAT:home\_MAT | 0.5279324 | 0.6562517 | 0.8963968 | 0.6562517 | 0.3409680 | 0.1127670 | 0.8504618 | 0.7901870 | 0.7054938 | 0.9497591 | 0.5799671 | 0.9578852 | 0.0020747 | 0.0065977 | 0.9081135 | 0.8839608 | 0.9081135 | 0.8738593 | 0.7901365 | 0.9156182 | 0.9081135 | 0.8319386 | 0.9772112 | 0.9896309 | 0.9972006 | 0.8504618 |
| garden\_MAT:home\_MAP | 0.9497591 | 0.9896309 | 0.9578852 | 0.9568635 | 0.7901870 | 0.9497591 | 0.6784637 | 0.8113073 | 0.9156182 | 0.8839608 | 0.9036287 | 0.5811460 | 0.9568635 | 0.8839608 | 0.9497591 | 0.9568635 | 0.9772112 | 0.9497591 | 0.9278257 | 0.9497591 | 0.6816003 | 0.2992035 | 0.5217715 | 0.5963105 | 0.8963968 | 0.9947381 |
| home\_MAP:garden\_MAP | 0.6562517 | 0.8319386 | 0.9375043 | 0.9497591 | 0.4066664 | 0.8504618 | 0.7094603 | 0.8243195 | 0.9972006 | 0.9497591 | 0.9497591 | 0.3409680 | 0.6562517 | 0.4386689 | 0.9497591 | 0.9081135 | 0.9972006 | 0.9896309 | 0.9660977 | 0.9578852 | 0.9578852 | 0.6562517 | 0.8839608 | 0.7142929 | 0.9081135 | 0.9997927 |
| home\_MAT:garden\_MAP | 0.3594856 | 0.8839608 | 0.6562517 | 0.7420479 | 0.0517640 | 0.6831969 | 0.3873276 | 0.4331327 | 0.6302118 | 0.9497591 | 0.5538317 | 0.8504618 | 0.0094126 | 0.1127670 | 0.2259822 | 0.6409043 | 0.9497591 | 0.9293429 | 0.9641533 | 0.9006483 | 0.9497591 | 0.9972006 | 0.9497591 | 0.8243195 | 0.5279324 | 0.4034730 |
| garden\_MAT:genetic\_PC1 | 0.8839608 | 0.8504618 | 0.8243195 | 0.9156182 | 0.5519882 | 0.7231307 | 0.9641533 | 0.9081135 | 0.8963968 | 0.9947381 | 0.9081135 | 0.6253987 | 0.3409680 | 0.5279324 | 0.4034730 | 0.9477242 | 0.8909461 | 0.6562517 | 0.0701601 | 0.2259822 | 0.9497591 | 0.9660977 | 0.7420479 | 0.9497591 | 0.2967393 | 0.0141326 |
| garden\_MAT:genetic\_PC2 | 0.9497591 | 0.8956653 | 0.7231307 | 0.7204388 | 0.0142116 | 0.2259822 | 0.7300443 | 0.8319386 | 0.7420479 | 0.7901870 | 0.8839608 | 0.9497591 | 0.0063268 | 0.0141326 | 0.8504618 | 0.6562517 | 0.9081135 | 0.9896309 | 0.9278257 | 0.9081135 | 0.3873276 | 0.7231307 | 0.7561167 | 0.9497591 | 0.9497591 | 0.9497591 |
| garden\_MAT:genetic\_PC3 | 0.9997927 | 0.9888578 | 0.9568635 | 0.9156182 | 0.7231307 | 0.9497591 | 0.9497591 | 0.9497591 | 0.8243195 | 0.4499788 | 0.6113780 | 0.9497591 | 0.4034730 | 0.5519882 | 0.8839608 | 0.9497591 | 0.9497591 | 0.8839608 | 0.9947381 | 0.9972006 | 0.0445429 | 0.2026187 | 0.9641533 | 0.9497591 | 0.7420479 | 0.5279324 |
| garden\_MAP:genetic\_PC1 | 0.9477242 | 0.9568635 | 0.9045102 | 0.9293429 | 0.9497591 | 0.9497591 | 0.7561167 | 0.3873276 | 0.8576184 | 0.9578852 | 0.9497591 | 0.9293429 | 0.0078227 | 0.0032772 | 0.8552363 | 0.8969622 | 0.7422474 | 0.9578852 | 0.8504618 | 0.9497591 | 0.8504618 | 0.9497591 | 0.6562517 | 0.4331327 | 0.0142672 | 0.0116704 |
| garden\_MAP:genetic\_PC2 | 0.9709058 | 0.9497591 | 0.9477242 | 0.9006483 | 0.3276007 | 0.8909461 | 0.9271710 | 0.9497591 | 0.8839608 | 0.9497591 | 0.9568635 | 0.9081135 | 0.0157332 | 0.1082516 | 0.9006483 | 0.2269557 | 0.9497591 | 0.6113780 | 0.9081135 | 0.8853214 | 0.6967308 | 0.7420479 | 0.9156182 | 0.9156182 | 0.8839608 | 0.9081135 |
| garden\_MAP:genetic\_PC3 | 0.8963968 | 0.9497591 | 0.8963968 | 0.9156182 | 0.7766957 | 0.9497591 | 0.9497591 | 0.9997927 | 0.9006483 | 0.8639394 | 0.5519882 | 0.8839608 | 0.1408440 | 0.4331327 | 0.7420479 | 0.7901870 | 0.9578852 | 0.9896309 | 0.7766957 | 0.8243195 | 0.6799905 | 0.5986666 | 0.9081135 | 0.7231307 | 0.2259822 | 0.1913357 |

##### 2.1.4 Test daylength effects on fall phenology

Daylength could be determining when genotypes set bud - check whether
the daylength of budset is the same for a given genotype, or varies
across gardens, which would suggest that other environmental factors
contribute to budset timing

```
# daylength of budset by genotype
# this varies across gardens, suggesting daylength is not the only factor controlling budset timing
ggplot(data = dat, aes(x = Genotype, y = daylength_Stage6_2023, color = k2_tricho))+
  geom_boxplot() +
    geom_jitter(width = 0.2, height = 0) +
    scale_color_gradient2(high = "darkolivegreen2", mid = "grey20", low = "dodgerblue2", midpoint = 0.5, name = 'P. trichocarpa\nancestry', guide = 'none') +
  facet_wrap(~transect, scales = 'free_x')
```

```
## Warning: Removed 782 rows containing non-finite outside the scale range
## (`stat_boxplot()`).
```

```
## Warning: Removed 782 rows containing missing values or values outside the scale range
## (`geom_point()`).
```

```
ggplot(data = dat, aes(x = Genotype, y = daylength_Stage8_2023, color = k2_tricho))+
  geom_boxplot() +
    geom_jitter(width = 0.2, height = 0) +
    scale_color_gradient2(high = "darkolivegreen2", mid = "grey20", low = "dodgerblue2", midpoint = 0.5, name = 'P. trichocarpa\nancestry', guide = 'none') +
  facet_wrap(~transect, scales = 'free_x')
```

```
## Warning: Removed 1081 rows containing non-finite outside the scale range
## (`stat_boxplot()`).
```

```
## Warning: Removed 1081 rows containing missing values or values outside the scale range
## (`geom_point()`).
```

```
# plot timing vs daylength by garden
ggplot(data = dat, aes(x = DOY_Stage6_2023, y = daylength_Stage6_2023, color = k2_tricho))+
    geom_vline(xintercept = 171, col = 'grey', lty = 2 )+  # day of solstice
    geom_point() +
    scale_color_gradient2(high = "darkolivegreen2", mid = "grey20", low = "dodgerblue2", midpoint = 0.5, name = 'P. trichocarpa\nancestry', guide = 'none') +
    facet_wrap(~MiniCG_Site, drop = T)
```

```
## Warning: Removed 782 rows containing missing values or values outside the scale range
## (`geom_point()`).
```

```
ggplot(data = dat, aes(x = DOY_last_budset_2023, y = daylength_last_budset_2023, color = k2_tricho))+
    geom_vline(xintercept = 171, col = 'grey', lty = 2 )+  # day of solstice
    geom_point() +
    scale_color_gradient2(high = "darkolivegreen2", mid = "grey20", low = "dodgerblue2", midpoint = 0.5, name = 'P. trichocarpa\nancestry', guide = 'none') +
    facet_wrap(~MiniCG_Site, drop = T)
```

```
## Warning: Removed 747 rows containing missing values or values outside the scale range
## (`geom_point()`).
```

```
ggplot(data = dat, aes(x = DOY_Stage6_2024, y = daylength_Stage6_2024, color = k2_tricho))+
  geom_vline(xintercept = 171, col = 'grey', lty = 2 )+  # day of solstice
    geom_point() +
    scale_color_gradient2(high = "darkolivegreen2", mid = "grey20", low = "dodgerblue2", midpoint = 0.5, name = 'P. trichocarpa\nancestry', guide = 'none') +
    facet_wrap(~MiniCG_Site_2024, drop = T)
```

```
## Warning: Removed 1027 rows containing missing values or values outside the scale range
## (`geom_point()`).
```

```
ggplot(data = dat, aes(x = DOY_last_budset_2024, y = daylength_last_budset_2024, color = k2_tricho))+
    geom_vline(xintercept = 171, col = 'grey', lty = 2 )+  # day of solstice
    geom_point() +
    scale_color_gradient2(high = "darkolivegreen2", mid = "grey20", low = "dodgerblue2", midpoint = 0.5, name = 'P. trichocarpa\nancestry', guide = 'none') +
    facet_wrap(~MiniCG_Site_2024, drop = T)
```

```
## Warning: Removed 1021 rows containing missing values or values outside the scale range
## (`geom_point()`).
```

#### 2.2 2024 data

##### 2.2.1 Setup

```
###############################################
# phenotypes
# all vars measured in 2024
vars <- c("DOY_Stage2_2024", "Stage2_cGDD_2024", "DOY_Stage3_2024", "Stage3_cGDD_2024", "DOY_Stage6_2024", "DOY_last_budset_2024",  "DOY_Stage7_2024", "stage7_presence_2024", "growing_season_days_2024")
###############################################

# setup variables and put into a dataframe

# set climate and phenotype variables
garden_MAT <- dat$garden_MAT_2024
garden_MAT_2 <- dat$garden_MAT_2024^2
home_MAT <- dat$provenance_MAT
home_MAT_2 <- dat$provenance_MAT^2
garden_MAP <- dat$garden_MAP_2024
garden_MAP_2 <- dat$garden_MAP_2024^2
home_MAP <- dat$provenance_MAP
home_MAP_2 <- dat$provenance_MAP^2
home_lat <- dat$provenance_latitude
garden_lat <- dat$garden_Latitude

# random effects
block <- as.character(interaction(dat$MiniCG_Site, dat$block, drop = T))
#block <- as.character(dat$block)
Pt <- dat$Pt
genotype <- as.character(dat$Genotype)
garden <- as.character(dat$MiniCG_Site)
genetic_PC1 <- dat$genetic_PC1
genetic_PC2 <- dat$genetic_PC2
genetic_PC3 <- dat$genetic_PC3

# put in df
df <- data.frame(dat[,vars], garden_MAT, garden_MAT_2, home_MAT, home_MAT_2, garden_MAP, garden_MAP_2, home_MAP, home_MAP_2, home_lat, garden_lat, Pt, genotype, garden, block, genetic_PC1, genetic_PC2, genetic_PC3)
rownames(df) <- dat$Unique_ID
str(df)
```

```
## 'data.frame':    1510 obs. of  26 variables:
##  $ DOY_Stage2_2024         : num  75 95 92 92 86 86 92 92 92 92 ...
##  $ Stage2_cGDD_2024        : num  418 614 584 584 531 ...
##  $ DOY_Stage3_2024         : num  92 99 95 95 92 95 99 95 95 95 ...
##  $ Stage3_cGDD_2024        : num  584 643 614 614 584 ...
##  $ DOY_Stage6_2024         : num  178 176 165 165 153 NA 165 178 165 165 ...
##  $ DOY_last_budset_2024    : num  242 243 242 243 242 243 242 243 242 243 ...
##  $ DOY_Stage7_2024         : num  NA 193 178 NA 165 176 NA 193 NA NA ...
##  $ stage7_presence_2024    : num  0 1 1 0 1 1 0 1 0 0 ...
##  $ growing_season_days_2024: num  167 148 150 151 156 157 150 151 150 151 ...
##  $ garden_MAT              : num  11.3 11.3 11.3 11.3 11.3 11.3 11.3 11.3 11.3 11.3 ...
##  $ garden_MAT_2            : num  128 128 128 128 128 ...
##  $ home_MAT                : num  2.3 2.3 3.8 3.8 5.6 5.6 3.9 3.9 1.2 1.2 ...
##  $ home_MAT_2              : num  5.29 5.29 14.44 14.44 31.36 ...
##  $ garden_MAP              : num  1427 1427 1427 1427 1427 ...
##  $ garden_MAP_2            : num  2036329 2036329 2036329 2036329 2036329 ...
##  $ home_MAP                : int  424 424 320 320 319 319 605 605 1154 1154 ...
##  $ home_MAP_2              : num  179776 179776 102400 102400 101761 ...
##  $ home_lat                : num  51.9 51.9 52.1 52.1 52 ...
##  $ garden_lat              : num  47.1 47.1 47.1 47.1 47.1 ...
##  $ Pt                      : num  0.571 0.571 0.908 0.908 0.864 ...
##  $ genotype                : chr  "206" "206" "210" "210" ...
##  $ garden                  : chr  "EVERGREEN" "EVERGREEN" "EVERGREEN" "EVERGREEN" ...
##  $ block                   : chr  "EVERGREEN.1" "EVERGREEN.2" "EVERGREEN.1" "EVERGREEN.2" ...
##  $ genetic_PC1             : num  0.00423 0.00423 -0.03207 -0.03207 -0.02393 ...
##  $ genetic_PC2             : num  0.02407 0.02407 -0.00722 -0.00722 0.01811 ...
##  $ genetic_PC3             : num  -0.0243 -0.0243 -0.0365 -0.0365 -0.0505 ...
```

##### 2.2.2 Models

```
dput(vars)
```

```
## c("DOY_Stage2_2024", "Stage2_cGDD_2024", "DOY_Stage3_2024", "Stage3_cGDD_2024", 
## "DOY_Stage6_2024", "DOY_last_budset_2024", "DOY_Stage7_2024", 
## "stage7_presence_2024", "growing_season_days_2024")
```

```
# ignore stage 7 presence here because it's binary
vars <- c("DOY_Stage2_2024", "Stage2_cGDD_2024", "DOY_Stage3_2024", "Stage3_cGDD_2024", 
"DOY_Stage6_2024",  "DOY_Stage7_2024", "DOY_last_budset_2024",
"growing_season_days_2024")

mods.24 <- list()

for(n in 1:length(vars)){
  
  var <- vars[n]
  
  print(paste('Model for:', var))

  
  #  full model with just linear interaction effects
    mod <- lmer(paste0(var, '~ garden_MAT*home_MAT + garden_MAT*home_MAP +   garden_MAP*home_MAP + garden_MAP*home_MAT + genetic_PC1*garden_MAT + genetic_PC2*garden_MAT + genetic_PC3*garden_MAT +  genetic_PC1*garden_MAP + genetic_PC2*garden_MAP + genetic_PC3*garden_MAP + (1 | genotype) + (1 | garden/block)'),
                        data = df
  )
  
  mod.info <- Anova(mod)
  
  print(mod.info)

    
  # save to list
  mods.24[[n]] <- mod
  
  p <- plot_model(mod, type = 'std', vline.color = "black", show.values = T, title = var)
  plot(p)
  
  
  # plots for significant factors
    pvals <- mod.info$`Pr(>Chisq)`
    if(min(pvals) <= 0.05){
      
      # get significant factors
      sig <- rownames(mod.info)[pvals <= 0.05]
      
      # plot for each significant factor
      for(f in 1:length(sig)){
        
        y <- var # y axis is always the trait
        
        # if it is an interaction effect, we need to include both variables in the plot
        if(grepl(':', sig[f])){
          
          # get two variables in interaction
          interaction_vars <- strsplit(sig[f], ':')[[1]]
          
          # env will start with 'garden' because it's always garden climate
          env <- grep('garden', interaction_vars, value = T)
          # the one without garden is the genetic effect, either PCs or home clim
          x <- grep('garden', interaction_vars, value = T, invert = T)
          
          p2 <- plot_int(trait = y, genetics = x, env = env, dat = df)
          #plot(p2)
          
          # if just a single variable, still plot gardens separately, but sort by MAT by default
        } else{
          
          # if single variable is garden climate, use that and plot boxplots
          if(grepl('garden', sig[f])){
            env <- sig[f]
            plot_env(trait = y, env = env, dat = df)
            
          # if single variable is a genotype characteristic, facet by garden sorted by MAT  
          } else {
            env <- 'garden_MAT'
          x <- sig[f]
           p2 <- plot_int(trait = y, genetics = x, env = env, dat = df)
          #plot(p2)
          }
        }
      }
    }
  
}
```

```
## [1] "Model for: DOY_Stage2_2024"
```

```
## Warning: Some predictor variables are on very different scales: consider
## rescaling
```

```
## Analysis of Deviance Table (Type II Wald chisquare tests)
## 
## Response: DOY_Stage2_2024
##                         Chisq Df Pr(>Chisq)  
## garden_MAT             1.7635  1    0.18419  
## home_MAT               5.3324  1    0.02093 *
## home_MAP               0.7223  1    0.39538  
## garden_MAP             0.6417  1    0.42311  
## genetic_PC1            0.1868  1    0.66560  
## genetic_PC2            1.4155  1    0.23415  
## genetic_PC3            0.0984  1    0.75375  
## garden_MAT:home_MAT    0.5947  1    0.44060  
## garden_MAT:home_MAP    0.5115  1    0.47451  
## home_MAP:garden_MAP    0.0427  1    0.83632  
## home_MAT:garden_MAP    0.4322  1    0.51093  
## garden_MAT:genetic_PC1 0.1564  1    0.69245  
## garden_MAT:genetic_PC2 0.0602  1    0.80613  
## garden_MAT:genetic_PC3 0.1927  1    0.66065  
## garden_MAP:genetic_PC1 0.2785  1    0.59770  
## garden_MAP:genetic_PC2 0.6038  1    0.43713  
## garden_MAP:genetic_PC3 0.6588  1    0.41699  
## ---
## Signif. codes:  0 '***' 0.001 '**' 0.01 '*' 0.05 '.' 0.1 ' ' 1
```

```
## `geom_smooth()` using formula = 'y ~ x'
```

```
## [1] "Model for: Stage2_cGDD_2024"
```

```
## Warning: Some predictor variables are on very different scales: consider
## rescaling
```

```
## Analysis of Deviance Table (Type II Wald chisquare tests)
## 
## Response: Stage2_cGDD_2024
##                         Chisq Df Pr(>Chisq)  
## garden_MAT             1.8018  1    0.17950  
## home_MAT               4.7553  1    0.02921 *
## home_MAP               0.2275  1    0.63336  
## garden_MAP             0.4435  1    0.50545  
## genetic_PC1            0.0239  1    0.87726  
## genetic_PC2            1.5283  1    0.21637  
## genetic_PC3            0.3221  1    0.57033  
## garden_MAT:home_MAT    0.1224  1    0.72648  
## garden_MAT:home_MAP    0.8647  1    0.35242  
## home_MAP:garden_MAP    1.8834  1    0.16994  
## home_MAT:garden_MAP    0.0677  1    0.79474  
## garden_MAT:genetic_PC1 0.3834  1    0.53581  
## garden_MAT:genetic_PC2 0.0040  1    0.94962  
## garden_MAT:genetic_PC3 0.8996  1    0.34288  
## garden_MAP:genetic_PC1 0.0411  1    0.83943  
## garden_MAP:genetic_PC2 0.9814  1    0.32186  
## garden_MAP:genetic_PC3 0.0594  1    0.80739  
## ---
## Signif. codes:  0 '***' 0.001 '**' 0.01 '*' 0.05 '.' 0.1 ' ' 1
```

```
## `geom_smooth()` using formula = 'y ~ x'
```

```
## [1] "Model for: DOY_Stage3_2024"
```

```
## Warning: Some predictor variables are on very different scales: consider
## rescaling
```

```
## boundary (singular) fit: see help('isSingular')
```

```
## Analysis of Deviance Table (Type II Wald chisquare tests)
## 
## Response: DOY_Stage3_2024
##                         Chisq Df Pr(>Chisq)
## garden_MAT             2.1080  1     0.1465
## home_MAT               2.4897  1     0.1146
## home_MAP               2.1263  1     0.1448
## garden_MAP             1.2073  1     0.2719
## genetic_PC1            0.1052  1     0.7457
## genetic_PC2            1.1093  1     0.2922
## genetic_PC3            0.0827  1     0.7737
## garden_MAT:home_MAT    0.9061  1     0.3411
## garden_MAT:home_MAP    0.0042  1     0.9485
## home_MAP:garden_MAP    0.1260  1     0.7226
## home_MAT:garden_MAP    0.0510  1     0.8214
## garden_MAT:genetic_PC1 1.0455  1     0.3065
## garden_MAT:genetic_PC2 0.7897  1     0.3742
## garden_MAT:genetic_PC3 0.1683  1     0.6816
## garden_MAP:genetic_PC1 0.4487  1     0.5029
## garden_MAP:genetic_PC2 0.0347  1     0.8522
## garden_MAP:genetic_PC3 0.0187  1     0.8913
```

```
## boundary (singular) fit: see help('isSingular')
```

```
## [1] "Model for: Stage3_cGDD_2024"
```

```
## Warning: Some predictor variables are on very different scales: consider
## rescaling
```

```
## boundary (singular) fit: see help('isSingular')
```

```
## Analysis of Deviance Table (Type II Wald chisquare tests)
## 
## Response: Stage3_cGDD_2024
##                         Chisq Df Pr(>Chisq)
## garden_MAT             2.5990  1     0.1069
## home_MAT               1.6544  1     0.1984
## home_MAP               2.1808  1     0.1397
## garden_MAP             0.4866  1     0.4855
## genetic_PC1            0.0381  1     0.8452
## genetic_PC2            1.0435  1     0.3070
## genetic_PC3            0.0309  1     0.8605
## garden_MAT:home_MAT    0.4097  1     0.5221
## garden_MAT:home_MAP    0.0272  1     0.8689
## home_MAP:garden_MAP    0.5371  1     0.4636
## home_MAT:garden_MAP    0.1714  1     0.6789
## garden_MAT:genetic_PC1 1.4664  1     0.2259
## garden_MAT:genetic_PC2 0.6178  1     0.4319
## garden_MAT:genetic_PC3 0.4290  1     0.5125
## garden_MAP:genetic_PC1 0.7884  1     0.3746
## garden_MAP:genetic_PC2 0.0001  1     0.9940
## garden_MAP:genetic_PC3 0.2840  1     0.5941
```

```
## boundary (singular) fit: see help('isSingular')
```

```
## [1] "Model for: DOY_Stage6_2024"
```

```
## Warning: Some predictor variables are on very different scales: consider
## rescaling
```

```
## Analysis of Deviance Table (Type II Wald chisquare tests)
## 
## Response: DOY_Stage6_2024
##                         Chisq Df Pr(>Chisq)  
## garden_MAT             0.0707  1    0.79026  
## home_MAT               3.4277  1    0.06411 .
## home_MAP               2.6103  1    0.10617  
## garden_MAP             0.0306  1    0.86109  
## genetic_PC1            3.2062  1    0.07336 .
## genetic_PC2            0.0221  1    0.88181  
## genetic_PC3            2.1719  1    0.14056  
## garden_MAT:home_MAT    0.9822  1    0.32165  
## garden_MAT:home_MAP    0.9447  1    0.33108  
## home_MAP:garden_MAP    2.4953  1    0.11419  
## home_MAT:garden_MAP    1.7241  1    0.18916  
## garden_MAT:genetic_PC1 0.2427  1    0.62229  
## garden_MAT:genetic_PC2 3.1344  1    0.07665 .
## garden_MAT:genetic_PC3 2.6185  1    0.10562  
## garden_MAP:genetic_PC1 0.7610  1    0.38300  
## garden_MAP:genetic_PC2 0.0793  1    0.77829  
## garden_MAP:genetic_PC3 3.6762  1    0.05520 .
## ---
## Signif. codes:  0 '***' 0.001 '**' 0.01 '*' 0.05 '.' 0.1 ' ' 1
```

```
## [1] "Model for: DOY_Stage7_2024"
```

```
## Warning: Some predictor variables are on very different scales: consider
## rescaling
```

```
## boundary (singular) fit: see help('isSingular')
```

```
## Analysis of Deviance Table (Type II Wald chisquare tests)
## 
## Response: DOY_Stage7_2024
##                         Chisq Df Pr(>Chisq)
## garden_MAT             0.0230  1     0.8796
## home_MAT               1.3622  1     0.2432
## home_MAP               2.1166  1     0.1457
## garden_MAP             0.0311  1     0.8600
## genetic_PC1            0.6791  1     0.4099
## genetic_PC2            0.1537  1     0.6951
## genetic_PC3            2.0829  1     0.1490
## garden_MAT:home_MAT    0.9257  1     0.3360
## garden_MAT:home_MAP    0.8083  1     0.3686
## home_MAP:garden_MAP    0.9700  1     0.3247
## home_MAT:garden_MAP    0.0340  1     0.8537
## garden_MAT:genetic_PC1 0.0110  1     0.9164
## garden_MAT:genetic_PC2 0.4655  1     0.4951
## garden_MAT:genetic_PC3 0.5814  1     0.4458
## garden_MAP:genetic_PC1 1.2308  1     0.2673
## garden_MAP:genetic_PC2 0.2533  1     0.6148
## garden_MAP:genetic_PC3 2.5356  1     0.1113
```

```
## boundary (singular) fit: see help('isSingular')
```

```
## [1] "Model for: DOY_last_budset_2024"
```

```
## Warning: Some predictor variables are on very different scales: consider
## rescaling
```

```
## boundary (singular) fit: see help('isSingular')
```

```
## Analysis of Deviance Table (Type II Wald chisquare tests)
## 
## Response: DOY_last_budset_2024
##                          Chisq Df Pr(>Chisq)    
## garden_MAT              0.1139  1  0.7357641    
## home_MAT                0.8161  1  0.3663182    
## home_MAP                0.0012  1  0.9728587    
## garden_MAP              0.0015  1  0.9691503    
## genetic_PC1            18.9169  1  1.365e-05 ***
## genetic_PC2            12.0350  1  0.0005221 ***
## genetic_PC3             2.5393  1  0.1110409    
## garden_MAT:home_MAT     0.0570  1  0.8112585    
## garden_MAT:home_MAP     0.0988  1  0.7532631    
## home_MAP:garden_MAP     0.0002  1  0.9896962    
## home_MAT:garden_MAP     0.8671  1  0.3517691    
## garden_MAT:genetic_PC1  1.4856  1  0.2228979    
## garden_MAT:genetic_PC2  2.4736  1  0.1157725    
## garden_MAT:genetic_PC3  5.1058  1  0.0238457 *  
## garden_MAP:genetic_PC1  0.0702  1  0.7910072    
## garden_MAP:genetic_PC2  0.4011  1  0.5265353    
## garden_MAP:genetic_PC3  0.4341  1  0.5099900    
## ---
## Signif. codes:  0 '***' 0.001 '**' 0.01 '*' 0.05 '.' 0.1 ' ' 1
```

```
## boundary (singular) fit: see help('isSingular')
```

```
## `geom_smooth()` using formula = 'y ~ x'
```

```
## `geom_smooth()` using formula = 'y ~ x'
```

```
## `geom_smooth()` using formula = 'y ~ x'
```

```
## [1] "Model for: growing_season_days_2024"
```

```
## Warning: Some predictor variables are on very different scales: consider
## rescaling
```

```
## boundary (singular) fit: see help('isSingular')
```

```
## Analysis of Deviance Table (Type II Wald chisquare tests)
## 
## Response: growing_season_days_2024
##                          Chisq Df Pr(>Chisq)    
## garden_MAT              0.3644  1  0.5460918    
## home_MAT                0.0117  1  0.9136873    
## home_MAP                1.2152  1  0.2703015    
## garden_MAP              0.0065  1  0.9359408    
## genetic_PC1            16.2749  1  5.479e-05 ***
## genetic_PC2            13.6488  1  0.0002204 ***
## genetic_PC3             0.8323  1  0.3615988    
## garden_MAT:home_MAT     0.0158  1  0.9000894    
## garden_MAT:home_MAP     0.1583  1  0.6906950    
## home_MAP:garden_MAP     0.1757  1  0.6750542    
## home_MAT:garden_MAP     0.1641  1  0.6854023    
## garden_MAT:genetic_PC1  2.2194  1  0.1362859    
## garden_MAT:genetic_PC2  2.0047  1  0.1568090    
## garden_MAT:genetic_PC3  3.4087  1  0.0648555 .  
## garden_MAP:genetic_PC1  0.0031  1  0.9556738    
## garden_MAP:genetic_PC2  0.4981  1  0.4803535    
## garden_MAP:genetic_PC3  0.1765  1  0.6743700    
## ---
## Signif. codes:  0 '***' 0.001 '**' 0.01 '*' 0.05 '.' 0.1 ' ' 1
```

```
## boundary (singular) fit: see help('isSingular')
```

```
## `geom_smooth()` using formula = 'y ~ x'
```

```
## Warning in qt((1 - level)/2, df): NaNs produced
```

```
## Warning in max(ids, na.rm = TRUE): no non-missing arguments to max; returning
## -Inf
```

```
## `geom_smooth()` using formula = 'y ~ x'
```

```
## Warning in qt((1 - level)/2, df): NaNs produced
## Warning in qt((1 - level)/2, df): no non-missing arguments to max; returning
## -Inf
```

```
#plot_models(mods.24,std.est = T, m.labels = vars, p.shape = T, spacing = 1, vline.color = 'black', colors = cols25())

names(mods.24) <- vars

# setup dfs for effect sizes and p-values of each factor
# use last full model run in loop to get effect names - models in mods object have some effects reduced
rows <- names(fixef(mod))[-1] # names of fixed effects, minus intercept
# reorder row so garden clim is first
rows <- rows[c(1,4,2,3,5:17)]
colms <- vars

# effect size
df.eff.24 <- as.data.frame(matrix(ncol = length(colms), nrow = length(rows)))
colnames(df.eff.24) <- colms
rownames(df.eff.24) <- rows
# standardized effect size
df.eff.std.24 <- as.data.frame(matrix(ncol = length(colms), nrow = length(rows)))
colnames(df.eff.std.24) <- colms
rownames(df.eff.std.24) <- rows
# p values
df.pval.24 <- as.data.frame(matrix(ncol = length(colms), nrow = length(rows)))
colnames(df.pval.24) <- colms
rownames(df.pval.24) <- rows
# all p values
df.pval.all.24 <- as.data.frame(matrix(ncol = length(colms), nrow = length(rows)))
colnames(df.pval.all.24) <- colms
rownames(df.pval.all.24) <- rows

# get variables and their p-values 

for(n in 1:length(vars)){
  
  var <- vars[[n]]
  mod <- mods.24[[n]]
  mod.info <- car::Anova(mod)
  # standardize coefficients
  eff.std <- effectsize::standardize_parameters(mod)
  
  # fixed effects, minus intercept
  fes <- names(fixef(mod)[-1])
  

  
  fes <- rownames(mod.info)
  
  # get their p-values and effect sizes for each fixed effect
    for(f in 1:length(fes)){
      
      fe <- fes[[f]]
      
      # check for significance
      pval <- mod.info[fe, 'Pr(>Chisq)']
      
      if(pval <= 0.05){
        df.eff.24[fe, var] <- fixef(mods.24[[var]])[fe]
        df.eff.std.24[fe,var] <- eff.std[eff.std$Parameter == fe, 'Std_Coefficient']
        df.pval.24[fe, var] <- mod.info[fe, 'Pr(>Chisq)']
      }
  }
}
```

```
## boundary (singular) fit: see help('isSingular')
## boundary (singular) fit: see help('isSingular')
## boundary (singular) fit: see help('isSingular')
## boundary (singular) fit: see help('isSingular')
## boundary (singular) fit: see help('isSingular')
```

```
# get all pvalues so they can be adjusted
for(n in 1:length(vars)){
  
  var <- vars[[n]]
  mod <- mods.24[[n]]
  mod.info <- car::Anova(mod)

  # loop through retained variables and get their p-values and effect sizes
    for(f in 1:nrow(df.pval.all.24)){
      
      fe <- rownames(df.pval.all.24)[f]
      # get pval 
         df.pval.all.24[fe, var] <- mod.info[fe, 'Pr(>Chisq)']
         
        # for car
         
  }
}

# adjust pvals
padj <- p.adjust(unlist(df.pval.all.24), method = 'BH')
df.pval.adj.24 <- as.data.frame(matrix(padj, ncol = ncol(df.pval.all.24)))
colnames(df.pval.adj.24) <- colnames(df.pval.all.24)
rownames(df.pval.adj.24) <- rownames(df.pval.all.24)

hist(unlist(df.pval.all.24))
```

```
hist(unlist(df.pval.adj.24))
```

```
plot(unlist(df.pval.all.24), unlist(df.pval.adj.24))
```

```
plot(unlist(df.eff.std.24), unlist(df.pval.all.24))
```

```
##################
# color scale based on all effect sizes
max_col <- max(unlist(df.eff.std.24), na.rm = T)
min_col <- min(unlist(df.eff.std.24), na.rm = T)

# make scale same at both ends
max_col <- max(abs(max_col), abs(min_col))
min_col <- min(abs(max_col), abs(min_col))*-1
max_ceil <- ceiling(max_col*10)/10 # take ceiling rounded to one decimal place - used for placing ticks on legend

colf <- colorRamp2(c('red', 'white', 'blue'), breaks = c(min_col, 0, max_col))

# raw pvals
# cell_text <- df.pval
# cell_text[is.na(df.pval)] <- ''
# cell_text[df.pval > 0.05] <- ''
# cell_text[df.pval <= 0.05 & df.pval > 0.01] <- '*'
# cell_text[df.pval <= 0.01 & df.pval > 0.001] <- '**'
# cell_text[df.pval<= 0.001] <- '***'
# cell_text <- t(cell_text)

# adjusted pvalues
cell_text <- df.pval.adj.24
cell_text[is.na(df.pval.adj.24)] <- ''
cell_text[df.pval.adj.24 > 0.05] <- ''
cell_text[df.pval.adj.24 <= 0.05 & df.pval.adj.24 > 0.01] <- '*'
cell_text[df.pval.adj.24 <= 0.01 & df.pval.adj.24 > 0.001] <- '**'
cell_text[df.pval.adj.24<= 0.001] <- '***'
cell_text <- t(cell_text)


# heatmap for 2024 traits
hm24 <- Heatmap(as.matrix(t(df.eff.std.24)),
        cluster_rows = F,
        cluster_columns = F,
        rect_gp = gpar(col = "black", lwd = 1),
        col = colf,
        na_col = 'grey80',
        row_names_side = 'left',
        column_names_side = 'top',
        column_names_rot = 90,
        column_names_centered = T,
        column_title = 'Predictors',
        column_title_gp = gpar(fontface = 'bold'),
        row_title_gp = gpar(fontface = 'bold'),
        row_title = 'Trait',
        #left_annotation = ha_row,
        heatmap_legend_param = list(title = 'Std. Effect Size', at = c(max_ceil*-1, 0, max_ceil)),
        cell_fun = function(j, i, x, y, width, height, fill) { # add text to each grid
                      grid.text(cell_text[i, j], x, y, gp = gpar(cex = 1.5))}
        )


ComplexHeatmap::draw(hm24, padding = unit(c(2, 30, 2, 15), "mm"))
```

##### 2.2.3 Print model tables - 2024

```
kable(df.eff.24, label = 'Effects for 2024 traits')
```

|  | DOY\_Stage2\_2024 | Stage2\_cGDD\_2024 | DOY\_Stage3\_2024 | Stage3\_cGDD\_2024 | DOY\_Stage6\_2024 | DOY\_Stage7\_2024 | DOY\_last\_budset\_2024 | growing\_season\_days\_2024 |
| --- | --- | --- | --- | --- | --- | --- | --- | --- |
| garden\_MAT | NA | NA | NA | NA | NA | NA | NA | NA |
| garden\_MAP | NA | NA | NA | NA | NA | NA | NA | NA |
| home\_MAT | 2.392197 | 14.19508 | NA | NA | NA | NA | NA | NA |
| home\_MAP | NA | NA | NA | NA | NA | NA | NA | NA |
| genetic\_PC1 | NA | NA | NA | NA | NA | NA | -459.9173 | -698.8311 |
| genetic\_PC2 | NA | NA | NA | NA | NA | NA | -687.0127 | -754.3902 |
| genetic\_PC3 | NA | NA | NA | NA | NA | NA | NA | NA |
| garden\_MAT:home\_MAT | NA | NA | NA | NA | NA | NA | NA | NA |
| garden\_MAT:home\_MAP | NA | NA | NA | NA | NA | NA | NA | NA |
| home\_MAP:garden\_MAP | NA | NA | NA | NA | NA | NA | NA | NA |
| home\_MAT:garden\_MAP | NA | NA | NA | NA | NA | NA | NA | NA |
| garden\_MAT:genetic\_PC1 | NA | NA | NA | NA | NA | NA | NA | NA |
| garden\_MAT:genetic\_PC2 | NA | NA | NA | NA | NA | NA | NA | NA |
| garden\_MAT:genetic\_PC3 | NA | NA | NA | NA | NA | NA | -52.3884 | NA |
| garden\_MAP:genetic\_PC1 | NA | NA | NA | NA | NA | NA | NA | NA |
| garden\_MAP:genetic\_PC2 | NA | NA | NA | NA | NA | NA | NA | NA |
| garden\_MAP:genetic\_PC3 | NA | NA | NA | NA | NA | NA | NA | NA |

```
kable(df.eff.std.24, label = 'Standardized effect sizes for 2024 traits')
```

|  | DOY\_Stage2\_2024 | Stage2\_cGDD\_2024 | DOY\_Stage3\_2024 | Stage3\_cGDD\_2024 | DOY\_Stage6\_2024 | DOY\_Stage7\_2024 | DOY\_last\_budset\_2024 | growing\_season\_days\_2024 |
| --- | --- | --- | --- | --- | --- | --- | --- | --- |
| garden\_MAT | NA | NA | NA | NA | NA | NA | NA | NA |
| garden\_MAP | NA | NA | NA | NA | NA | NA | NA | NA |
| home\_MAT | 0.1899271 | 0.1379996 | NA | NA | NA | NA | NA | NA |
| home\_MAP | NA | NA | NA | NA | NA | NA | NA | NA |
| genetic\_PC1 | NA | NA | NA | NA | NA | NA | -0.1845826 | -0.1781931 |
| genetic\_PC2 | NA | NA | NA | NA | NA | NA | -0.1369484 | -0.1484453 |
| genetic\_PC3 | NA | NA | NA | NA | NA | NA | NA | NA |
| garden\_MAT:home\_MAT | NA | NA | NA | NA | NA | NA | NA | NA |
| garden\_MAT:home\_MAP | NA | NA | NA | NA | NA | NA | NA | NA |
| home\_MAP:garden\_MAP | NA | NA | NA | NA | NA | NA | NA | NA |
| home\_MAT:garden\_MAP | NA | NA | NA | NA | NA | NA | NA | NA |
| garden\_MAT:genetic\_PC1 | NA | NA | NA | NA | NA | NA | NA | NA |
| garden\_MAT:genetic\_PC2 | NA | NA | NA | NA | NA | NA | NA | NA |
| garden\_MAT:genetic\_PC3 | NA | NA | NA | NA | NA | NA | -0.1127357 | NA |
| garden\_MAP:genetic\_PC1 | NA | NA | NA | NA | NA | NA | NA | NA |
| garden\_MAP:genetic\_PC2 | NA | NA | NA | NA | NA | NA | NA | NA |
| garden\_MAP:genetic\_PC3 | NA | NA | NA | NA | NA | NA | NA | NA |

```
kable(df.pval.all.24, label = 'p-values for 2024 traits')
```

|  | DOY\_Stage2\_2024 | Stage2\_cGDD\_2024 | DOY\_Stage3\_2024 | Stage3\_cGDD\_2024 | DOY\_Stage6\_2024 | DOY\_Stage7\_2024 | DOY\_last\_budset\_2024 | growing\_season\_days\_2024 |
| --- | --- | --- | --- | --- | --- | --- | --- | --- |
| garden\_MAT | 0.1841873 | 0.1794999 | 0.1465355 | 0.1069298 | 0.7902626 | 0.8795637 | 0.7357641 | 0.5460918 |
| garden\_MAP | 0.4231079 | 0.5054503 | 0.2718669 | 0.4854702 | 0.8610925 | 0.8600474 | 0.9691503 | 0.9359408 |
| home\_MAT | 0.0209324 | 0.0292084 | 0.1145911 | 0.1983576 | 0.0641124 | 0.2431627 | 0.3663182 | 0.9136873 |
| home\_MAP | 0.3953799 | 0.6333577 | 0.1447862 | 0.1397429 | 0.1061732 | 0.1457136 | 0.9728587 | 0.2703015 |
| genetic\_PC1 | 0.6655996 | 0.8772646 | 0.7456759 | 0.8451592 | 0.0733599 | 0.4099066 | 0.0000137 | 0.0000548 |
| genetic\_PC2 | 0.2341475 | 0.2163695 | 0.2922352 | 0.3070148 | 0.8818055 | 0.6950554 | 0.0005221 | 0.0002204 |
| genetic\_PC3 | 0.7537485 | 0.5703288 | 0.7737302 | 0.8605452 | 0.1405553 | 0.1489521 | 0.1110409 | 0.3615988 |
| garden\_MAT:home\_MAT | 0.4406000 | 0.7264785 | 0.3411471 | 0.5221102 | 0.3216494 | 0.3359872 | 0.8112585 | 0.9000894 |
| garden\_MAT:home\_MAP | 0.4745070 | 0.3524235 | 0.9484771 | 0.8689298 | 0.3310826 | 0.3686232 | 0.7532631 | 0.6906950 |
| home\_MAP:garden\_MAP | 0.8363225 | 0.1699439 | 0.7226024 | 0.4636354 | 0.1141885 | 0.3246759 | 0.9896962 | 0.6750542 |
| home\_MAT:garden\_MAP | 0.5109251 | 0.7947406 | 0.8213761 | 0.6788957 | 0.1891629 | 0.8536507 | 0.3517691 | 0.6854023 |
| garden\_MAT:genetic\_PC1 | 0.6924499 | 0.5358113 | 0.3065458 | 0.2259151 | 0.6222868 | 0.9163834 | 0.2228979 | 0.1362859 |
| garden\_MAT:genetic\_PC2 | 0.8061324 | 0.9496172 | 0.3742046 | 0.4318831 | 0.0766548 | 0.4950624 | 0.1157725 | 0.1568090 |
| garden\_MAT:genetic\_PC3 | 0.6606483 | 0.3428801 | 0.6816347 | 0.5124998 | 0.1056247 | 0.4457716 | 0.0238457 | 0.0648555 |
| garden\_MAP:genetic\_PC1 | 0.5976960 | 0.8394279 | 0.5029402 | 0.3745884 | 0.3830018 | 0.2672575 | 0.7910072 | 0.9556738 |
| garden\_MAP:genetic\_PC2 | 0.4371259 | 0.3218636 | 0.8522315 | 0.9940379 | 0.7782878 | 0.6147718 | 0.5265353 | 0.4803535 |
| garden\_MAP:genetic\_PC3 | 0.4169893 | 0.8073925 | 0.8912895 | 0.5941096 | 0.0551959 | 0.1113046 | 0.5099900 | 0.6743700 |

```
kable(df.pval.adj.24, label = 'Adjusted p-values for 2024 traits')
```

|  | DOY\_Stage2\_2024 | Stage2\_cGDD\_2024 | DOY\_Stage3\_2024 | Stage3\_cGDD\_2024 | DOY\_Stage6\_2024 | DOY\_Stage7\_2024 | DOY\_last\_budset\_2024 | growing\_season\_days\_2024 |
| --- | --- | --- | --- | --- | --- | --- | --- | --- |
| garden\_MAT | 0.8039425 | 0.8039425 | 0.7502774 | 0.7502774 | 0.9671416 | 0.9671416 | 0.9671416 | 0.9168949 |
| garden\_MAP | 0.9048498 | 0.9051944 | 0.8783452 | 0.9051944 | 0.9671416 | 0.9671416 | 0.9873790 | 0.9846336 |
| home\_MAT | 0.5405028 | 0.5674769 | 0.7502774 | 0.8174739 | 0.7502774 | 0.8702664 | 0.8783452 | 0.9736574 |
| home\_MAP | 0.8961945 | 0.9671416 | 0.7502774 | 0.7502774 | 0.7502774 | 0.7502774 | 0.9873790 | 0.8783452 |
| genetic\_PC1 | 0.9671416 | 0.9671416 | 0.9671416 | 0.9671416 | 0.7502774 | 0.9048498 | 0.0018569 | 0.0037254 |
| genetic\_PC2 | 0.8606502 | 0.8534569 | 0.8783452 | 0.8783452 | 0.9671416 | 0.9671416 | 0.0177515 | 0.0099904 |
| genetic\_PC3 | 0.9671416 | 0.9459112 | 0.9671416 | 0.9671416 | 0.7502774 | 0.7502774 | 0.7502774 | 0.8783452 |
| garden\_MAT:home\_MAT | 0.9048498 | 0.9671416 | 0.8783452 | 0.9064405 | 0.8783452 | 0.8783452 | 0.9671416 | 0.9715251 |
| garden\_MAT:home\_MAP | 0.9051944 | 0.8783452 | 0.9846336 | 0.9671416 | 0.8783452 | 0.8783452 | 0.9671416 | 0.9671416 |
| home\_MAP:garden\_MAP | 0.9671416 | 0.7969783 | 0.9671416 | 0.9051944 | 0.7502774 | 0.8783452 | 0.9940379 | 0.9671416 |
| home\_MAT:garden\_MAP | 0.9051944 | 0.9671416 | 0.9671416 | 0.9671416 | 0.8039425 | 0.9671416 | 0.8783452 | 0.9671416 |
| garden\_MAT:genetic\_PC1 | 0.9671416 | 0.9108791 | 0.8783452 | 0.8534569 | 0.9671416 | 0.9736574 | 0.8534569 | 0.7502774 |
| garden\_MAT:genetic\_PC2 | 0.9671416 | 0.9846336 | 0.8783452 | 0.9048498 | 0.7502774 | 0.9051944 | 0.7502774 | 0.7616435 |
| garden\_MAT:genetic\_PC3 | 0.9671416 | 0.8783452 | 0.9671416 | 0.9051944 | 0.7502774 | 0.9048498 | 0.5405028 | 0.7502774 |
| garden\_MAP:genetic\_PC1 | 0.9671416 | 0.9671416 | 0.9051944 | 0.8783452 | 0.8828515 | 0.8783452 | 0.9671416 | 0.9846336 |
| garden\_MAP:genetic\_PC2 | 0.9048498 | 0.8783452 | 0.9671416 | 0.9940379 | 0.9671416 | 0.9671416 | 0.9064405 | 0.9051944 |
| garden\_MAP:genetic\_PC3 | 0.9048498 | 0.9671416 | 0.9697230 | 0.9671416 | 0.7502774 | 0.7502774 | 0.9051944 | 0.9671416 |

### 3 Summary heatmap

```
# check that effects match
cbind(rownames(df.eff.std.23), rownames(df.eff.std.24))
```

```
##       [,1]                     [,2]                    
##  [1,] "garden_MAT"             "garden_MAT"            
##  [2,] "garden_MAP"             "garden_MAP"            
##  [3,] "home_MAT"               "home_MAT"              
##  [4,] "home_MAP"               "home_MAP"              
##  [5,] "genetic_PC1"            "genetic_PC1"           
##  [6,] "genetic_PC2"            "genetic_PC2"           
##  [7,] "genetic_PC3"            "genetic_PC3"           
##  [8,] "garden_MAT:home_MAT"    "garden_MAT:home_MAT"   
##  [9,] "garden_MAT:home_MAP"    "garden_MAT:home_MAP"   
## [10,] "home_MAP:garden_MAP"    "home_MAP:garden_MAP"   
## [11,] "home_MAT:garden_MAP"    "home_MAT:garden_MAP"   
## [12,] "garden_MAT:genetic_PC1" "garden_MAT:genetic_PC1"
## [13,] "garden_MAT:genetic_PC2" "garden_MAT:genetic_PC2"
## [14,] "garden_MAT:genetic_PC3" "garden_MAT:genetic_PC3"
## [15,] "garden_MAP:genetic_PC1" "garden_MAP:genetic_PC1"
## [16,] "garden_MAP:genetic_PC2" "garden_MAP:genetic_PC2"
## [17,] "garden_MAP:genetic_PC3" "garden_MAP:genetic_PC3"
```

```
# merge effect sizes
df.eff.both <- cbind.data.frame(df.eff.std.23, df.eff.std.24)
head(df.eff.both)
```

```
##             DOY_Stage2_2023 Stage2_cGDD_2023 DOY_Stage3_2023 Stage3_cGDD_2023
## garden_MAT       -0.5692812        0.6404178      -0.5512488        0.7093145
## garden_MAP               NA               NA              NA               NA
## home_MAT          0.1589187               NA       0.1714929               NA
## home_MAP                 NA               NA              NA               NA
## genetic_PC1              NA               NA              NA               NA
## genetic_PC2              NA               NA              NA               NA
##             DOY_Stage6_2023 daylength_Stage6_2023 DOY_Stage7_2023
## garden_MAT               NA                    NA              NA
## garden_MAP               NA                    NA              NA
## home_MAT          0.1827911                    NA              NA
## home_MAP                 NA                    NA              NA
## genetic_PC1      -0.1177713                    NA              NA
## genetic_PC2              NA                    NA              NA
##             daylength_Stage7_2023 DOY_last_budset_2023
## garden_MAT                     NA                   NA
## garden_MAP                     NA                   NA
## home_MAT                       NA            0.1312286
## home_MAP                       NA                   NA
## genetic_PC1                    NA                   NA
## genetic_PC2                    NA                   NA
##             daylength_last_budset_2023 growing_season_days_2023
## garden_MAT                          NA                       NA
## garden_MAP                          NA                       NA
## home_MAT                            NA                       NA
## home_MAP                            NA                       NA
## genetic_PC1                         NA                       NA
## genetic_PC2                         NA                       NA
##             leaf_thickness_avg_mm_2023 leaf_area_cm2_2023 leaf_mass_g_2023
## garden_MAT                          NA          0.4509394        0.4342514
## garden_MAP                          NA         -0.4740569       -0.4714874
## home_MAT                            NA                 NA        0.1274059
## home_MAP                            NA                 NA               NA
## genetic_PC1                         NA         -0.0994744               NA
## genetic_PC2                         NA                 NA               NA
##             LMA_g_m2_2023 lower_stomata_pore_length_mean_um
## garden_MAT             NA                                NA
## garden_MAP             NA                                NA
## home_MAT               NA                                NA
## home_MAP               NA                                NA
## genetic_PC1            NA                                NA
## genetic_PC2            NA                         0.2559665
##             upper_stomata_pore_length_mean_um lower_stomata_density_mm2
## garden_MAT                                 NA                        NA
## garden_MAP                                 NA                        NA
## home_MAT                                   NA                        NA
## home_MAP                            0.2385671                        NA
## genetic_PC1                                NA                 0.2464168
## genetic_PC2                                NA                        NA
##             upper_stomata_density_mm2 stomata_ratio licor_gsw licor_gbw
## garden_MAT                         NA            NA        NA        NA
## garden_MAP                  0.2535938            NA        NA        NA
## home_MAT                           NA            NA        NA        NA
## home_MAP                           NA            NA        NA        NA
## genetic_PC1                        NA            NA        NA        NA
## genetic_PC2                        NA            NA        NA        NA
##             licor_ETR  licor_Fs licor_Fm. licor_PhiPS2 DOY_Stage2_2024
## garden_MAT         NA        NA        NA           NA              NA
## garden_MAP         NA        NA        NA           NA              NA
## home_MAT           NA        NA        NA           NA       0.1899271
## home_MAP           NA        NA        NA           NA              NA
## genetic_PC1        NA        NA        NA           NA              NA
## genetic_PC2        NA 0.1165161        NA           NA              NA
##             Stage2_cGDD_2024 DOY_Stage3_2024 Stage3_cGDD_2024 DOY_Stage6_2024
## garden_MAT                NA              NA               NA              NA
## garden_MAP                NA              NA               NA              NA
## home_MAT           0.1379996              NA               NA              NA
## home_MAP                  NA              NA               NA              NA
## genetic_PC1               NA              NA               NA              NA
## genetic_PC2               NA              NA               NA              NA
##             DOY_Stage7_2024 DOY_last_budset_2024 growing_season_days_2024
## garden_MAT               NA                   NA                       NA
## garden_MAP               NA                   NA                       NA
## home_MAT                 NA                   NA                       NA
## home_MAP                 NA                   NA                       NA
## genetic_PC1              NA           -0.1845826               -0.1781931
## genetic_PC2              NA           -0.1369484               -0.1484453
```

```
# merge pvalues
df.pval.adj.both <- cbind.data.frame(df.pval.adj.23, df.pval.adj.24)

rownames(df.pval.adj.both) <- rownames(df.eff.both)
colnames(df.pval.adj.both) <- colnames(df.eff.both)

df.pval.both <- cbind.data.frame(df.pval.23, df.pval.24)
rownames(df.pval.both) <- rownames(df.eff.both)
colnames(df.pval.both) <- colnames(df.eff.both)

# reorder rows
dput(colnames(df.eff.both))
```

```
## c("DOY_Stage2_2023", "Stage2_cGDD_2023", "DOY_Stage3_2023", "Stage3_cGDD_2023", 
## "DOY_Stage6_2023", "daylength_Stage6_2023", "DOY_Stage7_2023", 
## "daylength_Stage7_2023", "DOY_last_budset_2023", "daylength_last_budset_2023", 
## "growing_season_days_2023", "leaf_thickness_avg_mm_2023", "leaf_area_cm2_2023", 
## "leaf_mass_g_2023", "LMA_g_m2_2023", "lower_stomata_pore_length_mean_um", 
## "upper_stomata_pore_length_mean_um", "lower_stomata_density_mm2", 
## "upper_stomata_density_mm2", "stomata_ratio", "licor_gsw", "licor_gbw", 
## "licor_ETR", "licor_Fs", "licor_Fm.", "licor_PhiPS2", "DOY_Stage2_2024", 
## "Stage2_cGDD_2024", "DOY_Stage3_2024", "Stage3_cGDD_2024", "DOY_Stage6_2024", 
## "DOY_Stage7_2024", "DOY_last_budset_2024", "growing_season_days_2024"
## )
```

```
ord <- c("growing_season_days_2023", "growing_season_days_2024", "DOY_Stage2_2023", "Stage2_cGDD_2023", "DOY_Stage2_2024", "Stage2_cGDD_2024", "DOY_Stage3_2023", "Stage3_cGDD_2023",  "DOY_Stage3_2024", "Stage3_cGDD_2024","DOY_Stage6_2023", "DOY_Stage6_2024", "DOY_Stage7_2023", "DOY_Stage7_2024", "DOY_last_budset_2023", "DOY_last_budset_2024", 
 "leaf_thickness_avg_mm_2023", "leaf_area_cm2_2023", 
"leaf_mass_g_2023", "LMA_g_m2_2023", "lower_stomata_pore_length_mean_um", 
"upper_stomata_pore_length_mean_um", "lower_stomata_density_mm2", 
"upper_stomata_density_mm2",  "licor_gsw", "stomata_ratio", 
"licor_ETR", "licor_Fm.", "licor_Fs", "licor_gbw",  
  "licor_PhiPS2"  
)

df.eff.both <- df.eff.both[,ord]
df.pval.adj.both <- df.pval.adj.both[,ord]


# clean up trait names

trait_names <- colnames(df.eff.both)
trait_names <- gsub('Stage2_cGDD', 'Stage2_Bud_Flush_cGDD', trait_names)
trait_names <- gsub('DOY_Stage2', 'Stage2_Bud_Flush_DOY', trait_names)
trait_names <- gsub('DOY_Stage3', 'Stage3_Leaf_Emergence_DOY', trait_names)
trait_names <- gsub('DOY_Stage6', 'Stage6_First_Budset_DOY', trait_names)
trait_names <- gsub('DOY_Stage7', 'Stage7_Lammas_Growth_DOY', trait_names)
#trait_names <- gsub('stage7_presence', 'Stage7_Lammas_Growth_presence', trait_names)
trait_names <- gsub('DOY_Stage8', 'Stage8_Budset_after_Lammas_DOY', trait_names)
trait_names <- gsub('DOY_last_budset', 'Stage8_Final_Budset_DOY', trait_names)
trait_names <- gsub('growing_season_days', 'Growing_Season_Days', trait_names)

trait_names <- gsub('licor_', '', trait_names)
trait_names <- gsub('Fm.', "Fm'", trait_names)
trait_names <- gsub('upper', "Adaxial", trait_names)
trait_names <- gsub('lower', "Abaxial", trait_names)
trait_names <- gsub('stomata_ratio', "Stomata_ratio", trait_names)
trait_names <- gsub('leaf', 'Leaf', trait_names)

# remove units
trait_names <- gsub('_cm2', '', trait_names)
trait_names <- gsub('_cm', '', trait_names)
trait_names <- gsub('_mm2', '', trait_names)
trait_names <- gsub('_m2', '', trait_names)
trait_names <- gsub('_mm', '', trait_names)
trait_names <- gsub('_um', '', trait_names)
trait_names <- gsub('_g', '', trait_names)

trait_names <- gsub('_', ' ', trait_names)

# check
cbind(colnames(df.eff.both), trait_names)
```

```
##                                           trait_names                       
##  [1,] "growing_season_days_2023"          "Growing Season Days 2023"        
##  [2,] "growing_season_days_2024"          "Growing Season Days 2024"        
##  [3,] "DOY_Stage2_2023"                   "Stage2 Bud Flush DOY 2023"       
##  [4,] "Stage2_cGDD_2023"                  "Stage2 Bud Flush cGDD 2023"      
##  [5,] "DOY_Stage2_2024"                   "Stage2 Bud Flush DOY 2024"       
##  [6,] "Stage2_cGDD_2024"                  "Stage2 Bud Flush cGDD 2024"      
##  [7,] "DOY_Stage3_2023"                   "Stage3 Leaf Emergence DOY 2023"  
##  [8,] "Stage3_cGDD_2023"                  "Stage3 cGDD 2023"                
##  [9,] "DOY_Stage3_2024"                   "Stage3 Leaf Emergence DOY 2024"  
## [10,] "Stage3_cGDD_2024"                  "Stage3 cGDD 2024"                
## [11,] "DOY_Stage6_2023"                   "Stage6 First Budset DOY 2023"    
## [12,] "DOY_Stage6_2024"                   "Stage6 First Budset DOY 2024"    
## [13,] "DOY_Stage7_2023"                   "Stage7 Lammas Growth DOY 2023"   
## [14,] "DOY_Stage7_2024"                   "Stage7 Lammas Growth DOY 2024"   
## [15,] "DOY_last_budset_2023"              "Stage8 Final Budset DOY 2023"    
## [16,] "DOY_last_budset_2024"              "Stage8 Final Budset DOY 2024"    
## [17,] "leaf_thickness_avg_mm_2023"        "Leaf thickness avg 2023"         
## [18,] "leaf_area_cm2_2023"                "Leaf area 2023"                  
## [19,] "leaf_mass_g_2023"                  "Leaf mass 2023"                  
## [20,] "LMA_g_m2_2023"                     "LMA 2023"                        
## [21,] "lower_stomata_pore_length_mean_um" "Abaxial stomata pore length mean"
## [22,] "upper_stomata_pore_length_mean_um" "Adaxial stomata pore length mean"
## [23,] "lower_stomata_density_mm2"         "Abaxial stomata density"         
## [24,] "upper_stomata_density_mm2"         "Adaxial stomata density"         
## [25,] "licor_gsw"                         "gsw"                             
## [26,] "stomata_ratio"                     "Stomata ratio"                   
## [27,] "licor_ETR"                         "ETR"                             
## [28,] "licor_Fm."                         "Fm'"                             
## [29,] "licor_Fs"                          "Fs"                              
## [30,] "licor_gbw"                         "gbw"                             
## [31,] "licor_PhiPS2"                      "PhiPS2"
```

```
# rename effects
effect_names <- rownames(df.eff.both)
effect_names <- gsub('home', 'Home', effect_names)
effect_names <- gsub('garden', 'Garden', effect_names)
effect_names <- gsub('genetic', 'Genetic', effect_names)
effect_names <- gsub(':', ' x ', effect_names)
effect_names <- gsub('_', ' ', effect_names)
# check
cbind(rownames(df.eff.both), effect_names)
```

```
##                                effect_names              
##  [1,] "garden_MAT"             "Garden MAT"              
##  [2,] "garden_MAP"             "Garden MAP"              
##  [3,] "home_MAT"               "Home MAT"                
##  [4,] "home_MAP"               "Home MAP"                
##  [5,] "genetic_PC1"            "Genetic PC1"             
##  [6,] "genetic_PC2"            "Genetic PC2"             
##  [7,] "genetic_PC3"            "Genetic PC3"             
##  [8,] "garden_MAT:home_MAT"    "Garden MAT x Home MAT"   
##  [9,] "garden_MAT:home_MAP"    "Garden MAT x Home MAP"   
## [10,] "home_MAP:garden_MAP"    "Home MAP x Garden MAP"   
## [11,] "home_MAT:garden_MAP"    "Home MAT x Garden MAP"   
## [12,] "garden_MAT:genetic_PC1" "Garden MAT x Genetic PC1"
## [13,] "garden_MAT:genetic_PC2" "Garden MAT x Genetic PC2"
## [14,] "garden_MAT:genetic_PC3" "Garden MAT x Genetic PC3"
## [15,] "garden_MAP:genetic_PC1" "Garden MAP x Genetic PC1"
## [16,] "garden_MAP:genetic_PC2" "Garden MAP x Genetic PC2"
## [17,] "garden_MAP:genetic_PC3" "Garden MAP x Genetic PC3"
```

```
# adjusted pvalues
cell_text <- df.pval.adj.both
cell_text[is.na(df.pval.adj.both)] <- ''
cell_text[df.pval.adj.both > 0.05] <- ''
cell_text[df.pval.adj.both <= 0.05 & df.pval.adj.both > 0.01] <- '*'
cell_text[df.pval.adj.both <= 0.01 & df.pval.adj.both > 0.001] <- '**'
cell_text[df.pval.adj.both<= 0.001] <- '***'
cell_text <- t(cell_text)

# color scale based on all effect sizes
max_col <- max(unlist(df.eff.both), na.rm = T)
min_col <- min(unlist(df.eff.both), na.rm = T)

# make scale same at both ends
max_col <- max(abs(max_col), abs(min_col))
min_col <- min(abs(max_col), abs(min_col))*-1
max_ceil <- ceiling(max_col*10)/10 # take ceiling rounded to one decimal place - used for placing ticks on legend

colf <- colorRamp2(c('red', 'white', 'blue'), breaks = c(min_col, 0, max_col))

# column annotation - GxE categories
col_cat <-  c(rep('E', 2), rep('G', 5), rep('GxE', 10))
ha_col <- columnAnnotation(Effect_Category = col_cat,
                        show_annotation_name = F,
                            col = list(Effect_Category = c('E' = "#015b58", 'G' = "#4760b8", 'GxE' = "#380069" )))


# row annotation - trait categories
cats <- c(rep('Phenology', 16),  rep('Leaf Morphology', 4), rep('Stomata', 6), rep('Photosynthesis', 5))
cbind(colnames(df.eff.both), cats)
```

```
##                                           cats             
##  [1,] "growing_season_days_2023"          "Phenology"      
##  [2,] "growing_season_days_2024"          "Phenology"      
##  [3,] "DOY_Stage2_2023"                   "Phenology"      
##  [4,] "Stage2_cGDD_2023"                  "Phenology"      
##  [5,] "DOY_Stage2_2024"                   "Phenology"      
##  [6,] "Stage2_cGDD_2024"                  "Phenology"      
##  [7,] "DOY_Stage3_2023"                   "Phenology"      
##  [8,] "Stage3_cGDD_2023"                  "Phenology"      
##  [9,] "DOY_Stage3_2024"                   "Phenology"      
## [10,] "Stage3_cGDD_2024"                  "Phenology"      
## [11,] "DOY_Stage6_2023"                   "Phenology"      
## [12,] "DOY_Stage6_2024"                   "Phenology"      
## [13,] "DOY_Stage7_2023"                   "Phenology"      
## [14,] "DOY_Stage7_2024"                   "Phenology"      
## [15,] "DOY_last_budset_2023"              "Phenology"      
## [16,] "DOY_last_budset_2024"              "Phenology"      
## [17,] "leaf_thickness_avg_mm_2023"        "Leaf Morphology"
## [18,] "leaf_area_cm2_2023"                "Leaf Morphology"
## [19,] "leaf_mass_g_2023"                  "Leaf Morphology"
## [20,] "LMA_g_m2_2023"                     "Leaf Morphology"
## [21,] "lower_stomata_pore_length_mean_um" "Stomata"        
## [22,] "upper_stomata_pore_length_mean_um" "Stomata"        
## [23,] "lower_stomata_density_mm2"         "Stomata"        
## [24,] "upper_stomata_density_mm2"         "Stomata"        
## [25,] "licor_gsw"                         "Stomata"        
## [26,] "stomata_ratio"                     "Stomata"        
## [27,] "licor_ETR"                         "Photosynthesis" 
## [28,] "licor_Fm."                         "Photosynthesis" 
## [29,] "licor_Fs"                          "Photosynthesis" 
## [30,] "licor_gbw"                         "Photosynthesis" 
## [31,] "licor_PhiPS2"                      "Photosynthesis"
```

```
# choose colors
ha_row <- rowAnnotation(Trait_Category = cats,
                        show_annotation_name = F,
                            col = list(Trait_Category = c('Phenology' = "#e2e260", 'Photosynthesis' = "#2d4030", 'Leaf Morphology' = "#516823", 'Stomata' =  "#88a2b9")))


# combined heatmap
hm_both <- Heatmap(as.matrix(t(df.eff.both)),
        cluster_rows = F,
        cluster_columns = F,
        rect_gp = gpar(col = "black", lwd = 1),
        col = colf,
        na_col = 'grey80',
        row_names_side = 'left',
        row_labels = trait_names,
        column_names_side = 'top',
        column_names_rot = 90,
        column_names_centered = F,
        column_title = 'Trait Predictors',
        column_title_gp = gpar(fontface = 'bold'),
        column_labels = effect_names,
        row_title_gp = gpar(fontface = 'bold'),
        row_title = 'Trait',
        left_annotation = ha_row,
        top_annotation = ha_col,
        heatmap_legend_param = list(title = 'Std. Effect Size', at = c(max_ceil*-1, 0, max_ceil)),
        cell_fun = function(j, i, x, y, width, height, fill) { # add text to each grid
                      grid.text(cell_text[i, j], x, y, gp = gpar(cex = 1.5))}
        )


#png(file = 'results/plasticity/GxE_full_models_heatmap_2023-2024.png', height = 10, width = 12, res = 300, units = 'in')
#pdf(file = 'results/plasticity/GxE_full_models_heatmap_2023-2024.pdf', height = 10, width = 12)
ComplexHeatmap::draw(hm_both, padding = unit(c(2, 30, 2, 15), "mm"))
```

```
#dev.off()


# save heatmap object and matrix used for it
# can use to combine with plasticity heatmap
gxe_hm <- list(effect = df.eff.both,
               pval = df.pval.both,
               pval_adj = df.pval.adj.both,
               hm_text = cell_text,
               row_annot = ha_row,
               col_annot = ha_col,
               heatmap = hm_both)


#save(gxe_hm, file = 'results/plasticity/GxE_traits_heatmap_data.Rdata')
```
