## Supplementary material for "Phenotypic plasticity evolved for climate variability constrains performance under climate warming": Analysis Rmarkdown Files: plasticity_vs_ancestry_and_climate_correlations.html

Plasticity variation by genotype


### Plasticity variation by genotype

###### Alayna Mead

#### 2026-03-12

- 1 Setup
- 2 Preliminary correlation
  matrix
- 3 Relationship between plasticity and
  home climate / species ancestry
  - 3.1 Statistical models
  - 3.2 Tables with results
  - 3.3 Plots of variables with that
    significantly predict trait plasticity
  - 3.4 Heatmap of home-climate
    relationships
- 4 Relationship between plasticity and
  home climate, adjusted for species ancestry
  - 4.1 Tables with results
- 5 Heatmaps summarizing significant
  relationships
  - 5.1 Plot plasticity heatmaps
    together
  - 5.2 Merge with GxE heatmap
  - 5.3 All heatmaps

### 1 Setup

```
library(ggplot2)
library(gridExtra) # for plot panels
library(cowplot) # for extracting legend from plots
library(viridis) # colors
```

```
## Loading required package: viridisLite
```

```
library(circlize) # color palette
```

```
## ========================================
## circlize version 0.4.16
## CRAN page: https://cran.r-project.org/package=circlize
## Github page: https://github.com/jokergoo/circlize
## Documentation: https://jokergoo.github.io/circlize_book/book/
## 
## If you use it in published research, please cite:
## Gu, Z. circlize implements and enhances circular visualization
##   in R. Bioinformatics 2014.
## 
## This message can be suppressed by:
##   suppressPackageStartupMessages(library(circlize))
## ========================================
```

```
library(bbmle) # AICtab
```

```
## Loading required package: stats4
```

```
library(ComplexHeatmap) # heatmaps
```

```
## Loading required package: grid
```

```
## ========================================
## ComplexHeatmap version 2.24.1
## Bioconductor page: http://bioconductor.org/packages/ComplexHeatmap/
## Github page: https://github.com/jokergoo/ComplexHeatmap
## Documentation: http://jokergoo.github.io/ComplexHeatmap-reference
## 
## If you use it in published research, please cite either one:
## - Gu, Z. Complex Heatmap Visualization. iMeta 2022.
## - Gu, Z. Complex heatmaps reveal patterns and correlations in multidimensional 
##     genomic data. Bioinformatics 2016.
## 
## 
## The new InteractiveComplexHeatmap package can directly export static 
## complex heatmaps into an interactive Shiny app with zero effort. Have a try!
## 
## This message can be suppressed by:
##   suppressPackageStartupMessages(library(ComplexHeatmap))
## ========================================
```

```
library(lm.beta) # standardize model coefficients
library(psych) # corPlot()
```

```
## 
## Attaching package: 'psych'
```

```
## The following objects are masked from 'package:ggplot2':
## 
##     %+%, alpha
```

```
library(knitr) # tables with kable()


# load plasticity data
load('data/clean/genotypePlasticityRDPI_2023-2024.Rdata')
dat <- rdpi
rm(rdpi)

# remove genotypes without genetic info
dat <- dat[! is.na(dat$k2_tricho),]
str(dat)
```

```
## 'data.frame':    44 obs. of  121 variables:
##  $ genotype                                                    : chr  "206" "210" "218" "233" ...
##  $ provenance_latitude                                         : num  51.9 52.1 52 53.3 55.5 ...
##  $ provenance_longitude                                        : num  -124 -123 -122 -122 -123 ...
##  $ provenance_MAT                                              : num  2.3 3.8 5.6 3.9 1.2 2.4 2.9 6.3 6.9 6.7 ...
##  $ provenance_MWMT                                             : num  13 15.6 17.5 15.4 13.5 15.2 13 17.6 18.8 18.5 ...
##  $ provenance_MCMT                                             : num  -9.6 -9.9 -8.9 -9.3 -12.2 -12.4 -6.9 -5.8 -5.8 -4.8 ...
##  $ provenance_TD                                               : num  22.6 25.5 26.4 24.7 25.7 27.6 19.9 23.4 24.6 23.3 ...
##  $ provenance_MAP                                              : int  424 320 319 605 1154 512 990 368 354 796 ...
##  $ provenance_MSP                                              : int  183 164 173 283 346 251 214 137 167 316 ...
##  $ provenance_AHM                                              : num  29.1 43.3 48.9 23 9.7 24.2 13 44.5 47.7 21 ...
##  $ provenance_SHM                                              : num  71.1 94.8 101.4 54.4 39.1 ...
##  $ provenance_DD_0                                             : int  1064 991 791 927 1393 1345 850 585 578 525 ...
##  $ provenance_DD5                                              : int  925 1321 1673 1261 913 1218 884 1681 1860 1756 ...
##  $ provenance_DD_18                                            : int  5696 5172 4590 5152 6113 5695 5501 4318 4177 4213 ...
##  $ provenance_DD18                                             : int  11 32 81 29 14 27 9 84 134 117 ...
##  $ provenance_NFFD                                             : int  121 145 171 162 140 150 136 179 188 191 ...
##  $ provenance_bFFP                                             : int  176 162 147 152 166 161 176 150 139 142 ...
##  $ provenance_eFFP                                             : int  239 249 261 256 248 252 252 264 267 273 ...
##  $ provenance_FFP                                              : int  62 87 114 104 82 92 76 113 128 131 ...
##  $ provenance_PAS                                              : int  178 102 83 211 676 203 547 114 87 188 ...
##  $ provenance_EMT                                              : num  -44.2 -42.6 -38.7 -39.7 -42.9 -44.3 -36.6 -34.5 -34 -32.4 ...
##  $ provenance_EXT                                              : num  33.4 36.2 37.3 33.9 31.4 34.1 32.7 38.3 39.1 38.4 ...
##  $ provenance_MAR                                              : num  11.5 11.3 12 11.2 10.4 11.2 13.5 10.3 11.5 13 ...
##  $ provenance_Eref                                             : int  581 651 737 577 485 556 553 761 788 752 ...
##  $ provenance_CMD                                              : int  346 449 515 243 106 261 256 559 562 286 ...
##  $ provenance_RH                                               : int  52 52 54 59 63 57 61 56 57 60 ...
##  $ ID                                                          : chr  "206_S39_L001" "210_S57_L001" "218_S47_L001" "233_S93_L002" ...
##  $ k2_balsam                                                   : num  0.4267 0.0974 0.1409 0.046 0.2821 ...
##  $ k2_tricho                                                   : num  0.573 0.903 0.859 0.954 0.718 ...
##  $ k3_balsam                                                   : num  0.38969 0.07606 0.11274 0.00001 0.24774 ...
##  $ k3_coastal_tricho                                           : num  0.505 0.41 0.582 0.778 0.638 ...
##  $ k3_interior_tricho                                          : num  0.105 0.514 0.305 0.222 0.114 ...
##  $ genetic_PC1                                                 : num  0.00423 -0.03207 -0.02393 -0.03496 -0.00995 ...
##  $ genetic_PC2                                                 : num  0.02407 -0.00722 0.01811 0.039 0.03374 ...
##  $ genetic_PC3                                                 : num  -0.0243 -0.0365 -0.0505 -0.0418 -0.0256 ...
##  $ genetic_PC4                                                 : num  0.0378 0.0122 0.046 0.0279 0.0399 ...
##  $ genetic_PC5                                                 : num  0.00732 0.02208 0.01169 -0.02046 -0.00303 ...
##  $ genetic_PC6                                                 : num  0.005351 0.010408 -0.000551 -0.000388 -0.0039 ...
##  $ provenance_climate_PC1                                      : num  0.00733 -0.00678 -0.01474 0.00574 0.03686 ...
##  $ provenance_climate_PC2                                      : num  0.0295 0.0417 0.0262 0.0018 -0.0499 ...
##  $ provenance_climate_PC3                                      : num  0.0445 0.0469 0.0398 0.0123 -0.0213 ...
##  $ provenance_climate_PC4                                      : num  -0.0558 -0.02824 0.01608 0.00353 -0.03972 ...
##  $ provenance_climate_PC5                                      : num  -0.04995 -0.00471 0.01885 -0.01826 0.03115 ...
##  $ provenance_climate_PC6                                      : num  -0.02587 -0.01913 0.0176 0.00291 0.03076 ...
##  $ provenance_climate_PC7                                      : num  -0.01978 -0.02843 -0.00546 0.017 0.01689 ...
##  $ provenance_climate_PC8                                      : num  -0.0198 -0.0408 -0.0323 -0.0238 -0.0208 ...
##  $ provenance_climate_PC9                                      : num  -0.03435 -0.04177 -0.04709 -0.0322 -0.00429 ...
##  $ color_Pt                                                    : chr  "#464B3CFF" "#A0C65FFF" "#94B55AFF" "#AFDB64FF" ...
##  $ color_k3                                                    : chr  "#1B8163" "#836913" "#4E941D" "#39C600" ...
##  $ interspecific_heterozygosity                                : num  0.383 0.165 0.213 0.139 0.257 ...
##  $ heterozygosity                                              : num  0.0996 0.0992 0.0925 0.0989 0.0942 ...
##  $ plasticity_rdpi_GrowthIncrement_2023_log                    : num  -6.321 0.177 0.166 0.166 -5.686 ...
##  $ plasticity_ppi_GrowthIncrement_2023_log                     : num  8.72 2.82 4.48 2.77 8.46 ...
##  $ plasticity_rdpi_DOY_Stage2_2023                             : num  0.11 0.137 0.182 0.276 0.055 ...
##  $ plasticity_ppi_DOY_Stage2_2023                              : num  48 48 56 96 38 48 49 95 38 106 ...
##  $ plasticity_rdpi_DOY_Stage3_2023                             : num  0.111 0.142 0.106 0.249 0.04 ...
##  $ plasticity_ppi_DOY_Stage3_2023                              : num  55 55 53 92 44 49 54 93 36 102 ...
##  $ plasticity_rdpi_DOY_Stage6_2023                             : num  0.0877 0.0636 0.0822 0.0645 0.0858 ...
##  $ plasticity_ppi_DOY_Stage6_2023                              : num  110 59 82 89 87 99 88 98 71 102 ...
##  $ plasticity_rdpi_DOY_last_budset_2023                        : num  0.0679 0.1361 0.0562 0.0911 0.1713 ...
##  $ plasticity_ppi_DOY_last_budset_2023                         : num  155 142 99 142 142 114 104 115 106 132 ...
##  $ plasticity_rdpi_stage7_presence_2023                        : num  1 1 1 1 1 1 1 1 1 1 ...
##  $ plasticity_ppi_stage7_presence_2023                         : num  1 1 1 1 1 1 1 1 1 1 ...
##  $ plasticity_rdpi_growing_season_days_2023                    : num  0.17 0.11 0.236 1 0.383 ...
##  $ plasticity_ppi_growing_season_days_2023                     : num  158 157 102 201 138 123 101 122 116 134 ...
##  $ plasticity_rdpi_DOY_Stage2_2024                             : num  0.1671 0.1869 0.1462 0.1348 0.0363 ...
##  $ plasticity_ppi_DOY_Stage2_2024                              : num  62 55 45 37 21 45 17 70 47 46 ...
##  $ plasticity_rdpi_DOY_Stage3_2024                             : num  0.1688 0.1792 0.1198 0.1246 0.0418 ...
##  $ plasticity_ppi_DOY_Stage3_2024                              : num  57 54 42 40 43 54 10 73 54 39 ...
##  $ plasticity_rdpi_DOY_Stage6_2024                             : num  0.0499 0.0474 0.0694 0.0186 0.1119 ...
##  $ plasticity_ppi_DOY_Stage6_2024                              : num  68 72 123 17 103 68 126 123 72 55 ...
##  $ plasticity_rdpi_DOY_Stage7_2023                             : num  0.0685 0.0358 0.079 0.051 NA ...
##  $ plasticity_ppi_DOY_Stage7_2023                              : num  101 46 57 76 92 39 70 78 70 89 ...
##  $ plasticity_rdpi_DOY_Stage7_2024                             : num  0.069 0.0187 0.0286 NA NA ...
##  $ plasticity_ppi_DOY_Stage7_2024                              : num  81 16 32 4 102 14 66 56 48 57 ...
##  $ plasticity_rdpi_DOY_last_budset_2024                        : num  0.0864 0.042 0.0641 0.0694 0.1003 ...
##  $ plasticity_ppi_DOY_last_budset_2024                         : num  131 85 115 82 131 81 122 83 64 108 ...
##  $ plasticity_rdpi_stage7_presence_2024                        : num  0.545 0.6 0.583 0.833 1 ...
##  $ plasticity_ppi_stage7_presence_2024                         : num  1 1 1 1 1 1 1 1 1 1 ...
##  $ plasticity_rdpi_growing_season_days_2024                    : num  0.2416 0.0911 0.1444 0.1478 0.25 ...
##  $ plasticity_ppi_growing_season_days_2024                     : num  135 88 118 91 112 93 105 94 64 124 ...
##  $ plasticity_rdpi_licor_gsw                                   : num  0.1445 0.086 0.0912 0.2825 0.206 ...
##  $ plasticity_ppi_licor_gsw                                    : num  0.303 0.177 0.213 0.239 0.518 ...
##  $ plasticity_rdpi_licor_gbw                                   : num  0.000388 0.000395 0.000351 0.000536 0.000506 ...
##  $ plasticity_ppi_licor_gbw                                    : num  0.00731 0.00683 0.00757 0.00806 0.00668 ...
##  $ plasticity_rdpi_licor_PhiPS2                                : num  0.29 0.179 0.235 0.235 0.224 ...
##  $ plasticity_ppi_licor_PhiPS2                                 : num  0.572 0.449 0.549 0.516 0.454 ...
##  $ plasticity_rdpi_licor_ETR                                   : num  0.452 0.548 0.521 0.264 0.437 ...
##  $ plasticity_ppi_licor_ETR                                    : num  172 392 310 273 235 ...
##  $ plasticity_rdpi_licor_Fs                                    : num  0.1526 0.082 0.1191 0.0726 0.1283 ...
##  $ plasticity_ppi_licor_Fs                                     : num  81.7 76.5 88.1 102.6 95.7 ...
##  $ plasticity_rdpi_licor_Fm.                                   : num  0.241 0.197 0.235 0.184 0.179 ...
##  $ plasticity_ppi_licor_Fm.                                    : num  280 241 318 247 251 ...
##  $ plasticity_rdpi_leaf_thickness_avg_mm_2023                  : num  0.0718 0.0654 0.0623 0.1003 0.0715 ...
##  $ plasticity_ppi_leaf_thickness_avg_mm_2023                   : num  0.092 0.138 0.31 0.116 0.138 0.164 0.148 0.154 0.196 0.134 ...
##  $ plasticity_rdpi_LMA_g_m2_2023                               : num  0.0783 0.1012 0.0811 0.1422 0.0975 ...
##  $ plasticity_ppi_LMA_g_m2_2023                                : num  0.765 0.851 0.591 0.748 0.415 ...
##  $ plasticity_rdpi_leaf_mass_g_2023_log                        : num  -0.262 -0.265 -0.15 -0.651 -0.511 ...
##  $ plasticity_ppi_leaf_mass_g_2023_log                         : num  2.58 3.33 2.2 3.45 2.83 ...
##   [list output truncated]
```

```
# print session info, including package versions
sessionInfo()
```

```
## R version 4.5.2 (2025-10-31)
## Platform: x86_64-pc-linux-gnu
## Running under: Arch Linux
## 
## Matrix products: default
## BLAS:   /usr/lib/libblas.so.3.12.0 
## LAPACK: /usr/lib/liblapack.so.3.12.0  LAPACK version 3.12.0
## 
## locale:
##  [1] LC_CTYPE=en_US.UTF-8       LC_NUMERIC=C              
##  [3] LC_TIME=en_US.UTF-8        LC_COLLATE=en_US.UTF-8    
##  [5] LC_MONETARY=en_US.UTF-8    LC_MESSAGES=en_US.UTF-8   
##  [7] LC_PAPER=en_US.UTF-8       LC_NAME=C                 
##  [9] LC_ADDRESS=C               LC_TELEPHONE=C            
## [11] LC_MEASUREMENT=en_US.UTF-8 LC_IDENTIFICATION=C       
## 
## time zone: US/Eastern
## tzcode source: system (glibc)
## 
## attached base packages:
## [1] grid      stats4    stats     graphics  grDevices datasets  utils    
## [8] methods   base     
## 
## other attached packages:
##  [1] knitr_1.50            psych_2.5.3           lm.beta_1.7-2        
##  [4] ComplexHeatmap_2.24.1 bbmle_1.0.25.1        circlize_0.4.16      
##  [7] viridis_0.6.5         viridisLite_0.4.2     cowplot_1.2.0        
## [10] gridExtra_2.3         ggplot2_3.5.2        
## 
## loaded via a namespace (and not attached):
##  [1] sass_0.4.10         generics_0.1.4      renv_0.17.3        
##  [4] shape_1.4.6.1       lattice_0.22-7      digest_0.6.37      
##  [7] magrittr_2.0.3      RColorBrewer_1.1-3  evaluate_1.0.3     
## [10] iterators_1.0.14    mvtnorm_1.3-3       fastmap_1.2.0      
## [13] foreach_1.5.2       doParallel_1.0.17   jsonlite_2.0.0     
## [16] Matrix_1.7-4        GlobalOptions_0.1.2 scales_1.3.0       
## [19] codetools_0.2-20    numDeriv_2016.8-1.1 jquerylib_0.1.4    
## [22] mnormt_2.1.1        cli_3.6.5           crayon_1.5.3       
## [25] rlang_1.1.6         munsell_0.5.1       withr_3.0.2        
## [28] cachem_1.1.0        yaml_2.3.10         parallel_4.5.2     
## [31] tools_4.5.2         bdsmatrix_1.3-7     dplyr_1.1.4        
## [34] colorspace_2.1-1    BiocGenerics_0.54.0 GetoptLong_1.0.5   
## [37] png_0.1-8           vctrs_0.6.5         R6_2.6.1           
## [40] matrixStats_1.5.0   lifecycle_1.0.4     S4Vectors_0.46.0   
## [43] IRanges_2.42.0      clue_0.3-66         MASS_7.3-65        
## [46] cluster_2.1.8.1     pkgconfig_2.0.3     pillar_1.10.2      
## [49] bslib_0.9.0         gtable_0.3.6        glue_1.8.0         
## [52] xfun_0.52           tibble_3.2.1        tidyselect_1.2.1   
## [55] rstudioapi_0.17.1   xtable_1.8-4        rjson_0.2.23       
## [58] nlme_3.1-168        htmltools_0.5.8.1   rmarkdown_2.29     
## [61] compiler_4.5.2
```

```
# markdown settings
knitr::opts_chunk$set(fig.width = 10, fig.height = 8)
```

```
# ggplot settings
theme_set(theme_bw(base_size = 16))
```

### 2 Preliminary correlation matrix

```
vars <- c("plasticity_rdpi_GrowthIncrement_2023_log", "plasticity_rdpi_DOY_Stage2_2023", "plasticity_rdpi_DOY_Stage3_2023", "plasticity_rdpi_DOY_last_budset_2023", "plasticity_rdpi_stage7_presence_2023", "plasticity_rdpi_growing_season_days_2023", "plasticity_rdpi_DOY_Stage2_2024", "plasticity_rdpi_DOY_Stage3_2024", "plasticity_rdpi_DOY_Stage6_2024", "plasticity_rdpi_DOY_last_budset_2024", "plasticity_rdpi_growing_season_days_2024", "plasticity_rdpi_licor_gsw", "plasticity_rdpi_licor_gbw", "plasticity_rdpi_licor_PhiPS2", "plasticity_rdpi_licor_ETR", "plasticity_rdpi_licor_Fs", "plasticity_rdpi_licor_Fm.", "plasticity_rdpi_leaf_thickness_avg_mm_2023", "plasticity_rdpi_LMA_g_m2_2023", "provenance_MAT",  "provenance_MAP", "provenance_TD", "provenance_latitude", "genetic_PC1", "genetic_PC2", "genetic_PC3")

# correlation matrix
cors <- cor(dat[,vars], use = 'pairwise.complete.obs')

# heatmap of correlations
rows <- c("plasticity_rdpi_GrowthIncrement_2023_log", "plasticity_rdpi_DOY_Stage2_2023", "plasticity_rdpi_DOY_Stage3_2023", "plasticity_rdpi_DOY_last_budset_2023", "plasticity_rdpi_stage7_presence_2023", "plasticity_rdpi_growing_season_days_2023", "plasticity_rdpi_DOY_Stage2_2024", "plasticity_rdpi_DOY_Stage3_2024", "plasticity_rdpi_DOY_Stage6_2024", "plasticity_rdpi_DOY_last_budset_2024", "plasticity_rdpi_growing_season_days_2024", "plasticity_rdpi_licor_gsw", "plasticity_rdpi_licor_gbw", "plasticity_rdpi_licor_PhiPS2", "plasticity_rdpi_licor_ETR", "plasticity_rdpi_licor_Fs", "plasticity_rdpi_licor_Fm.", "plasticity_rdpi_leaf_thickness_avg_mm_2023", "plasticity_rdpi_LMA_g_m2_2023")
columns <- c("provenance_MAT",  "provenance_MAP", "provenance_TD", "provenance_latitude", "genetic_PC1", "genetic_PC2", "genetic_PC3")

# trait plasticity vs predictors
corPlot(cors[rows, columns], MAR = 16, las = 2, symmetric = FALSE)
```

```
# correlations of predictors
corPlot(cors[columns, columns], MAR = 16, las = 2)
```

```
# correlations of trait plasticity
corPlot(cors[rows, rows], MAR = 16, las = 2)
```

### 3 Relationship between plasticity and home climate / species ancestry

#### 3.1 Statistical models

Test whether home climate (MAP, MAT, and TD) and genetic structure
(PCs 1-3) predict trait plasticity using a linear regression

```
# plasticity traits
traits <- c("plasticity_rdpi_growing_season_days_2023", "plasticity_rdpi_growing_season_days_2024", "plasticity_rdpi_DOY_Stage2_2023", "plasticity_rdpi_DOY_Stage2_2024","plasticity_rdpi_DOY_Stage3_2023", "plasticity_rdpi_DOY_Stage3_2024", "plasticity_rdpi_DOY_Stage6_2023",  "plasticity_rdpi_DOY_Stage6_2024", "plasticity_rdpi_DOY_Stage7_2023",  "plasticity_rdpi_DOY_Stage7_2024", "plasticity_rdpi_stage7_presence_2023", "plasticity_rdpi_stage7_presence_2024", "plasticity_rdpi_DOY_last_budset_2023", "plasticity_rdpi_DOY_last_budset_2024",  
 "plasticity_rdpi_leaf_thickness_avg_mm_2023", "plasticity_rdpi_leaf_area_cm2_2023_log","plasticity_rdpi_leaf_mass_g_2023_log", "plasticity_rdpi_LMA_g_m2_2023", "plasticity_rdpi_lower_stomata_pore_length_mean_um", "plasticity_rdpi_upper_stomata_pore_length_mean_um", "plasticity_rdpi_lower_stomata_density_mm2",
 "plasticity_rdpi_upper_stomata_density_mm2_log", "plasticity_rdpi_upper_stomata_presence",   "plasticity_rdpi_upper_stomata_density_over_total_density_log",
 "plasticity_rdpi_licor_gsw",
  "plasticity_rdpi_licor_ETR","plasticity_rdpi_licor_Fm.",  "plasticity_rdpi_licor_Fs", "plasticity_rdpi_licor_gbw",  "plasticity_rdpi_licor_PhiPS2")


# possible predictors of plasticity
preds <- c('genetic_PC1', 'genetic_PC2', 'genetic_PC3', 'provenance_MAT', 'provenance_MAP', 'provenance_TD')

# set up dataframes for saving model info (effects and pvals) within a list
res.lr <- list() # results for linear regressions
# setup template dataframe
df <- as.data.frame(matrix(ncol = length(traits), nrow = length(preds)))
colnames(df) <- traits
rownames(df) <- preds

# add duplicate dataframes to list for saving effect size and pvalues
res.lr$eff <- df
res.lr$eff.sig <- df
res.lr$eff.std <- df
res.lr$eff.std.sig <- df
res.lr$pvals <- df
res.lr$rsq <- df
res.lr$cors <- df
res.lr$cors.sig <- df

# loop through each trait/predictor combination, run a regression, and extract info from model

for(t in 1:length(traits)){
  
  trait <- traits[t]
  
  for(p in 1:length(preds)){
    
    pred <- preds[p]
    
    # model
    form <- paste0(trait, '~', pred)
    mod <- lm(form, data = dat)
    mod.info <- summary(mod)
    print(paste('Model:', trait, '~', pred))
    print(mod.info)
    
    res.lr$rsq[pred, trait] <- mod.info$adj.r.squared
    res.lr$pvals[pred, trait] <- mod.info$coefficients[pred, 'Pr(>|t|)']
    res.lr$eff[pred, trait] <- mod.info$coefficients[pred, 'Estimate']
    
    # standardize coefficients
    # coef for predictor is second object after intercept
    res.lr$eff.std[pred,trait] <- lm.beta(mod)$standardized.coefficients[2] 
    
    print(lm.beta(mod))
    
    # also do a correlation test - used to visualize 
    # (note - correlation coefficients are the same as standard effect sizes, so this wasn't really needed)
    cor <- cor.test(dat[,pred], dat[,trait])
    res.lr$cors[pred, trait] <- cor$estimate
    
    if(res.lr$pvals[pred, trait] <= 0.05){
      res.lr$eff.sig[pred, trait] <- mod.info$coefficients[pred, 'Estimate']
      res.lr$cors.sig[pred, trait] <- cor$estimate
      res.lr$eff.std.sig[pred,trait] <- lm.beta(mod)$standardized.coefficients[2]
    }
    
  }
}
```

```
## [1] "Model: plasticity_rdpi_growing_season_days_2023 ~ genetic_PC1"
## 
## Call:
## lm(formula = form, data = dat)
## 
## Residuals:
##      Min       1Q   Median       3Q      Max 
## -0.14594 -0.09714 -0.05486  0.02009  0.75838 
## 
## Coefficients:
##             Estimate Std. Error t value Pr(>|t|)    
## (Intercept)  0.27408    0.02803   9.780 2.17e-12 ***
## genetic_PC1  0.92861    0.79096   1.174    0.247    
## ---
## Signif. codes:  0 '***' 0.001 '**' 0.01 '*' 0.05 '.' 0.1 ' ' 1
## 
## Residual standard error: 0.1841 on 42 degrees of freedom
## Multiple R-squared:  0.03177,    Adjusted R-squared:  0.008722 
## F-statistic: 1.378 on 1 and 42 DF,  p-value: 0.247
## 
## 
## Call:
## lm(formula = form, data = dat)
## 
## Standardized Coefficients::
## (Intercept) genetic_PC1 
##          NA   0.1782551 
## 
## [1] "Model: plasticity_rdpi_growing_season_days_2023 ~ genetic_PC2"
## 
## Call:
## lm(formula = form, data = dat)
## 
## Residuals:
##      Min       1Q   Median       3Q      Max 
## -0.17328 -0.09270 -0.04586  0.01399  0.73575 
## 
## Coefficients:
##             Estimate Std. Error t value Pr(>|t|)    
## (Intercept)  0.26622    0.02938   9.062 1.95e-11 ***
## genetic_PC2  0.30309    0.77937   0.389    0.699    
## ---
## Signif. codes:  0 '***' 0.001 '**' 0.01 '*' 0.05 '.' 0.1 ' ' 1
## 
## Residual standard error: 0.1867 on 42 degrees of freedom
## Multiple R-squared:  0.003588,   Adjusted R-squared:  -0.02014 
## F-statistic: 0.1512 on 1 and 42 DF,  p-value: 0.6993
## 
## 
## Call:
## lm(formula = form, data = dat)
## 
## Standardized Coefficients::
## (Intercept) genetic_PC2 
##          NA   0.0598997 
## 
## [1] "Model: plasticity_rdpi_growing_season_days_2023 ~ genetic_PC3"
## 
## Call:
## lm(formula = form, data = dat)
## 
## Residuals:
##      Min       1Q   Median       3Q      Max 
## -0.17175 -0.09699 -0.04985  0.01300  0.72464 
## 
## Coefficients:
##             Estimate Std. Error t value Pr(>|t|)    
## (Intercept)  0.26903    0.02821   9.537 4.53e-12 ***
## genetic_PC3 -0.22090    0.72546  -0.304    0.762    
## ---
## Signif. codes:  0 '***' 0.001 '**' 0.01 '*' 0.05 '.' 0.1 ' ' 1
## 
## Residual standard error: 0.1869 on 42 degrees of freedom
## Multiple R-squared:  0.002203,   Adjusted R-squared:  -0.02155 
## F-statistic: 0.09272 on 1 and 42 DF,  p-value: 0.7623
## 
## 
## Call:
## lm(formula = form, data = dat)
## 
## Standardized Coefficients::
## (Intercept) genetic_PC3 
##          NA -0.04693312 
## 
## [1] "Model: plasticity_rdpi_growing_season_days_2023 ~ provenance_MAT"
## 
## Call:
## lm(formula = form, data = dat)
## 
## Residuals:
##      Min       1Q   Median       3Q      Max 
## -0.16001 -0.09767 -0.04850  0.02274  0.73746 
## 
## Coefficients:
##                 Estimate Std. Error t value Pr(>|t|)    
## (Intercept)     0.281542   0.038854   7.246 6.48e-09 ***
## provenance_MAT -0.004872   0.010827  -0.450    0.655    
## ---
## Signif. codes:  0 '***' 0.001 '**' 0.01 '*' 0.05 '.' 0.1 ' ' 1
## 
## Residual standard error: 0.1866 on 42 degrees of freedom
## Multiple R-squared:  0.004799,   Adjusted R-squared:  -0.0189 
## F-statistic: 0.2025 on 1 and 42 DF,  p-value: 0.655
## 
## 
## Call:
## lm(formula = form, data = dat)
## 
## Standardized Coefficients::
##    (Intercept) provenance_MAT 
##             NA    -0.06927216 
## 
## [1] "Model: plasticity_rdpi_growing_season_days_2023 ~ provenance_MAP"
## 
## Call:
## lm(formula = form, data = dat)
## 
## Residuals:
##      Min       1Q   Median       3Q      Max 
## -0.19857 -0.08394 -0.04685  0.02257  0.73534 
## 
## Coefficients:
##                 Estimate Std. Error t value Pr(>|t|)   
## (Intercept)    2.169e-01  7.940e-02   2.731  0.00918 **
## provenance_MAP 9.001e-05  1.271e-04   0.708  0.48266   
## ---
## Signif. codes:  0 '***' 0.001 '**' 0.01 '*' 0.05 '.' 0.1 ' ' 1
## 
## Residual standard error: 0.186 on 42 degrees of freedom
## Multiple R-squared:  0.0118, Adjusted R-squared:  -0.01172 
## F-statistic: 0.5017 on 1 and 42 DF,  p-value: 0.4827
## 
## 
## Call:
## lm(formula = form, data = dat)
## 
## Standardized Coefficients::
##    (Intercept) provenance_MAP 
##             NA      0.1086497 
## 
## [1] "Model: plasticity_rdpi_growing_season_days_2023 ~ provenance_TD"
## 
## Call:
## lm(formula = form, data = dat)
## 
## Residuals:
##      Min       1Q   Median       3Q      Max 
## -0.15941 -0.10291 -0.04536  0.01374  0.73372 
## 
## Coefficients:
##               Estimate Std. Error t value Pr(>|t|)
## (Intercept)   0.185085   0.196036   0.944    0.350
## provenance_TD 0.003287   0.007556   0.435    0.666
## 
## Residual standard error: 0.1867 on 42 degrees of freedom
## Multiple R-squared:  0.004486,   Adjusted R-squared:  -0.01922 
## F-statistic: 0.1893 on 1 and 42 DF,  p-value: 0.6658
## 
## 
## Call:
## lm(formula = form, data = dat)
## 
## Standardized Coefficients::
##   (Intercept) provenance_TD 
##            NA    0.06697586 
## 
## [1] "Model: plasticity_rdpi_growing_season_days_2024 ~ genetic_PC1"
## 
## Call:
## lm(formula = form, data = dat)
## 
## Residuals:
##      Min       1Q   Median       3Q      Max 
## -0.10335 -0.04393 -0.01862  0.04134  0.14893 
## 
## Coefficients:
##             Estimate Std. Error t value Pr(>|t|)    
## (Intercept)  0.19139    0.01061  18.046   <2e-16 ***
## genetic_PC1  0.70770    0.30443   2.325   0.0253 *  
## ---
## Signif. codes:  0 '***' 0.001 '**' 0.01 '*' 0.05 '.' 0.1 ' ' 1
## 
## Residual standard error: 0.06733 on 40 degrees of freedom
##   (2 observations deleted due to missingness)
## Multiple R-squared:  0.119,  Adjusted R-squared:  0.097 
## F-statistic: 5.404 on 1 and 40 DF,  p-value: 0.02525
## 
## 
## Call:
## lm(formula = form, data = dat)
## 
## Standardized Coefficients::
## (Intercept) genetic_PC1 
##          NA   0.3449963 
## 
## [1] "Model: plasticity_rdpi_growing_season_days_2024 ~ genetic_PC2"
## 
## Call:
## lm(formula = form, data = dat)
## 
## Residuals:
##      Min       1Q   Median       3Q      Max 
## -0.10152 -0.04536 -0.01289  0.03406  0.16164 
## 
## Coefficients:
##             Estimate Std. Error t value Pr(>|t|)    
## (Intercept)  0.18014    0.01093  16.486   <2e-16 ***
## genetic_PC2  0.60359    0.28771   2.098   0.0423 *  
## ---
## Signif. codes:  0 '***' 0.001 '**' 0.01 '*' 0.05 '.' 0.1 ' ' 1
## 
## Residual standard error: 0.06809 on 40 degrees of freedom
##   (2 observations deleted due to missingness)
## Multiple R-squared:  0.09913,    Adjusted R-squared:  0.0766 
## F-statistic: 4.401 on 1 and 40 DF,  p-value: 0.04227
## 
## 
## Call:
## lm(formula = form, data = dat)
## 
## Standardized Coefficients::
## (Intercept) genetic_PC2 
##          NA    0.314844 
## 
## [1] "Model: plasticity_rdpi_growing_season_days_2024 ~ genetic_PC3"
## 
## Call:
## lm(formula = form, data = dat)
## 
## Residuals:
##      Min       1Q   Median       3Q      Max 
## -0.11335 -0.04423 -0.01190  0.05343  0.15541 
## 
## Coefficients:
##             Estimate Std. Error t value Pr(>|t|)    
## (Intercept)  0.18698    0.01107   16.89   <2e-16 ***
## genetic_PC3  0.15846    0.28286    0.56    0.578    
## ---
## Signif. codes:  0 '***' 0.001 '**' 0.01 '*' 0.05 '.' 0.1 ' ' 1
## 
## Residual standard error: 0.07146 on 40 degrees of freedom
##   (2 observations deleted due to missingness)
## Multiple R-squared:  0.007784,   Adjusted R-squared:  -0.01702 
## F-statistic: 0.3138 on 1 and 40 DF,  p-value: 0.5785
## 
## 
## Call:
## lm(formula = form, data = dat)
## 
## Standardized Coefficients::
## (Intercept) genetic_PC3 
##          NA  0.08822956 
## 
## [1] "Model: plasticity_rdpi_growing_season_days_2024 ~ provenance_MAT"
## 
## Call:
## lm(formula = form, data = dat)
## 
## Residuals:
##      Min       1Q   Median       3Q      Max 
## -0.10331 -0.03769 -0.02096  0.05152  0.14858 
## 
## Coefficients:
##                 Estimate Std. Error t value Pr(>|t|)    
## (Intercept)     0.202230   0.016010  12.631 1.53e-15 ***
## provenance_MAT -0.005885   0.004395  -1.339    0.188    
## ---
## Signif. codes:  0 '***' 0.001 '**' 0.01 '*' 0.05 '.' 0.1 ' ' 1
## 
## Residual standard error: 0.07018 on 40 degrees of freedom
##   (2 observations deleted due to missingness)
## Multiple R-squared:  0.04291,    Adjusted R-squared:  0.01898 
## F-statistic: 1.793 on 1 and 40 DF,  p-value: 0.1881
## 
## 
## Call:
## lm(formula = form, data = dat)
## 
## Standardized Coefficients::
##    (Intercept) provenance_MAT 
##             NA     -0.2071505 
## 
## [1] "Model: plasticity_rdpi_growing_season_days_2024 ~ provenance_MAP"
## 
## Call:
## lm(formula = form, data = dat)
## 
## Residuals:
##      Min       1Q   Median       3Q      Max 
## -0.11657 -0.05438 -0.01085  0.04342  0.15523 
## 
## Coefficients:
##                 Estimate Std. Error t value Pr(>|t|)    
## (Intercept)    1.498e-01  3.090e-02   4.849 1.92e-05 ***
## provenance_MAP 6.182e-05  4.881e-05   1.267    0.213    
## ---
## Signif. codes:  0 '***' 0.001 '**' 0.01 '*' 0.05 '.' 0.1 ' ' 1
## 
## Residual standard error: 0.07034 on 40 degrees of freedom
##   (2 observations deleted due to missingness)
## Multiple R-squared:  0.03856,    Adjusted R-squared:  0.01452 
## F-statistic: 1.604 on 1 and 40 DF,  p-value: 0.2126
## 
## 
## Call:
## lm(formula = form, data = dat)
## 
## Standardized Coefficients::
##    (Intercept) provenance_MAP 
##             NA        0.19636 
## 
## [1] "Model: plasticity_rdpi_growing_season_days_2024 ~ provenance_TD"
## 
## Call:
## lm(formula = form, data = dat)
## 
## Residuals:
##      Min       1Q   Median       3Q      Max 
## -0.11095 -0.04428 -0.01260  0.05457  0.14346 
## 
## Coefficients:
##               Estimate Std. Error t value Pr(>|t|)
## (Intercept)   0.123995   0.081589   1.520    0.136
## provenance_TD 0.002461   0.003186   0.772    0.444
## 
## Residual standard error: 0.07121 on 40 degrees of freedom
##   (2 observations deleted due to missingness)
## Multiple R-squared:  0.01469,    Adjusted R-squared:  -0.009938 
## F-statistic: 0.5966 on 1 and 40 DF,  p-value: 0.4444
## 
## 
## Call:
## lm(formula = form, data = dat)
## 
## Standardized Coefficients::
##   (Intercept) provenance_TD 
##            NA     0.1212217 
## 
## [1] "Model: plasticity_rdpi_DOY_Stage2_2023 ~ genetic_PC1"
## 
## Call:
## lm(formula = form, data = dat)
## 
## Residuals:
##      Min       1Q   Median       3Q      Max 
## -0.09680 -0.06308 -0.01531  0.01121  0.24708 
## 
## Coefficients:
##             Estimate Std. Error t value Pr(>|t|)    
## (Intercept)  0.12746    0.01231  10.354 3.93e-13 ***
## genetic_PC1  0.04018    0.34743   0.116    0.908    
## ---
## Signif. codes:  0 '***' 0.001 '**' 0.01 '*' 0.05 '.' 0.1 ' ' 1
## 
## Residual standard error: 0.08086 on 42 degrees of freedom
## Multiple R-squared:  0.0003184,  Adjusted R-squared:  -0.02348 
## F-statistic: 0.01338 on 1 and 42 DF,  p-value: 0.9085
## 
## 
## Call:
## lm(formula = form, data = dat)
## 
## Standardized Coefficients::
## (Intercept) genetic_PC1 
##          NA  0.01784328 
## 
## [1] "Model: plasticity_rdpi_DOY_Stage2_2023 ~ genetic_PC2"
## 
## Call:
## lm(formula = form, data = dat)
## 
## Residuals:
##      Min       1Q   Median       3Q      Max 
## -0.09056 -0.06324 -0.01497  0.01199  0.24516 
## 
## Coefficients:
##             Estimate Std. Error t value Pr(>|t|)    
## (Intercept)   0.1258     0.0127   9.911 1.47e-12 ***
## genetic_PC2   0.1310     0.3369   0.389    0.699    
## ---
## Signif. codes:  0 '***' 0.001 '**' 0.01 '*' 0.05 '.' 0.1 ' ' 1
## 
## Residual standard error: 0.08073 on 42 degrees of freedom
## Multiple R-squared:  0.003585,   Adjusted R-squared:  -0.02014 
## F-statistic: 0.1511 on 1 and 42 DF,  p-value: 0.6994
## 
## 
## Call:
## lm(formula = form, data = dat)
## 
## Standardized Coefficients::
## (Intercept) genetic_PC2 
##          NA  0.05987321 
## 
## [1] "Model: plasticity_rdpi_DOY_Stage2_2023 ~ genetic_PC3"
## 
## Call:
## lm(formula = form, data = dat)
## 
## Residuals:
##      Min       1Q   Median       3Q      Max 
## -0.09766 -0.06189 -0.01628  0.01058  0.24859 
## 
## Coefficients:
##             Estimate Std. Error t value Pr(>|t|)    
## (Intercept)  0.12724    0.01221  10.423 3.21e-13 ***
## genetic_PC3 -0.01149    0.31395  -0.037    0.971    
## ---
## Signif. codes:  0 '***' 0.001 '**' 0.01 '*' 0.05 '.' 0.1 ' ' 1
## 
## Residual standard error: 0.08087 on 42 degrees of freedom
## Multiple R-squared:  3.187e-05,  Adjusted R-squared:  -0.02378 
## F-statistic: 0.001339 on 1 and 42 DF,  p-value: 0.971
## 
## 
## Call:
## lm(formula = form, data = dat)
## 
## Standardized Coefficients::
##  (Intercept)  genetic_PC3 
##           NA -0.005645249 
## 
## [1] "Model: plasticity_rdpi_DOY_Stage2_2023 ~ provenance_MAT"
## 
## Call:
## lm(formula = form, data = dat)
## 
## Residuals:
##      Min       1Q   Median       3Q      Max 
## -0.09758 -0.06059 -0.01593  0.01061  0.24954 
## 
## Coefficients:
##                 Estimate Std. Error t value Pr(>|t|)    
## (Intercept)    0.1261618  0.0168347   7.494 2.88e-09 ***
## provenance_MAT 0.0004453  0.0046911   0.095    0.925    
## ---
## Signif. codes:  0 '***' 0.001 '**' 0.01 '*' 0.05 '.' 0.1 ' ' 1
## 
## Residual standard error: 0.08086 on 42 degrees of freedom
## Multiple R-squared:  0.0002145,  Adjusted R-squared:  -0.02359 
## F-statistic: 0.00901 on 1 and 42 DF,  p-value: 0.9248
## 
## 
## Call:
## lm(formula = form, data = dat)
## 
## Standardized Coefficients::
##    (Intercept) provenance_MAT 
##             NA     0.01464524 
## 
## [1] "Model: plasticity_rdpi_DOY_Stage2_2023 ~ provenance_MAP"
## 
## Call:
## lm(formula = form, data = dat)
## 
## Residuals:
##      Min       1Q   Median       3Q      Max 
## -0.10042 -0.04697 -0.01582  0.01649  0.24811 
## 
## Coefficients:
##                  Estimate Std. Error t value Pr(>|t|)    
## (Intercept)     1.579e-01  3.416e-02   4.622  3.6e-05 ***
## provenance_MAP -5.236e-05  5.466e-05  -0.958    0.344    
## ---
## Signif. codes:  0 '***' 0.001 '**' 0.01 '*' 0.05 '.' 0.1 ' ' 1
## 
## Residual standard error: 0.08 on 42 degrees of freedom
## Multiple R-squared:  0.02138,    Adjusted R-squared:  -0.001926 
## F-statistic: 0.9174 on 1 and 42 DF,  p-value: 0.3437
## 
## 
## Call:
## lm(formula = form, data = dat)
## 
## Standardized Coefficients::
##    (Intercept) provenance_MAP 
##             NA     -0.1462021 
## 
## [1] "Model: plasticity_rdpi_DOY_Stage2_2023 ~ provenance_TD"
## 
## Call:
## lm(formula = form, data = dat)
## 
## Residuals:
##      Min       1Q   Median       3Q      Max 
## -0.09100 -0.06160 -0.01338  0.01618  0.25147 
## 
## Coefficients:
##               Estimate Std. Error t value Pr(>|t|)
## (Intercept)   0.045389   0.083970   0.541    0.592
## provenance_TD 0.003189   0.003237   0.985    0.330
## 
## Residual standard error: 0.07995 on 42 degrees of freedom
## Multiple R-squared:  0.02259,    Adjusted R-squared:  -0.0006814 
## F-statistic: 0.9707 on 1 and 42 DF,  p-value: 0.3301
## 
## 
## Call:
## lm(formula = form, data = dat)
## 
## Standardized Coefficients::
##   (Intercept) provenance_TD 
##            NA     0.1503006 
## 
## [1] "Model: plasticity_rdpi_DOY_Stage2_2024 ~ genetic_PC1"
## 
## Call:
## lm(formula = form, data = dat)
## 
## Residuals:
##      Min       1Q   Median       3Q      Max 
## -0.09181 -0.04819  0.01407  0.03774  0.10849 
## 
## Coefficients:
##             Estimate Std. Error t value Pr(>|t|)    
## (Intercept)  0.11780    0.00824   14.30   <2e-16 ***
## genetic_PC1  0.04381    0.23017    0.19     0.85    
## ---
## Signif. codes:  0 '***' 0.001 '**' 0.01 '*' 0.05 '.' 0.1 ' ' 1
## 
## Residual standard error: 0.05343 on 41 degrees of freedom
##   (1 observation deleted due to missingness)
## Multiple R-squared:  0.0008828,  Adjusted R-squared:  -0.02349 
## F-statistic: 0.03623 on 1 and 41 DF,  p-value: 0.85
## 
## 
## Call:
## lm(formula = form, data = dat)
## 
## Standardized Coefficients::
## (Intercept) genetic_PC1 
##          NA  0.02971159 
## 
## [1] "Model: plasticity_rdpi_DOY_Stage2_2024 ~ genetic_PC2"
## 
## Call:
## lm(formula = form, data = dat)
## 
## Residuals:
##      Min       1Q   Median       3Q      Max 
## -0.09119 -0.04852  0.01474  0.03737  0.10638 
## 
## Coefficients:
##              Estimate Std. Error t value Pr(>|t|)    
## (Intercept)  0.117835   0.008455  13.936   <2e-16 ***
## genetic_PC2 -0.026893   0.225163  -0.119    0.906    
## ---
## Signif. codes:  0 '***' 0.001 '**' 0.01 '*' 0.05 '.' 0.1 ' ' 1
## 
## Residual standard error: 0.05344 on 41 degrees of freedom
##   (1 observation deleted due to missingness)
## Multiple R-squared:  0.0003478,  Adjusted R-squared:  -0.02403 
## F-statistic: 0.01427 on 1 and 41 DF,  p-value: 0.9055
## 
## 
## Call:
## lm(formula = form, data = dat)
## 
## Standardized Coefficients::
## (Intercept) genetic_PC2 
##          NA -0.01864987 
## 
## [1] "Model: plasticity_rdpi_DOY_Stage2_2024 ~ genetic_PC3"
## 
## Call:
## lm(formula = form, data = dat)
## 
## Residuals:
##      Min       1Q   Median       3Q      Max 
## -0.09245 -0.04826  0.01246  0.03569  0.10265 
## 
## Coefficients:
##              Estimate Std. Error t value Pr(>|t|)    
## (Intercept)  0.117180   0.008144  14.388   <2e-16 ***
## genetic_PC3 -0.122438   0.210440  -0.582    0.564    
## ---
## Signif. codes:  0 '***' 0.001 '**' 0.01 '*' 0.05 '.' 0.1 ' ' 1
## 
## Residual standard error: 0.05323 on 41 degrees of freedom
##   (1 observation deleted due to missingness)
## Multiple R-squared:  0.008189,   Adjusted R-squared:  -0.016 
## F-statistic: 0.3385 on 1 and 41 DF,  p-value: 0.5639
## 
## 
## Call:
## lm(formula = form, data = dat)
## 
## Standardized Coefficients::
## (Intercept) genetic_PC3 
##          NA -0.09049214 
## 
## [1] "Model: plasticity_rdpi_DOY_Stage2_2024 ~ provenance_MAT"
## 
## Call:
## lm(formula = form, data = dat)
## 
## Residuals:
##      Min       1Q   Median       3Q      Max 
## -0.09335 -0.04823  0.01330  0.03226  0.09626 
## 
## Coefficients:
##                Estimate Std. Error t value Pr(>|t|)    
## (Intercept)    0.110180   0.011358   9.701 3.55e-12 ***
## provenance_MAT 0.002892   0.003131   0.924    0.361    
## ---
## Signif. codes:  0 '***' 0.001 '**' 0.01 '*' 0.05 '.' 0.1 ' ' 1
## 
## Residual standard error: 0.0529 on 41 degrees of freedom
##   (1 observation deleted due to missingness)
## Multiple R-squared:  0.02039,    Adjusted R-squared:  -0.003502 
## F-statistic: 0.8534 on 1 and 41 DF,  p-value: 0.361
## 
## 
## Call:
## lm(formula = form, data = dat)
## 
## Standardized Coefficients::
##    (Intercept) provenance_MAT 
##             NA      0.1427968 
## 
## [1] "Model: plasticity_rdpi_DOY_Stage2_2024 ~ provenance_MAP"
## 
## Call:
## lm(formula = form, data = dat)
## 
## Residuals:
##      Min       1Q   Median       3Q      Max 
## -0.08934 -0.04859  0.01058  0.04211  0.10164 
## 
## Coefficients:
##                  Estimate Std. Error t value Pr(>|t|)    
## (Intercept)     1.492e-01  2.241e-02    6.66 4.99e-08 ***
## provenance_MAP -5.388e-05  3.567e-05   -1.51    0.139    
## ---
## Signif. codes:  0 '***' 0.001 '**' 0.01 '*' 0.05 '.' 0.1 ' ' 1
## 
## Residual standard error: 0.05202 on 41 degrees of freedom
##   (1 observation deleted due to missingness)
## Multiple R-squared:  0.05271,    Adjusted R-squared:  0.02961 
## F-statistic: 2.282 on 1 and 41 DF,  p-value: 0.1386
## 
## 
## Call:
## lm(formula = form, data = dat)
## 
## Standardized Coefficients::
##    (Intercept) provenance_MAP 
##             NA     -0.2295972 
## 
## [1] "Model: plasticity_rdpi_DOY_Stage2_2024 ~ provenance_TD"
## 
## Call:
## lm(formula = form, data = dat)
## 
## Residuals:
##      Min       1Q   Median       3Q      Max 
## -0.09517 -0.04755  0.01570  0.03804  0.10494 
## 
## Coefficients:
##                 Estimate Std. Error t value Pr(>|t|)  
## (Intercept)    0.1372618  0.0562321   2.441   0.0191 *
## provenance_TD -0.0007689  0.0021721  -0.354   0.7252  
## ---
## Signif. codes:  0 '***' 0.001 '**' 0.01 '*' 0.05 '.' 0.1 ' ' 1
## 
## Residual standard error: 0.05337 on 41 degrees of freedom
##   (1 observation deleted due to missingness)
## Multiple R-squared:  0.003047,   Adjusted R-squared:  -0.02127 
## F-statistic: 0.1253 on 1 and 41 DF,  p-value: 0.7252
## 
## 
## Call:
## lm(formula = form, data = dat)
## 
## Standardized Coefficients::
##   (Intercept) provenance_TD 
##            NA   -0.05519913 
## 
## [1] "Model: plasticity_rdpi_DOY_Stage3_2023 ~ genetic_PC1"
## 
## Call:
## lm(formula = form, data = dat)
## 
## Residuals:
##       Min        1Q    Median        3Q       Max 
## -0.083532 -0.033608 -0.012272  0.008273  0.229541 
## 
## Coefficients:
##             Estimate Std. Error t value Pr(>|t|)    
## (Intercept)  0.12007    0.00977  12.290 1.69e-15 ***
## genetic_PC1  0.07765    0.27573   0.282     0.78    
## ---
## Signif. codes:  0 '***' 0.001 '**' 0.01 '*' 0.05 '.' 0.1 ' ' 1
## 
## Residual standard error: 0.06417 on 42 degrees of freedom
## Multiple R-squared:  0.001885,   Adjusted R-squared:  -0.02188 
## F-statistic: 0.0793 on 1 and 42 DF,  p-value: 0.7796
## 
## 
## Call:
## lm(formula = form, data = dat)
## 
## Standardized Coefficients::
## (Intercept) genetic_PC1 
##          NA  0.04341213 
## 
## [1] "Model: plasticity_rdpi_DOY_Stage3_2023 ~ genetic_PC2"
## 
## Call:
## lm(formula = form, data = dat)
## 
## Residuals:
##       Min        1Q    Median        3Q       Max 
## -0.081257 -0.033767 -0.009966  0.008301  0.229893 
## 
## Coefficients:
##             Estimate Std. Error t value Pr(>|t|)    
## (Intercept)  0.11895    0.01010  11.781 6.75e-15 ***
## genetic_PC2  0.06833    0.26787   0.255      0.8    
## ---
## Signif. codes:  0 '***' 0.001 '**' 0.01 '*' 0.05 '.' 0.1 ' ' 1
## 
## Residual standard error: 0.06418 on 42 degrees of freedom
## Multiple R-squared:  0.001547,   Adjusted R-squared:  -0.02223 
## F-statistic: 0.06508 on 1 and 42 DF,  p-value: 0.7999
## 
## 
## Call:
## lm(formula = form, data = dat)
## 
## Standardized Coefficients::
## (Intercept) genetic_PC2 
##          NA  0.03933217 
## 
## [1] "Model: plasticity_rdpi_DOY_Stage3_2023 ~ genetic_PC3"
## 
## Call:
## lm(formula = form, data = dat)
## 
## Residuals:
##       Min        1Q    Median        3Q       Max 
## -0.084525 -0.036729 -0.008865  0.005160  0.225172 
## 
## Coefficients:
##             Estimate Std. Error t value Pr(>|t|)    
## (Intercept) 0.119953   0.009665  12.412 1.22e-15 ***
## genetic_PC3 0.131038   0.248538   0.527    0.601    
## ---
## Signif. codes:  0 '***' 0.001 '**' 0.01 '*' 0.05 '.' 0.1 ' ' 1
## 
## Residual standard error: 0.06402 on 42 degrees of freedom
## Multiple R-squared:  0.006575,   Adjusted R-squared:  -0.01708 
## F-statistic: 0.278 on 1 and 42 DF,  p-value: 0.6008
## 
## 
## Call:
## lm(formula = form, data = dat)
## 
## Standardized Coefficients::
## (Intercept) genetic_PC3 
##          NA  0.08108631 
## 
## [1] "Model: plasticity_rdpi_DOY_Stage3_2023 ~ provenance_MAT"
## 
## Call:
## lm(formula = form, data = dat)
## 
## Residuals:
##       Min        1Q    Median        3Q       Max 
## -0.085298 -0.031124 -0.008929  0.006535  0.222236 
## 
## Coefficients:
##                 Estimate Std. Error t value Pr(>|t|)    
## (Intercept)     0.126404   0.013288   9.513 4.88e-12 ***
## provenance_MAT -0.002714   0.003703  -0.733    0.468    
## ---
## Signif. codes:  0 '***' 0.001 '**' 0.01 '*' 0.05 '.' 0.1 ' ' 1
## 
## Residual standard error: 0.06382 on 42 degrees of freedom
## Multiple R-squared:  0.01263,    Adjusted R-squared:  -0.01088 
## F-statistic: 0.5373 on 1 and 42 DF,  p-value: 0.4676
## 
## 
## Call:
## lm(formula = form, data = dat)
## 
## Standardized Coefficients::
##    (Intercept) provenance_MAT 
##             NA     -0.1123857 
## 
## [1] "Model: plasticity_rdpi_DOY_Stage3_2023 ~ provenance_MAP"
## 
## Call:
## lm(formula = form, data = dat)
## 
## Residuals:
##       Min        1Q    Median        3Q       Max 
## -0.088213 -0.030630 -0.013856  0.003263  0.231478 
## 
## Coefficients:
##                  Estimate Std. Error t value Pr(>|t|)    
## (Intercept)     1.537e-01  2.684e-02   5.726 9.83e-07 ***
## provenance_MAP -5.820e-05  4.296e-05  -1.355    0.183    
## ---
## Signif. codes:  0 '***' 0.001 '**' 0.01 '*' 0.05 '.' 0.1 ' ' 1
## 
## Residual standard error: 0.06287 on 42 degrees of freedom
## Multiple R-squared:  0.04187,    Adjusted R-squared:  0.01906 
## F-statistic: 1.835 on 1 and 42 DF,  p-value: 0.1827
## 
## 
## Call:
## lm(formula = form, data = dat)
## 
## Standardized Coefficients::
##    (Intercept) provenance_MAP 
##             NA     -0.2046212 
## 
## [1] "Model: plasticity_rdpi_DOY_Stage3_2023 ~ provenance_TD"
## 
## Call:
## lm(formula = form, data = dat)
## 
## Residuals:
##       Min        1Q    Median        3Q       Max 
## -0.079776 -0.035688 -0.014100  0.005645  0.234151 
## 
## Coefficients:
##               Estimate Std. Error t value Pr(>|t|)
## (Intercept)   0.053874   0.066674   0.808    0.424
## provenance_TD 0.002563   0.002570   0.997    0.324
## 
## Residual standard error: 0.06348 on 42 degrees of freedom
## Multiple R-squared:  0.02314,    Adjusted R-squared:  -0.0001208 
## F-statistic: 0.9948 on 1 and 42 DF,  p-value: 0.3243
## 
## 
## Call:
## lm(formula = form, data = dat)
## 
## Standardized Coefficients::
##   (Intercept) provenance_TD 
##            NA     0.1521111 
## 
## [1] "Model: plasticity_rdpi_DOY_Stage3_2024 ~ genetic_PC1"
## 
## Call:
## lm(formula = form, data = dat)
## 
## Residuals:
##       Min        1Q    Median        3Q       Max 
## -0.091163 -0.029813  0.006297  0.028074  0.114393 
## 
## Coefficients:
##             Estimate Std. Error t value Pr(>|t|)    
## (Intercept) 0.121333   0.007671  15.817   <2e-16 ***
## genetic_PC1 0.177658   0.214286   0.829    0.412    
## ---
## Signif. codes:  0 '***' 0.001 '**' 0.01 '*' 0.05 '.' 0.1 ' ' 1
## 
## Residual standard error: 0.04974 on 41 degrees of freedom
##   (1 observation deleted due to missingness)
## Multiple R-squared:  0.01649,    Adjusted R-squared:  -0.0075 
## F-statistic: 0.6874 on 1 and 41 DF,  p-value: 0.4119
## 
## 
## Call:
## lm(formula = form, data = dat)
## 
## Standardized Coefficients::
## (Intercept) genetic_PC1 
##          NA   0.1284073 
## 
## [1] "Model: plasticity_rdpi_DOY_Stage3_2024 ~ genetic_PC2"
## 
## Call:
## lm(formula = form, data = dat)
## 
## Residuals:
##       Min        1Q    Median        3Q       Max 
## -0.084759 -0.025548  0.007614  0.027079  0.106021 
## 
## Coefficients:
##              Estimate Std. Error t value Pr(>|t|)    
## (Intercept)  0.121313   0.007917   15.32   <2e-16 ***
## genetic_PC2 -0.092807   0.210819   -0.44    0.662    
## ---
## Signif. codes:  0 '***' 0.001 '**' 0.01 '*' 0.05 '.' 0.1 ' ' 1
## 
## Residual standard error: 0.05004 on 41 degrees of freedom
##   (1 observation deleted due to missingness)
## Multiple R-squared:  0.004704,   Adjusted R-squared:  -0.01957 
## F-statistic: 0.1938 on 1 and 41 DF,  p-value: 0.6621
## 
## 
## Call:
## lm(formula = form, data = dat)
## 
## Standardized Coefficients::
## (Intercept) genetic_PC2 
##          NA -0.06858932 
## 
## [1] "Model: plasticity_rdpi_DOY_Stage3_2024 ~ genetic_PC3"
## 
## Call:
## lm(formula = form, data = dat)
## 
## Residuals:
##       Min        1Q    Median        3Q       Max 
## -0.095050 -0.025290  0.008667  0.024101  0.098143 
## 
## Coefficients:
##              Estimate Std. Error t value Pr(>|t|)    
## (Intercept)  0.119585   0.007519  15.904   <2e-16 ***
## genetic_PC3 -0.253554   0.194284  -1.305    0.199    
## ---
## Signif. codes:  0 '***' 0.001 '**' 0.01 '*' 0.05 '.' 0.1 ' ' 1
## 
## Residual standard error: 0.04914 on 41 degrees of freedom
##   (1 observation deleted due to missingness)
## Multiple R-squared:  0.03988,    Adjusted R-squared:  0.01647 
## F-statistic: 1.703 on 1 and 41 DF,  p-value: 0.1991
## 
## 
## Call:
## lm(formula = form, data = dat)
## 
## Standardized Coefficients::
## (Intercept) genetic_PC3 
##          NA  -0.1997116 
## 
## [1] "Model: plasticity_rdpi_DOY_Stage3_2024 ~ provenance_MAT"
## 
## Call:
## lm(formula = form, data = dat)
## 
## Residuals:
##       Min        1Q    Median        3Q       Max 
## -0.090489 -0.028964  0.008519  0.025069  0.099466 
## 
## Coefficients:
##                Estimate Std. Error t value Pr(>|t|)    
## (Intercept)    0.115290   0.010708  10.766 1.61e-13 ***
## provenance_MAT 0.001995   0.002952   0.676    0.503    
## ---
## Signif. codes:  0 '***' 0.001 '**' 0.01 '*' 0.05 '.' 0.1 ' ' 1
## 
## Residual standard error: 0.04988 on 41 degrees of freedom
##   (1 observation deleted due to missingness)
## Multiple R-squared:  0.01102,    Adjusted R-squared:  -0.0131 
## F-statistic: 0.4567 on 1 and 41 DF,  p-value: 0.503
## 
## 
## Call:
## lm(formula = form, data = dat)
## 
## Standardized Coefficients::
##    (Intercept) provenance_MAT 
##             NA      0.1049602 
## 
## [1] "Model: plasticity_rdpi_DOY_Stage3_2024 ~ provenance_MAP"
## 
## Call:
## lm(formula = form, data = dat)
## 
## Residuals:
##      Min       1Q   Median       3Q      Max 
## -0.08974 -0.02661  0.00606  0.02539  0.09406 
## 
## Coefficients:
##                  Estimate Std. Error t value Pr(>|t|)    
## (Intercept)     1.716e-01  1.984e-02   8.647 8.72e-11 ***
## provenance_MAP -8.711e-05  3.159e-05  -2.758  0.00865 ** 
## ---
## Signif. codes:  0 '***' 0.001 '**' 0.01 '*' 0.05 '.' 0.1 ' ' 1
## 
## Residual standard error: 0.04606 on 41 degrees of freedom
##   (1 observation deleted due to missingness)
## Multiple R-squared:  0.1565, Adjusted R-squared:  0.1359 
## F-statistic: 7.605 on 1 and 41 DF,  p-value: 0.008652
## 
## 
## Call:
## lm(formula = form, data = dat)
## 
## Standardized Coefficients::
##    (Intercept) provenance_MAP 
##             NA     -0.3955644 
## 
## [1] "Model: plasticity_rdpi_DOY_Stage3_2024 ~ provenance_TD"
## 
## Call:
## lm(formula = form, data = dat)
## 
## Residuals:
##       Min        1Q    Median        3Q       Max 
## -0.093889 -0.024971  0.006404  0.024605  0.110230 
## 
## Coefficients:
##               Estimate Std. Error t value Pr(>|t|)
## (Intercept)   0.082364   0.052504   1.569    0.124
## provenance_TD 0.001484   0.002028   0.732    0.468
## 
## Residual standard error: 0.04983 on 41 degrees of freedom
##   (1 observation deleted due to missingness)
## Multiple R-squared:  0.01289,    Adjusted R-squared:  -0.01118 
## F-statistic: 0.5356 on 1 and 41 DF,  p-value: 0.4684
## 
## 
## Call:
## lm(formula = form, data = dat)
## 
## Standardized Coefficients::
##   (Intercept) provenance_TD 
##            NA     0.1135557 
## 
## [1] "Model: plasticity_rdpi_DOY_Stage6_2023 ~ genetic_PC1"
## 
## Call:
## lm(formula = form, data = dat)
## 
## Residuals:
##       Min        1Q    Median        3Q       Max 
## -0.034596 -0.013238  0.000868  0.010593  0.043963 
## 
## Coefficients:
##             Estimate Std. Error t value Pr(>|t|)    
## (Intercept) 0.086571   0.002693   32.15   <2e-16 ***
## genetic_PC1 0.202908   0.075993    2.67   0.0107 *  
## ---
## Signif. codes:  0 '***' 0.001 '**' 0.01 '*' 0.05 '.' 0.1 ' ' 1
## 
## Residual standard error: 0.01769 on 42 degrees of freedom
## Multiple R-squared:  0.1451, Adjusted R-squared:  0.1248 
## F-statistic: 7.129 on 1 and 42 DF,  p-value: 0.01074
## 
## 
## Call:
## lm(formula = form, data = dat)
## 
## Standardized Coefficients::
## (Intercept) genetic_PC1 
##          NA   0.3809366 
## 
## [1] "Model: plasticity_rdpi_DOY_Stage6_2023 ~ genetic_PC2"
## 
## Call:
## lm(formula = form, data = dat)
## 
## Residuals:
##       Min        1Q    Median        3Q       Max 
## -0.032178 -0.012586  0.000096  0.014981  0.050144 
## 
## Coefficients:
##              Estimate Std. Error t value Pr(>|t|)    
## (Intercept)  0.086095   0.002995  28.742   <2e-16 ***
## genetic_PC2 -0.049113   0.079473  -0.618     0.54    
## ---
## Signif. codes:  0 '***' 0.001 '**' 0.01 '*' 0.05 '.' 0.1 ' ' 1
## 
## Residual standard error: 0.01904 on 42 degrees of freedom
## Multiple R-squared:  0.009011,   Adjusted R-squared:  -0.01458 
## F-statistic: 0.3819 on 1 and 42 DF,  p-value: 0.5399
## 
## 
## Call:
## lm(formula = form, data = dat)
## 
## Standardized Coefficients::
## (Intercept) genetic_PC2 
##          NA -0.09492715 
## 
## [1] "Model: plasticity_rdpi_DOY_Stage6_2023 ~ genetic_PC3"
## 
## Call:
## lm(formula = form, data = dat)
## 
## Residuals:
##      Min       1Q   Median       3Q      Max 
## -0.04133 -0.01522  0.00351  0.01125  0.04127 
## 
## Coefficients:
##             Estimate Std. Error t value Pr(>|t|)    
## (Intercept) 0.085871   0.002745   31.29   <2e-16 ***
## genetic_PC3 0.149614   0.070580    2.12     0.04 *  
## ---
## Signif. codes:  0 '***' 0.001 '**' 0.01 '*' 0.05 '.' 0.1 ' ' 1
## 
## Residual standard error: 0.01818 on 42 degrees of freedom
## Multiple R-squared:  0.09665,    Adjusted R-squared:  0.07514 
## F-statistic: 4.494 on 1 and 42 DF,  p-value: 0.03997
## 
## 
## Call:
## lm(formula = form, data = dat)
## 
## Standardized Coefficients::
## (Intercept) genetic_PC3 
##          NA   0.3108837 
## 
## [1] "Model: plasticity_rdpi_DOY_Stage6_2023 ~ provenance_MAT"
## 
## Call:
## lm(formula = form, data = dat)
## 
## Residuals:
##       Min        1Q    Median        3Q       Max 
## -0.034414 -0.013206  0.000516  0.008326  0.038558 
## 
## Coefficients:
##                 Estimate Std. Error t value Pr(>|t|)    
## (Intercept)     0.092996   0.003619  25.698  < 2e-16 ***
## provenance_MAT -0.003002   0.001008  -2.977  0.00482 ** 
## ---
## Signif. codes:  0 '***' 0.001 '**' 0.01 '*' 0.05 '.' 0.1 ' ' 1
## 
## Residual standard error: 0.01738 on 42 degrees of freedom
## Multiple R-squared:  0.1742, Adjusted R-squared:  0.1546 
## F-statistic: 8.861 on 1 and 42 DF,  p-value: 0.004819
## 
## 
## Call:
## lm(formula = form, data = dat)
## 
## Standardized Coefficients::
##    (Intercept) provenance_MAT 
##             NA     -0.4173931 
## 
## [1] "Model: plasticity_rdpi_DOY_Stage6_2023 ~ provenance_MAP"
## 
## Call:
## lm(formula = form, data = dat)
## 
## Residuals:
##       Min        1Q    Median        3Q       Max 
## -0.032069 -0.013036  0.001253  0.014332  0.047907 
## 
## Coefficients:
##                  Estimate Std. Error t value Pr(>|t|)    
## (Intercept)     9.065e-02  8.124e-03  11.158 3.85e-14 ***
## provenance_MAP -8.688e-06  1.300e-05  -0.668    0.508    
## ---
## Signif. codes:  0 '***' 0.001 '**' 0.01 '*' 0.05 '.' 0.1 ' ' 1
## 
## Residual standard error: 0.01903 on 42 degrees of freedom
## Multiple R-squared:  0.01052,    Adjusted R-squared:  -0.01304 
## F-statistic: 0.4465 on 1 and 42 DF,  p-value: 0.5076
## 
## 
## Call:
## lm(formula = form, data = dat)
## 
## Standardized Coefficients::
##    (Intercept) provenance_MAP 
##             NA     -0.1025671 
## 
## [1] "Model: plasticity_rdpi_DOY_Stage6_2023 ~ provenance_TD"
## 
## Call:
## lm(formula = form, data = dat)
## 
## Residuals:
##       Min        1Q    Median        3Q       Max 
## -0.028617 -0.016481  0.000714  0.010601  0.044402 
## 
## Coefficients:
##                Estimate Std. Error t value Pr(>|t|)  
## (Intercept)   0.0501488  0.0193155   2.596   0.0129 *
## provenance_TD 0.0013795  0.0007445   1.853   0.0709 .
## ---
## Signif. codes:  0 '***' 0.001 '**' 0.01 '*' 0.05 '.' 0.1 ' ' 1
## 
## Residual standard error: 0.01839 on 42 degrees of freedom
## Multiple R-squared:  0.07556,    Adjusted R-squared:  0.05355 
## F-statistic: 3.433 on 1 and 42 DF,  p-value: 0.07094
## 
## 
## Call:
## lm(formula = form, data = dat)
## 
## Standardized Coefficients::
##   (Intercept) provenance_TD 
##            NA     0.2748821 
## 
## [1] "Model: plasticity_rdpi_DOY_Stage6_2024 ~ genetic_PC1"
## 
## Call:
## lm(formula = form, data = dat)
## 
## Residuals:
##       Min        1Q    Median        3Q       Max 
## -0.043690 -0.016016 -0.005762  0.004787  0.094603 
## 
## Coefficients:
##              Estimate Std. Error t value Pr(>|t|)    
## (Intercept)  0.061832   0.004269  14.483   <2e-16 ***
## genetic_PC1 -0.013046   0.119264  -0.109    0.913    
## ---
## Signif. codes:  0 '***' 0.001 '**' 0.01 '*' 0.05 '.' 0.1 ' ' 1
## 
## Residual standard error: 0.02768 on 41 degrees of freedom
##   (1 observation deleted due to missingness)
## Multiple R-squared:  0.0002918,  Adjusted R-squared:  -0.02409 
## F-statistic: 0.01197 on 1 and 41 DF,  p-value: 0.9134
## 
## 
## Call:
## lm(formula = form, data = dat)
## 
## Standardized Coefficients::
## (Intercept) genetic_PC1 
##          NA -0.01708103 
## 
## [1] "Model: plasticity_rdpi_DOY_Stage6_2024 ~ genetic_PC2"
## 
## Call:
## lm(formula = form, data = dat)
## 
## Residuals:
##       Min        1Q    Median        3Q       Max 
## -0.043785 -0.015962 -0.006446  0.005278  0.095812 
## 
## Coefficients:
##             Estimate Std. Error t value Pr(>|t|)    
## (Intercept)  0.06173    0.00438  14.096   <2e-16 ***
## genetic_PC2  0.01659    0.11663   0.142    0.888    
## ---
## Signif. codes:  0 '***' 0.001 '**' 0.01 '*' 0.05 '.' 0.1 ' ' 1
## 
## Residual standard error: 0.02768 on 41 degrees of freedom
##   (1 observation deleted due to missingness)
## Multiple R-squared:  0.0004932,  Adjusted R-squared:  -0.02388 
## F-statistic: 0.02023 on 1 and 41 DF,  p-value: 0.8876
## 
## 
## Call:
## lm(formula = form, data = dat)
## 
## Standardized Coefficients::
## (Intercept) genetic_PC2 
##          NA  0.02220888 
## 
## [1] "Model: plasticity_rdpi_DOY_Stage6_2024 ~ genetic_PC3"
## 
## Call:
## lm(formula = form, data = dat)
## 
## Residuals:
##       Min        1Q    Median        3Q       Max 
## -0.037389 -0.016805 -0.008562  0.010144  0.096317 
## 
## Coefficients:
##             Estimate Std. Error t value Pr(>|t|)    
## (Intercept) 0.062384   0.004134  15.090   <2e-16 ***
## genetic_PC3 0.152921   0.106820   1.432     0.16    
## ---
## Signif. codes:  0 '***' 0.001 '**' 0.01 '*' 0.05 '.' 0.1 ' ' 1
## 
## Residual standard error: 0.02702 on 41 degrees of freedom
##   (1 observation deleted due to missingness)
## Multiple R-squared:  0.04761,    Adjusted R-squared:  0.02438 
## F-statistic: 2.049 on 1 and 41 DF,  p-value: 0.1598
## 
## 
## Call:
## lm(formula = form, data = dat)
## 
## Standardized Coefficients::
## (Intercept) genetic_PC3 
##          NA   0.2181879 
## 
## [1] "Model: plasticity_rdpi_DOY_Stage6_2024 ~ provenance_MAT"
## 
## Call:
## lm(formula = form, data = dat)
## 
## Residuals:
##       Min        1Q    Median        3Q       Max 
## -0.041292 -0.016099 -0.009104  0.008588  0.094924 
## 
## Coefficients:
##                 Estimate Std. Error t value Pr(>|t|)    
## (Intercept)     0.065717   0.005884  11.169 5.18e-14 ***
## provenance_MAT -0.001494   0.001622  -0.921    0.362    
## ---
## Signif. codes:  0 '***' 0.001 '**' 0.01 '*' 0.05 '.' 0.1 ' ' 1
## 
## Residual standard error: 0.0274 on 41 degrees of freedom
##   (1 observation deleted due to missingness)
## Multiple R-squared:  0.02028,    Adjusted R-squared:  -0.003616 
## F-statistic: 0.8487 on 1 and 41 DF,  p-value: 0.3623
## 
## 
## Call:
## lm(formula = form, data = dat)
## 
## Standardized Coefficients::
##    (Intercept) provenance_MAT 
##             NA     -0.1424051 
## 
## [1] "Model: plasticity_rdpi_DOY_Stage6_2024 ~ provenance_MAP"
## 
## Call:
## lm(formula = form, data = dat)
## 
## Residuals:
##       Min        1Q    Median        3Q       Max 
## -0.043701 -0.015371 -0.008040  0.002638  0.095081 
## 
## Coefficients:
##                 Estimate Std. Error t value Pr(>|t|)    
## (Intercept)    4.862e-02  1.172e-02   4.150 0.000164 ***
## provenance_MAP 2.260e-05  1.865e-05   1.212 0.232583    
## ---
## Signif. codes:  0 '***' 0.001 '**' 0.01 '*' 0.05 '.' 0.1 ' ' 1
## 
## Residual standard error: 0.0272 on 41 degrees of freedom
##   (1 observation deleted due to missingness)
## Multiple R-squared:  0.03457,    Adjusted R-squared:  0.01102 
## F-statistic: 1.468 on 1 and 41 DF,  p-value: 0.2326
## 
## 
## Call:
## lm(formula = form, data = dat)
## 
## Standardized Coefficients::
##    (Intercept) provenance_MAP 
##             NA      0.1859285 
## 
## [1] "Model: plasticity_rdpi_DOY_Stage6_2024 ~ provenance_TD"
## 
## Call:
## lm(formula = form, data = dat)
## 
## Residuals:
##       Min        1Q    Median        3Q       Max 
## -0.044183 -0.015994 -0.006762  0.006949  0.092973 
## 
## Coefficients:
##                 Estimate Std. Error t value Pr(>|t|)   
## (Intercept)    0.0864828  0.0289136   2.991  0.00469 **
## provenance_TD -0.0009596  0.0011168  -0.859  0.39522   
## ---
## Signif. codes:  0 '***' 0.001 '**' 0.01 '*' 0.05 '.' 0.1 ' ' 1
## 
## Residual standard error: 0.02744 on 41 degrees of freedom
##   (1 observation deleted due to missingness)
## Multiple R-squared:  0.01769,    Adjusted R-squared:  -0.006272 
## F-statistic: 0.7382 on 1 and 41 DF,  p-value: 0.3952
## 
## 
## Call:
## lm(formula = form, data = dat)
## 
## Standardized Coefficients::
##   (Intercept) provenance_TD 
##            NA    -0.1329937 
## 
## [1] "Model: plasticity_rdpi_DOY_Stage7_2023 ~ genetic_PC1"
## 
## Call:
## lm(formula = form, data = dat)
## 
## Residuals:
##       Min        1Q    Median        3Q       Max 
## -0.060511 -0.018392 -0.005088  0.011563  0.116866 
## 
## Coefficients:
##             Estimate Std. Error t value Pr(>|t|)    
## (Intercept) 0.077661   0.006088  12.756 2.63e-14 ***
## genetic_PC1 0.235698   0.168209   1.401     0.17    
## ---
## Signif. codes:  0 '***' 0.001 '**' 0.01 '*' 0.05 '.' 0.1 ' ' 1
## 
## Residual standard error: 0.03397 on 33 degrees of freedom
##   (9 observations deleted due to missingness)
## Multiple R-squared:  0.05616,    Adjusted R-squared:  0.02756 
## F-statistic: 1.963 on 1 and 33 DF,  p-value: 0.1705
## 
## 
## Call:
## lm(formula = form, data = dat)
## 
## Standardized Coefficients::
## (Intercept) genetic_PC1 
##          NA   0.2369738 
## 
## [1] "Model: plasticity_rdpi_DOY_Stage7_2023 ~ genetic_PC2"
## 
## Call:
## lm(formula = form, data = dat)
## 
## Residuals:
##       Min        1Q    Median        3Q       Max 
## -0.045943 -0.017740 -0.007601  0.013029  0.121955 
## 
## Coefficients:
##             Estimate Std. Error t value Pr(>|t|)    
## (Intercept) 0.074261   0.005994  12.388 5.88e-14 ***
## genetic_PC2 0.075655   0.151588   0.499    0.621    
## ---
## Signif. codes:  0 '***' 0.001 '**' 0.01 '*' 0.05 '.' 0.1 ' ' 1
## 
## Residual standard error: 0.03483 on 33 degrees of freedom
##   (9 observations deleted due to missingness)
## Multiple R-squared:  0.007491,   Adjusted R-squared:  -0.02258 
## F-statistic: 0.2491 on 1 and 33 DF,  p-value: 0.621
## 
## 
## Call:
## lm(formula = form, data = dat)
## 
## Standardized Coefficients::
## (Intercept) genetic_PC2 
##          NA  0.08655277 
## 
## [1] "Model: plasticity_rdpi_DOY_Stage7_2023 ~ genetic_PC3"
## 
## Call:
## lm(formula = form, data = dat)
## 
## Residuals:
##       Min        1Q    Median        3Q       Max 
## -0.058460 -0.017708 -0.002776  0.013539  0.126716 
## 
## Coefficients:
##             Estimate Std. Error t value Pr(>|t|)    
## (Intercept) 0.076402   0.005847  13.066 1.35e-14 ***
## genetic_PC3 0.211311   0.149786   1.411    0.168    
## ---
## Signif. codes:  0 '***' 0.001 '**' 0.01 '*' 0.05 '.' 0.1 ' ' 1
## 
## Residual standard error: 0.03395 on 33 degrees of freedom
##   (9 observations deleted due to missingness)
## Multiple R-squared:  0.05688,    Adjusted R-squared:  0.0283 
## F-statistic:  1.99 on 1 and 33 DF,  p-value: 0.1677
## 
## 
## Call:
## lm(formula = form, data = dat)
## 
## Standardized Coefficients::
## (Intercept) genetic_PC3 
##          NA   0.2384941 
## 
## [1] "Model: plasticity_rdpi_DOY_Stage7_2023 ~ provenance_MAT"
## 
## Call:
## lm(formula = form, data = dat)
## 
## Residuals:
##       Min        1Q    Median        3Q       Max 
## -0.050830 -0.015282 -0.009727  0.013981  0.118226 
## 
## Coefficients:
##                 Estimate Std. Error t value Pr(>|t|)    
## (Intercept)     0.088788   0.009392   9.454 6.48e-11 ***
## provenance_MAT -0.004451   0.002398  -1.856   0.0723 .  
## ---
## Signif. codes:  0 '***' 0.001 '**' 0.01 '*' 0.05 '.' 0.1 ' ' 1
## 
## Residual standard error: 0.03327 on 33 degrees of freedom
##   (9 observations deleted due to missingness)
## Multiple R-squared:  0.09456,    Adjusted R-squared:  0.06712 
## F-statistic: 3.446 on 1 and 33 DF,  p-value: 0.07234
## 
## 
## Call:
## lm(formula = form, data = dat)
## 
## Standardized Coefficients::
##    (Intercept) provenance_MAT 
##             NA     -0.3075082 
## 
## [1] "Model: plasticity_rdpi_DOY_Stage7_2023 ~ provenance_MAP"
## 
## Call:
## lm(formula = form, data = dat)
## 
## Residuals:
##       Min        1Q    Median        3Q       Max 
## -0.045644 -0.020841 -0.002778  0.012882  0.121673 
## 
## Coefficients:
##                 Estimate Std. Error t value Pr(>|t|)   
## (Intercept)    5.504e-02  1.708e-02   3.223  0.00286 **
## provenance_MAP 3.435e-05  2.791e-05   1.231  0.22720   
## ---
## Signif. codes:  0 '***' 0.001 '**' 0.01 '*' 0.05 '.' 0.1 ' ' 1
## 
## Residual standard error: 0.03419 on 33 degrees of freedom
##   (9 observations deleted due to missingness)
## Multiple R-squared:  0.04387,    Adjusted R-squared:  0.0149 
## F-statistic: 1.514 on 1 and 33 DF,  p-value: 0.2272
## 
## 
## Call:
## lm(formula = form, data = dat)
## 
## Standardized Coefficients::
##    (Intercept) provenance_MAP 
##             NA      0.2094559 
## 
## [1] "Model: plasticity_rdpi_DOY_Stage7_2023 ~ provenance_TD"
## 
## Call:
## lm(formula = form, data = dat)
## 
## Residuals:
##       Min        1Q    Median        3Q       Max 
## -0.047058 -0.017751 -0.007278  0.009716  0.120703 
## 
## Coefficients:
##                Estimate Std. Error t value Pr(>|t|)
## (Intercept)   0.0579676  0.0556623   1.041    0.305
## provenance_TD 0.0006781  0.0022267   0.305    0.763
## 
## Residual standard error: 0.03491 on 33 degrees of freedom
##   (9 observations deleted due to missingness)
## Multiple R-squared:  0.002803,   Adjusted R-squared:  -0.02742 
## F-statistic: 0.09274 on 1 and 33 DF,  p-value: 0.7626
## 
## 
## Call:
## lm(formula = form, data = dat)
## 
## Standardized Coefficients::
##   (Intercept) provenance_TD 
##            NA    0.05293868 
## 
## [1] "Model: plasticity_rdpi_DOY_Stage7_2024 ~ genetic_PC1"
## 
## Call:
## lm(formula = form, data = dat)
## 
## Residuals:
##       Min        1Q    Median        3Q       Max 
## -0.038784 -0.015692 -0.002531  0.013868  0.076448 
## 
## Coefficients:
##             Estimate Std. Error t value Pr(>|t|)   
## (Intercept)  0.05533    0.01716   3.224  0.00531 **
## genetic_PC1 -0.06641    0.47768  -0.139  0.89116   
## ---
## Signif. codes:  0 '***' 0.001 '**' 0.01 '*' 0.05 '.' 0.1 ' ' 1
## 
## Residual standard error: 0.02697 on 16 degrees of freedom
##   (26 observations deleted due to missingness)
## Multiple R-squared:  0.001207,   Adjusted R-squared:  -0.06122 
## F-statistic: 0.01933 on 1 and 16 DF,  p-value: 0.8912
## 
## 
## Call:
## lm(formula = form, data = dat)
## 
## Standardized Coefficients::
## (Intercept) genetic_PC1 
##          NA -0.03473746 
## 
## [1] "Model: plasticity_rdpi_DOY_Stage7_2024 ~ genetic_PC2"
## 
## Call:
## lm(formula = form, data = dat)
## 
## Residuals:
##       Min        1Q    Median        3Q       Max 
## -0.039088 -0.017648 -0.002527  0.016607  0.074479 
## 
## Coefficients:
##              Estimate Std. Error t value Pr(>|t|)    
## (Intercept)  0.057326   0.006362   9.011 1.15e-07 ***
## genetic_PC2 -0.061308   0.163044  -0.376    0.712    
## ---
## Signif. codes:  0 '***' 0.001 '**' 0.01 '*' 0.05 '.' 0.1 ' ' 1
## 
## Residual standard error: 0.02687 on 16 degrees of freedom
##   (26 observations deleted due to missingness)
## Multiple R-squared:  0.00876,    Adjusted R-squared:  -0.05319 
## F-statistic: 0.1414 on 1 and 16 DF,  p-value: 0.7118
## 
## 
## Call:
## lm(formula = form, data = dat)
## 
## Standardized Coefficients::
## (Intercept) genetic_PC2 
##          NA -0.09359225 
## 
## [1] "Model: plasticity_rdpi_DOY_Stage7_2024 ~ genetic_PC3"
## 
## Call:
## lm(formula = form, data = dat)
## 
## Residuals:
##       Min        1Q    Median        3Q       Max 
## -0.034331 -0.016022 -0.004288  0.010456  0.073130 
## 
## Coefficients:
##             Estimate Std. Error t value Pr(>|t|)    
## (Intercept) 0.064861   0.007566   8.573 2.23e-07 ***
## genetic_PC3 0.324664   0.208703   1.556    0.139    
## ---
## Signif. codes:  0 '***' 0.001 '**' 0.01 '*' 0.05 '.' 0.1 ' ' 1
## 
## Residual standard error: 0.02516 on 16 degrees of freedom
##   (26 observations deleted due to missingness)
## Multiple R-squared:  0.1314, Adjusted R-squared:  0.07709 
## F-statistic:  2.42 on 1 and 16 DF,  p-value: 0.1394
## 
## 
## Call:
## lm(formula = form, data = dat)
## 
## Standardized Coefficients::
## (Intercept) genetic_PC3 
##          NA   0.3624602 
## 
## [1] "Model: plasticity_rdpi_DOY_Stage7_2024 ~ provenance_MAT"
## 
## Call:
## lm(formula = form, data = dat)
## 
## Residuals:
##       Min        1Q    Median        3Q       Max 
## -0.042248 -0.012703 -0.000636  0.005403  0.065347 
## 
## Coefficients:
##                 Estimate Std. Error t value Pr(>|t|)    
## (Intercept)     0.084030   0.014641   5.739 3.05e-05 ***
## provenance_MAT -0.006080   0.003095  -1.964   0.0671 .  
## ---
## Signif. codes:  0 '***' 0.001 '**' 0.01 '*' 0.05 '.' 0.1 ' ' 1
## 
## Residual standard error: 0.02423 on 16 degrees of freedom
##   (26 observations deleted due to missingness)
## Multiple R-squared:  0.1943, Adjusted R-squared:  0.1439 
## F-statistic: 3.858 on 1 and 16 DF,  p-value: 0.06714
## 
## 
## Call:
## lm(formula = form, data = dat)
## 
## Standardized Coefficients::
##    (Intercept) provenance_MAT 
##             NA     -0.4407564 
## 
## [1] "Model: plasticity_rdpi_DOY_Stage7_2024 ~ provenance_MAP"
## 
## Call:
## lm(formula = form, data = dat)
## 
## Residuals:
##       Min        1Q    Median        3Q       Max 
## -0.028843 -0.012549 -0.004341  0.005012  0.075902 
## 
## Coefficients:
##                 Estimate Std. Error t value Pr(>|t|)  
## (Intercept)    3.449e-02  1.910e-02   1.806   0.0898 .
## provenance_MAP 4.073e-05  3.199e-05   1.273   0.2211  
## ---
## Signif. codes:  0 '***' 0.001 '**' 0.01 '*' 0.05 '.' 0.1 ' ' 1
## 
## Residual standard error: 0.02572 on 16 degrees of freedom
##   (26 observations deleted due to missingness)
## Multiple R-squared:  0.09202,    Adjusted R-squared:  0.03527 
## F-statistic: 1.621 on 1 and 16 DF,  p-value: 0.2211
## 
## 
## Call:
## lm(formula = form, data = dat)
## 
## Standardized Coefficients::
##    (Intercept) provenance_MAP 
##             NA      0.3033432 
## 
## [1] "Model: plasticity_rdpi_DOY_Stage7_2024 ~ provenance_TD"
## 
## Call:
## lm(formula = form, data = dat)
## 
## Residuals:
##      Min       1Q   Median       3Q      Max 
## -0.02877 -0.01153 -0.00513  0.00626  0.07471 
## 
## Coefficients:
##                Estimate Std. Error t value Pr(>|t|)  
## (Intercept)    0.215319   0.095334   2.259   0.0382 *
## provenance_TD -0.006632   0.004000  -1.658   0.1168  
## ---
## Signif. codes:  0 '***' 0.001 '**' 0.01 '*' 0.05 '.' 0.1 ' ' 1
## 
## Residual standard error: 0.02493 on 16 degrees of freedom
##   (26 observations deleted due to missingness)
## Multiple R-squared:  0.1466, Adjusted R-squared:  0.09329 
## F-statistic: 2.749 on 1 and 16 DF,  p-value: 0.1168
## 
## 
## Call:
## lm(formula = form, data = dat)
## 
## Standardized Coefficients::
##   (Intercept) provenance_TD 
##            NA    -0.3829203 
## 
## [1] "Model: plasticity_rdpi_stage7_presence_2023 ~ genetic_PC1"
## 
## Call:
## lm(formula = form, data = dat)
## 
## Residuals:
##      Min       1Q   Median       3Q      Max 
## -0.60472  0.02906  0.03617  0.04902  0.06349 
## 
## Coefficients:
##             Estimate Std. Error t value Pr(>|t|)    
## (Intercept)  0.95818    0.02323  41.256   <2e-16 ***
## genetic_PC1 -0.33645    0.65547  -0.513     0.61    
## ---
## Signif. codes:  0 '***' 0.001 '**' 0.01 '*' 0.05 '.' 0.1 ' ' 1
## 
## Residual standard error: 0.1525 on 42 degrees of freedom
## Multiple R-squared:  0.006234,   Adjusted R-squared:  -0.01743 
## F-statistic: 0.2635 on 1 and 42 DF,  p-value: 0.6104
## 
## 
## Call:
## lm(formula = form, data = dat)
## 
## Standardized Coefficients::
## (Intercept) genetic_PC1 
##          NA -0.07895618 
## 
## [1] "Model: plasticity_rdpi_stage7_presence_2023 ~ genetic_PC2"
## 
## Call:
## lm(formula = form, data = dat)
## 
## Residuals:
##      Min       1Q   Median       3Q      Max 
## -0.60872  0.02699  0.03449  0.04940  0.06844 
## 
## Coefficients:
##             Estimate Std. Error t value Pr(>|t|)    
## (Intercept)  0.95464    0.02391  39.930   <2e-16 ***
## genetic_PC2  0.48339    0.63430   0.762     0.45    
## ---
## Signif. codes:  0 '***' 0.001 '**' 0.01 '*' 0.05 '.' 0.1 ' ' 1
## 
## Residual standard error: 0.152 on 42 degrees of freedom
## Multiple R-squared:  0.01364,    Adjusted R-squared:  -0.009845 
## F-statistic: 0.5808 on 1 and 42 DF,  p-value: 0.4503
## 
## 
## Call:
## lm(formula = form, data = dat)
## 
## Standardized Coefficients::
## (Intercept) genetic_PC2 
##          NA    0.116789 
## 
## [1] "Model: plasticity_rdpi_stage7_presence_2023 ~ genetic_PC3"
## 
## Call:
## lm(formula = form, data = dat)
## 
## Residuals:
##      Min       1Q   Median       3Q      Max 
## -0.63209  0.02830  0.03234  0.05109  0.06666 
## 
## Coefficients:
##             Estimate Std. Error t value Pr(>|t|)    
## (Intercept)  0.95916    0.02301  41.683   <2e-16 ***
## genetic_PC3 -0.34069    0.59175  -0.576    0.568    
## ---
## Signif. codes:  0 '***' 0.001 '**' 0.01 '*' 0.05 '.' 0.1 ' ' 1
## 
## Residual standard error: 0.1524 on 42 degrees of freedom
## Multiple R-squared:  0.00783,    Adjusted R-squared:  -0.01579 
## F-statistic: 0.3315 on 1 and 42 DF,  p-value: 0.5679
## 
## 
## Call:
## lm(formula = form, data = dat)
## 
## Standardized Coefficients::
## (Intercept) genetic_PC3 
##          NA -0.08848948 
## 
## [1] "Model: plasticity_rdpi_stage7_presence_2023 ~ provenance_MAT"
## 
## Call:
## lm(formula = form, data = dat)
## 
## Residuals:
##      Min       1Q   Median       3Q      Max 
## -0.62578  0.03615  0.03991  0.04215  0.04707 
## 
## Coefficients:
##                 Estimate Std. Error t value Pr(>|t|)    
## (Intercept)     0.963326   0.031849  30.246   <2e-16 ***
## provenance_MAT -0.001405   0.008875  -0.158    0.875    
## ---
## Signif. codes:  0 '***' 0.001 '**' 0.01 '*' 0.05 '.' 0.1 ' ' 1
## 
## Residual standard error: 0.153 on 42 degrees of freedom
## Multiple R-squared:  0.0005965,  Adjusted R-squared:  -0.0232 
## F-statistic: 0.02507 on 1 and 42 DF,  p-value: 0.875
## 
## 
## Call:
## lm(formula = form, data = dat)
## 
## Standardized Coefficients::
##    (Intercept) provenance_MAT 
##             NA    -0.02442302 
## 
## [1] "Model: plasticity_rdpi_stage7_presence_2023 ~ provenance_MAP"
## 
## Call:
## lm(formula = form, data = dat)
## 
## Residuals:
##      Min       1Q   Median       3Q      Max 
## -0.62943  0.03686  0.03911  0.04076  0.05288 
## 
## Coefficients:
##                  Estimate Std. Error t value Pr(>|t|)    
## (Intercept)     9.706e-01  6.531e-02  14.861   <2e-16 ***
## provenance_MAP -1.836e-05  1.045e-04  -0.176    0.861    
## ---
## Signif. codes:  0 '***' 0.001 '**' 0.01 '*' 0.05 '.' 0.1 ' ' 1
## 
## Residual standard error: 0.153 on 42 degrees of freedom
## Multiple R-squared:  0.000734,   Adjusted R-squared:  -0.02306 
## F-statistic: 0.03085 on 1 and 42 DF,  p-value: 0.8614
## 
## 
## Call:
## lm(formula = form, data = dat)
## 
## Standardized Coefficients::
##    (Intercept) provenance_MAP 
##             NA    -0.02709311 
## 
## [1] "Model: plasticity_rdpi_stage7_presence_2023 ~ provenance_TD"
## 
## Call:
## lm(formula = form, data = dat)
## 
## Residuals:
##      Min       1Q   Median       3Q      Max 
## -0.62822  0.03182  0.04206  0.04898  0.06333 
## 
## Coefficients:
##               Estimate Std. Error t value Pr(>|t|)    
## (Intercept)   0.856797   0.159911   5.358 3.31e-06 ***
## provenance_TD 0.004014   0.006164   0.651    0.518    
## ---
## Signif. codes:  0 '***' 0.001 '**' 0.01 '*' 0.05 '.' 0.1 ' ' 1
## 
## Residual standard error: 0.1523 on 42 degrees of freedom
## Multiple R-squared:  0.009995,   Adjusted R-squared:  -0.01358 
## F-statistic: 0.424 on 1 and 42 DF,  p-value: 0.5185
## 
## 
## Call:
## lm(formula = form, data = dat)
## 
## Standardized Coefficients::
##   (Intercept) provenance_TD 
##            NA    0.09997512 
## 
## [1] "Model: plasticity_rdpi_stage7_presence_2024 ~ genetic_PC1"
## 
## Call:
## lm(formula = form, data = dat)
## 
## Residuals:
##      Min       1Q   Median       3Q      Max 
## -0.45316 -0.16032 -0.00627  0.11681  0.42237 
## 
## Coefficients:
##             Estimate Std. Error t value Pr(>|t|)    
## (Intercept)  0.78506    0.03475  22.590  < 2e-16 ***
## genetic_PC1  4.34501    0.97080   4.476 5.95e-05 ***
## ---
## Signif. codes:  0 '***' 0.001 '**' 0.01 '*' 0.05 '.' 0.1 ' ' 1
## 
## Residual standard error: 0.2253 on 41 degrees of freedom
##   (1 observation deleted due to missingness)
## Multiple R-squared:  0.3282, Adjusted R-squared:  0.3118 
## F-statistic: 20.03 on 1 and 41 DF,  p-value: 5.946e-05
## 
## 
## Call:
## lm(formula = form, data = dat)
## 
## Standardized Coefficients::
## (Intercept) genetic_PC1 
##          NA   0.5729064 
## 
## [1] "Model: plasticity_rdpi_stage7_presence_2024 ~ genetic_PC2"
## 
## Call:
## lm(formula = form, data = dat)
## 
## Residuals:
##     Min      1Q  Median      3Q     Max 
## -0.5540 -0.2384  0.1000  0.2131  0.3597 
## 
## Coefficients:
##             Estimate Std. Error t value Pr(>|t|)    
## (Intercept)  0.74078    0.04171  17.759   <2e-16 ***
## genetic_PC2  2.10443    1.11076   1.895   0.0652 .  
## ---
## Signif. codes:  0 '***' 0.001 '**' 0.01 '*' 0.05 '.' 0.1 ' ' 1
## 
## Residual standard error: 0.2636 on 41 degrees of freedom
##   (1 observation deleted due to missingness)
## Multiple R-squared:  0.0805, Adjusted R-squared:  0.05807 
## F-statistic: 3.589 on 1 and 41 DF,  p-value: 0.06522
## 
## 
## Call:
## lm(formula = form, data = dat)
## 
## Standardized Coefficients::
## (Intercept) genetic_PC2 
##          NA   0.2837246 
## 
## [1] "Model: plasticity_rdpi_stage7_presence_2024 ~ genetic_PC3"
## 
## Call:
## lm(formula = form, data = dat)
## 
## Residuals:
##     Min      1Q  Median      3Q     Max 
## -0.5521 -0.1821  0.0951  0.2291  0.3196 
## 
## Coefficients:
##             Estimate Std. Error t value Pr(>|t|)    
## (Intercept)  0.76788    0.04045  18.985   <2e-16 ***
## genetic_PC3  1.91185    1.04507   1.829   0.0746 .  
## ---
## Signif. codes:  0 '***' 0.001 '**' 0.01 '*' 0.05 '.' 0.1 ' ' 1
## 
## Residual standard error: 0.2643 on 41 degrees of freedom
##   (1 observation deleted due to missingness)
## Multiple R-squared:  0.07547,    Adjusted R-squared:  0.05292 
## F-statistic: 3.347 on 1 and 41 DF,  p-value: 0.07462
## 
## 
## Call:
## lm(formula = form, data = dat)
## 
## Standardized Coefficients::
## (Intercept) genetic_PC3 
##          NA   0.2747115 
## 
## [1] "Model: plasticity_rdpi_stage7_presence_2024 ~ provenance_MAT"
## 
## Call:
## lm(formula = form, data = dat)
## 
## Residuals:
##      Min       1Q   Median       3Q      Max 
## -0.41209 -0.21086 -0.01176  0.21059  0.44323 
## 
## Coefficients:
##                Estimate Std. Error t value Pr(>|t|)    
## (Intercept)     0.90162    0.05022  17.952  < 2e-16 ***
## provenance_MAT -0.05474    0.01385  -3.954 0.000297 ***
## ---
## Signif. codes:  0 '***' 0.001 '**' 0.01 '*' 0.05 '.' 0.1 ' ' 1
## 
## Residual standard error: 0.2339 on 41 degrees of freedom
##   (1 observation deleted due to missingness)
## Multiple R-squared:  0.276,  Adjusted R-squared:  0.2584 
## F-statistic: 15.63 on 1 and 41 DF,  p-value: 0.0002975
## 
## 
## Call:
## lm(formula = form, data = dat)
## 
## Standardized Coefficients::
##    (Intercept) provenance_MAT 
##             NA      -0.525369 
## 
## [1] "Model: plasticity_rdpi_stage7_presence_2024 ~ provenance_MAP"
## 
## Call:
## lm(formula = form, data = dat)
## 
## Residuals:
##     Min      1Q  Median      3Q     Max 
## -0.6353 -0.1710  0.1238  0.2400  0.2719 
## 
## Coefficients:
##                Estimate Std. Error t value Pr(>|t|)    
## (Intercept)    0.671975   0.117455   5.721 1.08e-06 ***
## provenance_MAP 0.000153   0.000187   0.818    0.418    
## ---
## Signif. codes:  0 '***' 0.001 '**' 0.01 '*' 0.05 '.' 0.1 ' ' 1
## 
## Residual standard error: 0.2727 on 41 degrees of freedom
##   (1 observation deleted due to missingness)
## Multiple R-squared:  0.01606,    Adjusted R-squared:  -0.007934 
## F-statistic: 0.6694 on 1 and 41 DF,  p-value: 0.418
## 
## 
## Call:
## lm(formula = form, data = dat)
## 
## Standardized Coefficients::
##    (Intercept) provenance_MAP 
##             NA      0.1267468 
## 
## [1] "Model: plasticity_rdpi_stage7_presence_2024 ~ provenance_TD"
## 
## Call:
## lm(formula = form, data = dat)
## 
## Residuals:
##      Min       1Q   Median       3Q      Max 
## -0.55635 -0.17817  0.03504  0.21580  0.37501 
## 
## Coefficients:
##               Estimate Std. Error t value Pr(>|t|)  
## (Intercept)    0.14856    0.27303   0.544   0.5893  
## provenance_TD  0.02394    0.01055   2.270   0.0285 *
## ---
## Signif. codes:  0 '***' 0.001 '**' 0.01 '*' 0.05 '.' 0.1 ' ' 1
## 
## Residual standard error: 0.2591 on 41 degrees of freedom
##   (1 observation deleted due to missingness)
## Multiple R-squared:  0.1117, Adjusted R-squared:  0.08999 
## F-statistic: 5.154 on 1 and 41 DF,  p-value: 0.02852
## 
## 
## Call:
## lm(formula = form, data = dat)
## 
## Standardized Coefficients::
##   (Intercept) provenance_TD 
##            NA     0.3341572 
## 
## [1] "Model: plasticity_rdpi_DOY_last_budset_2023 ~ genetic_PC1"
## 
## Call:
## lm(formula = form, data = dat)
## 
## Residuals:
##       Min        1Q    Median        3Q       Max 
## -0.042238 -0.030050 -0.007304  0.010449  0.095019 
## 
## Coefficients:
##             Estimate Std. Error t value Pr(>|t|)    
## (Intercept) 0.098519   0.005563  17.709   <2e-16 ***
## genetic_PC1 0.004925   0.157005   0.031    0.975    
## ---
## Signif. codes:  0 '***' 0.001 '**' 0.01 '*' 0.05 '.' 0.1 ' ' 1
## 
## Residual standard error: 0.03654 on 42 degrees of freedom
## Multiple R-squared:  2.342e-05,  Adjusted R-squared:  -0.02379 
## F-statistic: 0.0009838 on 1 and 42 DF,  p-value: 0.9751
## 
## 
## Call:
## lm(formula = form, data = dat)
## 
## Standardized Coefficients::
## (Intercept) genetic_PC1 
##          NA 0.004839809 
## 
## [1] "Model: plasticity_rdpi_DOY_last_budset_2023 ~ genetic_PC2"
## 
## Call:
## lm(formula = form, data = dat)
## 
## Residuals:
##       Min        1Q    Median        3Q       Max 
## -0.041422 -0.029957 -0.007521  0.011884  0.097860 
## 
## Coefficients:
##              Estimate Std. Error t value Pr(>|t|)    
## (Intercept)  0.099828   0.005703  17.505   <2e-16 ***
## genetic_PC2 -0.123843   0.151302  -0.819    0.418    
## ---
## Signif. codes:  0 '***' 0.001 '**' 0.01 '*' 0.05 '.' 0.1 ' ' 1
## 
## Residual standard error: 0.03625 on 42 degrees of freedom
## Multiple R-squared:  0.0157, Adjusted R-squared:  -0.007735 
## F-statistic:  0.67 on 1 and 42 DF,  p-value: 0.4177
## 
## 
## Call:
## lm(formula = form, data = dat)
## 
## Standardized Coefficients::
## (Intercept) genetic_PC2 
##          NA  -0.1253041 
## 
## [1] "Model: plasticity_rdpi_DOY_last_budset_2023 ~ genetic_PC3"
## 
## Call:
## lm(formula = form, data = dat)
## 
## Residuals:
##      Min       1Q   Median       3Q      Max 
## -0.04556 -0.02588 -0.01004  0.01022  0.08829 
## 
## Coefficients:
##             Estimate Std. Error t value Pr(>|t|)    
## (Intercept) 0.098788   0.005448  18.133   <2e-16 ***
## genetic_PC3 0.144216   0.140100   1.029    0.309    
## ---
## Signif. codes:  0 '***' 0.001 '**' 0.01 '*' 0.05 '.' 0.1 ' ' 1
## 
## Residual standard error: 0.03609 on 42 degrees of freedom
## Multiple R-squared:  0.02461,    Adjusted R-squared:  0.001385 
## F-statistic:  1.06 on 1 and 42 DF,  p-value: 0.3092
## 
## 
## Call:
## lm(formula = form, data = dat)
## 
## Standardized Coefficients::
## (Intercept) genetic_PC3 
##          NA   0.1568705 
## 
## [1] "Model: plasticity_rdpi_DOY_last_budset_2023 ~ provenance_MAT"
## 
## Call:
## lm(formula = form, data = dat)
## 
## Residuals:
##      Min       1Q   Median       3Q      Max 
## -0.03730 -0.02379 -0.01335  0.01617  0.08171 
## 
## Coefficients:
##                 Estimate Std. Error t value Pr(>|t|)    
## (Intercept)     0.108343   0.007281  14.880   <2e-16 ***
## provenance_MAT -0.003979   0.002029  -1.961   0.0565 .  
## ---
## Signif. codes:  0 '***' 0.001 '**' 0.01 '*' 0.05 '.' 0.1 ' ' 1
## 
## Residual standard error: 0.03497 on 42 degrees of freedom
## Multiple R-squared:  0.08389,    Adjusted R-squared:  0.06207 
## F-statistic: 3.846 on 1 and 42 DF,  p-value: 0.05652
## 
## 
## Call:
## lm(formula = form, data = dat)
## 
## Standardized Coefficients::
##    (Intercept) provenance_MAT 
##             NA      -0.289631 
## 
## [1] "Model: plasticity_rdpi_DOY_last_budset_2023 ~ provenance_MAP"
## 
## Call:
## lm(formula = form, data = dat)
## 
## Residuals:
##       Min        1Q    Median        3Q       Max 
## -0.042293 -0.029725 -0.007172  0.010594  0.095132 
## 
## Coefficients:
##                 Estimate Std. Error t value Pr(>|t|)    
## (Intercept)    9.686e-02  1.560e-02   6.210 1.98e-07 ***
## provenance_MAP 2.793e-06  2.496e-05   0.112    0.911    
## ---
## Signif. codes:  0 '***' 0.001 '**' 0.01 '*' 0.05 '.' 0.1 ' ' 1
## 
## Residual standard error: 0.03653 on 42 degrees of freedom
## Multiple R-squared:  0.000298,   Adjusted R-squared:  -0.0235 
## F-statistic: 0.01252 on 1 and 42 DF,  p-value: 0.9114
## 
## 
## Call:
## lm(formula = form, data = dat)
## 
## Standardized Coefficients::
##    (Intercept) provenance_MAP 
##             NA     0.01726383 
## 
## [1] "Model: plasticity_rdpi_DOY_last_budset_2023 ~ provenance_TD"
## 
## Call:
## lm(formula = form, data = dat)
## 
## Residuals:
##      Min       1Q   Median       3Q      Max 
## -0.04358 -0.02766 -0.01050  0.01638  0.09699 
## 
## Coefficients:
##               Estimate Std. Error t value Pr(>|t|)
## (Intercept)   0.054100   0.037747   1.433    0.159
## provenance_TD 0.001729   0.001455   1.188    0.241
## 
## Residual standard error: 0.03594 on 42 degrees of freedom
## Multiple R-squared:  0.03253,    Adjusted R-squared:  0.009499 
## F-statistic: 1.412 on 1 and 42 DF,  p-value: 0.2413
## 
## 
## Call:
## lm(formula = form, data = dat)
## 
## Standardized Coefficients::
##   (Intercept) provenance_TD 
##            NA     0.1803705 
## 
## [1] "Model: plasticity_rdpi_DOY_last_budset_2024 ~ genetic_PC1"
## 
## Call:
## lm(formula = form, data = dat)
## 
## Residuals:
##       Min        1Q    Median        3Q       Max 
## -0.038806 -0.014752 -0.004971  0.011667  0.082321 
## 
## Coefficients:
##             Estimate Std. Error t value Pr(>|t|)    
## (Intercept) 0.083702   0.003971   21.08   <2e-16 ***
## genetic_PC1 0.207439   0.110919    1.87   0.0686 .  
## ---
## Signif. codes:  0 '***' 0.001 '**' 0.01 '*' 0.05 '.' 0.1 ' ' 1
## 
## Residual standard error: 0.02575 on 41 degrees of freedom
##   (1 observation deleted due to missingness)
## Multiple R-squared:  0.0786, Adjusted R-squared:  0.05613 
## F-statistic: 3.498 on 1 and 41 DF,  p-value: 0.06861
## 
## 
## Call:
## lm(formula = form, data = dat)
## 
## Standardized Coefficients::
## (Intercept) genetic_PC1 
##          NA   0.2803598 
## 
## [1] "Model: plasticity_rdpi_DOY_last_budset_2024 ~ genetic_PC2"
## 
## Call:
## lm(formula = form, data = dat)
## 
## Residuals:
##       Min        1Q    Median        3Q       Max 
## -0.038129 -0.019466 -0.002694  0.012724  0.065627 
## 
## Coefficients:
##             Estimate Std. Error t value Pr(>|t|)    
## (Intercept) 0.080495   0.004062  19.817   <2e-16 ***
## genetic_PC2 0.209586   0.108165   1.938   0.0596 .  
## ---
## Signif. codes:  0 '***' 0.001 '**' 0.01 '*' 0.05 '.' 0.1 ' ' 1
## 
## Residual standard error: 0.02567 on 41 degrees of freedom
##   (1 observation deleted due to missingness)
## Multiple R-squared:  0.08389,    Adjusted R-squared:  0.06155 
## F-statistic: 3.754 on 1 and 41 DF,  p-value: 0.05958
## 
## 
## Call:
## lm(formula = form, data = dat)
## 
## Standardized Coefficients::
## (Intercept) genetic_PC2 
##          NA   0.2896385 
## 
## [1] "Model: plasticity_rdpi_DOY_last_budset_2024 ~ genetic_PC3"
## 
## Call:
## lm(formula = form, data = dat)
## 
## Residuals:
##       Min        1Q    Median        3Q       Max 
## -0.044916 -0.014989 -0.005697  0.013212  0.077203 
## 
## Coefficients:
##             Estimate Std. Error t value Pr(>|t|)    
## (Intercept) 0.082866   0.004071   20.36   <2e-16 ***
## genetic_PC3 0.086255   0.105177    0.82    0.417    
## ---
## Signif. codes:  0 '***' 0.001 '**' 0.01 '*' 0.05 '.' 0.1 ' ' 1
## 
## Residual standard error: 0.0266 on 41 degrees of freedom
##   (1 observation deleted due to missingness)
## Multiple R-squared:  0.01614,    Adjusted R-squared:  -0.007858 
## F-statistic: 0.6726 on 1 and 41 DF,  p-value: 0.4169
## 
## 
## Call:
## lm(formula = form, data = dat)
## 
## Standardized Coefficients::
## (Intercept) genetic_PC3 
##          NA   0.1270393 
## 
## [1] "Model: plasticity_rdpi_DOY_last_budset_2024 ~ provenance_MAT"
## 
## Call:
## lm(formula = form, data = dat)
## 
## Residuals:
##       Min        1Q    Median        3Q       Max 
## -0.037086 -0.018511 -0.004894  0.010497  0.074490 
## 
## Coefficients:
##                 Estimate Std. Error t value Pr(>|t|)    
## (Intercept)     0.089695   0.005539  16.194   <2e-16 ***
## provenance_MAT -0.002781   0.001527  -1.821   0.0758 .  
## ---
## Signif. codes:  0 '***' 0.001 '**' 0.01 '*' 0.05 '.' 0.1 ' ' 1
## 
## Residual standard error: 0.0258 on 41 degrees of freedom
##   (1 observation deleted due to missingness)
## Multiple R-squared:  0.07486,    Adjusted R-squared:  0.0523 
## F-statistic: 3.318 on 1 and 41 DF,  p-value: 0.07584
## 
## 
## Call:
## lm(formula = form, data = dat)
## 
## Standardized Coefficients::
##    (Intercept) provenance_MAT 
##             NA     -0.2736125 
## 
## [1] "Model: plasticity_rdpi_DOY_last_budset_2024 ~ provenance_MAP"
## 
## Call:
## lm(formula = form, data = dat)
## 
## Residuals:
##       Min        1Q    Median        3Q       Max 
## -0.047042 -0.015314 -0.004318  0.010988  0.064679 
## 
## Coefficients:
##                 Estimate Std. Error t value Pr(>|t|)    
## (Intercept)    6.968e-02  1.135e-02   6.140 2.73e-07 ***
## provenance_MAP 2.198e-05  1.807e-05   1.216    0.231    
## ---
## Signif. codes:  0 '***' 0.001 '**' 0.01 '*' 0.05 '.' 0.1 ' ' 1
## 
## Residual standard error: 0.02635 on 41 degrees of freedom
##   (1 observation deleted due to missingness)
## Multiple R-squared:  0.03482,    Adjusted R-squared:  0.01128 
## F-statistic: 1.479 on 1 and 41 DF,  p-value: 0.2309
## 
## 
## Call:
## lm(formula = form, data = dat)
## 
## Standardized Coefficients::
##    (Intercept) provenance_MAP 
##             NA       0.186607 
## 
## [1] "Model: plasticity_rdpi_DOY_last_budset_2024 ~ provenance_TD"
## 
## Call:
## lm(formula = form, data = dat)
## 
## Residuals:
##       Min        1Q    Median        3Q       Max 
## -0.043585 -0.016896 -0.006085  0.012012  0.080679 
## 
## Coefficients:
##               Estimate Std. Error t value Pr(>|t|)  
## (Intercept)   0.050540   0.027805   1.818   0.0764 .
## provenance_TD 0.001251   0.001074   1.165   0.2507  
## ---
## Signif. codes:  0 '***' 0.001 '**' 0.01 '*' 0.05 '.' 0.1 ' ' 1
## 
## Residual standard error: 0.02639 on 41 degrees of freedom
##   (1 observation deleted due to missingness)
## Multiple R-squared:  0.03205,    Adjusted R-squared:  0.008438 
## F-statistic: 1.357 on 1 and 41 DF,  p-value: 0.2507
## 
## 
## Call:
## lm(formula = form, data = dat)
## 
## Standardized Coefficients::
##   (Intercept) provenance_TD 
##            NA     0.1790169 
## 
## [1] "Model: plasticity_rdpi_leaf_thickness_avg_mm_2023 ~ genetic_PC1"
## 
## Call:
## lm(formula = form, data = dat)
## 
## Residuals:
##      Min       1Q   Median       3Q      Max 
## -0.05122 -0.02871 -0.01608  0.01790  0.20652 
## 
## Coefficients:
##              Estimate Std. Error t value Pr(>|t|)    
## (Intercept)  0.093929   0.007227  12.997  2.6e-16 ***
## genetic_PC1 -0.294101   0.203953  -1.442    0.157    
## ---
## Signif. codes:  0 '***' 0.001 '**' 0.01 '*' 0.05 '.' 0.1 ' ' 1
## 
## Residual standard error: 0.04747 on 42 degrees of freedom
## Multiple R-squared:  0.04717,    Adjusted R-squared:  0.02449 
## F-statistic: 2.079 on 1 and 42 DF,  p-value: 0.1567
## 
## 
## Call:
## lm(formula = form, data = dat)
## 
## Standardized Coefficients::
## (Intercept) genetic_PC1 
##          NA  -0.2171949 
## 
## [1] "Model: plasticity_rdpi_leaf_thickness_avg_mm_2023 ~ genetic_PC2"
## 
## Call:
## lm(formula = form, data = dat)
## 
## Residuals:
##      Min       1Q   Median       3Q      Max 
## -0.06351 -0.02870 -0.01252  0.01446  0.20929 
## 
## Coefficients:
##              Estimate Std. Error t value Pr(>|t|)    
## (Intercept)  0.098305   0.007485  13.133   <2e-16 ***
## genetic_PC2 -0.271198   0.198586  -1.366    0.179    
## ---
## Signif. codes:  0 '***' 0.001 '**' 0.01 '*' 0.05 '.' 0.1 ' ' 1
## 
## Residual standard error: 0.04758 on 42 degrees of freedom
## Multiple R-squared:  0.04252,    Adjusted R-squared:  0.01972 
## F-statistic: 1.865 on 1 and 42 DF,  p-value: 0.1793
## 
## 
## Call:
## lm(formula = form, data = dat)
## 
## Standardized Coefficients::
## (Intercept) genetic_PC2 
##          NA  -0.2061953 
## 
## [1] "Model: plasticity_rdpi_leaf_thickness_avg_mm_2023 ~ genetic_PC3"
## 
## Call:
## lm(formula = form, data = dat)
## 
## Residuals:
##      Min       1Q   Median       3Q      Max 
## -0.06291 -0.02761 -0.01775  0.01413  0.21232 
## 
## Coefficients:
##              Estimate Std. Error t value Pr(>|t|)    
## (Intercept)  0.095073   0.007283  13.054 2.24e-16 ***
## genetic_PC3 -0.153453   0.187287  -0.819    0.417    
## ---
## Signif. codes:  0 '***' 0.001 '**' 0.01 '*' 0.05 '.' 0.1 ' ' 1
## 
## Residual standard error: 0.04824 on 42 degrees of freedom
## Multiple R-squared:  0.01573,    Adjusted R-squared:  -0.007702 
## F-statistic: 0.6713 on 1 and 42 DF,  p-value: 0.4172
## 
## 
## Call:
## lm(formula = form, data = dat)
## 
## Standardized Coefficients::
## (Intercept) genetic_PC3 
##          NA  -0.1254293 
## 
## [1] "Model: plasticity_rdpi_leaf_thickness_avg_mm_2023 ~ provenance_MAT"
## 
## Call:
## lm(formula = form, data = dat)
## 
## Residuals:
##      Min       1Q   Median       3Q      Max 
## -0.05762 -0.03003 -0.01499  0.01604  0.19754 
## 
## Coefficients:
##                Estimate Std. Error t value Pr(>|t|)    
## (Intercept)    0.084513   0.009827   8.600 8.27e-11 ***
## provenance_MAT 0.004393   0.002738   1.604    0.116    
## ---
## Signif. codes:  0 '***' 0.001 '**' 0.01 '*' 0.05 '.' 0.1 ' ' 1
## 
## Residual standard error: 0.0472 on 42 degrees of freedom
## Multiple R-squared:  0.05773,    Adjusted R-squared:  0.0353 
## F-statistic: 2.573 on 1 and 42 DF,  p-value: 0.1162
## 
## 
## Call:
## lm(formula = form, data = dat)
## 
## Standardized Coefficients::
##    (Intercept) provenance_MAT 
##             NA      0.2402774 
## 
## [1] "Model: plasticity_rdpi_leaf_thickness_avg_mm_2023 ~ provenance_MAP"
## 
## Call:
## lm(formula = form, data = dat)
## 
## Residuals:
##      Min       1Q   Median       3Q      Max 
## -0.05937 -0.02802 -0.01664  0.01114  0.21897 
## 
## Coefficients:
##                  Estimate Std. Error t value Pr(>|t|)    
## (Intercept)     1.013e-01  2.074e-02   4.884 1.55e-05 ***
## provenance_MAP -1.011e-05  3.319e-05  -0.305    0.762    
## ---
## Signif. codes:  0 '***' 0.001 '**' 0.01 '*' 0.05 '.' 0.1 ' ' 1
## 
## Residual standard error: 0.04857 on 42 degrees of freedom
## Multiple R-squared:  0.002203,   Adjusted R-squared:  -0.02155 
## F-statistic: 0.09274 on 1 and 42 DF,  p-value: 0.7622
## 
## 
## Call:
## lm(formula = form, data = dat)
## 
## Standardized Coefficients::
##    (Intercept) provenance_MAP 
##             NA     -0.0469394 
## 
## [1] "Model: plasticity_rdpi_leaf_thickness_avg_mm_2023 ~ provenance_TD"
## 
## Call:
## lm(formula = form, data = dat)
## 
## Residuals:
##      Min       1Q   Median       3Q      Max 
## -0.05625 -0.03007 -0.01736  0.01780  0.21427 
## 
## Coefficients:
##                Estimate Std. Error t value Pr(>|t|)  
## (Intercept)    0.129865   0.050787   2.557   0.0143 *
## provenance_TD -0.001343   0.001958  -0.686   0.4965  
## ---
## Signif. codes:  0 '***' 0.001 '**' 0.01 '*' 0.05 '.' 0.1 ' ' 1
## 
## Residual standard error: 0.04836 on 42 degrees of freedom
## Multiple R-squared:  0.01108,    Adjusted R-squared:  -0.01246 
## F-statistic: 0.4706 on 1 and 42 DF,  p-value: 0.4965
## 
## 
## Call:
## lm(formula = form, data = dat)
## 
## Standardized Coefficients::
##   (Intercept) provenance_TD 
##            NA    -0.1052675 
## 
## [1] "Model: plasticity_rdpi_leaf_area_cm2_2023_log ~ genetic_PC1"
## 
## Call:
## lm(formula = form, data = dat)
## 
## Residuals:
##      Min       1Q   Median       3Q      Max 
## -0.11278 -0.04254 -0.02372  0.01918  0.40844 
## 
## Coefficients:
##             Estimate Std. Error t value Pr(>|t|)    
## (Intercept)   0.1646     0.0143  11.517  1.4e-14 ***
## genetic_PC1   0.2077     0.4035   0.515    0.609    
## ---
## Signif. codes:  0 '***' 0.001 '**' 0.01 '*' 0.05 '.' 0.1 ' ' 1
## 
## Residual standard error: 0.0939 on 42 degrees of freedom
## Multiple R-squared:  0.006269,   Adjusted R-squared:  -0.01739 
## F-statistic: 0.2649 on 1 and 42 DF,  p-value: 0.6094
## 
## 
## Call:
## lm(formula = form, data = dat)
## 
## Standardized Coefficients::
## (Intercept) genetic_PC1 
##          NA  0.07917443 
## 
## [1] "Model: plasticity_rdpi_leaf_area_cm2_2023_log ~ genetic_PC2"
## 
## Call:
## lm(formula = form, data = dat)
## 
## Residuals:
##      Min       1Q   Median       3Q      Max 
## -0.10558 -0.03858 -0.01782  0.02599  0.35821 
## 
## Coefficients:
##             Estimate Std. Error t value Pr(>|t|)    
## (Intercept)   0.1558     0.0142  10.972 6.56e-14 ***
## genetic_PC2   0.7295     0.3767   1.937   0.0595 .  
## ---
## Signif. codes:  0 '***' 0.001 '**' 0.01 '*' 0.05 '.' 0.1 ' ' 1
## 
## Residual standard error: 0.09025 on 42 degrees of freedom
## Multiple R-squared:  0.08199,    Adjusted R-squared:  0.06013 
## F-statistic: 3.751 on 1 and 42 DF,  p-value: 0.05952
## 
## 
## Call:
## lm(formula = form, data = dat)
## 
## Standardized Coefficients::
## (Intercept) genetic_PC2 
##          NA   0.2863332 
## 
## [1] "Model: plasticity_rdpi_leaf_area_cm2_2023_log ~ genetic_PC3"
## 
## Call:
## lm(formula = form, data = dat)
## 
## Residuals:
##      Min       1Q   Median       3Q      Max 
## -0.13062 -0.05002 -0.01075  0.02103  0.32695 
## 
## Coefficients:
##             Estimate Std. Error t value Pr(>|t|)    
## (Intercept)  0.16566    0.01289  12.847 3.85e-16 ***
## genetic_PC3  0.99904    0.33160   3.013  0.00437 ** 
## ---
## Signif. codes:  0 '***' 0.001 '**' 0.01 '*' 0.05 '.' 0.1 ' ' 1
## 
## Residual standard error: 0.08542 on 42 degrees of freedom
## Multiple R-squared:  0.1777, Adjusted R-squared:  0.1581 
## F-statistic: 9.077 on 1 and 42 DF,  p-value: 0.004373
## 
## 
## Call:
## lm(formula = form, data = dat)
## 
## Standardized Coefficients::
## (Intercept) genetic_PC3 
##          NA   0.4215522 
## 
## [1] "Model: plasticity_rdpi_leaf_area_cm2_2023_log ~ provenance_MAT"
## 
## Call:
## lm(formula = form, data = dat)
## 
## Residuals:
##      Min       1Q   Median       3Q      Max 
## -0.10529 -0.04102 -0.02070  0.01627  0.38113 
## 
## Coefficients:
##                 Estimate Std. Error t value Pr(>|t|)    
## (Intercept)     0.200611   0.017779  11.283  2.7e-14 ***
## provenance_MAT -0.014944   0.004954  -3.016  0.00433 ** 
## ---
## Signif. codes:  0 '***' 0.001 '**' 0.01 '*' 0.05 '.' 0.1 ' ' 1
## 
## Residual standard error: 0.0854 on 42 degrees of freedom
## Multiple R-squared:  0.1781, Adjusted R-squared:  0.1585 
## F-statistic: 9.098 on 1 and 42 DF,  p-value: 0.004331
## 
## 
## Call:
## lm(formula = form, data = dat)
## 
## Standardized Coefficients::
##    (Intercept) provenance_MAT 
##             NA     -0.4219645 
## 
## [1] "Model: plasticity_rdpi_leaf_area_cm2_2023_log ~ provenance_MAP"
## 
## Call:
## lm(formula = form, data = dat)
## 
## Residuals:
##      Min       1Q   Median       3Q      Max 
## -0.10113 -0.04090 -0.02662  0.02510  0.40412 
## 
## Coefficients:
##                 Estimate Std. Error t value Pr(>|t|)    
## (Intercept)    1.604e-01  4.021e-02   3.989  0.00026 ***
## provenance_MAP 5.497e-06  6.436e-05   0.085  0.93234    
## ---
## Signif. codes:  0 '***' 0.001 '**' 0.01 '*' 0.05 '.' 0.1 ' ' 1
## 
## Residual standard error: 0.09419 on 42 degrees of freedom
## Multiple R-squared:  0.0001737,  Adjusted R-squared:  -0.02363 
## F-statistic: 0.007295 on 1 and 42 DF,  p-value: 0.9323
## 
## 
## Call:
## lm(formula = form, data = dat)
## 
## Standardized Coefficients::
##    (Intercept) provenance_MAP 
##             NA     0.01317772 
## 
## [1] "Model: plasticity_rdpi_leaf_area_cm2_2023_log ~ provenance_TD"
## 
## Call:
## lm(formula = form, data = dat)
## 
## Residuals:
##      Min       1Q   Median       3Q      Max 
## -0.11120 -0.04286 -0.01746  0.01929  0.41101 
## 
## Coefficients:
##                Estimate Std. Error t value Pr(>|t|)  
## (Intercept)   -0.046418   0.093351  -0.497   0.6216  
## provenance_TD  0.008181   0.003598   2.274   0.0282 *
## ---
## Signif. codes:  0 '***' 0.001 '**' 0.01 '*' 0.05 '.' 0.1 ' ' 1
## 
## Residual standard error: 0.08888 on 42 degrees of freedom
## Multiple R-squared:  0.1096, Adjusted R-squared:  0.08839 
## F-statistic: 5.169 on 1 and 42 DF,  p-value: 0.02817
## 
## 
## Call:
## lm(formula = form, data = dat)
## 
## Standardized Coefficients::
##   (Intercept) provenance_TD 
##            NA     0.3310394 
## 
## [1] "Model: plasticity_rdpi_leaf_mass_g_2023_log ~ genetic_PC1"
## 
## Call:
## lm(formula = form, data = dat)
## 
## Residuals:
##      Min       1Q   Median       3Q      Max 
## -0.50212 -0.03473  0.05745  0.09235  0.20644 
## 
## Coefficients:
##             Estimate Std. Error t value Pr(>|t|)    
## (Intercept) -0.30247    0.02438 -12.409 1.23e-15 ***
## genetic_PC1  1.38008    0.68793   2.006   0.0513 .  
## ---
## Signif. codes:  0 '***' 0.001 '**' 0.01 '*' 0.05 '.' 0.1 ' ' 1
## 
## Residual standard error: 0.1601 on 42 degrees of freedom
## Multiple R-squared:  0.08744,    Adjusted R-squared:  0.06572 
## F-statistic: 4.025 on 1 and 42 DF,  p-value: 0.05131
## 
## 
## Call:
## lm(formula = form, data = dat)
## 
## Standardized Coefficients::
## (Intercept) genetic_PC1 
##          NA    0.295709 
## 
## [1] "Model: plasticity_rdpi_leaf_mass_g_2023_log ~ genetic_PC2"
## 
## Call:
## lm(formula = form, data = dat)
## 
## Residuals:
##      Min       1Q   Median       3Q      Max 
## -0.51725 -0.04911  0.02488  0.12091  0.18896 
## 
## Coefficients:
##             Estimate Std. Error t value Pr(>|t|)    
## (Intercept) -0.29650    0.02544 -11.654 9.59e-15 ***
## genetic_PC2 -1.18916    0.67498  -1.762   0.0854 .  
## ---
## Signif. codes:  0 '***' 0.001 '**' 0.01 '*' 0.05 '.' 0.1 ' ' 1
## 
## Residual standard error: 0.1617 on 42 degrees of freedom
## Multiple R-squared:  0.06881,    Adjusted R-squared:  0.04664 
## F-statistic: 3.104 on 1 and 42 DF,  p-value: 0.08539
## 
## 
## Call:
## lm(formula = form, data = dat)
## 
## Standardized Coefficients::
## (Intercept) genetic_PC2 
##          NA  -0.2623254 
## 
## [1] "Model: plasticity_rdpi_leaf_mass_g_2023_log ~ genetic_PC3"
## 
## Call:
## lm(formula = form, data = dat)
## 
## Residuals:
##      Min       1Q   Median       3Q      Max 
## -0.56775 -0.01065  0.03065  0.09898  0.20788 
## 
## Coefficients:
##             Estimate Std. Error t value Pr(>|t|)    
## (Intercept) -0.30961    0.02528 -12.245 1.91e-15 ***
## genetic_PC3 -0.15050    0.65023  -0.231    0.818    
## ---
## Signif. codes:  0 '***' 0.001 '**' 0.01 '*' 0.05 '.' 0.1 ' ' 1
## 
## Residual standard error: 0.1675 on 42 degrees of freedom
## Multiple R-squared:  0.001274,   Adjusted R-squared:  -0.02251 
## F-statistic: 0.05357 on 1 and 42 DF,  p-value: 0.8181
## 
## 
## Call:
## lm(formula = form, data = dat)
## 
## Standardized Coefficients::
## (Intercept) genetic_PC3 
##          NA -0.03569234 
## 
## [1] "Model: plasticity_rdpi_leaf_mass_g_2023_log ~ provenance_MAT"
## 
## Call:
## lm(formula = form, data = dat)
## 
## Residuals:
##      Min       1Q   Median       3Q      Max 
## -0.56238 -0.01109  0.03310  0.10278  0.22142 
## 
## Coefficients:
##                 Estimate Std. Error t value Pr(>|t|)    
## (Intercept)    -0.316332   0.034857  -9.075 1.87e-11 ***
## provenance_MAT  0.002840   0.009713   0.292    0.771    
## ---
## Signif. codes:  0 '***' 0.001 '**' 0.01 '*' 0.05 '.' 0.1 ' ' 1
## 
## Residual standard error: 0.1674 on 42 degrees of freedom
## Multiple R-squared:  0.002031,   Adjusted R-squared:  -0.02173 
## F-statistic: 0.08547 on 1 and 42 DF,  p-value: 0.7715
## 
## 
## Call:
## lm(formula = form, data = dat)
## 
## Standardized Coefficients::
##    (Intercept) provenance_MAT 
##             NA     0.04506403 
## 
## [1] "Model: plasticity_rdpi_leaf_mass_g_2023_log ~ provenance_MAP"
## 
## Call:
## lm(formula = form, data = dat)
## 
## Residuals:
##      Min       1Q   Median       3Q      Max 
## -0.45570 -0.03875  0.03916  0.08171  0.27197 
## 
## Coefficients:
##                  Estimate Std. Error t value Pr(>|t|)  
## (Intercept)    -0.1572238  0.0670150  -2.346   0.0238 *
## provenance_MAP -0.0002601  0.0001073  -2.426   0.0197 *
## ---
## Signif. codes:  0 '***' 0.001 '**' 0.01 '*' 0.05 '.' 0.1 ' ' 1
## 
## Residual standard error: 0.157 on 42 degrees of freedom
## Multiple R-squared:  0.1229, Adjusted R-squared:  0.102 
## F-statistic: 5.883 on 1 and 42 DF,  p-value: 0.01966
## 
## 
## Call:
## lm(formula = form, data = dat)
## 
## Standardized Coefficients::
##    (Intercept) provenance_MAP 
##             NA      -0.350529 
## 
## [1] "Model: plasticity_rdpi_leaf_mass_g_2023_log ~ provenance_TD"
## 
## Call:
## lm(formula = form, data = dat)
## 
## Residuals:
##      Min       1Q   Median       3Q      Max 
## -0.50294 -0.02849  0.04758  0.09591  0.21927 
## 
## Coefficients:
##                Estimate Std. Error t value Pr(>|t|)   
## (Intercept)   -0.568181   0.171329  -3.316  0.00189 **
## provenance_TD  0.010083   0.006604   1.527  0.13431   
## ---
## Signif. codes:  0 '***' 0.001 '**' 0.01 '*' 0.05 '.' 0.1 ' ' 1
## 
## Residual standard error: 0.1631 on 42 degrees of freedom
## Multiple R-squared:  0.05258,    Adjusted R-squared:  0.03003 
## F-statistic: 2.331 on 1 and 42 DF,  p-value: 0.1343
## 
## 
## Call:
## lm(formula = form, data = dat)
## 
## Standardized Coefficients::
##   (Intercept) provenance_TD 
##            NA     0.2293133 
## 
## [1] "Model: plasticity_rdpi_LMA_g_m2_2023 ~ genetic_PC1"
## 
## Call:
## lm(formula = form, data = dat)
## 
## Residuals:
##       Min        1Q    Median        3Q       Max 
## -0.059622 -0.024392 -0.006093  0.023132  0.095806 
## 
## Coefficients:
##             Estimate Std. Error t value Pr(>|t|)    
## (Intercept)  0.11855    0.00543  21.833   <2e-16 ***
## genetic_PC1  0.39733    0.15325   2.593    0.013 *  
## ---
## Signif. codes:  0 '***' 0.001 '**' 0.01 '*' 0.05 '.' 0.1 ' ' 1
## 
## Residual standard error: 0.03567 on 42 degrees of freedom
## Multiple R-squared:  0.138,  Adjusted R-squared:  0.1174 
## F-statistic: 6.722 on 1 and 42 DF,  p-value: 0.01305
## 
## 
## Call:
## lm(formula = form, data = dat)
## 
## Standardized Coefficients::
## (Intercept) genetic_PC1 
##          NA   0.3714367 
## 
## [1] "Model: plasticity_rdpi_LMA_g_m2_2023 ~ genetic_PC2"
## 
## Call:
## lm(formula = form, data = dat)
## 
## Residuals:
##       Min        1Q    Median        3Q       Max 
## -0.067920 -0.030065 -0.006432  0.017777  0.121915 
## 
## Coefficients:
##              Estimate Std. Error t value Pr(>|t|)    
## (Intercept)  0.117147   0.006035  19.410   <2e-16 ***
## genetic_PC2 -0.051949   0.160124  -0.324    0.747    
## ---
## Signif. codes:  0 '***' 0.001 '**' 0.01 '*' 0.05 '.' 0.1 ' ' 1
## 
## Residual standard error: 0.03837 on 42 degrees of freedom
## Multiple R-squared:  0.0025, Adjusted R-squared:  -0.02125 
## F-statistic: 0.1053 on 1 and 42 DF,  p-value: 0.7472
## 
## 
## Call:
## lm(formula = form, data = dat)
## 
## Standardized Coefficients::
## (Intercept) genetic_PC2 
##          NA -0.04999781 
## 
## [1] "Model: plasticity_rdpi_LMA_g_m2_2023 ~ genetic_PC3"
## 
## Call:
## lm(formula = form, data = dat)
## 
## Residuals:
##       Min        1Q    Median        3Q       Max 
## -0.063235 -0.031299 -0.007204  0.014170  0.125629 
## 
## Coefficients:
##             Estimate Std. Error t value Pr(>|t|)    
## (Intercept) 0.116750   0.005779  20.202   <2e-16 ***
## genetic_PC3 0.080118   0.148618   0.539    0.593    
## ---
## Signif. codes:  0 '***' 0.001 '**' 0.01 '*' 0.05 '.' 0.1 ' ' 1
## 
## Residual standard error: 0.03828 on 42 degrees of freedom
## Multiple R-squared:  0.006872,   Adjusted R-squared:  -0.01677 
## F-statistic: 0.2906 on 1 and 42 DF,  p-value: 0.5927
## 
## 
## Call:
## lm(formula = form, data = dat)
## 
## Standardized Coefficients::
## (Intercept) genetic_PC3 
##          NA  0.08289663 
## 
## [1] "Model: plasticity_rdpi_LMA_g_m2_2023 ~ provenance_MAT"
## 
## Call:
## lm(formula = form, data = dat)
## 
## Residuals:
##       Min        1Q    Median        3Q       Max 
## -0.054963 -0.029729 -0.004664  0.020482  0.122993 
## 
## Coefficients:
##                 Estimate Std. Error t value Pr(>|t|)    
## (Intercept)     0.121929   0.007908   15.42   <2e-16 ***
## provenance_MAT -0.002158   0.002204   -0.98    0.333    
## ---
## Signif. codes:  0 '***' 0.001 '**' 0.01 '*' 0.05 '.' 0.1 ' ' 1
## 
## Residual standard error: 0.03798 on 42 degrees of freedom
## Multiple R-squared:  0.02233,    Adjusted R-squared:  -0.0009446 
## F-statistic: 0.9594 on 1 and 42 DF,  p-value: 0.3329
## 
## 
## Call:
## lm(formula = form, data = dat)
## 
## Standardized Coefficients::
##    (Intercept) provenance_MAT 
##             NA     -0.1494429 
## 
## [1] "Model: plasticity_rdpi_LMA_g_m2_2023 ~ provenance_MAP"
## 
## Call:
## lm(formula = form, data = dat)
## 
## Residuals:
##       Min        1Q    Median        3Q       Max 
## -0.065698 -0.029516 -0.005919  0.020528  0.123193 
## 
## Coefficients:
##                  Estimate Std. Error t value Pr(>|t|)    
## (Intercept)     1.219e-01  1.638e-02   7.443 3.41e-09 ***
## provenance_MAP -9.072e-06  2.621e-05  -0.346    0.731    
## ---
## Signif. codes:  0 '***' 0.001 '**' 0.01 '*' 0.05 '.' 0.1 ' ' 1
## 
## Residual standard error: 0.03836 on 42 degrees of freedom
## Multiple R-squared:  0.002844,   Adjusted R-squared:  -0.0209 
## F-statistic: 0.1198 on 1 and 42 DF,  p-value: 0.731
## 
## 
## Call:
## lm(formula = form, data = dat)
## 
## Standardized Coefficients::
##    (Intercept) provenance_MAP 
##             NA    -0.05333132 
## 
## [1] "Model: plasticity_rdpi_LMA_g_m2_2023 ~ provenance_TD"
## 
## Call:
## lm(formula = form, data = dat)
## 
## Residuals:
##       Min        1Q    Median        3Q       Max 
## -0.055452 -0.025448 -0.009403  0.013001  0.117023 
## 
## Coefficients:
##               Estimate Std. Error t value Pr(>|t|)  
## (Intercept)   0.031914   0.038123   0.837   0.4073  
## provenance_TD 0.003298   0.001469   2.244   0.0301 *
## ---
## Signif. codes:  0 '***' 0.001 '**' 0.01 '*' 0.05 '.' 0.1 ' ' 1
## 
## Residual standard error: 0.0363 on 42 degrees of freedom
## Multiple R-squared:  0.1071, Adjusted R-squared:  0.08582 
## F-statistic: 5.037 on 1 and 42 DF,  p-value: 0.03014
## 
## 
## Call:
## lm(formula = form, data = dat)
## 
## Standardized Coefficients::
##   (Intercept) provenance_TD 
##            NA     0.3272347 
## 
## [1] "Model: plasticity_rdpi_lower_stomata_pore_length_mean_um ~ genetic_PC1"
## 
## Call:
## lm(formula = form, data = dat)
## 
## Residuals:
##      Min       1Q   Median       3Q      Max 
## -0.05555 -0.03120 -0.01864  0.01621  0.14171 
## 
## Coefficients:
##             Estimate Std. Error t value Pr(>|t|)    
## (Intercept) 0.095877   0.007206  13.306   <2e-16 ***
## genetic_PC1 0.052589   0.203361   0.259    0.797    
## ---
## Signif. codes:  0 '***' 0.001 '**' 0.01 '*' 0.05 '.' 0.1 ' ' 1
## 
## Residual standard error: 0.04733 on 42 degrees of freedom
## Multiple R-squared:  0.00159,    Adjusted R-squared:  -0.02218 
## F-statistic: 0.06687 on 1 and 42 DF,  p-value: 0.7972
## 
## 
## Call:
## lm(formula = form, data = dat)
## 
## Standardized Coefficients::
## (Intercept) genetic_PC1 
##          NA  0.03987095 
## 
## [1] "Model: plasticity_rdpi_lower_stomata_pore_length_mean_um ~ genetic_PC2"
## 
## Call:
## lm(formula = form, data = dat)
## 
## Residuals:
##      Min       1Q   Median       3Q      Max 
## -0.05804 -0.03218 -0.01920  0.01413  0.14098 
## 
## Coefficients:
##             Estimate Std. Error t value Pr(>|t|)    
## (Intercept)  0.09588    0.00745  12.871 3.62e-16 ***
## genetic_PC2 -0.02469    0.19765  -0.125    0.901    
## ---
## Signif. codes:  0 '***' 0.001 '**' 0.01 '*' 0.05 '.' 0.1 ' ' 1
## 
## Residual standard error: 0.04736 on 42 degrees of freedom
## Multiple R-squared:  0.0003715,  Adjusted R-squared:  -0.02343 
## F-statistic: 0.01561 on 1 and 42 DF,  p-value: 0.9012
## 
## 
## Call:
## lm(formula = form, data = dat)
## 
## Standardized Coefficients::
## (Intercept) genetic_PC2 
##          NA -0.01927333 
## 
## [1] "Model: plasticity_rdpi_lower_stomata_pore_length_mean_um ~ genetic_PC3"
## 
## Call:
## lm(formula = form, data = dat)
## 
## Residuals:
##      Min       1Q   Median       3Q      Max 
## -0.06953 -0.02855 -0.01054  0.01431  0.12644 
## 
## Coefficients:
##              Estimate Std. Error t value Pr(>|t|)    
## (Intercept)  0.094949   0.006874  13.813   <2e-16 ***
## genetic_PC3 -0.328180   0.176773  -1.856   0.0704 .  
## ---
## Signif. codes:  0 '***' 0.001 '**' 0.01 '*' 0.05 '.' 0.1 ' ' 1
## 
## Residual standard error: 0.04553 on 42 degrees of freedom
## Multiple R-squared:  0.07584,    Adjusted R-squared:  0.05383 
## F-statistic: 3.447 on 1 and 42 DF,  p-value: 0.07041
## 
## 
## Call:
## lm(formula = form, data = dat)
## 
## Standardized Coefficients::
## (Intercept) genetic_PC3 
##          NA  -0.2753874 
## 
## [1] "Model: plasticity_rdpi_lower_stomata_pore_length_mean_um ~ provenance_MAT"
## 
## Call:
## lm(formula = form, data = dat)
## 
## Residuals:
##      Min       1Q   Median       3Q      Max 
## -0.05879 -0.03226 -0.01966  0.01491  0.14102 
## 
## Coefficients:
##                 Estimate Std. Error t value Pr(>|t|)    
## (Intercept)    0.0949615  0.0098601   9.631 3.41e-12 ***
## provenance_MAT 0.0002647  0.0027476   0.096    0.924    
## ---
## Signif. codes:  0 '***' 0.001 '**' 0.01 '*' 0.05 '.' 0.1 ' ' 1
## 
## Residual standard error: 0.04736 on 42 degrees of freedom
## Multiple R-squared:  0.0002209,  Adjusted R-squared:  -0.02358 
## F-statistic: 0.009281 on 1 and 42 DF,  p-value: 0.9237
## 
## 
## Call:
## lm(formula = form, data = dat)
## 
## Standardized Coefficients::
##    (Intercept) provenance_MAT 
##             NA     0.01486352 
## 
## [1] "Model: plasticity_rdpi_lower_stomata_pore_length_mean_um ~ provenance_MAP"
## 
## Call:
## lm(formula = form, data = dat)
## 
## Residuals:
##      Min       1Q   Median       3Q      Max 
## -0.06626 -0.03392 -0.01595  0.02518  0.13200 
## 
## Coefficients:
##                  Estimate Std. Error t value Pr(>|t|)    
## (Intercept)     1.185e-01  1.987e-02   5.965 4.46e-07 ***
## provenance_MAP -3.916e-05  3.180e-05  -1.232    0.225    
## ---
## Signif. codes:  0 '***' 0.001 '**' 0.01 '*' 0.05 '.' 0.1 ' ' 1
## 
## Residual standard error: 0.04653 on 42 degrees of freedom
## Multiple R-squared:  0.03487,    Adjusted R-squared:  0.01189 
## F-statistic: 1.517 on 1 and 42 DF,  p-value: 0.2249
## 
## 
## Call:
## lm(formula = form, data = dat)
## 
## Standardized Coefficients::
##    (Intercept) provenance_MAP 
##             NA     -0.1867248 
## 
## [1] "Model: plasticity_rdpi_lower_stomata_pore_length_mean_um ~ provenance_TD"
## 
## Call:
## lm(formula = form, data = dat)
## 
## Residuals:
##      Min       1Q   Median       3Q      Max 
## -0.05785 -0.03238 -0.01955  0.01483  0.14100 
## 
## Coefficients:
##                 Estimate Std. Error t value Pr(>|t|)  
## (Intercept)    9.643e-02  4.975e-02   1.938   0.0593 .
## provenance_TD -3.155e-05  1.917e-03  -0.016   0.9870  
## ---
## Signif. codes:  0 '***' 0.001 '**' 0.01 '*' 0.05 '.' 0.1 ' ' 1
## 
## Residual standard error: 0.04737 on 42 degrees of freedom
## Multiple R-squared:  6.444e-06,  Adjusted R-squared:  -0.0238 
## F-statistic: 0.0002707 on 1 and 42 DF,  p-value: 0.987
## 
## 
## Call:
## lm(formula = form, data = dat)
## 
## Standardized Coefficients::
##   (Intercept) provenance_TD 
##            NA  -0.002538602 
## 
## [1] "Model: plasticity_rdpi_upper_stomata_pore_length_mean_um ~ genetic_PC1"
## 
## Call:
## lm(formula = form, data = dat)
## 
## Residuals:
##       Min        1Q    Median        3Q       Max 
## -0.069773 -0.033252 -0.006083  0.013136  0.129368 
## 
## Coefficients:
##              Estimate Std. Error t value Pr(>|t|)    
## (Intercept)  0.123334   0.008551  14.424   <2e-16 ***
## genetic_PC1 -0.174993   0.260073  -0.673    0.505    
## ---
## Signif. codes:  0 '***' 0.001 '**' 0.01 '*' 0.05 '.' 0.1 ' ' 1
## 
## Residual standard error: 0.05094 on 36 degrees of freedom
##   (6 observations deleted due to missingness)
## Multiple R-squared:  0.01242,    Adjusted R-squared:  -0.01501 
## F-statistic: 0.4527 on 1 and 36 DF,  p-value: 0.5053
## 
## 
## Call:
## lm(formula = form, data = dat)
## 
## Standardized Coefficients::
## (Intercept) genetic_PC1 
##          NA  -0.1114451 
## 
## [1] "Model: plasticity_rdpi_upper_stomata_pore_length_mean_um ~ genetic_PC2"
## 
## Call:
## lm(formula = form, data = dat)
## 
## Residuals:
##      Min       1Q   Median       3Q      Max 
## -0.06097 -0.03841 -0.01155  0.02119  0.12061 
## 
## Coefficients:
##              Estimate Std. Error t value Pr(>|t|)    
## (Intercept)  0.128365   0.008648  14.843   <2e-16 ***
## genetic_PC2 -0.264295   0.215818  -1.225    0.229    
## ---
## Signif. codes:  0 '***' 0.001 '**' 0.01 '*' 0.05 '.' 0.1 ' ' 1
## 
## Residual standard error: 0.05022 on 36 degrees of freedom
##   (6 observations deleted due to missingness)
## Multiple R-squared:  0.03999,    Adjusted R-squared:  0.01333 
## F-statistic:   1.5 on 1 and 36 DF,  p-value: 0.2287
## 
## 
## Call:
## lm(formula = form, data = dat)
## 
## Standardized Coefficients::
## (Intercept) genetic_PC2 
##          NA  -0.1999805 
## 
## [1] "Model: plasticity_rdpi_upper_stomata_pore_length_mean_um ~ genetic_PC3"
## 
## Call:
## lm(formula = form, data = dat)
## 
## Residuals:
##       Min        1Q    Median        3Q       Max 
## -0.062584 -0.033765 -0.008951  0.010510  0.136167 
## 
## Coefficients:
##             Estimate Std. Error t value Pr(>|t|)    
## (Intercept) 0.124617   0.008289  15.033   <2e-16 ***
## genetic_PC3 0.112660   0.207374   0.543     0.59    
## ---
## Signif. codes:  0 '***' 0.001 '**' 0.01 '*' 0.05 '.' 0.1 ' ' 1
## 
## Residual standard error: 0.05105 on 36 degrees of freedom
##   (6 observations deleted due to missingness)
## Multiple R-squared:  0.008132,   Adjusted R-squared:  -0.01942 
## F-statistic: 0.2951 on 1 and 36 DF,  p-value: 0.5903
## 
## 
## Call:
## lm(formula = form, data = dat)
## 
## Standardized Coefficients::
## (Intercept) genetic_PC3 
##          NA  0.09017633 
## 
## [1] "Model: plasticity_rdpi_upper_stomata_pore_length_mean_um ~ provenance_MAT"
## 
## Call:
## lm(formula = form, data = dat)
## 
## Residuals:
##      Min       1Q   Median       3Q      Max 
## -0.06357 -0.03227 -0.01057  0.01406  0.13489 
## 
## Coefficients:
##                 Estimate Std. Error t value Pr(>|t|)    
## (Intercept)     0.127350   0.011602  10.976 4.87e-13 ***
## provenance_MAT -0.001053   0.003361  -0.313    0.756    
## ---
## Signif. codes:  0 '***' 0.001 '**' 0.01 '*' 0.05 '.' 0.1 ' ' 1
## 
## Residual standard error: 0.05119 on 36 degrees of freedom
##   (6 observations deleted due to missingness)
## Multiple R-squared:  0.002716,   Adjusted R-squared:  -0.02499 
## F-statistic: 0.09805 on 1 and 36 DF,  p-value: 0.756
## 
## 
## Call:
## lm(formula = form, data = dat)
## 
## Standardized Coefficients::
##    (Intercept) provenance_MAT 
##             NA    -0.05211726 
## 
## [1] "Model: plasticity_rdpi_upper_stomata_pore_length_mean_um ~ provenance_MAP"
## 
## Call:
## lm(formula = form, data = dat)
## 
## Residuals:
##      Min       1Q   Median       3Q      Max 
## -0.06504 -0.03220 -0.01072  0.01404  0.13365 
## 
## Coefficients:
##                  Estimate Std. Error t value Pr(>|t|)    
## (Intercept)     1.312e-01  2.298e-02   5.708 1.71e-06 ***
## provenance_MAP -1.067e-05  3.594e-05  -0.297    0.768    
## ---
## Signif. codes:  0 '***' 0.001 '**' 0.01 '*' 0.05 '.' 0.1 ' ' 1
## 
## Residual standard error: 0.0512 on 36 degrees of freedom
##   (6 observations deleted due to missingness)
## Multiple R-squared:  0.002443,   Adjusted R-squared:  -0.02527 
## F-statistic: 0.08818 on 1 and 36 DF,  p-value: 0.7682
## 
## 
## Call:
## lm(formula = form, data = dat)
## 
## Standardized Coefficients::
##    (Intercept) provenance_MAP 
##             NA    -0.04943175 
## 
## [1] "Model: plasticity_rdpi_upper_stomata_pore_length_mean_um ~ provenance_TD"
## 
## Call:
## lm(formula = form, data = dat)
## 
## Residuals:
##      Min       1Q   Median       3Q      Max 
## -0.06301 -0.03604 -0.01226  0.01688  0.13949 
## 
## Coefficients:
##               Estimate Std. Error t value Pr(>|t|)
## (Intercept)   0.055710   0.058666   0.950    0.349
## provenance_TD 0.002712   0.002280   1.189    0.242
## 
## Residual standard error: 0.05028 on 36 degrees of freedom
##   (6 observations deleted due to missingness)
## Multiple R-squared:  0.03781,    Adjusted R-squared:  0.01109 
## F-statistic: 1.415 on 1 and 36 DF,  p-value: 0.242
## 
## 
## Call:
## lm(formula = form, data = dat)
## 
## Standardized Coefficients::
##   (Intercept) provenance_TD 
##            NA     0.1944597 
## 
## [1] "Model: plasticity_rdpi_lower_stomata_density_mm2 ~ genetic_PC1"
## 
## Call:
## lm(formula = form, data = dat)
## 
## Residuals:
##     Min      1Q  Median      3Q     Max 
## -0.2354 -0.1676 -0.1065  0.1251  0.6399 
## 
## Coefficients:
##             Estimate Std. Error t value Pr(>|t|)    
## (Intercept)  0.34513    0.03613   9.552 4.33e-12 ***
## genetic_PC1  0.33271    1.01976   0.326    0.746    
## ---
## Signif. codes:  0 '***' 0.001 '**' 0.01 '*' 0.05 '.' 0.1 ' ' 1
## 
## Residual standard error: 0.2373 on 42 degrees of freedom
## Multiple R-squared:  0.002528,   Adjusted R-squared:  -0.02122 
## F-statistic: 0.1064 on 1 and 42 DF,  p-value: 0.7458
## 
## 
## Call:
## lm(formula = form, data = dat)
## 
## Standardized Coefficients::
## (Intercept) genetic_PC1 
##          NA  0.05028007 
## 
## [1] "Model: plasticity_rdpi_lower_stomata_density_mm2 ~ genetic_PC2"
## 
## Call:
## lm(formula = form, data = dat)
## 
## Residuals:
##      Min       1Q   Median       3Q      Max 
## -0.28474 -0.15371 -0.09797  0.10556  0.62909 
## 
## Coefficients:
##             Estimate Std. Error t value Pr(>|t|)    
## (Intercept)  0.36059    0.03622   9.955 1.28e-12 ***
## genetic_PC2 -1.58883    0.96099  -1.653    0.106    
## ---
## Signif. codes:  0 '***' 0.001 '**' 0.01 '*' 0.05 '.' 0.1 ' ' 1
## 
## Residual standard error: 0.2303 on 42 degrees of freedom
## Multiple R-squared:  0.06111,    Adjusted R-squared:  0.03875 
## F-statistic: 2.733 on 1 and 42 DF,  p-value: 0.1057
## 
## 
## Call:
## lm(formula = form, data = dat)
## 
## Standardized Coefficients::
## (Intercept) genetic_PC2 
##          NA  -0.2471969 
## 
## [1] "Model: plasticity_rdpi_lower_stomata_density_mm2 ~ genetic_PC3"
## 
## Call:
## lm(formula = form, data = dat)
## 
## Residuals:
##      Min       1Q   Median       3Q      Max 
## -0.29520 -0.19097 -0.07152  0.10969  0.62334 
## 
## Coefficients:
##             Estimate Std. Error t value Pr(>|t|)    
## (Intercept)  0.34095    0.03508   9.718 2.62e-12 ***
## genetic_PC3 -1.24707    0.90223  -1.382    0.174    
## ---
## Signif. codes:  0 '***' 0.001 '**' 0.01 '*' 0.05 '.' 0.1 ' ' 1
## 
## Residual standard error: 0.2324 on 42 degrees of freedom
## Multiple R-squared:  0.04351,    Adjusted R-squared:  0.02074 
## F-statistic:  1.91 on 1 and 42 DF,  p-value: 0.1742
## 
## 
## Call:
## lm(formula = form, data = dat)
## 
## Standardized Coefficients::
## (Intercept) genetic_PC3 
##          NA  -0.2085878 
## 
## [1] "Model: plasticity_rdpi_lower_stomata_density_mm2 ~ provenance_MAT"
## 
## Call:
## lm(formula = form, data = dat)
## 
## Residuals:
##     Min      1Q  Median      3Q     Max 
## -0.2758 -0.1851 -0.0580  0.1154  0.6621 
## 
## Coefficients:
##                Estimate Std. Error t value Pr(>|t|)    
## (Intercept)     0.29285    0.04816   6.081 3.04e-07 ***
## provenance_MAT  0.02046    0.01342   1.524    0.135    
## ---
## Signif. codes:  0 '***' 0.001 '**' 0.01 '*' 0.05 '.' 0.1 ' ' 1
## 
## Residual standard error: 0.2313 on 42 degrees of freedom
## Multiple R-squared:  0.05243,    Adjusted R-squared:  0.02987 
## F-statistic: 2.324 on 1 and 42 DF,  p-value: 0.1349
## 
## 
## Call:
## lm(formula = form, data = dat)
## 
## Standardized Coefficients::
##    (Intercept) provenance_MAT 
##             NA      0.2289695 
## 
## [1] "Model: plasticity_rdpi_lower_stomata_density_mm2 ~ provenance_MAP"
## 
## Call:
## lm(formula = form, data = dat)
## 
## Residuals:
##      Min       1Q   Median       3Q      Max 
## -0.28062 -0.17716 -0.08424  0.11502  0.64701 
## 
## Coefficients:
##                  Estimate Std. Error t value Pr(>|t|)    
## (Intercept)     0.4471725  0.1000036   4.472  5.8e-05 ***
## provenance_MAP -0.0001774  0.0001600  -1.108    0.274    
## ---
## Signif. codes:  0 '***' 0.001 '**' 0.01 '*' 0.05 '.' 0.1 ' ' 1
## 
## Residual standard error: 0.2342 on 42 degrees of freedom
## Multiple R-squared:  0.02841,    Adjusted R-squared:  0.005278 
## F-statistic: 1.228 on 1 and 42 DF,  p-value: 0.2741
## 
## 
## Call:
## lm(formula = form, data = dat)
## 
## Standardized Coefficients::
##    (Intercept) provenance_MAP 
##             NA     -0.1685554 
## 
## [1] "Model: plasticity_rdpi_lower_stomata_density_mm2 ~ provenance_TD"
## 
## Call:
## lm(formula = form, data = dat)
## 
## Residuals:
##     Min      1Q  Median      3Q     Max 
## -0.2399 -0.1712 -0.1068  0.1267  0.6524 
## 
## Coefficients:
##                Estimate Std. Error t value Pr(>|t|)
## (Intercept)   0.3194487  0.2495436   1.280    0.208
## provenance_TD 0.0009362  0.0096187   0.097    0.923
## 
## Residual standard error: 0.2376 on 42 degrees of freedom
## Multiple R-squared:  0.0002255,  Adjusted R-squared:  -0.02358 
## F-statistic: 0.009473 on 1 and 42 DF,  p-value: 0.9229
## 
## 
## Call:
## lm(formula = form, data = dat)
## 
## Standardized Coefficients::
##   (Intercept) provenance_TD 
##            NA    0.01501666 
## 
## [1] "Model: plasticity_rdpi_upper_stomata_density_mm2_log ~ genetic_PC1"
## 
## Call:
## lm(formula = form, data = dat)
## 
## Residuals:
##      Min       1Q   Median       3Q      Max 
## -123.458    3.802    6.106    7.691   15.274 
## 
## Coefficients:
##             Estimate Std. Error t value Pr(>|t|)  
## (Intercept)   -9.837      3.689  -2.666   0.0108 *
## genetic_PC1  -50.811    104.109  -0.488   0.6280  
## ---
## Signif. codes:  0 '***' 0.001 '**' 0.01 '*' 0.05 '.' 0.1 ' ' 1
## 
## Residual standard error: 24.23 on 42 degrees of freedom
## Multiple R-squared:  0.005639,   Adjusted R-squared:  -0.01804 
## F-statistic: 0.2382 on 1 and 42 DF,  p-value: 0.628
## 
## 
## Call:
## lm(formula = form, data = dat)
## 
## Standardized Coefficients::
## (Intercept) genetic_PC1 
##          NA -0.07509601 
## 
## [1] "Model: plasticity_rdpi_upper_stomata_density_mm2_log ~ genetic_PC2"
## 
## Call:
## lm(formula = form, data = dat)
## 
## Residuals:
##      Min       1Q   Median       3Q      Max 
## -124.070    0.362    4.539   10.741   17.352 
## 
## Coefficients:
##             Estimate Std. Error t value Pr(>|t|)   
## (Intercept)  -10.995      3.746  -2.935  0.00538 **
## genetic_PC2  130.923     99.377   1.317  0.19483   
## ---
## Signif. codes:  0 '***' 0.001 '**' 0.01 '*' 0.05 '.' 0.1 ' ' 1
## 
## Residual standard error: 23.81 on 42 degrees of freedom
## Multiple R-squared:  0.03968,    Adjusted R-squared:  0.01682 
## F-statistic: 1.736 on 1 and 42 DF,  p-value: 0.1948
## 
## 
## Call:
## lm(formula = form, data = dat)
## 
## Standardized Coefficients::
## (Intercept) genetic_PC2 
##          NA   0.1992105 
## 
## [1] "Model: plasticity_rdpi_upper_stomata_density_mm2_log ~ genetic_PC3"
## 
## Call:
## lm(formula = form, data = dat)
## 
## Residuals:
##      Min       1Q   Median       3Q      Max 
## -123.432   -0.568    6.198   10.101   15.404 
## 
## Coefficients:
##             Estimate Std. Error t value Pr(>|t|)  
## (Intercept)   -9.321      3.585  -2.600   0.0128 *
## genetic_PC3  129.692     92.182   1.407   0.1668  
## ---
## Signif. codes:  0 '***' 0.001 '**' 0.01 '*' 0.05 '.' 0.1 ' ' 1
## 
## Residual standard error: 23.74 on 42 degrees of freedom
## Multiple R-squared:  0.04501,    Adjusted R-squared:  0.02227 
## F-statistic: 1.979 on 1 and 42 DF,  p-value: 0.1668
## 
## 
## Call:
## lm(formula = form, data = dat)
## 
## Standardized Coefficients::
## (Intercept) genetic_PC3 
##          NA   0.2121492 
## 
## [1] "Model: plasticity_rdpi_upper_stomata_density_mm2_log ~ provenance_MAT"
## 
## Call:
## lm(formula = form, data = dat)
## 
## Residuals:
##      Min       1Q   Median       3Q      Max 
## -126.696    0.098    5.942    8.973   15.983 
## 
## Coefficients:
##                Estimate Std. Error t value Pr(>|t|)
## (Intercept)      -4.930      4.950  -0.996    0.325
## provenance_MAT   -1.881      1.379  -1.363    0.180
## 
## Residual standard error: 23.78 on 42 degrees of freedom
## Multiple R-squared:  0.04239,    Adjusted R-squared:  0.01959 
## F-statistic: 1.859 on 1 and 42 DF,  p-value: 0.18
## 
## 
## Call:
## lm(formula = form, data = dat)
## 
## Standardized Coefficients::
##    (Intercept) provenance_MAT 
##             NA     -0.2058831 
## 
## [1] "Model: plasticity_rdpi_upper_stomata_density_mm2_log ~ provenance_MAP"
## 
## Call:
## lm(formula = form, data = dat)
## 
## Residuals:
##      Min       1Q   Median       3Q      Max 
## -125.121    1.816    6.832    8.768   13.882 
## 
## Coefficients:
##                 Estimate Std. Error t value Pr(>|t|)  
## (Intercept)    -21.12871   10.19773  -2.072   0.0445 *
## provenance_MAP   0.01975    0.01632   1.210   0.2331  
## ---
## Signif. codes:  0 '***' 0.001 '**' 0.01 '*' 0.05 '.' 0.1 ' ' 1
## 
## Residual standard error: 23.89 on 42 degrees of freedom
## Multiple R-squared:  0.03368,    Adjusted R-squared:  0.01067 
## F-statistic: 1.464 on 1 and 42 DF,  p-value: 0.2331
## 
## 
## Call:
## lm(formula = form, data = dat)
## 
## Standardized Coefficients::
##    (Intercept) provenance_MAP 
##             NA      0.1835267 
## 
## [1] "Model: plasticity_rdpi_upper_stomata_density_mm2_log ~ provenance_TD"
## 
## Call:
## lm(formula = form, data = dat)
## 
## Residuals:
##      Min       1Q   Median       3Q      Max 
## -126.011    3.552    6.324    8.200   12.416 
## 
## Coefficients:
##               Estimate Std. Error t value Pr(>|t|)
## (Intercept)    -2.3344    25.4940  -0.092    0.927
## provenance_TD  -0.2824     0.9827  -0.287    0.775
## 
## Residual standard error: 24.27 on 42 degrees of freedom
## Multiple R-squared:  0.001962,   Adjusted R-squared:  -0.0218 
## F-statistic: 0.08259 on 1 and 42 DF,  p-value: 0.7752
## 
## 
## Call:
## lm(formula = form, data = dat)
## 
## Standardized Coefficients::
##   (Intercept) provenance_TD 
##            NA   -0.04429961 
## 
## [1] "Model: plasticity_rdpi_upper_stomata_presence ~ genetic_PC1"
## 
## Call:
## lm(formula = form, data = dat)
## 
## Residuals:
##     Min      1Q  Median      3Q     Max 
## -0.7444 -0.3740  0.1721  0.3130  0.4662 
## 
## Coefficients:
##             Estimate Std. Error t value Pr(>|t|)    
## (Intercept)   0.6590     0.0612  10.767  1.6e-13 ***
## genetic_PC1   2.6225     1.7751   1.477    0.147    
## ---
## Signif. codes:  0 '***' 0.001 '**' 0.01 '*' 0.05 '.' 0.1 ' ' 1
## 
## Residual standard error: 0.394 on 41 degrees of freedom
##   (1 observation deleted due to missingness)
## Multiple R-squared:  0.05055,    Adjusted R-squared:  0.02739 
## F-statistic: 2.183 on 1 and 41 DF,  p-value: 0.1472
## 
## 
## Call:
## lm(formula = form, data = dat)
## 
## Standardized Coefficients::
## (Intercept) genetic_PC1 
##          NA   0.2248249 
## 
## [1] "Model: plasticity_rdpi_upper_stomata_presence ~ genetic_PC2"
## 
## Call:
## lm(formula = form, data = dat)
## 
## Residuals:
##     Min      1Q  Median      3Q     Max 
## -0.6626 -0.3705  0.2295  0.3574  0.3810 
## 
## Coefficients:
##             Estimate Std. Error t value Pr(>|t|)    
## (Intercept)  0.64573    0.06447  10.015  1.4e-12 ***
## genetic_PC2 -0.35417    1.69180  -0.209    0.835    
## ---
## Signif. codes:  0 '***' 0.001 '**' 0.01 '*' 0.05 '.' 0.1 ' ' 1
## 
## Residual standard error: 0.4041 on 41 degrees of freedom
##   (1 observation deleted due to missingness)
## Multiple R-squared:  0.001068,   Adjusted R-squared:  -0.0233 
## F-statistic: 0.04382 on 1 and 41 DF,  p-value: 0.8352
## 
## 
## Call:
## lm(formula = form, data = dat)
## 
## Standardized Coefficients::
## (Intercept) genetic_PC2 
##          NA -0.03267637 
## 
## [1] "Model: plasticity_rdpi_upper_stomata_presence ~ genetic_PC3"
## 
## Call:
## lm(formula = form, data = dat)
## 
## Residuals:
##     Min      1Q  Median      3Q     Max 
## -0.7336 -0.3091  0.1598  0.2822  0.5011 
## 
## Coefficients:
##             Estimate Std. Error t value Pr(>|t|)    
## (Intercept)  0.63497    0.05905  10.753 1.67e-13 ***
## genetic_PC3 -2.95219    1.50216  -1.965   0.0562 .  
## ---
## Signif. codes:  0 '***' 0.001 '**' 0.01 '*' 0.05 '.' 0.1 ' ' 1
## 
## Residual standard error: 0.3865 on 41 degrees of freedom
##   (1 observation deleted due to missingness)
## Multiple R-squared:  0.08609,    Adjusted R-squared:  0.0638 
## F-statistic: 3.862 on 1 and 41 DF,  p-value: 0.05618
## 
## 
## Call:
## lm(formula = form, data = dat)
## 
## Standardized Coefficients::
## (Intercept) genetic_PC3 
##          NA  -0.2934184 
## 
## [1] "Model: plasticity_rdpi_upper_stomata_presence ~ provenance_MAT"
## 
## Call:
## lm(formula = form, data = dat)
## 
## Residuals:
##     Min      1Q  Median      3Q     Max 
## -0.6602 -0.4012  0.2200  0.3525  0.4376 
## 
## Coefficients:
##                Estimate Std. Error t value Pr(>|t|)    
## (Intercept)     0.68473    0.08858   7.730 1.57e-09 ***
## provenance_MAT -0.01653    0.02458  -0.672    0.505    
## ---
## Signif. codes:  0 '***' 0.001 '**' 0.01 '*' 0.05 '.' 0.1 ' ' 1
## 
## Residual standard error: 0.4021 on 41 degrees of freedom
##   (1 observation deleted due to missingness)
## Multiple R-squared:  0.0109, Adjusted R-squared:  -0.01322 
## F-statistic: 0.4519 on 1 and 41 DF,  p-value: 0.5052
## 
## 
## Call:
## lm(formula = form, data = dat)
## 
## Standardized Coefficients::
##    (Intercept) provenance_MAT 
##             NA     -0.1044076 
## 
## [1] "Model: plasticity_rdpi_upper_stomata_presence ~ provenance_MAP"
## 
## Call:
## lm(formula = form, data = dat)
## 
## Residuals:
##     Min      1Q  Median      3Q     Max 
## -0.7218 -0.3556  0.1687  0.3351  0.5498 
## 
## Coefficients:
##                  Estimate Std. Error t value Pr(>|t|)    
## (Intercept)     0.8419479  0.1727481   4.874 1.68e-05 ***
## provenance_MAP -0.0003395  0.0002744  -1.237    0.223    
## ---
## Signif. codes:  0 '***' 0.001 '**' 0.01 '*' 0.05 '.' 0.1 ' ' 1
## 
## Residual standard error: 0.397 on 41 degrees of freedom
##   (1 observation deleted due to missingness)
## Multiple R-squared:  0.036,  Adjusted R-squared:  0.01248 
## F-statistic: 1.531 on 1 and 41 DF,  p-value: 0.223
## 
## 
## Call:
## lm(formula = form, data = dat)
## 
## Standardized Coefficients::
##    (Intercept) provenance_MAP 
##             NA     -0.1897269 
## 
## [1] "Model: plasticity_rdpi_upper_stomata_presence ~ provenance_TD"
## 
## Call:
## lm(formula = form, data = dat)
## 
## Residuals:
##      Min       1Q   Median       3Q      Max 
## -0.78944 -0.37346  0.07696  0.34417  0.48645 
## 
## Coefficients:
##               Estimate Std. Error t value Pr(>|t|)
## (Intercept)    0.05316    0.45142   0.118    0.907
## provenance_TD  0.02314    0.01758   1.316    0.196
## 
## Residual standard error: 0.3961 on 41 degrees of freedom
##   (1 observation deleted due to missingness)
## Multiple R-squared:  0.04051,    Adjusted R-squared:  0.01711 
## F-statistic: 1.731 on 1 and 41 DF,  p-value: 0.1956
## 
## 
## Call:
## lm(formula = form, data = dat)
## 
## Standardized Coefficients::
##   (Intercept) provenance_TD 
##            NA     0.2012753 
## 
## [1] "Model: plasticity_rdpi_upper_stomata_density_over_total_density_log ~ genetic_PC1"
## 
## Call:
## lm(formula = form, data = dat)
## 
## Residuals:
##      Min       1Q   Median       3Q      Max 
## -0.50103 -0.04109  0.03670  0.08096  0.21068 
## 
## Coefficients:
##             Estimate Std. Error t value Pr(>|t|)    
## (Intercept) -0.26245    0.02357 -11.134 4.13e-14 ***
## genetic_PC1  0.82008    0.66526   1.233    0.225    
## ---
## Signif. codes:  0 '***' 0.001 '**' 0.01 '*' 0.05 '.' 0.1 ' ' 1
## 
## Residual standard error: 0.1548 on 42 degrees of freedom
## Multiple R-squared:  0.03492,    Adjusted R-squared:  0.01194 
## F-statistic:  1.52 on 1 and 42 DF,  p-value: 0.2245
## 
## 
## Call:
## lm(formula = form, data = dat)
## 
## Standardized Coefficients::
## (Intercept) genetic_PC1 
##          NA   0.1868628 
## 
## [1] "Model: plasticity_rdpi_upper_stomata_density_over_total_density_log ~ genetic_PC2"
## 
## Call:
## lm(formula = form, data = dat)
## 
## Residuals:
##      Min       1Q   Median       3Q      Max 
## -0.44282 -0.06986 -0.00068  0.11076  0.28490 
## 
## Coefficients:
##             Estimate Std. Error t value Pr(>|t|)    
## (Intercept) -0.27723    0.02411 -11.500 1.47e-14 ***
## genetic_PC2  0.99593    0.63956   1.557    0.127    
## ---
## Signif. codes:  0 '***' 0.001 '**' 0.01 '*' 0.05 '.' 0.1 ' ' 1
## 
## Residual standard error: 0.1532 on 42 degrees of freedom
## Multiple R-squared:  0.05458,    Adjusted R-squared:  0.03207 
## F-statistic: 2.425 on 1 and 42 DF,  p-value: 0.1269
## 
## 
## Call:
## lm(formula = form, data = dat)
## 
## Standardized Coefficients::
## (Intercept) genetic_PC2 
##          NA   0.2336335 
## 
## [1] "Model: plasticity_rdpi_upper_stomata_density_over_total_density_log ~ genetic_PC3"
## 
## Call:
## lm(formula = form, data = dat)
## 
## Residuals:
##      Min       1Q   Median       3Q      Max 
## -0.45001 -0.03974  0.01354  0.08856  0.26221 
## 
## Coefficients:
##             Estimate Std. Error t value Pr(>|t|)    
## (Intercept) -0.26574    0.02368 -11.220 3.23e-14 ***
## genetic_PC3  0.37613    0.60908   0.618     0.54    
## ---
## Signif. codes:  0 '***' 0.001 '**' 0.01 '*' 0.05 '.' 0.1 ' ' 1
## 
## Residual standard error: 0.1569 on 42 degrees of freedom
## Multiple R-squared:  0.008998,   Adjusted R-squared:  -0.0146 
## F-statistic: 0.3814 on 1 and 42 DF,  p-value: 0.5402
## 
## 
## Call:
## lm(formula = form, data = dat)
## 
## Standardized Coefficients::
## (Intercept) genetic_PC3 
##          NA  0.09485913 
## 
## [1] "Model: plasticity_rdpi_upper_stomata_density_over_total_density_log ~ provenance_MAT"
## 
## Call:
## lm(formula = form, data = dat)
## 
## Residuals:
##      Min       1Q   Median       3Q      Max 
## -0.46345 -0.04172  0.02981  0.08637  0.21888 
## 
## Coefficients:
##                 Estimate Std. Error t value Pr(>|t|)    
## (Intercept)    -0.235562   0.032072  -7.345 4.69e-09 ***
## provenance_MAT -0.012503   0.008937  -1.399    0.169    
## ---
## Signif. codes:  0 '***' 0.001 '**' 0.01 '*' 0.05 '.' 0.1 ' ' 1
## 
## Residual standard error: 0.1541 on 42 degrees of freedom
## Multiple R-squared:  0.04453,    Adjusted R-squared:  0.02178 
## F-statistic: 1.957 on 1 and 42 DF,  p-value: 0.1691
## 
## 
## Call:
## lm(formula = form, data = dat)
## 
## Standardized Coefficients::
##    (Intercept) provenance_MAT 
##             NA     -0.2110153 
## 
## [1] "Model: plasticity_rdpi_upper_stomata_density_over_total_density_log ~ provenance_MAP"
## 
## Call:
## lm(formula = form, data = dat)
## 
## Residuals:
##      Min       1Q   Median       3Q      Max 
## -0.45166 -0.03215  0.00761  0.10108  0.30045 
## 
## Coefficients:
##                 Estimate Std. Error t value Pr(>|t|)    
## (Intercept)    -0.357686   0.065586  -5.454 2.41e-06 ***
## provenance_MAP  0.000156   0.000105   1.486    0.145    
## ---
## Signif. codes:  0 '***' 0.001 '**' 0.01 '*' 0.05 '.' 0.1 ' ' 1
## 
## Residual standard error: 0.1536 on 42 degrees of freedom
## Multiple R-squared:  0.04995,    Adjusted R-squared:  0.02733 
## F-statistic: 2.208 on 1 and 42 DF,  p-value: 0.1448
## 
## 
## Call:
## lm(formula = form, data = dat)
## 
## Standardized Coefficients::
##    (Intercept) provenance_MAP 
##             NA      0.2234846 
## 
## [1] "Model: plasticity_rdpi_upper_stomata_density_over_total_density_log ~ provenance_TD"
## 
## Call:
## lm(formula = form, data = dat)
## 
## Residuals:
##      Min       1Q   Median       3Q      Max 
## -0.46575 -0.02966  0.02833  0.08761  0.25351 
## 
## Coefficients:
##                Estimate Std. Error t value Pr(>|t|)  
## (Intercept)   -0.299801   0.165440  -1.812   0.0771 .
## provenance_TD  0.001297   0.006377   0.203   0.8398  
## ---
## Signif. codes:  0 '***' 0.001 '**' 0.01 '*' 0.05 '.' 0.1 ' ' 1
## 
## Residual standard error: 0.1575 on 42 degrees of freedom
## Multiple R-squared:  0.0009835,  Adjusted R-squared:  -0.0228 
## F-statistic: 0.04135 on 1 and 42 DF,  p-value: 0.8398
## 
## 
## Call:
## lm(formula = form, data = dat)
## 
## Standardized Coefficients::
##   (Intercept) provenance_TD 
##            NA    0.03136111 
## 
## [1] "Model: plasticity_rdpi_licor_gsw ~ genetic_PC1"
## 
## Call:
## lm(formula = form, data = dat)
## 
## Residuals:
##      Min       1Q   Median       3Q      Max 
## -0.13294 -0.06052 -0.01463  0.05283  0.22167 
## 
## Coefficients:
##             Estimate Std. Error t value Pr(>|t|)    
## (Intercept)  0.22400    0.01273  17.594   <2e-16 ***
## genetic_PC1  0.78513    0.35931   2.185   0.0345 *  
## ---
## Signif. codes:  0 '***' 0.001 '**' 0.01 '*' 0.05 '.' 0.1 ' ' 1
## 
## Residual standard error: 0.08362 on 42 degrees of freedom
## Multiple R-squared:  0.1021, Adjusted R-squared:  0.0807 
## F-statistic: 4.775 on 1 and 42 DF,  p-value: 0.03451
## 
## 
## Call:
## lm(formula = form, data = dat)
## 
## Standardized Coefficients::
## (Intercept) genetic_PC1 
##          NA   0.3194947 
## 
## [1] "Model: plasticity_rdpi_licor_gsw ~ genetic_PC2"
## 
## Call:
## lm(formula = form, data = dat)
## 
## Residuals:
##       Min        1Q    Median        3Q       Max 
## -0.136792 -0.069753 -0.009806  0.059481  0.252857 
## 
## Coefficients:
##             Estimate Std. Error t value Pr(>|t|)    
## (Intercept)  0.22172    0.01386  16.003   <2e-16 ***
## genetic_PC2 -0.14961    0.36759  -0.407    0.686    
## ---
## Signif. codes:  0 '***' 0.001 '**' 0.01 '*' 0.05 '.' 0.1 ' ' 1
## 
## Residual standard error: 0.08807 on 42 degrees of freedom
## Multiple R-squared:  0.003929,   Adjusted R-squared:  -0.01979 
## F-statistic: 0.1657 on 1 and 42 DF,  p-value: 0.6861
## 
## 
## Call:
## lm(formula = form, data = dat)
## 
## Standardized Coefficients::
## (Intercept) genetic_PC2 
##          NA -0.06267879 
## 
## [1] "Model: plasticity_rdpi_licor_gsw ~ genetic_PC3"
## 
## Call:
## lm(formula = form, data = dat)
## 
## Residuals:
##      Min       1Q   Median       3Q      Max 
## -0.13676 -0.06995 -0.01289  0.05976  0.25513 
## 
## Coefficients:
##             Estimate Std. Error t value Pr(>|t|)    
## (Intercept)  0.21996    0.01331  16.521   <2e-16 ***
## genetic_PC3 -0.07727    0.34239  -0.226    0.823    
## ---
## Signif. codes:  0 '***' 0.001 '**' 0.01 '*' 0.05 '.' 0.1 ' ' 1
## 
## Residual standard error: 0.08819 on 42 degrees of freedom
## Multiple R-squared:  0.001211,   Adjusted R-squared:  -0.02257 
## F-statistic: 0.05093 on 1 and 42 DF,  p-value: 0.8226
## 
## 
## Call:
## lm(formula = form, data = dat)
## 
## Standardized Coefficients::
## (Intercept) genetic_PC3 
##          NA     -0.0348 
## 
## [1] "Model: plasticity_rdpi_licor_gsw ~ provenance_MAT"
## 
## Call:
## lm(formula = form, data = dat)
## 
## Residuals:
##      Min       1Q   Median       3Q      Max 
## -0.13305 -0.07010 -0.01497  0.05753  0.24958 
## 
## Coefficients:
##                  Estimate Std. Error t value Pr(>|t|)    
## (Intercept)     0.2220841  0.0183673  12.091 2.89e-15 ***
## provenance_MAT -0.0007961  0.0051182  -0.156    0.877    
## ---
## Signif. codes:  0 '***' 0.001 '**' 0.01 '*' 0.05 '.' 0.1 ' ' 1
## 
## Residual standard error: 0.08822 on 42 degrees of freedom
## Multiple R-squared:  0.0005757,  Adjusted R-squared:  -0.02322 
## F-statistic: 0.02419 on 1 and 42 DF,  p-value: 0.8771
## 
## 
## Call:
## lm(formula = form, data = dat)
## 
## Standardized Coefficients::
##    (Intercept) provenance_MAT 
##             NA    -0.02399425 
## 
## [1] "Model: plasticity_rdpi_licor_gsw ~ provenance_MAP"
## 
## Call:
## lm(formula = form, data = dat)
## 
## Residuals:
##      Min       1Q   Median       3Q      Max 
## -0.12367 -0.06982 -0.01571  0.05700  0.23477 
## 
## Coefficients:
##                 Estimate Std. Error t value Pr(>|t|)    
## (Intercept)    1.971e-01  3.749e-02   5.257  4.6e-06 ***
## provenance_MAP 3.942e-05  5.999e-05   0.657    0.515    
## ---
## Signif. codes:  0 '***' 0.001 '**' 0.01 '*' 0.05 '.' 0.1 ' ' 1
## 
## Residual standard error: 0.0878 on 42 degrees of freedom
## Multiple R-squared:  0.01017,    Adjusted R-squared:  -0.01339 
## F-statistic: 0.4317 on 1 and 42 DF,  p-value: 0.5147
## 
## 
## Call:
## lm(formula = form, data = dat)
## 
## Standardized Coefficients::
##    (Intercept) provenance_MAP 
##             NA      0.1008694 
## 
## [1] "Model: plasticity_rdpi_licor_gsw ~ provenance_TD"
## 
## Call:
## lm(formula = form, data = dat)
## 
## Residuals:
##      Min       1Q   Median       3Q      Max 
## -0.13375 -0.06952 -0.01207  0.05481  0.26027 
## 
## Coefficients:
##               Estimate Std. Error t value Pr(>|t|)  
## (Intercept)   0.168989   0.092340   1.830   0.0743 .
## provenance_TD 0.001991   0.003559   0.559   0.5788  
## ---
## Signif. codes:  0 '***' 0.001 '**' 0.01 '*' 0.05 '.' 0.1 ' ' 1
## 
## Residual standard error: 0.08792 on 42 degrees of freedom
## Multiple R-squared:  0.007397,   Adjusted R-squared:  -0.01624 
## F-statistic: 0.313 on 1 and 42 DF,  p-value: 0.5788
## 
## 
## Call:
## lm(formula = form, data = dat)
## 
## Standardized Coefficients::
##   (Intercept) provenance_TD 
##            NA    0.08600502 
## 
## [1] "Model: plasticity_rdpi_licor_ETR ~ genetic_PC1"
## 
## Call:
## lm(formula = form, data = dat)
## 
## Residuals:
##       Min        1Q    Median        3Q       Max 
## -0.272093 -0.062521 -0.007274  0.085373  0.244771 
## 
## Coefficients:
##             Estimate Std. Error t value Pr(>|t|)    
## (Intercept)  0.46672    0.02009  23.235   <2e-16 ***
## genetic_PC1 -0.19353    0.56689  -0.341    0.735    
## ---
## Signif. codes:  0 '***' 0.001 '**' 0.01 '*' 0.05 '.' 0.1 ' ' 1
## 
## Residual standard error: 0.1319 on 42 degrees of freedom
## Multiple R-squared:  0.002767,   Adjusted R-squared:  -0.02098 
## F-statistic: 0.1166 on 1 and 42 DF,  p-value: 0.7345
## 
## 
## Call:
## lm(formula = form, data = dat)
## 
## Standardized Coefficients::
## (Intercept) genetic_PC1 
##          NA -0.05260544 
## 
## [1] "Model: plasticity_rdpi_licor_ETR ~ genetic_PC2"
## 
## Call:
## lm(formula = form, data = dat)
## 
## Residuals:
##       Min        1Q    Median        3Q       Max 
## -0.276356 -0.074312  0.000283  0.083060  0.245917 
## 
## Coefficients:
##             Estimate Std. Error t value Pr(>|t|)    
## (Intercept)  0.47234    0.02063  22.896   <2e-16 ***
## genetic_PC2 -0.43297    0.54733  -0.791    0.433    
## ---
## Signif. codes:  0 '***' 0.001 '**' 0.01 '*' 0.05 '.' 0.1 ' ' 1
## 
## Residual standard error: 0.1311 on 42 degrees of freedom
## Multiple R-squared:  0.01468,    Adjusted R-squared:  -0.008779 
## F-statistic: 0.6258 on 1 and 42 DF,  p-value: 0.4334
## 
## 
## Call:
## lm(formula = form, data = dat)
## 
## Standardized Coefficients::
## (Intercept) genetic_PC2 
##          NA  -0.1211641 
## 
## [1] "Model: plasticity_rdpi_licor_ETR ~ genetic_PC3"
## 
## Call:
## lm(formula = form, data = dat)
## 
## Residuals:
##       Min        1Q    Median        3Q       Max 
## -0.287018 -0.073555  0.004389  0.084291  0.257437 
## 
## Coefficients:
##             Estimate Std. Error t value Pr(>|t|)    
## (Intercept)  0.46679    0.01977  23.606   <2e-16 ***
## genetic_PC3 -0.43313    0.50852  -0.852    0.399    
## ---
## Signif. codes:  0 '***' 0.001 '**' 0.01 '*' 0.05 '.' 0.1 ' ' 1
## 
## Residual standard error: 0.131 on 42 degrees of freedom
## Multiple R-squared:  0.01698,    Adjusted R-squared:  -0.006426 
## F-statistic: 0.7255 on 1 and 42 DF,  p-value: 0.3992
## 
## 
## Call:
## lm(formula = form, data = dat)
## 
## Standardized Coefficients::
## (Intercept) genetic_PC3 
##          NA  -0.1303054 
## 
## [1] "Model: plasticity_rdpi_licor_ETR ~ provenance_MAT"
## 
## Call:
## lm(formula = form, data = dat)
## 
## Residuals:
##       Min        1Q    Median        3Q       Max 
## -0.274273 -0.072903 -0.003044  0.089146  0.233243 
## 
## Coefficients:
##                Estimate Std. Error t value Pr(>|t|)    
## (Intercept)    0.458844   0.027434  16.725   <2e-16 ***
## provenance_MAT 0.003568   0.007645   0.467    0.643    
## ---
## Signif. codes:  0 '***' 0.001 '**' 0.01 '*' 0.05 '.' 0.1 ' ' 1
## 
## Residual standard error: 0.1318 on 42 degrees of freedom
## Multiple R-squared:  0.005159,   Adjusted R-squared:  -0.01853 
## F-statistic: 0.2178 on 1 and 42 DF,  p-value: 0.6431
## 
## 
## Call:
## lm(formula = form, data = dat)
## 
## Standardized Coefficients::
##    (Intercept) provenance_MAT 
##             NA     0.07182818 
## 
## [1] "Model: plasticity_rdpi_licor_ETR ~ provenance_MAP"
## 
## Call:
## lm(formula = form, data = dat)
## 
## Residuals:
##       Min        1Q    Median        3Q       Max 
## -0.274935 -0.071022 -0.001468  0.085927  0.270686 
## 
## Coefficients:
##                  Estimate Std. Error t value Pr(>|t|)    
## (Intercept)     5.050e-01  5.607e-02   9.006 2.32e-11 ***
## provenance_MAP -6.381e-05  8.973e-05  -0.711    0.481    
## ---
## Signif. codes:  0 '***' 0.001 '**' 0.01 '*' 0.05 '.' 0.1 ' ' 1
## 
## Residual standard error: 0.1313 on 42 degrees of freedom
## Multiple R-squared:  0.0119, Adjusted R-squared:  -0.01163 
## F-statistic: 0.5057 on 1 and 42 DF,  p-value: 0.481
## 
## 
## Call:
## lm(formula = form, data = dat)
## 
## Standardized Coefficients::
##    (Intercept) provenance_MAP 
##             NA     -0.1090701 
## 
## [1] "Model: plasticity_rdpi_licor_ETR ~ provenance_TD"
## 
## Call:
## lm(formula = form, data = dat)
## 
## Residuals:
##       Min        1Q    Median        3Q       Max 
## -0.275311 -0.062874 -0.008473  0.092486  0.232198 
## 
## Coefficients:
##                Estimate Std. Error t value Pr(>|t|)   
## (Intercept)   0.4573091  0.1387440   3.296    0.002 **
## provenance_TD 0.0004037  0.0053479   0.075    0.940   
## ---
## Signif. codes:  0 '***' 0.001 '**' 0.01 '*' 0.05 '.' 0.1 ' ' 1
## 
## Residual standard error: 0.1321 on 42 degrees of freedom
## Multiple R-squared:  0.0001357,  Adjusted R-squared:  -0.02367 
## F-statistic: 0.005699 on 1 and 42 DF,  p-value: 0.9402
## 
## 
## Call:
## lm(formula = form, data = dat)
## 
## Standardized Coefficients::
##   (Intercept) provenance_TD 
##            NA    0.01164755 
## 
## [1] "Model: plasticity_rdpi_licor_Fm. ~ genetic_PC1"
## 
## Call:
## lm(formula = form, data = dat)
## 
## Residuals:
##       Min        1Q    Median        3Q       Max 
## -0.135702 -0.027903 -0.001883  0.035649  0.102667 
## 
## Coefficients:
##              Estimate Std. Error t value Pr(>|t|)    
## (Intercept)  0.231249   0.008136  28.422   <2e-16 ***
## genetic_PC1 -0.298705   0.229624  -1.301      0.2    
## ---
## Signif. codes:  0 '***' 0.001 '**' 0.01 '*' 0.05 '.' 0.1 ' ' 1
## 
## Residual standard error: 0.05344 on 42 degrees of freedom
## Multiple R-squared:  0.03873,    Adjusted R-squared:  0.01584 
## F-statistic: 1.692 on 1 and 42 DF,  p-value: 0.2004
## 
## 
## Call:
## lm(formula = form, data = dat)
## 
## Standardized Coefficients::
## (Intercept) genetic_PC1 
##          NA  -0.1967993 
## 
## [1] "Model: plasticity_rdpi_licor_Fm. ~ genetic_PC2"
## 
## Call:
## lm(formula = form, data = dat)
## 
## Residuals:
##       Min        1Q    Median        3Q       Max 
## -0.131282 -0.034523  0.004171  0.031518  0.101371 
## 
## Coefficients:
##              Estimate Std. Error t value Pr(>|t|)    
## (Intercept)  0.235853   0.008407  28.055   <2e-16 ***
## genetic_PC2 -0.290166   0.223037  -1.301      0.2    
## ---
## Signif. codes:  0 '***' 0.001 '**' 0.01 '*' 0.05 '.' 0.1 ' ' 1
## 
## Residual standard error: 0.05344 on 42 degrees of freedom
## Multiple R-squared:  0.03874,    Adjusted R-squared:  0.01585 
## F-statistic: 1.693 on 1 and 42 DF,  p-value: 0.2004
## 
## 
## Call:
## lm(formula = form, data = dat)
## 
## Standardized Coefficients::
## (Intercept) genetic_PC2 
##          NA  -0.1968183 
## 
## [1] "Model: plasticity_rdpi_licor_Fm. ~ genetic_PC3"
## 
## Call:
## lm(formula = form, data = dat)
## 
## Residuals:
##       Min        1Q    Median        3Q       Max 
## -0.127188 -0.030279  0.002802  0.025207  0.097450 
## 
## Coefficients:
##              Estimate Std. Error t value Pr(>|t|)    
## (Intercept)  0.232157   0.008054  28.826   <2e-16 ***
## genetic_PC3 -0.280973   0.207115  -1.357    0.182    
## ---
## Signif. codes:  0 '***' 0.001 '**' 0.01 '*' 0.05 '.' 0.1 ' ' 1
## 
## Residual standard error: 0.05335 on 42 degrees of freedom
## Multiple R-squared:  0.04198,    Adjusted R-squared:  0.01917 
## F-statistic:  1.84 on 1 and 42 DF,  p-value: 0.1822
## 
## 
## Call:
## lm(formula = form, data = dat)
## 
## Standardized Coefficients::
## (Intercept) genetic_PC3 
##          NA  -0.2048878 
## 
## [1] "Model: plasticity_rdpi_licor_Fm. ~ provenance_MAT"
## 
## Call:
## lm(formula = form, data = dat)
## 
## Residuals:
##       Min        1Q    Median        3Q       Max 
## -0.126928 -0.027193  0.002535  0.035606  0.095914 
## 
## Coefficients:
##                Estimate Std. Error t value Pr(>|t|)    
## (Intercept)    0.222621   0.011120  20.020   <2e-16 ***
## provenance_MAT 0.004084   0.003099   1.318    0.195    
## ---
## Signif. codes:  0 '***' 0.001 '**' 0.01 '*' 0.05 '.' 0.1 ' ' 1
## 
## Residual standard error: 0.05341 on 42 degrees of freedom
## Multiple R-squared:  0.03971,    Adjusted R-squared:  0.01685 
## F-statistic: 1.737 on 1 and 42 DF,  p-value: 0.1947
## 
## 
## Call:
## lm(formula = form, data = dat)
## 
## Standardized Coefficients::
##    (Intercept) provenance_MAT 
##             NA      0.1992739 
## 
## [1] "Model: plasticity_rdpi_licor_Fm. ~ provenance_MAP"
## 
## Call:
## lm(formula = form, data = dat)
## 
## Residuals:
##       Min        1Q    Median        3Q       Max 
## -0.141297 -0.026155 -0.001721  0.031741  0.105666 
## 
## Coefficients:
##                  Estimate Std. Error t value Pr(>|t|)    
## (Intercept)     2.355e-01  2.327e-02  10.124 7.76e-13 ***
## provenance_MAP -4.820e-06  3.724e-05  -0.129    0.898    
## ---
## Signif. codes:  0 '***' 0.001 '**' 0.01 '*' 0.05 '.' 0.1 ' ' 1
## 
## Residual standard error: 0.0545 on 42 degrees of freedom
## Multiple R-squared:  0.0003988,  Adjusted R-squared:  -0.0234 
## F-statistic: 0.01676 on 1 and 42 DF,  p-value: 0.8976
## 
## 
## Call:
## lm(formula = form, data = dat)
## 
## Standardized Coefficients::
##    (Intercept) provenance_MAP 
##             NA    -0.01997042 
## 
## [1] "Model: plasticity_rdpi_licor_Fm. ~ provenance_TD"
## 
## Call:
## lm(formula = form, data = dat)
## 
## Residuals:
##       Min        1Q    Median        3Q       Max 
## -0.137870 -0.026007 -0.001896  0.032370  0.102262 
## 
## Coefficients:
##                Estimate Std. Error t value Pr(>|t|)    
## (Intercept)    0.261617   0.057068   4.584 4.06e-05 ***
## provenance_TD -0.001125   0.002200  -0.512    0.612    
## ---
## Signif. codes:  0 '***' 0.001 '**' 0.01 '*' 0.05 '.' 0.1 ' ' 1
## 
## Residual standard error: 0.05434 on 42 degrees of freedom
## Multiple R-squared:  0.006191,   Adjusted R-squared:  -0.01747 
## F-statistic: 0.2616 on 1 and 42 DF,  p-value: 0.6117
## 
## 
## Call:
## lm(formula = form, data = dat)
## 
## Standardized Coefficients::
##   (Intercept) provenance_TD 
##            NA   -0.07868296 
## 
## [1] "Model: plasticity_rdpi_licor_Fs ~ genetic_PC1"
## 
## Call:
## lm(formula = form, data = dat)
## 
## Residuals:
##       Min        1Q    Median        3Q       Max 
## -0.084266 -0.030592 -0.002018  0.031021  0.109175 
## 
## Coefficients:
##             Estimate Std. Error t value Pr(>|t|)    
## (Intercept) 0.132869   0.006451   20.60   <2e-16 ***
## genetic_PC1 0.320442   0.182068    1.76   0.0857 .  
## ---
## Signif. codes:  0 '***' 0.001 '**' 0.01 '*' 0.05 '.' 0.1 ' ' 1
## 
## Residual standard error: 0.04237 on 42 degrees of freedom
## Multiple R-squared:  0.06869,    Adjusted R-squared:  0.04651 
## F-statistic: 3.098 on 1 and 42 DF,  p-value: 0.08569
## 
## 
## Call:
## lm(formula = form, data = dat)
## 
## Standardized Coefficients::
## (Intercept) genetic_PC1 
##          NA   0.2620823 
## 
## [1] "Model: plasticity_rdpi_licor_Fs ~ genetic_PC2"
## 
## Call:
## lm(formula = form, data = dat)
## 
## Residuals:
##       Min        1Q    Median        3Q       Max 
## -0.080692 -0.025153 -0.004947  0.027406  0.113305 
## 
## Coefficients:
##             Estimate Std. Error t value Pr(>|t|)    
## (Intercept) 0.129540   0.006843  18.931   <2e-16 ***
## genetic_PC2 0.161800   0.181543   0.891    0.378    
## ---
## Signif. codes:  0 '***' 0.001 '**' 0.01 '*' 0.05 '.' 0.1 ' ' 1
## 
## Residual standard error: 0.0435 on 42 degrees of freedom
## Multiple R-squared:  0.01856,    Adjusted R-squared:  -0.004806 
## F-statistic: 0.7943 on 1 and 42 DF,  p-value: 0.3779
## 
## 
## Call:
## lm(formula = form, data = dat)
## 
## Standardized Coefficients::
## (Intercept) genetic_PC2 
##          NA   0.1362407 
## 
## [1] "Model: plasticity_rdpi_licor_Fs ~ genetic_PC3"
## 
## Call:
## lm(formula = form, data = dat)
## 
## Residuals:
##       Min        1Q    Median        3Q       Max 
## -0.089181 -0.023631 -0.000692  0.029214  0.099386 
## 
## Coefficients:
##             Estimate Std. Error t value Pr(>|t|)    
## (Intercept) 0.131981   0.006299  20.951   <2e-16 ***
## genetic_PC3 0.343692   0.161998   2.122   0.0398 *  
## ---
## Signif. codes:  0 '***' 0.001 '**' 0.01 '*' 0.05 '.' 0.1 ' ' 1
## 
## Residual standard error: 0.04173 on 42 degrees of freedom
## Multiple R-squared:  0.0968, Adjusted R-squared:  0.07529 
## F-statistic: 4.501 on 1 and 42 DF,  p-value: 0.03982
## 
## 
## Call:
## lm(formula = form, data = dat)
## 
## Standardized Coefficients::
## (Intercept) genetic_PC3 
##          NA   0.3111203 
## 
## [1] "Model: plasticity_rdpi_licor_Fs ~ provenance_MAT"
## 
## Call:
## lm(formula = form, data = dat)
## 
## Residuals:
##       Min        1Q    Median        3Q       Max 
## -0.090066 -0.029514  0.001309  0.026737  0.096596 
## 
## Coefficients:
##                 Estimate Std. Error t value Pr(>|t|)    
## (Intercept)     0.145894   0.008537  17.090   <2e-16 ***
## provenance_MAT -0.005904   0.002379  -2.482   0.0172 *  
## ---
## Signif. codes:  0 '***' 0.001 '**' 0.01 '*' 0.05 '.' 0.1 ' ' 1
## 
## Residual standard error: 0.041 on 42 degrees of freedom
## Multiple R-squared:  0.1279, Adjusted R-squared:  0.1071 
## F-statistic:  6.16 on 1 and 42 DF,  p-value: 0.01715
## 
## 
## Call:
## lm(formula = form, data = dat)
## 
## Standardized Coefficients::
##    (Intercept) provenance_MAT 
##             NA     -0.3576317 
## 
## [1] "Model: plasticity_rdpi_licor_Fs ~ provenance_MAP"
## 
## Call:
## lm(formula = form, data = dat)
## 
## Residuals:
##       Min        1Q    Median        3Q       Max 
## -0.076606 -0.027706 -0.001946  0.029981  0.114936 
## 
## Coefficients:
##                  Estimate Std. Error t value Pr(>|t|)    
## (Intercept)     1.415e-01  1.867e-02   7.578 2.19e-09 ***
## provenance_MAP -1.746e-05  2.988e-05  -0.584    0.562    
## ---
## Signif. codes:  0 '***' 0.001 '**' 0.01 '*' 0.05 '.' 0.1 ' ' 1
## 
## Residual standard error: 0.04373 on 42 degrees of freedom
## Multiple R-squared:  0.008068,   Adjusted R-squared:  -0.01555 
## F-statistic: 0.3416 on 1 and 42 DF,  p-value: 0.562
## 
## 
## Call:
## lm(formula = form, data = dat)
## 
## Standardized Coefficients::
##    (Intercept) provenance_MAP 
##             NA    -0.08981987 
## 
## [1] "Model: plasticity_rdpi_licor_Fs ~ provenance_TD"
## 
## Call:
## lm(formula = form, data = dat)
## 
## Residuals:
##       Min        1Q    Median        3Q       Max 
## -0.070574 -0.031587 -0.005447  0.030888  0.101239 
## 
## Coefficients:
##               Estimate Std. Error t value Pr(>|t|)   
## (Intercept)   0.008713   0.041968   0.208  0.83653   
## provenance_TD 0.004774   0.001618   2.951  0.00516 **
## ---
## Signif. codes:  0 '***' 0.001 '**' 0.01 '*' 0.05 '.' 0.1 ' ' 1
## 
## Residual standard error: 0.03996 on 42 degrees of freedom
## Multiple R-squared:  0.1717, Adjusted R-squared:  0.152 
## F-statistic: 8.709 on 1 and 42 DF,  p-value: 0.005161
## 
## 
## Call:
## lm(formula = form, data = dat)
## 
## Standardized Coefficients::
##   (Intercept) provenance_TD 
##            NA     0.4144225 
## 
## [1] "Model: plasticity_rdpi_licor_gbw ~ genetic_PC1"
## 
## Call:
## lm(formula = form, data = dat)
## 
## Residuals:
##        Min         1Q     Median         3Q        Max 
## -1.251e-04 -5.968e-05 -2.218e-05  3.146e-05  2.650e-04 
## 
## Coefficients:
##              Estimate Std. Error t value Pr(>|t|)    
## (Intercept) 4.103e-04  1.347e-05  30.464   <2e-16 ***
## genetic_PC1 1.013e-04  3.801e-04   0.266    0.791    
## ---
## Signif. codes:  0 '***' 0.001 '**' 0.01 '*' 0.05 '.' 0.1 ' ' 1
## 
## Residual standard error: 8.847e-05 on 42 degrees of freedom
## Multiple R-squared:  0.001688,   Adjusted R-squared:  -0.02208 
## F-statistic: 0.071 on 1 and 42 DF,  p-value: 0.7912
## 
## 
## Call:
## lm(formula = form, data = dat)
## 
## Standardized Coefficients::
## (Intercept) genetic_PC1 
##          NA   0.0410802 
## 
## [1] "Model: plasticity_rdpi_licor_gbw ~ genetic_PC2"
## 
## Call:
## lm(formula = form, data = dat)
## 
## Residuals:
##        Min         1Q     Median         3Q        Max 
## -1.190e-04 -6.315e-05 -2.385e-05  3.837e-05  2.620e-04 
## 
## Coefficients:
##              Estimate Std. Error t value Pr(>|t|)    
## (Intercept) 4.095e-04  1.393e-05  29.404   <2e-16 ***
## genetic_PC2 2.797e-05  3.695e-04   0.076     0.94    
## ---
## Signif. codes:  0 '***' 0.001 '**' 0.01 '*' 0.05 '.' 0.1 ' ' 1
## 
## Residual standard error: 8.854e-05 on 42 degrees of freedom
## Multiple R-squared:  0.0001364,  Adjusted R-squared:  -0.02367 
## F-statistic: 0.005729 on 1 and 42 DF,  p-value: 0.94
## 
## 
## Call:
## lm(formula = form, data = dat)
## 
## Standardized Coefficients::
## (Intercept) genetic_PC2 
##          NA  0.01167849 
## 
## [1] "Model: plasticity_rdpi_licor_gbw ~ genetic_PC3"
## 
## Call:
## lm(formula = form, data = dat)
## 
## Residuals:
##        Min         1Q     Median         3Q        Max 
## -1.182e-04 -6.238e-05 -2.295e-05  3.862e-05  2.630e-04 
## 
## Coefficients:
##              Estimate Std. Error t value Pr(>|t|)    
## (Intercept) 4.099e-04  1.336e-05   30.68   <2e-16 ***
## genetic_PC3 5.161e-05  3.436e-04    0.15    0.881    
## ---
## Signif. codes:  0 '***' 0.001 '**' 0.01 '*' 0.05 '.' 0.1 ' ' 1
## 
## Residual standard error: 8.852e-05 on 42 degrees of freedom
## Multiple R-squared:  0.0005367,  Adjusted R-squared:  -0.02326 
## F-statistic: 0.02256 on 1 and 42 DF,  p-value: 0.8813
## 
## 
## Call:
## lm(formula = form, data = dat)
## 
## Standardized Coefficients::
## (Intercept) genetic_PC3 
##          NA  0.02316785 
## 
## [1] "Model: plasticity_rdpi_licor_gbw ~ provenance_MAT"
## 
## Call:
## lm(formula = form, data = dat)
## 
## Residuals:
##        Min         1Q     Median         3Q        Max 
## -1.203e-04 -6.316e-05 -2.055e-05  2.930e-05  2.758e-04 
## 
## Coefficients:
##                  Estimate Std. Error t value Pr(>|t|)    
## (Intercept)     4.177e-04  1.835e-05  22.763   <2e-16 ***
## provenance_MAT -3.178e-06  5.113e-06  -0.621    0.538    
## ---
## Signif. codes:  0 '***' 0.001 '**' 0.01 '*' 0.05 '.' 0.1 ' ' 1
## 
## Residual standard error: 8.814e-05 on 42 degrees of freedom
## Multiple R-squared:  0.009111,   Adjusted R-squared:  -0.01448 
## F-statistic: 0.3862 on 1 and 42 DF,  p-value: 0.5377
## 
## 
## Call:
## lm(formula = form, data = dat)
## 
## Standardized Coefficients::
##    (Intercept) provenance_MAT 
##             NA    -0.09544969 
## 
## [1] "Model: plasticity_rdpi_licor_gbw ~ provenance_MAP"
## 
## Call:
## lm(formula = form, data = dat)
## 
## Residuals:
##        Min         1Q     Median         3Q        Max 
## -1.190e-04 -6.051e-05 -2.016e-05  4.307e-05  2.559e-04 
## 
## Coefficients:
##                 Estimate Std. Error t value Pr(>|t|)    
## (Intercept)    3.818e-04  3.752e-05   10.18 6.66e-13 ***
## provenance_MAP 4.802e-08  6.005e-08    0.80    0.428    
## ---
## Signif. codes:  0 '***' 0.001 '**' 0.01 '*' 0.05 '.' 0.1 ' ' 1
## 
## Residual standard error: 8.788e-05 on 42 degrees of freedom
## Multiple R-squared:  0.015,  Adjusted R-squared:  -0.008456 
## F-statistic: 0.6395 on 1 and 42 DF,  p-value: 0.4284
## 
## 
## Call:
## lm(formula = form, data = dat)
## 
## Standardized Coefficients::
##    (Intercept) provenance_MAP 
##             NA      0.1224611 
## 
## [1] "Model: plasticity_rdpi_licor_gbw ~ provenance_TD"
## 
## Call:
## lm(formula = form, data = dat)
## 
## Residuals:
##        Min         1Q     Median         3Q        Max 
## -1.189e-04 -6.320e-05 -2.481e-05  3.980e-05  2.598e-04 
## 
## Coefficients:
##                 Estimate Std. Error t value Pr(>|t|)    
## (Intercept)    4.225e-04  9.297e-05   4.545  4.6e-05 ***
## provenance_TD -4.949e-07  3.584e-06  -0.138    0.891    
## ---
## Signif. codes:  0 '***' 0.001 '**' 0.01 '*' 0.05 '.' 0.1 ' ' 1
## 
## Residual standard error: 8.852e-05 on 42 degrees of freedom
## Multiple R-squared:  0.0004539,  Adjusted R-squared:  -0.02334 
## F-statistic: 0.01907 on 1 and 42 DF,  p-value: 0.8908
## 
## 
## Call:
## lm(formula = form, data = dat)
## 
## Standardized Coefficients::
##   (Intercept) provenance_TD 
##            NA   -0.02130484 
## 
## [1] "Model: plasticity_rdpi_licor_PhiPS2 ~ genetic_PC1"
## 
## Call:
## lm(formula = form, data = dat)
## 
## Residuals:
##       Min        1Q    Median        3Q       Max 
## -0.120604 -0.028580  0.008035  0.037595  0.085443 
## 
## Coefficients:
##             Estimate Std. Error t value Pr(>|t|)    
## (Intercept) 0.256723   0.007646  33.576   <2e-16 ***
## genetic_PC1 0.060722   0.215785   0.281     0.78    
## ---
## Signif. codes:  0 '***' 0.001 '**' 0.01 '*' 0.05 '.' 0.1 ' ' 1
## 
## Residual standard error: 0.05022 on 42 degrees of freedom
## Multiple R-squared:  0.001882,   Adjusted R-squared:  -0.02188 
## F-statistic: 0.07919 on 1 and 42 DF,  p-value: 0.7798
## 
## 
## Call:
## lm(formula = form, data = dat)
## 
## Standardized Coefficients::
## (Intercept) genetic_PC1 
##          NA  0.04337995 
## 
## [1] "Model: plasticity_rdpi_licor_PhiPS2 ~ genetic_PC2"
## 
## Call:
## lm(formula = form, data = dat)
## 
## Residuals:
##       Min        1Q    Median        3Q       Max 
## -0.118321 -0.028871  0.009182  0.037267  0.085769 
## 
## Coefficients:
##              Estimate Std. Error t value Pr(>|t|)    
## (Intercept)  0.256677   0.007906  32.465   <2e-16 ***
## genetic_PC2 -0.023733   0.209761  -0.113     0.91    
## ---
## Signif. codes:  0 '***' 0.001 '**' 0.01 '*' 0.05 '.' 0.1 ' ' 1
## 
## Residual standard error: 0.05026 on 42 degrees of freedom
## Multiple R-squared:  0.0003047,  Adjusted R-squared:  -0.0235 
## F-statistic: 0.0128 on 1 and 42 DF,  p-value: 0.9105
## 
## 
## Call:
## lm(formula = form, data = dat)
## 
## Standardized Coefficients::
## (Intercept) genetic_PC2 
##          NA -0.01745541 
## 
## [1] "Model: plasticity_rdpi_licor_PhiPS2 ~ genetic_PC3"
## 
## Call:
## lm(formula = form, data = dat)
## 
## Residuals:
##       Min        1Q    Median        3Q       Max 
## -0.116203 -0.031245  0.008593  0.035397  0.088490 
## 
## Coefficients:
##             Estimate Std. Error t value Pr(>|t|)    
## (Intercept)  0.25630    0.00758  33.811   <2e-16 ***
## genetic_PC3 -0.05780    0.19494  -0.296    0.768    
## ---
## Signif. codes:  0 '***' 0.001 '**' 0.01 '*' 0.05 '.' 0.1 ' ' 1
## 
## Residual standard error: 0.05021 on 42 degrees of freedom
## Multiple R-squared:  0.002089,   Adjusted R-squared:  -0.02167 
## F-statistic: 0.08791 on 1 and 42 DF,  p-value: 0.7683
## 
## 
## Call:
## lm(formula = form, data = dat)
## 
## Standardized Coefficients::
## (Intercept) genetic_PC3 
##          NA -0.04570307 
## 
## [1] "Model: plasticity_rdpi_licor_PhiPS2 ~ provenance_MAT"
## 
## Call:
## lm(formula = form, data = dat)
## 
## Residuals:
##       Min        1Q    Median        3Q       Max 
## -0.119334 -0.029060  0.008064  0.037103  0.084165 
## 
## Coefficients:
##                  Estimate Std. Error t value Pr(>|t|)    
## (Intercept)     0.2569991  0.0104643   24.56   <2e-16 ***
## provenance_MAT -0.0002332  0.0029160   -0.08    0.937    
## ---
## Signif. codes:  0 '***' 0.001 '**' 0.01 '*' 0.05 '.' 0.1 ' ' 1
## 
## Residual standard error: 0.05026 on 42 degrees of freedom
## Multiple R-squared:  0.0001523,  Adjusted R-squared:  -0.02365 
## F-statistic: 0.006396 on 1 and 42 DF,  p-value: 0.9366
## 
## 
## Call:
## lm(formula = form, data = dat)
## 
## Standardized Coefficients::
##    (Intercept) provenance_MAT 
##             NA    -0.01233921 
## 
## [1] "Model: plasticity_rdpi_licor_PhiPS2 ~ provenance_MAP"
## 
## Call:
## lm(formula = form, data = dat)
## 
## Residuals:
##       Min        1Q    Median        3Q       Max 
## -0.118330 -0.029397  0.006907  0.037884  0.083278 
## 
## Coefficients:
##                 Estimate Std. Error t value Pr(>|t|)    
## (Intercept)    2.548e-01  2.146e-02  11.872 5.27e-15 ***
## provenance_MAP 2.845e-06  3.434e-05   0.083    0.934    
## ---
## Signif. codes:  0 '***' 0.001 '**' 0.01 '*' 0.05 '.' 0.1 ' ' 1
## 
## Residual standard error: 0.05026 on 42 degrees of freedom
## Multiple R-squared:  0.0001633,  Adjusted R-squared:  -0.02364 
## F-statistic: 0.00686 on 1 and 42 DF,  p-value: 0.9344
## 
## 
## Call:
## lm(formula = form, data = dat)
## 
## Standardized Coefficients::
##    (Intercept) provenance_MAP 
##             NA     0.01277954 
## 
## [1] "Model: plasticity_rdpi_licor_PhiPS2 ~ provenance_TD"
## 
## Call:
## lm(formula = form, data = dat)
## 
## Residuals:
##       Min        1Q    Median        3Q       Max 
## -0.116861 -0.026918  0.005274  0.039658  0.090134 
## 
## Coefficients:
##               Estimate Std. Error t value Pr(>|t|)    
## (Intercept)   0.195166   0.051922   3.759 0.000521 ***
## provenance_TD 0.002386   0.002001   1.192 0.239905    
## ---
## Signif. codes:  0 '***' 0.001 '**' 0.01 '*' 0.05 '.' 0.1 ' ' 1
## 
## Residual standard error: 0.04944 on 42 degrees of freedom
## Multiple R-squared:  0.03273,    Adjusted R-squared:  0.009699 
## F-statistic: 1.421 on 1 and 42 DF,  p-value: 0.2399
## 
## 
## Call:
## lm(formula = form, data = dat)
## 
## Standardized Coefficients::
##   (Intercept) provenance_TD 
##            NA     0.1809132
```

```
hist(unlist(res.lr$eff), breaks = 20, main = 'Effect Sizes')
```

```
hist(unlist(res.lr$eff.std), breaks = 20, main = 'Standardized Effect Sizes')
```

```
hist(unlist(res.lr$cors), breaks = 20, main = 'correlations') # same as std effect sizes
```

```
hist(unlist(res.lr$cors.sig), main= 'signficant correlations')
```

```
# adjust p-values

padj <- p.adjust(unlist(res.lr$pvals), method = 'BH')
res.lr$padj <- as.data.frame(matrix(padj, ncol = ncol(res.lr$pvals)))
colnames(res.lr$padj) <- colnames(res.lr$pvals)
rownames(res.lr$padj) <- rownames(res.lr$pvals)

plot(unlist(res.lr$pvals), unlist(res.lr$padj), xlab = 'pvals', ylab = 'adjusted pvals')
```

```
sum(res.lr$pvals < 0.05) # 19
```

```
## [1] 19
```

```
sum(res.lr$pvals < 0.01) # 7
```

```
## [1] 7
```

```
sum(res.lr$padj < 0.05) # 2
```

```
## [1] 2
```

```
sum(res.lr$padj < 0.1) # 2
```

```
## [1] 2
```

#### 3.2 Tables with results

```
kable(res.lr$eff.std.sig, label = 'Standardized effects for significant relationships between plasticity and climate/genetics')
```

|  | plasticity\_rdpi\_growing\_season\_days\_2023 | plasticity\_rdpi\_growing\_season\_days\_2024 | plasticity\_rdpi\_DOY\_Stage2\_2023 | plasticity\_rdpi\_DOY\_Stage2\_2024 | plasticity\_rdpi\_DOY\_Stage3\_2023 | plasticity\_rdpi\_DOY\_Stage3\_2024 | plasticity\_rdpi\_DOY\_Stage6\_2023 | plasticity\_rdpi\_DOY\_Stage6\_2024 | plasticity\_rdpi\_DOY\_Stage7\_2023 | plasticity\_rdpi\_DOY\_Stage7\_2024 | plasticity\_rdpi\_stage7\_presence\_2023 | plasticity\_rdpi\_stage7\_presence\_2024 | plasticity\_rdpi\_DOY\_last\_budset\_2023 | plasticity\_rdpi\_DOY\_last\_budset\_2024 | plasticity\_rdpi\_leaf\_thickness\_avg\_mm\_2023 | plasticity\_rdpi\_leaf\_area\_cm2\_2023\_log | plasticity\_rdpi\_leaf\_mass\_g\_2023\_log | plasticity\_rdpi\_LMA\_g\_m2\_2023 | plasticity\_rdpi\_lower\_stomata\_pore\_length\_mean\_um | plasticity\_rdpi\_upper\_stomata\_pore\_length\_mean\_um | plasticity\_rdpi\_lower\_stomata\_density\_mm2 | plasticity\_rdpi\_upper\_stomata\_density\_mm2\_log | plasticity\_rdpi\_upper\_stomata\_presence | plasticity\_rdpi\_upper\_stomata\_density\_over\_total\_density\_log | plasticity\_rdpi\_licor\_gsw | plasticity\_rdpi\_licor\_ETR | plasticity\_rdpi\_licor\_Fm. | plasticity\_rdpi\_licor\_Fs | plasticity\_rdpi\_licor\_gbw | plasticity\_rdpi\_licor\_PhiPS2 |
| --- | --- | --- | --- | --- | --- | --- | --- | --- | --- | --- | --- | --- | --- | --- | --- | --- | --- | --- | --- | --- | --- | --- | --- | --- | --- | --- | --- | --- | --- | --- |
| genetic\_PC1 | NA | 0.3449963 | NA | NA | NA | NA | 0.3809366 | NA | NA | NA | NA | 0.5729064 | NA | NA | NA | NA | NA | 0.3714367 | NA | NA | NA | NA | NA | NA | 0.3194947 | NA | NA | NA | NA | NA |
| genetic\_PC2 | NA | 0.3148440 | NA | NA | NA | NA | NA | NA | NA | NA | NA | NA | NA | NA | NA | NA | NA | NA | NA | NA | NA | NA | NA | NA | NA | NA | NA | NA | NA | NA |
| genetic\_PC3 | NA | NA | NA | NA | NA | NA | 0.3108837 | NA | NA | NA | NA | NA | NA | NA | NA | 0.4215522 | NA | NA | NA | NA | NA | NA | NA | NA | NA | NA | NA | 0.3111203 | NA | NA |
| provenance\_MAT | NA | NA | NA | NA | NA | NA | -0.4173931 | NA | NA | NA | NA | -0.5253690 | NA | NA | NA | -0.4219645 | NA | NA | NA | NA | NA | NA | NA | NA | NA | NA | NA | -0.3576317 | NA | NA |
| provenance\_MAP | NA | NA | NA | NA | NA | -0.3955644 | NA | NA | NA | NA | NA | NA | NA | NA | NA | NA | -0.350529 | NA | NA | NA | NA | NA | NA | NA | NA | NA | NA | NA | NA | NA |
| provenance\_TD | NA | NA | NA | NA | NA | NA | NA | NA | NA | NA | NA | 0.3341572 | NA | NA | NA | 0.3310394 | NA | 0.3272347 | NA | NA | NA | NA | NA | NA | NA | NA | NA | 0.4144225 | NA | NA |

```
kable(res.lr$pvals, label = 'Unadjusted p-values for relationships between plasticity and climate/genetics' )
```

|  | plasticity\_rdpi\_growing\_season\_days\_2023 | plasticity\_rdpi\_growing\_season\_days\_2024 | plasticity\_rdpi\_DOY\_Stage2\_2023 | plasticity\_rdpi\_DOY\_Stage2\_2024 | plasticity\_rdpi\_DOY\_Stage3\_2023 | plasticity\_rdpi\_DOY\_Stage3\_2024 | plasticity\_rdpi\_DOY\_Stage6\_2023 | plasticity\_rdpi\_DOY\_Stage6\_2024 | plasticity\_rdpi\_DOY\_Stage7\_2023 | plasticity\_rdpi\_DOY\_Stage7\_2024 | plasticity\_rdpi\_stage7\_presence\_2023 | plasticity\_rdpi\_stage7\_presence\_2024 | plasticity\_rdpi\_DOY\_last\_budset\_2023 | plasticity\_rdpi\_DOY\_last\_budset\_2024 | plasticity\_rdpi\_leaf\_thickness\_avg\_mm\_2023 | plasticity\_rdpi\_leaf\_area\_cm2\_2023\_log | plasticity\_rdpi\_leaf\_mass\_g\_2023\_log | plasticity\_rdpi\_LMA\_g\_m2\_2023 | plasticity\_rdpi\_lower\_stomata\_pore\_length\_mean\_um | plasticity\_rdpi\_upper\_stomata\_pore\_length\_mean\_um | plasticity\_rdpi\_lower\_stomata\_density\_mm2 | plasticity\_rdpi\_upper\_stomata\_density\_mm2\_log | plasticity\_rdpi\_upper\_stomata\_presence | plasticity\_rdpi\_upper\_stomata\_density\_over\_total\_density\_log | plasticity\_rdpi\_licor\_gsw | plasticity\_rdpi\_licor\_ETR | plasticity\_rdpi\_licor\_Fm. | plasticity\_rdpi\_licor\_Fs | plasticity\_rdpi\_licor\_gbw | plasticity\_rdpi\_licor\_PhiPS2 |
| --- | --- | --- | --- | --- | --- | --- | --- | --- | --- | --- | --- | --- | --- | --- | --- | --- | --- | --- | --- | --- | --- | --- | --- | --- | --- | --- | --- | --- | --- | --- |
| genetic\_PC1 | 0.2469982 | 0.0252504 | 0.9084763 | 0.8499890 | 0.7796268 | 0.4118649 | 0.0107384 | 0.9134285 | 0.1704821 | 0.8911584 | 0.6104333 | 0.0000595 | 0.9751263 | 0.0686096 | 0.1567162 | 0.6094436 | 0.0513106 | 0.0130491 | 0.7972079 | 0.5053315 | 0.7458443 | 0.6280486 | 0.1472082 | 0.2245352 | 0.0345122 | 0.7345085 | 0.2004036 | 0.0856883 | 0.7911926 | 0.7797861 |
| genetic\_PC2 | 0.6993202 | 0.0422736 | 0.6994469 | 0.9055115 | 0.7998917 | 0.6620880 | 0.5399195 | 0.8875861 | 0.6210335 | 0.7118415 | 0.4502624 | 0.0652158 | 0.4176835 | 0.0595752 | 0.1793199 | 0.0595235 | 0.0853877 | 0.7472239 | 0.9011761 | 0.2286717 | 0.1057205 | 0.1948341 | 0.8352175 | 0.1269237 | 0.6860710 | 0.4333528 | 0.2003594 | 0.3778722 | 0.9400248 | 0.9104577 |
| genetic\_PC3 | 0.7622530 | 0.5784690 | 0.9709885 | 0.5638739 | 0.6008042 | 0.1991465 | 0.0399748 | 0.1598448 | 0.1676754 | 0.1393539 | 0.5678659 | 0.0746193 | 0.3091923 | 0.4169055 | 0.4172147 | 0.0043732 | 0.8180812 | 0.5926732 | 0.0704063 | 0.5902914 | 0.1742152 | 0.1668114 | 0.0561789 | 0.5402112 | 0.8225554 | 0.3991904 | 0.1821544 | 0.0398158 | 0.8813375 | 0.7683099 |
| provenance\_MAT | 0.6550132 | 0.1880717 | 0.9248281 | 0.3609891 | 0.4676382 | 0.5029581 | 0.0048189 | 0.3623216 | 0.0723359 | 0.0671355 | 0.8749584 | 0.0002975 | 0.0565226 | 0.0758352 | 0.1161709 | 0.0043310 | 0.7714620 | 0.3329415 | 0.9237109 | 0.7559896 | 0.1349067 | 0.1799939 | 0.5052217 | 0.1691435 | 0.8771367 | 0.6431227 | 0.1946891 | 0.0171541 | 0.5376805 | 0.9366383 |
| provenance\_MAP | 0.4826572 | 0.2126431 | 0.3436500 | 0.1385876 | 0.1827363 | 0.0086521 | 0.5076394 | 0.2325832 | 0.2271989 | 0.2210758 | 0.8614147 | 0.4179876 | 0.9114365 | 0.2308526 | 0.7622222 | 0.9323423 | 0.0196605 | 0.7309812 | 0.2248837 | 0.7682087 | 0.2740728 | 0.2330679 | 0.2230088 | 0.1447663 | 0.5147273 | 0.4809547 | 0.8976204 | 0.5620363 | 0.4284091 | 0.9343821 |
| provenance\_TD | 0.6657679 | 0.4444357 | 0.3301425 | 0.7251629 | 0.3242830 | 0.4684258 | 0.0709443 | 0.3952220 | 0.7626314 | 0.1167792 | 0.5184805 | 0.0285222 | 0.2413421 | 0.2507173 | 0.4964683 | 0.0281670 | 0.1343058 | 0.0301413 | 0.9869516 | 0.2420430 | 0.9229273 | 0.7752373 | 0.1955789 | 0.8398487 | 0.5788266 | 0.9401834 | 0.6116731 | 0.0051611 | 0.8908194 | 0.2399055 |

```
kable(res.lr$padj, label = 'Adjusted p-values for relationships between plasticity and climate/genetics' )
```

|  | plasticity\_rdpi\_growing\_season\_days\_2023 | plasticity\_rdpi\_growing\_season\_days\_2024 | plasticity\_rdpi\_DOY\_Stage2\_2023 | plasticity\_rdpi\_DOY\_Stage2\_2024 | plasticity\_rdpi\_DOY\_Stage3\_2023 | plasticity\_rdpi\_DOY\_Stage3\_2024 | plasticity\_rdpi\_DOY\_Stage6\_2023 | plasticity\_rdpi\_DOY\_Stage6\_2024 | plasticity\_rdpi\_DOY\_Stage7\_2023 | plasticity\_rdpi\_DOY\_Stage7\_2024 | plasticity\_rdpi\_stage7\_presence\_2023 | plasticity\_rdpi\_stage7\_presence\_2024 | plasticity\_rdpi\_DOY\_last\_budset\_2023 | plasticity\_rdpi\_DOY\_last\_budset\_2024 | plasticity\_rdpi\_leaf\_thickness\_avg\_mm\_2023 | plasticity\_rdpi\_leaf\_area\_cm2\_2023\_log | plasticity\_rdpi\_leaf\_mass\_g\_2023\_log | plasticity\_rdpi\_LMA\_g\_m2\_2023 | plasticity\_rdpi\_lower\_stomata\_pore\_length\_mean\_um | plasticity\_rdpi\_upper\_stomata\_pore\_length\_mean\_um | plasticity\_rdpi\_lower\_stomata\_density\_mm2 | plasticity\_rdpi\_upper\_stomata\_density\_mm2\_log | plasticity\_rdpi\_upper\_stomata\_presence | plasticity\_rdpi\_upper\_stomata\_density\_over\_total\_density\_log | plasticity\_rdpi\_licor\_gsw | plasticity\_rdpi\_licor\_ETR | plasticity\_rdpi\_licor\_Fm. | plasticity\_rdpi\_licor\_Fs | plasticity\_rdpi\_licor\_gbw | plasticity\_rdpi\_licor\_PhiPS2 |
| --- | --- | --- | --- | --- | --- | --- | --- | --- | --- | --- | --- | --- | --- | --- | --- | --- | --- | --- | --- | --- | --- | --- | --- | --- | --- | --- | --- | --- | --- | --- |
| genetic\_PC1 | 0.5849958 | 0.3616961 | 0.9561187 | 0.9561187 | 0.9548401 | 0.8090082 | 0.2416146 | 0.9561187 | 0.5809032 | 0.9561187 | 0.9024685 | 0.0107027 | 0.9805739 | 0.4265731 | 0.5809032 | 0.9024685 | 0.4265731 | 0.2609820 | 0.9561187 | 0.8620292 | 0.9548401 | 0.9116834 | 0.5809032 | 0.5809032 | 0.3882625 | 0.9548401 | 0.5809032 | 0.4536439 | 0.9561187 | 0.9548401 |
| genetic\_PC2 | 0.9548401 | 0.4004869 | 0.9548401 | 0.9561187 | 0.9561187 | 0.9362362 | 0.8760182 | 0.9561187 | 0.9088295 | 0.9548401 | 0.8355384 | 0.4265731 | 0.8090082 | 0.4265731 | 0.5809032 | 0.4265731 | 0.4536439 | 0.9548401 | 0.9561187 | 0.5809032 | 0.5437052 | 0.5809032 | 0.9561187 | 0.5809032 | 0.9548401 | 0.8210894 | 0.5809032 | 0.7908953 | 0.9561187 | 0.9561187 |
| genetic\_PC3 | 0.9548401 | 0.8981792 | 0.9805739 | 0.8966304 | 0.9024685 | 0.5809032 | 0.3997485 | 0.5809032 | 0.5809032 | 0.5809032 | 0.8966304 | 0.4265731 | 0.7044889 | 0.8090082 | 0.8090082 | 0.1548324 | 0.9561187 | 0.9024685 | 0.4265731 | 0.9024685 | 0.5809032 | 0.5809032 | 0.4265731 | 0.8760182 | 0.9561187 | 0.8090082 | 0.5809032 | 0.3997485 | 0.9561187 | 0.9548401 |
| provenance\_MAT | 0.9357331 | 0.5809032 | 0.9561187 | 0.7672692 | 0.8516832 | 0.8620292 | 0.1548324 | 0.7672692 | 0.4265731 | 0.4265731 | 0.9561187 | 0.0267748 | 0.4265731 | 0.4265731 | 0.5681149 | 0.1548324 | 0.9548401 | 0.7308471 | 0.9561187 | 0.9548401 | 0.5809032 | 0.5809032 | 0.8620292 | 0.5809032 | 0.9561187 | 0.9260966 | 0.5809032 | 0.3087744 | 0.8760182 | 0.9561187 |
| provenance\_MAP | 0.8601811 | 0.5809032 | 0.7452651 | 0.5809032 | 0.5809032 | 0.2224833 | 0.8620292 | 0.5809032 | 0.5809032 | 0.5809032 | 0.9561187 | 0.8090082 | 0.9561187 | 0.5809032 | 0.9548401 | 0.9561187 | 0.3217174 | 0.9548401 | 0.5809032 | 0.9548401 | 0.6324758 | 0.5809032 | 0.5809032 | 0.5809032 | 0.8641342 | 0.8601811 | 0.9561187 | 0.8966304 | 0.8203579 | 0.9561187 |
| provenance\_TD | 0.9362362 | 0.8333170 | 0.7308471 | 0.9548401 | 0.7296366 | 0.8516832 | 0.4265731 | 0.8090082 | 0.9548401 | 0.5681149 | 0.8641342 | 0.3616961 | 0.5809032 | 0.5860923 | 0.8620292 | 0.3616961 | 0.5809032 | 0.3616961 | 0.9869516 | 0.5809032 | 0.9561187 | 0.9548401 | 0.5809032 | 0.9561187 | 0.8981792 | 0.9561187 | 0.9024685 | 0.1548324 | 0.9561187 | 0.5809032 |

#### 3.3 Plots of variables with that significantly predict trait plasticity

```
# set up clean trait names
trait_names <- colnames(res.lr$eff.std.sig)

trait_names <- gsub('plasticity_rdpi_', '', trait_names)

trait_names <- gsub('Stage2_cGDD', 'Stage2_Bud_Flush_cGDD', trait_names)
trait_names <- gsub('DOY_Stage2', 'Stage2_Bud_Flush_DOY', trait_names)
trait_names <- gsub('DOY_Stage3', 'Stage3_Leaf_Emergence_DOY', trait_names)
trait_names <- gsub('DOY_Stage6', 'Stage6_First_Budset_DOY', trait_names)
trait_names <- gsub('DOY_Stage7', 'Stage7_Lammas_Growth_DOY', trait_names)
trait_names <- gsub('stage7_presence', 'Stage7_Lammas_Growth_presence', trait_names)
trait_names <- gsub('DOY_Stage8', 'Stage8_Budset_after_Lammas_DOY', trait_names)
trait_names <- gsub('DOY_last_budset', 'Stage8_Final_Budset_DOY', trait_names)
trait_names <- gsub('growing_season_days', 'Growing_Season_Days', trait_names)

trait_names <- gsub('licor_', '', trait_names)
trait_names <- gsub('Fm.', "Fm'", trait_names)
trait_names <- gsub('upper', "Adaxial", trait_names)
trait_names <- gsub('lower', "Abaxial", trait_names)
trait_names <- gsub('stomata_ratio', "Stomata_ratio", trait_names)
trait_names <- gsub('leaf', 'Leaf', trait_names)
trait_names <- gsub('Phi', 'Φ', trait_names)

# remove units
trait_names <- gsub('_cm2', '', trait_names)
trait_names <- gsub('_cm', '', trait_names)
trait_names <- gsub('_mm2', '', trait_names)
trait_names <- gsub('_m2', '', trait_names)
trait_names <- gsub('_mm', '', trait_names)
trait_names <- gsub('_um', '', trait_names)
trait_names <- gsub('_g', '', trait_names)

trait_names <- gsub('_', ' ', trait_names)

# check
cbind(colnames(res.lr$eff.std.sig), trait_names)
```

```
##                                                                     
##  [1,] "plasticity_rdpi_growing_season_days_2023"                    
##  [2,] "plasticity_rdpi_growing_season_days_2024"                    
##  [3,] "plasticity_rdpi_DOY_Stage2_2023"                             
##  [4,] "plasticity_rdpi_DOY_Stage2_2024"                             
##  [5,] "plasticity_rdpi_DOY_Stage3_2023"                             
##  [6,] "plasticity_rdpi_DOY_Stage3_2024"                             
##  [7,] "plasticity_rdpi_DOY_Stage6_2023"                             
##  [8,] "plasticity_rdpi_DOY_Stage6_2024"                             
##  [9,] "plasticity_rdpi_DOY_Stage7_2023"                             
## [10,] "plasticity_rdpi_DOY_Stage7_2024"                             
## [11,] "plasticity_rdpi_stage7_presence_2023"                        
## [12,] "plasticity_rdpi_stage7_presence_2024"                        
## [13,] "plasticity_rdpi_DOY_last_budset_2023"                        
## [14,] "plasticity_rdpi_DOY_last_budset_2024"                        
## [15,] "plasticity_rdpi_leaf_thickness_avg_mm_2023"                  
## [16,] "plasticity_rdpi_leaf_area_cm2_2023_log"                      
## [17,] "plasticity_rdpi_leaf_mass_g_2023_log"                        
## [18,] "plasticity_rdpi_LMA_g_m2_2023"                               
## [19,] "plasticity_rdpi_lower_stomata_pore_length_mean_um"           
## [20,] "plasticity_rdpi_upper_stomata_pore_length_mean_um"           
## [21,] "plasticity_rdpi_lower_stomata_density_mm2"                   
## [22,] "plasticity_rdpi_upper_stomata_density_mm2_log"               
## [23,] "plasticity_rdpi_upper_stomata_presence"                      
## [24,] "plasticity_rdpi_upper_stomata_density_over_total_density_log"
## [25,] "plasticity_rdpi_licor_gsw"                                   
## [26,] "plasticity_rdpi_licor_ETR"                                   
## [27,] "plasticity_rdpi_licor_Fm."                                   
## [28,] "plasticity_rdpi_licor_Fs"                                    
## [29,] "plasticity_rdpi_licor_gbw"                                   
## [30,] "plasticity_rdpi_licor_PhiPS2"                                
##       trait_names                                     
##  [1,] "Growing Season Days 2023"                      
##  [2,] "Growing Season Days 2024"                      
##  [3,] "Stage2 Bud Flush DOY 2023"                     
##  [4,] "Stage2 Bud Flush DOY 2024"                     
##  [5,] "Stage3 Leaf Emergence DOY 2023"                
##  [6,] "Stage3 Leaf Emergence DOY 2024"                
##  [7,] "Stage6 First Budset DOY 2023"                  
##  [8,] "Stage6 First Budset DOY 2024"                  
##  [9,] "Stage7 Lammas Growth DOY 2023"                 
## [10,] "Stage7 Lammas Growth DOY 2024"                 
## [11,] "Stage7 Lammas Growth presence 2023"            
## [12,] "Stage7 Lammas Growth presence 2024"            
## [13,] "Stage8 Final Budset DOY 2023"                  
## [14,] "Stage8 Final Budset DOY 2024"                  
## [15,] "Leaf thickness avg 2023"                       
## [16,] "Leaf area 2023 log"                            
## [17,] "Leaf mass 2023 log"                            
## [18,] "LMA 2023"                                      
## [19,] "Abaxial stomata pore length mean"              
## [20,] "Adaxial stomata pore length mean"              
## [21,] "Abaxial stomata density"                       
## [22,] "Adaxial stomata density log"                   
## [23,] "Adaxial stomata presence"                      
## [24,] "Adaxial stomata density over total density log"
## [25,] "gsw"                                           
## [26,] "ETR"                                           
## [27,] "Fm'"                                           
## [28,] "Fs"                                            
## [29,] "gbw"                                           
## [30,] "ΦPS2"
```

```
# make df to join them
trait_names_df <- cbind.data.frame(colnames(res.lr$eff.std.sig), trait_names)
rownames(trait_names_df) <- trait_names_df[,1]
######################

# plot significant relationships for each predictor (row)

# use matrix with significant effects to select variables
# rename matrix for simplicity
sig <- res.lr$eff.std.sig

# loop through predictors
for(p in 1:nrow(sig)){
  
  xvar <- rownames(sig)[p]
  
  # get significant values
  yvars <- colnames(sig)[! is.na(sig[p,])]
  
  # loop through traits
  plots <- list()
  for(t in 1:length(yvars)){
    
    yvar <- yvars[t]
    
    # get correlation and pval
    cortext <- round(sig[xvar, yvar], 2)
    ptext <- round(res.lr$pvals[xvar, yvar], 3)
    if(ptext == '0'){
      ptext <- '<0.001'
    } else{
      ptext <- paste0('= ', ptext)
    }
    
    
    # make main title pretty
    maintitle <- trait_names_df[yvar, 2]
    # wrap long variables
    maintitle <- paste(strwrap(maintitle, width = 22), collapse = '\n')
    
    # make subtitle
    subtitle <- paste('R = ', cortext, '\n', 'p ', ptext, sep = '')
    
    # make xlabel pretty
    xtitle <- paste(toupper(substr(xvar, 1, 1)), substr(xvar, 2, nchar(xvar)), sep="")
    xtitle <- gsub('_', ' ', xtitle)
    
    p1 <- ggplot(data = dat, aes_string(x = xvar, y = yvar, color = 'k2_tricho'))+
      geom_point(size = 5, na.rm = TRUE) +
      scale_color_gradient2(high = "darkolivegreen2", mid = "grey20", low = "dodgerblue2", midpoint = 0.5, name = 'P. trichocarpa\nancestry', guide = F) +
      geom_smooth(method = lm, na.rm = TRUE, col = 'black')+
      ggtitle(yvar) +
      ylab('Plasticity (RDPI)') +
      xlab(xtitle) +
      ggtitle(maintitle, sub = subtitle) #+
      # theme(plot.title = element_text(size = 16),
      #       plot.subtitle = element_text(size = 14))
  
    plots[[t]] <- p1
    names(plots)[t] <- yvar
    
  }
  
  # save set of plots with predictor name
  assign(paste0('plots.', xvar), plots)
  
  # add legend to plot panel so we don't need a legend for each individual plot
  # do this by extracting the legend from a simplified plot using cowplot
  # then add to the end of the list of plots
  leg_plot <- ggplot(data = dat, aes_string(x = xvar, y = yvar, color = 'k2_tricho'))+
    geom_point()+
      scale_color_gradient2(high = "darkolivegreen2", mid = "grey20", low = "dodgerblue2", midpoint = 0.5, name = 'P. trichocarpa\nancestry')
  
  legend <- cowplot::get_legend(leg_plot)
  plots[[t+1]] <- legend
  p2 <- grid.arrange(grobs = plots, ncol = 3)
  #plot(p2)
  
}
```

```
## Warning: `aes_string()` was deprecated in ggplot2 3.0.0.
## ℹ Please use tidy evaluation idioms with `aes()`.
## ℹ See also `vignette("ggplot2-in-packages")` for more information.
## This warning is displayed once every 8 hours.
## Call `lifecycle::last_lifecycle_warnings()` to see where this warning was
## generated.
```

```
## `geom_smooth()` using formula = 'y ~ x'
```

```
## Warning: The `guide` argument in `scale_*()` cannot be `FALSE`. This was deprecated in
## ggplot2 3.3.4.
## ℹ Please use "none" instead.
## This warning is displayed once every 8 hours.
## Call `lifecycle::last_lifecycle_warnings()` to see where this warning was
## generated.
```

```
## `geom_smooth()` using formula = 'y ~ x'
## `geom_smooth()` using formula = 'y ~ x'
## `geom_smooth()` using formula = 'y ~ x'
## `geom_smooth()` using formula = 'y ~ x'
```

```
## Warning: Removed 2 rows containing missing values or values outside the scale range
## (`geom_point()`).
```

```
## `geom_smooth()` using formula = 'y ~ x'
```

```
## `geom_smooth()` using formula = 'y ~ x'
## `geom_smooth()` using formula = 'y ~ x'
## `geom_smooth()` using formula = 'y ~ x'
```

```
## `geom_smooth()` using formula = 'y ~ x'
## `geom_smooth()` using formula = 'y ~ x'
## `geom_smooth()` using formula = 'y ~ x'
## `geom_smooth()` using formula = 'y ~ x'
```

```
## `geom_smooth()` using formula = 'y ~ x'
## `geom_smooth()` using formula = 'y ~ x'
```

```
## `geom_smooth()` using formula = 'y ~ x'
## `geom_smooth()` using formula = 'y ~ x'
## `geom_smooth()` using formula = 'y ~ x'
## `geom_smooth()` using formula = 'y ~ x'
```

```
# PC1 plots - for main text

leg_plot <- 
  ggplot(data = dat, aes_string(x = xvar, y = yvar, color = 'k2_tricho'))+
  geom_point(na.rm = TRUE)+
  scale_color_gradient2(high = "darkolivegreen2", mid = "grey20", low = "dodgerblue2", midpoint = 0.5, name = expression(paste('% ', italic("P. trichocarpa"))))

legend <- cowplot::get_legend(leg_plot)
plots.genetic_PC1[[length(plots.genetic_PC1)+1]] <- legend
p2 <- grid.arrange(grobs = plots.genetic_PC1, ncol = 3)
```

```
## `geom_smooth()` using formula = 'y ~ x'
## `geom_smooth()` using formula = 'y ~ x'
## `geom_smooth()` using formula = 'y ~ x'
## `geom_smooth()` using formula = 'y ~ x'
## `geom_smooth()` using formula = 'y ~ x'
```

```
plot(p2)
#ggsave(p2, file = 'results/plasticity/plasticityRDPI_by_geneticPC1_significant_bigText.png', height = 8, width = 12)


# all other plots together for supplement
p3 <- grid.arrange(grobs = c(plots.genetic_PC2, plots.genetic_PC3, plots.provenance_MAP, plots.provenance_MAT, plots.provenance_TD, legend$grobs[1]), ncol = 4)
```

```
## `geom_smooth()` using formula = 'y ~ x'
## `geom_smooth()` using formula = 'y ~ x'
## `geom_smooth()` using formula = 'y ~ x'
## `geom_smooth()` using formula = 'y ~ x'
## `geom_smooth()` using formula = 'y ~ x'
## `geom_smooth()` using formula = 'y ~ x'
## `geom_smooth()` using formula = 'y ~ x'
## `geom_smooth()` using formula = 'y ~ x'
## `geom_smooth()` using formula = 'y ~ x'
## `geom_smooth()` using formula = 'y ~ x'
## `geom_smooth()` using formula = 'y ~ x'
## `geom_smooth()` using formula = 'y ~ x'
## `geom_smooth()` using formula = 'y ~ x'
## `geom_smooth()` using formula = 'y ~ x'
```

```
plot(p3)
#ggsave(p3, file = 'results/plasticity/plasticityRDPI_by_allExceptAncestry_significant_bigText.png', height = 16, width = 16)
```

#### 3.4 Heatmap of home-climate relationships

```
# heatmap params
text_cex <- 1
rot <- 90 # angle for column labels

leg_title <- 14 # font size for legend title
leg_label <- 12 # font size for legend labels

########################################################

# Heatmap of correlations
# colors for heatmap
colf <- colorRampPalette(c('red', 'white', 'blue'))

colnames(res.lr$cors.sig) <- gsub('plasticity_rdpi_', '', colnames(res.lr$cors.sig))
colnames(res.lr$padj) <- gsub('plasticity_rdpi_', '', colnames(res.lr$padj))

# trait categories

# row annotation - trait categories
cats <- c(rep('Phenology', 14),  rep('Leaf Morphology', 4), rep('Stomata', 7), rep('Photosynthesis', 5))
cbind(colnames(res.lr$cors.sig), cats)
```

```
##                                                      cats             
##  [1,] "growing_season_days_2023"                     "Phenology"      
##  [2,] "growing_season_days_2024"                     "Phenology"      
##  [3,] "DOY_Stage2_2023"                              "Phenology"      
##  [4,] "DOY_Stage2_2024"                              "Phenology"      
##  [5,] "DOY_Stage3_2023"                              "Phenology"      
##  [6,] "DOY_Stage3_2024"                              "Phenology"      
##  [7,] "DOY_Stage6_2023"                              "Phenology"      
##  [8,] "DOY_Stage6_2024"                              "Phenology"      
##  [9,] "DOY_Stage7_2023"                              "Phenology"      
## [10,] "DOY_Stage7_2024"                              "Phenology"      
## [11,] "stage7_presence_2023"                         "Phenology"      
## [12,] "stage7_presence_2024"                         "Phenology"      
## [13,] "DOY_last_budset_2023"                         "Phenology"      
## [14,] "DOY_last_budset_2024"                         "Phenology"      
## [15,] "leaf_thickness_avg_mm_2023"                   "Leaf Morphology"
## [16,] "leaf_area_cm2_2023_log"                       "Leaf Morphology"
## [17,] "leaf_mass_g_2023_log"                         "Leaf Morphology"
## [18,] "LMA_g_m2_2023"                                "Leaf Morphology"
## [19,] "lower_stomata_pore_length_mean_um"            "Stomata"        
## [20,] "upper_stomata_pore_length_mean_um"            "Stomata"        
## [21,] "lower_stomata_density_mm2"                    "Stomata"        
## [22,] "upper_stomata_density_mm2_log"                "Stomata"        
## [23,] "upper_stomata_presence"                       "Stomata"        
## [24,] "upper_stomata_density_over_total_density_log" "Stomata"        
## [25,] "licor_gsw"                                    "Stomata"        
## [26,] "licor_ETR"                                    "Photosynthesis" 
## [27,] "licor_Fm."                                    "Photosynthesis" 
## [28,] "licor_Fs"                                     "Photosynthesis" 
## [29,] "licor_gbw"                                    "Photosynthesis" 
## [30,] "licor_PhiPS2"                                 "Photosynthesis"
```

```
ha_row <- rowAnnotation(Trait_Category = cats,
                        show_annotation_name = F,
                            col = list(Trait_Category = c('Phenology' = "#e2e260", 'Photosynthesis' = "#2d4030", 'Leaf Morphology' = "#516823", 'Stomata' =  "#88a2b9")))

# cell text showing significance
# cell_text <- res.lr$pvals
# cell_text[res.lr$pvals > 0.05] <- ''
# cell_text[res.lr$pvals <= 0.05 & res.lr$pvals > 0.01] <- '*'
# cell_text[res.lr$pvals <= 0.01 & res.lr$pvals > 0.001] <- '**'
# cell_text[res.lr$pvals<= 0.001] <- '***'
# cell_text <- t(cell_text)

# or just run for adjusted pvals

cell_text <- res.lr$padj
cell_text[res.lr$padj > 0.05] <- ''
cell_text[res.lr$padj <= 0.05 & res.lr$padj > 0.01] <- '*'
cell_text[res.lr$padj <= 0.01 & res.lr$padj > 0.001] <- '**'
cell_text[res.lr$padj<= 0.001] <- '***'
cell_text <- t(cell_text)

############################
# heatmap by standardized effect size

# color scale based on all effect sizes
max_col <- max(unlist(res.lr$eff.std.sig), na.rm = T)
min_col <- min(unlist(res.lr$eff.std.sig), na.rm = T)
colf <- colorRamp2(c('red', 'white', 'blue'), breaks = c(min_col, 0, max_col))


hm1 <- Heatmap(as.matrix(t(res.lr$eff.std.sig)),
        cluster_rows = F,
        cluster_columns = F,
        col = colf,
        row_names_side = 'left',
        column_names_side = 'top',
        column_names_rot = 90,
        column_names_centered = T,
        column_title = 'Predictors',
        row_title = 'Trait plasticity',
        left_annotation = ha_row,
        heatmap_legend_param = list(title = 'Std. Effect Size'),
        cell_fun = function(j, i, x, y, width, height, fill) { # add text to each grid
                      grid.text(cell_text[i, j], x, y, gp = gpar(cex = 1.5))})

ComplexHeatmap::draw(hm1, padding = unit(c(2, 30, 2, 15), "mm"))
```

### 4 Relationship between plasticity and home climate, adjusted for species ancestry

```
##################################
# models estimating climate effects while controlling for ancestry

# possible predictors of plasticity
preds <- c('provenance_MAT', 'provenance_MAP', 'provenance_TD')

# set up dataframes for saving info within a list
res.lr.pt <- list() # results for linear regressions
# setup template dataframe
df <- as.data.frame(matrix(ncol = length(traits), nrow = length(preds)))
colnames(df) <- traits
rownames(df) <- preds

# add duplicate dataframes to list for saving effect size and pvalues
res.lr.pt$eff <- df
res.lr.pt$eff.sig <- df
res.lr.pt$eff.std <- df
res.lr.pt$eff.std.sig <- df
res.lr.pt$pvals <- df
res.lr.pt$rsq <- df
res.lr.pt$cors <- df
res.lr.pt$cors.sig <- df

# loop through each trait/predictor combination, run a regression, and extract info from model


for(t in 1:length(traits)){
  
  trait <- traits[t]
  
  for(p in 1:length(preds)){
    
    pred <- preds[p]
    
    # model
    form <- paste0(trait, '~', pred, '+ k2_tricho')
    mod <- lm(form, data = dat)
    mod.info <- summary(mod)
    print(paste('Model:', trait, '~', pred))
    print(mod.info)
    
    # save model info
    res.lr.pt$rsq[pred, trait] <- mod.info$adj.r.squared
    res.lr.pt$pvals[pred, trait] <- mod.info$coefficients[pred, 'Pr(>|t|)']
    res.lr.pt$eff[pred, trait] <- mod.info$coefficients[pred, 'Estimate']
    
    # standardize coefficients
    # coef for predictor is second object after intercept
    res.lr.pt$eff.std[pred,trait] <- lm.beta(mod)$standardized.coefficients[2] 
    print(lm.beta(mod))
    
    # also do a correlation test - used to visualize 
    cor <- cor.test(dat[,pred], dat[,trait])
    res.lr.pt$cors[pred, trait] <- cor$estimate
    
    if(res.lr.pt$pvals[pred, trait] <= 0.05){
      res.lr.pt$eff.sig[pred, trait] <- mod.info$coefficients[pred, 'Estimate']
      res.lr.pt$cors.sig[pred, trait] <- cor$estimate
      res.lr.pt$eff.std.sig[pred,trait] <- lm.beta(mod)$standardized.coefficients[2]
    }
  }
}
```

```
## [1] "Model: plasticity_rdpi_growing_season_days_2023 ~ provenance_MAT"
## 
## Call:
## lm(formula = form, data = dat)
## 
## Residuals:
##      Min       1Q   Median       3Q      Max 
## -0.14822 -0.10107 -0.04840  0.01581  0.75878 
## 
## Coefficients:
##                 Estimate Std. Error t value Pr(>|t|)    
## (Intercept)     0.334863   0.063504   5.273 4.64e-06 ***
## provenance_MAT  0.005222   0.014403   0.363    0.719    
## k2_tricho      -0.119511   0.112684  -1.061    0.295    
## ---
## Signif. codes:  0 '***' 0.001 '**' 0.01 '*' 0.05 '.' 0.1 ' ' 1
## 
## Residual standard error: 0.1863 on 41 degrees of freedom
## Multiple R-squared:  0.03137,    Adjusted R-squared:  -0.01588 
## F-statistic: 0.664 on 2 and 41 DF,  p-value: 0.5202
## 
## 
## Call:
## lm(formula = form, data = dat)
## 
## Standardized Coefficients::
##    (Intercept) provenance_MAT      k2_tricho 
##             NA     0.07424151    -0.21718770 
## 
## [1] "Model: plasticity_rdpi_growing_season_days_2023 ~ provenance_MAP"
## 
## Call:
## lm(formula = form, data = dat)
## 
## Residuals:
##      Min       1Q   Median       3Q      Max 
## -0.17617 -0.10160 -0.04226  0.02492  0.76178 
## 
## Coefficients:
##                  Estimate Std. Error t value Pr(>|t|)   
## (Intercept)     0.2671325  0.0874721   3.054  0.00396 **
## provenance_MAP  0.0001313  0.0001298   1.012  0.31758   
## k2_tricho      -0.1136066  0.0862329  -1.317  0.19501   
## ---
## Signif. codes:  0 '***' 0.001 '**' 0.01 '*' 0.05 '.' 0.1 ' ' 1
## 
## Residual standard error: 0.1844 on 41 degrees of freedom
## Multiple R-squared:  0.05194,    Adjusted R-squared:  0.005692 
## F-statistic: 1.123 on 2 and 41 DF,  p-value: 0.3351
## 
## 
## Call:
## lm(formula = form, data = dat)
## 
## Standardized Coefficients::
##    (Intercept) provenance_MAP      k2_tricho 
##             NA      0.1585564     -0.2064578 
## 
## [1] "Model: plasticity_rdpi_growing_season_days_2023 ~ provenance_TD"
## 
## Call:
## lm(formula = form, data = dat)
## 
## Residuals:
##      Min       1Q   Median       3Q      Max 
## -0.15023 -0.09313 -0.04944  0.01894  0.76152 
## 
## Coefficients:
##                Estimate Std. Error t value Pr(>|t|)
## (Intercept)    0.421630   0.298340   1.413    0.165
## provenance_TD -0.003026   0.009646  -0.314    0.755
## k2_tricho     -0.113644   0.108148  -1.051    0.299
## 
## Residual standard error: 0.1864 on 41 degrees of freedom
## Multiple R-squared:  0.03059,    Adjusted R-squared:  -0.01669 
## F-statistic: 0.647 on 2 and 41 DF,  p-value: 0.5289
## 
## 
## Call:
## lm(formula = form, data = dat)
## 
## Standardized Coefficients::
##   (Intercept) provenance_TD     k2_tricho 
##            NA   -0.06164977   -0.20652609 
## 
## [1] "Model: plasticity_rdpi_growing_season_days_2024 ~ provenance_MAT"
## 
## Call:
## lm(formula = form, data = dat)
## 
## Residuals:
##      Min       1Q   Median       3Q      Max 
## -0.10408 -0.04294 -0.02034  0.04357  0.14389 
## 
## Coefficients:
##                  Estimate Std. Error t value Pr(>|t|)    
## (Intercept)     0.2361008  0.0245860   9.603 7.96e-12 ***
## provenance_MAT  0.0002056  0.0054776   0.038   0.9702    
## k2_tricho      -0.0744375  0.0417767  -1.782   0.0826 .  
## ---
## Signif. codes:  0 '***' 0.001 '**' 0.01 '*' 0.05 '.' 0.1 ' ' 1
## 
## Residual standard error: 0.06835 on 39 degrees of freedom
##   (2 observations deleted due to missingness)
## Multiple R-squared:  0.115,  Adjusted R-squared:  0.06957 
## F-statistic: 2.533 on 2 and 39 DF,  p-value: 0.09243
## 
## 
## Call:
## lm(formula = form, data = dat)
## 
## Standardized Coefficients::
##    (Intercept) provenance_MAT      k2_tricho 
##             NA     0.00723708    -0.34352395 
## 
## [1] "Model: plasticity_rdpi_growing_season_days_2024 ~ provenance_MAP"
## 
## Call:
## lm(formula = form, data = dat)
## 
## Residuals:
##      Min       1Q   Median       3Q      Max 
## -0.09661 -0.04367 -0.02035  0.02183  0.16780 
## 
## Coefficients:
##                  Estimate Std. Error t value Pr(>|t|)    
## (Intercept)     1.926e-01  3.289e-02   5.856  8.2e-07 ***
## provenance_MAP  8.699e-05  4.638e-05   1.876   0.0682 .  
## k2_tricho      -8.558e-02  3.192e-02  -2.681   0.0107 *  
## ---
## Signif. codes:  0 '***' 0.001 '**' 0.01 '*' 0.05 '.' 0.1 ' ' 1
## 
## Residual standard error: 0.06546 on 39 degrees of freedom
##   (2 observations deleted due to missingness)
## Multiple R-squared:  0.1882, Adjusted R-squared:  0.1465 
## F-statistic: 4.519 on 2 and 39 DF,  p-value: 0.01717
## 
## 
## Call:
## lm(formula = form, data = dat)
## 
## Standardized Coefficients::
##    (Intercept) provenance_MAP      k2_tricho 
##             NA      0.2763284     -0.3949567 
## 
## [1] "Model: plasticity_rdpi_growing_season_days_2024 ~ provenance_TD"
## 
## Call:
## lm(formula = form, data = dat)
## 
## Residuals:
##      Min       1Q   Median       3Q      Max 
## -0.09593 -0.04204 -0.02247  0.03848  0.14432 
## 
## Coefficients:
##                Estimate Std. Error t value Pr(>|t|)  
## (Intercept)    0.299899   0.111934   2.679   0.0107 *
## provenance_TD -0.002168   0.003706  -0.585   0.5620  
## k2_tricho     -0.086652   0.039562  -2.190   0.0345 *
## ---
## Signif. codes:  0 '***' 0.001 '**' 0.01 '*' 0.05 '.' 0.1 ' ' 1
## 
## Residual standard error: 0.06805 on 39 degrees of freedom
##   (2 observations deleted due to missingness)
## Multiple R-squared:  0.1226, Adjusted R-squared:  0.07763 
## F-statistic: 2.725 on 2 and 39 DF,  p-value: 0.07801
## 
## 
## Call:
## lm(formula = form, data = dat)
## 
## Standardized Coefficients::
##   (Intercept) provenance_TD     k2_tricho 
##            NA    -0.1067856    -0.3998933 
## 
## [1] "Model: plasticity_rdpi_DOY_Stage2_2023 ~ provenance_MAT"
## 
## Call:
## lm(formula = form, data = dat)
## 
## Residuals:
##      Min       1Q   Median       3Q      Max 
## -0.09552 -0.06023 -0.01601  0.01232  0.25000 
## 
## Coefficients:
##                 Estimate Std. Error t value Pr(>|t|)    
## (Intercept)     0.131609   0.027869   4.722 2.73e-05 ***
## provenance_MAT  0.001477   0.006321   0.234    0.816    
## k2_tricho      -0.012210   0.049452  -0.247    0.806    
## ---
## Signif. codes:  0 '***' 0.001 '**' 0.01 '*' 0.05 '.' 0.1 ' ' 1
## 
## Residual standard error: 0.08178 on 41 degrees of freedom
## Multiple R-squared:  0.001699,   Adjusted R-squared:  -0.047 
## F-statistic: 0.03488 on 2 and 41 DF,  p-value: 0.9657
## 
## 
## Call:
## lm(formula = form, data = dat)
## 
## Standardized Coefficients::
##    (Intercept) provenance_MAT      k2_tricho 
##             NA     0.04856225    -0.05132861 
## 
## [1] "Model: plasticity_rdpi_DOY_Stage2_2023 ~ provenance_MAP"
## 
## Call:
## lm(formula = form, data = dat)
## 
## Residuals:
##      Min       1Q   Median       3Q      Max 
## -0.10116 -0.04751 -0.01541  0.01539  0.24913 
## 
## Coefficients:
##                  Estimate Std. Error t value Pr(>|t|)    
## (Intercept)     1.561e-01  3.841e-02   4.063 0.000213 ***
## provenance_MAP -5.384e-05  5.701e-05  -0.944 0.350523    
## k2_tricho       4.068e-03  3.787e-02   0.107 0.914979    
## ---
## Signif. codes:  0 '***' 0.001 '**' 0.01 '*' 0.05 '.' 0.1 ' ' 1
## 
## Residual standard error: 0.08096 on 41 degrees of freedom
## Multiple R-squared:  0.02165,    Adjusted R-squared:  -0.02607 
## F-statistic: 0.4537 on 2 and 41 DF,  p-value: 0.6385
## 
## 
## Call:
## lm(formula = form, data = dat)
## 
## Standardized Coefficients::
##    (Intercept) provenance_MAP      k2_tricho 
##             NA    -0.15033587     0.01710085 
## 
## [1] "Model: plasticity_rdpi_DOY_Stage2_2023 ~ provenance_TD"
## 
## Call:
## lm(formula = form, data = dat)
## 
## Residuals:
##      Min       1Q   Median       3Q      Max 
## -0.09234 -0.05938 -0.01591  0.01512  0.26036 
## 
## Coefficients:
##                Estimate Std. Error t value Pr(>|t|)
## (Intercept)   -0.014765   0.128900  -0.115    0.909
## provenance_TD  0.004794   0.004168   1.150    0.257
## k2_tricho      0.028900   0.046726   0.619    0.540
## 
## Residual standard error: 0.08055 on 41 degrees of freedom
## Multiple R-squared:  0.03163,    Adjusted R-squared:  -0.01561 
## F-statistic: 0.6695 on 2 and 41 DF,  p-value: 0.5175
## 
## 
## Call:
## lm(formula = form, data = dat)
## 
## Standardized Coefficients::
##   (Intercept) provenance_TD     k2_tricho 
##            NA     0.2259683     0.1214948 
## 
## [1] "Model: plasticity_rdpi_DOY_Stage2_2024 ~ provenance_MAT"
## 
## Call:
## lm(formula = form, data = dat)
## 
## Residuals:
##      Min       1Q   Median       3Q      Max 
## -0.09255 -0.04861  0.01219  0.03726  0.10380 
## 
## Coefficients:
##                 Estimate Std. Error t value Pr(>|t|)    
## (Intercept)     0.124878   0.018084   6.905 2.54e-08 ***
## provenance_MAT  0.005769   0.004169   1.384    0.174    
## k2_tricho      -0.033455   0.032053  -1.044    0.303    
## ---
## Signif. codes:  0 '***' 0.001 '**' 0.01 '*' 0.05 '.' 0.1 ' ' 1
## 
## Residual standard error: 0.05284 on 40 degrees of freedom
##   (1 observation deleted due to missingness)
## Multiple R-squared:  0.04636,    Adjusted R-squared:  -0.00132 
## F-statistic: 0.9723 on 2 and 40 DF,  p-value: 0.387
## 
## 
## Call:
## lm(formula = form, data = dat)
## 
## Standardized Coefficients::
##    (Intercept) provenance_MAT      k2_tricho 
##             NA      0.2848110     -0.2148002 
## 
## [1] "Model: plasticity_rdpi_DOY_Stage2_2024 ~ provenance_MAP"
## 
## Call:
## lm(formula = form, data = dat)
## 
## Residuals:
##      Min       1Q   Median       3Q      Max 
## -0.08868 -0.04871  0.01286  0.04111  0.10012 
## 
## Coefficients:
##                  Estimate Std. Error t value Pr(>|t|)    
## (Intercept)     1.471e-01  2.523e-02   5.831 8.16e-07 ***
## provenance_MAP -5.553e-05  3.716e-05  -1.494    0.143    
## k2_tricho       4.609e-03  2.466e-02   0.187    0.853    
## ---
## Signif. codes:  0 '***' 0.001 '**' 0.01 '*' 0.05 '.' 0.1 ' ' 1
## 
## Residual standard error: 0.05264 on 40 degrees of freedom
##   (1 observation deleted due to missingness)
## Multiple R-squared:  0.05354,    Adjusted R-squared:  0.006218 
## F-statistic: 1.131 on 2 and 40 DF,  p-value: 0.3327
## 
## 
## Call:
## lm(formula = form, data = dat)
## 
## Standardized Coefficients::
##    (Intercept) provenance_MAP      k2_tricho 
##             NA    -0.23661245     0.02959282 
## 
## [1] "Model: plasticity_rdpi_DOY_Stage2_2024 ~ provenance_TD"
## 
## Call:
## lm(formula = form, data = dat)
## 
## Residuals:
##      Min       1Q   Median       3Q      Max 
## -0.09241 -0.05028  0.01353  0.03614  0.10830 
## 
## Coefficients:
##                Estimate Std. Error t value Pr(>|t|)  
## (Intercept)    0.169235   0.086292   1.961   0.0568 .
## provenance_TD -0.001622   0.002795  -0.580   0.5650  
## k2_tricho     -0.015373   0.031251  -0.492   0.6255  
## ---
## Signif. codes:  0 '***' 0.001 '**' 0.01 '*' 0.05 '.' 0.1 ' ' 1
## 
## Residual standard error: 0.05387 on 40 degrees of freedom
##   (1 observation deleted due to missingness)
## Multiple R-squared:  0.009042,   Adjusted R-squared:  -0.04051 
## F-statistic: 0.1825 on 2 and 40 DF,  p-value: 0.8339
## 
## 
## Call:
## lm(formula = form, data = dat)
## 
## Standardized Coefficients::
##   (Intercept) provenance_TD     k2_tricho 
##            NA   -0.11641803   -0.09870412 
## 
## [1] "Model: plasticity_rdpi_DOY_Stage3_2023 ~ provenance_MAT"
## 
## Call:
## lm(formula = form, data = dat)
## 
## Residuals:
##       Min        1Q    Median        3Q       Max 
## -0.086270 -0.032241 -0.008605  0.006711  0.222022 
## 
## Coefficients:
##                 Estimate Std. Error t value Pr(>|t|)    
## (Intercept)     0.123826   0.022008   5.626 1.47e-06 ***
## provenance_MAT -0.003202   0.004992  -0.641    0.525    
## k2_tricho       0.005778   0.039052   0.148    0.883    
## ---
## Signif. codes:  0 '***' 0.001 '**' 0.01 '*' 0.05 '.' 0.1 ' ' 1
## 
## Residual standard error: 0.06458 on 41 degrees of freedom
## Multiple R-squared:  0.01316,    Adjusted R-squared:  -0.03498 
## F-statistic: 0.2733 on 2 and 41 DF,  p-value: 0.7622
## 
## 
## Call:
## lm(formula = form, data = dat)
## 
## Standardized Coefficients::
##    (Intercept) provenance_MAT      k2_tricho 
##             NA    -0.13259476     0.03058353 
## 
## [1] "Model: plasticity_rdpi_DOY_Stage3_2023 ~ provenance_MAP"
## 
## Call:
## lm(formula = form, data = dat)
## 
## Residuals:
##       Min        1Q    Median        3Q       Max 
## -0.087938 -0.030884 -0.013906  0.002945  0.231100 
## 
## Coefficients:
##                  Estimate Std. Error t value Pr(>|t|)    
## (Intercept)     1.544e-01  3.019e-02   5.113 7.78e-06 ***
## provenance_MAP -5.765e-05  4.481e-05  -1.287    0.205    
## k2_tricho      -1.519e-03  2.976e-02  -0.051    0.960    
## ---
## Signif. codes:  0 '***' 0.001 '**' 0.01 '*' 0.05 '.' 0.1 ' ' 1
## 
## Residual standard error: 0.06363 on 41 degrees of freedom
## Multiple R-squared:  0.04193,    Adjusted R-squared:  -0.004804 
## F-statistic: 0.8972 on 2 and 41 DF,  p-value: 0.4156
## 
## 
## Call:
## lm(formula = form, data = dat)
## 
## Standardized Coefficients::
##    (Intercept) provenance_MAP      k2_tricho 
##             NA   -0.202677790   -0.008039746 
## 
## [1] "Model: plasticity_rdpi_DOY_Stage3_2023 ~ provenance_TD"
## 
## Call:
## lm(formula = form, data = dat)
## 
## Residuals:
##       Min        1Q    Median        3Q       Max 
## -0.080496 -0.034285 -0.011839  0.004564  0.237732 
## 
## Coefficients:
##               Estimate Std. Error t value Pr(>|t|)
## (Intercept)   0.029652   0.102705   0.289    0.774
## provenance_TD 0.003210   0.003321   0.967    0.339
## k2_tricho     0.011637   0.037230   0.313    0.756
## 
## Residual standard error: 0.06418 on 41 degrees of freedom
## Multiple R-squared:  0.02546,    Adjusted R-squared:  -0.02208 
## F-statistic: 0.5356 on 2 and 41 DF,  p-value: 0.5894
## 
## 
## Call:
## lm(formula = form, data = dat)
## 
## Standardized Coefficients::
##   (Intercept) provenance_TD     k2_tricho 
##            NA    0.19047264    0.06159478 
## 
## [1] "Model: plasticity_rdpi_DOY_Stage3_2024 ~ provenance_MAT"
## 
## Call:
## lm(formula = form, data = dat)
## 
## Residuals:
##       Min        1Q    Median        3Q       Max 
## -0.087636 -0.027775  0.003276  0.024622  0.100903 
## 
## Coefficients:
##                 Estimate Std. Error t value Pr(>|t|)    
## (Intercept)     0.138553   0.016622   8.335 2.79e-10 ***
## provenance_MAT  0.006548   0.003832   1.709   0.0952 .  
## k2_tricho      -0.052951   0.029463  -1.797   0.0799 .  
## ---
## Signif. codes:  0 '***' 0.001 '**' 0.01 '*' 0.05 '.' 0.1 ' ' 1
## 
## Residual standard error: 0.04857 on 40 degrees of freedom
##   (1 observation deleted due to missingness)
## Multiple R-squared:  0.08491,    Adjusted R-squared:  0.03915 
## F-statistic: 1.856 on 2 and 40 DF,  p-value: 0.1695
## 
## 
## Call:
## lm(formula = form, data = dat)
## 
## Standardized Coefficients::
##    (Intercept) provenance_MAT      k2_tricho 
##             NA      0.3445043     -0.3623165 
## 
## [1] "Model: plasticity_rdpi_DOY_Stage3_2024 ~ provenance_MAP"
## 
## Call:
## lm(formula = form, data = dat)
## 
## Residuals:
##       Min        1Q    Median        3Q       Max 
## -0.091435 -0.026133  0.006927  0.025451  0.096153 
## 
## Coefficients:
##                  Estimate Std. Error t value Pr(>|t|)    
## (Intercept)     1.744e-01  2.233e-02   7.810 1.44e-09 ***
## provenance_MAP -8.485e-05  3.288e-05  -2.580   0.0136 *  
## k2_tricho      -6.314e-03  2.182e-02  -0.289   0.7738    
## ---
## Signif. codes:  0 '***' 0.001 '**' 0.01 '*' 0.05 '.' 0.1 ' ' 1
## 
## Residual standard error: 0.04659 on 40 degrees of freedom
##   (1 observation deleted due to missingness)
## Multiple R-squared:  0.1582, Adjusted R-squared:  0.1161 
## F-statistic:  3.76 on 2 and 40 DF,  p-value: 0.0319
## 
## 
## Call:
## lm(formula = form, data = dat)
## 
## Standardized Coefficients::
##    (Intercept) provenance_MAP      k2_tricho 
##             NA    -0.38532238    -0.04320439 
## 
## [1] "Model: plasticity_rdpi_DOY_Stage3_2024 ~ provenance_TD"
## 
## Call:
## lm(formula = form, data = dat)
## 
## Residuals:
##       Min        1Q    Median        3Q       Max 
## -0.090034 -0.027687  0.005774  0.027050  0.113552 
## 
## Coefficients:
##                 Estimate Std. Error t value Pr(>|t|)
## (Intercept)    0.1140375  0.0805397   1.416    0.165
## provenance_TD  0.0006395  0.0026086   0.245    0.808
## k2_tricho     -0.0152289  0.0291682  -0.522    0.604
## 
## Residual standard error: 0.05028 on 40 degrees of freedom
##   (1 observation deleted due to missingness)
## Multiple R-squared:  0.01958,    Adjusted R-squared:  -0.02944 
## F-statistic: 0.3993 on 2 and 40 DF,  p-value: 0.6734
## 
## 
## Call:
## lm(formula = form, data = dat)
## 
## Standardized Coefficients::
##   (Intercept) provenance_TD     k2_tricho 
##            NA    0.04892566   -0.10420401 
## 
## [1] "Model: plasticity_rdpi_DOY_Stage6_2023 ~ provenance_MAT"
## 
## Call:
## lm(formula = form, data = dat)
## 
## Residuals:
##       Min        1Q    Median        3Q       Max 
## -0.029692 -0.011786  0.000377  0.009027  0.039179 
## 
## Coefficients:
##                 Estimate Std. Error t value Pr(>|t|)    
## (Intercept)     0.098256   0.005905  16.640   <2e-16 ***
## provenance_MAT -0.002006   0.001339  -1.498    0.142    
## k2_tricho      -0.011789   0.010478  -1.125    0.267    
## ---
## Signif. codes:  0 '***' 0.001 '**' 0.01 '*' 0.05 '.' 0.1 ' ' 1
## 
## Residual standard error: 0.01733 on 41 degrees of freedom
## Multiple R-squared:  0.199,  Adjusted R-squared:  0.1599 
## F-statistic: 5.091 on 2 and 41 DF,  p-value: 0.01059
## 
## 
## Call:
## lm(formula = form, data = dat)
## 
## Standardized Coefficients::
##    (Intercept) provenance_MAT      k2_tricho 
##             NA     -0.2789347     -0.2095372 
## 
## [1] "Model: plasticity_rdpi_DOY_Stage6_2023 ~ provenance_MAP"
## 
## Call:
## lm(formula = form, data = dat)
## 
## Residuals:
##       Min        1Q    Median        3Q       Max 
## -0.035555 -0.013405  0.001477  0.010339  0.043596 
## 
## Coefficients:
##                  Estimate Std. Error t value Pr(>|t|)    
## (Intercept)     1.004e-01  8.443e-03  11.892 7.17e-15 ***
## provenance_MAP -6.623e-07  1.253e-05  -0.053   0.9581    
## k2_tricho      -2.205e-02  8.323e-03  -2.650   0.0114 *  
## ---
## Signif. codes:  0 '***' 0.001 '**' 0.01 '*' 0.05 '.' 0.1 ' ' 1
## 
## Residual standard error: 0.01779 on 41 degrees of freedom
## Multiple R-squared:  0.1552, Adjusted R-squared:  0.114 
## F-statistic: 3.765 on 2 and 41 DF,  p-value: 0.03153
## 
## 
## Call:
## lm(formula = form, data = dat)
## 
## Standardized Coefficients::
##    (Intercept) provenance_MAP      k2_tricho 
##             NA   -0.007818833   -0.391962190 
## 
## [1] "Model: plasticity_rdpi_DOY_Stage6_2023 ~ provenance_TD"
## 
## Call:
## lm(formula = form, data = dat)
## 
## Residuals:
##       Min        1Q    Median        3Q       Max 
## -0.035018 -0.013313  0.001363  0.010227  0.043255 
## 
## Coefficients:
##                 Estimate Std. Error t value Pr(>|t|)   
## (Intercept)    0.0927473  0.0284541   3.260  0.00225 **
## provenance_TD  0.0002426  0.0009200   0.264  0.79335   
## k2_tricho     -0.0204658  0.0103146  -1.984  0.05396 . 
## ---
## Signif. codes:  0 '***' 0.001 '**' 0.01 '*' 0.05 '.' 0.1 ' ' 1
## 
## Residual standard error: 0.01778 on 41 degrees of freedom
## Multiple R-squared:  0.1565, Adjusted R-squared:  0.1154 
## F-statistic: 3.805 on 2 and 41 DF,  p-value: 0.03049
## 
## 
## Call:
## lm(formula = form, data = dat)
## 
## Standardized Coefficients::
##   (Intercept) provenance_TD     k2_tricho 
##            NA    0.04833853   -0.36374671 
## 
## [1] "Model: plasticity_rdpi_DOY_Stage6_2024 ~ provenance_MAT"
## 
## Call:
## lm(formula = form, data = dat)
## 
## Residuals:
##       Min        1Q    Median        3Q       Max 
## -0.044084 -0.014766 -0.008802  0.008844  0.090494 
## 
## Coefficients:
##                 Estimate Std. Error t value Pr(>|t|)    
## (Intercept)     0.058874   0.009393   6.268 1.98e-07 ***
## provenance_MAT -0.002833   0.002165  -1.309    0.198    
## k2_tricho       0.015575   0.016648   0.936    0.355    
## ---
## Signif. codes:  0 '***' 0.001 '**' 0.01 '*' 0.05 '.' 0.1 ' ' 1
## 
## Residual standard error: 0.02745 on 40 degrees of freedom
##   (1 observation deleted due to missingness)
## Multiple R-squared:  0.04126,    Adjusted R-squared:  -0.006679 
## F-statistic: 0.8607 on 2 and 40 DF,  p-value: 0.4306
## 
## 
## Call:
## lm(formula = form, data = dat)
## 
## Standardized Coefficients::
##    (Intercept) provenance_MAT      k2_tricho 
##             NA     -0.2700433      0.1930561 
## 
## [1] "Model: plasticity_rdpi_DOY_Stage6_2024 ~ provenance_MAP"
## 
## Call:
## lm(formula = form, data = dat)
## 
## Residuals:
##       Min        1Q    Median        3Q       Max 
## -0.042971 -0.016142 -0.007601  0.002856  0.095792 
## 
## Coefficients:
##                  Estimate Std. Error t value Pr(>|t|)    
## (Intercept)     4.976e-02  1.320e-02   3.771 0.000526 ***
## provenance_MAP  2.350e-05  1.943e-05   1.210 0.233504    
## k2_tricho      -2.527e-03  1.290e-02  -0.196 0.845652    
## ---
## Signif. codes:  0 '***' 0.001 '**' 0.01 '*' 0.05 '.' 0.1 ' ' 1
## 
## Residual standard error: 0.02753 on 40 degrees of freedom
##   (1 observation deleted due to missingness)
## Multiple R-squared:  0.0355, Adjusted R-squared:  -0.01273 
## F-statistic: 0.736 on 2 and 40 DF,  p-value: 0.4854
## 
## 
## Call:
## lm(formula = form, data = dat)
## 
## Standardized Coefficients::
##    (Intercept) provenance_MAP      k2_tricho 
##             NA     0.19335282    -0.03131817 
## 
## [1] "Model: plasticity_rdpi_DOY_Stage6_2024 ~ provenance_TD"
## 
## Call:
## lm(formula = form, data = dat)
## 
## Residuals:
##       Min        1Q    Median        3Q       Max 
## -0.042007 -0.017285 -0.008334  0.008142  0.094421 
## 
## Coefficients:
##                Estimate Std. Error t value Pr(>|t|)  
## (Intercept)    0.105017   0.044333   2.369   0.0228 *
## provenance_TD -0.001454   0.001436  -1.013   0.3174  
## k2_tricho     -0.008912   0.016056  -0.555   0.5820  
## ---
## Signif. codes:  0 '***' 0.001 '**' 0.01 '*' 0.05 '.' 0.1 ' ' 1
## 
## Residual standard error: 0.02768 on 40 degrees of freedom
##   (1 observation deleted due to missingness)
## Multiple R-squared:  0.0252, Adjusted R-squared:  -0.02355 
## F-statistic: 0.5169 on 2 and 40 DF,  p-value: 0.6003
## 
## 
## Call:
## lm(formula = form, data = dat)
## 
## Standardized Coefficients::
##   (Intercept) provenance_TD     k2_tricho 
##            NA    -0.2015038    -0.1104599 
## 
## [1] "Model: plasticity_rdpi_DOY_Stage7_2023 ~ provenance_MAT"
## 
## Call:
## lm(formula = form, data = dat)
## 
## Residuals:
##       Min        1Q    Median        3Q       Max 
## -0.054577 -0.016526 -0.009045  0.013555  0.117026 
## 
## Coefficients:
##                 Estimate Std. Error t value Pr(>|t|)    
## (Intercept)     0.092784   0.013935   6.658 1.64e-07 ***
## provenance_MAT -0.003742   0.003028  -1.236    0.225    
## k2_tricho      -0.008622   0.021964  -0.393    0.697    
## ---
## Signif. codes:  0 '***' 0.001 '**' 0.01 '*' 0.05 '.' 0.1 ' ' 1
## 
## Residual standard error: 0.0337 on 32 degrees of freedom
##   (9 observations deleted due to missingness)
## Multiple R-squared:  0.0989, Adjusted R-squared:  0.04258 
## F-statistic: 1.756 on 2 and 32 DF,  p-value: 0.189
## 
## 
## Call:
## lm(formula = form, data = dat)
## 
## Standardized Coefficients::
##    (Intercept) provenance_MAT      k2_tricho 
##             NA    -0.25850202    -0.08210137 
## 
## [1] "Model: plasticity_rdpi_DOY_Stage7_2023 ~ provenance_MAP"
## 
## Call:
## lm(formula = form, data = dat)
## 
## Residuals:
##       Min        1Q    Median        3Q       Max 
## -0.065633 -0.017718 -0.009241  0.018213  0.114930 
## 
## Coefficients:
##                  Estimate Std. Error t value Pr(>|t|)    
## (Intercept)     7.120e-02  1.883e-02   3.780 0.000646 ***
## provenance_MAP  4.632e-05  2.784e-05   1.664 0.105850    
## k2_tricho      -3.195e-02  1.783e-02  -1.792 0.082571 .  
## ---
## Signif. codes:  0 '***' 0.001 '**' 0.01 '*' 0.05 '.' 0.1 ' ' 1
## 
## Residual standard error: 0.0331 on 32 degrees of freedom
##   (9 observations deleted due to missingness)
## Multiple R-squared:  0.1311, Adjusted R-squared:  0.07678 
## F-statistic: 2.414 on 2 and 32 DF,  p-value: 0.1056
## 
## 
## Call:
## lm(formula = form, data = dat)
## 
## Standardized Coefficients::
##    (Intercept) provenance_MAP      k2_tricho 
##             NA      0.2824816     -0.3042104 
## 
## [1] "Model: plasticity_rdpi_DOY_Stage7_2023 ~ provenance_TD"
## 
## Call:
## lm(formula = form, data = dat)
## 
## Residuals:
##       Min        1Q    Median        3Q       Max 
## -0.064268 -0.018909 -0.006425  0.015414  0.118453 
## 
## Coefficients:
##                Estimate Std. Error t value Pr(>|t|)  
## (Intercept)    0.158896   0.084203   1.887   0.0683 .
## provenance_TD -0.002302   0.002888  -0.797   0.4313  
## k2_tricho     -0.037214   0.023679  -1.572   0.1259  
## ---
## Signif. codes:  0 '***' 0.001 '**' 0.01 '*' 0.05 '.' 0.1 ' ' 1
## 
## Residual standard error: 0.03416 on 32 degrees of freedom
##   (9 observations deleted due to missingness)
## Multiple R-squared:  0.07426,    Adjusted R-squared:  0.0164 
## F-statistic: 1.283 on 2 and 32 DF,  p-value: 0.291
## 
## 
## Call:
## lm(formula = form, data = dat)
## 
## Standardized Coefficients::
##   (Intercept) provenance_TD     k2_tricho 
##            NA    -0.1797011    -0.3543718 
## 
## [1] "Model: plasticity_rdpi_DOY_Stage7_2024 ~ provenance_MAT"
## 
## Call:
## lm(formula = form, data = dat)
## 
## Residuals:
##       Min        1Q    Median        3Q       Max 
## -0.042607 -0.008463 -0.004831  0.002165  0.058779 
## 
## Coefficients:
##                 Estimate Std. Error t value Pr(>|t|)  
## (Intercept)     0.021009   0.046017   0.457   0.6545  
## provenance_MAT -0.008606   0.003473  -2.478   0.0256 *
## k2_tricho       0.080860   0.056172   1.440   0.1705  
## ---
## Signif. codes:  0 '***' 0.001 '**' 0.01 '*' 0.05 '.' 0.1 ' ' 1
## 
## Residual standard error: 0.02345 on 15 degrees of freedom
##   (26 observations deleted due to missingness)
## Multiple R-squared:  0.2921, Adjusted R-squared:  0.1977 
## F-statistic: 3.094 on 2 and 15 DF,  p-value: 0.07498
## 
## 
## Call:
## lm(formula = form, data = dat)
## 
## Standardized Coefficients::
##    (Intercept) provenance_MAT      k2_tricho 
##             NA     -0.6239377      0.3624293 
## 
## [1] "Model: plasticity_rdpi_DOY_Stage7_2024 ~ provenance_MAP"
## 
## Call:
## lm(formula = form, data = dat)
## 
## Residuals:
##       Min        1Q    Median        3Q       Max 
## -0.027369 -0.015543 -0.002758  0.004632  0.076429 
## 
## Coefficients:
##                  Estimate Std. Error t value Pr(>|t|)
## (Intercept)     5.640e-02  5.069e-02   1.113    0.283
## provenance_MAP  4.926e-05  3.751e-05   1.313    0.209
## k2_tricho      -2.920e-02  6.232e-02  -0.469    0.646
## 
## Residual standard error: 0.02637 on 15 degrees of freedom
##   (26 observations deleted due to missingness)
## Multiple R-squared:  0.1051, Adjusted R-squared:  -0.0142 
## F-statistic: 0.881 on 2 and 15 DF,  p-value: 0.4348
## 
## 
## Call:
## lm(formula = form, data = dat)
## 
## Standardized Coefficients::
##    (Intercept) provenance_MAP      k2_tricho 
##             NA      0.3668459     -0.1308963 
## 
## [1] "Model: plasticity_rdpi_DOY_Stage7_2024 ~ provenance_TD"
## 
## Call:
## lm(formula = form, data = dat)
## 
## Residuals:
##       Min        1Q    Median        3Q       Max 
## -0.028086 -0.011559 -0.005515  0.005996  0.074888 
## 
## Coefficients:
##                Estimate Std. Error t value Pr(>|t|)  
## (Intercept)    0.228397   0.122676   1.862   0.0823 .
## provenance_TD -0.006807   0.004241  -1.605   0.1294  
## k2_tricho     -0.009746   0.054636  -0.178   0.8608  
## ---
## Signif. codes:  0 '***' 0.001 '**' 0.01 '*' 0.05 '.' 0.1 ' ' 1
## 
## Residual standard error: 0.02572 on 15 degrees of freedom
##   (26 observations deleted due to missingness)
## Multiple R-squared:  0.1484, Adjusted R-squared:  0.03489 
## F-statistic: 1.307 on 2 and 15 DF,  p-value: 0.2997
## 
## 
## Call:
## lm(formula = form, data = dat)
## 
## Standardized Coefficients::
##   (Intercept) provenance_TD     k2_tricho 
##            NA   -0.39300812   -0.04368333 
## 
## [1] "Model: plasticity_rdpi_stage7_presence_2023 ~ provenance_MAT"
## 
## Call:
## lm(formula = form, data = dat)
## 
## Residuals:
##      Min       1Q   Median       3Q      Max 
## -0.59522  0.03003  0.03648  0.04681  0.07234 
## 
## Coefficients:
##                 Estimate Std. Error t value Pr(>|t|)    
## (Intercept)     0.940733   0.052576  17.893   <2e-16 ***
## provenance_MAT -0.005682   0.011925  -0.477    0.636    
## k2_tricho       0.050639   0.093293   0.543    0.590    
## ---
## Signif. codes:  0 '***' 0.001 '**' 0.01 '*' 0.05 '.' 0.1 ' ' 1
## 
## Residual standard error: 0.1543 on 41 degrees of freedom
## Multiple R-squared:  0.007727,   Adjusted R-squared:  -0.04068 
## F-statistic: 0.1596 on 2 and 41 DF,  p-value: 0.853
## 
## 
## Call:
## lm(formula = form, data = dat)
## 
## Standardized Coefficients::
##    (Intercept) provenance_MAT      k2_tricho 
##             NA    -0.09876316     0.11250332 
## 
## [1] "Model: plasticity_rdpi_stage7_presence_2023 ~ provenance_MAP"
## 
## Call:
## lm(formula = form, data = dat)
## 
## Residuals:
##      Min       1Q   Median       3Q      Max 
## -0.61653  0.03185  0.03745  0.04539  0.05942 
## 
## Coefficients:
##                  Estimate Std. Error t value Pr(>|t|)    
## (Intercept)     9.592e-01  7.335e-02  13.078 3.22e-16 ***
## provenance_MAP -2.772e-05  1.089e-04  -0.255    0.800    
## k2_tricho       2.571e-02  7.231e-02   0.356    0.724    
## ---
## Signif. codes:  0 '***' 0.001 '**' 0.01 '*' 0.05 '.' 0.1 ' ' 1
## 
## Residual standard error: 0.1546 on 41 degrees of freedom
## Multiple R-squared:  0.003807,   Adjusted R-squared:  -0.04479 
## F-statistic: 0.07834 on 2 and 41 DF,  p-value: 0.9248
## 
## 
## Call:
## lm(formula = form, data = dat)
## 
## Standardized Coefficients::
##    (Intercept) provenance_MAP      k2_tricho 
##             NA    -0.04090300     0.05712983 
## 
## [1] "Model: plasticity_rdpi_stage7_presence_2023 ~ provenance_TD"
## 
## Call:
## lm(formula = form, data = dat)
## 
## Residuals:
##      Min       1Q   Median       3Q      Max 
## -0.58510  0.02499  0.03499  0.04296  0.10840 
## 
## Coefficients:
##               Estimate Std. Error t value Pr(>|t|)   
## (Intercept)   0.689187   0.244166   2.823  0.00732 **
## provenance_TD 0.008487   0.007894   1.075  0.28864   
## k2_tricho     0.080525   0.088510   0.910  0.36825   
## ---
## Signif. codes:  0 '***' 0.001 '**' 0.01 '*' 0.05 '.' 0.1 ' ' 1
## 
## Residual standard error: 0.1526 on 41 degrees of freedom
## Multiple R-squared:  0.02959,    Adjusted R-squared:  -0.01775 
## F-statistic: 0.625 on 2 and 41 DF,  p-value: 0.5403
## 
## 
## Call:
## lm(formula = form, data = dat)
## 
## Standardized Coefficients::
##   (Intercept) provenance_TD     k2_tricho 
##            NA     0.2113956     0.1789008 
## 
## [1] "Model: plasticity_rdpi_stage7_presence_2024 ~ provenance_MAT"
## 
## Call:
## lm(formula = form, data = dat)
## 
## Residuals:
##      Min       1Q   Median       3Q      Max 
## -0.38527 -0.18347  0.00386  0.10048  0.44898 
## 
## Coefficients:
##                Estimate Std. Error t value Pr(>|t|)    
## (Intercept)     1.03511    0.07638  13.552   <2e-16 ***
## provenance_MAT -0.02861    0.01761  -1.625   0.1120    
## k2_tricho      -0.30384    0.13538  -2.244   0.0304 *  
## ---
## Signif. codes:  0 '***' 0.001 '**' 0.01 '*' 0.05 '.' 0.1 ' ' 1
## 
## Residual standard error: 0.2232 on 40 degrees of freedom
##   (1 observation deleted due to missingness)
## Multiple R-squared:  0.357,  Adjusted R-squared:  0.3248 
## F-statistic:  11.1 on 2 and 40 DF,  p-value: 0.000146
## 
## 
## Call:
## lm(formula = form, data = dat)
## 
## Standardized Coefficients::
##    (Intercept) provenance_MAT      k2_tricho 
##             NA     -0.2746109     -0.3792781 
## 
## [1] "Model: plasticity_rdpi_stage7_presence_2024 ~ provenance_MAP"
## 
## Call:
## lm(formula = form, data = dat)
## 
## Residuals:
##      Min       1Q   Median       3Q      Max 
## -0.48332 -0.15607  0.01104  0.11408  0.48210 
## 
## Coefficients:
##                  Estimate Std. Error t value Pr(>|t|)    
## (Intercept)     0.8972086  0.1045425   8.582 1.31e-10 ***
## provenance_MAP  0.0003322  0.0001539   2.158    0.037 *  
## k2_tricho      -0.5015498  0.1021666  -4.909 1.58e-05 ***
## ---
## Signif. codes:  0 '***' 0.001 '**' 0.01 '*' 0.05 '.' 0.1 ' ' 1
## 
## Residual standard error: 0.2181 on 40 degrees of freedom
##   (1 observation deleted due to missingness)
## Multiple R-squared:  0.386,  Adjusted R-squared:  0.3553 
## F-statistic: 12.57 on 2 and 40 DF,  p-value: 5.8e-05
## 
## 
## Call:
## lm(formula = form, data = dat)
## 
## Standardized Coefficients::
##    (Intercept) provenance_MAP      k2_tricho 
##             NA      0.2751622     -0.6260660 
## 
## [1] "Model: plasticity_rdpi_stage7_presence_2024 ~ provenance_TD"
## 
## Call:
## lm(formula = form, data = dat)
## 
## Residuals:
##      Min       1Q   Median       3Q      Max 
## -0.46614 -0.17463  0.00623  0.12842  0.39164 
## 
## Coefficients:
##                Estimate Std. Error t value Pr(>|t|)   
## (Intercept)    1.105999   0.369071   2.997  0.00467 **
## provenance_TD -0.001594   0.011954  -0.133  0.89460   
## k2_tricho     -0.460346   0.133662  -3.444  0.00136 **
## ---
## Signif. codes:  0 '***' 0.001 '**' 0.01 '*' 0.05 '.' 0.1 ' ' 1
## 
## Residual standard error: 0.2304 on 40 degrees of freedom
##   (1 observation deleted due to missingness)
## Multiple R-squared:  0.3148, Adjusted R-squared:  0.2806 
## F-statistic:  9.19 on 2 and 40 DF,  p-value: 0.0005198
## 
## 
## Call:
## lm(formula = form, data = dat)
## 
## Standardized Coefficients::
##   (Intercept) provenance_TD     k2_tricho 
##            NA   -0.02224554   -0.57463323 
## 
## [1] "Model: plasticity_rdpi_DOY_last_budset_2023 ~ provenance_MAT"
## 
## Call:
## lm(formula = form, data = dat)
## 
## Residuals:
##      Min       1Q   Median       3Q      Max 
## -0.04247 -0.02185 -0.01094  0.01892  0.08053 
## 
## Coefficients:
##                 Estimate Std. Error t value Pr(>|t|)    
## (Intercept)     0.094190   0.011735   8.026 6.11e-10 ***
## provenance_MAT -0.006658   0.002662  -2.502   0.0165 *  
## k2_tricho       0.031722   0.020824   1.523   0.1353    
## ---
## Signif. codes:  0 '***' 0.001 '**' 0.01 '*' 0.05 '.' 0.1 ' ' 1
## 
## Residual standard error: 0.03444 on 41 degrees of freedom
## Multiple R-squared:  0.133,  Adjusted R-squared:  0.09067 
## F-statistic: 3.144 on 2 and 41 DF,  p-value: 0.05368
## 
## 
## Call:
## lm(formula = form, data = dat)
## 
## Standardized Coefficients::
##    (Intercept) provenance_MAT      k2_tricho 
##             NA     -0.4846574      0.2951449 
## 
## [1] "Model: plasticity_rdpi_DOY_last_budset_2023 ~ provenance_MAP"
## 
## Call:
## lm(formula = form, data = dat)
## 
## Residuals:
##       Min        1Q    Median        3Q       Max 
## -0.041341 -0.029643 -0.007799  0.010462  0.094302 
## 
## Coefficients:
##                  Estimate Std. Error t value Pr(>|t|)    
## (Intercept)     9.834e-02  1.754e-02   5.608 1.56e-06 ***
## provenance_MAP  4.010e-06  2.603e-05   0.154    0.878    
## k2_tricho      -3.342e-03  1.729e-02  -0.193    0.848    
## ---
## Signif. codes:  0 '***' 0.001 '**' 0.01 '*' 0.05 '.' 0.1 ' ' 1
## 
## Residual standard error: 0.03696 on 41 degrees of freedom
## Multiple R-squared:  0.001209,   Adjusted R-squared:  -0.04751 
## F-statistic: 0.02481 on 2 and 41 DF,  p-value: 0.9755
## 
## 
## Call:
## lm(formula = form, data = dat)
## 
## Standardized Coefficients::
##    (Intercept) provenance_MAP      k2_tricho 
##             NA     0.02478112    -0.03109813 
## 
## [1] "Model: plasticity_rdpi_DOY_last_budset_2023 ~ provenance_TD"
## 
## Call:
## lm(formula = form, data = dat)
## 
## Residuals:
##       Min        1Q    Median        3Q       Max 
## -0.047324 -0.024433 -0.009568  0.017656  0.101709 
## 
## Coefficients:
##               Estimate Std. Error t value Pr(>|t|)
## (Intercept)    0.02222    0.05784   0.384    0.703
## provenance_TD  0.00258    0.00187   1.380    0.175
## k2_tricho      0.01532    0.02097   0.731    0.469
## 
## Residual standard error: 0.03614 on 41 degrees of freedom
## Multiple R-squared:  0.04496,    Adjusted R-squared:  -0.001623 
## F-statistic: 0.9652 on 2 and 41 DF,  p-value: 0.3894
## 
## 
## Call:
## lm(formula = form, data = dat)
## 
## Standardized Coefficients::
##   (Intercept) provenance_TD     k2_tricho 
##            NA     0.2691222     0.1425030 
## 
## [1] "Model: plasticity_rdpi_DOY_last_budset_2024 ~ provenance_MAT"
## 
## Call:
## lm(formula = form, data = dat)
## 
## Residuals:
##       Min        1Q    Median        3Q       Max 
## -0.035275 -0.016091 -0.004084  0.011592  0.078620 
## 
## Coefficients:
##                 Estimate Std. Error t value Pr(>|t|)    
## (Intercept)     0.095524   0.008859  10.782 2.11e-13 ***
## provenance_MAT -0.001640   0.002042  -0.803    0.427    
## k2_tricho      -0.013268   0.015703  -0.845    0.403    
## ---
## Signif. codes:  0 '***' 0.001 '**' 0.01 '*' 0.05 '.' 0.1 ' ' 1
## 
## Residual standard error: 0.02589 on 40 degrees of freedom
##   (1 observation deleted due to missingness)
## Multiple R-squared:  0.09109,    Adjusted R-squared:  0.04564 
## F-statistic: 2.004 on 2 and 40 DF,  p-value: 0.1481
## 
## 
## Call:
## lm(formula = form, data = dat)
## 
## Standardized Coefficients::
##    (Intercept) provenance_MAT      k2_tricho 
##             NA     -0.1613758     -0.1697609 
## 
## [1] "Model: plasticity_rdpi_DOY_last_budset_2024 ~ provenance_MAP"
## 
## Call:
## lm(formula = form, data = dat)
## 
## Residuals:
##       Min        1Q    Median        3Q       Max 
## -0.037847 -0.018200 -0.004755  0.006973  0.069919 
## 
## Coefficients:
##                  Estimate Std. Error t value Pr(>|t|)    
## (Intercept)     8.161e-02  1.204e-02   6.776 3.85e-08 ***
## provenance_MAP  3.146e-05  1.774e-05   1.774   0.0837 .  
## k2_tricho      -2.656e-02  1.177e-02  -2.256   0.0296 *  
## ---
## Signif. codes:  0 '***' 0.001 '**' 0.01 '*' 0.05 '.' 0.1 ' ' 1
## 
## Residual standard error: 0.02513 on 40 degrees of freedom
##   (1 observation deleted due to missingness)
## Multiple R-squared:  0.1438, Adjusted R-squared:  0.101 
## F-statistic: 3.359 on 2 and 40 DF,  p-value: 0.04484
## 
## 
## Call:
## lm(formula = form, data = dat)
## 
## Standardized Coefficients::
##    (Intercept) provenance_MAP      k2_tricho 
##             NA      0.2671568     -0.3397861 
## 
## [1] "Model: plasticity_rdpi_DOY_last_budset_2024 ~ provenance_TD"
## 
## Call:
## lm(formula = form, data = dat)
## 
## Residuals:
##       Min        1Q    Median        3Q       Max 
## -0.039933 -0.014489 -0.003981  0.012678  0.081184 
## 
## Coefficients:
##                 Estimate Std. Error t value Pr(>|t|)  
## (Intercept)    0.0942403  0.0418020   2.254   0.0297 *
## provenance_TD  0.0000858  0.0013540   0.063   0.9498  
## k2_tricho     -0.0210115  0.0151390  -1.388   0.1728  
## ---
## Signif. codes:  0 '***' 0.001 '**' 0.01 '*' 0.05 '.' 0.1 ' ' 1
## 
## Residual standard error: 0.02609 on 40 degrees of freedom
##   (1 observation deleted due to missingness)
## Multiple R-squared:  0.07652,    Adjusted R-squared:  0.03035 
## F-statistic: 1.657 on 2 and 40 DF,  p-value: 0.2035
## 
## 
## Call:
## lm(formula = form, data = dat)
## 
## Standardized Coefficients::
##   (Intercept) provenance_TD     k2_tricho 
##            NA    0.01227478   -0.26884072 
## 
## [1] "Model: plasticity_rdpi_leaf_thickness_avg_mm_2023 ~ provenance_MAT"
## 
## Call:
## lm(formula = form, data = dat)
## 
## Residuals:
##      Min       1Q   Median       3Q      Max 
## -0.05056 -0.03213 -0.01375  0.01929  0.19828 
## 
## Coefficients:
##                Estimate Std. Error t value Pr(>|t|)    
## (Intercept)    0.077366   0.016219   4.770 2.34e-05 ***
## provenance_MAT 0.003040   0.003679   0.826    0.413    
## k2_tricho      0.016018   0.028780   0.557    0.581    
## ---
## Signif. codes:  0 '***' 0.001 '**' 0.01 '*' 0.05 '.' 0.1 ' ' 1
## 
## Residual standard error: 0.04759 on 41 degrees of freedom
## Multiple R-squared:  0.0648, Adjusted R-squared:  0.01918 
## F-statistic:  1.42 on 2 and 41 DF,  p-value: 0.2533
## 
## 
## Call:
## lm(formula = form, data = dat)
## 
## Standardized Coefficients::
##    (Intercept) provenance_MAT      k2_tricho 
##             NA      0.1662747      0.1119926 
## 
## [1] "Model: plasticity_rdpi_leaf_thickness_avg_mm_2023 ~ provenance_MAP"
## 
## Call:
## lm(formula = form, data = dat)
## 
## Residuals:
##      Min       1Q   Median       3Q      Max 
## -0.04800 -0.02861 -0.01173  0.01341  0.20816 
## 
## Coefficients:
##                  Estimate Std. Error t value Pr(>|t|)    
## (Intercept)     0.0856193  0.0226402   3.782 0.000498 ***
## provenance_MAP -0.0000230  0.0000336  -0.684 0.497521    
## k2_tricho       0.0354264  0.0223195   1.587 0.120142    
## ---
## Signif. codes:  0 '***' 0.001 '**' 0.01 '*' 0.05 '.' 0.1 ' ' 1
## 
## Residual standard error: 0.04772 on 41 degrees of freedom
## Multiple R-squared:  0.05997,    Adjusted R-squared:  0.01411 
## F-statistic: 1.308 on 2 and 41 DF,  p-value: 0.2815
## 
## 
## Call:
## lm(formula = form, data = dat)
## 
## Standardized Coefficients::
##    (Intercept) provenance_MAP      k2_tricho 
##             NA     -0.1068114      0.2476832 
## 
## [1] "Model: plasticity_rdpi_leaf_thickness_avg_mm_2023 ~ provenance_TD"
## 
## Call:
## lm(formula = form, data = dat)
## 
## Residuals:
##      Min       1Q   Median       3Q      Max 
## -0.05249 -0.03095 -0.01379  0.01577  0.20710 
## 
## Coefficients:
##                Estimate Std. Error t value Pr(>|t|)
## (Intercept)   0.0538448  0.0767275   0.702    0.487
## provenance_TD 0.0006859  0.0024808   0.276    0.784
## k2_tricho     0.0365229  0.0278136   1.313    0.196
## 
## Residual standard error: 0.04794 on 41 degrees of freedom
## Multiple R-squared:  0.05099,    Adjusted R-squared:  0.0047 
## F-statistic: 1.102 on 2 and 41 DF,  p-value: 0.342
## 
## 
## Call:
## lm(formula = form, data = dat)
## 
## Standardized Coefficients::
##   (Intercept) provenance_TD     k2_tricho 
##            NA    0.05376552    0.25534937 
## 
## [1] "Model: plasticity_rdpi_leaf_area_cm2_2023_log ~ provenance_MAT"
## 
## Call:
## lm(formula = form, data = dat)
## 
## Residuals:
##      Min       1Q   Median       3Q      Max 
## -0.09581 -0.04526 -0.01942  0.01973  0.33621 
## 
## Coefficients:
##                 Estimate Std. Error t value Pr(>|t|)    
## (Intercept)     0.150556   0.027750   5.425 2.83e-06 ***
## provenance_MAT -0.024420   0.006294  -3.880 0.000372 ***
## k2_tricho       0.112190   0.049242   2.278 0.027984 *  
## ---
## Signif. codes:  0 '***' 0.001 '**' 0.01 '*' 0.05 '.' 0.1 ' ' 1
## 
## Residual standard error: 0.08143 on 41 degrees of freedom
## Multiple R-squared:  0.2704, Adjusted R-squared:  0.2348 
## F-statistic: 7.599 on 2 and 41 DF,  p-value: 0.001559
## 
## 
## Call:
## lm(formula = form, data = dat)
## 
## Standardized Coefficients::
##    (Intercept) provenance_MAT      k2_tricho 
##             NA     -0.6895291      0.4049212 
## 
## [1] "Model: plasticity_rdpi_leaf_area_cm2_2023_log ~ provenance_MAP"
## 
## Call:
## lm(formula = form, data = dat)
## 
## Residuals:
##      Min       1Q   Median       3Q      Max 
## -0.10994 -0.04094 -0.02648  0.01737  0.40778 
## 
## Coefficients:
##                  Estimate Std. Error t value Pr(>|t|)    
## (Intercept)     1.674e-01  4.516e-02   3.707  0.00062 ***
## provenance_MAP  1.127e-05  6.702e-05   0.168  0.86732    
## k2_tricho      -1.586e-02  4.452e-02  -0.356  0.72351    
## ---
## Signif. codes:  0 '***' 0.001 '**' 0.01 '*' 0.05 '.' 0.1 ' ' 1
## 
## Residual standard error: 0.09518 on 41 degrees of freedom
## Multiple R-squared:  0.003258,   Adjusted R-squared:  -0.04536 
## F-statistic: 0.06701 on 2 and 41 DF,  p-value: 0.9353
## 
## 
## Call:
## lm(formula = form, data = dat)
## 
## Standardized Coefficients::
##    (Intercept) provenance_MAP      k2_tricho 
##             NA     0.02701350    -0.05723694 
## 
## [1] "Model: plasticity_rdpi_leaf_area_cm2_2023_log ~ provenance_TD"
## 
## Call:
## lm(formula = form, data = dat)
## 
## Residuals:
##      Min       1Q   Median       3Q      Max 
## -0.12333 -0.05149 -0.01693  0.03871  0.39522 
## 
## Coefficients:
##               Estimate Std. Error t value Pr(>|t|)  
## (Intercept)   -0.19289    0.14074  -1.371   0.1780  
## provenance_TD  0.01209    0.00455   2.657   0.0112 *
## k2_tricho      0.07037    0.05102   1.379   0.1753  
## ---
## Signif. codes:  0 '***' 0.001 '**' 0.01 '*' 0.05 '.' 0.1 ' ' 1
## 
## Residual standard error: 0.08794 on 41 degrees of freedom
## Multiple R-squared:  0.1491, Adjusted R-squared:  0.1076 
## F-statistic: 3.591 on 2 and 41 DF,  p-value: 0.03654
## 
## 
## Call:
## lm(formula = form, data = dat)
## 
## Standardized Coefficients::
##   (Intercept) provenance_TD     k2_tricho 
##            NA     0.4892213     0.2539828 
## 
## [1] "Model: plasticity_rdpi_leaf_mass_g_2023_log ~ provenance_MAT"
## 
## Call:
## lm(formula = form, data = dat)
## 
## Residuals:
##      Min       1Q   Median       3Q      Max 
## -0.47431 -0.03897  0.03975  0.11530  0.18065 
## 
## Coefficients:
##                Estimate Std. Error t value Pr(>|t|)    
## (Intercept)    -0.18913    0.05201  -3.636 0.000764 ***
## provenance_MAT  0.02692    0.01180   2.282 0.027752 *  
## k2_tricho      -0.28510    0.09229  -3.089 0.003597 ** 
## ---
## Signif. codes:  0 '***' 0.001 '**' 0.01 '*' 0.05 '.' 0.1 ' ' 1
## 
## Residual standard error: 0.1526 on 41 degrees of freedom
## Multiple R-squared:  0.1905, Adjusted R-squared:  0.151 
## F-statistic: 4.823 on 2 and 41 DF,  p-value: 0.01315
## 
## 
## Call:
## lm(formula = form, data = dat)
## 
## Standardized Coefficients::
##    (Intercept) provenance_MAT      k2_tricho 
##             NA      0.4272099     -0.5783238 
## 
## [1] "Model: plasticity_rdpi_leaf_mass_g_2023_log ~ provenance_MAP"
## 
## Call:
## lm(formula = form, data = dat)
## 
## Residuals:
##      Min       1Q   Median       3Q      Max 
## -0.47827 -0.05319  0.05388  0.08030  0.22030 
## 
## Coefficients:
##                  Estimate Std. Error t value Pr(>|t|)  
## (Intercept)    -0.1082751  0.0733102  -1.477   0.1473  
## provenance_MAP -0.0002199  0.0001088  -2.021   0.0498 *
## k2_tricho      -0.1106286  0.0722717  -1.531   0.1335  
## ---
## Signif. codes:  0 '***' 0.001 '**' 0.01 '*' 0.05 '.' 0.1 ' ' 1
## 
## Residual standard error: 0.1545 on 41 degrees of freedom
## Multiple R-squared:  0.1703, Adjusted R-squared:  0.1298 
## F-statistic: 4.207 on 2 and 41 DF,  p-value: 0.02178
## 
## 
## Call:
## lm(formula = form, data = dat)
## 
## Standardized Coefficients::
##    (Intercept) provenance_MAP      k2_tricho 
##             NA     -0.2962824     -0.2244115 
## 
## [1] "Model: plasticity_rdpi_leaf_mass_g_2023_log ~ provenance_TD"
## 
## Call:
## lm(formula = form, data = dat)
## 
## Residuals:
##      Min       1Q   Median       3Q      Max 
## -0.49998 -0.05023  0.05476  0.09584  0.19946 
## 
## Coefficients:
##                Estimate Std. Error t value Pr(>|t|)
## (Intercept)   -0.311346   0.258825  -1.203    0.236
## provenance_TD  0.003228   0.008368   0.386    0.702
## k2_tricho     -0.123392   0.093824  -1.315    0.196
## 
## Residual standard error: 0.1617 on 41 degrees of freedom
## Multiple R-squared:  0.09093,    Adjusted R-squared:  0.04659 
## F-statistic: 2.051 on 2 and 41 DF,  p-value: 0.1416
## 
## 
## Call:
## lm(formula = form, data = dat)
## 
## Standardized Coefficients::
##   (Intercept) provenance_TD     k2_tricho 
##            NA    0.07342324   -0.25030292 
## 
## [1] "Model: plasticity_rdpi_LMA_g_m2_2023 ~ provenance_MAT"
## 
## Call:
## lm(formula = form, data = dat)
## 
## Residuals:
##       Min        1Q    Median        3Q       Max 
## -0.058823 -0.028143 -0.006063  0.026984  0.088269 
## 
## Coefficients:
##                 Estimate Std. Error t value Pr(>|t|)    
## (Intercept)     0.146121   0.012200  11.977 5.71e-15 ***
## provenance_MAT  0.002421   0.002767   0.875   0.3867    
## k2_tricho      -0.054222   0.021649  -2.505   0.0163 *  
## ---
## Signif. codes:  0 '***' 0.001 '**' 0.01 '*' 0.05 '.' 0.1 ' ' 1
## 
## Residual standard error: 0.0358 on 41 degrees of freedom
## Multiple R-squared:  0.1521, Adjusted R-squared:  0.1107 
## F-statistic: 3.676 on 2 and 41 DF,  p-value: 0.034
## 
## 
## Call:
## lm(formula = form, data = dat)
## 
## Standardized Coefficients::
##    (Intercept) provenance_MAT      k2_tricho 
##             NA      0.1676495     -0.4798747 
## 
## [1] "Model: plasticity_rdpi_LMA_g_m2_2023 ~ provenance_MAP"
## 
## Call:
## lm(formula = form, data = dat)
## 
## Residuals:
##       Min        1Q    Median        3Q       Max 
## -0.059206 -0.024422 -0.006224  0.023075  0.095491 
## 
## Coefficients:
##                  Estimate Std. Error t value Pr(>|t|)    
## (Intercept)     1.408e-01  1.713e-02   8.219 3.32e-10 ***
## provenance_MAP  6.484e-06  2.543e-05   0.255   0.8000    
## k2_tricho      -4.275e-02  1.689e-02  -2.531   0.0153 *  
## ---
## Signif. codes:  0 '***' 0.001 '**' 0.01 '*' 0.05 '.' 0.1 ' ' 1
## 
## Residual standard error: 0.03611 on 41 degrees of freedom
## Multiple R-squared:  0.1376, Adjusted R-squared:  0.09553 
## F-statistic: 3.271 on 2 and 41 DF,  p-value: 0.04809
## 
## 
## Call:
## lm(formula = form, data = dat)
## 
## Standardized Coefficients::
##    (Intercept) provenance_MAP      k2_tricho 
##             NA     0.03811658    -0.37830882 
## 
## [1] "Model: plasticity_rdpi_LMA_g_m2_2023 ~ provenance_TD"
## 
## Call:
## lm(formula = form, data = dat)
## 
## Residuals:
##       Min        1Q    Median        3Q       Max 
## -0.055155 -0.022649 -0.009782  0.013318  0.100294 
## 
## Coefficients:
##                Estimate Std. Error t value Pr(>|t|)
## (Intercept)    0.095422   0.057306   1.665    0.104
## provenance_TD  0.001603   0.001853   0.865    0.392
## k2_tricho     -0.030512   0.020773  -1.469    0.150
## 
## Residual standard error: 0.03581 on 41 degrees of freedom
## Multiple R-squared:  0.1517, Adjusted R-squared:  0.1103 
## F-statistic: 3.666 on 2 and 41 DF,  p-value: 0.03428
## 
## 
## Call:
## lm(formula = form, data = dat)
## 
## Standardized Coefficients::
##   (Intercept) provenance_TD     k2_tricho 
##            NA      0.159056     -0.270034 
## 
## [1] "Model: plasticity_rdpi_lower_stomata_pore_length_mean_um ~ provenance_MAT"
## 
## Call:
## lm(formula = form, data = dat)
## 
## Residuals:
##      Min       1Q   Median       3Q      Max 
## -0.05852 -0.03268 -0.01863  0.01792  0.14263 
## 
## Coefficients:
##                 Estimate Std. Error t value Pr(>|t|)    
## (Intercept)     0.100446   0.016299   6.163 2.54e-07 ***
## provenance_MAT  0.001303   0.003697   0.352    0.726    
## k2_tricho      -0.012293   0.028922  -0.425    0.673    
## ---
## Signif. codes:  0 '***' 0.001 '**' 0.01 '*' 0.05 '.' 0.1 ' ' 1
## 
## Residual standard error: 0.04783 on 41 degrees of freedom
## Multiple R-squared:  0.004607,   Adjusted R-squared:  -0.04395 
## F-statistic: 0.09488 on 2 and 41 DF,  p-value: 0.9097
## 
## 
## Call:
## lm(formula = form, data = dat)
## 
## Standardized Coefficients::
##    (Intercept) provenance_MAT      k2_tricho 
##             NA     0.07316614    -0.08823278 
## 
## [1] "Model: plasticity_rdpi_lower_stomata_pore_length_mean_um ~ provenance_MAP"
## 
## Call:
## lm(formula = form, data = dat)
## 
## Residuals:
##      Min       1Q   Median       3Q      Max 
## -0.06659 -0.03407 -0.01614  0.02513  0.13184 
## 
## Coefficients:
##                  Estimate Std. Error t value Pr(>|t|)    
## (Intercept)     1.182e-01  2.235e-02   5.288 4.42e-06 ***
## provenance_MAP -3.945e-05  3.316e-05  -1.189    0.241    
## k2_tricho       7.769e-04  2.203e-02   0.035    0.972    
## ---
## Signif. codes:  0 '***' 0.001 '**' 0.01 '*' 0.05 '.' 0.1 ' ' 1
## 
## Residual standard error: 0.0471 on 41 degrees of freedom
## Multiple R-squared:  0.0349, Adjusted R-squared:  -0.01218 
## F-statistic: 0.7412 on 2 and 41 DF,  p-value: 0.4828
## 
## 
## Call:
## lm(formula = form, data = dat)
## 
## Standardized Coefficients::
##    (Intercept) provenance_MAP      k2_tricho 
##             NA   -0.188072827    0.005576568 
## 
## [1] "Model: plasticity_rdpi_lower_stomata_pore_length_mean_um ~ provenance_TD"
## 
## Call:
## lm(formula = form, data = dat)
## 
## Residuals:
##      Min       1Q   Median       3Q      Max 
## -0.05578 -0.03162 -0.01812  0.01628  0.14219 
## 
## Coefficients:
##                 Estimate Std. Error t value Pr(>|t|)
## (Intercept)    0.1160719  0.0766117   1.515    0.137
## provenance_TD -0.0005558  0.0024770  -0.224    0.824
## k2_tricho     -0.0094383  0.0277716  -0.340    0.736
## 
## Residual standard error: 0.04787 on 41 degrees of freedom
## Multiple R-squared:  0.002816,   Adjusted R-squared:  -0.04583 
## F-statistic: 0.05788 on 2 and 41 DF,  p-value: 0.9438
## 
## 
## Call:
## lm(formula = form, data = dat)
## 
## Standardized Coefficients::
##   (Intercept) provenance_TD     k2_tricho 
##            NA   -0.04472995   -0.06774400 
## 
## [1] "Model: plasticity_rdpi_upper_stomata_pore_length_mean_um ~ provenance_MAT"
## 
## Call:
## lm(formula = form, data = dat)
## 
## Residuals:
##      Min       1Q   Median       3Q      Max 
## -0.07109 -0.03561 -0.01080  0.01637  0.13709 
## 
## Coefficients:
##                 Estimate Std. Error t value Pr(>|t|)    
## (Intercept)     0.106004   0.021553   4.918 2.05e-05 ***
## provenance_MAT -0.004518   0.004463  -1.012    0.318    
## k2_tricho       0.043052   0.036712   1.173    0.249    
## ---
## Signif. codes:  0 '***' 0.001 '**' 0.01 '*' 0.05 '.' 0.1 ' ' 1
## 
## Residual standard error: 0.05092 on 35 degrees of freedom
##   (6 observations deleted due to missingness)
## Multiple R-squared:  0.04042,    Adjusted R-squared:  -0.01441 
## F-statistic: 0.7372 on 2 and 35 DF,  p-value: 0.4858
## 
## 
## Call:
## lm(formula = form, data = dat)
## 
## Standardized Coefficients::
##    (Intercept) provenance_MAT      k2_tricho 
##             NA     -0.2237107      0.2591314 
## 
## [1] "Model: plasticity_rdpi_upper_stomata_pore_length_mean_um ~ provenance_MAP"
## 
## Call:
## lm(formula = form, data = dat)
## 
## Residuals:
##       Min        1Q    Median        3Q       Max 
## -0.070633 -0.030102 -0.009313  0.019255  0.127676 
## 
## Coefficients:
##                  Estimate Std. Error t value Pr(>|t|)    
## (Intercept)     1.202e-01  2.741e-02   4.385 0.000101 ***
## provenance_MAP -1.694e-05  3.712e-05  -0.456 0.650895    
## k2_tricho       2.138e-02  2.856e-02   0.749 0.459105    
## ---
## Signif. codes:  0 '***' 0.001 '**' 0.01 '*' 0.05 '.' 0.1 ' ' 1
## 
## Residual standard error: 0.05151 on 35 degrees of freedom
##   (6 observations deleted due to missingness)
## Multiple R-squared:  0.01816,    Adjusted R-squared:  -0.03794 
## F-statistic: 0.3237 on 2 and 35 DF,  p-value: 0.7256
## 
## 
## Call:
## lm(formula = form, data = dat)
## 
## Standardized Coefficients::
##    (Intercept) provenance_MAP      k2_tricho 
##             NA    -0.07847209     0.12869973 
## 
## [1] "Model: plasticity_rdpi_upper_stomata_pore_length_mean_um ~ provenance_TD"
## 
## Call:
## lm(formula = form, data = dat)
## 
## Residuals:
##      Min       1Q   Median       3Q      Max 
## -0.07621 -0.02649 -0.01227  0.01498  0.12980 
## 
## Coefficients:
##                Estimate Std. Error t value Pr(>|t|)  
## (Intercept)   -0.059950   0.086245  -0.695   0.4916  
## provenance_TD  0.005660   0.002761   2.050   0.0479 *
## k2_tricho      0.058754   0.032898   1.786   0.0828 .
## ---
## Signif. codes:  0 '***' 0.001 '**' 0.01 '*' 0.05 '.' 0.1 ' ' 1
## 
## Residual standard error: 0.04882 on 35 degrees of freedom
##   (6 observations deleted due to missingness)
## Multiple R-squared:  0.1182, Adjusted R-squared:  0.06779 
## F-statistic: 2.345 on 2 and 35 DF,  p-value: 0.1107
## 
## 
## Call:
## lm(formula = form, data = dat)
## 
## Standardized Coefficients::
##   (Intercept) provenance_TD     k2_tricho 
##            NA     0.4058783     0.3536408 
## 
## [1] "Model: plasticity_rdpi_lower_stomata_density_mm2 ~ provenance_MAT"
## 
## Call:
## lm(formula = form, data = dat)
## 
## Residuals:
##     Min      1Q  Median      3Q     Max 
## -0.3541 -0.1546 -0.0653  0.1179  0.5484 
## 
## Coefficients:
##                Estimate Std. Error t value Pr(>|t|)    
## (Intercept)     0.40790    0.07649   5.333 3.82e-06 ***
## provenance_MAT  0.04224    0.01735   2.435   0.0193 *  
## k2_tricho      -0.25786    0.13572  -1.900   0.0645 .  
## ---
## Signif. codes:  0 '***' 0.001 '**' 0.01 '*' 0.05 '.' 0.1 ' ' 1
## 
## Residual standard error: 0.2244 on 41 degrees of freedom
## Multiple R-squared:  0.1291, Adjusted R-squared:  0.08662 
## F-statistic: 3.039 on 2 and 41 DF,  p-value: 0.05879
## 
## 
## Call:
## lm(formula = form, data = dat)
## 
## Standardized Coefficients::
##    (Intercept) provenance_MAT      k2_tricho 
##             NA      0.4727468     -0.3689224 
## 
## [1] "Model: plasticity_rdpi_lower_stomata_density_mm2 ~ provenance_MAP"
## 
## Call:
## lm(formula = form, data = dat)
## 
## Residuals:
##      Min       1Q   Median       3Q      Max 
## -0.27818 -0.17312 -0.08554  0.11722  0.64179 
## 
## Coefficients:
##                  Estimate Std. Error t value Pr(>|t|)    
## (Intercept)     0.4523606  0.1124650   4.022 0.000242 ***
## provenance_MAP -0.0001731  0.0001669  -1.037 0.305788    
## k2_tricho      -0.0117256  0.1108717  -0.106 0.916290    
## ---
## Signif. codes:  0 '***' 0.001 '**' 0.01 '*' 0.05 '.' 0.1 ' ' 1
## 
## Residual standard error: 0.237 on 41 degrees of freedom
## Multiple R-squared:  0.02868,    Adjusted R-squared:  -0.01871 
## F-statistic: 0.6052 on 2 and 41 DF,  p-value: 0.5508
## 
## 
## Call:
## lm(formula = form, data = dat)
## 
## Standardized Coefficients::
##    (Intercept) provenance_MAP      k2_tricho 
##             NA    -0.16450028    -0.01677565 
## 
## [1] "Model: plasticity_rdpi_lower_stomata_density_mm2 ~ provenance_TD"
## 
## Call:
## lm(formula = form, data = dat)
## 
## Residuals:
##     Min      1Q  Median      3Q     Max 
## -0.2331 -0.1697 -0.1094  0.1269  0.6406 
## 
## Coefficients:
##                Estimate Std. Error t value Pr(>|t|)
## (Intercept)    0.431604   0.384151   1.124    0.268
## provenance_TD -0.002057   0.012421  -0.166    0.869
## k2_tricho     -0.053883   0.139254  -0.387    0.701
## 
## Residual standard error: 0.24 on 41 degrees of freedom
## Multiple R-squared:  0.003863,   Adjusted R-squared:  -0.04473 
## F-statistic: 0.0795 on 2 and 41 DF,  p-value: 0.9237
## 
## 
## Call:
## lm(formula = form, data = dat)
## 
## Standardized Coefficients::
##   (Intercept) provenance_TD     k2_tricho 
##            NA   -0.03299518   -0.07708958 
## 
## [1] "Model: plasticity_rdpi_upper_stomata_density_mm2_log ~ provenance_MAT"
## 
## Call:
## lm(formula = form, data = dat)
## 
## Residuals:
##      Min       1Q   Median       3Q      Max 
## -108.744   -3.178    2.641    8.876   30.718 
## 
## Coefficients:
##                Estimate Std. Error t value Pr(>|t|)  
## (Intercept)     -18.374      7.760  -2.368   0.0227 *
## provenance_MAT   -4.426      1.760  -2.515   0.0159 *
## k2_tricho        30.134     13.770   2.188   0.0344 *
## ---
## Signif. codes:  0 '***' 0.001 '**' 0.01 '*' 0.05 '.' 0.1 ' ' 1
## 
## Residual standard error: 22.77 on 41 degrees of freedom
## Multiple R-squared:  0.1425, Adjusted R-squared:  0.1007 
## F-statistic: 3.408 on 2 and 41 DF,  p-value: 0.04274
## 
## 
## Call:
## lm(formula = form, data = dat)
## 
## Standardized Coefficients::
##    (Intercept) provenance_MAT      k2_tricho 
##             NA     -0.4844902      0.4216326 
## 
## [1] "Model: plasticity_rdpi_upper_stomata_density_mm2_log ~ provenance_MAP"
## 
## Call:
## lm(formula = form, data = dat)
## 
## Residuals:
##      Min       1Q   Median       3Q      Max 
## -122.625    1.309    6.127    9.800   16.379 
## 
## Coefficients:
##                 Estimate Std. Error t value Pr(>|t|)  
## (Intercept)    -23.04733   11.44944  -2.013   0.0507 .
## provenance_MAP   0.01817    0.01699   1.069   0.2912  
## k2_tricho        4.33624   11.28724   0.384   0.7028  
## ---
## Signif. codes:  0 '***' 0.001 '**' 0.01 '*' 0.05 '.' 0.1 ' ' 1
## 
## Residual standard error: 24.13 on 41 degrees of freedom
## Multiple R-squared:  0.03715,    Adjusted R-squared:  -0.00982 
## F-statistic: 0.7909 on 2 and 41 DF,  p-value: 0.4602
## 
## 
## Call:
## lm(formula = form, data = dat)
## 
## Standardized Coefficients::
##    (Intercept) provenance_MAP      k2_tricho 
##             NA     0.16886058     0.06067197 
## 
## [1] "Model: plasticity_rdpi_upper_stomata_density_mm2_log ~ provenance_TD"
## 
## Call:
## lm(formula = form, data = dat)
## 
## Residuals:
##      Min       1Q   Median       3Q      Max 
## -121.739    3.586    5.563    7.922   17.023 
## 
## Coefficients:
##               Estimate Std. Error t value Pr(>|t|)
## (Intercept)   -20.2944    39.1413  -0.518    0.607
## provenance_TD   0.1969     1.2655   0.156    0.877
## k2_tricho       8.6286    14.1887   0.608    0.546
## 
## Residual standard error: 24.46 on 41 degrees of freedom
## Multiple R-squared:  0.01088,    Adjusted R-squared:  -0.03737 
## F-statistic: 0.2256 on 2 and 41 DF,  p-value: 0.799
## 
## 
## Call:
## lm(formula = form, data = dat)
## 
## Standardized Coefficients::
##   (Intercept) provenance_TD     k2_tricho 
##            NA    0.03089159    0.12072979 
## 
## [1] "Model: plasticity_rdpi_upper_stomata_presence ~ provenance_MAT"
## 
## Call:
## lm(formula = form, data = dat)
## 
## Residuals:
##     Min      1Q  Median      3Q     Max 
## -0.8088 -0.3495  0.1120  0.3256  0.4866 
## 
## Coefficients:
##                Estimate Std. Error t value Pr(>|t|)    
## (Intercept)     0.87126    0.14062   6.196 2.51e-07 ***
## provenance_MAT  0.01593    0.03082   0.517   0.6080    
## k2_tricho      -0.40411    0.23992  -1.684   0.0999 .  
## ---
## Signif. codes:  0 '***' 0.001 '**' 0.01 '*' 0.05 '.' 0.1 ' ' 1
## 
## Residual standard error: 0.3934 on 40 degrees of freedom
##   (1 observation deleted due to missingness)
## Multiple R-squared:  0.07641,    Adjusted R-squared:  0.03023 
## F-statistic: 1.655 on 2 and 40 DF,  p-value: 0.204
## 
## 
## Call:
## lm(formula = form, data = dat)
## 
## Standardized Coefficients::
##    (Intercept) provenance_MAT      k2_tricho 
##             NA      0.1006725     -0.3279693 
## 
## [1] "Model: plasticity_rdpi_upper_stomata_presence ~ provenance_MAP"
## 
## Call:
## lm(formula = form, data = dat)
## 
## Residuals:
##     Min      1Q  Median      3Q     Max 
## -0.7783 -0.3103  0.1550  0.2961  0.5547 
## 
## Coefficients:
##                  Estimate Std. Error t value Pr(>|t|)    
## (Intercept)     0.9848389  0.1940344   5.076  9.3e-06 ***
## provenance_MAP -0.0002516  0.0002761  -0.911    0.368    
## k2_tricho      -0.2904410  0.1901095  -1.528    0.134    
## ---
## Signif. codes:  0 '***' 0.001 '**' 0.01 '*' 0.05 '.' 0.1 ' ' 1
## 
## Residual standard error: 0.3907 on 40 degrees of freedom
##   (1 observation deleted due to missingness)
## Multiple R-squared:  0.08915,    Adjusted R-squared:  0.0436 
## F-statistic: 1.957 on 2 and 40 DF,  p-value: 0.1545
## 
## 
## Call:
## lm(formula = form, data = dat)
## 
## Standardized Coefficients::
##    (Intercept) provenance_MAP      k2_tricho 
##             NA     -0.1406034     -0.2357167 
## 
## [1] "Model: plasticity_rdpi_upper_stomata_presence ~ provenance_TD"
## 
## Call:
## lm(formula = form, data = dat)
## 
## Residuals:
##     Min      1Q  Median      3Q     Max 
## -0.7670 -0.3254  0.1495  0.2988  0.4954 
## 
## Coefficients:
##                Estimate Std. Error t value Pr(>|t|)
## (Intercept)    0.611704   0.646844   0.946    0.350
## provenance_TD  0.008421   0.021363   0.394    0.696
## k2_tricho     -0.274716   0.229003  -1.200    0.237
## 
## Residual standard error: 0.394 on 40 degrees of freedom
##   (1 observation deleted due to missingness)
## Multiple R-squared:  0.07383,    Adjusted R-squared:  0.02752 
## F-statistic: 1.594 on 2 and 40 DF,  p-value: 0.2157
## 
## 
## Call:
## lm(formula = form, data = dat)
## 
## Standardized Coefficients::
##   (Intercept) provenance_TD     k2_tricho 
##            NA    0.07325941   -0.22295441 
## 
## [1] "Model: plasticity_rdpi_upper_stomata_density_over_total_density_log ~ provenance_MAT"
## 
## Call:
## lm(formula = form, data = dat)
## 
## Residuals:
##      Min       1Q   Median       3Q      Max 
## -0.47351 -0.04147  0.03416  0.07899  0.21784 
## 
## Coefficients:
##                Estimate Std. Error t value Pr(>|t|)    
## (Intercept)    -0.22539    0.05310  -4.245 0.000122 ***
## provenance_MAT -0.01058    0.01204  -0.878 0.384893    
## k2_tricho      -0.02280    0.09422  -0.242 0.809949    
## ---
## Signif. codes:  0 '***' 0.001 '**' 0.01 '*' 0.05 '.' 0.1 ' ' 1
## 
## Residual standard error: 0.1558 on 41 degrees of freedom
## Multiple R-squared:  0.04589,    Adjusted R-squared:  -0.000651 
## F-statistic: 0.986 on 2 and 41 DF,  p-value: 0.3817
## 
## 
## Call:
## lm(formula = form, data = dat)
## 
## Standardized Coefficients::
##    (Intercept) provenance_MAT      k2_tricho 
##             NA    -0.17850877    -0.04919411 
## 
## [1] "Model: plasticity_rdpi_upper_stomata_density_over_total_density_log ~ provenance_MAP"
## 
## Call:
## lm(formula = form, data = dat)
## 
## Residuals:
##      Min       1Q   Median       3Q      Max 
## -0.50011 -0.04712  0.02442  0.08869  0.23772 
## 
## Coefficients:
##                  Estimate Std. Error t value Pr(>|t|)    
## (Intercept)    -0.3095060  0.0717228  -4.315 9.81e-05 ***
## provenance_MAP  0.0001956  0.0001064   1.837   0.0734 .  
## k2_tricho      -0.1088915  0.0707067  -1.540   0.1312    
## ---
## Signif. codes:  0 '***' 0.001 '**' 0.01 '*' 0.05 '.' 0.1 ' ' 1
## 
## Residual standard error: 0.1512 on 41 degrees of freedom
## Multiple R-squared:  0.1019, Adjusted R-squared:  0.05809 
## F-statistic: 2.326 on 2 and 41 DF,  p-value: 0.1105
## 
## 
## Call:
## lm(formula = form, data = dat)
## 
## Standardized Coefficients::
##    (Intercept) provenance_MAP      k2_tricho 
##             NA      0.2802660     -0.2348976 
## 
## [1] "Model: plasticity_rdpi_upper_stomata_density_over_total_density_log ~ provenance_TD"
## 
## Call:
## lm(formula = form, data = dat)
## 
## Residuals:
##      Min       1Q   Median       3Q      Max 
## -0.49019 -0.03585  0.03144  0.07755  0.24252 
## 
## Coefficients:
##                Estimate Std. Error t value Pr(>|t|)
## (Intercept)   -0.067106   0.250558  -0.268    0.790
## provenance_TD -0.004914   0.008101  -0.607    0.548
## k2_tricho     -0.111795   0.090827  -1.231    0.225
## 
## Residual standard error: 0.1566 on 41 degrees of freedom
## Multiple R-squared:  0.03658,    Adjusted R-squared:  -0.01041 
## F-statistic: 0.7784 on 2 and 41 DF,  p-value: 0.4658
## 
## 
## Call:
## lm(formula = form, data = dat)
## 
## Standardized Coefficients::
##   (Intercept) provenance_TD     k2_tricho 
##            NA    -0.1188351    -0.2411606 
## 
## [1] "Model: plasticity_rdpi_licor_gsw ~ provenance_MAT"
## 
## Call:
## lm(formula = form, data = dat)
## 
## Residuals:
##      Min       1Q   Median       3Q      Max 
## -0.13517 -0.05267 -0.01382  0.06376  0.21463 
## 
## Coefficients:
##                 Estimate Std. Error t value Pr(>|t|)    
## (Intercept)     0.286846   0.027617  10.386 4.76e-13 ***
## provenance_MAT  0.011464   0.006264   1.830  0.07451 .  
## k2_tricho      -0.145153   0.049005  -2.962  0.00507 ** 
## ---
## Signif. codes:  0 '***' 0.001 '**' 0.01 '*' 0.05 '.' 0.1 ' ' 1
## 
## Residual standard error: 0.08104 on 41 degrees of freedom
## Multiple R-squared:  0.1767, Adjusted R-squared:  0.1366 
## F-statistic: 4.401 on 2 and 41 DF,  p-value: 0.01856
## 
## 
## Call:
## lm(formula = form, data = dat)
## 
## Standardized Coefficients::
##    (Intercept) provenance_MAT      k2_tricho 
##             NA      0.3455137     -0.5591981 
## 
## [1] "Model: plasticity_rdpi_licor_gsw ~ provenance_MAP"
## 
## Call:
## lm(formula = form, data = dat)
## 
## Residuals:
##      Min       1Q   Median       3Q      Max 
## -0.13006 -0.06022 -0.02615  0.05428  0.21420 
## 
## Coefficients:
##                  Estimate Std. Error t value Pr(>|t|)    
## (Intercept)     2.404e-01  3.920e-02   6.132  2.8e-07 ***
## provenance_MAP  7.506e-05  5.818e-05   1.290   0.2043    
## k2_tricho      -9.794e-02  3.865e-02  -2.534   0.0152 *  
## ---
## Signif. codes:  0 '***' 0.001 '**' 0.01 '*' 0.05 '.' 0.1 ' ' 1
## 
## Residual standard error: 0.08263 on 41 degrees of freedom
## Multiple R-squared:  0.1442, Adjusted R-squared:  0.1025 
## F-statistic: 3.455 on 2 and 41 DF,  p-value: 0.04105
## 
## 
## Call:
## lm(formula = form, data = dat)
## 
## Standardized Coefficients::
##    (Intercept) provenance_MAP      k2_tricho 
##             NA      0.1920782     -0.3773197 
## 
## [1] "Model: plasticity_rdpi_licor_gsw ~ provenance_TD"
## 
## Call:
## lm(formula = form, data = dat)
## 
## Residuals:
##      Min       1Q   Median       3Q      Max 
## -0.10934 -0.06214 -0.01118  0.05728  0.20812 
## 
## Coefficients:
##                Estimate Std. Error t value Pr(>|t|)   
## (Intercept)    0.413774   0.133090   3.109  0.00341 **
## provenance_TD -0.004542   0.004303  -1.055  0.29741   
## k2_tricho     -0.117603   0.048245  -2.438  0.01921 * 
## ---
## Signif. codes:  0 '***' 0.001 '**' 0.01 '*' 0.05 '.' 0.1 ' ' 1
## 
## Residual standard error: 0.08316 on 41 degrees of freedom
## Multiple R-squared:  0.133,  Adjusted R-squared:  0.09075 
## F-statistic: 3.146 on 2 and 41 DF,  p-value: 0.05358
## 
## 
## Call:
## lm(formula = form, data = dat)
## 
## Standardized Coefficients::
##   (Intercept) provenance_TD     k2_tricho 
##            NA     -0.196164     -0.453061 
## 
## [1] "Model: plasticity_rdpi_licor_ETR ~ provenance_MAT"
## 
## Call:
## lm(formula = form, data = dat)
## 
## Residuals:
##       Min        1Q    Median        3Q       Max 
## -0.274386 -0.073068 -0.002912  0.089357  0.232713 
## 
## Coefficients:
##                  Estimate Std. Error t value Pr(>|t|)    
## (Intercept)     0.4592411  0.0454497  10.104 1.08e-12 ***
## provenance_MAT  0.0036430  0.0103085   0.353    0.726    
## k2_tricho      -0.0008897  0.0806479  -0.011    0.991    
## ---
## Signif. codes:  0 '***' 0.001 '**' 0.01 '*' 0.05 '.' 0.1 ' ' 1
## 
## Residual standard error: 0.1334 on 41 degrees of freedom
## Multiple R-squared:  0.005162,   Adjusted R-squared:  -0.04337 
## F-statistic: 0.1064 on 2 and 41 DF,  p-value: 0.8993
## 
## 
## Call:
## lm(formula = form, data = dat)
## 
## Standardized Coefficients::
##    (Intercept) provenance_MAT      k2_tricho 
##             NA    0.073341115   -0.002289608 
## 
## [1] "Model: plasticity_rdpi_licor_ETR ~ provenance_MAP"
## 
## Call:
## lm(formula = form, data = dat)
## 
## Residuals:
##       Min        1Q    Median        3Q       Max 
## -0.270704 -0.062582 -0.007301  0.075689  0.277356 
## 
## Coefficients:
##                  Estimate Std. Error t value Pr(>|t|)    
## (Intercept)     4.917e-01  6.289e-02   7.819 1.18e-09 ***
## provenance_MAP -7.470e-05  9.333e-05  -0.800    0.428    
## k2_tricho       2.994e-02  6.200e-02   0.483    0.632    
## ---
## Signif. codes:  0 '***' 0.001 '**' 0.01 '*' 0.05 '.' 0.1 ' ' 1
## 
## Residual standard error: 0.1325 on 41 degrees of freedom
## Multiple R-squared:  0.01748,    Adjusted R-squared:  -0.03044 
## F-statistic: 0.3648 on 2 and 41 DF,  p-value: 0.6966
## 
## 
## Call:
## lm(formula = form, data = dat)
## 
## Standardized Coefficients::
##    (Intercept) provenance_MAP      k2_tricho 
##             NA     -0.1276928      0.0770398 
## 
## [1] "Model: plasticity_rdpi_licor_ETR ~ provenance_TD"
## 
## Call:
## lm(formula = form, data = dat)
## 
## Residuals:
##       Min        1Q    Median        3Q       Max 
## -0.272441 -0.059742 -0.009152  0.086412  0.248993 
## 
## Coefficients:
##               Estimate Std. Error t value Pr(>|t|)  
## (Intercept)   0.386710   0.213475   1.812   0.0774 .
## provenance_TD 0.002288   0.006902   0.331   0.7420  
## k2_tricho     0.033918   0.077384   0.438   0.6635  
## ---
## Signif. codes:  0 '***' 0.001 '**' 0.01 '*' 0.05 '.' 0.1 ' ' 1
## 
## Residual standard error: 0.1334 on 41 degrees of freedom
## Multiple R-squared:  0.004799,   Adjusted R-squared:  -0.04375 
## F-statistic: 0.09885 on 2 and 41 DF,  p-value: 0.9061
## 
## 
## Call:
## lm(formula = form, data = dat)
## 
## Standardized Coefficients::
##   (Intercept) provenance_TD     k2_tricho 
##            NA    0.06600787    0.08728294 
## 
## [1] "Model: plasticity_rdpi_licor_Fm. ~ provenance_MAT"
## 
## Call:
## lm(formula = form, data = dat)
## 
## Residuals:
##      Min       1Q   Median       3Q      Max 
## -0.12932 -0.02720 -0.00041  0.03510  0.09607 
## 
## Coefficients:
##                Estimate Std. Error t value Pr(>|t|)    
## (Intercept)    0.214158   0.018347  11.673  1.3e-14 ***
## provenance_MAT 0.002482   0.004161   0.596    0.554    
## k2_tricho      0.018968   0.032556   0.583    0.563    
## ---
## Signif. codes:  0 '***' 0.001 '**' 0.01 '*' 0.05 '.' 0.1 ' ' 1
## 
## Residual standard error: 0.05384 on 41 degrees of freedom
## Multiple R-squared:  0.0476, Adjusted R-squared:  0.001137 
## F-statistic: 1.024 on 2 and 41 DF,  p-value: 0.368
## 
## 
## Call:
## lm(formula = form, data = dat)
## 
## Standardized Coefficients::
##    (Intercept) provenance_MAT      k2_tricho 
##             NA      0.1210986      0.1183073 
## 
## [1] "Model: plasticity_rdpi_licor_Fm. ~ provenance_MAP"
## 
## Call:
## lm(formula = form, data = dat)
## 
## Residuals:
##       Min        1Q    Median        3Q       Max 
## -0.137325 -0.028234 -0.004439  0.036271  0.104024 
## 
## Coefficients:
##                  Estimate Std. Error t value Pr(>|t|)    
## (Intercept)     2.202e-01  2.559e-02   8.607 9.89e-11 ***
## provenance_MAP -1.741e-05  3.798e-05  -0.458    0.649    
## k2_tricho       3.459e-02  2.523e-02   1.371    0.178    
## ---
## Signif. codes:  0 '***' 0.001 '**' 0.01 '*' 0.05 '.' 0.1 ' ' 1
## 
## Residual standard error: 0.05393 on 41 degrees of freedom
## Multiple R-squared:  0.04423,    Adjusted R-squared:  -0.002391 
## F-statistic: 0.9487 on 2 and 41 DF,  p-value: 0.3956
## 
## 
## Call:
## lm(formula = form, data = dat)
## 
## Standardized Coefficients::
##    (Intercept) provenance_MAP      k2_tricho 
##             NA    -0.07212616     0.21576195 
## 
## [1] "Model: plasticity_rdpi_licor_Fm. ~ provenance_TD"
## 
## Call:
## lm(formula = form, data = dat)
## 
## Residuals:
##      Min       1Q   Median       3Q      Max 
## -0.13714 -0.02809 -0.00078  0.03687  0.10599 
## 
## Coefficients:
##               Estimate Std. Error t value Pr(>|t|)  
## (Intercept)   0.180210   0.086384   2.086   0.0432 *
## provenance_TD 0.001047   0.002793   0.375   0.7096  
## k2_tricho     0.039111   0.031314   1.249   0.2188  
## ---
## Signif. codes:  0 '***' 0.001 '**' 0.01 '*' 0.05 '.' 0.1 ' ' 1
## 
## Residual standard error: 0.05398 on 41 degrees of freedom
## Multiple R-squared:  0.04262,    Adjusted R-squared:  -0.004084 
## F-statistic: 0.9126 on 2 and 41 DF,  p-value: 0.4095
## 
## 
## Call:
## lm(formula = form, data = dat)
## 
## Standardized Coefficients::
##   (Intercept) provenance_TD     k2_tricho 
##            NA    0.07324831    0.24394649 
## 
## [1] "Model: plasticity_rdpi_licor_Fs ~ provenance_MAT"
## 
## Call:
## lm(formula = form, data = dat)
## 
## Residuals:
##       Min        1Q    Median        3Q       Max 
## -0.090179 -0.029448  0.001243  0.026846  0.096676 
## 
## Coefficients:
##                 Estimate Std. Error t value Pr(>|t|)    
## (Intercept)     0.146569   0.014142  10.364 5.08e-13 ***
## provenance_MAT -0.005776   0.003208  -1.801   0.0791 .  
## k2_tricho      -0.001514   0.025094  -0.060   0.9522    
## ---
## Signif. codes:  0 '***' 0.001 '**' 0.01 '*' 0.05 '.' 0.1 ' ' 1
## 
## Residual standard error: 0.0415 on 41 degrees of freedom
## Multiple R-squared:  0.128,  Adjusted R-squared:  0.08544 
## F-statistic: 3.009 on 2 and 41 DF,  p-value: 0.06037
## 
## 
## Call:
## lm(formula = form, data = dat)
## 
## Standardized Coefficients::
##    (Intercept) provenance_MAT      k2_tricho 
##             NA    -0.34988724    -0.01172008 
## 
## [1] "Model: plasticity_rdpi_licor_Fs ~ provenance_MAP"
## 
## Call:
## lm(formula = form, data = dat)
## 
## Residuals:
##       Min        1Q    Median        3Q       Max 
## -0.084246 -0.031354 -0.003446  0.030490  0.109006 
## 
## Coefficients:
##                  Estimate Std. Error t value Pr(>|t|)    
## (Intercept)     1.549e-01  2.044e-02   7.578 2.55e-09 ***
## provenance_MAP -6.422e-06  3.034e-05  -0.212    0.833    
## k2_tricho      -3.034e-02  2.015e-02  -1.506    0.140    
## ---
## Signif. codes:  0 '***' 0.001 '**' 0.01 '*' 0.05 '.' 0.1 ' ' 1
## 
## Residual standard error: 0.04309 on 41 degrees of freedom
## Multiple R-squared:  0.06004,    Adjusted R-squared:  0.01419 
## F-statistic: 1.309 on 2 and 41 DF,  p-value: 0.281
## 
## 
## Call:
## lm(formula = form, data = dat)
## 
## Standardized Coefficients::
##    (Intercept) provenance_MAP      k2_tricho 
##             NA    -0.03302946    -0.23493504 
## 
## [1] "Model: plasticity_rdpi_licor_Fs ~ provenance_TD"
## 
## Call:
## lm(formula = form, data = dat)
## 
## Residuals:
##       Min        1Q    Median        3Q       Max 
## -0.069473 -0.031302 -0.005684  0.030405  0.101418 
## 
## Coefficients:
##               Estimate Std. Error t value Pr(>|t|)  
## (Intercept)   0.002044   0.064709   0.032   0.9750  
## provenance_TD 0.004952   0.002092   2.367   0.0227 *
## k2_tricho     0.003204   0.023457   0.137   0.8920  
## ---
## Signif. codes:  0 '***' 0.001 '**' 0.01 '*' 0.05 '.' 0.1 ' ' 1
## 
## Residual standard error: 0.04043 on 41 degrees of freedom
## Multiple R-squared:  0.1721, Adjusted R-squared:  0.1317 
## F-statistic: 4.262 on 2 and 41 DF,  p-value: 0.02081
## 
## 
## Call:
## lm(formula = form, data = dat)
## 
## Standardized Coefficients::
##   (Intercept) provenance_TD     k2_tricho 
##            NA    0.42987321    0.02480831 
## 
## [1] "Model: plasticity_rdpi_licor_gbw ~ provenance_MAT"
## 
## Call:
## lm(formula = form, data = dat)
## 
## Residuals:
##        Min         1Q     Median         3Q        Max 
## -1.167e-04 -6.262e-05 -2.012e-05  3.131e-05  2.761e-04 
## 
## Coefficients:
##                  Estimate Std. Error t value Pr(>|t|)    
## (Intercept)     4.147e-04  3.039e-05  13.644   <2e-16 ***
## provenance_MAT -3.744e-06  6.894e-06  -0.543    0.590    
## k2_tricho       6.705e-06  5.393e-05   0.124    0.902    
## ---
## Signif. codes:  0 '***' 0.001 '**' 0.01 '*' 0.05 '.' 0.1 ' ' 1
## 
## Residual standard error: 8.919e-05 on 41 degrees of freedom
## Multiple R-squared:  0.009484,   Adjusted R-squared:  -0.03883 
## F-statistic: 0.1963 on 2 and 41 DF,  p-value: 0.8225
## 
## 
## Call:
## lm(formula = form, data = dat)
## 
## Standardized Coefficients::
##    (Intercept) provenance_MAT      k2_tricho 
##             NA    -0.11246211     0.02574589 
## 
## [1] "Model: plasticity_rdpi_licor_gbw ~ provenance_MAP"
## 
## Call:
## lm(formula = form, data = dat)
## 
## Residuals:
##        Min         1Q     Median         3Q        Max 
## -1.310e-04 -6.068e-05 -1.664e-05  3.085e-05  2.625e-04 
## 
## Coefficients:
##                  Estimate Std. Error t value Pr(>|t|)    
## (Intercept)     3.913e-04  4.206e-05   9.304 1.17e-11 ***
## provenance_MAP  5.589e-08  6.242e-08   0.895    0.376    
## k2_tricho      -2.162e-05  4.147e-05  -0.521    0.605    
## ---
## Signif. codes:  0 '***' 0.001 '**' 0.01 '*' 0.05 '.' 0.1 ' ' 1
## 
## Residual standard error: 8.865e-05 on 41 degrees of freedom
## Multiple R-squared:  0.02149,    Adjusted R-squared:  -0.02625 
## F-statistic: 0.4501 on 2 and 41 DF,  p-value: 0.6407
## 
## 
## Call:
## lm(formula = form, data = dat)
## 
## Standardized Coefficients::
##    (Intercept) provenance_MAP      k2_tricho 
##             NA      0.1425294     -0.0830204 
## 
## [1] "Model: plasticity_rdpi_licor_gbw ~ provenance_TD"
## 
## Call:
## lm(formula = form, data = dat)
## 
## Residuals:
##        Min         1Q     Median         3Q        Max 
## -1.318e-04 -5.924e-05 -2.244e-05  4.289e-05  2.650e-04 
## 
## Coefficients:
##                 Estimate Std. Error t value Pr(>|t|)   
## (Intercept)    4.773e-04  1.429e-04   3.339   0.0018 **
## provenance_TD -1.956e-06  4.621e-06  -0.423   0.6743   
## k2_tricho     -2.631e-05  5.181e-05  -0.508   0.6143   
## ---
## Signif. codes:  0 '***' 0.001 '**' 0.01 '*' 0.05 '.' 0.1 ' ' 1
## 
## Residual standard error: 8.932e-05 on 41 degrees of freedom
## Multiple R-squared:  0.006701,   Adjusted R-squared:  -0.04175 
## F-statistic: 0.1383 on 2 and 41 DF,  p-value: 0.8713
## 
## 
## Call:
## lm(formula = form, data = dat)
## 
## Standardized Coefficients::
##   (Intercept) provenance_TD     k2_tricho 
##            NA   -0.08422081   -0.10102022 
## 
## [1] "Model: plasticity_rdpi_licor_PhiPS2 ~ provenance_MAT"
## 
## Call:
## lm(formula = form, data = dat)
## 
## Residuals:
##       Min        1Q    Median        3Q       Max 
## -0.119708 -0.028687  0.008714  0.037819  0.085722 
## 
## Coefficients:
##                  Estimate Std. Error t value Pr(>|t|)    
## (Intercept)     0.2592436  0.0173305  14.959   <2e-16 ***
## provenance_MAT  0.0001917  0.0039308   0.049    0.961    
## k2_tricho      -0.0050308  0.0307521  -0.164    0.871    
## ---
## Signif. codes:  0 '***' 0.001 '**' 0.01 '*' 0.05 '.' 0.1 ' ' 1
## 
## Residual standard error: 0.05086 on 41 degrees of freedom
## Multiple R-squared:  0.0008045,  Adjusted R-squared:  -0.04794 
## F-statistic: 0.0165 on 2 and 41 DF,  p-value: 0.9836
## 
## 
## Call:
## lm(formula = form, data = dat)
## 
## Standardized Coefficients::
##    (Intercept) provenance_MAT      k2_tricho 
##             NA     0.01014398    -0.03402514 
## 
## [1] "Model: plasticity_rdpi_licor_PhiPS2 ~ provenance_MAP"
## 
## Call:
## lm(formula = form, data = dat)
## 
## Residuals:
##       Min        1Q    Median        3Q       Max 
## -0.119532 -0.028972  0.006922  0.038493  0.084064 
## 
## Coefficients:
##                  Estimate Std. Error t value Pr(>|t|)    
## (Intercept)     2.569e-01  2.412e-02  10.648 2.25e-13 ***
## provenance_MAP  4.583e-06  3.581e-05   0.128    0.899    
## k2_tricho      -4.776e-03  2.378e-02  -0.201    0.842    
## ---
## Signif. codes:  0 '***' 0.001 '**' 0.01 '*' 0.05 '.' 0.1 ' ' 1
## 
## Residual standard error: 0.05085 on 41 degrees of freedom
## Multiple R-squared:  0.001146,   Adjusted R-squared:  -0.04758 
## F-statistic: 0.02351 on 2 and 41 DF,  p-value: 0.9768
## 
## 
## Call:
## lm(formula = form, data = dat)
## 
## Standardized Coefficients::
##    (Intercept) provenance_MAP      k2_tricho 
##             NA     0.02058704    -0.03229865 
## 
## [1] "Model: plasticity_rdpi_licor_PhiPS2 ~ provenance_TD"
## 
## Call:
## lm(formula = form, data = dat)
## 
## Residuals:
##       Min        1Q    Median        3Q       Max 
## -0.109780 -0.028940  0.008827  0.038643  0.087067 
## 
## Coefficients:
##               Estimate Std. Error t value Pr(>|t|)  
## (Intercept)   0.152253   0.079581   1.913   0.0627 .
## provenance_TD 0.003531   0.002573   1.372   0.1774  
## k2_tricho     0.020617   0.028848   0.715   0.4789  
## ---
## Signif. codes:  0 '***' 0.001 '**' 0.01 '*' 0.05 '.' 0.1 ' ' 1
## 
## Residual standard error: 0.04973 on 41 degrees of freedom
## Multiple R-squared:  0.04463,    Adjusted R-squared:  -0.001973 
## F-statistic: 0.9577 on 2 and 41 DF,  p-value: 0.3922
## 
## 
## Call:
## lm(formula = form, data = dat)
## 
## Standardized Coefficients::
##   (Intercept) provenance_TD     k2_tricho 
##            NA     0.2677557     0.1394376
```

```
# adjust p-values

padj <- p.adjust(unlist(res.lr.pt$pvals), method = 'BH')
res.lr.pt$padj <- as.data.frame(matrix(padj, ncol = ncol(res.lr.pt$pvals)))
colnames(res.lr.pt$padj) <- colnames(res.lr.pt$pvals)
rownames(res.lr.pt$padj) <- rownames(res.lr.pt$pvals)

plot(unlist(res.lr.pt$pvals), unlist(res.lr.pt$padj))
```

```
sum(res.lr.pt$pvals < 0.05) # 12
```

```
## [1] 12
```

```
sum(res.lr.pt$pvals < 0.01) # 1
```

```
## [1] 1
```

```
sum(res.lr.pt$padj < 0.05) # 1
```

```
## [1] 1
```

```
sum(res.lr.pt$padj < 0.1) # 1
```

```
## [1] 1
```

```
###################
# heatmap by standardized effect size

colnames(res.lr.pt$cors.sig) <- gsub('plasticity_rdpi_', '', colnames(res.lr.pt$cors.sig))
colnames(res.lr.pt$padj) <- gsub('plasticity_rdpi_', '', colnames(res.lr.pt$padj))


# row annotation - trait categories
cats <- c(rep('Phenology', 14),  rep('Leaf Morphology', 4), rep('Stomata', 7), rep('Photosynthesis', 5))
cbind(colnames(res.lr.pt$cors.sig), cats)
```

```
##                                                      cats             
##  [1,] "growing_season_days_2023"                     "Phenology"      
##  [2,] "growing_season_days_2024"                     "Phenology"      
##  [3,] "DOY_Stage2_2023"                              "Phenology"      
##  [4,] "DOY_Stage2_2024"                              "Phenology"      
##  [5,] "DOY_Stage3_2023"                              "Phenology"      
##  [6,] "DOY_Stage3_2024"                              "Phenology"      
##  [7,] "DOY_Stage6_2023"                              "Phenology"      
##  [8,] "DOY_Stage6_2024"                              "Phenology"      
##  [9,] "DOY_Stage7_2023"                              "Phenology"      
## [10,] "DOY_Stage7_2024"                              "Phenology"      
## [11,] "stage7_presence_2023"                         "Phenology"      
## [12,] "stage7_presence_2024"                         "Phenology"      
## [13,] "DOY_last_budset_2023"                         "Phenology"      
## [14,] "DOY_last_budset_2024"                         "Phenology"      
## [15,] "leaf_thickness_avg_mm_2023"                   "Leaf Morphology"
## [16,] "leaf_area_cm2_2023_log"                       "Leaf Morphology"
## [17,] "leaf_mass_g_2023_log"                         "Leaf Morphology"
## [18,] "LMA_g_m2_2023"                                "Leaf Morphology"
## [19,] "lower_stomata_pore_length_mean_um"            "Stomata"        
## [20,] "upper_stomata_pore_length_mean_um"            "Stomata"        
## [21,] "lower_stomata_density_mm2"                    "Stomata"        
## [22,] "upper_stomata_density_mm2_log"                "Stomata"        
## [23,] "upper_stomata_presence"                       "Stomata"        
## [24,] "upper_stomata_density_over_total_density_log" "Stomata"        
## [25,] "licor_gsw"                                    "Stomata"        
## [26,] "licor_ETR"                                    "Photosynthesis" 
## [27,] "licor_Fm."                                    "Photosynthesis" 
## [28,] "licor_Fs"                                     "Photosynthesis" 
## [29,] "licor_gbw"                                    "Photosynthesis" 
## [30,] "licor_PhiPS2"                                 "Photosynthesis"
```

```
ha_row <- rowAnnotation(Trait_Category = cats,
                        show_annotation_name = F,
                            col = list(Trait_Category = c('Phenology' = "#e2e260", 'Photosynthesis' = "#2d4030", 'Leaf Morphology' = "#516823", 'Stomata' =  "#88a2b9")))

# cell text showing significance
# cell_text <- res.lr.pt$pvals
# cell_text[res.lr.pt$pvals > 0.05] <- ''
# cell_text[res.lr.pt$pvals <= 0.05 & res.lr.pt$pvals > 0.01] <- '*'
# cell_text[res.lr.pt$pvals <= 0.01 & res.lr.pt$pvals > 0.001] <- '**'
# cell_text[res.lr.pt$pvals<= 0.001] <- '***'
# cell_text <- t(cell_text)

# or just run for adjusted pvals

cell_text <- res.lr.pt$padj
cell_text[res.lr.pt$padj > 0.05] <- ''
cell_text[res.lr.pt$padj <= 0.05 & res.lr.pt$padj > 0.01] <- '*'
cell_text[res.lr.pt$padj <= 0.01 & res.lr.pt$padj > 0.001] <- '**'
cell_text[res.lr.pt$padj<= 0.001] <- '***'
cell_text <- t(cell_text)

# heatmap by standardized effect size

# color scale based on all effect sizes
max_col <- max(unlist(res.lr.pt$eff.std.sig), na.rm = T)
min_col <- min(unlist(res.lr.pt$eff.std.sig), na.rm = T)
colf <- colorRamp2(c('red', 'white', 'blue'), breaks = c(min_col, 0, max_col))


hm2 <- Heatmap(as.matrix(t(res.lr.pt$eff.std.sig)),
        cluster_rows = F,
        cluster_columns = F,
        #col = colf(100),
        col = colf,
        row_names_side = 'left',
        column_names_side = 'top',
        column_names_rot = 90,
        column_names_centered = T,
        column_title = 'Predictors',
        row_title = 'Trait plasticity',
        left_annotation = ha_row,
        heatmap_legend_param = list(title = 'Std. Effect Size', at = c(-0.6, 0, 0.6)),
        cell_fun = function(j, i, x, y, width, height, fill) { # add text to each grid
                      grid.text(cell_text[i, j], x, y, gp = gpar(cex = 1.5))})

ComplexHeatmap::draw(hm2, padding = unit(c(2, 30, 2, 15), "mm"))
```

#### 4.1 Tables with results

```
kable(res.lr.pt$eff.std.sig, label = 'Standardized effects for significant relationships between plasticity and climate, controlling for ancestry')
```

|  | plasticity\_rdpi\_growing\_season\_days\_2023 | plasticity\_rdpi\_growing\_season\_days\_2024 | plasticity\_rdpi\_DOY\_Stage2\_2023 | plasticity\_rdpi\_DOY\_Stage2\_2024 | plasticity\_rdpi\_DOY\_Stage3\_2023 | plasticity\_rdpi\_DOY\_Stage3\_2024 | plasticity\_rdpi\_DOY\_Stage6\_2023 | plasticity\_rdpi\_DOY\_Stage6\_2024 | plasticity\_rdpi\_DOY\_Stage7\_2023 | plasticity\_rdpi\_DOY\_Stage7\_2024 | plasticity\_rdpi\_stage7\_presence\_2023 | plasticity\_rdpi\_stage7\_presence\_2024 | plasticity\_rdpi\_DOY\_last\_budset\_2023 | plasticity\_rdpi\_DOY\_last\_budset\_2024 | plasticity\_rdpi\_leaf\_thickness\_avg\_mm\_2023 | plasticity\_rdpi\_leaf\_area\_cm2\_2023\_log | plasticity\_rdpi\_leaf\_mass\_g\_2023\_log | plasticity\_rdpi\_LMA\_g\_m2\_2023 | plasticity\_rdpi\_lower\_stomata\_pore\_length\_mean\_um | plasticity\_rdpi\_upper\_stomata\_pore\_length\_mean\_um | plasticity\_rdpi\_lower\_stomata\_density\_mm2 | plasticity\_rdpi\_upper\_stomata\_density\_mm2\_log | plasticity\_rdpi\_upper\_stomata\_presence | plasticity\_rdpi\_upper\_stomata\_density\_over\_total\_density\_log | plasticity\_rdpi\_licor\_gsw | plasticity\_rdpi\_licor\_ETR | plasticity\_rdpi\_licor\_Fm. | plasticity\_rdpi\_licor\_Fs | plasticity\_rdpi\_licor\_gbw | plasticity\_rdpi\_licor\_PhiPS2 |
| --- | --- | --- | --- | --- | --- | --- | --- | --- | --- | --- | --- | --- | --- | --- | --- | --- | --- | --- | --- | --- | --- | --- | --- | --- | --- | --- | --- | --- | --- | --- |
| provenance\_MAT | NA | NA | NA | NA | NA | NA | NA | NA | NA | -0.6239377 | NA | NA | -0.4846574 | NA | NA | -0.6895291 | 0.4272099 | NA | NA | NA | 0.4727468 | -0.4844902 | NA | NA | NA | NA | NA | NA | NA | NA |
| provenance\_MAP | NA | NA | NA | NA | NA | -0.3853224 | NA | NA | NA | NA | NA | 0.2751622 | NA | NA | NA | NA | -0.2962824 | NA | NA | NA | NA | NA | NA | NA | NA | NA | NA | NA | NA | NA |
| provenance\_TD | NA | NA | NA | NA | NA | NA | NA | NA | NA | NA | NA | NA | NA | NA | NA | 0.4892213 | NA | NA | NA | 0.4058783 | NA | NA | NA | NA | NA | NA | NA | 0.4298732 | NA | NA |

```
kable(res.lr$pvals, label = 'Unadjusted p-values for relationships between plasticity and climate, controlling for ancestry' )
```

|  | plasticity\_rdpi\_growing\_season\_days\_2023 | plasticity\_rdpi\_growing\_season\_days\_2024 | plasticity\_rdpi\_DOY\_Stage2\_2023 | plasticity\_rdpi\_DOY\_Stage2\_2024 | plasticity\_rdpi\_DOY\_Stage3\_2023 | plasticity\_rdpi\_DOY\_Stage3\_2024 | plasticity\_rdpi\_DOY\_Stage6\_2023 | plasticity\_rdpi\_DOY\_Stage6\_2024 | plasticity\_rdpi\_DOY\_Stage7\_2023 | plasticity\_rdpi\_DOY\_Stage7\_2024 | plasticity\_rdpi\_stage7\_presence\_2023 | plasticity\_rdpi\_stage7\_presence\_2024 | plasticity\_rdpi\_DOY\_last\_budset\_2023 | plasticity\_rdpi\_DOY\_last\_budset\_2024 | plasticity\_rdpi\_leaf\_thickness\_avg\_mm\_2023 | plasticity\_rdpi\_leaf\_area\_cm2\_2023\_log | plasticity\_rdpi\_leaf\_mass\_g\_2023\_log | plasticity\_rdpi\_LMA\_g\_m2\_2023 | plasticity\_rdpi\_lower\_stomata\_pore\_length\_mean\_um | plasticity\_rdpi\_upper\_stomata\_pore\_length\_mean\_um | plasticity\_rdpi\_lower\_stomata\_density\_mm2 | plasticity\_rdpi\_upper\_stomata\_density\_mm2\_log | plasticity\_rdpi\_upper\_stomata\_presence | plasticity\_rdpi\_upper\_stomata\_density\_over\_total\_density\_log | plasticity\_rdpi\_licor\_gsw | plasticity\_rdpi\_licor\_ETR | plasticity\_rdpi\_licor\_Fm. | plasticity\_rdpi\_licor\_Fs | plasticity\_rdpi\_licor\_gbw | plasticity\_rdpi\_licor\_PhiPS2 |
| --- | --- | --- | --- | --- | --- | --- | --- | --- | --- | --- | --- | --- | --- | --- | --- | --- | --- | --- | --- | --- | --- | --- | --- | --- | --- | --- | --- | --- | --- | --- |
| genetic\_PC1 | 0.2469982 | 0.0252504 | 0.9084763 | 0.8499890 | 0.7796268 | 0.4118649 | 0.0107384 | 0.9134285 | 0.1704821 | 0.8911584 | 0.6104333 | 0.0000595 | 0.9751263 | 0.0686096 | 0.1567162 | 0.6094436 | 0.0513106 | 0.0130491 | 0.7972079 | 0.5053315 | 0.7458443 | 0.6280486 | 0.1472082 | 0.2245352 | 0.0345122 | 0.7345085 | 0.2004036 | 0.0856883 | 0.7911926 | 0.7797861 |
| genetic\_PC2 | 0.6993202 | 0.0422736 | 0.6994469 | 0.9055115 | 0.7998917 | 0.6620880 | 0.5399195 | 0.8875861 | 0.6210335 | 0.7118415 | 0.4502624 | 0.0652158 | 0.4176835 | 0.0595752 | 0.1793199 | 0.0595235 | 0.0853877 | 0.7472239 | 0.9011761 | 0.2286717 | 0.1057205 | 0.1948341 | 0.8352175 | 0.1269237 | 0.6860710 | 0.4333528 | 0.2003594 | 0.3778722 | 0.9400248 | 0.9104577 |
| genetic\_PC3 | 0.7622530 | 0.5784690 | 0.9709885 | 0.5638739 | 0.6008042 | 0.1991465 | 0.0399748 | 0.1598448 | 0.1676754 | 0.1393539 | 0.5678659 | 0.0746193 | 0.3091923 | 0.4169055 | 0.4172147 | 0.0043732 | 0.8180812 | 0.5926732 | 0.0704063 | 0.5902914 | 0.1742152 | 0.1668114 | 0.0561789 | 0.5402112 | 0.8225554 | 0.3991904 | 0.1821544 | 0.0398158 | 0.8813375 | 0.7683099 |
| provenance\_MAT | 0.6550132 | 0.1880717 | 0.9248281 | 0.3609891 | 0.4676382 | 0.5029581 | 0.0048189 | 0.3623216 | 0.0723359 | 0.0671355 | 0.8749584 | 0.0002975 | 0.0565226 | 0.0758352 | 0.1161709 | 0.0043310 | 0.7714620 | 0.3329415 | 0.9237109 | 0.7559896 | 0.1349067 | 0.1799939 | 0.5052217 | 0.1691435 | 0.8771367 | 0.6431227 | 0.1946891 | 0.0171541 | 0.5376805 | 0.9366383 |
| provenance\_MAP | 0.4826572 | 0.2126431 | 0.3436500 | 0.1385876 | 0.1827363 | 0.0086521 | 0.5076394 | 0.2325832 | 0.2271989 | 0.2210758 | 0.8614147 | 0.4179876 | 0.9114365 | 0.2308526 | 0.7622222 | 0.9323423 | 0.0196605 | 0.7309812 | 0.2248837 | 0.7682087 | 0.2740728 | 0.2330679 | 0.2230088 | 0.1447663 | 0.5147273 | 0.4809547 | 0.8976204 | 0.5620363 | 0.4284091 | 0.9343821 |
| provenance\_TD | 0.6657679 | 0.4444357 | 0.3301425 | 0.7251629 | 0.3242830 | 0.4684258 | 0.0709443 | 0.3952220 | 0.7626314 | 0.1167792 | 0.5184805 | 0.0285222 | 0.2413421 | 0.2507173 | 0.4964683 | 0.0281670 | 0.1343058 | 0.0301413 | 0.9869516 | 0.2420430 | 0.9229273 | 0.7752373 | 0.1955789 | 0.8398487 | 0.5788266 | 0.9401834 | 0.6116731 | 0.0051611 | 0.8908194 | 0.2399055 |

```
kable(res.lr$padj, label = 'Adjusted p-values for relationships between plasticity and climate, controlling for ancestry' )
```

|  | growing\_season\_days\_2023 | growing\_season\_days\_2024 | DOY\_Stage2\_2023 | DOY\_Stage2\_2024 | DOY\_Stage3\_2023 | DOY\_Stage3\_2024 | DOY\_Stage6\_2023 | DOY\_Stage6\_2024 | DOY\_Stage7\_2023 | DOY\_Stage7\_2024 | stage7\_presence\_2023 | stage7\_presence\_2024 | DOY\_last\_budset\_2023 | DOY\_last\_budset\_2024 | leaf\_thickness\_avg\_mm\_2023 | leaf\_area\_cm2\_2023\_log | leaf\_mass\_g\_2023\_log | LMA\_g\_m2\_2023 | lower\_stomata\_pore\_length\_mean\_um | upper\_stomata\_pore\_length\_mean\_um | lower\_stomata\_density\_mm2 | upper\_stomata\_density\_mm2\_log | upper\_stomata\_presence | upper\_stomata\_density\_over\_total\_density\_log | licor\_gsw | licor\_ETR | licor\_Fm. | licor\_Fs | licor\_gbw | licor\_PhiPS2 |
| --- | --- | --- | --- | --- | --- | --- | --- | --- | --- | --- | --- | --- | --- | --- | --- | --- | --- | --- | --- | --- | --- | --- | --- | --- | --- | --- | --- | --- | --- | --- |
| genetic\_PC1 | 0.5849958 | 0.3616961 | 0.9561187 | 0.9561187 | 0.9548401 | 0.8090082 | 0.2416146 | 0.9561187 | 0.5809032 | 0.9561187 | 0.9024685 | 0.0107027 | 0.9805739 | 0.4265731 | 0.5809032 | 0.9024685 | 0.4265731 | 0.2609820 | 0.9561187 | 0.8620292 | 0.9548401 | 0.9116834 | 0.5809032 | 0.5809032 | 0.3882625 | 0.9548401 | 0.5809032 | 0.4536439 | 0.9561187 | 0.9548401 |
| genetic\_PC2 | 0.9548401 | 0.4004869 | 0.9548401 | 0.9561187 | 0.9561187 | 0.9362362 | 0.8760182 | 0.9561187 | 0.9088295 | 0.9548401 | 0.8355384 | 0.4265731 | 0.8090082 | 0.4265731 | 0.5809032 | 0.4265731 | 0.4536439 | 0.9548401 | 0.9561187 | 0.5809032 | 0.5437052 | 0.5809032 | 0.9561187 | 0.5809032 | 0.9548401 | 0.8210894 | 0.5809032 | 0.7908953 | 0.9561187 | 0.9561187 |
| genetic\_PC3 | 0.9548401 | 0.8981792 | 0.9805739 | 0.8966304 | 0.9024685 | 0.5809032 | 0.3997485 | 0.5809032 | 0.5809032 | 0.5809032 | 0.8966304 | 0.4265731 | 0.7044889 | 0.8090082 | 0.8090082 | 0.1548324 | 0.9561187 | 0.9024685 | 0.4265731 | 0.9024685 | 0.5809032 | 0.5809032 | 0.4265731 | 0.8760182 | 0.9561187 | 0.8090082 | 0.5809032 | 0.3997485 | 0.9561187 | 0.9548401 |
| provenance\_MAT | 0.9357331 | 0.5809032 | 0.9561187 | 0.7672692 | 0.8516832 | 0.8620292 | 0.1548324 | 0.7672692 | 0.4265731 | 0.4265731 | 0.9561187 | 0.0267748 | 0.4265731 | 0.4265731 | 0.5681149 | 0.1548324 | 0.9548401 | 0.7308471 | 0.9561187 | 0.9548401 | 0.5809032 | 0.5809032 | 0.8620292 | 0.5809032 | 0.9561187 | 0.9260966 | 0.5809032 | 0.3087744 | 0.8760182 | 0.9561187 |
| provenance\_MAP | 0.8601811 | 0.5809032 | 0.7452651 | 0.5809032 | 0.5809032 | 0.2224833 | 0.8620292 | 0.5809032 | 0.5809032 | 0.5809032 | 0.9561187 | 0.8090082 | 0.9561187 | 0.5809032 | 0.9548401 | 0.9561187 | 0.3217174 | 0.9548401 | 0.5809032 | 0.9548401 | 0.6324758 | 0.5809032 | 0.5809032 | 0.5809032 | 0.8641342 | 0.8601811 | 0.9561187 | 0.8966304 | 0.8203579 | 0.9561187 |
| provenance\_TD | 0.9362362 | 0.8333170 | 0.7308471 | 0.9548401 | 0.7296366 | 0.8516832 | 0.4265731 | 0.8090082 | 0.9548401 | 0.5681149 | 0.8641342 | 0.3616961 | 0.5809032 | 0.5860923 | 0.8620292 | 0.3616961 | 0.5809032 | 0.3616961 | 0.9869516 | 0.5809032 | 0.9561187 | 0.9548401 | 0.5809032 | 0.9561187 | 0.8981792 | 0.9561187 | 0.9024685 | 0.1548324 | 0.9561187 | 0.5809032 |

### 5 Heatmaps summarizing significant relationships

#### 5.1 Plot plasticity heatmaps together

```
########## plot together

# rename traits

trait_names <- colnames(res.lr$eff.std.sig)
trait_names <- gsub('plasticity_rdpi_', '', trait_names)
trait_names <- gsub('Stage2_cGDD', 'Stage2_Bud_Flush_cGDD', trait_names)
trait_names <- gsub('DOY_Stage2', 'Stage2_Bud_Flush_DOY', trait_names)
trait_names <- gsub('DOY_Stage3', 'Stage3_Leaf_Emergence_DOY', trait_names)
trait_names <- gsub('DOY_Stage6', 'Stage6_First_Budset_DOY', trait_names)
trait_names <- gsub('DOY_Stage7', 'Stage7_Lammas_Growth_DOY', trait_names)
trait_names <- gsub('stage7_presence', 'Stage7_Lammas_Growth_presence', trait_names)
trait_names <- gsub('DOY_Stage8', 'Stage8_Budset_after_Lammas_DOY', trait_names)
trait_names <- gsub('DOY_last_budset', 'Stage8_Final_Budset_DOY', trait_names)
trait_names <- gsub('growing_season_days', 'Growing_Season_Days', trait_names)
trait_names <- gsub('_', ' ', trait_names)

effect_names1 <- rownames(res.lr$eff.std.sig)
effect_names1 <- gsub('genetic', 'Genetic', effect_names1)
effect_names1 <- gsub('provenance', 'Home', effect_names1)
effect_names1 <- gsub('_', ' ', effect_names1)


########

#draw(hm2 + hm4, padding = unit(c(2, 30, 2, 15), "mm"))

# use same color scale for both heatmaps

# color scale based on all effect sizes
max_col <- max(c(unlist(res.lr$eff.std.sig), unlist(res.lr.pt$eff.std.sig)), na.rm = T)
min_col <- min(c(unlist(res.lr$eff.std.sig), unlist(res.lr.pt$eff.std.sig)), na.rm = T)

# make scale same at both ends
max_col <- max(abs(max_col), abs(min_col))
min_col <- min(abs(max_col), abs(min_col))*-1
max_ceil <- ceiling(max_col*10)/10 # take ceiling rounded to one decimal place - used for placing ticks on legend

colf <- colorRamp2(c('red', 'white', 'blue'), breaks = c(min_col, 0, max_col))


# standardized effect sizes for simple linear regressions

colnames(res.lr$eff.std.sig) <- gsub('plasticity_rdpi_', '', colnames(res.lr$eff.std.sig))

# row annotation - trait categories
cats <- c(rep('Phenology', 14),  rep('Leaf Morphology', 4), rep('Stomata', 7), rep('Photosynthesis', 5))
cbind(colnames(res.lr$cors.sig), cats)
```

```
##                                                      cats             
##  [1,] "growing_season_days_2023"                     "Phenology"      
##  [2,] "growing_season_days_2024"                     "Phenology"      
##  [3,] "DOY_Stage2_2023"                              "Phenology"      
##  [4,] "DOY_Stage2_2024"                              "Phenology"      
##  [5,] "DOY_Stage3_2023"                              "Phenology"      
##  [6,] "DOY_Stage3_2024"                              "Phenology"      
##  [7,] "DOY_Stage6_2023"                              "Phenology"      
##  [8,] "DOY_Stage6_2024"                              "Phenology"      
##  [9,] "DOY_Stage7_2023"                              "Phenology"      
## [10,] "DOY_Stage7_2024"                              "Phenology"      
## [11,] "stage7_presence_2023"                         "Phenology"      
## [12,] "stage7_presence_2024"                         "Phenology"      
## [13,] "DOY_last_budset_2023"                         "Phenology"      
## [14,] "DOY_last_budset_2024"                         "Phenology"      
## [15,] "leaf_thickness_avg_mm_2023"                   "Leaf Morphology"
## [16,] "leaf_area_cm2_2023_log"                       "Leaf Morphology"
## [17,] "leaf_mass_g_2023_log"                         "Leaf Morphology"
## [18,] "LMA_g_m2_2023"                                "Leaf Morphology"
## [19,] "lower_stomata_pore_length_mean_um"            "Stomata"        
## [20,] "upper_stomata_pore_length_mean_um"            "Stomata"        
## [21,] "lower_stomata_density_mm2"                    "Stomata"        
## [22,] "upper_stomata_density_mm2_log"                "Stomata"        
## [23,] "upper_stomata_presence"                       "Stomata"        
## [24,] "upper_stomata_density_over_total_density_log" "Stomata"        
## [25,] "licor_gsw"                                    "Stomata"        
## [26,] "licor_ETR"                                    "Photosynthesis" 
## [27,] "licor_Fm."                                    "Photosynthesis" 
## [28,] "licor_Fs"                                     "Photosynthesis" 
## [29,] "licor_gbw"                                    "Photosynthesis" 
## [30,] "licor_PhiPS2"                                 "Photosynthesis"
```

```
ha_row <- rowAnnotation(Trait_Category = cats,
                        show_annotation_name = F,
                            col = list(Trait_Category = c('Phenology' = "#e2e260", 'Photosynthesis' = "#2d4030", 'Leaf Morphology' = "#516823", 'Stomata' =  "#88a2b9")))

# cell text showing significance
cell_text <- res.lr$padj
cell_text[res.lr$padj > 0.05] <- ''
cell_text[res.lr$padj <= 0.05 & res.lr$padj > 0.01] <- '*'
cell_text[res.lr$padj <= 0.01 & res.lr$padj > 0.001] <- '**'
cell_text[res.lr$padj<= 0.001] <- '***'
cell_text <- t(cell_text)


hm.lr <- Heatmap(as.matrix(t(res.lr$eff.std.sig)),
        cluster_rows = F,
        cluster_columns = F,
        rect_gp = gpar(col = "black", lwd = 1),
        col = colf,
        na_col = 'grey80',
        row_names_side = 'left',
        column_names_side = 'top',
        column_names_rot = 90,
        column_names_centered = T,
        column_title = 'Plasticity Predictors',
        column_labels = effect_names1,
        column_title_gp = gpar(fontface = 'bold'),
        row_title_gp = gpar(fontface = 'bold'),
        row_title = 'Trait plasticity',
        row_labels = trait_names,
        left_annotation = ha_row,
        row_names_max_width = unit(8, 'cm'),
        heatmap_legend_param = list(title = 'Std. Effect Size', at = c(max_ceil*-1, 0, max_ceil)),
        cell_fun = function(j, i, x, y, width, height, fill) { # add text to each grid
                      grid.text(cell_text[i, j], x, y, gp = gpar(cex = 1.5))})

ComplexHeatmap::draw(hm.lr, padding = unit(c(2, 30, 2, 15), "mm"))
```

```
# linear regression, climate controlling for ancestry

effect_names2 <- rownames(res.lr.pt$eff.std.sig)
effect_names2 <- gsub('provenance', 'Home', effect_names2)
effect_names2 <- gsub('_', ' ', effect_names2)

# cell text showing significance
cell_text <- res.lr.pt$padj
cell_text[res.lr.pt$padj > 0.05] <- ''
cell_text[res.lr.pt$padj <= 0.05 & res.lr.pt$padj > 0.01] <- '*'
cell_text[res.lr.pt$padj <= 0.01 & res.lr.pt$padj > 0.001] <- '**'
cell_text[res.lr.pt$padj<= 0.001] <- '***'
cell_text <- t(cell_text)


hm.lr.pt <- Heatmap(as.matrix(t(res.lr.pt$eff.std.sig)),
        cluster_rows = F,
        cluster_columns = F,
        rect_gp = gpar(col = "black", lwd = 1),
        col = colf,
        na_col = 'grey80',
        row_names_side = 'left',
        column_names_side = 'top',
        column_names_rot = 90,
        column_names_centered = T,
        column_title = paste0('Plasticity Predictors', '\n', '(controlling for ancestry)'),
        column_labels = effect_names2,
        column_title_gp = gpar(fontface = 'bold'),
        row_title_gp = gpar(fontface = 'bold'),
        row_title = 'Trait plasticity',
        row_labels = trait_names,
        left_annotation = ha_row,
        #heatmap_legend_param = list(title = 'Std. Effect Size', at = c(max_ceil*-1, 0, max_ceil)),
        show_heatmap_legend = F, # legend is the same, so don't plot it
        cell_fun = function(j, i, x, y, width, height, fill) { # add text to each grid
                      grid.text(cell_text[i, j], x, y, gp = gpar(cex = 1.5))})

ComplexHeatmap::draw(hm.lr.pt, padding = unit(c(2, 30, 2, 15), "mm"))
```

```
#png(file = 'results/plasticity/heatmap_plasticity_vs_ancestry_and_climate_padjusted.png', height = 8, width = 10, res = 300, units = 'in')
draw(hm.lr + hm.lr.pt, padding = unit(c(2, 15, 2, 15), "mm"))
```

```
#dev.off()
```

#### 5.2 Merge with GxE heatmap

```
# gxe_hm
# GxE plot has cGDD but not presence/absences traits (stage7 and upper stomata)
load('results/plasticity/GxE_traits_heatmap_data.Rdata')


str(gxe_hm$effect)
```

```
## 'data.frame':    17 obs. of  31 variables:
##  $ growing_season_days_2023         : logi  NA NA NA NA NA NA ...
##  $ growing_season_days_2024         : num  NA NA NA NA -0.178 ...
##  $ DOY_Stage2_2023                  : num  -0.569 NA 0.159 NA NA ...
##  $ Stage2_cGDD_2023                 : num  0.64 NA NA NA NA ...
##  $ DOY_Stage2_2024                  : num  NA NA 0.19 NA NA ...
##  $ Stage2_cGDD_2024                 : num  NA NA 0.138 NA NA ...
##  $ DOY_Stage3_2023                  : num  -0.551 NA 0.171 NA NA ...
##  $ Stage3_cGDD_2023                 : num  0.709 NA NA NA NA ...
##  $ DOY_Stage3_2024                  : logi  NA NA NA NA NA NA ...
##  $ Stage3_cGDD_2024                 : logi  NA NA NA NA NA NA ...
##  $ DOY_Stage6_2023                  : num  NA NA 0.183 NA -0.118 ...
##  $ DOY_Stage6_2024                  : logi  NA NA NA NA NA NA ...
##  $ DOY_Stage7_2023                  : logi  NA NA NA NA NA NA ...
##  $ DOY_Stage7_2024                  : logi  NA NA NA NA NA NA ...
##  $ DOY_last_budset_2023             : num  NA NA 0.131 NA NA ...
##  $ DOY_last_budset_2024             : num  NA NA NA NA -0.185 ...
##  $ leaf_thickness_avg_mm_2023       : num  NA NA NA NA NA ...
##  $ leaf_area_cm2_2023               : num  0.4509 -0.4741 NA NA -0.0995 ...
##  $ leaf_mass_g_2023                 : num  0.434 -0.471 0.127 NA NA ...
##  $ LMA_g_m2_2023                    : num  NA NA NA NA NA NA NA NA NA NA ...
##  $ lower_stomata_pore_length_mean_um: num  NA NA NA NA NA ...
##  $ upper_stomata_pore_length_mean_um: num  NA NA NA 0.239 NA ...
##  $ lower_stomata_density_mm2        : num  NA NA NA NA 0.246 ...
##  $ upper_stomata_density_mm2        : num  NA 0.254 NA NA NA ...
##  $ licor_gsw                        : num  NA NA NA NA NA NA NA NA NA NA ...
##  $ stomata_ratio                    : num  NA NA NA NA NA ...
##  $ licor_ETR                        : logi  NA NA NA NA NA NA ...
##  $ licor_Fm.                        : num  NA NA NA NA NA NA NA NA NA NA ...
##  $ licor_Fs                         : num  NA NA NA NA NA ...
##  $ licor_gbw                        : num  NA NA NA NA NA ...
##  $ licor_PhiPS2                     : num  NA NA NA NA NA NA NA NA NA NA ...
```

```
str(res.lr$eff.std.sig)
```

```
## 'data.frame':    6 obs. of  30 variables:
##  $ growing_season_days_2023                    : logi  NA NA NA NA NA NA
##  $ growing_season_days_2024                    : num  0.345 0.315 NA NA NA ...
##  $ DOY_Stage2_2023                             : logi  NA NA NA NA NA NA
##  $ DOY_Stage2_2024                             : logi  NA NA NA NA NA NA
##  $ DOY_Stage3_2023                             : logi  NA NA NA NA NA NA
##  $ DOY_Stage3_2024                             : num  NA NA NA NA -0.396 ...
##  $ DOY_Stage6_2023                             : num  0.381 NA 0.311 -0.417 NA ...
##  $ DOY_Stage6_2024                             : logi  NA NA NA NA NA NA
##  $ DOY_Stage7_2023                             : logi  NA NA NA NA NA NA
##  $ DOY_Stage7_2024                             : logi  NA NA NA NA NA NA
##  $ stage7_presence_2023                        : logi  NA NA NA NA NA NA
##  $ stage7_presence_2024                        : num  0.573 NA NA -0.525 NA ...
##  $ DOY_last_budset_2023                        : logi  NA NA NA NA NA NA
##  $ DOY_last_budset_2024                        : logi  NA NA NA NA NA NA
##  $ leaf_thickness_avg_mm_2023                  : logi  NA NA NA NA NA NA
##  $ leaf_area_cm2_2023_log                      : num  NA NA 0.422 -0.422 NA ...
##  $ leaf_mass_g_2023_log                        : num  NA NA NA NA -0.351 ...
##  $ LMA_g_m2_2023                               : num  0.371 NA NA NA NA ...
##  $ lower_stomata_pore_length_mean_um           : logi  NA NA NA NA NA NA
##  $ upper_stomata_pore_length_mean_um           : logi  NA NA NA NA NA NA
##  $ lower_stomata_density_mm2                   : logi  NA NA NA NA NA NA
##  $ upper_stomata_density_mm2_log               : logi  NA NA NA NA NA NA
##  $ upper_stomata_presence                      : logi  NA NA NA NA NA NA
##  $ upper_stomata_density_over_total_density_log: logi  NA NA NA NA NA NA
##  $ licor_gsw                                   : num  0.319 NA NA NA NA ...
##  $ licor_ETR                                   : logi  NA NA NA NA NA NA
##  $ licor_Fm.                                   : logi  NA NA NA NA NA NA
##  $ licor_Fs                                    : num  NA NA 0.311 -0.358 NA ...
##  $ licor_gbw                                   : logi  NA NA NA NA NA NA
##  $ licor_PhiPS2                                : logi  NA NA NA NA NA NA
```

```
# effect size dataframes
df1 <- gxe_hm$effect
df2 <- res.lr$eff.std.sig
df3 <- res.lr.pt$eff.std.sig

dim(df1)
```

```
## [1] 17 31
```

```
dim(df2)
```

```
## [1]  6 30
```

```
dim(df3)
```

```
## [1]  3 30
```

```
# pvalue dataframes
pval1 <- gxe_hm$pval_adj
pval2 <- res.lr$padj
pval3 <- res.lr.pt$padj

# whitch traits don't match between the two datasets?
colnames(df2)[! colnames(df2) %in% colnames(df1)]
```

```
## [1] "stage7_presence_2023"                        
## [2] "stage7_presence_2024"                        
## [3] "leaf_area_cm2_2023_log"                      
## [4] "leaf_mass_g_2023_log"                        
## [5] "upper_stomata_density_mm2_log"               
## [6] "upper_stomata_presence"                      
## [7] "upper_stomata_density_over_total_density_log"
```

```
colnames(df1)[! colnames(df1) %in% colnames(df2)]
```

```
## [1] "Stage2_cGDD_2023"          "Stage2_cGDD_2024"         
## [3] "Stage3_cGDD_2023"          "Stage3_cGDD_2024"         
## [5] "leaf_area_cm2_2023"        "leaf_mass_g_2023"         
## [7] "upper_stomata_density_mm2" "stomata_ratio"
```

```
# remove plasticity from column names
colnames(df2) <- gsub('plasticity_rdpi_', '', colnames(df2))
colnames(df3) <- gsub('plasticity_rdpi_', '', colnames(df3))
colnames(pval2) <- gsub('plasticity_rdpi_', '', colnames(pval2))
colnames(pval3) <- gsub('plasticity_rdpi_', '', colnames(pval3))

# insert empty column for stage 7 presence in GxE heatmap
df1$stage7_presence_2023 <- NA
df1$stage7_presence_2024 <- NA
df1$upper_stomata_presence  <- NA

pval1$stage7_presence_2023 <- NA
pval1$stage7_presence_2024 <- NA
pval1$upper_stomata_presence  <- NA

# insert empty column for GDD for plasticity heatmap

df2$Stage2_cGDD_2023 <- NA
df2$Stage2_cGDD_2024 <- NA
df2$Stage3_cGDD_2023 <- NA
df2$Stage3_cGDD_2024 <- NA

df3$Stage2_cGDD_2023 <- NA
df3$Stage2_cGDD_2024 <- NA
df3$Stage3_cGDD_2023 <- NA
df3$Stage3_cGDD_2024 <- NA

# pvals
pval2$Stage2_cGDD_2023 <- NA
pval2$Stage2_cGDD_2024 <- NA
pval2$Stage3_cGDD_2023 <- NA
pval2$Stage3_cGDD_2024 <- NA

pval3$Stage2_cGDD_2023 <- NA
pval3$Stage2_cGDD_2024 <- NA
pval3$Stage3_cGDD_2023 <- NA
pval3$Stage3_cGDD_2024 <- NA

cbind(colnames(df1), colnames(df2))
```

```
##       [,1]                               
##  [1,] "growing_season_days_2023"         
##  [2,] "growing_season_days_2024"         
##  [3,] "DOY_Stage2_2023"                  
##  [4,] "Stage2_cGDD_2023"                 
##  [5,] "DOY_Stage2_2024"                  
##  [6,] "Stage2_cGDD_2024"                 
##  [7,] "DOY_Stage3_2023"                  
##  [8,] "Stage3_cGDD_2023"                 
##  [9,] "DOY_Stage3_2024"                  
## [10,] "Stage3_cGDD_2024"                 
## [11,] "DOY_Stage6_2023"                  
## [12,] "DOY_Stage6_2024"                  
## [13,] "DOY_Stage7_2023"                  
## [14,] "DOY_Stage7_2024"                  
## [15,] "DOY_last_budset_2023"             
## [16,] "DOY_last_budset_2024"             
## [17,] "leaf_thickness_avg_mm_2023"       
## [18,] "leaf_area_cm2_2023"               
## [19,] "leaf_mass_g_2023"                 
## [20,] "LMA_g_m2_2023"                    
## [21,] "lower_stomata_pore_length_mean_um"
## [22,] "upper_stomata_pore_length_mean_um"
## [23,] "lower_stomata_density_mm2"        
## [24,] "upper_stomata_density_mm2"        
## [25,] "licor_gsw"                        
## [26,] "stomata_ratio"                    
## [27,] "licor_ETR"                        
## [28,] "licor_Fm."                        
## [29,] "licor_Fs"                         
## [30,] "licor_gbw"                        
## [31,] "licor_PhiPS2"                     
## [32,] "stage7_presence_2023"             
## [33,] "stage7_presence_2024"             
## [34,] "upper_stomata_presence"           
##       [,2]                                          
##  [1,] "growing_season_days_2023"                    
##  [2,] "growing_season_days_2024"                    
##  [3,] "DOY_Stage2_2023"                             
##  [4,] "DOY_Stage2_2024"                             
##  [5,] "DOY_Stage3_2023"                             
##  [6,] "DOY_Stage3_2024"                             
##  [7,] "DOY_Stage6_2023"                             
##  [8,] "DOY_Stage6_2024"                             
##  [9,] "DOY_Stage7_2023"                             
## [10,] "DOY_Stage7_2024"                             
## [11,] "stage7_presence_2023"                        
## [12,] "stage7_presence_2024"                        
## [13,] "DOY_last_budset_2023"                        
## [14,] "DOY_last_budset_2024"                        
## [15,] "leaf_thickness_avg_mm_2023"                  
## [16,] "leaf_area_cm2_2023_log"                      
## [17,] "leaf_mass_g_2023_log"                        
## [18,] "LMA_g_m2_2023"                               
## [19,] "lower_stomata_pore_length_mean_um"           
## [20,] "upper_stomata_pore_length_mean_um"           
## [21,] "lower_stomata_density_mm2"                   
## [22,] "upper_stomata_density_mm2_log"               
## [23,] "upper_stomata_presence"                      
## [24,] "upper_stomata_density_over_total_density_log"
## [25,] "licor_gsw"                                   
## [26,] "licor_ETR"                                   
## [27,] "licor_Fm."                                   
## [28,] "licor_Fs"                                    
## [29,] "licor_gbw"                                   
## [30,] "licor_PhiPS2"                                
## [31,] "Stage2_cGDD_2023"                            
## [32,] "Stage2_cGDD_2024"                            
## [33,] "Stage3_cGDD_2023"                            
## [34,] "Stage3_cGDD_2024"
```

```
# reorder

dput(colnames(df1))
```

```
## c("growing_season_days_2023", "growing_season_days_2024", "DOY_Stage2_2023", 
## "Stage2_cGDD_2023", "DOY_Stage2_2024", "Stage2_cGDD_2024", "DOY_Stage3_2023", 
## "Stage3_cGDD_2023", "DOY_Stage3_2024", "Stage3_cGDD_2024", "DOY_Stage6_2023", 
## "DOY_Stage6_2024", "DOY_Stage7_2023", "DOY_Stage7_2024", "DOY_last_budset_2023", 
## "DOY_last_budset_2024", "leaf_thickness_avg_mm_2023", "leaf_area_cm2_2023", 
## "leaf_mass_g_2023", "LMA_g_m2_2023", "lower_stomata_pore_length_mean_um", 
## "upper_stomata_pore_length_mean_um", "lower_stomata_density_mm2", 
## "upper_stomata_density_mm2", "licor_gsw", "stomata_ratio", "licor_ETR", 
## "licor_Fm.", "licor_Fs", "licor_gbw", "licor_PhiPS2", "stage7_presence_2023", 
## "stage7_presence_2024", "upper_stomata_presence")
```

```
ord1 <- c("growing_season_days_2023", "growing_season_days_2024", "DOY_Stage2_2023", 
"Stage2_cGDD_2023", "DOY_Stage2_2024", "Stage2_cGDD_2024", "DOY_Stage3_2023", 
"Stage3_cGDD_2023", "DOY_Stage3_2024", "Stage3_cGDD_2024", "DOY_Stage6_2023", 
"DOY_Stage6_2024", "DOY_Stage7_2023", "DOY_Stage7_2024", "stage7_presence_2023", 
"stage7_presence_2024", "DOY_last_budset_2023", 
"DOY_last_budset_2024", "leaf_thickness_avg_mm_2023", "leaf_area_cm2_2023", 
"leaf_mass_g_2023", "LMA_g_m2_2023", "lower_stomata_pore_length_mean_um", 
"upper_stomata_pore_length_mean_um", "lower_stomata_density_mm2", 
"upper_stomata_density_mm2", "upper_stomata_presence", "stomata_ratio", "licor_gsw", "licor_ETR", "licor_Fm.", 
"licor_Fs", "licor_gbw", "licor_PhiPS2")


dput(colnames(df2))
```

```
## c("growing_season_days_2023", "growing_season_days_2024", "DOY_Stage2_2023", 
## "DOY_Stage2_2024", "DOY_Stage3_2023", "DOY_Stage3_2024", "DOY_Stage6_2023", 
## "DOY_Stage6_2024", "DOY_Stage7_2023", "DOY_Stage7_2024", "stage7_presence_2023", 
## "stage7_presence_2024", "DOY_last_budset_2023", "DOY_last_budset_2024", 
## "leaf_thickness_avg_mm_2023", "leaf_area_cm2_2023_log", "leaf_mass_g_2023_log", 
## "LMA_g_m2_2023", "lower_stomata_pore_length_mean_um", "upper_stomata_pore_length_mean_um", 
## "lower_stomata_density_mm2", "upper_stomata_density_mm2_log", 
## "upper_stomata_presence", "upper_stomata_density_over_total_density_log", 
## "licor_gsw", "licor_ETR", "licor_Fm.", "licor_Fs", "licor_gbw", 
## "licor_PhiPS2", "Stage2_cGDD_2023", "Stage2_cGDD_2024", "Stage3_cGDD_2023", 
## "Stage3_cGDD_2024")
```

```
ord2 <- c("growing_season_days_2023", "growing_season_days_2024", "DOY_Stage2_2023", "Stage2_cGDD_2023",
"DOY_Stage2_2024",  "Stage2_cGDD_2024", "DOY_Stage3_2023","Stage3_cGDD_2023", "DOY_Stage3_2024", "Stage3_cGDD_2024", "DOY_Stage6_2023", 
"DOY_Stage6_2024", "DOY_Stage7_2023", "DOY_Stage7_2024", "stage7_presence_2023", 
"stage7_presence_2024", "DOY_last_budset_2023", "DOY_last_budset_2024", 
"leaf_thickness_avg_mm_2023", "leaf_area_cm2_2023_log", "leaf_mass_g_2023_log",
"LMA_g_m2_2023", "lower_stomata_pore_length_mean_um", "upper_stomata_pore_length_mean_um", 
"lower_stomata_density_mm2", "upper_stomata_density_mm2_log", 
"upper_stomata_presence", "upper_stomata_density_over_total_density_log", "licor_gsw", 
"licor_ETR", "licor_Fm.", "licor_Fs", "licor_gbw", 
"licor_PhiPS2")

# check that they are now in the same order
cbind(ord1, ord2)
```

```
##       ord1                               
##  [1,] "growing_season_days_2023"         
##  [2,] "growing_season_days_2024"         
##  [3,] "DOY_Stage2_2023"                  
##  [4,] "Stage2_cGDD_2023"                 
##  [5,] "DOY_Stage2_2024"                  
##  [6,] "Stage2_cGDD_2024"                 
##  [7,] "DOY_Stage3_2023"                  
##  [8,] "Stage3_cGDD_2023"                 
##  [9,] "DOY_Stage3_2024"                  
## [10,] "Stage3_cGDD_2024"                 
## [11,] "DOY_Stage6_2023"                  
## [12,] "DOY_Stage6_2024"                  
## [13,] "DOY_Stage7_2023"                  
## [14,] "DOY_Stage7_2024"                  
## [15,] "stage7_presence_2023"             
## [16,] "stage7_presence_2024"             
## [17,] "DOY_last_budset_2023"             
## [18,] "DOY_last_budset_2024"             
## [19,] "leaf_thickness_avg_mm_2023"       
## [20,] "leaf_area_cm2_2023"               
## [21,] "leaf_mass_g_2023"                 
## [22,] "LMA_g_m2_2023"                    
## [23,] "lower_stomata_pore_length_mean_um"
## [24,] "upper_stomata_pore_length_mean_um"
## [25,] "lower_stomata_density_mm2"        
## [26,] "upper_stomata_density_mm2"        
## [27,] "upper_stomata_presence"           
## [28,] "stomata_ratio"                    
## [29,] "licor_gsw"                        
## [30,] "licor_ETR"                        
## [31,] "licor_Fm."                        
## [32,] "licor_Fs"                         
## [33,] "licor_gbw"                        
## [34,] "licor_PhiPS2"                     
##       ord2                                          
##  [1,] "growing_season_days_2023"                    
##  [2,] "growing_season_days_2024"                    
##  [3,] "DOY_Stage2_2023"                             
##  [4,] "Stage2_cGDD_2023"                            
##  [5,] "DOY_Stage2_2024"                             
##  [6,] "Stage2_cGDD_2024"                            
##  [7,] "DOY_Stage3_2023"                             
##  [8,] "Stage3_cGDD_2023"                            
##  [9,] "DOY_Stage3_2024"                             
## [10,] "Stage3_cGDD_2024"                            
## [11,] "DOY_Stage6_2023"                             
## [12,] "DOY_Stage6_2024"                             
## [13,] "DOY_Stage7_2023"                             
## [14,] "DOY_Stage7_2024"                             
## [15,] "stage7_presence_2023"                        
## [16,] "stage7_presence_2024"                        
## [17,] "DOY_last_budset_2023"                        
## [18,] "DOY_last_budset_2024"                        
## [19,] "leaf_thickness_avg_mm_2023"                  
## [20,] "leaf_area_cm2_2023_log"                      
## [21,] "leaf_mass_g_2023_log"                        
## [22,] "LMA_g_m2_2023"                               
## [23,] "lower_stomata_pore_length_mean_um"           
## [24,] "upper_stomata_pore_length_mean_um"           
## [25,] "lower_stomata_density_mm2"                   
## [26,] "upper_stomata_density_mm2_log"               
## [27,] "upper_stomata_presence"                      
## [28,] "upper_stomata_density_over_total_density_log"
## [29,] "licor_gsw"                                   
## [30,] "licor_ETR"                                   
## [31,] "licor_Fm."                                   
## [32,] "licor_Fs"                                    
## [33,] "licor_gbw"                                   
## [34,] "licor_PhiPS2"
```

```
# reorder
df1 <- df1[,ord1]
pval1 <- pval1[,ord1]

df2 <- df2[,ord2]
pval2 <- pval2[,ord2]
df3 <- df3[,ord2]
pval3 <- pval3[,ord2]
```

#### 5.3 All heatmaps

```
# heatmap params
text_cex <- 1
rot <- 90 # angle for column labels

leg_title <- 14 # font size for legend title
leg_label <- 12 # font size for legend labels

########################################################
# GxE

# rename traits

trait_names <- colnames(df1)
trait_names <- gsub('Stage2_cGDD', 'Stage2_Bud_Flush_cGDD', trait_names)
trait_names <- gsub('DOY_Stage2', 'Stage2_Bud_Flush_DOY', trait_names)
trait_names <- gsub('DOY_Stage3', 'Stage3_Leaf_Emergence_DOY', trait_names)
trait_names <- gsub('DOY_Stage6', 'Stage6_First_Budset_DOY', trait_names)
trait_names <- gsub('DOY_Stage7', 'Stage7_Lammas_Growth_DOY', trait_names)
#trait_names <- gsub('stage7_presence', 'Stage7_Lammas_Growth_presence', trait_names)
trait_names <- gsub('DOY_Stage8', 'Stage8_Budset_after_Lammas_DOY', trait_names)
trait_names <- gsub('DOY_last_budset', 'Stage8_Final_Budset_DOY', trait_names)
trait_names <- gsub('growing_season_days', 'Growing_Season_Days', trait_names)

trait_names <- gsub('licor_', '', trait_names)
trait_names <- gsub('Fm.', "Fm'", trait_names)
trait_names <- gsub('upper', "Adaxial", trait_names)
trait_names <- gsub('lower', "Abaxial", trait_names)
trait_names <- gsub('stomata_ratio', "Stomata_ratio", trait_names)
trait_names <- gsub('leaf', 'Leaf', trait_names)

# remove units
trait_names <- gsub('_cm2', '', trait_names)
trait_names <- gsub('_cm', '', trait_names)
trait_names <- gsub('_mm2', '', trait_names)
trait_names <- gsub('_m2', '', trait_names)
trait_names <- gsub('_mm', '', trait_names)
trait_names <- gsub('_um', '', trait_names)
trait_names <- gsub('_g', '', trait_names)

trait_names <- gsub('_', ' ', trait_names)

# check
cbind(colnames(df1), trait_names)
```

```
##                                           trait_names                       
##  [1,] "growing_season_days_2023"          "Growing Season Days 2023"        
##  [2,] "growing_season_days_2024"          "Growing Season Days 2024"        
##  [3,] "DOY_Stage2_2023"                   "Stage2 Bud Flush DOY 2023"       
##  [4,] "Stage2_cGDD_2023"                  "Stage2 Bud Flush cGDD 2023"      
##  [5,] "DOY_Stage2_2024"                   "Stage2 Bud Flush DOY 2024"       
##  [6,] "Stage2_cGDD_2024"                  "Stage2 Bud Flush cGDD 2024"      
##  [7,] "DOY_Stage3_2023"                   "Stage3 Leaf Emergence DOY 2023"  
##  [8,] "Stage3_cGDD_2023"                  "Stage3 cGDD 2023"                
##  [9,] "DOY_Stage3_2024"                   "Stage3 Leaf Emergence DOY 2024"  
## [10,] "Stage3_cGDD_2024"                  "Stage3 cGDD 2024"                
## [11,] "DOY_Stage6_2023"                   "Stage6 First Budset DOY 2023"    
## [12,] "DOY_Stage6_2024"                   "Stage6 First Budset DOY 2024"    
## [13,] "DOY_Stage7_2023"                   "Stage7 Lammas Growth DOY 2023"   
## [14,] "DOY_Stage7_2024"                   "Stage7 Lammas Growth DOY 2024"   
## [15,] "stage7_presence_2023"              "stage7 presence 2023"            
## [16,] "stage7_presence_2024"              "stage7 presence 2024"            
## [17,] "DOY_last_budset_2023"              "Stage8 Final Budset DOY 2023"    
## [18,] "DOY_last_budset_2024"              "Stage8 Final Budset DOY 2024"    
## [19,] "leaf_thickness_avg_mm_2023"        "Leaf thickness avg 2023"         
## [20,] "leaf_area_cm2_2023"                "Leaf area 2023"                  
## [21,] "leaf_mass_g_2023"                  "Leaf mass 2023"                  
## [22,] "LMA_g_m2_2023"                     "LMA 2023"                        
## [23,] "lower_stomata_pore_length_mean_um" "Abaxial stomata pore length mean"
## [24,] "upper_stomata_pore_length_mean_um" "Adaxial stomata pore length mean"
## [25,] "lower_stomata_density_mm2"         "Abaxial stomata density"         
## [26,] "upper_stomata_density_mm2"         "Adaxial stomata density"         
## [27,] "upper_stomata_presence"            "Adaxial stomata presence"        
## [28,] "stomata_ratio"                     "Stomata ratio"                   
## [29,] "licor_gsw"                         "gsw"                             
## [30,] "licor_ETR"                         "ETR"                             
## [31,] "licor_Fm."                         "Fm'"                             
## [32,] "licor_Fs"                          "Fs"                              
## [33,] "licor_gbw"                         "gbw"                             
## [34,] "licor_PhiPS2"                      "PhiPS2"
```

```
# rename effects
effect_names <- rownames(df1)
effect_names <- gsub('home', 'Home', effect_names)
effect_names <- gsub('garden', 'Garden', effect_names)
effect_names <- gsub('pc', 'Genetic PC', effect_names)
effect_names <- gsub(':', ' x ', effect_names)
effect_names <- gsub('_', ' ', effect_names)
# check
cbind(rownames(df1), effect_names)
```

```
##                                effect_names              
##  [1,] "garden_MAT"             "Garden MAT"              
##  [2,] "garden_MAP"             "Garden MAP"              
##  [3,] "home_MAT"               "Home MAT"                
##  [4,] "home_MAP"               "Home MAP"                
##  [5,] "genetic_PC1"            "genetic PC1"             
##  [6,] "genetic_PC2"            "genetic PC2"             
##  [7,] "genetic_PC3"            "genetic PC3"             
##  [8,] "garden_MAT:home_MAT"    "Garden MAT x Home MAT"   
##  [9,] "garden_MAT:home_MAP"    "Garden MAT x Home MAP"   
## [10,] "home_MAP:garden_MAP"    "Home MAP x Garden MAP"   
## [11,] "home_MAT:garden_MAP"    "Home MAT x Garden MAP"   
## [12,] "garden_MAT:genetic_PC1" "Garden MAT x genetic PC1"
## [13,] "garden_MAT:genetic_PC2" "Garden MAT x genetic PC2"
## [14,] "garden_MAT:genetic_PC3" "Garden MAT x genetic PC3"
## [15,] "garden_MAP:genetic_PC1" "Garden MAP x genetic PC1"
## [16,] "garden_MAP:genetic_PC2" "Garden MAP x genetic PC2"
## [17,] "garden_MAP:genetic_PC3" "Garden MAP x genetic PC3"
```

```
# adjusted pvalues
cell_text <- pval1
cell_text[pval1 > 0.05] <- ''
cell_text[pval1 <= 0.05 & pval1 > 0.01] <- '*'
cell_text[pval1 <= 0.01 & pval1 > 0.001] <- '**'
cell_text[pval1<= 0.001] <- '***'
cell_text <- t(cell_text)

# use same color scale for all heatmaps

# color scale based on all effect sizes
max_col <- max(c(unlist(df1), unlist(df2), unlist(df3)), na.rm = T)
min_col <- min(c(unlist(df1), unlist(df2), unlist(df3)), na.rm = T)

# make scale same at both ends
max_col <- max(abs(max_col), abs(min_col))
min_col <- min(abs(max_col), abs(min_col))*-1
max_ceil <- ceiling(max_col*10)/10 # take ceiling rounded to one decimal place - used for placing ticks on legend

colf <- colorRamp2(c('red', 'white', 'blue'), breaks = c(min_col, 0, max_col))

# column annotation - GxE categories

col_cat <-  c(rep('E', 2), rep('G', 5), rep('GxE', 10))
ha_col <- columnAnnotation(Effect_Category = col_cat,
                        show_annotation_name = F,
                        col = list(Effect_Category = c('E' = "#015b58", 'G' = "#4760b8", 'GxE' = "#380069" )),
                        annotation_legend_param = list(title = 'Effect Category', title_gp = gpar(fontsize = leg_title, fontface = "bold"), labels_gp = gpar(fontsize = leg_label)))


# row annotation - trait categories
cats <- c(rep('Phenology', 18),  rep('Leaf Morphology', 4), rep('Stomata', 7), rep('Photosynthesis', 5))
cbind(trait_names, cats)
```

```
##       trait_names                        cats             
##  [1,] "Growing Season Days 2023"         "Phenology"      
##  [2,] "Growing Season Days 2024"         "Phenology"      
##  [3,] "Stage2 Bud Flush DOY 2023"        "Phenology"      
##  [4,] "Stage2 Bud Flush cGDD 2023"       "Phenology"      
##  [5,] "Stage2 Bud Flush DOY 2024"        "Phenology"      
##  [6,] "Stage2 Bud Flush cGDD 2024"       "Phenology"      
##  [7,] "Stage3 Leaf Emergence DOY 2023"   "Phenology"      
##  [8,] "Stage3 cGDD 2023"                 "Phenology"      
##  [9,] "Stage3 Leaf Emergence DOY 2024"   "Phenology"      
## [10,] "Stage3 cGDD 2024"                 "Phenology"      
## [11,] "Stage6 First Budset DOY 2023"     "Phenology"      
## [12,] "Stage6 First Budset DOY 2024"     "Phenology"      
## [13,] "Stage7 Lammas Growth DOY 2023"    "Phenology"      
## [14,] "Stage7 Lammas Growth DOY 2024"    "Phenology"      
## [15,] "stage7 presence 2023"             "Phenology"      
## [16,] "stage7 presence 2024"             "Phenology"      
## [17,] "Stage8 Final Budset DOY 2023"     "Phenology"      
## [18,] "Stage8 Final Budset DOY 2024"     "Phenology"      
## [19,] "Leaf thickness avg 2023"          "Leaf Morphology"
## [20,] "Leaf area 2023"                   "Leaf Morphology"
## [21,] "Leaf mass 2023"                   "Leaf Morphology"
## [22,] "LMA 2023"                         "Leaf Morphology"
## [23,] "Abaxial stomata pore length mean" "Stomata"        
## [24,] "Adaxial stomata pore length mean" "Stomata"        
## [25,] "Abaxial stomata density"          "Stomata"        
## [26,] "Adaxial stomata density"          "Stomata"        
## [27,] "Adaxial stomata presence"         "Stomata"        
## [28,] "Stomata ratio"                    "Stomata"        
## [29,] "gsw"                              "Stomata"        
## [30,] "ETR"                              "Photosynthesis" 
## [31,] "Fm'"                              "Photosynthesis" 
## [32,] "Fs"                               "Photosynthesis" 
## [33,] "gbw"                              "Photosynthesis" 
## [34,] "PhiPS2"                           "Photosynthesis"
```

```
ha_row <- rowAnnotation(Trait_Category = cats,
                        show_annotation_name = F,
                            col = list(Trait_Category = c('Phenology' = "#e2e260", 'Photosynthesis' = "#2d4030", 'Leaf Morphology' = "#516823", 'Stomata' =  "#88a2b9")),
                        annotation_legend_param = list(title = 'Trait Category', title_gp = gpar(fontsize = leg_title, fontface = "bold"), labels_gp = gpar(fontsize = leg_label)))


# row annotation - trait categories

hm1 <- Heatmap(as.matrix(t(df1)),
        cluster_rows = F,
        cluster_columns = F,
        rect_gp = gpar(col = "black", lwd = 1),
        col = colf,
        na_col = 'grey80',
        row_names_side = 'left',
        row_labels = trait_names,
        column_names_side = 'top',
        column_names_rot = rot,
        column_names_centered = F,
        column_title = 'Trait Predictors',
        column_title_gp = gpar(fontface = 'bold', fontsize = leg_title),
        column_labels = effect_names,
        row_title_gp = gpar(fontface = 'bold', fontsize = leg_title),
        row_title = 'Trait',
        left_annotation = ha_row,
        top_annotation = ha_col,
        heatmap_legend_param = list(title = 'Std. Effect Size',  at = c(max_ceil*-1, 0, max_ceil), title_gp = gpar(fontsize = leg_title, fontface = "bold"), labels_gp = gpar(fontsize = leg_label)),
        cell_fun = function(j, i, x, y, width, height, fill) { # add text to each grid
                      grid.text(cell_text[i, j], x, y, gp = gpar(cex = text_cex))}
        )

ComplexHeatmap::draw(hm1, padding = unit(c(2, 30, 2, 15), "mm"))
```

```
########## plot together

effect_names1 <- rownames(df2)
effect_names1 <- gsub('genetic', 'Genetic', effect_names1)
effect_names1 <- gsub('provenance', 'Home', effect_names1)
effect_names1 <- gsub('_', ' ', effect_names1)


########


# standardized effect sizes for simple linear regressions

# cell text showing significance
cell_text <- pval2
cell_text[pval2 > 0.05] <- ''
cell_text[pval2 <= 0.05 & pval2 > 0.01] <- '*'
cell_text[pval2 <= 0.01 & pval2 > 0.001] <- '**'
cell_text[pval2 <= 0.001] <- '***'
cell_text <- t(cell_text)


hm2 <- Heatmap(as.matrix(t(df2)),
        cluster_rows = F,
        cluster_columns = F,
        rect_gp = gpar(col = "black", lwd = 1),
        col = colf,
        na_col = 'grey80',
        row_names_side = 'left',
        column_names_side = 'top',
        column_names_rot = rot,
        column_names_centered = F,
        column_title = 'Plasticity\nPredictors',
        column_labels = effect_names1,
        column_title_gp = gpar(fontface = 'bold', fontsize = leg_title),
        row_title_gp = gpar(fontface = 'bold', fontsize = leg_title),
        row_title = 'Trait plasticity',
        row_labels = trait_names,
        left_annotation = ha_row,
        row_names_max_width = unit(8, 'cm'),
        #heatmap_legend_param = list(title = 'Std. Effect Size', at = c(max_ceil*-1, 0, max_ceil)),
        show_heatmap_legend = F, # legend is the same, so don't plot it
        cell_fun = function(j, i, x, y, width, height, fill) { # add text to each grid
                      grid.text(cell_text[i, j], x, y, gp = gpar(cex = text_cex))})

ComplexHeatmap::draw(hm2, padding = unit(c(2, 30, 2, 15), "mm"))
```

```
ComplexHeatmap::draw(hm1 + hm2, padding = unit(c(2, 30, 2, 15), "mm"))
```

```
# linear regression, climate controlling for ancestry

effect_names2 <- rownames(df3)
effect_names2 <- gsub('provenance', 'Home', effect_names2)
effect_names2 <- gsub('_', ' ', effect_names2)

# cell text showing significance
cell_text <- pval3
cell_text[pval3 > 0.05] <- ''
cell_text[pval3 <= 0.05 & pval3 > 0.01] <- '*'
cell_text[pval3 <= 0.01 & pval3 > 0.001] <- '**'
cell_text[pval3<= 0.001] <- '***'
cell_text <- t(cell_text)


hm3 <- Heatmap(as.matrix(t(df3)),
        cluster_rows = F,
        cluster_columns = F,
        rect_gp = gpar(col = "black", lwd = 1),
        col = colf,
        na_col = 'grey80',
        row_names_side = 'left',
        column_names_side = 'top',
        column_names_rot = rot,
        column_names_centered = F,
        column_title = paste0('Plasticity', '\n', 'Predictors', '\n', '(controlling for ancestry)'),
        column_labels = effect_names2,
        column_title_gp = gpar(fontface = 'bold', fontsize = leg_title),
        row_title_gp = gpar(fontface = 'bold', fontsize = leg_title),
        row_title = 'Trait plasticity',
        row_labels = trait_names,
        left_annotation = ha_row,
        #heatmap_legend_param = list(title = 'Std. Effect Size', at = c(max_ceil*-1, 0, max_ceil)),
        show_heatmap_legend = F, # legend is the same, so don't plot it
        cell_fun = function(j, i, x, y, width, height, fill) { # add text to each grid
                      grid.text(cell_text[i, j], x, y, gp = gpar(cex = text_cex))})

ComplexHeatmap::draw(hm3, padding = unit(c(2, 30, 2, 15), "mm"))
```

```
# the combined heatmap!

#png(file = 'results/plasticity/heatmap_ALL_GxE_plasticity_vs_ancestry_and_climate_padjusted.png', height = 10, width = 16, res = 300, units = 'in')
#pdf(file = 'results/plasticity/heatmap_ALL_GxE_plasticity_vs_ancestry_and_climate_padjusted.pdf', height = 10, width = 16)

# alternatively - rotated column names
#png(file = 'results/plasticity/heatmap_ALL_GxE_plasticity_vs_ancestry_and_climate_padjusted_columnNamesRotated.png', height = 10, width = 18, res = 300, units = 'in')
#pdf(file = 'results/plasticity/heatmap_ALL_GxE_plasticity_vs_ancestry_and_climate_padjusted_columnNamesRotated.pdf', height = 10, width = 16)

draw(hm1 + hm2 + hm3, padding = unit(c(2, 15, 2, 15), "mm"))
```

```
#dev.off()
```
