## Supplementary material for "Phenotypic plasticity evolved for climate variability constrains performance under climate warming": Analysis Rmarkdown Files: README.md

# Scripts for PopUp Poplar Plasticity Analyses

Code for **"Phenotypic plasticity evolved for climate variability constrains performance under climate warming"**

Alayna Mead*, Michelle Zavala-Paez, Joie R. Beasley-Bennett, Andrew C. Bleich, Ian P. Clancy-Mallue, Dylan G. Fischer, Julie M. Golightly, Kristina M. Hufford, Lee A. Kalcsits, Sara K. Klopf, Jesse R. Lasky, Jared M. LeBoldus, David B. Lowry, Nora Mitchell, Emily V. Moran, Jason P. Sexton, Kelsey L. Søndreli, Matthew C. Fitzpatrick, Jason Holliday, Stephen Keller, Jill A. Hamilton



All scripts are R markdown files.

To recreate an R environment with package versions used here, the the renv.lock file is included in the 'code' directory. See details for using the renv package [here](https://rstudio.github.io/renv/articles/renv.html).


## calculate_plasticity_metrics.Rmd

Calculates plasticity metrics (including RDPI) from phenotypic data.

input: mini_garden_phenotypic_and_climate_data_2021-2024.Rdata

output: genotypePlasticityRDPI_2023-2024.Rdata


## variance_partitioning_GxE.Rmd

Determines the proportion of variance in a trait that is explained by genotype (G), environment (E), genotype x environment (GxE), block nested within garden, and the residuals.

input: mini_garden_phenotypic_and_climate_data_2021-2024.Rdata

output: Figure 3

## GxE_models_all_traits.Rmd

Tests for GxE effects on phenotypic traits. Unlike variance partitioning script, here we test the effects of specific climate and genetic variables.

input: mini_garden_phenotypic_and_climate_data_2021-2024.Rdata

output: Figure 4A
GxE_traits_heatmap_data.Rdata (used to merge heatmaps in next script)


## plasticity_vs_ancestry_and_climate_correlations.Rmd

Tests for relationships between trait plasticity and genotype characteristics, including home climate and genetic structure.
Plots a summary of model results as a heatmap.

input: genotypePlasticityRDPI_2023-2024.Rdata

output: Figure 4B and C (statistical models) + combined Figure 4, Figure 5, Figure S4

## test-for-trait-fitness_associations.Rmd

Test whether traits and their plasticity predict growth in each garden and summarize relationships across garden climates.

inputs: mini_garden_phenotypic_and_climate_data_2021-2024.Rdata, genotypePlasticityRDPI_2023-2024.Rdata

outputs: Figure 6, Figures S5-S11
