## Supplementary material for "Phenotypic plasticity evolved for climate variability constrains performance under climate warming": Analysis Rmarkdown Files: test_for_trait-fitness_associations.html


### Test for trait-fitness associations

###### Alayna Mead

#### 2026-03-12

- 1 Setup
  - 1.1 Functions
  - 1.2 Setup for climate color
    palettes
  - 1.3 Setup trait labels
- 2 Statistical models for traits
  - 2.1 Do trait values predict growth in
    2023?
    - 2.1.1 Garden MAT
    - 2.1.2 Garden MAP
    - 2.1.3 Garden TD
  - 2.2 Do trait values predict growth in
    2024?
    - 2.2.1 Garden MAT
    - 2.2.2 Garden MAP
    - 2.2.3 Garden TD
  - 2.3 Combine plots for both
    years
  - 2.4 Do trait values predict
    end-of-year survival?
- 3 Statistical models for
  plasticity
  - 3.1 Does trait plasticity (RDPI)
    predict growth in 2023?
    - 3.1.1 Garden MAT
    - 3.1.2 Garden MAP
    - 3.1.3 Garden TD
  - 3.2 Does trait plasticity (RDPI)
    predict growth in 2024?
    - 3.2.1 Garden MAT
    - 3.2.2 Garden MAP
    - 3.2.3 Garden TD
  - 3.3 Plasticity plots for both
    years
- 4 Is there a tradeoff between
  plasticity and traits?

Test whether traits and their plasticity predict growth in each
garden and summarize relationships across garden climates.

### 1 Setup

```
library(ggplot2)
library(gridExtra) # arrange plots
library(cowplot) # arrange plots
library(ggrepel) # text labels for gardens
library(ggnewscale) # multiple color scales
library(lme4) # models
```

```
## Loading required package: Matrix
```

```
library(sjPlot) # model plots
```

```
## 
## Attaching package: 'sjPlot'
```

```
## The following objects are masked from 'package:cowplot':
## 
##     plot_grid, save_plot
```

```
library(car) # Anova
```

```
## Loading required package: carData
```

```
sessionInfo()
```

```
## R version 4.5.2 (2025-10-31)
## Platform: x86_64-pc-linux-gnu
## Running under: Arch Linux
## 
## Matrix products: default
## BLAS:   /usr/lib/libblas.so.3.12.0 
## LAPACK: /usr/lib/liblapack.so.3.12.0  LAPACK version 3.12.0
## 
## locale:
##  [1] LC_CTYPE=en_US.UTF-8       LC_NUMERIC=C              
##  [3] LC_TIME=en_US.UTF-8        LC_COLLATE=en_US.UTF-8    
##  [5] LC_MONETARY=en_US.UTF-8    LC_MESSAGES=en_US.UTF-8   
##  [7] LC_PAPER=en_US.UTF-8       LC_NAME=C                 
##  [9] LC_ADDRESS=C               LC_TELEPHONE=C            
## [11] LC_MEASUREMENT=en_US.UTF-8 LC_IDENTIFICATION=C       
## 
## time zone: US/Eastern
## tzcode source: system (glibc)
## 
## attached base packages:
## [1] stats     graphics  grDevices datasets  utils     methods   base     
## 
## other attached packages:
##  [1] car_3.1-3        carData_3.0-5    sjPlot_2.8.17    lme4_1.1-37     
##  [5] Matrix_1.7-4     ggnewscale_0.5.1 ggrepel_0.9.6    cowplot_1.2.0   
##  [9] gridExtra_2.3    ggplot2_3.5.2   
## 
## loaded via a namespace (and not attached):
##  [1] tidyr_1.3.1        sass_0.4.10        generics_0.1.4     renv_0.17.3       
##  [5] lattice_0.22-7     digest_0.6.37      magrittr_2.0.3     evaluate_1.0.3    
##  [9] grid_4.5.2         fastmap_1.2.0      jsonlite_2.0.0     ggeffects_2.2.1   
## [13] Formula_1.2-5      purrr_1.0.4        scales_1.3.0       jquerylib_0.1.4   
## [17] abind_1.4-8        reformulas_0.4.0   Rdpack_2.6.4       cli_3.6.5         
## [21] sjmisc_2.8.10      rlang_1.1.6        rbibutils_2.3      performance_0.15.1
## [25] munsell_0.5.1      splines_4.5.2      withr_3.0.2        cachem_1.1.0      
## [29] yaml_2.3.10        datawizard_1.2.0   sjstats_0.19.0     tools_4.5.2       
## [33] nloptr_2.2.1       minqa_1.2.8        dplyr_1.1.4        colorspace_2.1-1  
## [37] sjlabelled_1.2.0   boot_1.3-32        vctrs_0.6.5        R6_2.6.1          
## [41] lifecycle_1.0.4    MASS_7.3-65        insight_1.4.2      pkgconfig_2.0.3   
## [45] pillar_1.10.2      bslib_0.9.0        gtable_0.3.6       glue_1.8.0        
## [49] Rcpp_1.0.14        xfun_0.52          tibble_3.2.1       tidyselect_1.2.1  
## [53] rstudioapi_0.17.1  knitr_1.50         htmltools_0.5.8.1  nlme_3.1-168      
## [57] rmarkdown_2.29     compiler_4.5.2
```

```
# load data
load('data/clean/mini_garden_phenotypic_and_climate_data_2021-2024.Rdata')


# set negative heights to NA
dat$GrowthIncrement_2024[dat$GrowthIncrement_2024 < 0] <- NA
dat$GrowthIncrement_2023[dat$GrowthIncrement_2023 < 0] <- NA
dat$GrowthIncrement_2022[dat$GrowthIncrement_2022 < 0] <- NA
dat$GrowthIncrement_2021[dat$GrowthIncrement_2021 < 0] <- NA

# remove NA genotypes and those without genetic info
dat <- dat[! is.na(dat$Genotype),]
dat <- dat[! is.na(dat$k2_tricho),]

# estimate mortality at beginning of spring
# if it was marked alive in fall but did not have a height next spring, mark as dead
dat$Survival_spring_2024 <- NA
dat$Survival_spring_2024[dat$Survival_09_2023 == 1 & is.na(dat$PreFlush_Height_cm_2024)] <- 0
dat$Survival_spring_2024[dat$Survival_09_2023 == 1 & ! is.na(dat$PreFlush_Height_cm_2024)] <- 1

table(dat$Survival_spring_2024, dat$MiniCG_Site)
```

```
##    
##     OLLU LOCK VA UCM SU NWMO PENN WYO MORTON MSU OSU SWMN WI ID NDSU EVERGREEN
##   0    0    0 67   0  0    0    0   0      3   0  48    5  3  1    0         0
##   1    0    0  0   0 83    0   72  64     47  76   0   76 45 70    0        85
##    
##     WSU
##   0  99
##   1   0
```

```
# load plasticity values - named rdpi
load('data/clean/genotypePlasticityRDPI_2023-2024.Rdata')

# print session info, including package versions
sessionInfo()
```

```
## R version 4.5.2 (2025-10-31)
## Platform: x86_64-pc-linux-gnu
## Running under: Arch Linux
## 
## Matrix products: default
## BLAS:   /usr/lib/libblas.so.3.12.0 
## LAPACK: /usr/lib/liblapack.so.3.12.0  LAPACK version 3.12.0
## 
## locale:
##  [1] LC_CTYPE=en_US.UTF-8       LC_NUMERIC=C              
##  [3] LC_TIME=en_US.UTF-8        LC_COLLATE=en_US.UTF-8    
##  [5] LC_MONETARY=en_US.UTF-8    LC_MESSAGES=en_US.UTF-8   
##  [7] LC_PAPER=en_US.UTF-8       LC_NAME=C                 
##  [9] LC_ADDRESS=C               LC_TELEPHONE=C            
## [11] LC_MEASUREMENT=en_US.UTF-8 LC_IDENTIFICATION=C       
## 
## time zone: US/Eastern
## tzcode source: system (glibc)
## 
## attached base packages:
## [1] stats     graphics  grDevices datasets  utils     methods   base     
## 
## other attached packages:
##  [1] car_3.1-3        carData_3.0-5    sjPlot_2.8.17    lme4_1.1-37     
##  [5] Matrix_1.7-4     ggnewscale_0.5.1 ggrepel_0.9.6    cowplot_1.2.0   
##  [9] gridExtra_2.3    ggplot2_3.5.2   
## 
## loaded via a namespace (and not attached):
##  [1] tidyr_1.3.1        sass_0.4.10        generics_0.1.4     renv_0.17.3       
##  [5] lattice_0.22-7     digest_0.6.37      magrittr_2.0.3     evaluate_1.0.3    
##  [9] grid_4.5.2         fastmap_1.2.0      jsonlite_2.0.0     ggeffects_2.2.1   
## [13] Formula_1.2-5      purrr_1.0.4        scales_1.3.0       jquerylib_0.1.4   
## [17] abind_1.4-8        reformulas_0.4.0   Rdpack_2.6.4       cli_3.6.5         
## [21] sjmisc_2.8.10      rlang_1.1.6        rbibutils_2.3      performance_0.15.1
## [25] munsell_0.5.1      splines_4.5.2      withr_3.0.2        cachem_1.1.0      
## [29] yaml_2.3.10        datawizard_1.2.0   sjstats_0.19.0     tools_4.5.2       
## [33] nloptr_2.2.1       minqa_1.2.8        dplyr_1.1.4        colorspace_2.1-1  
## [37] sjlabelled_1.2.0   boot_1.3-32        vctrs_0.6.5        R6_2.6.1          
## [41] lifecycle_1.0.4    MASS_7.3-65        insight_1.4.2      pkgconfig_2.0.3   
## [45] pillar_1.10.2      bslib_0.9.0        gtable_0.3.6       glue_1.8.0        
## [49] Rcpp_1.0.14        xfun_0.52          tibble_3.2.1       tidyselect_1.2.1  
## [53] rstudioapi_0.17.1  knitr_1.50         htmltools_0.5.8.1  nlme_3.1-168      
## [57] rmarkdown_2.29     compiler_4.5.2
```

```
# setup markdown
knitr::opts_chunk$set(fig.width = 14, fig.height = 12)
```

```
# setup ggplot
theme_set(
  theme_bw(base_size = 12) +
    theme(plot.title = element_text(size = 12))
)
```

```
# look at relationships among garden climate variables

# garden climates
par(mfrow = c(1, 3))
plot(dat$garden_climate_2023_PC1, dat$garden_climate_2023_PC2, type = 'n')
text(dat$garden_climate_2023_PC1, dat$garden_climate_2023_PC2, labels = dat$MiniCG_Site)

plot(dat$garden_MAT_2023, dat$garden_MAP_2023, type = 'n')
text(dat$garden_MAT_2023, dat$garden_MAP_2023, labels = dat$MiniCG_Site)

plot(dat$garden_MAT_2023, dat$garden_TD_2023, type = 'n')
text(dat$garden_MAT_2023, dat$garden_TD_2023, labels = dat$MiniCG_Site)
```

```
par(mfrow = c(1, 1)) # reset


# plot traits vs fitness for each garden separately

traits <- c("DOY_last_budset_2023", "growing_season_days_2023", "licor_gsw", "licor_gbw", "licor_ETR", "licor_PhiPS2", "licor_Fs", "licor_Fm.", "leaf_thickness_avg_mm_2023", "LMA_g_m2_2023", "leaf_mass_g_2023", "leaf_area_cm2_2023", "lower_stomata_pore_length_mean_um", "upper_stomata_presence", "upper_stomata_pore_length_mean_um",  "upper_stomata_density_mm2", "lower_stomata_density_mm2", "stomata_density_both", "upper_stomata_density_over_total_density", "stomata_ratio")

dat$log_GrowthIncrement_2023 <- log(dat$GrowthIncrement_2023+1)
```

#### 1.1 Functions

```
# function to convert numeric pvalues to text (used for plot titles)
pval_stars <- function(pval){
  
  if(pval > 0.05){
    return('NS')
  } else if(pval <= 0.05 & pval > 0.01){
    return('*')
  } else if(pval <= 0.01 & pval > 0.001){
    return('**')
  } else if(pval <= 0.001){
    return('***')
  }
  
}
```

#### 1.2 Setup for climate color palettes

```
# get climate ranges across both years, so the same color scale is used for 2023 and 2024 plots
# this is used to set 'limits' in scale_color_gradient2() 


# MAT
sites.23 <- c("SWMN", "WI", "ID", "WYO", "EVERGREEN", "MSU", "OSU", "MORTON", "WSU", "PENN", "VA", "SU")
sites.24 <- c("SWMN", "WI", "ID", "WYO", "EVERGREEN", "MSU", "MORTON", "PENN", "SU")

sites.mat.23 <- dat$garden_MAT_2023[match(sites.23, dat$MiniCG_Site)]
sites.mat.24 <- dat$garden_MAT_2024[match(sites.24, dat$MiniCG_Site)]

range_mat <- range(c(sites.mat.23, sites.mat.24))

# MAP
sites.map.23 <- dat$garden_MAP_2023[match(sites.23, dat$MiniCG_Site)]
sites.map.24 <- dat$garden_MAP_2024[match(sites.24, dat$MiniCG_Site)]

range_map <- range(c(sites.map.23, sites.map.24))

# TD
sites.td.23 <- dat$garden_TD_2023[match(sites.23, dat$MiniCG_Site)]
sites.td.24 <- dat$garden_TD_2024[match(sites.24, dat$MiniCG_Site)]

range_td <- range(c(sites.td.23, sites.td.24))
```

#### 1.3 Setup trait labels

```
traits <- c("DOY_Stage2_2023", "DOY_Stage3_2023", "DOY_Stage6_2023", "DOY_Stage7_2023", "stage7_presence_2023", "DOY_last_budset_2023", "growing_season_days_2023", "licor_gsw", "licor_gbw", "licor_ETR", "licor_PhiPS2", "licor_Fs", "licor_Fm.", "leaf_thickness_avg_mm_2023", "LMA_g_m2_2023", "leaf_mass_g_2023", "leaf_area_cm2_2023", "lower_stomata_pore_length_mean_um", "upper_stomata_presence", "upper_stomata_pore_length_mean_um",  "upper_stomata_density_mm2", "lower_stomata_density_mm2", "stomata_ratio", "DOY_Stage2_2024", "DOY_Stage3_2024", "DOY_Stage6_2024", "DOY_Stage7_2024", "stage7_presence_2024", "DOY_last_budset_2024", "growing_season_days_2024", "plasticity_rdpi_GrowthIncrement_2023_log", "plasticity_rdpi_DOY_Stage2_2023", "plasticity_rdpi_DOY_Stage3_2023", "plasticity_rdpi_DOY_Stage6_2023", "plasticity_rdpi_DOY_last_budset_2023", "plasticity_rdpi_stage7_presence_2023", "plasticity_rdpi_growing_season_days_2023", "plasticity_rdpi_DOY_Stage7_2023", "plasticity_rdpi_licor_gsw", "plasticity_rdpi_licor_gbw", "plasticity_rdpi_licor_PhiPS2", "plasticity_rdpi_licor_ETR", "plasticity_rdpi_licor_Fs", "plasticity_rdpi_licor_Fm.", "plasticity_rdpi_leaf_thickness_avg_mm_2023", "plasticity_rdpi_LMA_g_m2_2023", "plasticity_rdpi_leaf_mass_g_2023_log", "plasticity_rdpi_leaf_area_cm2_2023_log", "plasticity_rdpi_lower_stomata_count", "plasticity_rdpi_lower_stomata_pore_length_mean_um", "plasticity_rdpi_upper_stomata_presence", "plasticity_rdpi_upper_stomata_count_log", "plasticity_rdpi_upper_stomata_pore_length_mean_um", "plasticity_rdpi_upper_stomata_density_mm2_log", "plasticity_rdpi_lower_stomata_density_mm2", "plasticity_rdpi_stomata_density_both", "plasticity_rdpi_upper_stomata_density_over_total_density_log", "plasticity_rdpi_stomata_ratio_log", "plasticity_rdpi_DOY_Stage2_2024", "plasticity_rdpi_DOY_Stage3_2024", 
"plasticity_rdpi_DOY_Stage6_2024", "plasticity_rdpi_DOY_Stage7_2024", 
"plasticity_rdpi_DOY_last_budset_2024", "plasticity_rdpi_stage7_presence_2024", 
"plasticity_rdpi_growing_season_days_2024")

trait_names <- traits
################

trait_names <- gsub('Stage2_cGDD', 'Stage2_Bud_Flush_cGDD', trait_names)
trait_names <- gsub('DOY_Stage2', 'Stage2_Bud_Flush_DOY', trait_names)
trait_names <- gsub('DOY_Stage3', 'Stage3_Leaf_Emergence_DOY', trait_names)
trait_names <- gsub('DOY_Stage6', 'Stage6_First_Budset_DOY', trait_names)
trait_names <- gsub('DOY_Stage7', 'Stage7_Lammas_Growth_DOY', trait_names)
trait_names <- gsub('stage7_presence', 'Stage7_Lammas_Growth_presence', trait_names)
trait_names <- gsub('DOY_Stage8', 'Stage8_Budset_after_Lammas_DOY', trait_names)
trait_names <- gsub('DOY_last_budset', 'Stage8_Final_Budset_DOY', trait_names)
trait_names <- gsub('growing_season_days', 'Growing_Season_Days', trait_names)

trait_names <- gsub('licor_', '', trait_names)
trait_names <- gsub('Fm.', "Fm'", trait_names)
trait_names <- gsub('upper', "Adaxial", trait_names)
trait_names <- gsub('lower', "Abaxial", trait_names)
trait_names <- gsub('stomata_ratio', "Stomata_ratio", trait_names)
trait_names <- gsub('leaf', 'Leaf', trait_names)
trait_names <- gsub('Phi', 'Φ', trait_names)

# remove units
trait_names <- gsub('_cm2', '', trait_names)
trait_names <- gsub('_cm', '', trait_names)
trait_names <- gsub('_mm2', '', trait_names)
trait_names <- gsub('_m2', '', trait_names)
trait_names <- gsub('_mm', '', trait_names)
trait_names <- gsub('_um', '', trait_names)
trait_names <- gsub('_g_', ' ', trait_names)

trait_names <- gsub('plasticity_rdpi_', 'RDPI_', trait_names)

trait_names <- gsub('_', ' ', trait_names)

# check
cbind(traits, trait_names)
```

```
##       traits                                                        
##  [1,] "DOY_Stage2_2023"                                             
##  [2,] "DOY_Stage3_2023"                                             
##  [3,] "DOY_Stage6_2023"                                             
##  [4,] "DOY_Stage7_2023"                                             
##  [5,] "stage7_presence_2023"                                        
##  [6,] "DOY_last_budset_2023"                                        
##  [7,] "growing_season_days_2023"                                    
##  [8,] "licor_gsw"                                                   
##  [9,] "licor_gbw"                                                   
## [10,] "licor_ETR"                                                   
## [11,] "licor_PhiPS2"                                                
## [12,] "licor_Fs"                                                    
## [13,] "licor_Fm."                                                   
## [14,] "leaf_thickness_avg_mm_2023"                                  
## [15,] "LMA_g_m2_2023"                                               
## [16,] "leaf_mass_g_2023"                                            
## [17,] "leaf_area_cm2_2023"                                          
## [18,] "lower_stomata_pore_length_mean_um"                           
## [19,] "upper_stomata_presence"                                      
## [20,] "upper_stomata_pore_length_mean_um"                           
## [21,] "upper_stomata_density_mm2"                                   
## [22,] "lower_stomata_density_mm2"                                   
## [23,] "stomata_ratio"                                               
## [24,] "DOY_Stage2_2024"                                             
## [25,] "DOY_Stage3_2024"                                             
## [26,] "DOY_Stage6_2024"                                             
## [27,] "DOY_Stage7_2024"                                             
## [28,] "stage7_presence_2024"                                        
## [29,] "DOY_last_budset_2024"                                        
## [30,] "growing_season_days_2024"                                    
## [31,] "plasticity_rdpi_GrowthIncrement_2023_log"                    
## [32,] "plasticity_rdpi_DOY_Stage2_2023"                             
## [33,] "plasticity_rdpi_DOY_Stage3_2023"                             
## [34,] "plasticity_rdpi_DOY_Stage6_2023"                             
## [35,] "plasticity_rdpi_DOY_last_budset_2023"                        
## [36,] "plasticity_rdpi_stage7_presence_2023"                        
## [37,] "plasticity_rdpi_growing_season_days_2023"                    
## [38,] "plasticity_rdpi_DOY_Stage7_2023"                             
## [39,] "plasticity_rdpi_licor_gsw"                                   
## [40,] "plasticity_rdpi_licor_gbw"                                   
## [41,] "plasticity_rdpi_licor_PhiPS2"                                
## [42,] "plasticity_rdpi_licor_ETR"                                   
## [43,] "plasticity_rdpi_licor_Fs"                                    
## [44,] "plasticity_rdpi_licor_Fm."                                   
## [45,] "plasticity_rdpi_leaf_thickness_avg_mm_2023"                  
## [46,] "plasticity_rdpi_LMA_g_m2_2023"                               
## [47,] "plasticity_rdpi_leaf_mass_g_2023_log"                        
## [48,] "plasticity_rdpi_leaf_area_cm2_2023_log"                      
## [49,] "plasticity_rdpi_lower_stomata_count"                         
## [50,] "plasticity_rdpi_lower_stomata_pore_length_mean_um"           
## [51,] "plasticity_rdpi_upper_stomata_presence"                      
## [52,] "plasticity_rdpi_upper_stomata_count_log"                     
## [53,] "plasticity_rdpi_upper_stomata_pore_length_mean_um"           
## [54,] "plasticity_rdpi_upper_stomata_density_mm2_log"               
## [55,] "plasticity_rdpi_lower_stomata_density_mm2"                   
## [56,] "plasticity_rdpi_stomata_density_both"                        
## [57,] "plasticity_rdpi_upper_stomata_density_over_total_density_log"
## [58,] "plasticity_rdpi_stomata_ratio_log"                           
## [59,] "plasticity_rdpi_DOY_Stage2_2024"                             
## [60,] "plasticity_rdpi_DOY_Stage3_2024"                             
## [61,] "plasticity_rdpi_DOY_Stage6_2024"                             
## [62,] "plasticity_rdpi_DOY_Stage7_2024"                             
## [63,] "plasticity_rdpi_DOY_last_budset_2024"                        
## [64,] "plasticity_rdpi_stage7_presence_2024"                        
## [65,] "plasticity_rdpi_growing_season_days_2024"                    
##       trait_names                                          
##  [1,] "Stage2 Bud Flush DOY 2023"                          
##  [2,] "Stage3 Leaf Emergence DOY 2023"                     
##  [3,] "Stage6 First Budset DOY 2023"                       
##  [4,] "Stage7 Lammas Growth DOY 2023"                      
##  [5,] "Stage7 Lammas Growth presence 2023"                 
##  [6,] "Stage8 Final Budset DOY 2023"                       
##  [7,] "Growing Season Days 2023"                           
##  [8,] "gsw"                                                
##  [9,] "gbw"                                                
## [10,] "ETR"                                                
## [11,] "ΦPS2"                                               
## [12,] "Fs"                                                 
## [13,] "Fm'"                                                
## [14,] "Leaf thickness avg 2023"                            
## [15,] "LMA 2023"                                           
## [16,] "Leaf mass 2023"                                     
## [17,] "Leaf area 2023"                                     
## [18,] "Abaxial stomata pore length mean"                   
## [19,] "Adaxial stomata presence"                           
## [20,] "Adaxial stomata pore length mean"                   
## [21,] "Adaxial stomata density"                            
## [22,] "Abaxial stomata density"                            
## [23,] "Stomata ratio"                                      
## [24,] "Stage2 Bud Flush DOY 2024"                          
## [25,] "Stage3 Leaf Emergence DOY 2024"                     
## [26,] "Stage6 First Budset DOY 2024"                       
## [27,] "Stage7 Lammas Growth DOY 2024"                      
## [28,] "Stage7 Lammas Growth presence 2024"                 
## [29,] "Stage8 Final Budset DOY 2024"                       
## [30,] "Growing Season Days 2024"                           
## [31,] "RDPI GrowthIncrement 2023 log"                      
## [32,] "RDPI Stage2 Bud Flush DOY 2023"                     
## [33,] "RDPI Stage3 Leaf Emergence DOY 2023"                
## [34,] "RDPI Stage6 First Budset DOY 2023"                  
## [35,] "RDPI Stage8 Final Budset DOY 2023"                  
## [36,] "RDPI Stage7 Lammas Growth presence 2023"            
## [37,] "RDPI Growing Season Days 2023"                      
## [38,] "RDPI Stage7 Lammas Growth DOY 2023"                 
## [39,] "RDPI gsw"                                           
## [40,] "RDPI gbw"                                           
## [41,] "RDPI ΦPS2"                                          
## [42,] "RDPI ETR"                                           
## [43,] "RDPI Fs"                                            
## [44,] "RDPI Fm'"                                           
## [45,] "RDPI Leaf thickness avg 2023"                       
## [46,] "RDPI LMA 2023"                                      
## [47,] "RDPI Leaf mass 2023 log"                            
## [48,] "RDPI Leaf area 2023 log"                            
## [49,] "RDPI Abaxial stomata count"                         
## [50,] "RDPI Abaxial stomata pore length mean"              
## [51,] "RDPI Adaxial stomata presence"                      
## [52,] "RDPI Adaxial stomata count log"                     
## [53,] "RDPI Adaxial stomata pore length mean"              
## [54,] "RDPI Adaxial stomata density log"                   
## [55,] "RDPI Abaxial stomata density"                       
## [56,] "RDPI stomata density both"                          
## [57,] "RDPI Adaxial stomata density over total density log"
## [58,] "RDPI Stomata ratio log"                             
## [59,] "RDPI Stage2 Bud Flush DOY 2024"                     
## [60,] "RDPI Stage3 Leaf Emergence DOY 2024"                
## [61,] "RDPI Stage6 First Budset DOY 2024"                  
## [62,] "RDPI Stage7 Lammas Growth DOY 2024"                 
## [63,] "RDPI Stage8 Final Budset DOY 2024"                  
## [64,] "RDPI Stage7 Lammas Growth presence 2024"            
## [65,] "RDPI Growing Season Days 2024"
```

```
# make df to join them
trait_names_df <- cbind.data.frame(traits, trait_names)
rownames(trait_names_df) <- trait_names_df[,1]
```

### 2 Statistical models for traits

Test whether traits predict growth in each garden, and whether the
trait-growth relationship varies by garden climate.

#### 2.1 Do trait values predict growth in 2023?

```
# get 2023 traits

traits <- c("DOY_Stage2_2023", "DOY_Stage3_2023", "DOY_Stage6_2023", "DOY_Stage7_2023", "stage7_presence_2023", "DOY_last_budset_2023", "growing_season_days_2023", "licor_gsw", "licor_gbw", "licor_ETR", "licor_PhiPS2", "licor_Fs", "licor_Fm.", "leaf_thickness_avg_mm_2023", "LMA_g_m2_2023", "leaf_mass_g_2023", "leaf_area_cm2_2023", "lower_stomata_pore_length_mean_um", "upper_stomata_presence", "upper_stomata_pore_length_mean_um",  "upper_stomata_density_mm2", "lower_stomata_density_mm2", "stomata_ratio")


# do these traits predict growth increment?
```

##### 2.1.1 Garden MAT

Test whether garden climate, traits, and their interaction predict
growth. An interaction effect suggests that the relationship between the
trait and growth varies across garden climates.

This analysis is pretty much repeated for traits, plasticity, each
year, and three climate variables (MAT, MAP, TD). If modifying this code
in the future - it would probably have been cleaner to create functions
instead of repeating loops for each combination.

```
# reorder gardens by garden MAT
tmp <- aggregate(dat$garden_MAT_2023, by = list(dat$MiniCG_Site_2023), mean)
ord <- tmp$Group.1[order(tmp$x, decreasing = T)]
dat$MiniCG_Site_2023 <- factor(dat$MiniCG_Site_2023, levels = ord)

# sites <- levels(dat$MiniCG_Site_2023)
sites <- c("SWMN", "WI", "ID", "WYO", "EVERGREEN", "MSU", "OSU", "MORTON", "WSU", "PENN", "VA", "SU")

sites.mat <- dat$garden_MAT_2023[match(sites, dat$MiniCG_Site)]


# list to save results
mods.mat <- list()
# list to save summary plot for each trait with significant effects on fitness
plots.summ <- list()

# loop through traits
for(n in 1:length(traits)){
  
  trait <- traits[n]
  
  print(paste('Model for:', trait))
  
  mod <- lmer(paste0('GrowthIncrement_2023 ~', trait,  '* garden_MAT_2023 + Pt + (1 | MiniCG_Site/block) + (1 | Genotype)'),
              data = dat)
  
  print(car::Anova(mod))
  
  p1 <- plot_model(mod, vline.color = "black", show.values = T, type = 'std', 
                   title = paste('2023 Growth ~ ', trait, sep = ''))
  plot(p1)

  mods.mat[[n]] <- mod
  names(mods.mat)[n] <- trait
  
  #####################
  # posthoc tests
  # if either the climate or trait x climate are significant, run posthoc tests for each garden
  mod.info <- Anova(mod)
  # if any coefficients are significant, plot by garden
  # need to change indexes if model is changed!
  # 1 is trait effect, 2 is garden climate effect, 4 is trait x climate effect.
  # Skip 3 which is ancestry - we already know this predicts height.
  
  if(mod.info$`Pr(>Chisq)`[1] <= 0.05 | mod.info$`Pr(>Chisq)`[2] <= 0.05 | mod.info$`Pr(>Chisq)`[4] <=0.05){
    
    # list to save plots for each site
    plots.sub <- list()
    
    # dataframe to save outputs from posthoc tests
    df.posthoc <- as.data.frame(matrix(ncol = 3, nrow = length(sites)))
    rownames(df.posthoc) <- sites
    colnames(df.posthoc) <- c('mat', 'slope', 'pval')
    df.posthoc$mat <- sites.mat
    
    for(s in 1:length(sites)){
      
      site <- sites[s]
      dat.sub <- subset(dat, MiniCG_Site_2023 == site)
      # skip if site is missing data
      if(sum(!is.na(dat.sub[,trait])) > 10){
        mod.sub <- lmer(paste0('GrowthIncrement_2023 ~', trait, '+ Pt + (1 | block) + (1 | Genotype)'),
                        data = dat.sub)
        # get pval for trait
        mod.sub.info <- Anova(mod.sub)
        df.posthoc[site, 'pval'] <- mod.sub.info$`Pr(>Chisq)`[1]
        
        # get slopes from fixed effect of trait (effect on growth)
        df.posthoc[site, 'slope'] <- fixef(mod.sub)[2]
        
        # make title pretty
        maintitle <- paste(site, '\n', trait_names_df[trait, 'trait_names'], '\n', 'p = ', round(df.posthoc$pval[s], 3), sep = '')
        maintitle <- paste(strwrap(maintitle, width = 30), collapse = '\n')   
        
        # plot data for individual garden
        p <- ggplot(data = dat.sub, aes_string(y = 'GrowthIncrement_2023', x = trait, color = 'k2_tricho'), na.rm = TRUE) +
          geom_point(na.rm = TRUE) +
          scale_color_gradient2(high = "darkolivegreen2", mid = "grey20", low = "dodgerblue2", midpoint = 0.5, name = 'P. trichocarpa\nancestry', guide = 'none') +
          ggtitle(maintitle) +
          geom_smooth(color = 'black', lty = ifelse(df.posthoc$pval[s] <= 0.05, 1,2), method = 'lm', formula = y ~ x, na.rm = TRUE) +
          theme(axis.text.x = element_text(angle = 90, vjust = 0.5, hjust=1, size = 12)) +
          theme(axis.title.x = element_blank())
        
        #plot(p)
        plots.sub[[s]] <- p
      }
    }

    # pvals are adjusted for summary plot
    df.posthoc$pvals.adj <- p.adjust(df.posthoc$pval, method = 'BH') 
    
    # plot panel with relationship for each garden
    # remove empty plots
    plots.sub <- plots.sub[lengths(plots.sub)>0]
    # plot in grid
    p1 <- grid.arrange(grobs = plots.sub, ncol = 4)
    plot(p1)
    
    # setup text showing significance
    sig_text <- paste0('Trait ', pval_stars(mod.info[1, 'Pr(>Chisq)']), '\n',
           'Climate ', pval_stars(mod.info[2, 'Pr(>Chisq)']), '\n',
           'Trait x Climate ', pval_stars(mod.info[4, 'Pr(>Chisq)']))
    
    # wrap title
    maintitle <- trait_names_df[trait, 'trait_names']
    maintitle <- paste(strwrap(maintitle, width = 30), collapse = '\n')
    
    
    # plot slope of fitness relationship vs climate
    p2 <- ggplot(df.posthoc, aes(x = mat, y = slope), na.rm = TRUE) +
      geom_smooth(method = 'lm', col = 'black', na.rm = TRUE) +
      geom_hline(yintercept = 0, lty = 2) +
      geom_point(size = 5, stroke = 2, shape = ifelse(df.posthoc$pvals.adj <= 0.05, 16, 1), aes(col = mat), na.rm = TRUE) +
      scale_color_gradient2(high = "red2", mid = "grey", low = "blue2", midpoint = mean(range(sites.mat)), limits = range_mat, name = 'Garden MAT (°C)') +
      ggtitle(maintitle,
              subtitle = sig_text) +
      xlab('Garden MAT') +
      ylab('Effect on Growth Increment') +
      geom_text_repel(label = rownames(df.posthoc), aes(x = mat, y = slope), box.padding = 0.5, min.segment.length = 1, size = 4) +
        theme(plot.title = element_text(size = 15),
        plot.subtitle = element_text(size = 10),
        legend.position = 'none')
    plot(p2)
    
    plots.summ[[n]] <- p2
    names(plots.summ)[n] <- trait
    
    first <- FALSE # if we have made a plot, the next one isn't first

    
    # add legend
    leg_plot <- ggplot(df.posthoc, aes(x = mat, y = slope),  na.rm = TRUE) +
      geom_point(aes(color = mat))+
      scale_color_gradient2(high = "red2", mid = "grey", low = "blue2", midpoint = mean(range(sites.mat)), limits = range_mat, name = 'Garden MAT (°C)') +
      theme(legend.position = 'right')
    
    legend <- cowplot::get_legend(leg_plot)
    
  }
}
```

```
## [1] "Model for: DOY_Stage2_2023"
## Analysis of Deviance Table (Type II Wald chisquare tests)
## 
## Response: GrowthIncrement_2023
##                                   Chisq Df Pr(>Chisq)    
## DOY_Stage2_2023                  3.8925  1     0.0485 *  
## garden_MAT_2023                  1.5308  1     0.2160    
## Pt                              41.6229  1  1.107e-10 ***
## DOY_Stage2_2023:garden_MAT_2023  0.0754  1     0.7837    
## ---
## Signif. codes:  0 '***' 0.001 '**' 0.01 '*' 0.05 '.' 0.1 ' ' 1
```

```
## Warning: `aes_string()` was deprecated in ggplot2 3.0.0.
## ℹ Please use tidy evaluation idioms with `aes()`.
## ℹ See also `vignette("ggplot2-in-packages")` for more information.
## This warning is displayed once every 8 hours.
## Call `lifecycle::last_lifecycle_warnings()` to see where this warning was
## generated.
```

```
## boundary (singular) fit: see help('isSingular')
## boundary (singular) fit: see help('isSingular')
## boundary (singular) fit: see help('isSingular')
## boundary (singular) fit: see help('isSingular')
## boundary (singular) fit: see help('isSingular')
## boundary (singular) fit: see help('isSingular')
## boundary (singular) fit: see help('isSingular')
```

```
## `geom_smooth()` using formula = 'y ~ x'
```

```
## [1] "Model for: DOY_Stage3_2023"
## Analysis of Deviance Table (Type II Wald chisquare tests)
## 
## Response: GrowthIncrement_2023
##                                   Chisq Df Pr(>Chisq)    
## DOY_Stage3_2023                  1.7104  1     0.1909    
## garden_MAT_2023                  1.2916  1     0.2558    
## Pt                              32.8993  1  9.706e-09 ***
## DOY_Stage3_2023:garden_MAT_2023  0.0474  1     0.8276    
## ---
## Signif. codes:  0 '***' 0.001 '**' 0.01 '*' 0.05 '.' 0.1 ' ' 1
```

```
## [1] "Model for: DOY_Stage6_2023"
```

```
## Warning: Some predictor variables are on very different scales: consider
## rescaling
```

```
## Analysis of Deviance Table (Type II Wald chisquare tests)
## 
## Response: GrowthIncrement_2023
##                                   Chisq Df Pr(>Chisq)    
## DOY_Stage6_2023                 19.8936  1  8.187e-06 ***
## garden_MAT_2023                  1.7694  1     0.1835    
## Pt                              26.0263  1  3.368e-07 ***
## DOY_Stage6_2023:garden_MAT_2023 43.6591  1  3.909e-11 ***
## ---
## Signif. codes:  0 '***' 0.001 '**' 0.01 '*' 0.05 '.' 0.1 ' ' 1
```

```
## boundary (singular) fit: see help('isSingular')
## boundary (singular) fit: see help('isSingular')
## boundary (singular) fit: see help('isSingular')
## boundary (singular) fit: see help('isSingular')
## boundary (singular) fit: see help('isSingular')
## boundary (singular) fit: see help('isSingular')
```

```
## `geom_smooth()` using formula = 'y ~ x'
```

```
## Warning: Removed 1 row containing missing values or values outside the scale range
## (`geom_text_repel()`).
```

```
## Warning: Removed 1 row containing missing values or values outside the scale range
## (`geom_point()`).
```

```
## [1] "Model for: DOY_Stage7_2023"
```

```
## Warning: Some predictor variables are on very different scales: consider
## rescaling
```

```
## Analysis of Deviance Table (Type II Wald chisquare tests)
## 
## Response: GrowthIncrement_2023
##                                   Chisq Df Pr(>Chisq)    
## DOY_Stage7_2023                  1.4487  1    0.22873    
## garden_MAT_2023                  2.2428  1    0.13424    
## Pt                              36.4338  1  1.579e-09 ***
## DOY_Stage7_2023:garden_MAT_2023  2.7283  1    0.09858 .  
## ---
## Signif. codes:  0 '***' 0.001 '**' 0.01 '*' 0.05 '.' 0.1 ' ' 1
```

```
## [1] "Model for: stage7_presence_2023"
## Analysis of Deviance Table (Type II Wald chisquare tests)
## 
## Response: GrowthIncrement_2023
##                                        Chisq Df Pr(>Chisq)    
## stage7_presence_2023                  0.0309  1     0.8606    
## garden_MAT_2023                       1.9148  1     0.1664    
## Pt                                   46.1723  1  1.083e-11 ***
## stage7_presence_2023:garden_MAT_2023  0.0791  1     0.7786    
## ---
## Signif. codes:  0 '***' 0.001 '**' 0.01 '*' 0.05 '.' 0.1 ' ' 1
```

```
## [1] "Model for: DOY_last_budset_2023"
```

```
## Warning: Some predictor variables are on very different scales: consider
## rescaling
```

```
## Analysis of Deviance Table (Type II Wald chisquare tests)
## 
## Response: GrowthIncrement_2023
##                                        Chisq Df Pr(>Chisq)    
## DOY_last_budset_2023                  3.0145  1    0.08252 .  
## garden_MAT_2023                       1.6792  1    0.19503    
## Pt                                   45.2962  1  1.694e-11 ***
## DOY_last_budset_2023:garden_MAT_2023  0.1092  1    0.74110    
## ---
## Signif. codes:  0 '***' 0.001 '**' 0.01 '*' 0.05 '.' 0.1 ' ' 1
```

```
## [1] "Model for: growing_season_days_2023"
```

```
## Warning: Some predictor variables are on very different scales: consider
## rescaling
```

```
## Analysis of Deviance Table (Type II Wald chisquare tests)
## 
## Response: GrowthIncrement_2023
##                                            Chisq Df Pr(>Chisq)    
## growing_season_days_2023                  0.7836  1     0.3760    
## garden_MAT_2023                           1.9689  1     0.1606    
## Pt                                       47.3821  1  5.841e-12 ***
## growing_season_days_2023:garden_MAT_2023  1.0001  1     0.3173    
## ---
## Signif. codes:  0 '***' 0.001 '**' 0.01 '*' 0.05 '.' 0.1 ' ' 1
```

```
## [1] "Model for: licor_gsw"
## Analysis of Deviance Table (Type II Wald chisquare tests)
## 
## Response: GrowthIncrement_2023
##                             Chisq Df Pr(>Chisq)    
## licor_gsw                  1.7189  1    0.18983    
## garden_MAT_2023            5.3976  1    0.02016 *  
## Pt                        53.5563  1  2.513e-13 ***
## licor_gsw:garden_MAT_2023  6.2099  1    0.01270 *  
## ---
## Signif. codes:  0 '***' 0.001 '**' 0.01 '*' 0.05 '.' 0.1 ' ' 1
```

```
## boundary (singular) fit: see help('isSingular')
## boundary (singular) fit: see help('isSingular')
## boundary (singular) fit: see help('isSingular')
## boundary (singular) fit: see help('isSingular')
## boundary (singular) fit: see help('isSingular')
```

```
## `geom_smooth()` using formula = 'y ~ x'
```

```
## Warning: Removed 3 rows containing missing values or values outside the scale range
## (`geom_text_repel()`).
```

```
## Warning: Removed 3 rows containing missing values or values outside the scale range
## (`geom_point()`).
```

```
## [1] "Model for: licor_gbw"
```

```
## Warning: Some predictor variables are on very different scales: consider
## rescaling
```

```
## Analysis of Deviance Table (Type II Wald chisquare tests)
## 
## Response: GrowthIncrement_2023
##                             Chisq Df Pr(>Chisq)    
## licor_gbw                  0.4976  1   0.480538    
## garden_MAT_2023            7.3727  1   0.006622 ** 
## Pt                        54.8304  1  1.314e-13 ***
## licor_gbw:garden_MAT_2023  3.3325  1   0.067925 .  
## ---
## Signif. codes:  0 '***' 0.001 '**' 0.01 '*' 0.05 '.' 0.1 ' ' 1
```

```
## Warning: Some predictor variables are on very different scales: consider
## rescaling
```

```
## boundary (singular) fit: see help('isSingular')
```

```
## Warning: Some predictor variables are on very different scales: consider
## rescaling
```

```
## Warning: Some predictor variables are on very different scales: consider
## rescaling
```

```
## boundary (singular) fit: see help('isSingular')
```

```
## Warning: Some predictor variables are on very different scales: consider
## rescaling
```

```
## boundary (singular) fit: see help('isSingular')
```

```
## Warning: Some predictor variables are on very different scales: consider
## rescaling
## Warning: Some predictor variables are on very different scales: consider
## rescaling
## Warning: Some predictor variables are on very different scales: consider
## rescaling
```

```
## boundary (singular) fit: see help('isSingular')
```

```
## Warning: Some predictor variables are on very different scales: consider
## rescaling
```

```
## boundary (singular) fit: see help('isSingular')
```

```
## Warning: Some predictor variables are on very different scales: consider
## rescaling
```

```
## `geom_smooth()` using formula = 'y ~ x'
```

```
## Warning: Removed 3 rows containing missing values or values outside the scale range
## (`geom_text_repel()`).
```

```
## Warning: Removed 3 rows containing missing values or values outside the scale range
## (`geom_point()`).
```

```
## [1] "Model for: licor_ETR"
```

```
## Warning: Some predictor variables are on very different scales: consider
## rescaling
```

```
## Analysis of Deviance Table (Type II Wald chisquare tests)
## 
## Response: GrowthIncrement_2023
##                             Chisq Df Pr(>Chisq)    
## licor_ETR                  0.4531  1    0.50085    
## garden_MAT_2023            5.1133  1    0.02374 *  
## Pt                        57.7889  1  2.918e-14 ***
## licor_ETR:garden_MAT_2023  2.5508  1    0.11024    
## ---
## Signif. codes:  0 '***' 0.001 '**' 0.01 '*' 0.05 '.' 0.1 ' ' 1
```

```
## boundary (singular) fit: see help('isSingular')
## boundary (singular) fit: see help('isSingular')
## boundary (singular) fit: see help('isSingular')
## boundary (singular) fit: see help('isSingular')
## boundary (singular) fit: see help('isSingular')
```

```
## `geom_smooth()` using formula = 'y ~ x'
```

```
## Warning: Removed 3 rows containing missing values or values outside the scale range
## (`geom_text_repel()`).
## Removed 3 rows containing missing values or values outside the scale range
## (`geom_point()`).
```

```
## [1] "Model for: licor_PhiPS2"
## Analysis of Deviance Table (Type II Wald chisquare tests)
## 
## Response: GrowthIncrement_2023
##                                Chisq Df Pr(>Chisq)    
## licor_PhiPS2                  7.6867  1   0.005563 ** 
## garden_MAT_2023               4.6699  1   0.030696 *  
## Pt                           52.9076  1  3.496e-13 ***
## licor_PhiPS2:garden_MAT_2023  6.2200  1   0.012631 *  
## ---
## Signif. codes:  0 '***' 0.001 '**' 0.01 '*' 0.05 '.' 0.1 ' ' 1
```

```
## boundary (singular) fit: see help('isSingular')
## boundary (singular) fit: see help('isSingular')
## boundary (singular) fit: see help('isSingular')
## boundary (singular) fit: see help('isSingular')
## boundary (singular) fit: see help('isSingular')
## boundary (singular) fit: see help('isSingular')
```

```
## `geom_smooth()` using formula = 'y ~ x'
```

```
## Warning: Removed 3 rows containing missing values or values outside the scale range
## (`geom_text_repel()`).
## Removed 3 rows containing missing values or values outside the scale range
## (`geom_point()`).
```

```
## [1] "Model for: licor_Fs"
```

```
## Warning: Some predictor variables are on very different scales: consider
## rescaling
```

```
## Analysis of Deviance Table (Type II Wald chisquare tests)
## 
## Response: GrowthIncrement_2023
##                            Chisq Df Pr(>Chisq)    
## licor_Fs                  5.1500  1    0.02325 *  
## garden_MAT_2023           5.6229  1    0.01773 *  
## Pt                       49.2533  1   2.25e-12 ***
## licor_Fs:garden_MAT_2023  8.7299  1    0.00313 ** 
## ---
## Signif. codes:  0 '***' 0.001 '**' 0.01 '*' 0.05 '.' 0.1 ' ' 1
```

```
## boundary (singular) fit: see help('isSingular')
## boundary (singular) fit: see help('isSingular')
## boundary (singular) fit: see help('isSingular')
## boundary (singular) fit: see help('isSingular')
## boundary (singular) fit: see help('isSingular')
```

```
## `geom_smooth()` using formula = 'y ~ x'
```

```
## Warning: Removed 3 rows containing missing values or values outside the scale range
## (`geom_text_repel()`).
## Removed 3 rows containing missing values or values outside the scale range
## (`geom_point()`).
```

```
## [1] "Model for: licor_Fm."
```

```
## Warning: Some predictor variables are on very different scales: consider
## rescaling
```

```
## Analysis of Deviance Table (Type II Wald chisquare tests)
## 
## Response: GrowthIncrement_2023
##                             Chisq Df Pr(>Chisq)    
## licor_Fm.                  1.1663  1    0.28017    
## garden_MAT_2023            5.2396  1    0.02208 *  
## Pt                        55.4530  1  9.572e-14 ***
## licor_Fm.:garden_MAT_2023  0.2044  1    0.65119    
## ---
## Signif. codes:  0 '***' 0.001 '**' 0.01 '*' 0.05 '.' 0.1 ' ' 1
```

```
## boundary (singular) fit: see help('isSingular')
## boundary (singular) fit: see help('isSingular')
## boundary (singular) fit: see help('isSingular')
```

```
## Warning in checkConv(attr(opt, "derivs"), opt$par, ctrl = control$checkConv, :
## unable to evaluate scaled gradient
```

```
## Warning in checkConv(attr(opt, "derivs"), opt$par, ctrl = control$checkConv, :
## Model failed to converge: degenerate Hessian with 1 negative eigenvalues
```

```
## boundary (singular) fit: see help('isSingular')
## boundary (singular) fit: see help('isSingular')
```

```
## `geom_smooth()` using formula = 'y ~ x'
```

```
## Warning: Removed 3 rows containing missing values or values outside the scale range
## (`geom_text_repel()`).
```

```
## Warning: Removed 3 rows containing missing values or values outside the scale range
## (`geom_point()`).
```

```
## [1] "Model for: leaf_thickness_avg_mm_2023"
## Analysis of Deviance Table (Type II Wald chisquare tests)
## 
## Response: GrowthIncrement_2023
##                                              Chisq Df Pr(>Chisq)    
## leaf_thickness_avg_mm_2023                  4.5407  1    0.03310 *  
## garden_MAT_2023                             5.1296  1    0.02352 *  
## Pt                                         50.2927  1  1.324e-12 ***
## leaf_thickness_avg_mm_2023:garden_MAT_2023  1.1717  1    0.27905    
## ---
## Signif. codes:  0 '***' 0.001 '**' 0.01 '*' 0.05 '.' 0.1 ' ' 1
```

```
## boundary (singular) fit: see help('isSingular')
## boundary (singular) fit: see help('isSingular')
## boundary (singular) fit: see help('isSingular')
## boundary (singular) fit: see help('isSingular')
## boundary (singular) fit: see help('isSingular')
## boundary (singular) fit: see help('isSingular')
```

```
## `geom_smooth()` using formula = 'y ~ x'
```

```
## Warning: Removed 2 rows containing missing values or values outside the scale range
## (`geom_text_repel()`).
```

```
## Warning: Removed 2 rows containing missing values or values outside the scale range
## (`geom_point()`).
```

```
## [1] "Model for: LMA_g_m2_2023"
## Analysis of Deviance Table (Type II Wald chisquare tests)
## 
## Response: GrowthIncrement_2023
##                                 Chisq Df Pr(>Chisq)    
## LMA_g_m2_2023                  0.7845  1  0.3757743    
## garden_MAT_2023                2.2556  1  0.1331340    
## Pt                            36.9798  1  1.194e-09 ***
## LMA_g_m2_2023:garden_MAT_2023 10.8909  1  0.0009664 ***
## ---
## Signif. codes:  0 '***' 0.001 '**' 0.01 '*' 0.05 '.' 0.1 ' ' 1
```

```
## boundary (singular) fit: see help('isSingular')
## boundary (singular) fit: see help('isSingular')
## boundary (singular) fit: see help('isSingular')
## boundary (singular) fit: see help('isSingular')
## boundary (singular) fit: see help('isSingular')
## boundary (singular) fit: see help('isSingular')
## boundary (singular) fit: see help('isSingular')
## boundary (singular) fit: see help('isSingular')
```

```
## `geom_smooth()` using formula = 'y ~ x'
```

```
## Warning: Removed 1 row containing missing values or values outside the scale range
## (`geom_text_repel()`).
```

```
## Warning: Removed 1 row containing missing values or values outside the scale range
## (`geom_point()`).
```

```
## [1] "Model for: leaf_mass_g_2023"
## Analysis of Deviance Table (Type II Wald chisquare tests)
## 
## Response: GrowthIncrement_2023
##                                    Chisq Df Pr(>Chisq)    
## leaf_mass_g_2023                 30.8620  1  2.770e-08 ***
## garden_MAT_2023                   2.3942  1     0.1218    
## Pt                               28.6488  1  8.677e-08 ***
## leaf_mass_g_2023:garden_MAT_2023  0.4674  1     0.4942    
## ---
## Signif. codes:  0 '***' 0.001 '**' 0.01 '*' 0.05 '.' 0.1 ' ' 1
```

```
## boundary (singular) fit: see help('isSingular')
## boundary (singular) fit: see help('isSingular')
## boundary (singular) fit: see help('isSingular')
## boundary (singular) fit: see help('isSingular')
## boundary (singular) fit: see help('isSingular')
## boundary (singular) fit: see help('isSingular')
## boundary (singular) fit: see help('isSingular')
## boundary (singular) fit: see help('isSingular')
```

```
## `geom_smooth()` using formula = 'y ~ x'
```

```
## Warning: Removed 1 row containing missing values or values outside the scale range
## (`geom_text_repel()`).
## Removed 1 row containing missing values or values outside the scale range
## (`geom_point()`).
```

```
## [1] "Model for: leaf_area_cm2_2023"
## Analysis of Deviance Table (Type II Wald chisquare tests)
## 
## Response: GrowthIncrement_2023
##                                      Chisq Df Pr(>Chisq)    
## leaf_area_cm2_2023                 42.3726  1  7.544e-11 ***
## garden_MAT_2023                     2.4768  1     0.1155    
## Pt                                 28.4318  1  9.706e-08 ***
## leaf_area_cm2_2023:garden_MAT_2023  0.8130  1     0.3672    
## ---
## Signif. codes:  0 '***' 0.001 '**' 0.01 '*' 0.05 '.' 0.1 ' ' 1
```

```
## boundary (singular) fit: see help('isSingular')
## boundary (singular) fit: see help('isSingular')
## boundary (singular) fit: see help('isSingular')
## boundary (singular) fit: see help('isSingular')
## boundary (singular) fit: see help('isSingular')
## boundary (singular) fit: see help('isSingular')
## boundary (singular) fit: see help('isSingular')
## boundary (singular) fit: see help('isSingular')
## boundary (singular) fit: see help('isSingular')
```

```
## `geom_smooth()` using formula = 'y ~ x'
```

```
## Warning: Removed 1 row containing missing values or values outside the scale range
## (`geom_text_repel()`).
## Removed 1 row containing missing values or values outside the scale range
## (`geom_point()`).
```

```
## [1] "Model for: lower_stomata_pore_length_mean_um"
## Analysis of Deviance Table (Type II Wald chisquare tests)
## 
## Response: GrowthIncrement_2023
##                                                     Chisq Df Pr(>Chisq)    
## lower_stomata_pore_length_mean_um                  1.3532  1     0.2447    
## garden_MAT_2023                                    2.3328  1     0.1267    
## Pt                                                39.5515  1  3.195e-10 ***
## lower_stomata_pore_length_mean_um:garden_MAT_2023  2.3722  1     0.1235    
## ---
## Signif. codes:  0 '***' 0.001 '**' 0.01 '*' 0.05 '.' 0.1 ' ' 1
```

```
## [1] "Model for: upper_stomata_presence"
## Analysis of Deviance Table (Type II Wald chisquare tests)
## 
## Response: GrowthIncrement_2023
##                                          Chisq Df Pr(>Chisq)    
## upper_stomata_presence                  3.0230  1    0.08209 .  
## garden_MAT_2023                         2.2603  1    0.13273    
## Pt                                     40.5682  1  1.899e-10 ***
## upper_stomata_presence:garden_MAT_2023  0.2542  1    0.61412    
## ---
## Signif. codes:  0 '***' 0.001 '**' 0.01 '*' 0.05 '.' 0.1 ' ' 1
```

```
## [1] "Model for: upper_stomata_pore_length_mean_um"
## Analysis of Deviance Table (Type II Wald chisquare tests)
## 
## Response: GrowthIncrement_2023
##                                                     Chisq Df Pr(>Chisq)    
## upper_stomata_pore_length_mean_um                  4.0279  1    0.04475 *  
## garden_MAT_2023                                    2.2697  1    0.13193    
## Pt                                                20.1073  1  7.322e-06 ***
## upper_stomata_pore_length_mean_um:garden_MAT_2023  0.0322  1    0.85757    
## ---
## Signif. codes:  0 '***' 0.001 '**' 0.01 '*' 0.05 '.' 0.1 ' ' 1
```

```
## boundary (singular) fit: see help('isSingular')
## boundary (singular) fit: see help('isSingular')
## boundary (singular) fit: see help('isSingular')
## boundary (singular) fit: see help('isSingular')
## boundary (singular) fit: see help('isSingular')
## boundary (singular) fit: see help('isSingular')
## boundary (singular) fit: see help('isSingular')
## boundary (singular) fit: see help('isSingular')
```

```
## `geom_smooth()` using formula = 'y ~ x'
```

```
## Warning: Removed 1 row containing missing values or values outside the scale range
## (`geom_text_repel()`).
## Removed 1 row containing missing values or values outside the scale range
## (`geom_point()`).
```

```
## [1] "Model for: upper_stomata_density_mm2"
## Analysis of Deviance Table (Type II Wald chisquare tests)
## 
## Response: GrowthIncrement_2023
##                                             Chisq Df Pr(>Chisq)    
## upper_stomata_density_mm2                  1.3444  1     0.2463    
## garden_MAT_2023                            2.6237  1     0.1053    
## Pt                                        34.0893  1  5.264e-09 ***
## upper_stomata_density_mm2:garden_MAT_2023  0.0010  1     0.9749    
## ---
## Signif. codes:  0 '***' 0.001 '**' 0.01 '*' 0.05 '.' 0.1 ' ' 1
```

```
## [1] "Model for: lower_stomata_density_mm2"
```

```
## Warning: Some predictor variables are on very different scales: consider
## rescaling
```

```
## boundary (singular) fit: see help('isSingular')
```

```
## Analysis of Deviance Table (Type II Wald chisquare tests)
## 
## Response: GrowthIncrement_2023
##                                             Chisq Df Pr(>Chisq)    
## lower_stomata_density_mm2                  4.7020  1    0.03013 *  
## garden_MAT_2023                            2.4031  1    0.12109    
## Pt                                        36.8563  1  1.272e-09 ***
## lower_stomata_density_mm2:garden_MAT_2023  0.5163  1    0.47241    
## ---
## Signif. codes:  0 '***' 0.001 '**' 0.01 '*' 0.05 '.' 0.1 ' ' 1
```

```
## boundary (singular) fit: see help('isSingular')
```

```
## boundary (singular) fit: see help('isSingular')
## boundary (singular) fit: see help('isSingular')
## boundary (singular) fit: see help('isSingular')
## boundary (singular) fit: see help('isSingular')
## boundary (singular) fit: see help('isSingular')
## boundary (singular) fit: see help('isSingular')
## boundary (singular) fit: see help('isSingular')
```

```
## `geom_smooth()` using formula = 'y ~ x'
```

```
## Warning: Removed 1 row containing missing values or values outside the scale range
## (`geom_text_repel()`).
## Removed 1 row containing missing values or values outside the scale range
## (`geom_point()`).
```

```
## [1] "Model for: stomata_ratio"
```

```
## boundary (singular) fit: see help('isSingular')
```

```
## Analysis of Deviance Table (Type II Wald chisquare tests)
## 
## Response: GrowthIncrement_2023
##                                 Chisq Df Pr(>Chisq)    
## stomata_ratio                  2.9648  1    0.08510 .  
## garden_MAT_2023                3.0758  1    0.07946 .  
## Pt                            33.3459  1  7.714e-09 ***
## stomata_ratio:garden_MAT_2023  0.4019  1    0.52610    
## ---
## Signif. codes:  0 '***' 0.001 '**' 0.01 '*' 0.05 '.' 0.1 ' ' 1
```

```
## boundary (singular) fit: see help('isSingular')
```

```
plots.summ <- plots.summ[lengths(plots.summ)>0]

# add legend to beginning of plot list

plots2 <- plots.summ
plots2[[length(plots.summ)+1]] <- legend

p3 <- grid.arrange(grobs = plots2, ncol = 4)
```

```
## `geom_smooth()` using formula = 'y ~ x'
```

```
## `geom_smooth()` using formula = 'y ~ x'
```

```
## Warning: Removed 1 row containing missing values or values outside the scale range
## (`geom_text_repel()`).
```

```
## `geom_smooth()` using formula = 'y ~ x'
```

```
## Warning: Removed 3 rows containing missing values or values outside the scale range
## (`geom_text_repel()`).
```

```
## `geom_smooth()` using formula = 'y ~ x'
```

```
## Warning: Removed 3 rows containing missing values or values outside the scale range
## (`geom_text_repel()`).
```

```
## `geom_smooth()` using formula = 'y ~ x'
```

```
## Warning: Removed 3 rows containing missing values or values outside the scale range
## (`geom_text_repel()`).
```

```
## `geom_smooth()` using formula = 'y ~ x'
```

```
## Warning: Removed 3 rows containing missing values or values outside the scale range
## (`geom_text_repel()`).
```

```
## `geom_smooth()` using formula = 'y ~ x'
```

```
## Warning: Removed 3 rows containing missing values or values outside the scale range
## (`geom_text_repel()`).
```

```
## `geom_smooth()` using formula = 'y ~ x'
```

```
## Warning: Removed 3 rows containing missing values or values outside the scale range
## (`geom_text_repel()`).
```

```
## `geom_smooth()` using formula = 'y ~ x'
```

```
## Warning: Removed 2 rows containing missing values or values outside the scale range
## (`geom_text_repel()`).
```

```
## `geom_smooth()` using formula = 'y ~ x'
```

```
## Warning: Removed 1 row containing missing values or values outside the scale range
## (`geom_text_repel()`).
```

```
## `geom_smooth()` using formula = 'y ~ x'
```

```
## Warning: Removed 1 row containing missing values or values outside the scale range
## (`geom_text_repel()`).
```

```
## `geom_smooth()` using formula = 'y ~ x'
```

```
## Warning: Removed 1 row containing missing values or values outside the scale range
## (`geom_text_repel()`).
```

```
## `geom_smooth()` using formula = 'y ~ x'
```

```
## Warning: Removed 1 row containing missing values or values outside the scale range
## (`geom_text_repel()`).
```

```
## `geom_smooth()` using formula = 'y ~ x'
```

```
## Warning: Removed 1 row containing missing values or values outside the scale range
## (`geom_text_repel()`).
```

```
plot(p3)

#ggsave(file = 'results/fitness/fitnessEffects_summaryPlot_traitxMAT_2023.png', p3, height = 12, width = 14)
#ggsave(file = 'results/fitness/fitnessEffects_summaryPlot_traitxMAT_2023.pdf', p3, height = 12, width = 14)


# rename plots list for later use
plots.trait.mat.23 <- plots.summ
```

##### 2.1.2 Garden MAP

```
###################

# with garden MAP

# reorder gardens by garden MAP
tmp <- aggregate(dat$garden_MAP_2023, by = list(dat$MiniCG_Site_2023), mean)
ord <- tmp$Group.1[order(tmp$x, decreasing = T)]
dat$MiniCG_Site_2023 <- factor(dat$MiniCG_Site_2023, levels = ord)

sites.map <- dat$garden_MAP_2023[match(sites, dat$MiniCG_Site)]


# list to save results
mods.map <- list()
# list to save summary plot for each trait with significant effects on fitness
plots.summ <- list()

# loop through traits
for(n in 1:length(traits)){
  
  trait <- traits[n]
  
  print(paste('Model for:', trait))

  
  mod <- lmer(paste0('GrowthIncrement_2023 ~', trait,  '* garden_MAP_2023 + Pt + (1 | MiniCG_Site/block) + (1 | Genotype)'),
              data = dat)
  
  print(car::Anova(mod))
  
  p1 <- plot_model(mod, vline.color = "black", show.values = T, type = 'std', 
                   title = paste('2023 Growth ~ ', trait, sep = ''))
  plot(p1)

  mods.map[[n]] <- mod
  names(mods.map)[n] <- trait
  
  #####################
  # posthoc tests
  # if either the climate or trait x climate are significant, run posthoc tests for each garden
  mod.info <- Anova(mod)
  # if any coefficients are significant, plot by garden
  # need to change indexes if model is changed!
  # 1 is trait effect, 2 is garden climate effect, 4 is trait x climate effect.
  # Skip 3 which is ancestry - we already know this predicts height.
  
  if(mod.info$`Pr(>Chisq)`[1] <= 0.05 | mod.info$`Pr(>Chisq)`[2] <= 0.05 | mod.info$`Pr(>Chisq)`[4] <=0.05){
    
    # list to save plots for each site
    plots.sub <- list()
    
    # dataframe to save outputs from posthoc tests
    df.posthoc <- as.data.frame(matrix(ncol = 3, nrow = length(sites)))
    rownames(df.posthoc) <- sites
    colnames(df.posthoc) <- c('map', 'slope', 'pval')
    df.posthoc$map <- sites.map
    
    for(s in 1:length(sites)){
      
      site <- sites[s]
      dat.sub <- subset(dat, MiniCG_Site_2023 == site)
      # skip if site is missing data
      if(sum(!is.na(dat.sub[,trait])) > 10){
        mod.sub <- lmer(paste0('GrowthIncrement_2023 ~', trait, '+ Pt + (1 | block) + (1 | Genotype)'),
                        data = dat.sub)
        # get pval for trait
        mod.sub.info <- Anova(mod.sub)
        df.posthoc[site, 'pval'] <- mod.sub.info$`Pr(>Chisq)`[1]
        
        # get slopes from fixed effect of trait (effect on growth)
        df.posthoc[site, 'slope'] <- fixef(mod.sub)[2]
        
        # make title pretty
        maintitle <- paste(site, '\n', trait_names_df[trait, 'trait_names'], '\n', 'p = ', round(df.posthoc$pval[s], 3), sep = '')
        maintitle <- paste(strwrap(maintitle, width = 30), collapse = '\n')  
        
        # plot
        p <- ggplot(data = dat.sub, aes_string(y = 'GrowthIncrement_2023', x = trait, color = 'k2_tricho'), na.rm  = TRUE) +
          geom_point(na.rm = TRUE) +
          scale_color_gradient2(high = "darkolivegreen2", mid = "grey20", low = "dodgerblue2", midpoint = 0.5, name = 'P. trichocarpa\nancestry', guide = 'none') +
          ggtitle(maintitle) +
          geom_smooth(color = 'black', lty = ifelse(df.posthoc$pval[s] <= 0.05, 1,2), method = 'lm', formula = y ~ x, na.rm = TRUE) +
          theme(axis.text.x = element_text(angle = 90, vjust = 0.5, hjust=1, size = 12)) +
          theme(axis.title.x = element_blank())
        
        #plot(p)
        plots.sub[[s]] <- p
      }
    }

    # pvals are adjusted for summary plot
    df.posthoc$pvals.adj <- p.adjust(df.posthoc$pval, method = 'BH') 
    
    # plot panel with relationship for each garden
    # remove empty plots
    plots.sub <- plots.sub[lengths(plots.sub)>0]
    # plot in grid
    p1 <- grid.arrange(grobs = plots.sub, ncol = 4)
    plot(p1)
    
    # setup text showing significance
    sig_text <- paste0('Trait ', pval_stars(mod.info[1, 'Pr(>Chisq)']), '\n',
                       'Climate ', pval_stars(mod.info[2, 'Pr(>Chisq)']), '\n',
                       'Trait x Climate ', pval_stars(mod.info[4, 'Pr(>Chisq)']))
    
     # wrap title
    maintitle <- trait_names_df[trait, 'trait_names']
    maintitle <- paste(strwrap(maintitle, width = 30), collapse = '\n')
    
    # plot slope of fitness relationship vs climate
    p2 <- ggplot(df.posthoc, aes(x = map, y = slope), na.rm  = TRUE) +
            geom_smooth(method = 'lm', col = 'black', na.rm = TRUE) +
      geom_hline(yintercept = 0, lty = 2) +
      geom_point(size = 5, stroke = 2, shape = ifelse(df.posthoc$pvals.adj <= 0.05, 16, 1), aes(col = map), na.rm = TRUE) +
      scale_color_gradient2(high = "steelblue2",  mid = 'grey', low = "sienna3", midpoint = mean(range(sites.map)), limits = range_map,  name = 'Garden MAP') +
      ggtitle(maintitle,
              subtitle = sig_text) +
      xlab('Garden MAP') +
      ylab('Effect on Growth Increment') +
      geom_text_repel(label = rownames(df.posthoc), aes(x = map, y = slope), box.padding = 0.5, min.segment.length = 1, size = 4) +
      theme(plot.title = element_text(size = 15),
        plot.subtitle = element_text(size = 10),
        legend.position = 'none')
    plot(p2)
    
    plots.summ[[n]] <- p2
    names(plots.summ)[n] <- trait
    
    # add legend
    leg_plot <- ggplot(df.posthoc, aes(x = map, y = slope), na.rm  = TRUE) +
      geom_point(aes(color = map))+
      scale_color_gradient2(high = "steelblue2",  mid = 'grey', low = "sienna3", midpoint = mean(range(sites.map)), limits = range_map, name = 'Garden MAP (mm)') +
      theme(legend.position = 'right')
    
    legend <- cowplot::get_legend(leg_plot)
    
  }
}
```

```
## [1] "Model for: DOY_Stage2_2023"
```

```
## Warning: Some predictor variables are on very different scales: consider
## rescaling
```

```
## Analysis of Deviance Table (Type II Wald chisquare tests)
## 
## Response: GrowthIncrement_2023
##                                   Chisq Df Pr(>Chisq)    
## DOY_Stage2_2023                  5.9507  1    0.01471 *  
## garden_MAP_2023                  1.8602  1    0.17261    
## Pt                              42.8075  1   6.04e-11 ***
## DOY_Stage2_2023:garden_MAP_2023  0.0005  1    0.98257    
## ---
## Signif. codes:  0 '***' 0.001 '**' 0.01 '*' 0.05 '.' 0.1 ' ' 1
```

```
## boundary (singular) fit: see help('isSingular')
## boundary (singular) fit: see help('isSingular')
## boundary (singular) fit: see help('isSingular')
## boundary (singular) fit: see help('isSingular')
## boundary (singular) fit: see help('isSingular')
## boundary (singular) fit: see help('isSingular')
## boundary (singular) fit: see help('isSingular')
```

```
## `geom_smooth()` using formula = 'y ~ x'
```

```
## [1] "Model for: DOY_Stage3_2023"
```

```
## Warning: Some predictor variables are on very different scales: consider
## rescaling
```

```
## Analysis of Deviance Table (Type II Wald chisquare tests)
## 
## Response: GrowthIncrement_2023
##                                   Chisq Df Pr(>Chisq)    
## DOY_Stage3_2023                  3.5046  1     0.0612 .  
## garden_MAP_2023                  2.1109  1     0.1463    
## Pt                              34.6762  1  3.894e-09 ***
## DOY_Stage3_2023:garden_MAP_2023  0.0015  1     0.9692    
## ---
## Signif. codes:  0 '***' 0.001 '**' 0.01 '*' 0.05 '.' 0.1 ' ' 1
```

```
## [1] "Model for: DOY_Stage6_2023"
```

```
## Warning: Some predictor variables are on very different scales: consider
## rescaling
```

```
## Analysis of Deviance Table (Type II Wald chisquare tests)
## 
## Response: GrowthIncrement_2023
##                                   Chisq Df Pr(>Chisq)    
## DOY_Stage6_2023                 17.2409  1  3.293e-05 ***
## garden_MAP_2023                  0.7168  1  0.3971821    
## Pt                              38.1646  1  6.502e-10 ***
## DOY_Stage6_2023:garden_MAP_2023 13.7642  1  0.0002073 ***
## ---
## Signif. codes:  0 '***' 0.001 '**' 0.01 '*' 0.05 '.' 0.1 ' ' 1
```

```
## boundary (singular) fit: see help('isSingular')
## boundary (singular) fit: see help('isSingular')
## boundary (singular) fit: see help('isSingular')
## boundary (singular) fit: see help('isSingular')
## boundary (singular) fit: see help('isSingular')
## boundary (singular) fit: see help('isSingular')
```

```
## `geom_smooth()` using formula = 'y ~ x'
```

```
## Warning: Removed 1 row containing missing values or values outside the scale range
## (`geom_text_repel()`).
```

```
## Warning: Removed 1 row containing missing values or values outside the scale range
## (`geom_point()`).
```

```
## [1] "Model for: DOY_Stage7_2023"
```

```
## Warning: Some predictor variables are on very different scales: consider
## rescaling
```

```
## Analysis of Deviance Table (Type II Wald chisquare tests)
## 
## Response: GrowthIncrement_2023
##                                   Chisq Df Pr(>Chisq)    
## DOY_Stage7_2023                  1.5836  1     0.2082    
## garden_MAP_2023                  0.3129  1     0.5759    
## Pt                              36.1446  1  1.832e-09 ***
## DOY_Stage7_2023:garden_MAP_2023  0.8284  1     0.3627    
## ---
## Signif. codes:  0 '***' 0.001 '**' 0.01 '*' 0.05 '.' 0.1 ' ' 1
```

```
## [1] "Model for: stage7_presence_2023"
```

```
## Warning: Some predictor variables are on very different scales: consider
## rescaling
```

```
## Analysis of Deviance Table (Type II Wald chisquare tests)
## 
## Response: GrowthIncrement_2023
##                                        Chisq Df Pr(>Chisq)    
## stage7_presence_2023                  0.0052  1     0.9424    
## garden_MAP_2023                       1.5473  1     0.2135    
## Pt                                   46.6358  1  8.548e-12 ***
## stage7_presence_2023:garden_MAP_2023  0.0549  1     0.8148    
## ---
## Signif. codes:  0 '***' 0.001 '**' 0.01 '*' 0.05 '.' 0.1 ' ' 1
```

```
## [1] "Model for: DOY_last_budset_2023"
```

```
## Warning: Some predictor variables are on very different scales: consider
## rescaling
```

```
## Analysis of Deviance Table (Type II Wald chisquare tests)
## 
## Response: GrowthIncrement_2023
##                                        Chisq Df Pr(>Chisq)    
## DOY_last_budset_2023                  4.6481  1    0.03109 *  
## garden_MAP_2023                       0.8691  1    0.35120    
## Pt                                   44.1103  1  3.104e-11 ***
## DOY_last_budset_2023:garden_MAP_2023 51.8013  1  6.141e-13 ***
## ---
## Signif. codes:  0 '***' 0.001 '**' 0.01 '*' 0.05 '.' 0.1 ' ' 1
```

```
## boundary (singular) fit: see help('isSingular')
## boundary (singular) fit: see help('isSingular')
## boundary (singular) fit: see help('isSingular')
## boundary (singular) fit: see help('isSingular')
## boundary (singular) fit: see help('isSingular')
## boundary (singular) fit: see help('isSingular')
```

```
## `geom_smooth()` using formula = 'y ~ x'
```

```
## [1] "Model for: growing_season_days_2023"
```

```
## Warning: Some predictor variables are on very different scales: consider
## rescaling
```

```
## Analysis of Deviance Table (Type II Wald chisquare tests)
## 
## Response: GrowthIncrement_2023
##                                            Chisq Df Pr(>Chisq)    
## growing_season_days_2023                  1.1601  1    0.28145    
## garden_MAP_2023                           1.3973  1    0.23718    
## Pt                                       48.5682  1   3.19e-12 ***
## growing_season_days_2023:garden_MAP_2023  2.9739  1    0.08462 .  
## ---
## Signif. codes:  0 '***' 0.001 '**' 0.01 '*' 0.05 '.' 0.1 ' ' 1
```

```
## [1] "Model for: licor_gsw"
```

```
## Warning: Some predictor variables are on very different scales: consider
## rescaling
```

```
## Analysis of Deviance Table (Type II Wald chisquare tests)
## 
## Response: GrowthIncrement_2023
##                             Chisq Df Pr(>Chisq)    
## licor_gsw                  1.6789  1    0.19507    
## garden_MAP_2023            2.0401  1    0.15320    
## Pt                        53.6036  1  2.453e-13 ***
## licor_gsw:garden_MAP_2023  4.5093  1    0.03371 *  
## ---
## Signif. codes:  0 '***' 0.001 '**' 0.01 '*' 0.05 '.' 0.1 ' ' 1
```

```
## boundary (singular) fit: see help('isSingular')
## boundary (singular) fit: see help('isSingular')
## boundary (singular) fit: see help('isSingular')
## boundary (singular) fit: see help('isSingular')
## boundary (singular) fit: see help('isSingular')
```

```
## `geom_smooth()` using formula = 'y ~ x'
```

```
## Warning: Removed 3 rows containing missing values or values outside the scale range
## (`geom_text_repel()`).
```

```
## Warning: Removed 3 rows containing missing values or values outside the scale range
## (`geom_point()`).
```

```
## [1] "Model for: licor_gbw"
```

```
## Warning: Some predictor variables are on very different scales: consider
## rescaling
```

```
## Analysis of Deviance Table (Type II Wald chisquare tests)
## 
## Response: GrowthIncrement_2023
##                             Chisq Df Pr(>Chisq)    
## licor_gbw                  4.3283  1  0.0374833 *  
## garden_MAP_2023           11.8331  1  0.0005819 ***
## Pt                        56.4518  1  5.759e-14 ***
## licor_gbw:garden_MAP_2023 17.0712  1  3.600e-05 ***
## ---
## Signif. codes:  0 '***' 0.001 '**' 0.01 '*' 0.05 '.' 0.1 ' ' 1
```

```
## Warning: Some predictor variables are on very different scales: consider
## rescaling
```

```
## boundary (singular) fit: see help('isSingular')
```

```
## Warning: Some predictor variables are on very different scales: consider
## rescaling
```

```
## Warning: Some predictor variables are on very different scales: consider
## rescaling
```

```
## boundary (singular) fit: see help('isSingular')
```

```
## Warning: Some predictor variables are on very different scales: consider
## rescaling
```

```
## boundary (singular) fit: see help('isSingular')
```

```
## Warning: Some predictor variables are on very different scales: consider
## rescaling
## Warning: Some predictor variables are on very different scales: consider
## rescaling
## Warning: Some predictor variables are on very different scales: consider
## rescaling
```

```
## boundary (singular) fit: see help('isSingular')
```

```
## Warning: Some predictor variables are on very different scales: consider
## rescaling
```

```
## boundary (singular) fit: see help('isSingular')
```

```
## Warning: Some predictor variables are on very different scales: consider
## rescaling
```

```
## `geom_smooth()` using formula = 'y ~ x'
```

```
## Warning: Removed 3 rows containing missing values or values outside the scale range
## (`geom_text_repel()`).
```

```
## Warning: Removed 3 rows containing missing values or values outside the scale range
## (`geom_point()`).
```

```
## [1] "Model for: licor_ETR"
```

```
## Warning: Some predictor variables are on very different scales: consider
## rescaling
```

```
## Analysis of Deviance Table (Type II Wald chisquare tests)
## 
## Response: GrowthIncrement_2023
##                             Chisq Df Pr(>Chisq)    
## licor_ETR                  0.4717  1     0.4922    
## garden_MAP_2023            1.9926  1     0.1581    
## Pt                        56.2021  1  6.539e-14 ***
## licor_ETR:garden_MAP_2023  1.2007  1     0.2732    
## ---
## Signif. codes:  0 '***' 0.001 '**' 0.01 '*' 0.05 '.' 0.1 ' ' 1
```

```
## [1] "Model for: licor_PhiPS2"
```

```
## Warning: Some predictor variables are on very different scales: consider
## rescaling
```

```
## Analysis of Deviance Table (Type II Wald chisquare tests)
## 
## Response: GrowthIncrement_2023
##                                Chisq Df Pr(>Chisq)    
## licor_PhiPS2                  7.3857  1  0.0065743 ** 
## garden_MAP_2023               1.5046  1  0.2199650    
## Pt                           58.0149  1  2.601e-14 ***
## licor_PhiPS2:garden_MAP_2023 14.2108  1  0.0001634 ***
## ---
## Signif. codes:  0 '***' 0.001 '**' 0.01 '*' 0.05 '.' 0.1 ' ' 1
```

```
## boundary (singular) fit: see help('isSingular')
## boundary (singular) fit: see help('isSingular')
## boundary (singular) fit: see help('isSingular')
## boundary (singular) fit: see help('isSingular')
## boundary (singular) fit: see help('isSingular')
## boundary (singular) fit: see help('isSingular')
```

```
## `geom_smooth()` using formula = 'y ~ x'
```

```
## Warning: Removed 3 rows containing missing values or values outside the scale range
## (`geom_text_repel()`).
## Removed 3 rows containing missing values or values outside the scale range
## (`geom_point()`).
```

```
## [1] "Model for: licor_Fs"
```

```
## Warning: Some predictor variables are on very different scales: consider
## rescaling
```

```
## Analysis of Deviance Table (Type II Wald chisquare tests)
## 
## Response: GrowthIncrement_2023
##                            Chisq Df Pr(>Chisq)    
## licor_Fs                  4.9367  1   0.026293 *  
## garden_MAP_2023           2.0783  1   0.149406    
## Pt                       51.2115  1  8.293e-13 ***
## licor_Fs:garden_MAP_2023  7.7090  1   0.005494 ** 
## ---
## Signif. codes:  0 '***' 0.001 '**' 0.01 '*' 0.05 '.' 0.1 ' ' 1
```

```
## boundary (singular) fit: see help('isSingular')
## boundary (singular) fit: see help('isSingular')
## boundary (singular) fit: see help('isSingular')
## boundary (singular) fit: see help('isSingular')
## boundary (singular) fit: see help('isSingular')
```

```
## `geom_smooth()` using formula = 'y ~ x'
```

```
## Warning: Removed 3 rows containing missing values or values outside the scale range
## (`geom_text_repel()`).
## Removed 3 rows containing missing values or values outside the scale range
## (`geom_point()`).
```

```
## [1] "Model for: licor_Fm."
```

```
## Warning: Some predictor variables are on very different scales: consider
## rescaling
```

```
## Analysis of Deviance Table (Type II Wald chisquare tests)
## 
## Response: GrowthIncrement_2023
##                             Chisq Df Pr(>Chisq)    
## licor_Fm.                  1.0034  1     0.3165    
## garden_MAP_2023            1.9720  1     0.1602    
## Pt                        56.4627  1  5.727e-14 ***
## licor_Fm.:garden_MAP_2023  2.6734  1     0.1020    
## ---
## Signif. codes:  0 '***' 0.001 '**' 0.01 '*' 0.05 '.' 0.1 ' ' 1
```

```
## [1] "Model for: leaf_thickness_avg_mm_2023"
```

```
## Warning: Some predictor variables are on very different scales: consider
## rescaling
```

```
## Analysis of Deviance Table (Type II Wald chisquare tests)
## 
## Response: GrowthIncrement_2023
##                                              Chisq Df Pr(>Chisq)    
## leaf_thickness_avg_mm_2023                  4.5301  1     0.0333 *  
## garden_MAP_2023                             1.2757  1     0.2587    
## Pt                                         52.0428  1   5.43e-13 ***
## leaf_thickness_avg_mm_2023:garden_MAP_2023  0.6735  1     0.4118    
## ---
## Signif. codes:  0 '***' 0.001 '**' 0.01 '*' 0.05 '.' 0.1 ' ' 1
```

```
## boundary (singular) fit: see help('isSingular')
## boundary (singular) fit: see help('isSingular')
## boundary (singular) fit: see help('isSingular')
## boundary (singular) fit: see help('isSingular')
## boundary (singular) fit: see help('isSingular')
## boundary (singular) fit: see help('isSingular')
```

```
## `geom_smooth()` using formula = 'y ~ x'
```

```
## Warning: Removed 2 rows containing missing values or values outside the scale range
## (`geom_text_repel()`).
```

```
## Warning: Removed 2 rows containing missing values or values outside the scale range
## (`geom_point()`).
```

```
## [1] "Model for: LMA_g_m2_2023"
```

```
## Warning: Some predictor variables are on very different scales: consider
## rescaling
```

```
## Analysis of Deviance Table (Type II Wald chisquare tests)
## 
## Response: GrowthIncrement_2023
##                                 Chisq Df Pr(>Chisq)    
## LMA_g_m2_2023                  0.6998  1     0.4028    
## garden_MAP_2023                0.8165  1     0.3662    
## Pt                            34.8051  1  3.644e-09 ***
## LMA_g_m2_2023:garden_MAP_2023 46.2932  1  1.018e-11 ***
## ---
## Signif. codes:  0 '***' 0.001 '**' 0.01 '*' 0.05 '.' 0.1 ' ' 1
```

```
## boundary (singular) fit: see help('isSingular')
## boundary (singular) fit: see help('isSingular')
## boundary (singular) fit: see help('isSingular')
## boundary (singular) fit: see help('isSingular')
## boundary (singular) fit: see help('isSingular')
## boundary (singular) fit: see help('isSingular')
## boundary (singular) fit: see help('isSingular')
## boundary (singular) fit: see help('isSingular')
```

```
## `geom_smooth()` using formula = 'y ~ x'
```

```
## Warning: Removed 1 row containing missing values or values outside the scale range
## (`geom_text_repel()`).
```

```
## Warning: Removed 1 row containing missing values or values outside the scale range
## (`geom_point()`).
```

```
## [1] "Model for: leaf_mass_g_2023"
```

```
## Warning: Some predictor variables are on very different scales: consider
## rescaling
```

```
## Analysis of Deviance Table (Type II Wald chisquare tests)
## 
## Response: GrowthIncrement_2023
##                                    Chisq Df Pr(>Chisq)    
## leaf_mass_g_2023                 31.4261  1  2.072e-08 ***
## garden_MAP_2023                   0.7349  1  0.3913143    
## Pt                               25.6309  1  4.134e-07 ***
## leaf_mass_g_2023:garden_MAP_2023 12.6906  1  0.0003675 ***
## ---
## Signif. codes:  0 '***' 0.001 '**' 0.01 '*' 0.05 '.' 0.1 ' ' 1
```

```
## boundary (singular) fit: see help('isSingular')
## boundary (singular) fit: see help('isSingular')
## boundary (singular) fit: see help('isSingular')
## boundary (singular) fit: see help('isSingular')
## boundary (singular) fit: see help('isSingular')
## boundary (singular) fit: see help('isSingular')
## boundary (singular) fit: see help('isSingular')
## boundary (singular) fit: see help('isSingular')
```

```
## `geom_smooth()` using formula = 'y ~ x'
```

```
## Warning: Removed 1 row containing missing values or values outside the scale range
## (`geom_text_repel()`).
## Removed 1 row containing missing values or values outside the scale range
## (`geom_point()`).
```

```
## [1] "Model for: leaf_area_cm2_2023"
```

```
## Warning: Some predictor variables are on very different scales: consider
## rescaling
```

```
## Analysis of Deviance Table (Type II Wald chisquare tests)
## 
## Response: GrowthIncrement_2023
##                                      Chisq Df Pr(>Chisq)    
## leaf_area_cm2_2023                 41.9644  1  9.295e-11 ***
## garden_MAP_2023                     0.7772  1    0.37800    
## Pt                                 27.1642  1  1.869e-07 ***
## leaf_area_cm2_2023:garden_MAP_2023  4.2986  1    0.03814 *  
## ---
## Signif. codes:  0 '***' 0.001 '**' 0.01 '*' 0.05 '.' 0.1 ' ' 1
```

```
## boundary (singular) fit: see help('isSingular')
## boundary (singular) fit: see help('isSingular')
## boundary (singular) fit: see help('isSingular')
## boundary (singular) fit: see help('isSingular')
## boundary (singular) fit: see help('isSingular')
## boundary (singular) fit: see help('isSingular')
## boundary (singular) fit: see help('isSingular')
## boundary (singular) fit: see help('isSingular')
## boundary (singular) fit: see help('isSingular')
```

```
## `geom_smooth()` using formula = 'y ~ x'
```

```
## Warning: Removed 1 row containing missing values or values outside the scale range
## (`geom_text_repel()`).
## Removed 1 row containing missing values or values outside the scale range
## (`geom_point()`).
```

```
## [1] "Model for: lower_stomata_pore_length_mean_um"
```

```
## Warning: Some predictor variables are on very different scales: consider
## rescaling
```

```
## Analysis of Deviance Table (Type II Wald chisquare tests)
## 
## Response: GrowthIncrement_2023
##                                                     Chisq Df Pr(>Chisq)    
## lower_stomata_pore_length_mean_um                  1.1729  1   0.278800    
## garden_MAP_2023                                    0.8342  1   0.361057    
## Pt                                                40.4779  1  1.989e-10 ***
## lower_stomata_pore_length_mean_um:garden_MAP_2023  9.7699  1   0.001774 ** 
## ---
## Signif. codes:  0 '***' 0.001 '**' 0.01 '*' 0.05 '.' 0.1 ' ' 1
```

```
## boundary (singular) fit: see help('isSingular')
## boundary (singular) fit: see help('isSingular')
## boundary (singular) fit: see help('isSingular')
## boundary (singular) fit: see help('isSingular')
## boundary (singular) fit: see help('isSingular')
## boundary (singular) fit: see help('isSingular')
## boundary (singular) fit: see help('isSingular')
```

```
## `geom_smooth()` using formula = 'y ~ x'
```

```
## Warning: Removed 1 row containing missing values or values outside the scale range
## (`geom_text_repel()`).
## Removed 1 row containing missing values or values outside the scale range
## (`geom_point()`).
```

```
## [1] "Model for: upper_stomata_presence"
```

```
## Warning: Some predictor variables are on very different scales: consider
## rescaling
```

```
## Analysis of Deviance Table (Type II Wald chisquare tests)
## 
## Response: GrowthIncrement_2023
##                                          Chisq Df Pr(>Chisq)    
## upper_stomata_presence                  2.9228  1    0.08734 .  
## garden_MAP_2023                         0.8862  1    0.34650    
## Pt                                     41.6979  1  1.065e-10 ***
## upper_stomata_presence:garden_MAP_2023  1.5958  1    0.20650    
## ---
## Signif. codes:  0 '***' 0.001 '**' 0.01 '*' 0.05 '.' 0.1 ' ' 1
```

```
## [1] "Model for: upper_stomata_pore_length_mean_um"
```

```
## Warning: Some predictor variables are on very different scales: consider
## rescaling
```

```
## Analysis of Deviance Table (Type II Wald chisquare tests)
## 
## Response: GrowthIncrement_2023
##                                                     Chisq Df Pr(>Chisq)    
## upper_stomata_pore_length_mean_um                  3.5774  1    0.05857 .  
## garden_MAP_2023                                    1.0154  1    0.31361    
## Pt                                                21.1560  1  4.234e-06 ***
## upper_stomata_pore_length_mean_um:garden_MAP_2023  4.7487  1    0.02932 *  
## ---
## Signif. codes:  0 '***' 0.001 '**' 0.01 '*' 0.05 '.' 0.1 ' ' 1
```

```
## boundary (singular) fit: see help('isSingular')
## boundary (singular) fit: see help('isSingular')
## boundary (singular) fit: see help('isSingular')
## boundary (singular) fit: see help('isSingular')
## boundary (singular) fit: see help('isSingular')
## boundary (singular) fit: see help('isSingular')
## boundary (singular) fit: see help('isSingular')
## boundary (singular) fit: see help('isSingular')
```

```
## `geom_smooth()` using formula = 'y ~ x'
```

```
## Warning: Removed 1 row containing missing values or values outside the scale range
## (`geom_text_repel()`).
## Removed 1 row containing missing values or values outside the scale range
## (`geom_point()`).
```

```
## [1] "Model for: upper_stomata_density_mm2"
```

```
## Warning: Some predictor variables are on very different scales: consider
## rescaling
```

```
## Analysis of Deviance Table (Type II Wald chisquare tests)
## 
## Response: GrowthIncrement_2023
##                                             Chisq Df Pr(>Chisq)    
## upper_stomata_density_mm2                  1.1892  1     0.2755    
## garden_MAP_2023                            0.7046  1     0.4012    
## Pt                                        34.9964  1  3.303e-09 ***
## upper_stomata_density_mm2:garden_MAP_2023  1.8799  1     0.1703    
## ---
## Signif. codes:  0 '***' 0.001 '**' 0.01 '*' 0.05 '.' 0.1 ' ' 1
```

```
## [1] "Model for: lower_stomata_density_mm2"
```

```
## Warning: Some predictor variables are on very different scales: consider
## rescaling
```

```
## boundary (singular) fit: see help('isSingular')
```

```
## Analysis of Deviance Table (Type II Wald chisquare tests)
## 
## Response: GrowthIncrement_2023
##                                             Chisq Df Pr(>Chisq)    
## lower_stomata_density_mm2                  5.1521  1    0.02322 *  
## garden_MAP_2023                            0.8867  1    0.34637    
## Pt                                        37.7668  1  7.973e-10 ***
## lower_stomata_density_mm2:garden_MAP_2023  0.0439  1    0.83408    
## ---
## Signif. codes:  0 '***' 0.001 '**' 0.01 '*' 0.05 '.' 0.1 ' ' 1
```

```
## boundary (singular) fit: see help('isSingular')
```

```
## boundary (singular) fit: see help('isSingular')
## boundary (singular) fit: see help('isSingular')
## boundary (singular) fit: see help('isSingular')
## boundary (singular) fit: see help('isSingular')
## boundary (singular) fit: see help('isSingular')
## boundary (singular) fit: see help('isSingular')
## boundary (singular) fit: see help('isSingular')
```

```
## `geom_smooth()` using formula = 'y ~ x'
```

```
## Warning: Removed 1 row containing missing values or values outside the scale range
## (`geom_text_repel()`).
## Removed 1 row containing missing values or values outside the scale range
## (`geom_point()`).
```

```
## [1] "Model for: stomata_ratio"
```

```
## Warning: Some predictor variables are on very different scales: consider
## rescaling
```

```
## boundary (singular) fit: see help('isSingular')
```

```
## Analysis of Deviance Table (Type II Wald chisquare tests)
## 
## Response: GrowthIncrement_2023
##                                 Chisq Df Pr(>Chisq)    
## stomata_ratio                  2.7825  1     0.0953 .  
## garden_MAP_2023                0.4909  1     0.4835    
## Pt                            34.1986  1  4.977e-09 ***
## stomata_ratio:garden_MAP_2023  1.4236  1     0.2328    
## ---
## Signif. codes:  0 '***' 0.001 '**' 0.01 '*' 0.05 '.' 0.1 ' ' 1
```

```
## boundary (singular) fit: see help('isSingular')
```

```
# significant interaction for DPY stage6, DOY last budset, gsw, gbw, PhiPS2, Fs, LMA, leaf mass, leaf area, lower pore length, upper pore length
plots.summ <- plots.summ[lengths(plots.summ)>0]

# add legend to beginning of plot list

plots2 <- plots.summ
plots2[[length(plots.summ)+1]] <- legend

p3 <- grid.arrange(grobs = plots2, ncol = 4)
```

```
## `geom_smooth()` using formula = 'y ~ x'
```

```
## `geom_smooth()` using formula = 'y ~ x'
```

```
## Warning: Removed 1 row containing missing values or values outside the scale range
## (`geom_text_repel()`).
```

```
## `geom_smooth()` using formula = 'y ~ x'
## `geom_smooth()` using formula = 'y ~ x'
```

```
## Warning: Removed 3 rows containing missing values or values outside the scale range
## (`geom_text_repel()`).
```

```
## `geom_smooth()` using formula = 'y ~ x'
```

```
## Warning: Removed 3 rows containing missing values or values outside the scale range
## (`geom_text_repel()`).
```

```
## `geom_smooth()` using formula = 'y ~ x'
```

```
## Warning: Removed 3 rows containing missing values or values outside the scale range
## (`geom_text_repel()`).
```

```
## `geom_smooth()` using formula = 'y ~ x'
```

```
## Warning: Removed 3 rows containing missing values or values outside the scale range
## (`geom_text_repel()`).
```

```
## `geom_smooth()` using formula = 'y ~ x'
```

```
## Warning: Removed 2 rows containing missing values or values outside the scale range
## (`geom_text_repel()`).
```

```
## `geom_smooth()` using formula = 'y ~ x'
```

```
## Warning: Removed 1 row containing missing values or values outside the scale range
## (`geom_text_repel()`).
```

```
## `geom_smooth()` using formula = 'y ~ x'
```

```
## Warning: Removed 1 row containing missing values or values outside the scale range
## (`geom_text_repel()`).
```

```
## `geom_smooth()` using formula = 'y ~ x'
```

```
## Warning: Removed 1 row containing missing values or values outside the scale range
## (`geom_text_repel()`).
```

```
## `geom_smooth()` using formula = 'y ~ x'
```

```
## Warning: Removed 1 row containing missing values or values outside the scale range
## (`geom_text_repel()`).
```

```
## `geom_smooth()` using formula = 'y ~ x'
```

```
## Warning: Removed 1 row containing missing values or values outside the scale range
## (`geom_text_repel()`).
```

```
## `geom_smooth()` using formula = 'y ~ x'
```

```
## Warning: Removed 1 row containing missing values or values outside the scale range
## (`geom_text_repel()`).
```

```
## Warning: ggrepel: 5 unlabeled data points (too many overlaps). Consider
## increasing max.overlaps
```

```
plot(p3)
```

```
## Warning: ggrepel: 5 unlabeled data points (too many overlaps). Consider
## increasing max.overlaps
```

```
#ggsave(file = 'results/fitness/fitnessEffects_summaryPlot_traitxMAP_2023.png', p3, height = 12, width = 14)
#ggsave(file = 'results/fitness/fitnessEffects_summaryPlot_traitxMAP_2023.pdf', p3, height = 12, width = 14)

# rename plots list for later use
plots.trait.map.23 <- plots.summ
```

##### 2.1.3 Garden TD

```
###################


# with garden TD

# reorder gardens by garden TD
tmp <- aggregate(dat$garden_TD_2023, by = list(dat$MiniCG_Site_2023), mean)
ord <- tmp$Group.1[order(tmp$x, decreasing = T)]
dat$MiniCG_Site_2023 <- factor(dat$MiniCG_Site_2023, levels = ord)

sites.td <- dat$garden_TD_2023[match(sites, dat$MiniCG_Site)]


# list to save results
mods.td <- list()
# list to save summary plot for each trait with significant effects on fitness
plots.summ <- list()

# loop through traits
for(n in 1:length(traits)){
  
  trait <- traits[n]
  
  print(paste('Model for:', trait))

  
  mod <- lmer(paste0('GrowthIncrement_2023 ~', trait,  '* garden_TD_2023 + Pt + (1 | MiniCG_Site/block) + (1 | Genotype)'),
              data = dat)
  
  print(car::Anova(mod))
  
  p1 <- plot_model(mod, vline.color = "black", show.values = T, type = 'std', 
                   title = paste('2023 Growth ~ ', trait, sep = ''))
  plot(p1)

  mods.td[[n]] <- mod
  names(mods.td)[n] <- trait
  
  #####################
  # posthoc tests
  # if either the climate or trait x climate are significant, run posthoc tests for each garden
  mod.info <- Anova(mod)
  # if any coefficients are significant, plot by garden
  # need to change indexes if model is changed!
  # 1 is trait effect, 2 is garden climate effect, 4 is trait x climate effect.
  # Skip 3 which is ancestry - we already know this predicts height.
  
  if(mod.info$`Pr(>Chisq)`[1] <= 0.05 | mod.info$`Pr(>Chisq)`[2] <= 0.05 | mod.info$`Pr(>Chisq)`[4] <=0.05){
    
    # list to save plots for each site
    plots.sub <- list()
    
    # dataframe to save outputs from posthoc tests
    df.posthoc <- as.data.frame(matrix(ncol = 3, nrow = length(sites)))
    rownames(df.posthoc) <- sites
    colnames(df.posthoc) <- c('td', 'slope', 'pval')
    df.posthoc$td <- sites.td
    
    for(s in 1:length(sites)){
      
      site <- sites[s]
      dat.sub <- subset(dat, MiniCG_Site_2023 == site)
      # skip if site is missing data
      if(sum(!is.na(dat.sub[,trait])) > 10){
        mod.sub <- lmer(paste0('GrowthIncrement_2023 ~', trait, '+ Pt + (1 | block) + (1 | Genotype)'),
                        data = dat.sub)
        # get pval for trait
        mod.sub.info <- Anova(mod.sub)
        df.posthoc[site, 'pval'] <- mod.sub.info$`Pr(>Chisq)`[1]
        
        # get slopes from fixed effect of trait (effect on growth)
        df.posthoc[site, 'slope'] <- fixef(mod.sub)[2]
        
        # make title pretty
        maintitle <- paste(site, '\n', trait_names_df[trait, 'trait_names'], '\n', 'p = ', round(df.posthoc$pval[s], 3), sep = '')
        maintitle <- paste(strwrap(maintitle, width = 30), collapse = '\n')  
        
        # plot
        p <- ggplot(data = dat.sub, aes_string(y = 'GrowthIncrement_2023', x = trait, color = 'k2_tricho'), na.rm  = TRUE) +
          geom_point(na.rm = TRUE) +
          scale_color_gradient2(high = "darkolivegreen2", mid = "grey20", low = "dodgerblue2", midpoint = 0.5, name = 'P. trichocarpa\nancestry', guide = 'none') +
          ggtitle(maintitle) +
          geom_smooth(color = 'black', lty = ifelse(df.posthoc$pval[s] <= 0.05, 1,2), method = 'lm', formula = y ~ x, na.rm = TRUE) +
          theme(axis.text.x = element_text(angle = 90, vjust = 0.5, hjust=1, size = 12)) +
          theme(axis.title.x = element_blank())
        
        #plot(p)
        plots.sub[[s]] <- p
      }
    }

    # pvals are adjusted for summary plot
    df.posthoc$pvals.adj <- p.adjust(df.posthoc$pval, method = 'BH') 
    
    # plot panel with relationship for each garden
    # remove empty plots
    plots.sub <- plots.sub[lengths(plots.sub)>0]
    # plot in grid
    p1 <- grid.arrange(grobs = plots.sub, ncol = 4)
    plot(p1)
    
    # setup text showing significance
    sig_text <- paste0('Trait ', pval_stars(mod.info[1, 'Pr(>Chisq)']), '\n',
                       'Climate ', pval_stars(mod.info[2, 'Pr(>Chisq)']), '\n',
                       'Trait x Climate ', pval_stars(mod.info[4, 'Pr(>Chisq)']))
    
     # wrap title
    maintitle <- trait_names_df[trait, 'trait_names']
    maintitle <- paste(strwrap(maintitle, width = 30), collapse = '\n')
    
    # plot slope of fitness relationship vs climate
    p2 <- ggplot(df.posthoc, aes(x = td, y = slope), na.rm  = TRUE) +
      geom_smooth(method = 'lm', col = 'black', na.rm = TRUE) +
      geom_hline(yintercept = 0, lty = 2) +
      geom_point(size = 5, stroke = 2, shape = ifelse(df.posthoc$pvals.adj <= 0.05, 16, 1), aes(col = td), na.rm = TRUE) +
      scale_color_gradient2(high = "orchid4",  mid = 'grey', low = "palegreen4", midpoint = mean(range(sites.td)), limits = range_td, name = 'Garden TD (°C)') +
      ggtitle(maintitle,
              subtitle = sig_text) +
      xlab('Garden TD') +
      ylab('Effect on Growth Increment') +
      geom_text_repel(label = rownames(df.posthoc), aes(x = td, y = slope), box.padding = 0.5, min.segment.length = 1, size = 4) +
      theme(plot.title = element_text(size = 15),
            plot.subtitle = element_text(size = 10),
            legend.position = 'none')
    plot(p2)
    
    plots.summ[[n]] <- p2
    names(plots.summ)[n] <- trait
    
     # add legend
    leg_plot <- ggplot(df.posthoc, aes(x = td, y = slope), na.rm  = TRUE) +
      geom_point(aes(color = td))+
      scale_color_gradient2(high = "orchid4",  mid = 'grey', low = "palegreen4", midpoint = mean(range(sites.td)), limits = range_td, name = 'Garden TD (°C)') +
      theme(legend.position = 'right')
    
    legend <- cowplot::get_legend(leg_plot)
    
  }
}
```

```
## [1] "Model for: DOY_Stage2_2023"
```

```
## Warning: Some predictor variables are on very different scales: consider
## rescaling
```

```
## Analysis of Deviance Table (Type II Wald chisquare tests)
## 
## Response: GrowthIncrement_2023
##                                  Chisq Df Pr(>Chisq)    
## DOY_Stage2_2023                 4.9744  1    0.02573 *  
## garden_TD_2023                  0.0056  1    0.94040    
## Pt                             41.7348  1  1.045e-10 ***
## DOY_Stage2_2023:garden_TD_2023  0.1093  1    0.74092    
## ---
## Signif. codes:  0 '***' 0.001 '**' 0.01 '*' 0.05 '.' 0.1 ' ' 1
```

```
## boundary (singular) fit: see help('isSingular')
## boundary (singular) fit: see help('isSingular')
## boundary (singular) fit: see help('isSingular')
## boundary (singular) fit: see help('isSingular')
## boundary (singular) fit: see help('isSingular')
## boundary (singular) fit: see help('isSingular')
## boundary (singular) fit: see help('isSingular')
```

```
## `geom_smooth()` using formula = 'y ~ x'
```

```
## [1] "Model for: DOY_Stage3_2023"
```

```
## Warning: Some predictor variables are on very different scales: consider
## rescaling
```

```
## Analysis of Deviance Table (Type II Wald chisquare tests)
## 
## Response: GrowthIncrement_2023
##                                  Chisq Df Pr(>Chisq)    
## DOY_Stage3_2023                 2.5868  1     0.1078    
## garden_TD_2023                  0.0122  1     0.9121    
## Pt                             33.8287  1  6.018e-09 ***
## DOY_Stage3_2023:garden_TD_2023  0.0642  1     0.8000    
## ---
## Signif. codes:  0 '***' 0.001 '**' 0.01 '*' 0.05 '.' 0.1 ' ' 1
```

```
## [1] "Model for: DOY_Stage6_2023"
```

```
## Warning: Some predictor variables are on very different scales: consider
## rescaling
```

```
## Analysis of Deviance Table (Type II Wald chisquare tests)
## 
## Response: GrowthIncrement_2023
##                                  Chisq Df Pr(>Chisq)    
## DOY_Stage6_2023                17.4138  1  3.006e-05 ***
## garden_TD_2023                  0.1285  1     0.7200    
## Pt                             34.2299  1  4.897e-09 ***
## DOY_Stage6_2023:garden_TD_2023  0.9594  1     0.3273    
## ---
## Signif. codes:  0 '***' 0.001 '**' 0.01 '*' 0.05 '.' 0.1 ' ' 1
```

```
## boundary (singular) fit: see help('isSingular')
## boundary (singular) fit: see help('isSingular')
## boundary (singular) fit: see help('isSingular')
## boundary (singular) fit: see help('isSingular')
## boundary (singular) fit: see help('isSingular')
## boundary (singular) fit: see help('isSingular')
```

```
## `geom_smooth()` using formula = 'y ~ x'
```

```
## Warning: Removed 1 row containing missing values or values outside the scale range
## (`geom_text_repel()`).
```

```
## Warning: Removed 1 row containing missing values or values outside the scale range
## (`geom_point()`).
```

```
## [1] "Model for: DOY_Stage7_2023"
```

```
## Warning: Some predictor variables are on very different scales: consider
## rescaling
```

```
## Analysis of Deviance Table (Type II Wald chisquare tests)
## 
## Response: GrowthIncrement_2023
##                                  Chisq Df Pr(>Chisq)    
## DOY_Stage7_2023                 1.7807  1     0.1821    
## garden_TD_2023                  0.0201  1     0.8872    
## Pt                             37.2515  1  1.038e-09 ***
## DOY_Stage7_2023:garden_TD_2023  2.6725  1     0.1021    
## ---
## Signif. codes:  0 '***' 0.001 '**' 0.01 '*' 0.05 '.' 0.1 ' ' 1
```

```
## [1] "Model for: stage7_presence_2023"
## Analysis of Deviance Table (Type II Wald chisquare tests)
## 
## Response: GrowthIncrement_2023
##                                       Chisq Df Pr(>Chisq)    
## stage7_presence_2023                 0.0029  1     0.9571    
## garden_TD_2023                       0.0141  1     0.9056    
## Pt                                  46.4612  1  9.345e-12 ***
## stage7_presence_2023:garden_TD_2023  0.0022  1     0.9624    
## ---
## Signif. codes:  0 '***' 0.001 '**' 0.01 '*' 0.05 '.' 0.1 ' ' 1
```

```
## [1] "Model for: DOY_last_budset_2023"
```

```
## Warning: Some predictor variables are on very different scales: consider
## rescaling
```

```
## Analysis of Deviance Table (Type II Wald chisquare tests)
## 
## Response: GrowthIncrement_2023
##                                       Chisq Df Pr(>Chisq)    
## DOY_last_budset_2023                 3.2845  1    0.06994 .  
## garden_TD_2023                       0.0107  1    0.91762    
## Pt                                  46.0244  1  1.168e-11 ***
## DOY_last_budset_2023:garden_TD_2023  1.7587  1    0.18479    
## ---
## Signif. codes:  0 '***' 0.001 '**' 0.01 '*' 0.05 '.' 0.1 ' ' 1
```

```
## [1] "Model for: growing_season_days_2023"
```

```
## Warning: Some predictor variables are on very different scales: consider
## rescaling
```

```
## Analysis of Deviance Table (Type II Wald chisquare tests)
## 
## Response: GrowthIncrement_2023
##                                           Chisq Df Pr(>Chisq)    
## growing_season_days_2023                 0.9177  1     0.3381    
## garden_TD_2023                           0.0143  1     0.9047    
## Pt                                      46.8202  1  7.781e-12 ***
## growing_season_days_2023:garden_TD_2023  0.1332  1     0.7151    
## ---
## Signif. codes:  0 '***' 0.001 '**' 0.01 '*' 0.05 '.' 0.1 ' ' 1
```

```
## [1] "Model for: licor_gsw"
## Analysis of Deviance Table (Type II Wald chisquare tests)
## 
## Response: GrowthIncrement_2023
##                            Chisq Df Pr(>Chisq)    
## licor_gsw                 1.7280  1    0.18867    
## garden_TD_2023            1.3307  1    0.24868    
## Pt                       55.1468  1  1.119e-13 ***
## licor_gsw:garden_TD_2023  4.9585  1    0.02596 *  
## ---
## Signif. codes:  0 '***' 0.001 '**' 0.01 '*' 0.05 '.' 0.1 ' ' 1
```

```
## boundary (singular) fit: see help('isSingular')
## boundary (singular) fit: see help('isSingular')
## boundary (singular) fit: see help('isSingular')
## boundary (singular) fit: see help('isSingular')
## boundary (singular) fit: see help('isSingular')
```

```
## `geom_smooth()` using formula = 'y ~ x'
```

```
## Warning: Removed 3 rows containing missing values or values outside the scale range
## (`geom_text_repel()`).
```

```
## Warning: Removed 3 rows containing missing values or values outside the scale range
## (`geom_point()`).
```

```
## [1] "Model for: licor_gbw"
```

```
## Warning: Some predictor variables are on very different scales: consider
## rescaling
```

```
## Analysis of Deviance Table (Type II Wald chisquare tests)
## 
## Response: GrowthIncrement_2023
##                            Chisq Df Pr(>Chisq)    
## licor_gbw                 0.0141  1     0.9056    
## garden_TD_2023            1.1683  1     0.2798    
## Pt                       54.5203  1  1.538e-13 ***
## licor_gbw:garden_TD_2023  0.2660  1     0.6060    
## ---
## Signif. codes:  0 '***' 0.001 '**' 0.01 '*' 0.05 '.' 0.1 ' ' 1
```

```
## [1] "Model for: licor_ETR"
```

```
## Warning: Some predictor variables are on very different scales: consider
## rescaling
```

```
## Analysis of Deviance Table (Type II Wald chisquare tests)
## 
## Response: GrowthIncrement_2023
##                            Chisq Df Pr(>Chisq)    
## licor_ETR                 0.5795  1    0.44650    
## garden_TD_2023            1.1521  1    0.28311    
## Pt                       58.5249  1  2.007e-14 ***
## licor_ETR:garden_TD_2023  2.9621  1    0.08524 .  
## ---
## Signif. codes:  0 '***' 0.001 '**' 0.01 '*' 0.05 '.' 0.1 ' ' 1
```

```
## [1] "Model for: licor_PhiPS2"
## Analysis of Deviance Table (Type II Wald chisquare tests)
## 
## Response: GrowthIncrement_2023
##                               Chisq Df Pr(>Chisq)    
## licor_PhiPS2                 8.1950  1   0.004201 ** 
## garden_TD_2023               1.5481  1   0.213413    
## Pt                          56.2212  1  6.476e-14 ***
## licor_PhiPS2:garden_TD_2023  0.0730  1   0.787001    
## ---
## Signif. codes:  0 '***' 0.001 '**' 0.01 '*' 0.05 '.' 0.1 ' ' 1
```

```
## boundary (singular) fit: see help('isSingular')
## boundary (singular) fit: see help('isSingular')
## boundary (singular) fit: see help('isSingular')
## boundary (singular) fit: see help('isSingular')
## boundary (singular) fit: see help('isSingular')
## boundary (singular) fit: see help('isSingular')
```

```
## `geom_smooth()` using formula = 'y ~ x'
```

```
## Warning: Removed 3 rows containing missing values or values outside the scale range
## (`geom_text_repel()`).
## Removed 3 rows containing missing values or values outside the scale range
## (`geom_point()`).
```

```
## [1] "Model for: licor_Fs"
```

```
## Warning: Some predictor variables are on very different scales: consider
## rescaling
```

```
## Analysis of Deviance Table (Type II Wald chisquare tests)
## 
## Response: GrowthIncrement_2023
##                           Chisq Df Pr(>Chisq)    
## licor_Fs                 4.9545  1    0.02602 *  
## garden_TD_2023           1.1995  1    0.27343    
## Pt                      52.2240  1  4.952e-13 ***
## licor_Fs:garden_TD_2023  1.8112  1    0.17837    
## ---
## Signif. codes:  0 '***' 0.001 '**' 0.01 '*' 0.05 '.' 0.1 ' ' 1
```

```
## boundary (singular) fit: see help('isSingular')
## boundary (singular) fit: see help('isSingular')
## boundary (singular) fit: see help('isSingular')
## boundary (singular) fit: see help('isSingular')
## boundary (singular) fit: see help('isSingular')
```

```
## `geom_smooth()` using formula = 'y ~ x'
```

```
## Warning: Removed 3 rows containing missing values or values outside the scale range
## (`geom_text_repel()`).
## Removed 3 rows containing missing values or values outside the scale range
## (`geom_point()`).
```

```
## [1] "Model for: licor_Fm."
```

```
## Warning: Some predictor variables are on very different scales: consider
## rescaling
```

```
## Analysis of Deviance Table (Type II Wald chisquare tests)
## 
## Response: GrowthIncrement_2023
##                            Chisq Df Pr(>Chisq)    
## licor_Fm.                 1.4895  1     0.2223    
## garden_TD_2023            1.3571  1     0.2440    
## Pt                       57.0002  1  4.358e-14 ***
## licor_Fm.:garden_TD_2023  1.1096  1     0.2922    
## ---
## Signif. codes:  0 '***' 0.001 '**' 0.01 '*' 0.05 '.' 0.1 ' ' 1
```

```
## [1] "Model for: leaf_thickness_avg_mm_2023"
## Analysis of Deviance Table (Type II Wald chisquare tests)
## 
## Response: GrowthIncrement_2023
##                                             Chisq Df Pr(>Chisq)    
## leaf_thickness_avg_mm_2023                 4.5947  1    0.03207 *  
## garden_TD_2023                             0.4991  1    0.47991    
## Pt                                        50.8713  1  9.862e-13 ***
## leaf_thickness_avg_mm_2023:garden_TD_2023  1.4352  1    0.23091    
## ---
## Signif. codes:  0 '***' 0.001 '**' 0.01 '*' 0.05 '.' 0.1 ' ' 1
```

```
## boundary (singular) fit: see help('isSingular')
## boundary (singular) fit: see help('isSingular')
## boundary (singular) fit: see help('isSingular')
## boundary (singular) fit: see help('isSingular')
## boundary (singular) fit: see help('isSingular')
## boundary (singular) fit: see help('isSingular')
```

```
## `geom_smooth()` using formula = 'y ~ x'
```

```
## Warning: Removed 2 rows containing missing values or values outside the scale range
## (`geom_text_repel()`).
```

```
## Warning: Removed 2 rows containing missing values or values outside the scale range
## (`geom_point()`).
```

```
## [1] "Model for: LMA_g_m2_2023"
## Analysis of Deviance Table (Type II Wald chisquare tests)
## 
## Response: GrowthIncrement_2023
##                                Chisq Df Pr(>Chisq)    
## LMA_g_m2_2023                 0.7031  1  0.4017356    
## garden_TD_2023                0.2031  1  0.6522166    
## Pt                           36.3641  1  1.637e-09 ***
## LMA_g_m2_2023:garden_TD_2023 11.9185  1  0.0005558 ***
## ---
## Signif. codes:  0 '***' 0.001 '**' 0.01 '*' 0.05 '.' 0.1 ' ' 1
```

```
## boundary (singular) fit: see help('isSingular')
## boundary (singular) fit: see help('isSingular')
## boundary (singular) fit: see help('isSingular')
## boundary (singular) fit: see help('isSingular')
## boundary (singular) fit: see help('isSingular')
## boundary (singular) fit: see help('isSingular')
## boundary (singular) fit: see help('isSingular')
## boundary (singular) fit: see help('isSingular')
```

```
## `geom_smooth()` using formula = 'y ~ x'
```

```
## Warning: Removed 1 row containing missing values or values outside the scale range
## (`geom_text_repel()`).
```

```
## Warning: Removed 1 row containing missing values or values outside the scale range
## (`geom_point()`).
```

```
## [1] "Model for: leaf_mass_g_2023"
## Analysis of Deviance Table (Type II Wald chisquare tests)
## 
## Response: GrowthIncrement_2023
##                                   Chisq Df Pr(>Chisq)    
## leaf_mass_g_2023                30.7851  1  2.882e-08 ***
## garden_TD_2023                   0.2935  1     0.5880    
## Pt                              28.4132  1  9.799e-08 ***
## leaf_mass_g_2023:garden_TD_2023  0.9803  1     0.3221    
## ---
## Signif. codes:  0 '***' 0.001 '**' 0.01 '*' 0.05 '.' 0.1 ' ' 1
```

```
## boundary (singular) fit: see help('isSingular')
## boundary (singular) fit: see help('isSingular')
## boundary (singular) fit: see help('isSingular')
## boundary (singular) fit: see help('isSingular')
## boundary (singular) fit: see help('isSingular')
## boundary (singular) fit: see help('isSingular')
## boundary (singular) fit: see help('isSingular')
## boundary (singular) fit: see help('isSingular')
```

```
## `geom_smooth()` using formula = 'y ~ x'
```

```
## Warning: Removed 1 row containing missing values or values outside the scale range
## (`geom_text_repel()`).
## Removed 1 row containing missing values or values outside the scale range
## (`geom_point()`).
```

```
## [1] "Model for: leaf_area_cm2_2023"
```

```
## Warning: Some predictor variables are on very different scales: consider
## rescaling
```

```
## Analysis of Deviance Table (Type II Wald chisquare tests)
## 
## Response: GrowthIncrement_2023
##                                     Chisq Df Pr(>Chisq)    
## leaf_area_cm2_2023                42.2661  1  7.966e-11 ***
## garden_TD_2023                     0.3719  1     0.5420    
## Pt                                28.7041  1  8.432e-08 ***
## leaf_area_cm2_2023:garden_TD_2023  0.2213  1     0.6381    
## ---
## Signif. codes:  0 '***' 0.001 '**' 0.01 '*' 0.05 '.' 0.1 ' ' 1
```

```
## boundary (singular) fit: see help('isSingular')
## boundary (singular) fit: see help('isSingular')
## boundary (singular) fit: see help('isSingular')
## boundary (singular) fit: see help('isSingular')
## boundary (singular) fit: see help('isSingular')
## boundary (singular) fit: see help('isSingular')
## boundary (singular) fit: see help('isSingular')
## boundary (singular) fit: see help('isSingular')
## boundary (singular) fit: see help('isSingular')
```

```
## `geom_smooth()` using formula = 'y ~ x'
```

```
## Warning: Removed 1 row containing missing values or values outside the scale range
## (`geom_text_repel()`).
## Removed 1 row containing missing values or values outside the scale range
## (`geom_point()`).
```

```
## [1] "Model for: lower_stomata_pore_length_mean_um"
## Analysis of Deviance Table (Type II Wald chisquare tests)
## 
## Response: GrowthIncrement_2023
##                                                    Chisq Df Pr(>Chisq)    
## lower_stomata_pore_length_mean_um                 1.2532  1    0.26294    
## garden_TD_2023                                    0.2413  1    0.62330    
## Pt                                               40.4756  1  1.991e-10 ***
## lower_stomata_pore_length_mean_um:garden_TD_2023  3.0914  1    0.07871 .  
## ---
## Signif. codes:  0 '***' 0.001 '**' 0.01 '*' 0.05 '.' 0.1 ' ' 1
```

```
## [1] "Model for: upper_stomata_presence"
## Analysis of Deviance Table (Type II Wald chisquare tests)
## 
## Response: GrowthIncrement_2023
##                                         Chisq Df Pr(>Chisq)    
## upper_stomata_presence                 2.9399  1    0.08642 .  
## garden_TD_2023                         0.2228  1    0.63692    
## Pt                                    41.3322  1  1.284e-10 ***
## upper_stomata_presence:garden_TD_2023  0.3100  1    0.57768    
## ---
## Signif. codes:  0 '***' 0.001 '**' 0.01 '*' 0.05 '.' 0.1 ' ' 1
```

```
## [1] "Model for: upper_stomata_pore_length_mean_um"
## Analysis of Deviance Table (Type II Wald chisquare tests)
## 
## Response: GrowthIncrement_2023
##                                                    Chisq Df Pr(>Chisq)    
## upper_stomata_pore_length_mean_um                 3.9045  1    0.04816 *  
## garden_TD_2023                                    0.3183  1    0.57261    
## Pt                                               20.4540  1  6.108e-06 ***
## upper_stomata_pore_length_mean_um:garden_TD_2023  0.8359  1    0.36057    
## ---
## Signif. codes:  0 '***' 0.001 '**' 0.01 '*' 0.05 '.' 0.1 ' ' 1
```

```
## boundary (singular) fit: see help('isSingular')
## boundary (singular) fit: see help('isSingular')
## boundary (singular) fit: see help('isSingular')
## boundary (singular) fit: see help('isSingular')
## boundary (singular) fit: see help('isSingular')
## boundary (singular) fit: see help('isSingular')
## boundary (singular) fit: see help('isSingular')
## boundary (singular) fit: see help('isSingular')
```

```
## `geom_smooth()` using formula = 'y ~ x'
```

```
## Warning: Removed 1 row containing missing values or values outside the scale range
## (`geom_text_repel()`).
## Removed 1 row containing missing values or values outside the scale range
## (`geom_point()`).
```

```
## [1] "Model for: upper_stomata_density_mm2"
```

```
## Warning: Some predictor variables are on very different scales: consider
## rescaling
```

```
## Analysis of Deviance Table (Type II Wald chisquare tests)
## 
## Response: GrowthIncrement_2023
##                                            Chisq Df Pr(>Chisq)    
## upper_stomata_density_mm2                 1.2281  1     0.2678    
## garden_TD_2023                            0.2784  1     0.5977    
## Pt                                       34.3217  1  4.671e-09 ***
## upper_stomata_density_mm2:garden_TD_2023  0.0824  1     0.7741    
## ---
## Signif. codes:  0 '***' 0.001 '**' 0.01 '*' 0.05 '.' 0.1 ' ' 1
```

```
## [1] "Model for: lower_stomata_density_mm2"
```

```
## Warning: Some predictor variables are on very different scales: consider
## rescaling
```

```
## boundary (singular) fit: see help('isSingular')
```

```
## Analysis of Deviance Table (Type II Wald chisquare tests)
## 
## Response: GrowthIncrement_2023
##                                            Chisq Df Pr(>Chisq)    
## lower_stomata_density_mm2                 4.9624  1     0.0259 *  
## garden_TD_2023                            0.3538  1     0.5520    
## Pt                                       37.9769  1  7.159e-10 ***
## lower_stomata_density_mm2:garden_TD_2023  0.3233  1     0.5696    
## ---
## Signif. codes:  0 '***' 0.001 '**' 0.01 '*' 0.05 '.' 0.1 ' ' 1
```

```
## boundary (singular) fit: see help('isSingular')
```

```
## boundary (singular) fit: see help('isSingular')
## boundary (singular) fit: see help('isSingular')
## boundary (singular) fit: see help('isSingular')
## boundary (singular) fit: see help('isSingular')
## boundary (singular) fit: see help('isSingular')
## boundary (singular) fit: see help('isSingular')
## boundary (singular) fit: see help('isSingular')
```

```
## `geom_smooth()` using formula = 'y ~ x'
```

```
## Warning: Removed 1 row containing missing values or values outside the scale range
## (`geom_text_repel()`).
## Removed 1 row containing missing values or values outside the scale range
## (`geom_point()`).
```

```
## [1] "Model for: stomata_ratio"
```

```
## boundary (singular) fit: see help('isSingular')
```

```
## Analysis of Deviance Table (Type II Wald chisquare tests)
## 
## Response: GrowthIncrement_2023
##                                Chisq Df Pr(>Chisq)    
## stomata_ratio                 2.8283  1    0.09262 .  
## garden_TD_2023                0.3353  1    0.56257    
## Pt                           33.8831  1  5.852e-09 ***
## stomata_ratio:garden_TD_2023  0.0140  1    0.90567    
## ---
## Signif. codes:  0 '***' 0.001 '**' 0.01 '*' 0.05 '.' 0.1 ' ' 1
```

```
## boundary (singular) fit: see help('isSingular')
```

```
# significant interaction for DPY stage6, DOY last budset, gsw, gbw, PhiPS2, Fs, LMA, leaf mass, leaf area, lower pore length, upper pore length
plots.summ <- plots.summ[lengths(plots.summ)>0]

# add legend to beginning of plot list

plots2 <- plots.summ
plots2[[length(plots.summ)+1]] <- legend

p3 <- grid.arrange(grobs = plots2, ncol = 4)
```

```
## `geom_smooth()` using formula = 'y ~ x'
```

```
## `geom_smooth()` using formula = 'y ~ x'
```

```
## Warning: Removed 1 row containing missing values or values outside the scale range
## (`geom_text_repel()`).
```

```
## `geom_smooth()` using formula = 'y ~ x'
```

```
## Warning: Removed 3 rows containing missing values or values outside the scale range
## (`geom_text_repel()`).
```

```
## `geom_smooth()` using formula = 'y ~ x'
```

```
## Warning: Removed 3 rows containing missing values or values outside the scale range
## (`geom_text_repel()`).
```

```
## `geom_smooth()` using formula = 'y ~ x'
```

```
## Warning: Removed 3 rows containing missing values or values outside the scale range
## (`geom_text_repel()`).
```

```
## `geom_smooth()` using formula = 'y ~ x'
```

```
## Warning: Removed 2 rows containing missing values or values outside the scale range
## (`geom_text_repel()`).
```

```
## `geom_smooth()` using formula = 'y ~ x'
```

```
## Warning: Removed 1 row containing missing values or values outside the scale range
## (`geom_text_repel()`).
```

```
## `geom_smooth()` using formula = 'y ~ x'
```

```
## Warning: Removed 1 row containing missing values or values outside the scale range
## (`geom_text_repel()`).
```

```
## `geom_smooth()` using formula = 'y ~ x'
```

```
## Warning: Removed 1 row containing missing values or values outside the scale range
## (`geom_text_repel()`).
```

```
## `geom_smooth()` using formula = 'y ~ x'
```

```
## Warning: Removed 1 row containing missing values or values outside the scale range
## (`geom_text_repel()`).
```

```
## `geom_smooth()` using formula = 'y ~ x'
```

```
## Warning: Removed 1 row containing missing values or values outside the scale range
## (`geom_text_repel()`).
```

```
plot(p3)

#ggsave(file = 'results/fitness/fitnessEffects_summaryPlot_traitxTD_2023.png', p3, height = 10, width = 14)
#ggsave(file = 'results/fitness/fitnessEffects_summaryPlot_traitxTD_2023.pdf', p3, height = 10, width = 14)

# rename plots list for later use
plots.trait.td.23 <- plots.summ
```

#### 2.2 Do trait values predict growth in 2024?

```
# get 2024 traits
traits <- c("DOY_Stage2_2024", "DOY_Stage3_2024", "DOY_Stage6_2024", "DOY_Stage7_2024", "stage7_presence_2024", "DOY_last_budset_2024", "growing_season_days_2024")


# SWMN was only checked once for stage6 - remove these points
dat[dat$MiniCG_Site == 'SWMN', 'DOY_Stage6_2024'] <- NA
dat[dat$MiniCG_Site == 'SWMN', 'DOY_last_budset_2024'] <- NA

# do these traits predict growth increment?
```

##### 2.2.1 Garden MAT

```
# reorder gardens by garden MAT
tmp <- aggregate(dat$garden_MAT_2024, by = list(dat$MiniCG_Site_2024), mean)
ord <- tmp$Group.1[order(tmp$x, decreasing = T)]
dat$MiniCG_Site_2024 <- factor(dat$MiniCG_Site_2024, levels = ord)

# sites <- levels(dat$MiniCG_Site_2024)
sites <- c("WI", "ID", "WYO", "EVERGREEN", "MSU", "MORTON", "PENN", "SU")

sites.mat <- dat$garden_MAT_2024[match(sites, dat$MiniCG_Site)]

# list to save results
mods.mat <- list()
# list to save summary plot for each trait with significant effects on fitness
plots.summ <- list()


# loop through traits
for(n in 1:length(traits)){
  
  trait <- traits[n]
  
  print(paste('Model for:', trait))
  
  
  mod <- lmer(paste0('GrowthIncrement_2024 ~', trait,  '* garden_MAT_2024 + Pt + (1 | MiniCG_Site/block) + (1 | Genotype)'),
              data = dat)
  
  print(car::Anova(mod))
  
  p1 <- plot_model(mod, vline.color = "black", show.values = T, type = 'std', 
                   title = paste('2024 Growth ~ ', trait, sep = ''))
  plot(p1)

  mods.mat[[n]] <- mod
  names(mods.mat)[n] <- trait
  
  #####################
  # posthoc tests
  # if either the climate or trait x climate are significant, run posthoc tests for each garden
  mod.info <- Anova(mod)
  # if any coefficients are significant, plot by garden
  # need to change indexes if model is changed!
  # 1 is trait effect, 2 is garden climate effect, 4 is trait x climate effect.
  # Skip 3 which is ancestry - we already know this predicts height.
  
  if(mod.info$`Pr(>Chisq)`[1] <= 0.05 | mod.info$`Pr(>Chisq)`[2] <= 0.05 | mod.info$`Pr(>Chisq)`[4] <=0.05){
    
    # list to save plots for each site
    plots.sub <- list()
    
    # dataframe to save outputs from posthoc tests
    df.posthoc <- as.data.frame(matrix(ncol = 3, nrow = length(sites)))
    rownames(df.posthoc) <- sites
    colnames(df.posthoc) <- c('mat', 'slope', 'pval')
    df.posthoc$mat <- sites.mat
    
    for(s in 1:length(sites)){
      
      site <- sites[s]
      dat.sub <- subset(dat, MiniCG_Site_2024 == site)
      # skip if site is missing data
      if(sum(!is.na(dat.sub[,trait])) > 10){
        mod.sub <- lmer(paste0('GrowthIncrement_2024 ~', trait, '+ Pt + (1 | block) + (1 | Genotype)'),
                        data = dat.sub)
        # get pval for trait
        mod.sub.info <- Anova(mod.sub)
        df.posthoc[site, 'pval'] <- mod.sub.info$`Pr(>Chisq)`[1]
        
        # get slopes from fixed effect of trait (effect on growth)
        df.posthoc[site, 'slope'] <- fixef(mod.sub)[2]
        
        # make main title pretty
        maintitle <- paste(site, '\n', trait_names_df[trait, 'trait_names'], '\n', 'p = ', round(df.posthoc$pval[s], 3), sep = '')
        maintitle <- paste(strwrap(maintitle, width = 30), collapse = '\n')
        
        # plot
        p <- ggplot(data = dat.sub, aes_string(y = 'GrowthIncrement_2024', x = trait, color = 'k2_tricho'), na.rm  = TRUE) +
          geom_point(na.rm = TRUE) +
          scale_color_gradient2(high = "darkolivegreen2", mid = "grey20", low = "dodgerblue2", midpoint = 0.5, name = 'P. trichocarpa\nancestry', guide = 'none') +
          ggtitle(maintitle) +
          geom_smooth(color = 'black', lty = ifelse(df.posthoc$pval[s] <= 0.05, 1,2), method = 'lm', formula = y ~ x, na.rm = TRUE) +
          theme(axis.text.x = element_text(angle = 90, vjust = 0.5, hjust=1, size = 12)) +
          theme(axis.title.x = element_blank())
        
        #plot(p)
        plots.sub[[s]] <- p
      }
    }
    # pvals are adjusted for summary plot
    df.posthoc$pvals.adj <- p.adjust(df.posthoc$pval, method = 'BH') 
    
    # plot panel with relationship for each garden
    # remove empty plots
    plots.sub <- plots.sub[lengths(plots.sub)>0]
    # plot in grid
    p1 <- grid.arrange(grobs = plots.sub, ncol = 4)
    plot(p1)
    
    # setup text showing significance
    sig_text <- paste0('Trait ', pval_stars(mod.info[1, 'Pr(>Chisq)']), '\n',
           'Climate ', pval_stars(mod.info[2, 'Pr(>Chisq)']), '\n',
           'Trait x Climate ', pval_stars(mod.info[4, 'Pr(>Chisq)']))
    # wrap title
    maintitle <- trait_names_df[trait, 'trait_names']
    maintitle <- paste(strwrap(maintitle, width = 30), collapse = '\n')
    
    # legend position
    #only plot for first graph
    if(first == TRUE){
      leg_pos <- 'left'
    }else{
      leg_pos <- 'none'
    }
    
    # plot slope of fitness relationship vs climate
    p2 <- ggplot(df.posthoc, aes(x = mat, y = slope), na.rm  = TRUE) +
      geom_smooth(method = 'lm', col = 'black', na.rm = TRUE) +
      geom_hline(yintercept = 0, lty = 2) +
      geom_point(size = 5, stroke = 2, shape = ifelse(df.posthoc$pvals.adj <= 0.05, 16, 1), aes(col = mat), na.rm = TRUE) +
      scale_color_gradient2(high = "red2", mid = "grey", low = "blue2", midpoint = mean(range(sites.mat)), limits = range_mat, name = 'Garden MAT (°C)') +
      ggtitle(maintitle,
              subtitle = sig_text) +
      xlab('Garden MAT') +
      ylab('Effect on Growth Increment') +
      geom_text_repel(label = rownames(df.posthoc), aes(x = mat, y = slope), box.padding = 0.5, min.segment.length = 1, size = 4) +
      theme(plot.title = element_text(size = 15),
            plot.subtitle = element_text(size = 10),
            legend.position = 'none'
      )
    plot(p2)
    
    plots.summ[[n]] <- p2
    names(plots.summ)[n] <- trait
    
    # add legend
    leg_plot <- ggplot(df.posthoc, aes(x = mat, y = slope), na.rm  = TRUE) +
      geom_point(aes(color = mat))+
      scale_color_gradient2(high = "red2", mid = "grey", low = "blue2", midpoint = mean(range(sites.mat)), name = 'Garden MAT (°C)', limits = range_mat) +
      theme(legend.position = 'right')
    
    legend <- cowplot::get_legend(leg_plot)
    
  }
}
```

```
## [1] "Model for: DOY_Stage2_2024"
## Analysis of Deviance Table (Type II Wald chisquare tests)
## 
## Response: GrowthIncrement_2024
##                                   Chisq Df Pr(>Chisq)    
## DOY_Stage2_2024                  0.0018  1     0.9659    
## garden_MAT_2024                  1.5473  1     0.2135    
## Pt                              29.4663  1   5.69e-08 ***
## DOY_Stage2_2024:garden_MAT_2024  0.2268  1     0.6339    
## ---
## Signif. codes:  0 '***' 0.001 '**' 0.01 '*' 0.05 '.' 0.1 ' ' 1
```

```
## [1] "Model for: DOY_Stage3_2024"
## Analysis of Deviance Table (Type II Wald chisquare tests)
## 
## Response: GrowthIncrement_2024
##                                   Chisq Df Pr(>Chisq)    
## DOY_Stage3_2024                  0.5296  1     0.4668    
## garden_MAT_2024                  0.7808  1     0.3769    
## Pt                              35.3612  1  2.739e-09 ***
## DOY_Stage3_2024:garden_MAT_2024  1.1136  1     0.2913    
## ---
## Signif. codes:  0 '***' 0.001 '**' 0.01 '*' 0.05 '.' 0.1 ' ' 1
```

```
## [1] "Model for: DOY_Stage6_2024"
```

```
## Warning: Some predictor variables are on very different scales: consider
## rescaling
```

```
## Analysis of Deviance Table (Type II Wald chisquare tests)
## 
## Response: GrowthIncrement_2024
##                                   Chisq Df Pr(>Chisq)    
## DOY_Stage6_2024                 18.2497  1  1.938e-05 ***
## garden_MAT_2024                  0.3487  1     0.5549    
## Pt                              29.2383  1  6.400e-08 ***
## DOY_Stage6_2024:garden_MAT_2024  0.0528  1     0.8183    
## ---
## Signif. codes:  0 '***' 0.001 '**' 0.01 '*' 0.05 '.' 0.1 ' ' 1
```

```
## boundary (singular) fit: see help('isSingular')
## boundary (singular) fit: see help('isSingular')
## boundary (singular) fit: see help('isSingular')
## boundary (singular) fit: see help('isSingular')
## boundary (singular) fit: see help('isSingular')
## boundary (singular) fit: see help('isSingular')
```

```
## `geom_smooth()` using formula = 'y ~ x'
```

```
## Warning: Removed 1 row containing missing values or values outside the scale range
## (`geom_text_repel()`).
```

```
## Warning: Removed 1 row containing missing values or values outside the scale range
## (`geom_point()`).
```

```
## [1] "Model for: DOY_Stage7_2024"
```

```
## Warning: Some predictor variables are on very different scales: consider
## rescaling
```

```
## boundary (singular) fit: see help('isSingular')
```

```
## Analysis of Deviance Table (Type II Wald chisquare tests)
## 
## Response: GrowthIncrement_2024
##                                  Chisq Df Pr(>Chisq)  
## DOY_Stage7_2024                 1.0105  1     0.3148  
## garden_MAT_2024                 1.3197  1     0.2506  
## Pt                              5.8814  1     0.0153 *
## DOY_Stage7_2024:garden_MAT_2024 0.0249  1     0.8747  
## ---
## Signif. codes:  0 '***' 0.001 '**' 0.01 '*' 0.05 '.' 0.1 ' ' 1
```

```
## boundary (singular) fit: see help('isSingular')
```

```
## [1] "Model for: stage7_presence_2024"
## Analysis of Deviance Table (Type II Wald chisquare tests)
## 
## Response: GrowthIncrement_2024
##                                        Chisq Df Pr(>Chisq)    
## stage7_presence_2024                  1.5294  1     0.2162    
## garden_MAT_2024                       0.9435  1     0.3314    
## Pt                                   28.0951  1  1.155e-07 ***
## stage7_presence_2024:garden_MAT_2024  1.5185  1     0.2178    
## ---
## Signif. codes:  0 '***' 0.001 '**' 0.01 '*' 0.05 '.' 0.1 ' ' 1
```

```
## [1] "Model for: DOY_last_budset_2024"
```

```
## Warning: Some predictor variables are on very different scales: consider
## rescaling
```

```
## Analysis of Deviance Table (Type II Wald chisquare tests)
## 
## Response: GrowthIncrement_2024
##                                        Chisq Df Pr(>Chisq)    
## DOY_last_budset_2024                  0.8315  1   0.361847    
## garden_MAT_2024                       1.4576  1   0.227305    
## Pt                                   26.7330  1  2.336e-07 ***
## DOY_last_budset_2024:garden_MAT_2024 10.1902  1   0.001412 ** 
## ---
## Signif. codes:  0 '***' 0.001 '**' 0.01 '*' 0.05 '.' 0.1 ' ' 1
```

```
## boundary (singular) fit: see help('isSingular')
## boundary (singular) fit: see help('isSingular')
## boundary (singular) fit: see help('isSingular')
## boundary (singular) fit: see help('isSingular')
```

```
## `geom_smooth()` using formula = 'y ~ x'
```

```
## Warning: Removed 1 row containing missing values or values outside the scale range
## (`geom_text_repel()`).
## Removed 1 row containing missing values or values outside the scale range
## (`geom_point()`).
```

```
## [1] "Model for: growing_season_days_2024"
```

```
## Warning: Some predictor variables are on very different scales: consider
## rescaling
```

```
## Analysis of Deviance Table (Type II Wald chisquare tests)
## 
## Response: GrowthIncrement_2024
##                                            Chisq Df Pr(>Chisq)    
## growing_season_days_2024                  1.3821  1   0.239738    
## garden_MAT_2024                           1.2762  1   0.258612    
## Pt                                       24.5900  1  7.092e-07 ***
## growing_season_days_2024:garden_MAT_2024  8.1319  1   0.004349 ** 
## ---
## Signif. codes:  0 '***' 0.001 '**' 0.01 '*' 0.05 '.' 0.1 ' ' 1
```

```
## boundary (singular) fit: see help('isSingular')
## boundary (singular) fit: see help('isSingular')
## boundary (singular) fit: see help('isSingular')
## boundary (singular) fit: see help('isSingular')
```

```
## `geom_smooth()` using formula = 'y ~ x'
```

```
## Warning: Removed 1 row containing missing values or values outside the scale range
## (`geom_text_repel()`).
## Removed 1 row containing missing values or values outside the scale range
## (`geom_point()`).
```

```
# remove empty plots
plots.summ <- plots.summ[lengths(plots.summ)>0]

# add legend to beginning of plot list
plots.summ[[length(plots.summ)+1]] <- legend

p3 <- grid.arrange(grobs = plots.summ, ncol = 4)
```

```
## `geom_smooth()` using formula = 'y ~ x'
```

```
## Warning: Removed 1 row containing missing values or values outside the scale range
## (`geom_text_repel()`).
```

```
## `geom_smooth()` using formula = 'y ~ x'
```

```
## Warning: Removed 1 row containing missing values or values outside the scale range
## (`geom_text_repel()`).
```

```
## `geom_smooth()` using formula = 'y ~ x'
```

```
## Warning: Removed 1 row containing missing values or values outside the scale range
## (`geom_text_repel()`).
```

```
plot(p3)
#ggsave(file = 'results/fitness/fitnessEffects_summaryPlot_traitxMAT_2024.png', p3, height = 4, width = 12)
#ggsave(file = 'results/fitness/fitnessEffects_summaryPlot_traitxMAT_2024.pdf', p3, height = 4, width = 12)

# rename plots list for later use
plots.trait.mat.24 <- plots.summ
```

##### 2.2.2 Garden MAP

```
###################


# with garden MAP

# reorder gardens by garden MAP
tmp <- aggregate(dat$garden_MAP_2024, by = list(dat$MiniCG_Site_2024), mean)
ord <- tmp$Group.1[order(tmp$x, decreasing = T)]
dat$MiniCG_Site_2024 <- factor(dat$MiniCG_Site_2024, levels = ord)

sites.map <- dat$garden_MAP_2024[match(sites, dat$MiniCG_Site)]


# list to save results
mods.map <- list()
# list to save summary plot for each trait with significant effects on fitness
plots.summ <- list()

# loop through traits
for(n in 1:length(traits)){
  
  trait <- traits[n]
  
  print(paste('Model for:', trait))
  
  
  mod <- lmer(paste0('GrowthIncrement_2024 ~', trait,  '* garden_MAP_2024 + Pt + (1 | MiniCG_Site/block) + (1 | Genotype)'),
              data = dat)
  
  print(car::Anova(mod))
  
  p1 <- plot_model(mod, vline.color = "black", show.values = T, type = 'std', 
                   title = paste('2024 Growth ~ ', trait, sep = ''))
  plot(p1)

  mods.map[[n]] <- mod
  names(mods.map)[n] <- trait
  
  #####################
  # posthoc tests
  # if either the climate or trait x climate are significant, run posthoc tests for each garden
  mod.info <- Anova(mod)
  # if any coefficients are significant, plot by garden
  # need to change indexes if model is changed!
  # 1 is trait effect, 2 is garden climate effect, 4 is trait x climate effect.
  # Skip 3 which is ancestry - we already know this predicts height.
  
  if(mod.info$`Pr(>Chisq)`[1] <= 0.05 | mod.info$`Pr(>Chisq)`[2] <= 0.05 | mod.info$`Pr(>Chisq)`[4] <=0.05){
    
    # list to save plots for each site
    plots.sub <- list()
    
    # dataframe to save outputs from posthoc tests
    df.posthoc <- as.data.frame(matrix(ncol = 3, nrow = length(sites)))
    rownames(df.posthoc) <- sites
    colnames(df.posthoc) <- c('map', 'slope', 'pval')
    df.posthoc$map <- sites.map
    
    # make main title pretty
    maintitle <- paste(site, '\n', trait_names_df[trait, 'trait_names'], '\n', 'p = ', round(df.posthoc$pval[s], 3), sep = '')
    maintitle <- paste(strwrap(maintitle, width = 30), collapse = '\n')
    
    for(s in 1:length(sites)){
      
      site <- sites[s]
      dat.sub <- subset(dat, MiniCG_Site_2024 == site)
      # skip if site is missing data
      if(sum(!is.na(dat.sub[,trait])) > 10){
        mod.sub <- lmer(paste0('GrowthIncrement_2024 ~', trait, '+ Pt + (1 | block) + (1 | Genotype)'),
                        data = dat.sub)
        # get pval for trait
        mod.sub.info <- Anova(mod.sub)
        df.posthoc[site, 'pval'] <- mod.sub.info$`Pr(>Chisq)`[1]
        
        # get slopes from fixed effect of trait (effect on growth)
        df.posthoc[site, 'slope'] <- fixef(mod.sub)[2]
        
        # plot
        p <- ggplot(data = dat.sub, aes_string(y = 'GrowthIncrement_2024', x = trait, color = 'k2_tricho'), na.rm  = TRUE) +
          geom_point(na.rm = TRUE) +
          scale_color_gradient2(high = "darkolivegreen2", mid = "grey20", low = "dodgerblue2", midpoint = 0.5, name = 'P. trichocarpa\nancestry', guide = 'none') +
         ggtitle(maintitle) +
          geom_smooth(color = 'black', lty = ifelse(df.posthoc$pval[s] <= 0.05, 1,2), method = 'lm', formula = y ~ x, na.rm = TRUE) +
          theme(axis.text.x = element_text(angle = 90, vjust = 0.5, hjust=1, size = 12)) +
          theme(axis.title.x = element_blank())
        
        #plot(p)
        plots.sub[[s]] <- p
      }
    }

    # pvals are adjusted for summary plot
    df.posthoc$pvals.adj <- p.adjust(df.posthoc$pval, method = 'BH') 
    
    # plot panel with relationship for each garden
    # remove empty plots
    plots.sub <- plots.sub[lengths(plots.sub)>0]
    # plot in grid
    p1 <- grid.arrange(grobs = plots.sub, ncol = 4)
    plot(p1)
    
    # setup text showing significance
    sig_text <- paste0('Trait ', pval_stars(mod.info[1, 'Pr(>Chisq)']), '\n',
                       'Climate ', pval_stars(mod.info[2, 'Pr(>Chisq)']), '\n',
                       'Trait x Climate ', pval_stars(mod.info[4, 'Pr(>Chisq)']))
    
    # wrap title
    maintitle <- trait_names_df[trait, 'trait_names']
    maintitle <- paste(strwrap(maintitle, width = 30), collapse = '\n')
    
    # plot slope of fitness relationship vs climate
    p2 <- ggplot(df.posthoc, aes(x = map, y = slope), na.rm  = TRUE) +
      geom_smooth(method = 'lm', col = 'black', na.rm = TRUE) +
      geom_hline(yintercept = 0, lty = 2) +
      geom_point(size = 5, stroke = 2, shape = ifelse(df.posthoc$pvals.adj <= 0.05, 16, 1), aes(col = map), na.rm = TRUE) +
      scale_color_gradient2(high = "steelblue2",  mid = 'grey', low = "sienna3", midpoint = mean(range(sites.map)), limits = range_map, name = 'Garden MAP (mm)') +
      ggtitle(maintitle,
              subtitle = sig_text)+
      xlab('Garden MAP') +
      ylab('Effect on Growth Increment') +
      geom_text_repel(label = rownames(df.posthoc), aes(x = map, y = slope), box.padding = 0.5, min.segment.length = 1, size = 4) +
      theme(plot.title = element_text(size = 15),
            plot.subtitle = element_text(size = 10),
            legend.position = 'none')
    plot(p2)
    
    # add legend
    leg_plot <- ggplot(df.posthoc, aes(x = map, y = slope), na.rm  = TRUE) +
      geom_point(aes(color = map), na.rm = TRUE)+
      scale_color_gradient2(high = "steelblue2",  mid = 'grey', low = "sienna3", midpoint = mean(range(sites.map)), limits = range_map, name = 'Garden MAP (mm)') +
      theme(legend.position = 'right')
    
    legend <- cowplot::get_legend(leg_plot)
    
    
    plots.summ[[n]] <- p2
    names(plots.summ)[n] <- trait
    
  }
}
```

```
## [1] "Model for: DOY_Stage2_2024"
```

```
## Warning: Some predictor variables are on very different scales: consider
## rescaling
```

```
## Analysis of Deviance Table (Type II Wald chisquare tests)
## 
## Response: GrowthIncrement_2024
##                                   Chisq Df Pr(>Chisq)    
## DOY_Stage2_2024                  0.3603  1     0.5483    
## garden_MAP_2024                  0.0179  1     0.8937    
## Pt                              29.9274  1  4.485e-08 ***
## DOY_Stage2_2024:garden_MAP_2024  0.1394  1     0.7089    
## ---
## Signif. codes:  0 '***' 0.001 '**' 0.01 '*' 0.05 '.' 0.1 ' ' 1
```

```
## [1] "Model for: DOY_Stage3_2024"
```

```
## Warning: Some predictor variables are on very different scales: consider
## rescaling
```

```
## Analysis of Deviance Table (Type II Wald chisquare tests)
## 
## Response: GrowthIncrement_2024
##                                   Chisq Df Pr(>Chisq)    
## DOY_Stage3_2024                  1.4493  1     0.2286    
## garden_MAP_2024                  0.1204  1     0.7286    
## Pt                              35.6052  1  2.416e-09 ***
## DOY_Stage3_2024:garden_MAP_2024  0.3088  1     0.5784    
## ---
## Signif. codes:  0 '***' 0.001 '**' 0.01 '*' 0.05 '.' 0.1 ' ' 1
```

```
## [1] "Model for: DOY_Stage6_2024"
```

```
## Warning: Some predictor variables are on very different scales: consider
## rescaling
```

```
## Analysis of Deviance Table (Type II Wald chisquare tests)
## 
## Response: GrowthIncrement_2024
##                                   Chisq Df Pr(>Chisq)    
## DOY_Stage6_2024                 18.8342  1  1.426e-05 ***
## garden_MAP_2024                  0.0005  1     0.9818    
## Pt                              29.3432  1  6.063e-08 ***
## DOY_Stage6_2024:garden_MAP_2024  0.0110  1     0.9164    
## ---
## Signif. codes:  0 '***' 0.001 '**' 0.01 '*' 0.05 '.' 0.1 ' ' 1
```

```
## boundary (singular) fit: see help('isSingular')
## boundary (singular) fit: see help('isSingular')
## boundary (singular) fit: see help('isSingular')
## boundary (singular) fit: see help('isSingular')
## boundary (singular) fit: see help('isSingular')
## boundary (singular) fit: see help('isSingular')
```

```
## `geom_smooth()` using formula = 'y ~ x'
```

```
## Warning: Removed 1 row containing missing values or values outside the scale range
## (`geom_text_repel()`).
```

```
## [1] "Model for: DOY_Stage7_2024"
```

```
## Warning: Some predictor variables are on very different scales: consider
## rescaling
```

```
## boundary (singular) fit: see help('isSingular')
```

```
## Analysis of Deviance Table (Type II Wald chisquare tests)
## 
## Response: GrowthIncrement_2024
##                                  Chisq Df Pr(>Chisq)  
## DOY_Stage7_2024                 0.6215  1    0.43051  
## garden_MAP_2024                 0.2562  1    0.61272  
## Pt                              5.4877  1    0.01915 *
## DOY_Stage7_2024:garden_MAP_2024 0.3152  1    0.57453  
## ---
## Signif. codes:  0 '***' 0.001 '**' 0.01 '*' 0.05 '.' 0.1 ' ' 1
```

```
## boundary (singular) fit: see help('isSingular')
```

```
## [1] "Model for: stage7_presence_2024"
```

```
## Warning: Some predictor variables are on very different scales: consider
## rescaling
```

```
## Analysis of Deviance Table (Type II Wald chisquare tests)
## 
## Response: GrowthIncrement_2024
##                                        Chisq Df Pr(>Chisq)    
## stage7_presence_2024                  2.5001  1     0.1138    
## garden_MAP_2024                       0.0577  1     0.8102    
## Pt                                   25.7076  1  3.973e-07 ***
## stage7_presence_2024:garden_MAP_2024  2.6526  1     0.1034    
## ---
## Signif. codes:  0 '***' 0.001 '**' 0.01 '*' 0.05 '.' 0.1 ' ' 1
```

```
## [1] "Model for: DOY_last_budset_2024"
```

```
## Warning: Some predictor variables are on very different scales: consider
## rescaling
```

```
## Analysis of Deviance Table (Type II Wald chisquare tests)
## 
## Response: GrowthIncrement_2024
##                                        Chisq Df Pr(>Chisq)    
## DOY_last_budset_2024                  1.2430  1     0.2649    
## garden_MAP_2024                       0.0008  1     0.9769    
## Pt                                   30.6522  1  3.087e-08 ***
## DOY_last_budset_2024:garden_MAP_2024  1.1707  1     0.2793    
## ---
## Signif. codes:  0 '***' 0.001 '**' 0.01 '*' 0.05 '.' 0.1 ' ' 1
```

```
## [1] "Model for: growing_season_days_2024"
```

```
## Warning: Some predictor variables are on very different scales: consider
## rescaling
```

```
## Analysis of Deviance Table (Type II Wald chisquare tests)
## 
## Response: GrowthIncrement_2024
##                                            Chisq Df Pr(>Chisq)    
## growing_season_days_2024                  1.7461  1     0.1864    
## garden_MAP_2024                           0.0017  1     0.9670    
## Pt                                       28.2078  1   1.09e-07 ***
## growing_season_days_2024:garden_MAP_2024  1.4574  1     0.2273    
## ---
## Signif. codes:  0 '***' 0.001 '**' 0.01 '*' 0.05 '.' 0.1 ' ' 1
```

```
# significant interaction for DOY stage6, DOY last budset, gsw, gbw, PhiPS2, Fs, LMA, leaf mass, leaf area, lower pore length, upper pore length
plots.summ <- plots.summ[lengths(plots.summ)>0]

# add legend to beginning of plot list

plots.summ[[length(plots.summ)+1]] <- legend

# there is only one significant trait

p3 <- grid.arrange(grobs = plots.summ, ncol = 2)
```

```
## `geom_smooth()` using formula = 'y ~ x'
```

```
## Warning: Removed 1 row containing missing values or values outside the scale range
## (`geom_text_repel()`).
```

```
plot(p3)

#ggsave(file = 'results/fitness/fitnessEffects_summaryPlot_traitxMAP_2024.png', p3, height = 4, width = 6)
#ggsave(file = 'results/fitness/fitnessEffects_summaryPlot_traitxMAP_2024.pdf', p3, height = 4, width = 6)

# rename plots list for later use
plots.trait.map.24 <- plots.summ
```

##### 2.2.3 Garden TD

```
###################

# with garden TD

# reorder gardens by garden TD
tmp <- aggregate(dat$garden_TD_2024, by = list(dat$MiniCG_Site_2024), mean)
ord <- tmp$Group.1[order(tmp$x, decreasing = T)]
dat$MiniCG_Site_2024 <- factor(dat$MiniCG_Site_2024, levels = ord)

sites.td <- dat$garden_TD_2024[match(sites, dat$MiniCG_Site)]


# list to save results
mods.td <- list()
# list to save summary plot for each trait with significant effects on fitness
plots.summ <- list()

# loop through traits
for(n in 1:length(traits)){
  
  trait <- traits[n]
  
  print(paste('Model for:', trait))
  
  mod <- lmer(paste0('GrowthIncrement_2024 ~', trait,  '* garden_TD_2024 + Pt + (1 | MiniCG_Site/block) + (1 | Genotype)'),
              data = dat)
  
  print(car::Anova(mod))
  
  p1 <- plot_model(mod, vline.color = "black", show.values = T, type = 'std', 
                   title = paste('2024 Growth ~ ', trait, sep = ''))
  plot(p1)

  mods.td[[n]] <- mod
  names(mods.td)[n] <- trait
  
  #####################
  # posthoc tests
  # if either the climate or trait x climate are significant, run posthoc tests for each garden
  mod.info <- Anova(mod)
  # if any coefficients are significant, plot by garden
  # need to change indexes if model is changed!
  # 1 is trait effect, 2 is garden climate effect, 4 is trait x climate effect.
  # Skip 3 which is ancestry - we already know this predicts height.
  
  if(mod.info$`Pr(>Chisq)`[1] <= 0.05 | mod.info$`Pr(>Chisq)`[2] <= 0.05 | mod.info$`Pr(>Chisq)`[4] <=0.05){
    
    # list to save plots for each site
    plots.sub <- list()
    
    # dataframe to save outputs from posthoc tests
    df.posthoc <- as.data.frame(matrix(ncol = 3, nrow = length(sites)))
    rownames(df.posthoc) <- sites
    colnames(df.posthoc) <- c('td', 'slope', 'pval')
    df.posthoc$td <- sites.td
    
    # make main title pretty
    maintitle <- paste(site, '\n', trait_names_df[trait, 'trait_names'], '\n', 'p = ', round(df.posthoc$pval[s], 3), sep = '')
    maintitle <- paste(strwrap(maintitle, width = 30), collapse = '\n')
    
    for(s in 1:length(sites)){
      
      site <- sites[s]
      dat.sub <- subset(dat, MiniCG_Site_2024 == site)
      # skip if site is missing data
      if(sum(!is.na(dat.sub[,trait])) > 10){
        mod.sub <- lmer(paste0('GrowthIncrement_2024 ~', trait, '+ Pt + (1 | block) + (1 | Genotype)'),
                        data = dat.sub)
        # get pval for trait
        mod.sub.info <- Anova(mod.sub)
        df.posthoc[site, 'pval'] <- mod.sub.info$`Pr(>Chisq)`[1]
        
        # get slopes from fixed effect of trait (effect on growth)
        df.posthoc[site, 'slope'] <- fixef(mod.sub)[2]
        
        # plot
        p <- ggplot(data = dat.sub, aes_string(y = 'GrowthIncrement_2024', x = trait, color = 'k2_tricho'), na.rm  = TRUE) +
          geom_point(na.rm = TRUE) +
          scale_color_gradient2(high = "darkolivegreen2", mid = "grey20", low = "dodgerblue2", midpoint = 0.5, name = 'P. trichocarpa\nancestry', guide = 'none') +
         ggtitle(maintitle) +
          geom_smooth(color = 'black', lty = ifelse(df.posthoc$pval[s] <= 0.05, 1,2), method = 'lm', formula = y ~ x, na.rm = TRUE)
          theme(axis.text.x = element_text(angle = 90, vjust = 0.5, hjust=1, size = 12)) +
          theme(axis.title.x = element_blank())
        
        #plot(p)
        plots.sub[[s]] <- p
      }
    }

    # pvals are adjusted for summary plot
    df.posthoc$pvals.adj <- p.adjust(df.posthoc$pval, method = 'BH') 
    
    # plot panel with relationship for each garden
    # remove empty plots
    plots.sub <- plots.sub[lengths(plots.sub)>0]
    # plot in grid
    p1 <- grid.arrange(grobs = plots.sub, ncol = 4)
    plot(p1)
    
    # setup text showing significance
    sig_text <- paste0('Trait ', pval_stars(mod.info[1, 'Pr(>Chisq)']), '\n',
                       'Climate ', pval_stars(mod.info[2, 'Pr(>Chisq)']), '\n',
                       'Trait x Climate ', pval_stars(mod.info[4, 'Pr(>Chisq)']))
    
    # wrap title
    maintitle <- trait_names_df[trait, 'trait_names']
    maintitle <- paste(strwrap(maintitle, width = 30), collapse = '\n')
    
    
    # plot slope of fitness relationship vs climate
    p2 <- ggplot(df.posthoc, aes(x = td, y = slope), na.rm  = TRUE) +
      geom_point(size = 5, stroke = 2, shape = ifelse(df.posthoc$pvals.adj <= 0.05, 16, 1), aes(color = td), na.rm = TRUE) +
      geom_smooth(method = 'lm', col = 'black', na.rm = TRUE) +
      geom_hline(yintercept = 0, lty = 2) +
      scale_color_gradient2(high = "orchid4",  mid = 'grey', low = "palegreen4", midpoint = mean(range(sites.td)), limits = range_td, name = 'Garden TD (°C)') +
      ggtitle(maintitle,
              subtitle = sig_text)+
      xlab('Garden TD') +
      ylab('Effect on Growth Increment') +
      geom_text_repel(label = rownames(df.posthoc), aes(x = td, y = slope), box.padding = 0.5, min.segment.length = 1, size = 4) +
      theme(plot.title = element_text(size = 15),
            plot.subtitle = element_text(size = 10),
            legend.position = 'none')
    plot(p2)
    
    plots.summ[[n]] <- p2
    names(plots.summ)[n] <- trait
    
    
     # add legend
    leg_plot <- ggplot(df.posthoc, aes(x = td, y = slope), na.rm  = TRUE) +
      geom_point(aes(color = td))+
      scale_color_gradient2(high = "orchid4",  mid = 'grey', low = "palegreen4", midpoint = mean(range(sites.td)), limits = range_td, name = 'Garden TD (°C)')  +
      theme(legend.position = 'right')
    
    legend <- cowplot::get_legend(leg_plot)
  }
}
```

```
## [1] "Model for: DOY_Stage2_2024"
```

```
## Warning: Some predictor variables are on very different scales: consider
## rescaling
```

```
## Analysis of Deviance Table (Type II Wald chisquare tests)
## 
## Response: GrowthIncrement_2024
##                                  Chisq Df Pr(>Chisq)    
## DOY_Stage2_2024                 0.4058  1     0.5241    
## garden_TD_2024                  0.0154  1     0.9012    
## Pt                             30.5684  1  3.223e-08 ***
## DOY_Stage2_2024:garden_TD_2024  0.1648  1     0.6848    
## ---
## Signif. codes:  0 '***' 0.001 '**' 0.01 '*' 0.05 '.' 0.1 ' ' 1
```

```
## [1] "Model for: DOY_Stage3_2024"
```

```
## Warning: Some predictor variables are on very different scales: consider
## rescaling
```

```
## Analysis of Deviance Table (Type II Wald chisquare tests)
## 
## Response: GrowthIncrement_2024
##                                  Chisq Df Pr(>Chisq)    
## DOY_Stage3_2024                 1.3898  1     0.2384    
## garden_TD_2024                  0.0467  1     0.8289    
## Pt                             35.4874  1  2.567e-09 ***
## DOY_Stage3_2024:garden_TD_2024  0.0055  1     0.9408    
## ---
## Signif. codes:  0 '***' 0.001 '**' 0.01 '*' 0.05 '.' 0.1 ' ' 1
```

```
## [1] "Model for: DOY_Stage6_2024"
```

```
## Warning: Some predictor variables are on very different scales: consider
## rescaling
```

```
## Analysis of Deviance Table (Type II Wald chisquare tests)
## 
## Response: GrowthIncrement_2024
##                                  Chisq Df Pr(>Chisq)    
## DOY_Stage6_2024                17.0479  1  3.645e-05 ***
## garden_TD_2024                  0.5233  1     0.4694    
## Pt                             29.7001  1  5.043e-08 ***
## DOY_Stage6_2024:garden_TD_2024  2.6444  1     0.1039    
## ---
## Signif. codes:  0 '***' 0.001 '**' 0.01 '*' 0.05 '.' 0.1 ' ' 1
```

```
## boundary (singular) fit: see help('isSingular')
## boundary (singular) fit: see help('isSingular')
## boundary (singular) fit: see help('isSingular')
## boundary (singular) fit: see help('isSingular')
## boundary (singular) fit: see help('isSingular')
## boundary (singular) fit: see help('isSingular')
```

```
## `geom_smooth()` using formula = 'y ~ x'
```

```
## Warning: Removed 1 row containing missing values or values outside the scale range
## (`geom_text_repel()`).
```

```
## Warning: Removed 1 row containing missing values or values outside the scale range
## (`geom_point()`).
```

```
## [1] "Model for: DOY_Stage7_2024"
```

```
## Warning: Some predictor variables are on very different scales: consider
## rescaling
```

```
## Analysis of Deviance Table (Type II Wald chisquare tests)
## 
## Response: GrowthIncrement_2024
##                                 Chisq Df Pr(>Chisq)  
## DOY_Stage7_2024                0.7884  1    0.37458  
## garden_TD_2024                 0.0308  1    0.86058  
## Pt                             6.0887  1    0.01361 *
## DOY_Stage7_2024:garden_TD_2024 0.4983  1    0.48026  
## ---
## Signif. codes:  0 '***' 0.001 '**' 0.01 '*' 0.05 '.' 0.1 ' ' 1
```

```
## [1] "Model for: stage7_presence_2024"
## Analysis of Deviance Table (Type II Wald chisquare tests)
## 
## Response: GrowthIncrement_2024
##                                       Chisq Df Pr(>Chisq)    
## stage7_presence_2024                 2.3418  1     0.1259    
## garden_TD_2024                       0.0033  1     0.9539    
## Pt                                  27.0035  1  2.031e-07 ***
## stage7_presence_2024:garden_TD_2024  0.7688  1     0.3806    
## ---
## Signif. codes:  0 '***' 0.001 '**' 0.01 '*' 0.05 '.' 0.1 ' ' 1
```

```
## [1] "Model for: DOY_last_budset_2024"
```

```
## Warning: Some predictor variables are on very different scales: consider
## rescaling
```

```
## Analysis of Deviance Table (Type II Wald chisquare tests)
## 
## Response: GrowthIncrement_2024
##                                       Chisq Df Pr(>Chisq)    
## DOY_last_budset_2024                 0.9750  1    0.32344    
## garden_TD_2024                       0.0001  1    0.99146    
## Pt                                  27.4689  1  1.596e-07 ***
## DOY_last_budset_2024:garden_TD_2024  4.4175  1    0.03557 *  
## ---
## Signif. codes:  0 '***' 0.001 '**' 0.01 '*' 0.05 '.' 0.1 ' ' 1
```

```
## boundary (singular) fit: see help('isSingular')
## boundary (singular) fit: see help('isSingular')
## boundary (singular) fit: see help('isSingular')
## boundary (singular) fit: see help('isSingular')
```

```
## `geom_smooth()` using formula = 'y ~ x'
```

```
## Warning: Removed 1 row containing missing values or values outside the scale range
## (`geom_text_repel()`).
## Removed 1 row containing missing values or values outside the scale range
## (`geom_point()`).
```

```
## [1] "Model for: growing_season_days_2024"
```

```
## Warning: Some predictor variables are on very different scales: consider
## rescaling
```

```
## Analysis of Deviance Table (Type II Wald chisquare tests)
## 
## Response: GrowthIncrement_2024
##                                           Chisq Df Pr(>Chisq)    
## growing_season_days_2024                 1.5712  1     0.2100    
## garden_TD_2024                           0.0146  1     0.9037    
## Pt                                      24.5168  1  7.367e-07 ***
## growing_season_days_2024:garden_TD_2024  2.0862  1     0.1486    
## ---
## Signif. codes:  0 '***' 0.001 '**' 0.01 '*' 0.05 '.' 0.1 ' ' 1
```

```
plots.summ <- plots.summ[lengths(plots.summ)>0]
# add legend to beginning of plot list

plots.summ[[length(plots.summ)+1]] <- legend

# there are two significant traits
p3 <- grid.arrange(grobs = plots.summ, ncol = 3)
```

```
## `geom_smooth()` using formula = 'y ~ x'
```

```
## Warning: Removed 1 row containing missing values or values outside the scale range
## (`geom_text_repel()`).
```

```
## `geom_smooth()` using formula = 'y ~ x'
```

```
## Warning: Removed 1 row containing missing values or values outside the scale range
## (`geom_text_repel()`).
```

```
plot(p3)
#ggsave(file = 'results/fitness/fitnessEffects_summaryPlot_traitxTD_2024.png', p3, height = 4, width = 9)
#ggsave(file = 'results/fitness/fitnessEffects_summaryPlot_traitxTD_2024.pdf', p3, height = 4, width = 9)

# rename plots list for later use
plots.trait.td.24 <- plots.summ
```

#### 2.3 Combine plots for both years

```
# MAT
plots.trait.mat <- c(plots.trait.mat.23, plots.trait.mat.24)
plots.trait.mat <- plots.trait.mat[lengths(plots.trait.mat)>0]

# Because of ggrepel for garden labels, it will not plot if the window is too small
# To fix this when plotting within Rstudio, create a new window and expand to a larger size.
#dev.new()
p.mat <- grid.arrange(grobs = plots.trait.mat, ncol = 4)
```

```
## `geom_smooth()` using formula = 'y ~ x'
## `geom_smooth()` using formula = 'y ~ x'
```

```
## Warning: Removed 1 row containing missing values or values outside the scale range
## (`geom_text_repel()`).
```

```
## `geom_smooth()` using formula = 'y ~ x'
```

```
## Warning: Removed 3 rows containing missing values or values outside the scale range
## (`geom_text_repel()`).
```

```
## `geom_smooth()` using formula = 'y ~ x'
```

```
## Warning: Removed 3 rows containing missing values or values outside the scale range
## (`geom_text_repel()`).
```

```
## `geom_smooth()` using formula = 'y ~ x'
```

```
## Warning: Removed 3 rows containing missing values or values outside the scale range
## (`geom_text_repel()`).
```

```
## `geom_smooth()` using formula = 'y ~ x'
```

```
## Warning: Removed 3 rows containing missing values or values outside the scale range
## (`geom_text_repel()`).
```

```
## `geom_smooth()` using formula = 'y ~ x'
```

```
## Warning: Removed 3 rows containing missing values or values outside the scale range
## (`geom_text_repel()`).
```

```
## `geom_smooth()` using formula = 'y ~ x'
```

```
## Warning: Removed 3 rows containing missing values or values outside the scale range
## (`geom_text_repel()`).
```

```
## `geom_smooth()` using formula = 'y ~ x'
```

```
## Warning: Removed 2 rows containing missing values or values outside the scale range
## (`geom_text_repel()`).
```

```
## `geom_smooth()` using formula = 'y ~ x'
```

```
## Warning: Removed 1 row containing missing values or values outside the scale range
## (`geom_text_repel()`).
```

```
## `geom_smooth()` using formula = 'y ~ x'
```

```
## Warning: Removed 1 row containing missing values or values outside the scale range
## (`geom_text_repel()`).
```

```
## `geom_smooth()` using formula = 'y ~ x'
```

```
## Warning: Removed 1 row containing missing values or values outside the scale range
## (`geom_text_repel()`).
```

```
## `geom_smooth()` using formula = 'y ~ x'
```

```
## Warning: Removed 1 row containing missing values or values outside the scale range
## (`geom_text_repel()`).
```

```
## `geom_smooth()` using formula = 'y ~ x'
```

```
## Warning: Removed 1 row containing missing values or values outside the scale range
## (`geom_text_repel()`).
```

```
## `geom_smooth()` using formula = 'y ~ x'
```

```
## Warning: Removed 1 row containing missing values or values outside the scale range
## (`geom_text_repel()`).
```

```
## `geom_smooth()` using formula = 'y ~ x'
```

```
## Warning: Removed 1 row containing missing values or values outside the scale range
## (`geom_text_repel()`).
```

```
## `geom_smooth()` using formula = 'y ~ x'
```

```
## Warning: Removed 1 row containing missing values or values outside the scale range
## (`geom_text_repel()`).
```

```
plot(p.mat)

#ggsave(file = 'results/fitness/fitnessEffects_summaryPlot_traitxMAT.png', p.mat, height = 16, width = 14)
#ggsave(file = 'results/fitness/fitnessEffects_summaryPlot_traitxMAT.pdf', p.mat, height = 16, width = 14)

# MAP

plots.trait.map <- c(plots.trait.map.23, plots.trait.map.24)
plots.trait.map <- plots.trait.map[lengths(plots.trait.map)>0]

#dev.new()
p.map <- grid.arrange(grobs = plots.trait.map, ncol = 4)
```

```
## `geom_smooth()` using formula = 'y ~ x'
## `geom_smooth()` using formula = 'y ~ x'
```

```
## Warning: Removed 1 row containing missing values or values outside the scale range
## (`geom_text_repel()`).
```

```
## `geom_smooth()` using formula = 'y ~ x'
## `geom_smooth()` using formula = 'y ~ x'
```

```
## Warning: Removed 3 rows containing missing values or values outside the scale range
## (`geom_text_repel()`).
```

```
## `geom_smooth()` using formula = 'y ~ x'
```

```
## Warning: Removed 3 rows containing missing values or values outside the scale range
## (`geom_text_repel()`).
```

```
## `geom_smooth()` using formula = 'y ~ x'
```

```
## Warning: Removed 3 rows containing missing values or values outside the scale range
## (`geom_text_repel()`).
```

```
## `geom_smooth()` using formula = 'y ~ x'
```

```
## Warning: Removed 3 rows containing missing values or values outside the scale range
## (`geom_text_repel()`).
```

```
## `geom_smooth()` using formula = 'y ~ x'
```

```
## Warning: Removed 2 rows containing missing values or values outside the scale range
## (`geom_text_repel()`).
```

```
## `geom_smooth()` using formula = 'y ~ x'
```

```
## Warning: Removed 1 row containing missing values or values outside the scale range
## (`geom_text_repel()`).
```

```
## `geom_smooth()` using formula = 'y ~ x'
```

```
## Warning: Removed 1 row containing missing values or values outside the scale range
## (`geom_text_repel()`).
```

```
## `geom_smooth()` using formula = 'y ~ x'
```

```
## Warning: Removed 1 row containing missing values or values outside the scale range
## (`geom_text_repel()`).
```

```
## `geom_smooth()` using formula = 'y ~ x'
```

```
## Warning: Removed 1 row containing missing values or values outside the scale range
## (`geom_text_repel()`).
```

```
## `geom_smooth()` using formula = 'y ~ x'
```

```
## Warning: Removed 1 row containing missing values or values outside the scale range
## (`geom_text_repel()`).
```

```
## `geom_smooth()` using formula = 'y ~ x'
```

```
## Warning: Removed 1 row containing missing values or values outside the scale range
## (`geom_text_repel()`).
```

```
## `geom_smooth()` using formula = 'y ~ x'
```

```
## Warning: Removed 1 row containing missing values or values outside the scale range
## (`geom_text_repel()`).
```

```
## Warning: ggrepel: 5 unlabeled data points (too many overlaps). Consider
## increasing max.overlaps
```

```
plot(p.map)
```

```
## Warning: ggrepel: 5 unlabeled data points (too many overlaps). Consider
## increasing max.overlaps
```

```
#ggsave(file = 'results/fitness/fitnessEffects_summaryPlot_traitxMAP.png', p.map, height = 14, width = 14)
#ggsave(file = 'results/fitness/fitnessEffects_summaryPlot_traitxMAP.pdf', p.map, height = 14, width = 14)

# TD

plots.trait.td <- c(plots.trait.td.23, plots.trait.td.24)
plots.trait.td <- plots.trait.td[lengths(plots.trait.td)>0]

#dev.new()
p.td <- grid.arrange(grobs = plots.trait.td, ncol = 4)
```

```
## `geom_smooth()` using formula = 'y ~ x'
## `geom_smooth()` using formula = 'y ~ x'
```

```
## Warning: Removed 1 row containing missing values or values outside the scale range
## (`geom_text_repel()`).
```

```
## `geom_smooth()` using formula = 'y ~ x'
```

```
## Warning: Removed 3 rows containing missing values or values outside the scale range
## (`geom_text_repel()`).
```

```
## `geom_smooth()` using formula = 'y ~ x'
```

```
## Warning: Removed 3 rows containing missing values or values outside the scale range
## (`geom_text_repel()`).
```

```
## `geom_smooth()` using formula = 'y ~ x'
```

```
## Warning: Removed 3 rows containing missing values or values outside the scale range
## (`geom_text_repel()`).
```

```
## `geom_smooth()` using formula = 'y ~ x'
```

```
## Warning: Removed 2 rows containing missing values or values outside the scale range
## (`geom_text_repel()`).
```

```
## `geom_smooth()` using formula = 'y ~ x'
```

```
## Warning: Removed 1 row containing missing values or values outside the scale range
## (`geom_text_repel()`).
```

```
## `geom_smooth()` using formula = 'y ~ x'
```

```
## Warning: Removed 1 row containing missing values or values outside the scale range
## (`geom_text_repel()`).
```

```
## `geom_smooth()` using formula = 'y ~ x'
```

```
## Warning: Removed 1 row containing missing values or values outside the scale range
## (`geom_text_repel()`).
```

```
## `geom_smooth()` using formula = 'y ~ x'
```

```
## Warning: Removed 1 row containing missing values or values outside the scale range
## (`geom_text_repel()`).
```

```
## `geom_smooth()` using formula = 'y ~ x'
```

```
## Warning: Removed 1 row containing missing values or values outside the scale range
## (`geom_text_repel()`).
```

```
## `geom_smooth()` using formula = 'y ~ x'
```

```
## Warning: Removed 1 row containing missing values or values outside the scale range
## (`geom_text_repel()`).
```

```
## `geom_smooth()` using formula = 'y ~ x'
```

```
## Warning: Removed 1 row containing missing values or values outside the scale range
## (`geom_text_repel()`).
```

```
plot(p.td)

#ggsave(file = 'results/fitness/fitnessEffects_summaryPlot_traitxTD.png', p.td, height = 14, width = 14)
#ggsave(file = 'results/fitness/fitnessEffects_summaryPlot_traitxTD.pdf', p.td, height = 14, width = 14)
```

#### 2.4 Do trait values predict end-of-year survival?

Mostly, no - it seems that there is not enough mortality during one
season to predict how trait values affect it

Most of these models did not converge,likely because there was little
mortality within the year, so below code is not run

```
table(dat$Survival_09_2023, dat$MiniCG_Site)

# do these traits predict survival?

# reorder gardens by climate MAT

tmp <- aggregate(dat$garden_MAT_2023, by = list(dat$MiniCG_Site_2023), mean)
ord <- tmp$Group.1[order(tmp$x, decreasing = T)]
dat$MiniCG_Site_2023 <- factor(dat$MiniCG_Site_2023, levels = ord)

sites <- c("SWMN", "WI", "ID", "WYO", "EVERGREEN", "MSU", "OSU", "MORTON", "WSU", "PENN", "VA", "SU")

sites.mat <- dat$garden_MAT_2023[match(sites, dat$MiniCG_Site)]

mods.mat <- list()
for(n in 1:length(traits)){
    
    trait <- traits[n]
    
     print(paste('Model for:', trait))
    
    mod <- glmer(paste0('Survival_09_2023 ~', trait,  '* garden_MAT_2023 + (1 | MiniCG_Site/block) + (1 | Genotype)'),
                data = dat,
                family = binomial
    )
    
    print(car::Anova(mod))
    
    p1 <- plot_model(mod, vline.color = "black", show.values = T, type = 'std')
    plot(p1)
    #plot_model(mod, type = 're')
    
    mods.mat[n] <- mod
    names(mods.mat)[n] <- trait
    
    # plot data
    p2 <- ggplot(data = dat, aes_string(y = 'Survival_09_2023', x = trait, color = 'k2_tricho'), na.rm  = TRUE) +
    geom_point(na.rm = TRUE) +
    scale_color_gradient2(high = "darkolivegreen2", mid = "grey20", low = "dodgerblue2", midpoint = 0.5, name = 'P. trichocarpa\nancestry') +
    ggtitle(trait) +
    stat_smooth(color = 'black', method = "glm", method.args = list(family = "binomial")) +
    facet_wrap(~ MiniCG_Site_2023, drop = T, scales = 'free') +
    theme(axis.text.x = element_text(angle = 90, vjust = 0.5, hjust=1, size = 12)) +
    theme(axis.title.x = element_blank())

  plot(p2)
    
}

#plot_models(mods.pc1, vline.color = "black", show.values = T, std.est = T, m.labels = names(mods.pc1))


# with MAP

# reorder gardens by MAP
tmp <- aggregate(dat$garden_MAP_2023, by = list(dat$MiniCG_Site_2023), mean)
ord <- tmp$Group.1[order(tmp$x, decreasing = T)]
dat$MiniCG_Site_2023 <- factor(dat$MiniCG_Site_2023, levels = ord)

sites <- c("SWMN", "WI", "ID", "WYO", "EVERGREEN", "MSU", "OSU", "MORTON", "WSU", "PENN", "VA", "SU")

sites.map <- dat$garden_MAP_2023[match(sites, dat$MiniCG_Site)]


mods.map <- list()

for(n in 1:length(traits)){
    
    trait <- traits[n]
    
    mod <- glmer(paste0('Survival_09_2023 ~', trait,  '* garden_climate_2023_MAP + (1 | MiniCG_Site/block) + (1 | Genotype)'),
                data = dat,
                family = binomial
    )
    
    print(car::Anova(mod))
    
    p1 <- plot_model(mod, vline.color = "black", show.values = T, type = 'std')
    plot(p1)
    #plot_model(mod, type = 're')
    
      mods.map[n] <- mod
    names(mods.map)[n] <- trait
    
        # plot data
    p2 <- ggplot(data = dat, aes_string(y = 'Survival_09_2023', x = trait, color = 'k2_tricho'), na.rm  = TRUE) +
    geom_point(na.rm = TRUE) +
    scale_color_gradient2(high = "darkolivegreen2", mid = "grey20", low = "dodgerblue2", midpoint = 0.5, name = 'P. trichocarpa\nancestry') +
    ggtitle(trait) +
    stat_smooth(color = 'black', method = "glm", method.args = list(family = "binomial")) +
    facet_wrap(~ MiniCG_Site_2023, drop = T, scales = 'free') +
    theme(axis.text.x = element_text(angle = 90, vjust = 0.5, hjust=1, size = 12)) +
    theme(axis.title.x = element_blank())

  plot(p2)
    
}

#plot_models(mods.pc2, vline.color = "black", show.values = T, std.est = T, m.labels = names(mods.pc2))
```

### 3 Statistical models for plasticity

Test whether trait plasticity predicts growth in each garden, and
whether the plasticity-growth relationship varies by garden climate.

#### 3.1 Does trait plasticity (RDPI) predict growth in 2023?

```
# merge plasticity values with main df

# get plasticity traits
plast <- grep('rdpi', colnames(rdpi), value = T)
# remove 2024 traits
plast <- plast[grep('2024', plast, invert = T)]

# add empty columns
dat[,plast] <- NA

# fill in plasticity columns for each genotype
for(n in 1:nrow(rdpi)){
  
  geno <- rownames(rdpi)[n]
  dat[dat$Genotype == geno, plast] <- rdpi[geno,plast]
  
}


traits <- plast

# get rid of redundant stomata traits 
traits <- traits[! traits %in% c("plasticity_rdpi_lower_stomata_count", "plasticity_rdpi_upper_stomata_count_log", "plasticity_rdpi_upper_stomata_density_over_total_density_log", "plasticity_rdpi_stomata_density_both")]

# does plasticity in these traits predict growth increment?
```

##### 3.1.1 Garden MAT

```
# reorder gardens by MAT
tmp <- aggregate(dat$garden_MAT_2023, by = list(dat$MiniCG_Site_2023), mean)
ord <- tmp$Group.1[order(tmp$x, decreasing = T)]
dat$MiniCG_Site_2023 <- factor(dat$MiniCG_Site_2023, levels = ord)

sites <- levels(dat$MiniCG_Site_2023)
# remove UCM - height not measured
sites <- sites[! sites == 'UCM']

sites.mat <- dat$garden_MAT_2023[match(sites, dat$MiniCG_Site)]


# to save results
mods.mat <- list()

# list to save summary plot for each trait with significant effects on fitness
plots.summ <- list()

for(n in 1:length(traits)){
  
  trait <- traits[n]
  
  print(paste('Model for:', trait))
  
  mod <- lmer(paste0('GrowthIncrement_2023 ~', trait,  '* garden_MAT_2023 + Pt + (1 | MiniCG_Site/block) + (1 | Genotype)'),
              data = dat
  )
  
  print(car::Anova(mod))
  
  p1 <- plot_model(mod, vline.color = "black", show.values = T, type = 'std', 
                   title = paste('2023 Growth ~ ', trait, sep = ''))
  plot(p1)
  #plot_model(mod, type = 're')
  
  mods.mat[[n]] <- mod
  names(mods.mat)[n] <- trait
  
  #####################
  # posthoc tests
  # if either the climate or trait x climate are significant, run posthoc tests for each garden
  mod.info <- Anova(mod)
  # if any coefficients are significant, plot by garden
  # need to change indexes if model is changed!
  if(mod.info$`Pr(>Chisq)`[1] <= 0.05 | mod.info$`Pr(>Chisq)`[2] <= 0.05 | mod.info$`Pr(>Chisq)`[4] <=0.05){
    
# list to save plots for each site
    plots.sub <- list()
    
    # dataframe to save outputs from posthoc tests
    df.posthoc <- as.data.frame(matrix(ncol = 3, nrow = length(sites)))
    rownames(df.posthoc) <- sites
    colnames(df.posthoc) <- c('mat', 'slope', 'pval')
    df.posthoc$mat <- sites.mat
    
    for(s in 1:length(sites)){
      
      site <- sites[s]
      dat.sub <- subset(dat, MiniCG_Site_2023 == site)
      # skip if site is missing data
      if(sum(!is.na(dat.sub[,trait])) > 10){
        mod.sub <- lmer(paste0('GrowthIncrement_2023 ~', trait, '+ Pt + (1 | block) + (1 | Genotype)'),
                        data = dat.sub)
        # get pval for trait
        mod.sub.info <- Anova(mod.sub)
        df.posthoc[site, 'pval'] <- mod.sub.info$`Pr(>Chisq)`[1]
        
        # get slopes from fixed effect of trait (effect on growth)
        df.posthoc[site, 'slope'] <- fixef(mod.sub)[2]
        
        # make title pretty
        maintitle <- paste(site, '\n', trait_names_df[trait, 'trait_names'], '\n', 'p = ', round(df.posthoc$pval[s], 3), sep = '')
        maintitle <- paste(strwrap(maintitle, width = 26), collapse = '\n')   
        
        # plot
        p <- ggplot(data = dat.sub, aes_string(y = 'GrowthIncrement_2023', x = trait, color = 'k2_tricho'), na.rm  = TRUE) +
          geom_point(na.rm = TRUE) +
          scale_color_gradient2(high = "darkolivegreen2", mid = "grey20", low = "dodgerblue2", midpoint = 0.5, name = 'P. trichocarpa\nancestry', guide = 'none') +
          ggtitle(maintitle) +
          geom_smooth(color = 'black', lty = ifelse(df.posthoc$pval[s] <= 0.05, 1,2), method = 'lm', formula = y ~ x, na.rm = TRUE) +
          theme(axis.text.x = element_text(angle = 90, vjust = 0.5, hjust=1, size = 12)) +
          theme(axis.title.x = element_blank())
        
        #plot(p)
        plots.sub[[s]] <- p
        
      }
    }

    # pvals are adjusted for summary plot
    df.posthoc$pvals.adj <- p.adjust(df.posthoc$pval, method = 'BH') 
    
    # plot panel with relationship for each garden
    # remove empty plots
    plots.sub <- plots.sub[lengths(plots.sub)>0]
    # plot in grid
    p1 <- grid.arrange(grobs = plots.sub, ncol = 4)
    #plot(p1)
    
    # setup text showing significance
    sig_text <- paste0('Plasticity ', pval_stars(mod.info[1, 'Pr(>Chisq)']), '\n',
                       'Climate ', pval_stars(mod.info[2, 'Pr(>Chisq)']), '\n',
                       'Plasticity x Climate ', pval_stars(mod.info[4, 'Pr(>Chisq)']))
    
    # wrap title
    maintitle <- trait_names_df[trait, 'trait_names']
    maintitle <- paste(strwrap(maintitle, width = 26), collapse = '\n')
    
    # plot slope of fitness relationship vs climate
    p2 <- ggplot(df.posthoc, aes(x = mat, y = slope), na.rm  = TRUE) +
      geom_smooth(method = 'lm', col = 'black', na.rm = TRUE) +
      geom_hline(yintercept = 0, lty = 2) +
      geom_point(size = 5, stroke = 2, shape = ifelse(df.posthoc$pvals.adj <= 0.05, 16, 1), aes(col = mat), na.rm = TRUE) +
      scale_color_gradient2(high = "red2", mid = "grey", low = "blue2", midpoint = mean(range(sites.mat)), limits = range_mat, name = 'Garden MAT (°C)') +
      ggtitle(maintitle,
              subtitle = sig_text) +
      xlab('Garden MAT') +
      ylab('Effect on Growth Increment') +
      geom_text_repel(label = rownames(df.posthoc), aes(x = mat, y = slope), box.padding = 0.5, min.segment.length = 1, size = 4) +
        theme(plot.title = element_text(size = 15),
        plot.subtitle = element_text(size = 10),
        legend.position = 'none')
    plot(p2)
    
    plots.summ[[n]] <- p2
    names(plots.summ)[n] <- trait
    
    # add legend
    leg_plot <- ggplot(df.posthoc, aes(x = mat, y = slope), na.rm  = TRUE) +
      geom_point(aes(color = mat))+
      scale_color_gradient2(high = "red2", mid = "grey", low = "blue2", midpoint = mean(range(sites.mat)), limits = range_mat, name = 'Garden MAT (°C)') +
      theme(legend.position = 'right')
    
    legend <- cowplot::get_legend(leg_plot)
    
  }
}
```

```
## [1] "Model for: plasticity_rdpi_GrowthIncrement_2023_log"
## Analysis of Deviance Table (Type II Wald chisquare tests)
## 
## Response: GrowthIncrement_2023
##                                                            Chisq Df Pr(>Chisq)
## plasticity_rdpi_GrowthIncrement_2023_log                  0.5529  1  0.4571218
## garden_MAT_2023                                           2.6586  1  0.1029917
## Pt                                                       33.2616  1  8.056e-09
## plasticity_rdpi_GrowthIncrement_2023_log:garden_MAT_2023 11.1628  1  0.0008345
##                                                             
## plasticity_rdpi_GrowthIncrement_2023_log                    
## garden_MAT_2023                                             
## Pt                                                       ***
## plasticity_rdpi_GrowthIncrement_2023_log:garden_MAT_2023 ***
## ---
## Signif. codes:  0 '***' 0.001 '**' 0.01 '*' 0.05 '.' 0.1 ' ' 1
```

```
## boundary (singular) fit: see help('isSingular')
## boundary (singular) fit: see help('isSingular')
## boundary (singular) fit: see help('isSingular')
## boundary (singular) fit: see help('isSingular')
## boundary (singular) fit: see help('isSingular')
## boundary (singular) fit: see help('isSingular')
```

```
## `geom_smooth()` using formula = 'y ~ x'
```

```
## [1] "Model for: plasticity_rdpi_DOY_Stage2_2023"
## Analysis of Deviance Table (Type II Wald chisquare tests)
## 
## Response: GrowthIncrement_2023
##                                                   Chisq Df Pr(>Chisq)    
## plasticity_rdpi_DOY_Stage2_2023                  0.1785  1     0.6727    
## garden_MAT_2023                                  2.6562  1     0.1031    
## Pt                                              37.1309  1  1.105e-09 ***
## plasticity_rdpi_DOY_Stage2_2023:garden_MAT_2023  0.0214  1     0.8836    
## ---
## Signif. codes:  0 '***' 0.001 '**' 0.01 '*' 0.05 '.' 0.1 ' ' 1
```

```
## [1] "Model for: plasticity_rdpi_DOY_Stage3_2023"
## Analysis of Deviance Table (Type II Wald chisquare tests)
## 
## Response: GrowthIncrement_2023
##                                                   Chisq Df Pr(>Chisq)    
## plasticity_rdpi_DOY_Stage3_2023                  0.5801  1     0.4463    
## garden_MAT_2023                                  2.6570  1     0.1031    
## Pt                                              37.9710  1   7.18e-10 ***
## plasticity_rdpi_DOY_Stage3_2023:garden_MAT_2023  0.8630  1     0.3529    
## ---
## Signif. codes:  0 '***' 0.001 '**' 0.01 '*' 0.05 '.' 0.1 ' ' 1
```

```
## [1] "Model for: plasticity_rdpi_DOY_Stage6_2023"
## Analysis of Deviance Table (Type II Wald chisquare tests)
## 
## Response: GrowthIncrement_2023
##                                                   Chisq Df Pr(>Chisq)    
## plasticity_rdpi_DOY_Stage6_2023                  0.0440  1  0.8338252    
## garden_MAT_2023                                  2.6646  1  0.1026013    
## Pt                                              28.3909  1  9.913e-08 ***
## plasticity_rdpi_DOY_Stage6_2023:garden_MAT_2023 14.8456  1  0.0001167 ***
## ---
## Signif. codes:  0 '***' 0.001 '**' 0.01 '*' 0.05 '.' 0.1 ' ' 1
```

```
## boundary (singular) fit: see help('isSingular')
## boundary (singular) fit: see help('isSingular')
## boundary (singular) fit: see help('isSingular')
## boundary (singular) fit: see help('isSingular')
## boundary (singular) fit: see help('isSingular')
## boundary (singular) fit: see help('isSingular')
```

```
## `geom_smooth()` using formula = 'y ~ x'
```

```
## [1] "Model for: plasticity_rdpi_DOY_last_budset_2023"
## Analysis of Deviance Table (Type II Wald chisquare tests)
## 
## Response: GrowthIncrement_2023
##                                                        Chisq Df Pr(>Chisq)    
## plasticity_rdpi_DOY_last_budset_2023                  1.8989  1     0.1682    
## garden_MAT_2023                                       2.6470  1     0.1037    
## Pt                                                   38.5158  1  5.431e-10 ***
## plasticity_rdpi_DOY_last_budset_2023:garden_MAT_2023  1.0211  1     0.3123    
## ---
## Signif. codes:  0 '***' 0.001 '**' 0.01 '*' 0.05 '.' 0.1 ' ' 1
```

```
## [1] "Model for: plasticity_rdpi_stage7_presence_2023"
## Analysis of Deviance Table (Type II Wald chisquare tests)
## 
## Response: GrowthIncrement_2023
##                                                        Chisq Df Pr(>Chisq)    
## plasticity_rdpi_stage7_presence_2023                  0.0078  1     0.9294    
## garden_MAT_2023                                       2.6564  1     0.1031    
## Pt                                                   36.3972  1  1.609e-09 ***
## plasticity_rdpi_stage7_presence_2023:garden_MAT_2023  0.0063  1     0.9367    
## ---
## Signif. codes:  0 '***' 0.001 '**' 0.01 '*' 0.05 '.' 0.1 ' ' 1
```

```
## [1] "Model for: plasticity_rdpi_growing_season_days_2023"
## Analysis of Deviance Table (Type II Wald chisquare tests)
## 
## Response: GrowthIncrement_2023
##                                                            Chisq Df Pr(>Chisq)
## plasticity_rdpi_growing_season_days_2023                  2.4326  1   0.118834
## garden_MAT_2023                                           2.6733  1   0.102045
## Pt                                                       36.2570  1  1.729e-09
## plasticity_rdpi_growing_season_days_2023:garden_MAT_2023  9.8295  1   0.001717
##                                                             
## plasticity_rdpi_growing_season_days_2023                    
## garden_MAT_2023                                             
## Pt                                                       ***
## plasticity_rdpi_growing_season_days_2023:garden_MAT_2023 ** 
## ---
## Signif. codes:  0 '***' 0.001 '**' 0.01 '*' 0.05 '.' 0.1 ' ' 1
```

```
## boundary (singular) fit: see help('isSingular')
## boundary (singular) fit: see help('isSingular')
## boundary (singular) fit: see help('isSingular')
## boundary (singular) fit: see help('isSingular')
## boundary (singular) fit: see help('isSingular')
## boundary (singular) fit: see help('isSingular')
## boundary (singular) fit: see help('isSingular')
```

```
## `geom_smooth()` using formula = 'y ~ x'
```

```
## [1] "Model for: plasticity_rdpi_DOY_Stage7_2023"
## Analysis of Deviance Table (Type II Wald chisquare tests)
## 
## Response: GrowthIncrement_2023
##                                                   Chisq Df Pr(>Chisq)    
## plasticity_rdpi_DOY_Stage7_2023                  0.5461  1   0.459928    
## garden_MAT_2023                                  3.0410  1   0.081187 .  
## Pt                                              21.9881  1  2.743e-06 ***
## plasticity_rdpi_DOY_Stage7_2023:garden_MAT_2023  7.8311  1   0.005135 ** 
## ---
## Signif. codes:  0 '***' 0.001 '**' 0.01 '*' 0.05 '.' 0.1 ' ' 1
```

```
## boundary (singular) fit: see help('isSingular')
## boundary (singular) fit: see help('isSingular')
## boundary (singular) fit: see help('isSingular')
## boundary (singular) fit: see help('isSingular')
## boundary (singular) fit: see help('isSingular')
## boundary (singular) fit: see help('isSingular')
## boundary (singular) fit: see help('isSingular')
```

```
## `geom_smooth()` using formula = 'y ~ x'
```

```
## [1] "Model for: plasticity_rdpi_licor_gsw"
## Analysis of Deviance Table (Type II Wald chisquare tests)
## 
## Response: GrowthIncrement_2023
##                                             Chisq Df Pr(>Chisq)    
## plasticity_rdpi_licor_gsw                  0.3620  1     0.5474    
## garden_MAT_2023                            2.6546  1     0.1032    
## Pt                                        30.7963  1  2.866e-08 ***
## plasticity_rdpi_licor_gsw:garden_MAT_2023  2.4193  1     0.1198    
## ---
## Signif. codes:  0 '***' 0.001 '**' 0.01 '*' 0.05 '.' 0.1 ' ' 1
```

```
## [1] "Model for: plasticity_rdpi_licor_gbw"
```

```
## Warning: Some predictor variables are on very different scales: consider
## rescaling
```

```
## Analysis of Deviance Table (Type II Wald chisquare tests)
## 
## Response: GrowthIncrement_2023
##                                             Chisq Df Pr(>Chisq)    
## plasticity_rdpi_licor_gbw                  0.2336  1     0.6289    
## garden_MAT_2023                            2.6544  1     0.1033    
## Pt                                        36.7642  1  1.333e-09 ***
## plasticity_rdpi_licor_gbw:garden_MAT_2023  0.2448  1     0.6208    
## ---
## Signif. codes:  0 '***' 0.001 '**' 0.01 '*' 0.05 '.' 0.1 ' ' 1
```

```
## [1] "Model for: plasticity_rdpi_licor_PhiPS2"
## Analysis of Deviance Table (Type II Wald chisquare tests)
## 
## Response: GrowthIncrement_2023
##                                                Chisq Df Pr(>Chisq)    
## plasticity_rdpi_licor_PhiPS2                  2.0205  1    0.15519    
## garden_MAT_2023                               2.6392  1    0.10425    
## Pt                                           38.5390  1  5.367e-10 ***
## plasticity_rdpi_licor_PhiPS2:garden_MAT_2023  4.8265  1    0.02803 *  
## ---
## Signif. codes:  0 '***' 0.001 '**' 0.01 '*' 0.05 '.' 0.1 ' ' 1
```

```
## boundary (singular) fit: see help('isSingular')
## boundary (singular) fit: see help('isSingular')
## boundary (singular) fit: see help('isSingular')
## boundary (singular) fit: see help('isSingular')
```

```
## `geom_smooth()` using formula = 'y ~ x'
```

```
## [1] "Model for: plasticity_rdpi_licor_ETR"
## Analysis of Deviance Table (Type II Wald chisquare tests)
## 
## Response: GrowthIncrement_2023
##                                             Chisq Df Pr(>Chisq)    
## plasticity_rdpi_licor_ETR                  0.2392  1    0.62482    
## garden_MAT_2023                            2.6530  1    0.10336    
## Pt                                        37.1970  1  1.068e-09 ***
## plasticity_rdpi_licor_ETR:garden_MAT_2023  3.3561  1    0.06695 .  
## ---
## Signif. codes:  0 '***' 0.001 '**' 0.01 '*' 0.05 '.' 0.1 ' ' 1
```

```
## [1] "Model for: plasticity_rdpi_licor_Fs"
## Analysis of Deviance Table (Type II Wald chisquare tests)
## 
## Response: GrowthIncrement_2023
##                                            Chisq Df Pr(>Chisq)    
## plasticity_rdpi_licor_Fs                  2.0362  1   0.153591    
## garden_MAT_2023                           2.6540  1   0.103289    
## Pt                                       33.1504  1   8.53e-09 ***
## plasticity_rdpi_licor_Fs:garden_MAT_2023  9.7015  1   0.001841 ** 
## ---
## Signif. codes:  0 '***' 0.001 '**' 0.01 '*' 0.05 '.' 0.1 ' ' 1
```

```
## boundary (singular) fit: see help('isSingular')
## boundary (singular) fit: see help('isSingular')
## boundary (singular) fit: see help('isSingular')
## boundary (singular) fit: see help('isSingular')
```

```
## `geom_smooth()` using formula = 'y ~ x'
```

```
## [1] "Model for: plasticity_rdpi_licor_Fm."
## Analysis of Deviance Table (Type II Wald chisquare tests)
## 
## Response: GrowthIncrement_2023
##                                             Chisq Df Pr(>Chisq)    
## plasticity_rdpi_licor_Fm.                  0.1196  1     0.7295    
## garden_MAT_2023                            2.6495  1     0.1036    
## Pt                                        36.2473  1  1.738e-09 ***
## plasticity_rdpi_licor_Fm.:garden_MAT_2023  0.1987  1     0.6558    
## ---
## Signif. codes:  0 '***' 0.001 '**' 0.01 '*' 0.05 '.' 0.1 ' ' 1
```

```
## [1] "Model for: plasticity_rdpi_leaf_thickness_avg_mm_2023"
## Analysis of Deviance Table (Type II Wald chisquare tests)
## 
## Response: GrowthIncrement_2023
##                                                              Chisq Df
## plasticity_rdpi_leaf_thickness_avg_mm_2023                  0.9235  1
## garden_MAT_2023                                             2.6388  1
## Pt                                                         32.5231  1
## plasticity_rdpi_leaf_thickness_avg_mm_2023:garden_MAT_2023 10.6043  1
##                                                            Pr(>Chisq)    
## plasticity_rdpi_leaf_thickness_avg_mm_2023                   0.336569    
## garden_MAT_2023                                              0.104283    
## Pt                                                          1.178e-08 ***
## plasticity_rdpi_leaf_thickness_avg_mm_2023:garden_MAT_2023   0.001128 ** 
## ---
## Signif. codes:  0 '***' 0.001 '**' 0.01 '*' 0.05 '.' 0.1 ' ' 1
```

```
## boundary (singular) fit: see help('isSingular')
## boundary (singular) fit: see help('isSingular')
## boundary (singular) fit: see help('isSingular')
## boundary (singular) fit: see help('isSingular')
## boundary (singular) fit: see help('isSingular')
## boundary (singular) fit: see help('isSingular')
```

```
## `geom_smooth()` using formula = 'y ~ x'
```

```
## [1] "Model for: plasticity_rdpi_LMA_g_m2_2023"
## Analysis of Deviance Table (Type II Wald chisquare tests)
## 
## Response: GrowthIncrement_2023
##                                                 Chisq Df Pr(>Chisq)    
## plasticity_rdpi_LMA_g_m2_2023                  0.4501  1    0.50228    
## garden_MAT_2023                                2.6586  1    0.10299    
## Pt                                            31.2131  1  2.312e-08 ***
## plasticity_rdpi_LMA_g_m2_2023:garden_MAT_2023  6.5300  1    0.01061 *  
## ---
## Signif. codes:  0 '***' 0.001 '**' 0.01 '*' 0.05 '.' 0.1 ' ' 1
```

```
## boundary (singular) fit: see help('isSingular')
## boundary (singular) fit: see help('isSingular')
## boundary (singular) fit: see help('isSingular')
## boundary (singular) fit: see help('isSingular')
## boundary (singular) fit: see help('isSingular')
```

```
## `geom_smooth()` using formula = 'y ~ x'
```

```
## [1] "Model for: plasticity_rdpi_leaf_mass_g_2023_log"
## Analysis of Deviance Table (Type II Wald chisquare tests)
## 
## Response: GrowthIncrement_2023
##                                                        Chisq Df Pr(>Chisq)    
## plasticity_rdpi_leaf_mass_g_2023_log                  1.8169  1    0.17769    
## garden_MAT_2023                                       2.6775  1    0.10178    
## Pt                                                   40.1383  1  2.366e-10 ***
## plasticity_rdpi_leaf_mass_g_2023_log:garden_MAT_2023  5.1633  1    0.02307 *  
## ---
## Signif. codes:  0 '***' 0.001 '**' 0.01 '*' 0.05 '.' 0.1 ' ' 1
```

```
## boundary (singular) fit: see help('isSingular')
## boundary (singular) fit: see help('isSingular')
## boundary (singular) fit: see help('isSingular')
## boundary (singular) fit: see help('isSingular')
## boundary (singular) fit: see help('isSingular')
## boundary (singular) fit: see help('isSingular')
```

```
## `geom_smooth()` using formula = 'y ~ x'
```

```
## [1] "Model for: plasticity_rdpi_leaf_area_cm2_2023_log"
## Analysis of Deviance Table (Type II Wald chisquare tests)
## 
## Response: GrowthIncrement_2023
##                                                          Chisq Df Pr(>Chisq)
## plasticity_rdpi_leaf_area_cm2_2023_log                  8.3841  1   0.003785
## garden_MAT_2023                                         2.6360  1   0.104465
## Pt                                                     43.5708  1  4.089e-11
## plasticity_rdpi_leaf_area_cm2_2023_log:garden_MAT_2023  7.2532  1   0.007077
##                                                           
## plasticity_rdpi_leaf_area_cm2_2023_log                 ** 
## garden_MAT_2023                                           
## Pt                                                     ***
## plasticity_rdpi_leaf_area_cm2_2023_log:garden_MAT_2023 ** 
## ---
## Signif. codes:  0 '***' 0.001 '**' 0.01 '*' 0.05 '.' 0.1 ' ' 1
```

```
## boundary (singular) fit: see help('isSingular')
## boundary (singular) fit: see help('isSingular')
## boundary (singular) fit: see help('isSingular')
## boundary (singular) fit: see help('isSingular')
## boundary (singular) fit: see help('isSingular')
```

```
## `geom_smooth()` using formula = 'y ~ x'
```

```
## [1] "Model for: plasticity_rdpi_lower_stomata_pore_length_mean_um"
## Analysis of Deviance Table (Type II Wald chisquare tests)
## 
## Response: GrowthIncrement_2023
##                                                                     Chisq Df
## plasticity_rdpi_lower_stomata_pore_length_mean_um                  2.1319  1
## garden_MAT_2023                                                    2.6692  1
## Pt                                                                39.6028  1
## plasticity_rdpi_lower_stomata_pore_length_mean_um:garden_MAT_2023  1.3154  1
##                                                                   Pr(>Chisq)
## plasticity_rdpi_lower_stomata_pore_length_mean_um                     0.1443
## garden_MAT_2023                                                       0.1023
## Pt                                                                 3.112e-10
## plasticity_rdpi_lower_stomata_pore_length_mean_um:garden_MAT_2023     0.2514
##                                                                      
## plasticity_rdpi_lower_stomata_pore_length_mean_um                    
## garden_MAT_2023                                                      
## Pt                                                                ***
## plasticity_rdpi_lower_stomata_pore_length_mean_um:garden_MAT_2023    
## ---
## Signif. codes:  0 '***' 0.001 '**' 0.01 '*' 0.05 '.' 0.1 ' ' 1
```

```
## [1] "Model for: plasticity_rdpi_upper_stomata_presence"
## Analysis of Deviance Table (Type II Wald chisquare tests)
## 
## Response: GrowthIncrement_2023
##                                                          Chisq Df Pr(>Chisq)
## plasticity_rdpi_upper_stomata_presence                  2.1745  1     0.1403
## garden_MAT_2023                                         2.6710  1     0.1022
## Pt                                                     39.9846  1   2.56e-10
## plasticity_rdpi_upper_stomata_presence:garden_MAT_2023  0.4674  1     0.4942
##                                                           
## plasticity_rdpi_upper_stomata_presence                    
## garden_MAT_2023                                           
## Pt                                                     ***
## plasticity_rdpi_upper_stomata_presence:garden_MAT_2023    
## ---
## Signif. codes:  0 '***' 0.001 '**' 0.01 '*' 0.05 '.' 0.1 ' ' 1
```

```
## [1] "Model for: plasticity_rdpi_upper_stomata_pore_length_mean_um"
## Analysis of Deviance Table (Type II Wald chisquare tests)
## 
## Response: GrowthIncrement_2023
##                                                                     Chisq Df
## plasticity_rdpi_upper_stomata_pore_length_mean_um                  0.0176  1
## garden_MAT_2023                                                    2.7384  1
## Pt                                                                29.7554  1
## plasticity_rdpi_upper_stomata_pore_length_mean_um:garden_MAT_2023  0.0469  1
##                                                                   Pr(>Chisq)
## plasticity_rdpi_upper_stomata_pore_length_mean_um                    0.89454
## garden_MAT_2023                                                      0.09796
## Pt                                                                 4.901e-08
## plasticity_rdpi_upper_stomata_pore_length_mean_um:garden_MAT_2023    0.82854
##                                                                      
## plasticity_rdpi_upper_stomata_pore_length_mean_um                    
## garden_MAT_2023                                                   .  
## Pt                                                                ***
## plasticity_rdpi_upper_stomata_pore_length_mean_um:garden_MAT_2023    
## ---
## Signif. codes:  0 '***' 0.001 '**' 0.01 '*' 0.05 '.' 0.1 ' ' 1
```

```
## [1] "Model for: plasticity_rdpi_upper_stomata_density_mm2_log"
## Analysis of Deviance Table (Type II Wald chisquare tests)
## 
## Response: GrowthIncrement_2023
##                                                                 Chisq Df
## plasticity_rdpi_upper_stomata_density_mm2_log                  0.0598  1
## garden_MAT_2023                                                2.6566  1
## Pt                                                            36.6838  1
## plasticity_rdpi_upper_stomata_density_mm2_log:garden_MAT_2023  0.0366  1
##                                                               Pr(>Chisq)    
## plasticity_rdpi_upper_stomata_density_mm2_log                     0.8067    
## garden_MAT_2023                                                   0.1031    
## Pt                                                             1.389e-09 ***
## plasticity_rdpi_upper_stomata_density_mm2_log:garden_MAT_2023     0.8482    
## ---
## Signif. codes:  0 '***' 0.001 '**' 0.01 '*' 0.05 '.' 0.1 ' ' 1
```

```
## [1] "Model for: plasticity_rdpi_lower_stomata_density_mm2"
## Analysis of Deviance Table (Type II Wald chisquare tests)
## 
## Response: GrowthIncrement_2023
##                                                             Chisq Df Pr(>Chisq)
## plasticity_rdpi_lower_stomata_density_mm2                  6.9185  1   0.008531
## garden_MAT_2023                                            2.6451  1   0.103870
## Pt                                                        44.5898  1   2.43e-11
## plasticity_rdpi_lower_stomata_density_mm2:garden_MAT_2023  0.0122  1   0.912122
##                                                              
## plasticity_rdpi_lower_stomata_density_mm2                 ** 
## garden_MAT_2023                                              
## Pt                                                        ***
## plasticity_rdpi_lower_stomata_density_mm2:garden_MAT_2023    
## ---
## Signif. codes:  0 '***' 0.001 '**' 0.01 '*' 0.05 '.' 0.1 ' ' 1
```

```
## boundary (singular) fit: see help('isSingular')
## boundary (singular) fit: see help('isSingular')
## boundary (singular) fit: see help('isSingular')
## boundary (singular) fit: see help('isSingular')
## boundary (singular) fit: see help('isSingular')
## boundary (singular) fit: see help('isSingular')
```

```
## `geom_smooth()` using formula = 'y ~ x'
```

```
## [1] "Model for: plasticity_rdpi_stomata_ratio_log"
## Analysis of Deviance Table (Type II Wald chisquare tests)
## 
## Response: GrowthIncrement_2023
##                                                     Chisq Df Pr(>Chisq)    
## plasticity_rdpi_stomata_ratio_log                  1.6148  1     0.2038    
## garden_MAT_2023                                    2.6478  1     0.1037    
## Pt                                                37.8004  1  7.837e-10 ***
## plasticity_rdpi_stomata_ratio_log:garden_MAT_2023  0.1474  1     0.7010    
## ---
## Signif. codes:  0 '***' 0.001 '**' 0.01 '*' 0.05 '.' 0.1 ' ' 1
```

```
plots.summ <- plots.summ[lengths(plots.summ)>0]

# add legend to beginning of plot list

plots2 <- plots.summ
plots2[[length(plots.summ)+1]] <- legend

p3 <- grid.arrange(grobs = plots2, ncol = 4)
```

```
## `geom_smooth()` using formula = 'y ~ x'
```

```
## `geom_smooth()` using formula = 'y ~ x'
## `geom_smooth()` using formula = 'y ~ x'
## `geom_smooth()` using formula = 'y ~ x'
## `geom_smooth()` using formula = 'y ~ x'
## `geom_smooth()` using formula = 'y ~ x'
## `geom_smooth()` using formula = 'y ~ x'
## `geom_smooth()` using formula = 'y ~ x'
## `geom_smooth()` using formula = 'y ~ x'
## `geom_smooth()` using formula = 'y ~ x'
## `geom_smooth()` using formula = 'y ~ x'
```

```
plot(p3)

#ggsave(file = 'results/fitness/fitnessEffects_summaryPlot_traitPlasticityxMAT_2023.png', p3, height = 12, width = 14)
#ggsave(file = 'results/fitness/fitnessEffects_summaryPlot_traitPlasticityxMAT_2023.pdf', p3, height = 12, width = 18)

# rename plots list for later use
plots.plast.mat.23 <- plots.summ
```

##### 3.1.2 Garden MAP

```
################################################
# with MAP

# reorder gardens by MAP
tmp <- aggregate(dat$garden_MAP_2023, by = list(dat$MiniCG_Site_2023), mean)
ord <- tmp$Group.1[order(tmp$x, decreasing = T)]
dat$MiniCG_Site_2023 <- factor(dat$MiniCG_Site_2023, levels = ord)

sites <- levels(dat$MiniCG_Site_2023)
# remove UCM - height not measured
sites <- sites[! sites == 'UCM']

sites.map <- dat$garden_MAP_2023[match(sites, dat$MiniCG_Site)]


# to save results
mods.map <- list()

# list to save summary plot for each trait with significant effects on fitness
plots.summ <- list()

for(n in 1:length(traits)){
  
  trait <- traits[n]
  
  mod <- lmer(paste0('GrowthIncrement_2023 ~', trait,  '* garden_MAP_2023 + Pt + (1 | MiniCG_Site/block) + (1 | Genotype)'),
              data = dat
  )
  
  print(car::Anova(mod))
  
  p1 <- plot_model(mod, vline.color = "black", show.values = T, type = 'std', 
                   title = paste('2023 Growth ~ ', trait, sep = ''))
  plot(p1)
  #plot_model(mod, type = 're')
  
  mods.map[[n]] <- mod
  names(mods.map)[n] <- trait
  
  #####################
  # posthoc tests
  # if either the climate or trait x climate are significant, run posthoc tests for each garden
  mod.info <- Anova(mod)
  # if any coefficients are significant, plot by garden
  # need to change indexes if model is changed!
  if(mod.info$`Pr(>Chisq)`[1] <= 0.05 | mod.info$`Pr(>Chisq)`[2] <= 0.05 | mod.info$`Pr(>Chisq)`[4] <=0.05){
    
# list to save plots for each site
    plots.sub <- list()
    
    
    # dataframe to save outputs from posthoc tests
    df.posthoc <- as.data.frame(matrix(ncol = 3, nrow = length(sites)))
    rownames(df.posthoc) <- sites
    colnames(df.posthoc) <- c('map', 'slope', 'pval')
    df.posthoc$map <- sites.map
    
    for(s in 1:length(sites)){
      
      site <- sites[s]
      dat.sub <- subset(dat, MiniCG_Site_2023 == site)
      # skip if site is missing data
      if(sum(!is.na(dat.sub[,trait])) > 10){
        mod.sub <- lmer(paste0('GrowthIncrement_2023 ~', trait, '+ Pt + (1 | block) + (1 | Genotype)'),
                        data = dat.sub)
        # get pval for trait
        mod.sub.info <- Anova(mod.sub)
        df.posthoc[site, 'pval'] <- mod.sub.info$`Pr(>Chisq)`[1]
        
        # get slopes from fixed effect of trait (effect on growth)
        df.posthoc[site, 'slope'] <- fixef(mod.sub)[2]
        
        # make title pretty
        maintitle <- paste(site, '\n', trait_names_df[trait, 'trait_names'], '\n', 'p = ', round(df.posthoc$pval[s], 3), sep = '')
        maintitle <- paste(strwrap(maintitle, width = 26), collapse = '\n') 
        
        # plot
        p <- ggplot(data = dat.sub, aes_string(y = 'GrowthIncrement_2023', x = trait, color = 'k2_tricho'), na.rm  = TRUE) +
          geom_point(na.rm = TRUE) +
          scale_color_gradient2(high = "darkolivegreen2", mid = "grey20", low = "dodgerblue2", midpoint = 0.5, name = 'P. trichocarpa\nancestry', guide = 'none') +
          ggtitle(maintitle) +
          geom_smooth(color = 'black', lty = ifelse(df.posthoc$pval[s] <= 0.05, 1,2), method = 'lm', formula = y ~ x, na.rm = TRUE) +
          theme(axis.text.x = element_text(angle = 90, vjust = 0.5, hjust=1, size = 12)) +
          theme(axis.title.x = element_blank())
        
        #plot(p)
        plots.sub[[s]] <- p
        
      }
    }

    # pvals are adjusted for summary plot
    df.posthoc$pvals.adj <- p.adjust(df.posthoc$pval, method = 'BH') 
    
    # plot panel with relationship for each garden
    # remove empty plots
    plots.sub <- plots.sub[lengths(plots.sub)>0]
    # plot in grid
    p1 <- grid.arrange(grobs = plots.sub, ncol = 4)
    #plot(p1)
    
    # setup text showing significance
    sig_text <- paste0('Plasticity ', pval_stars(mod.info[1, 'Pr(>Chisq)']), '\n',
                       'Climate ', pval_stars(mod.info[2, 'Pr(>Chisq)']), '\n',
                       'Plasticity x Climate ', pval_stars(mod.info[4, 'Pr(>Chisq)']))
    
    # wrap title
    maintitle <- trait_names_df[trait, 'trait_names']
    maintitle <- paste(strwrap(maintitle, width = 26), collapse = '\n')
    
    # plot slope of fitness relationship vs climate
    p2 <- ggplot(df.posthoc, aes(x = map, y = slope), na.rm  = TRUE) +
      geom_smooth(method = 'lm', col = 'black', na.rm = TRUE) +
      geom_hline(yintercept = 0, lty = 2) +
      geom_point(size = 5, stroke = 2, shape = ifelse(df.posthoc$pvals.adj <= 0.05, 16, 1), aes(col = map), na.rm = TRUE) +
      scale_color_gradient2(high = "steelblue2",  mid = 'grey', low = "sienna3", midpoint = mean(range(sites.map)), limits = range_map, name = 'Garden MAP (mm)') +
      ggtitle(maintitle,
              subtitle = sig_text) +
      xlab('Garden MAP') +
      ylab('Effect on Growth Increment') +
      geom_text_repel(label = rownames(df.posthoc), aes(x = map, y = slope), box.padding = 0.5, min.segment.length = 1, size = 4) +
        theme(plot.title = element_text(size = 15),
        plot.subtitle = element_text(size = 10),
        legend.position = 'none')
    plot(p2)
    
    plots.summ[[n]] <- p2
    names(plots.summ)[n] <- trait
    
     # add legend
    leg_plot <- ggplot(df.posthoc, aes(x = map, y = slope), na.rm  = TRUE) +
      geom_point(aes(color = map))+
      scale_color_gradient2(high = "steelblue2",  mid = 'grey', low = "sienna3", midpoint = mean(range(sites.map)), limits = range_map, name = 'Garden MAP (mm)') +
      theme(legend.position = 'right')
    
    legend <- cowplot::get_legend(leg_plot)
    
  }
}
```

```
## Warning: Some predictor variables are on very different scales: consider
## rescaling
```

```
## Analysis of Deviance Table (Type II Wald chisquare tests)
## 
## Response: GrowthIncrement_2023
##                                                            Chisq Df Pr(>Chisq)
## plasticity_rdpi_GrowthIncrement_2023_log                  0.5752  1    0.44820
## garden_MAP_2023                                           0.9839  1    0.32125
## Pt                                                       33.1575  1  8.499e-09
## plasticity_rdpi_GrowthIncrement_2023_log:garden_MAP_2023  3.5345  1    0.06011
##                                                             
## plasticity_rdpi_GrowthIncrement_2023_log                    
## garden_MAP_2023                                             
## Pt                                                       ***
## plasticity_rdpi_GrowthIncrement_2023_log:garden_MAP_2023 .  
## ---
## Signif. codes:  0 '***' 0.001 '**' 0.01 '*' 0.05 '.' 0.1 ' ' 1
```

```
## Warning: Some predictor variables are on very different scales: consider
## rescaling
```

```
## Analysis of Deviance Table (Type II Wald chisquare tests)
## 
## Response: GrowthIncrement_2023
##                                                   Chisq Df Pr(>Chisq)    
## plasticity_rdpi_DOY_Stage2_2023                  0.1781  1     0.6730    
## garden_MAP_2023                                  0.9733  1     0.3239    
## Pt                                              37.5095  1  9.097e-10 ***
## plasticity_rdpi_DOY_Stage2_2023:garden_MAP_2023  1.1473  1     0.2841    
## ---
## Signif. codes:  0 '***' 0.001 '**' 0.01 '*' 0.05 '.' 0.1 ' ' 1
```

```
## Warning: Some predictor variables are on very different scales: consider
## rescaling
```

```
## Analysis of Deviance Table (Type II Wald chisquare tests)
## 
## Response: GrowthIncrement_2023
##                                                   Chisq Df Pr(>Chisq)    
## plasticity_rdpi_DOY_Stage3_2023                  0.5786  1     0.4469    
## garden_MAP_2023                                  0.9736  1     0.3238    
## Pt                                              37.3903  1   9.67e-10 ***
## plasticity_rdpi_DOY_Stage3_2023:garden_MAP_2023  1.7280  1     0.1887    
## ---
## Signif. codes:  0 '***' 0.001 '**' 0.01 '*' 0.05 '.' 0.1 ' ' 1
```

```
## Warning: Some predictor variables are on very different scales: consider
## rescaling
```

```
## Analysis of Deviance Table (Type II Wald chisquare tests)
## 
## Response: GrowthIncrement_2023
##                                                   Chisq Df Pr(>Chisq)    
## plasticity_rdpi_DOY_Stage6_2023                  0.0441  1     0.8336    
## garden_MAP_2023                                  0.9734  1     0.3238    
## Pt                                              29.6420  1  5.197e-08 ***
## plasticity_rdpi_DOY_Stage6_2023:garden_MAP_2023  0.0094  1     0.9229    
## ---
## Signif. codes:  0 '***' 0.001 '**' 0.01 '*' 0.05 '.' 0.1 ' ' 1
```

```
## Warning: Some predictor variables are on very different scales: consider
## rescaling
```

```
## Analysis of Deviance Table (Type II Wald chisquare tests)
## 
## Response: GrowthIncrement_2023
##                                                        Chisq Df Pr(>Chisq)    
## plasticity_rdpi_DOY_last_budset_2023                  1.9074  1     0.1673    
## garden_MAP_2023                                       0.9704  1     0.3246    
## Pt                                                   38.2651  1  6.176e-10 ***
## plasticity_rdpi_DOY_last_budset_2023:garden_MAP_2023  0.7105  1     0.3993    
## ---
## Signif. codes:  0 '***' 0.001 '**' 0.01 '*' 0.05 '.' 0.1 ' ' 1
```

```
## Warning: Some predictor variables are on very different scales: consider
## rescaling
```

```
## Analysis of Deviance Table (Type II Wald chisquare tests)
## 
## Response: GrowthIncrement_2023
##                                                        Chisq Df Pr(>Chisq)    
## plasticity_rdpi_stage7_presence_2023                  0.0069  1     0.9338    
## garden_MAP_2023                                       0.9822  1     0.3217    
## Pt                                                   37.1360  1  1.102e-09 ***
## plasticity_rdpi_stage7_presence_2023:garden_MAP_2023  1.3347  1     0.2480    
## ---
## Signif. codes:  0 '***' 0.001 '**' 0.01 '*' 0.05 '.' 0.1 ' ' 1
```

```
## Warning: Some predictor variables are on very different scales: consider
## rescaling
```

```
## Analysis of Deviance Table (Type II Wald chisquare tests)
## 
## Response: GrowthIncrement_2023
##                                                            Chisq Df Pr(>Chisq)
## plasticity_rdpi_growing_season_days_2023                  2.3781  1     0.1230
## garden_MAP_2023                                           0.9647  1     0.3260
## Pt                                                       34.8606  1  3.542e-09
## plasticity_rdpi_growing_season_days_2023:garden_MAP_2023  1.1423  1     0.2852
##                                                             
## plasticity_rdpi_growing_season_days_2023                    
## garden_MAP_2023                                             
## Pt                                                       ***
## plasticity_rdpi_growing_season_days_2023:garden_MAP_2023    
## ---
## Signif. codes:  0 '***' 0.001 '**' 0.01 '*' 0.05 '.' 0.1 ' ' 1
```

```
## Warning: Some predictor variables are on very different scales: consider
## rescaling
```

```
## Analysis of Deviance Table (Type II Wald chisquare tests)
## 
## Response: GrowthIncrement_2023
##                                                   Chisq Df Pr(>Chisq)    
## plasticity_rdpi_DOY_Stage7_2023                  0.5508  1     0.4580    
## garden_MAP_2023                                  0.8598  1     0.3538    
## Pt                                              22.8448  1  1.756e-06 ***
## plasticity_rdpi_DOY_Stage7_2023:garden_MAP_2023  0.0321  1     0.8579    
## ---
## Signif. codes:  0 '***' 0.001 '**' 0.01 '*' 0.05 '.' 0.1 ' ' 1
```

```
## Warning: Some predictor variables are on very different scales: consider
## rescaling
```

```
## Analysis of Deviance Table (Type II Wald chisquare tests)
## 
## Response: GrowthIncrement_2023
##                                             Chisq Df Pr(>Chisq)    
## plasticity_rdpi_licor_gsw                  0.3608  1     0.5481    
## garden_MAP_2023                            0.9745  1     0.3236    
## Pt                                        30.8929  1  2.727e-08 ***
## plasticity_rdpi_licor_gsw:garden_MAP_2023  0.4510  1     0.5019    
## ---
## Signif. codes:  0 '***' 0.001 '**' 0.01 '*' 0.05 '.' 0.1 ' ' 1
```

```
## Warning: Some predictor variables are on very different scales: consider
## rescaling
```

```
## Analysis of Deviance Table (Type II Wald chisquare tests)
## 
## Response: GrowthIncrement_2023
##                                             Chisq Df Pr(>Chisq)    
## plasticity_rdpi_licor_gbw                  0.2319  1    0.63013    
## garden_MAP_2023                            0.9588  1    0.32749    
## Pt                                        36.2163  1  1.766e-09 ***
## plasticity_rdpi_licor_gbw:garden_MAP_2023  6.2223  1    0.01262 *  
## ---
## Signif. codes:  0 '***' 0.001 '**' 0.01 '*' 0.05 '.' 0.1 ' ' 1
```

```
## Warning: Some predictor variables are on very different scales: consider
## rescaling
## Warning: Some predictor variables are on very different scales: consider
## rescaling
```

```
## boundary (singular) fit: see help('isSingular')
```

```
## Warning: Some predictor variables are on very different scales: consider
## rescaling
## Warning: Some predictor variables are on very different scales: consider
## rescaling
```

```
## boundary (singular) fit: see help('isSingular')
```

```
## Warning: Some predictor variables are on very different scales: consider
## rescaling
```

```
## boundary (singular) fit: see help('isSingular')
```

```
## Warning: Some predictor variables are on very different scales: consider
## rescaling
## Warning: Some predictor variables are on very different scales: consider
## rescaling
## Warning: Some predictor variables are on very different scales: consider
## rescaling
```

```
## boundary (singular) fit: see help('isSingular')
```

```
## Warning: Some predictor variables are on very different scales: consider
## rescaling
## Warning: Some predictor variables are on very different scales: consider
## rescaling
## Warning: Some predictor variables are on very different scales: consider
## rescaling
## Warning: Some predictor variables are on very different scales: consider
## rescaling
```

```
## `geom_smooth()` using formula = 'y ~ x'
```

```
## Warning: Some predictor variables are on very different scales: consider
## rescaling
```

```
## Analysis of Deviance Table (Type II Wald chisquare tests)
## 
## Response: GrowthIncrement_2023
##                                                Chisq Df Pr(>Chisq)    
## plasticity_rdpi_licor_PhiPS2                  2.0097  1     0.1563    
## garden_MAP_2023                               0.9806  1     0.3221    
## Pt                                           38.3947  1  5.779e-10 ***
## plasticity_rdpi_licor_PhiPS2:garden_MAP_2023  0.4994  1     0.4798    
## ---
## Signif. codes:  0 '***' 0.001 '**' 0.01 '*' 0.05 '.' 0.1 ' ' 1
```

```
## Warning: Some predictor variables are on very different scales: consider
## rescaling
```

```
## Analysis of Deviance Table (Type II Wald chisquare tests)
## 
## Response: GrowthIncrement_2023
##                                             Chisq Df Pr(>Chisq)    
## plasticity_rdpi_licor_ETR                  0.2400  1    0.62422    
## garden_MAP_2023                            0.9612  1    0.32689    
## Pt                                        37.0991  1  1.123e-09 ***
## plasticity_rdpi_licor_ETR:garden_MAP_2023  2.9975  1    0.08339 .  
## ---
## Signif. codes:  0 '***' 0.001 '**' 0.01 '*' 0.05 '.' 0.1 ' ' 1
```

```
## Warning: Some predictor variables are on very different scales: consider
## rescaling
```

```
## Analysis of Deviance Table (Type II Wald chisquare tests)
## 
## Response: GrowthIncrement_2023
##                                            Chisq Df Pr(>Chisq)    
## plasticity_rdpi_licor_Fs                  1.9987  1     0.1574    
## garden_MAP_2023                           0.9679  1     0.3252    
## Pt                                       32.8887  1  9.759e-09 ***
## plasticity_rdpi_licor_Fs:garden_MAP_2023  0.2996  1     0.5841    
## ---
## Signif. codes:  0 '***' 0.001 '**' 0.01 '*' 0.05 '.' 0.1 ' ' 1
```

```
## Warning: Some predictor variables are on very different scales: consider
## rescaling
```

```
## Analysis of Deviance Table (Type II Wald chisquare tests)
## 
## Response: GrowthIncrement_2023
##                                             Chisq Df Pr(>Chisq)    
## plasticity_rdpi_licor_Fm.                  0.1260  1     0.7226    
## garden_MAP_2023                            0.9761  1     0.3232    
## Pt                                        36.3768  1  1.626e-09 ***
## plasticity_rdpi_licor_Fm.:garden_MAP_2023  0.6335  1     0.4261    
## ---
## Signif. codes:  0 '***' 0.001 '**' 0.01 '*' 0.05 '.' 0.1 ' ' 1
```

```
## Warning: Some predictor variables are on very different scales: consider
## rescaling
```

```
## Analysis of Deviance Table (Type II Wald chisquare tests)
## 
## Response: GrowthIncrement_2023
##                                                              Chisq Df
## plasticity_rdpi_leaf_thickness_avg_mm_2023                  0.9421  1
## garden_MAP_2023                                             0.9760  1
## Pt                                                         33.1088  1
## plasticity_rdpi_leaf_thickness_avg_mm_2023:garden_MAP_2023  0.3057  1
##                                                            Pr(>Chisq)    
## plasticity_rdpi_leaf_thickness_avg_mm_2023                     0.3317    
## garden_MAP_2023                                                0.3232    
## Pt                                                          8.714e-09 ***
## plasticity_rdpi_leaf_thickness_avg_mm_2023:garden_MAP_2023     0.5803    
## ---
## Signif. codes:  0 '***' 0.001 '**' 0.01 '*' 0.05 '.' 0.1 ' ' 1
```

```
## Warning: Some predictor variables are on very different scales: consider
## rescaling
```

```
## Analysis of Deviance Table (Type II Wald chisquare tests)
## 
## Response: GrowthIncrement_2023
##                                                 Chisq Df Pr(>Chisq)    
## plasticity_rdpi_LMA_g_m2_2023                  0.4483  1     0.5032    
## garden_MAP_2023                                0.9762  1     0.3231    
## Pt                                            30.8288  1  2.818e-08 ***
## plasticity_rdpi_LMA_g_m2_2023:garden_MAP_2023  1.2015  1     0.2730    
## ---
## Signif. codes:  0 '***' 0.001 '**' 0.01 '*' 0.05 '.' 0.1 ' ' 1
```

```
## Warning: Some predictor variables are on very different scales: consider
## rescaling
```

```
## Analysis of Deviance Table (Type II Wald chisquare tests)
## 
## Response: GrowthIncrement_2023
##                                                        Chisq Df Pr(>Chisq)    
## plasticity_rdpi_leaf_mass_g_2023_log                  1.7924  1     0.1806    
## garden_MAP_2023                                       0.9626  1     0.3265    
## Pt                                                   39.3618  1  3.521e-10 ***
## plasticity_rdpi_leaf_mass_g_2023_log:garden_MAP_2023  0.7071  1     0.4004    
## ---
## Signif. codes:  0 '***' 0.001 '**' 0.01 '*' 0.05 '.' 0.1 ' ' 1
```

```
## Warning: Some predictor variables are on very different scales: consider
## rescaling
```

```
## Analysis of Deviance Table (Type II Wald chisquare tests)
## 
## Response: GrowthIncrement_2023
##                                                          Chisq Df Pr(>Chisq)
## plasticity_rdpi_leaf_area_cm2_2023_log                  8.4815  1   0.003588
## garden_MAP_2023                                         0.9587  1   0.327519
## Pt                                                     42.8839  1  5.809e-11
## plasticity_rdpi_leaf_area_cm2_2023_log:garden_MAP_2023  4.6673  1   0.030743
##                                                           
## plasticity_rdpi_leaf_area_cm2_2023_log                 ** 
## garden_MAP_2023                                           
## Pt                                                     ***
## plasticity_rdpi_leaf_area_cm2_2023_log:garden_MAP_2023 *  
## ---
## Signif. codes:  0 '***' 0.001 '**' 0.01 '*' 0.05 '.' 0.1 ' ' 1
```

```
## boundary (singular) fit: see help('isSingular')
## boundary (singular) fit: see help('isSingular')
## boundary (singular) fit: see help('isSingular')
## boundary (singular) fit: see help('isSingular')
## boundary (singular) fit: see help('isSingular')
```

```
## `geom_smooth()` using formula = 'y ~ x'
```

```
## Warning: Some predictor variables are on very different scales: consider
## rescaling
```

```
## Analysis of Deviance Table (Type II Wald chisquare tests)
## 
## Response: GrowthIncrement_2023
##                                                                     Chisq Df
## plasticity_rdpi_lower_stomata_pore_length_mean_um                  2.1219  1
## garden_MAP_2023                                                    0.9749  1
## Pt                                                                39.5414  1
## plasticity_rdpi_lower_stomata_pore_length_mean_um:garden_MAP_2023  0.0015  1
##                                                                   Pr(>Chisq)
## plasticity_rdpi_lower_stomata_pore_length_mean_um                     0.1452
## garden_MAP_2023                                                       0.3235
## Pt                                                                 3.212e-10
## plasticity_rdpi_lower_stomata_pore_length_mean_um:garden_MAP_2023     0.9688
##                                                                      
## plasticity_rdpi_lower_stomata_pore_length_mean_um                    
## garden_MAP_2023                                                      
## Pt                                                                ***
## plasticity_rdpi_lower_stomata_pore_length_mean_um:garden_MAP_2023    
## ---
## Signif. codes:  0 '***' 0.001 '**' 0.01 '*' 0.05 '.' 0.1 ' ' 1
```

```
## Warning: Some predictor variables are on very different scales: consider
## rescaling
```

```
## Analysis of Deviance Table (Type II Wald chisquare tests)
## 
## Response: GrowthIncrement_2023
##                                                          Chisq Df Pr(>Chisq)
## plasticity_rdpi_upper_stomata_presence                  2.2119  1     0.1369
## garden_MAP_2023                                         0.9749  1     0.3235
## Pt                                                     40.6112  1  1.857e-10
## plasticity_rdpi_upper_stomata_presence:garden_MAP_2023  0.7176  1     0.3969
##                                                           
## plasticity_rdpi_upper_stomata_presence                    
## garden_MAP_2023                                           
## Pt                                                     ***
## plasticity_rdpi_upper_stomata_presence:garden_MAP_2023    
## ---
## Signif. codes:  0 '***' 0.001 '**' 0.01 '*' 0.05 '.' 0.1 ' ' 1
```

```
## Warning: Some predictor variables are on very different scales: consider
## rescaling
```

```
## Analysis of Deviance Table (Type II Wald chisquare tests)
## 
## Response: GrowthIncrement_2023
##                                                                     Chisq Df
## plasticity_rdpi_upper_stomata_pore_length_mean_um                  0.0213  1
## garden_MAP_2023                                                    0.9220  1
## Pt                                                                29.9325  1
## plasticity_rdpi_upper_stomata_pore_length_mean_um:garden_MAP_2023  0.0186  1
##                                                                   Pr(>Chisq)
## plasticity_rdpi_upper_stomata_pore_length_mean_um                     0.8839
## garden_MAP_2023                                                       0.3369
## Pt                                                                 4.473e-08
## plasticity_rdpi_upper_stomata_pore_length_mean_um:garden_MAP_2023     0.8915
##                                                                      
## plasticity_rdpi_upper_stomata_pore_length_mean_um                    
## garden_MAP_2023                                                      
## Pt                                                                ***
## plasticity_rdpi_upper_stomata_pore_length_mean_um:garden_MAP_2023    
## ---
## Signif. codes:  0 '***' 0.001 '**' 0.01 '*' 0.05 '.' 0.1 ' ' 1
```

```
## Warning: Some predictor variables are on very different scales: consider
## rescaling
```

```
## Analysis of Deviance Table (Type II Wald chisquare tests)
## 
## Response: GrowthIncrement_2023
##                                                                 Chisq Df
## plasticity_rdpi_upper_stomata_density_mm2_log                  0.0577  1
## garden_MAP_2023                                                0.9713  1
## Pt                                                            36.7658  1
## plasticity_rdpi_upper_stomata_density_mm2_log:garden_MAP_2023  0.6516  1
##                                                               Pr(>Chisq)    
## plasticity_rdpi_upper_stomata_density_mm2_log                     0.8102    
## garden_MAP_2023                                                   0.3243    
## Pt                                                             1.332e-09 ***
## plasticity_rdpi_upper_stomata_density_mm2_log:garden_MAP_2023     0.4195    
## ---
## Signif. codes:  0 '***' 0.001 '**' 0.01 '*' 0.05 '.' 0.1 ' ' 1
```

```
## Warning: Some predictor variables are on very different scales: consider
## rescaling
```

```
## Analysis of Deviance Table (Type II Wald chisquare tests)
## 
## Response: GrowthIncrement_2023
##                                                             Chisq Df Pr(>Chisq)
## plasticity_rdpi_lower_stomata_density_mm2                  6.8955  1   0.008641
## garden_MAP_2023                                            0.9745  1   0.323563
## Pt                                                        44.1905  1  2.979e-11
## plasticity_rdpi_lower_stomata_density_mm2:garden_MAP_2023  1.4818  1   0.223498
##                                                              
## plasticity_rdpi_lower_stomata_density_mm2                 ** 
## garden_MAP_2023                                              
## Pt                                                        ***
## plasticity_rdpi_lower_stomata_density_mm2:garden_MAP_2023    
## ---
## Signif. codes:  0 '***' 0.001 '**' 0.01 '*' 0.05 '.' 0.1 ' ' 1
```

```
## boundary (singular) fit: see help('isSingular')
## boundary (singular) fit: see help('isSingular')
## boundary (singular) fit: see help('isSingular')
## boundary (singular) fit: see help('isSingular')
## boundary (singular) fit: see help('isSingular')
## boundary (singular) fit: see help('isSingular')
```

```
## `geom_smooth()` using formula = 'y ~ x'
```

```
## Warning: Some predictor variables are on very different scales: consider
## rescaling
```

```
## Analysis of Deviance Table (Type II Wald chisquare tests)
## 
## Response: GrowthIncrement_2023
##                                                     Chisq Df Pr(>Chisq)    
## plasticity_rdpi_stomata_ratio_log                  1.6334  1     0.2012    
## garden_MAP_2023                                    0.9674  1     0.3253    
## Pt                                                38.0592  1  6.863e-10 ***
## plasticity_rdpi_stomata_ratio_log:garden_MAP_2023  2.5257  1     0.1120    
## ---
## Signif. codes:  0 '***' 0.001 '**' 0.01 '*' 0.05 '.' 0.1 ' ' 1
```

```
plots.summ <- plots.summ[lengths(plots.summ)>0]

# add legend to beginning of plot list

plots2 <- plots.summ
plots2[[length(plots.summ)+1]] <- legend

p3 <- grid.arrange(grobs = plots2, ncol = 4)
```

```
## `geom_smooth()` using formula = 'y ~ x'
```

```
## `geom_smooth()` using formula = 'y ~ x'
## `geom_smooth()` using formula = 'y ~ x'
```

```
plot(p3)

#ggsave(file = 'results/fitness/fitnessEffects_summaryPlot_traitPlasticityxMAP_2023.png', p3, height = 6, width = 14)
#ggsave(file = 'results/fitness/fitnessEffects_summaryPlot_traitPlasticityxMAP_2023.pdf', p3, height = 6, width = 14)

# rename plots list for later use
plots.plast.map.23 <- plots.summ
```

##### 3.1.3 Garden TD

```
################################################
# with TD

# reorder gardens by TD
tmp <- aggregate(dat$garden_TD_2023, by = list(dat$MiniCG_Site_2023), mean)
ord <- tmp$Group.1[order(tmp$x, decreasing = T)]
dat$MiniCG_Site_2023 <- factor(dat$MiniCG_Site_2023, levels = ord)

sites <- levels(dat$MiniCG_Site_2023)
# remove UCM - height not measured
sites <- sites[! sites == 'UCM']

sites.td <- dat$garden_TD_2023[match(sites, dat$MiniCG_Site)]


# to save results
mods.td <- list()

# list to save summary plot for each trait with significant effects on fitness
plots.summ <- list()

for(n in 1:length(traits)){
  
  trait <- traits[n]
  
  mod <- lmer(paste0('GrowthIncrement_2023 ~', trait,  '* garden_TD_2023 + Pt + (1 | MiniCG_Site/block) + (1 | Genotype)'),
              data = dat
  )
  
  print(car::Anova(mod))
  
  p1 <- plot_model(mod, vline.color = "black", show.values = T, type = 'std', 
                   title = paste('2023 Growth ~ ', trait, sep = ''))
  plot(p1)
  #plot_model(mod, type = 're')
  
  mods.td[[n]] <- mod
  names(mods.td)[n] <- trait
  
  #####################
  # posthoc tests
  # if either the climate or trait x climate are significant, run posthoc tests for each garden
  mod.info <- Anova(mod)
  # if any coefficients are significant, plot by garden
  # need to change indexes if model is changed!
  if(mod.info$`Pr(>Chisq)`[1] <= 0.05 | mod.info$`Pr(>Chisq)`[2] <= 0.05 | mod.info$`Pr(>Chisq)`[4] <=0.05){
    
# list to save plots for each site
    plots.sub <- list()
    
    
    # dataframe to save outputs from posthoc tests
    df.posthoc <- as.data.frame(matrix(ncol = 3, nrow = length(sites)))
    rownames(df.posthoc) <- sites
    colnames(df.posthoc) <- c('td', 'slope', 'pval')
    df.posthoc$td <- sites.td
    
    for(s in 1:length(sites)){
      
      site <- sites[s]
      dat.sub <- subset(dat, MiniCG_Site_2023 == site)
      # skip if site is missing data
      if(sum(!is.na(dat.sub[,trait])) > 10){
        mod.sub <- lmer(paste0('GrowthIncrement_2023 ~', trait, '+ Pt + (1 | block) + (1 | Genotype)'),
                        data = dat.sub)
        # get pval for trait
        mod.sub.info <- Anova(mod.sub)
        df.posthoc[site, 'pval'] <- mod.sub.info$`Pr(>Chisq)`[1]
        
        # get slopes from fixed effect of trait (effect on growth)
        df.posthoc[site, 'slope'] <- fixef(mod.sub)[2]
        
        # make title pretty
        maintitle <- paste(site, '\n', trait_names_df[trait, 'trait_names'], '\n', 'p = ', round(df.posthoc$pval[s], 3), sep = '')
        maintitle <- paste(strwrap(maintitle, width = 26), collapse = '\n') 
        
        # plot
        p <- ggplot(data = dat.sub, aes_string(y = 'GrowthIncrement_2023', x = trait, color = 'k2_tricho', na.rm  = TRUE)) +
          geom_point(na.rm = TRUE) +
          scale_color_gradient2(high = "darkolivegreen2", mid = "grey20", low = "dodgerblue2", midpoint = 0.5, name = 'P. trichocarpa\nancestry', guide = 'none') +
          ggtitle(maintitle) +
          geom_smooth(color = 'black', lty = ifelse(df.posthoc$pval[s] <= 0.05, 1,2), method = 'lm', formula = y ~x, na.rm = TRUE) +
          theme(axis.text.x = element_text(angle = 90, vjust = 0.5, hjust=1, size = 12)) +
          theme(axis.title.x = element_blank())
        
        #plot(p)
        plots.sub[[s]] <- p
        
      }
    }

    # pvals are adjusted for summary plot
    df.posthoc$pvals.adj <- p.adjust(df.posthoc$pval, method = 'BH') 
    
    # plot panel with relationship for each garden
    # remove empty plots
    plots.sub <- plots.sub[lengths(plots.sub)>0]
    # plot in grid
    p1 <- grid.arrange(grobs = plots.sub, ncol = 4)
    #plot(p1)
    
    # setup text showing significance
    sig_text <- paste0('Plasticity ', pval_stars(mod.info[1, 'Pr(>Chisq)']), '\n',
                       'Climate ', pval_stars(mod.info[2, 'Pr(>Chisq)']), '\n',
                       'Plasticity x Climate ', pval_stars(mod.info[4, 'Pr(>Chisq)']))
    
    # wrap title
    maintitle <- trait_names_df[trait, 'trait_names']
    maintitle <- paste(strwrap(maintitle, width = 26), collapse = '\n')
    
    # plot slope of fitness relationship vs climate
    p2 <- ggplot(df.posthoc, aes(x = td, y = slope), na.rm  = TRUE) +
      geom_smooth(method = 'lm', col = 'black', na.rm = TRUE) +
      geom_hline(yintercept = 0, lty = 2) +
      geom_point(size = 5, stroke = 2, shape = ifelse(df.posthoc$pvals.adj <= 0.05, 16, 1), aes(col = td), na.rm = TRUE) +
      scale_color_gradient2("orchid4",  mid = 'grey', low = "palegreen4", midpoint = mean(range(sites.td)), limits= range_td, name = 'Garden TD (°C)') +
      ggtitle(maintitle,
              subtitle = sig_text) +
      xlab('Garden TD') +
      ylab('Effect on Growth Increment') +
      geom_text_repel(label = rownames(df.posthoc), aes(x = td, y = slope), box.padding = 0.5, min.segment.length = 1, size = 4) +
        theme(plot.title = element_text(size = 15),
        plot.subtitle = element_text(size = 10),
        legend.position = 'none')
    plot(p2)
    
    plots.summ[[n]] <- p2
    names(plots.summ)[n] <- trait
    
    # add legend
    leg_plot <- ggplot(df.posthoc, aes(x = td, y = slope), na.rm  = TRUE) +
      geom_point(aes(color = td))+
      scale_color_gradient2("orchid4",  mid = 'grey', low = "palegreen4", midpoint = mean(range(sites.td)), limits= range_td, name = 'Garden TD (°C)') +
      theme(legend.position = 'right')
    
    legend <- cowplot::get_legend(leg_plot)
    
  }
}
```

```
## Analysis of Deviance Table (Type II Wald chisquare tests)
## 
## Response: GrowthIncrement_2023
##                                                           Chisq Df Pr(>Chisq)
## plasticity_rdpi_GrowthIncrement_2023_log                 0.5617  1     0.4536
## garden_TD_2023                                           0.0301  1     0.8622
## Pt                                                      32.8478  1  9.967e-09
## plasticity_rdpi_GrowthIncrement_2023_log:garden_TD_2023  0.3819  1     0.5366
##                                                            
## plasticity_rdpi_GrowthIncrement_2023_log                   
## garden_TD_2023                                             
## Pt                                                      ***
## plasticity_rdpi_GrowthIncrement_2023_log:garden_TD_2023    
## ---
## Signif. codes:  0 '***' 0.001 '**' 0.01 '*' 0.05 '.' 0.1 ' ' 1
```

```
## Analysis of Deviance Table (Type II Wald chisquare tests)
## 
## Response: GrowthIncrement_2023
##                                                  Chisq Df Pr(>Chisq)    
## plasticity_rdpi_DOY_Stage2_2023                 0.1776  1     0.6734    
## garden_TD_2023                                  0.0304  1     0.8616    
## Pt                                             37.3926  1  9.659e-10 ***
## plasticity_rdpi_DOY_Stage2_2023:garden_TD_2023  0.1835  1     0.6683    
## ---
## Signif. codes:  0 '***' 0.001 '**' 0.01 '*' 0.05 '.' 0.1 ' ' 1
```

```
## Analysis of Deviance Table (Type II Wald chisquare tests)
## 
## Response: GrowthIncrement_2023
##                                                  Chisq Df Pr(>Chisq)    
## plasticity_rdpi_DOY_Stage3_2023                 0.5744  1     0.4485    
## garden_TD_2023                                  0.0306  1     0.8611    
## Pt                                             37.5024  1   9.13e-10 ***
## plasticity_rdpi_DOY_Stage3_2023:garden_TD_2023  0.1743  1     0.6763    
## ---
## Signif. codes:  0 '***' 0.001 '**' 0.01 '*' 0.05 '.' 0.1 ' ' 1
```

```
## Analysis of Deviance Table (Type II Wald chisquare tests)
## 
## Response: GrowthIncrement_2023
##                                                  Chisq Df Pr(>Chisq)    
## plasticity_rdpi_DOY_Stage6_2023                 0.0441  1    0.83368    
## garden_TD_2023                                  0.0306  1    0.86120    
## Pt                                             29.3909  1  5.915e-08 ***
## plasticity_rdpi_DOY_Stage6_2023:garden_TD_2023  3.1520  1    0.07583 .  
## ---
## Signif. codes:  0 '***' 0.001 '**' 0.01 '*' 0.05 '.' 0.1 ' ' 1
```

```
## Analysis of Deviance Table (Type II Wald chisquare tests)
## 
## Response: GrowthIncrement_2023
##                                                       Chisq Df Pr(>Chisq)    
## plasticity_rdpi_DOY_last_budset_2023                 1.8974  1     0.1684    
## garden_TD_2023                                       0.0299  1     0.8626    
## Pt                                                  38.3495  1  5.914e-10 ***
## plasticity_rdpi_DOY_last_budset_2023:garden_TD_2023  0.0304  1     0.8617    
## ---
## Signif. codes:  0 '***' 0.001 '**' 0.01 '*' 0.05 '.' 0.1 ' ' 1
```

```
## Analysis of Deviance Table (Type II Wald chisquare tests)
## 
## Response: GrowthIncrement_2023
##                                                       Chisq Df Pr(>Chisq)    
## plasticity_rdpi_stage7_presence_2023                 0.0071  1     0.9328    
## garden_TD_2023                                       0.0305  1     0.8614    
## Pt                                                  36.5313  1  1.502e-09 ***
## plasticity_rdpi_stage7_presence_2023:garden_TD_2023  0.2054  1     0.6504    
## ---
## Signif. codes:  0 '***' 0.001 '**' 0.01 '*' 0.05 '.' 0.1 ' ' 1
```

```
## Analysis of Deviance Table (Type II Wald chisquare tests)
## 
## Response: GrowthIncrement_2023
##                                                           Chisq Df Pr(>Chisq)
## plasticity_rdpi_growing_season_days_2023                 2.3755  1     0.1233
## garden_TD_2023                                           0.0310  1     0.8603
## Pt                                                      34.8915  1  3.486e-09
## plasticity_rdpi_growing_season_days_2023:garden_TD_2023  1.0787  1     0.2990
##                                                            
## plasticity_rdpi_growing_season_days_2023                   
## garden_TD_2023                                             
## Pt                                                      ***
## plasticity_rdpi_growing_season_days_2023:garden_TD_2023    
## ---
## Signif. codes:  0 '***' 0.001 '**' 0.01 '*' 0.05 '.' 0.1 ' ' 1
```

```
## Analysis of Deviance Table (Type II Wald chisquare tests)
## 
## Response: GrowthIncrement_2023
##                                                  Chisq Df Pr(>Chisq)    
## plasticity_rdpi_DOY_Stage7_2023                 0.5495  1     0.4585    
## garden_TD_2023                                  0.0687  1     0.7932    
## Pt                                             22.8531  1  1.749e-06 ***
## plasticity_rdpi_DOY_Stage7_2023:garden_TD_2023  0.5712  1     0.4498    
## ---
## Signif. codes:  0 '***' 0.001 '**' 0.01 '*' 0.05 '.' 0.1 ' ' 1
```

```
## Analysis of Deviance Table (Type II Wald chisquare tests)
## 
## Response: GrowthIncrement_2023
##                                            Chisq Df Pr(>Chisq)    
## plasticity_rdpi_licor_gsw                 0.3605  1     0.5482    
## garden_TD_2023                            0.0306  1     0.8610    
## Pt                                       31.0210  1  2.552e-08 ***
## plasticity_rdpi_licor_gsw:garden_TD_2023  1.7079  1     0.1913    
## ---
## Signif. codes:  0 '***' 0.001 '**' 0.01 '*' 0.05 '.' 0.1 ' ' 1
```

```
## Warning: Some predictor variables are on very different scales: consider
## rescaling
```

```
## Analysis of Deviance Table (Type II Wald chisquare tests)
## 
## Response: GrowthIncrement_2023
##                                            Chisq Df Pr(>Chisq)    
## plasticity_rdpi_licor_gbw                 0.2315  1     0.6304    
## garden_TD_2023                            0.0303  1     0.8618    
## Pt                                       36.5481  1  1.489e-09 ***
## plasticity_rdpi_licor_gbw:garden_TD_2023  1.4447  1     0.2294    
## ---
## Signif. codes:  0 '***' 0.001 '**' 0.01 '*' 0.05 '.' 0.1 ' ' 1
```

```
## Analysis of Deviance Table (Type II Wald chisquare tests)
## 
## Response: GrowthIncrement_2023
##                                               Chisq Df Pr(>Chisq)    
## plasticity_rdpi_licor_PhiPS2                 2.0071  1     0.1566    
## garden_TD_2023                               0.0302  1     0.8620    
## Pt                                          38.4810  1  5.529e-10 ***
## plasticity_rdpi_licor_PhiPS2:garden_TD_2023  0.7884  1     0.3746    
## ---
## Signif. codes:  0 '***' 0.001 '**' 0.01 '*' 0.05 '.' 0.1 ' ' 1
```

```
## Analysis of Deviance Table (Type II Wald chisquare tests)
## 
## Response: GrowthIncrement_2023
##                                            Chisq Df Pr(>Chisq)    
## plasticity_rdpi_licor_ETR                 0.2437  1     0.6216    
## garden_TD_2023                            0.0300  1     0.8625    
## Pt                                       37.3587  1  9.828e-10 ***
## plasticity_rdpi_licor_ETR:garden_TD_2023  0.7132  1     0.3984    
## ---
## Signif. codes:  0 '***' 0.001 '**' 0.01 '*' 0.05 '.' 0.1 ' ' 1
```

```
## Analysis of Deviance Table (Type II Wald chisquare tests)
## 
## Response: GrowthIncrement_2023
##                                           Chisq Df Pr(>Chisq)    
## plasticity_rdpi_licor_Fs                 1.9883  1    0.15852    
## garden_TD_2023                           0.0301  1    0.86236    
## Pt                                      32.8876  1  9.765e-09 ***
## plasticity_rdpi_licor_Fs:garden_TD_2023  3.7175  1    0.05384 .  
## ---
## Signif. codes:  0 '***' 0.001 '**' 0.01 '*' 0.05 '.' 0.1 ' ' 1
```

```
## Analysis of Deviance Table (Type II Wald chisquare tests)
## 
## Response: GrowthIncrement_2023
##                                            Chisq Df Pr(>Chisq)    
## plasticity_rdpi_licor_Fm.                 0.1239  1     0.7248    
## garden_TD_2023                            0.0303  1     0.8618    
## Pt                                       36.3714  1  1.631e-09 ***
## plasticity_rdpi_licor_Fm.:garden_TD_2023  0.1449  1     0.7035    
## ---
## Signif. codes:  0 '***' 0.001 '**' 0.01 '*' 0.05 '.' 0.1 ' ' 1
```

```
## Analysis of Deviance Table (Type II Wald chisquare tests)
## 
## Response: GrowthIncrement_2023
##                                                             Chisq Df Pr(>Chisq)
## plasticity_rdpi_leaf_thickness_avg_mm_2023                 0.9363  1     0.3332
## garden_TD_2023                                             0.0300  1     0.8625
## Pt                                                        32.9646  1  9.385e-09
## plasticity_rdpi_leaf_thickness_avg_mm_2023:garden_TD_2023  2.6233  1     0.1053
##                                                              
## plasticity_rdpi_leaf_thickness_avg_mm_2023                   
## garden_TD_2023                                               
## Pt                                                        ***
## plasticity_rdpi_leaf_thickness_avg_mm_2023:garden_TD_2023    
## ---
## Signif. codes:  0 '***' 0.001 '**' 0.01 '*' 0.05 '.' 0.1 ' ' 1
```

```
## Analysis of Deviance Table (Type II Wald chisquare tests)
## 
## Response: GrowthIncrement_2023
##                                                Chisq Df Pr(>Chisq)    
## plasticity_rdpi_LMA_g_m2_2023                 0.4461  1     0.5042    
## garden_TD_2023                                0.0301  1     0.8622    
## Pt                                           30.6680  1  3.062e-08 ***
## plasticity_rdpi_LMA_g_m2_2023:garden_TD_2023  0.5272  1     0.4678    
## ---
## Signif. codes:  0 '***' 0.001 '**' 0.01 '*' 0.05 '.' 0.1 ' ' 1
```

```
## Analysis of Deviance Table (Type II Wald chisquare tests)
## 
## Response: GrowthIncrement_2023
##                                                       Chisq Df Pr(>Chisq)    
## plasticity_rdpi_leaf_mass_g_2023_log                 1.7985  1     0.1799    
## garden_TD_2023                                       0.0302  1     0.8620    
## Pt                                                  39.7050  1  2.954e-10 ***
## plasticity_rdpi_leaf_mass_g_2023_log:garden_TD_2023  1.3757  1     0.2408    
## ---
## Signif. codes:  0 '***' 0.001 '**' 0.01 '*' 0.05 '.' 0.1 ' ' 1
```

```
## Analysis of Deviance Table (Type II Wald chisquare tests)
## 
## Response: GrowthIncrement_2023
##                                                         Chisq Df Pr(>Chisq)    
## plasticity_rdpi_leaf_area_cm2_2023_log                 8.4386  1   0.003673 ** 
## garden_TD_2023                                         0.0290  1   0.864838    
## Pt                                                    43.0225  1  5.411e-11 ***
## plasticity_rdpi_leaf_area_cm2_2023_log:garden_TD_2023  0.9007  1   0.342602    
## ---
## Signif. codes:  0 '***' 0.001 '**' 0.01 '*' 0.05 '.' 0.1 ' ' 1
```

```
## boundary (singular) fit: see help('isSingular')
## boundary (singular) fit: see help('isSingular')
## boundary (singular) fit: see help('isSingular')
## boundary (singular) fit: see help('isSingular')
## boundary (singular) fit: see help('isSingular')
```

```
## `geom_smooth()` using formula = 'y ~ x'
```

```
## Analysis of Deviance Table (Type II Wald chisquare tests)
## 
## Response: GrowthIncrement_2023
##                                                                    Chisq Df
## plasticity_rdpi_lower_stomata_pore_length_mean_um                 2.1235  1
## garden_TD_2023                                                    0.0321  1
## Pt                                                               39.4880  1
## plasticity_rdpi_lower_stomata_pore_length_mean_um:garden_TD_2023  0.0312  1
##                                                                  Pr(>Chisq)    
## plasticity_rdpi_lower_stomata_pore_length_mean_um                    0.1451    
## garden_TD_2023                                                       0.8579    
## Pt                                                                3.301e-10 ***
## plasticity_rdpi_lower_stomata_pore_length_mean_um:garden_TD_2023     0.8598    
## ---
## Signif. codes:  0 '***' 0.001 '**' 0.01 '*' 0.05 '.' 0.1 ' ' 1
```

```
## Analysis of Deviance Table (Type II Wald chisquare tests)
## 
## Response: GrowthIncrement_2023
##                                                         Chisq Df Pr(>Chisq)    
## plasticity_rdpi_upper_stomata_presence                 2.1784  1     0.1400    
## garden_TD_2023                                         0.0328  1     0.8562    
## Pt                                                    40.1005  1  2.412e-10 ***
## plasticity_rdpi_upper_stomata_presence:garden_TD_2023  0.4255  1     0.5142    
## ---
## Signif. codes:  0 '***' 0.001 '**' 0.01 '*' 0.05 '.' 0.1 ' ' 1
```

```
## Analysis of Deviance Table (Type II Wald chisquare tests)
## 
## Response: GrowthIncrement_2023
##                                                                    Chisq Df
## plasticity_rdpi_upper_stomata_pore_length_mean_um                 0.0201  1
## garden_TD_2023                                                    0.0332  1
## Pt                                                               30.0126  1
## plasticity_rdpi_upper_stomata_pore_length_mean_um:garden_TD_2023  0.5080  1
##                                                                  Pr(>Chisq)    
## plasticity_rdpi_upper_stomata_pore_length_mean_um                    0.8873    
## garden_TD_2023                                                       0.8553    
## Pt                                                                4.293e-08 ***
## plasticity_rdpi_upper_stomata_pore_length_mean_um:garden_TD_2023     0.4760    
## ---
## Signif. codes:  0 '***' 0.001 '**' 0.01 '*' 0.05 '.' 0.1 ' ' 1
```

```
## Warning: Some predictor variables are on very different scales: consider
## rescaling
```

```
## Analysis of Deviance Table (Type II Wald chisquare tests)
## 
## Response: GrowthIncrement_2023
##                                                                Chisq Df
## plasticity_rdpi_upper_stomata_density_mm2_log                 0.0588  1
## garden_TD_2023                                                0.0305  1
## Pt                                                           36.8385  1
## plasticity_rdpi_upper_stomata_density_mm2_log:garden_TD_2023  0.6452  1
##                                                              Pr(>Chisq)    
## plasticity_rdpi_upper_stomata_density_mm2_log                    0.8083    
## garden_TD_2023                                                   0.8615    
## Pt                                                            1.283e-09 ***
## plasticity_rdpi_upper_stomata_density_mm2_log:garden_TD_2023     0.4218    
## ---
## Signif. codes:  0 '***' 0.001 '**' 0.01 '*' 0.05 '.' 0.1 ' ' 1
```

```
## Analysis of Deviance Table (Type II Wald chisquare tests)
## 
## Response: GrowthIncrement_2023
##                                                            Chisq Df Pr(>Chisq)
## plasticity_rdpi_lower_stomata_density_mm2                 6.9193  1   0.008527
## garden_TD_2023                                            0.0299  1   0.862768
## Pt                                                       44.4178  1  2.653e-11
## plasticity_rdpi_lower_stomata_density_mm2:garden_TD_2023  1.1091  1   0.292269
##                                                             
## plasticity_rdpi_lower_stomata_density_mm2                ** 
## garden_TD_2023                                              
## Pt                                                       ***
## plasticity_rdpi_lower_stomata_density_mm2:garden_TD_2023    
## ---
## Signif. codes:  0 '***' 0.001 '**' 0.01 '*' 0.05 '.' 0.1 ' ' 1
```

```
## boundary (singular) fit: see help('isSingular')
## boundary (singular) fit: see help('isSingular')
## boundary (singular) fit: see help('isSingular')
## boundary (singular) fit: see help('isSingular')
## boundary (singular) fit: see help('isSingular')
## boundary (singular) fit: see help('isSingular')
```

```
## `geom_smooth()` using formula = 'y ~ x'
```

```
## Analysis of Deviance Table (Type II Wald chisquare tests)
## 
## Response: GrowthIncrement_2023
##                                                    Chisq Df Pr(>Chisq)    
## plasticity_rdpi_stomata_ratio_log                 1.6249  1    0.20241    
## garden_TD_2023                                    0.0297  1    0.86306    
## Pt                                               37.8086  1  7.804e-10 ***
## plasticity_rdpi_stomata_ratio_log:garden_TD_2023  2.9584  1    0.08543 .  
## ---
## Signif. codes:  0 '***' 0.001 '**' 0.01 '*' 0.05 '.' 0.1 ' ' 1
```

```
plots.summ <- plots.summ[lengths(plots.summ)>0]

# add legend to beginning of plot list

plots2 <- plots.summ
plots2[[length(plots.summ)+1]] <- legend

p3 <- grid.arrange(grobs = plots2, ncol = 3)
```

```
## `geom_smooth()` using formula = 'y ~ x'
```

```
## `geom_smooth()` using formula = 'y ~ x'
```

```
plot(p3)

#ggsave(file = 'results/fitness/fitnessEffects_summaryPlot_traitPlasticityxTD_2023.png', p3, height = 6, width = 14)
#ggsave(file = 'results/fitness/fitnessEffects_summaryPlot_traitPlasticityxTD_2023.pdf', p3, height = 6, width = 14)

# rename plots list for later use
plots.plast.td.23 <- plots.summ
```

#### 3.2 Does trait plasticity (RDPI) predict growth in 2024?

```
# merge plasticity values with main df

# get plasticity traits
plast <- grep('rdpi', colnames(rdpi), value = T)
plast <- grep('2024', plast, value = T)

# add empty columns
dat[,plast] <- NA

# fill in plasticity columns for each genotype
for(n in 1:nrow(rdpi)){
  
  geno <- rownames(rdpi)[n]
  dat[dat$Genotype == geno, plast] <- rdpi[geno,plast]
  
}

traits <- plast

# does plasticity in these traits predict growth increment?
```

##### 3.2.1 Garden MAT

```
# reorder gardens by MAT
tmp <- aggregate(dat$garden_MAT_2024, by = list(dat$MiniCG_Site_2024), mean)
ord <- tmp$Group.1[order(tmp$x, decreasing = T)]
dat$MiniCG_Site_2024 <- factor(dat$MiniCG_Site_2024, levels = ord)

# sites <- levels(dat$MiniCG_Site_2024)
sites <- c("WI", "ID", "WYO", "EVERGREEN", "MSU", "OSU", "MORTON", "WSU", "PENN", "VA", "SU")

sites.mat <- dat$garden_MAT_2024[match(sites, dat$MiniCG_Site)]

#sites <- levels(dat$MiniCG_Site_2024)
# remove UCM - height not measured
#sites <- sites[! sites == 'UCM']

# to save results
mods.mat <- list()

# list to save summary plot for each trait with significant effects on fitness
plots.summ <- list()

for(n in 1:length(traits)){
  
  trait <- traits[n]
  
  mod <- lmer(paste0('GrowthIncrement_2024 ~', trait,  '* garden_MAT_2024 + Pt + (1 | MiniCG_Site/block) + (1 | Genotype)'),
              data = dat
  )
  
  print(car::Anova(mod))
  
  p1 <- plot_model(mod, vline.color = "black", show.values = T, type = 'std', 
                   title = paste('2024 Growth ~ ', trait, sep = ''))
  plot(p1)
  #plot_model(mod, type = 're')
  
  mods.mat[[n]] <- mod
  names(mods.mat)[n] <- trait
  
  #####################
  # posthoc tests
  # if either the climate or trait x climate are significant, run posthoc tests for each garden
  mod.info <- Anova(mod)
  # if any coefficients are significant, plot by garden
  # need to change indexes if model is changed!
  if(mod.info$`Pr(>Chisq)`[1] <= 0.05 | mod.info$`Pr(>Chisq)`[2] <= 0.05 | mod.info$`Pr(>Chisq)`[4] <=0.05){
    
# list to save plots for each site
    plots.sub <- list()
    
    # dataframe to save outputs from posthoc tests
    df.posthoc <- as.data.frame(matrix(ncol = 3, nrow = length(sites)))
    rownames(df.posthoc) <- sites
    colnames(df.posthoc) <- c('mat', 'slope', 'pval')
    df.posthoc$mat <- sites.mat
    
    for(s in 1:length(sites)){
      
      site <- sites[s]
      dat.sub <- subset(dat, MiniCG_Site_2024 == site)
      # skip if site is missing data
      if(sum(!is.na(dat.sub[,trait])) > 10){
        
        mod.sub <- lmer(paste0('GrowthIncrement_2024 ~', trait, '+ Pt + (1 | block) + (1 | Genotype)'),
                        data = dat.sub)
        # get pval for trait
        mod.sub.info <- Anova(mod.sub)
        df.posthoc[site, 'pval'] <- mod.sub.info$`Pr(>Chisq)`[1]
        
        # get slopes from fixed effect of trait (effect on growth)
        df.posthoc[site, 'slope'] <- fixef(mod.sub)[2]
        
        # make title pretty
        maintitle <- paste(site, '\n', trait_names_df[trait, 'trait_names'], '\n', 'p = ', round(df.posthoc$pval[s], 3), sep = '')
        maintitle <- paste(strwrap(maintitle, width = 26), collapse = '\n') 
        
        # plot
        p <- ggplot(data = dat.sub, aes_string(y = 'GrowthIncrement_2024', x = trait, color = 'k2_tricho'), na.rm  = TRUE) +
          geom_point(na.rm = TRUE) +
          scale_color_gradient2(high = "darkolivegreen2", mid = "grey20", low = "dodgerblue2", midpoint = 0.5, name = 'P. trichocarpa\nancestry', guide = 'none') +
          ggtitle(maintitle) +
          geom_smooth(color = 'black', lty = ifelse(df.posthoc$pval[s] <= 0.05, 1,2), method = 'lm', formula = y ~ x, na.rm = TRUE) +
          theme(axis.text.x = element_text(angle = 90, vjust = 0.5, hjust=1, size = 12)) +
          theme(axis.title.x = element_blank())
        
        #plot(p)
        plots.sub[[s]] <- p
        
      }
    }

    # pvals are adjusted for summary plot
    df.posthoc$pvals.adj <- p.adjust(df.posthoc$pval, method = 'BH') 
    
    # plot panel with relationship for each garden
    # remove empty plots
    plots.sub <- plots.sub[lengths(plots.sub)>0]
    # plot in grid
    p1 <- grid.arrange(grobs = plots.sub, ncol = 4)
    #plot(p1)
    
    # setup text showing significance
    sig_text <- paste0('Plasticity ', pval_stars(mod.info[1, 'Pr(>Chisq)']), '\n',
                       'Climate ', pval_stars(mod.info[2, 'Pr(>Chisq)']), '\n',
                       'Plasticity x Climate ', pval_stars(mod.info[4, 'Pr(>Chisq)']))
    
    # wrap title
    maintitle <- trait_names_df[trait, 'trait_names']
    maintitle <- paste(strwrap(maintitle, width = 26), collapse = '\n')
    
    # plot slope of fitness relationship vs climate
    p2 <- ggplot(df.posthoc, aes(x = mat, y = slope), na.rm  = TRUE) +
      geom_smooth(method = 'lm', col = 'black', na.rm = TRUE) +
      geom_hline(yintercept = 0, lty = 2) +
      geom_point(size = 5, stroke = 2, shape = ifelse(df.posthoc$pvals.adj <= 0.05, 16, 1), aes(col = mat), na.rm = TRUE) +
      scale_color_gradient2(high = "red2", mid = "grey", low = "blue2", midpoint = mean(range(sites.mat)), limits = range_mat, name = 'Garden MAT (°C)') +
      ggtitle(maintitle,
              subtitle = sig_text) +
      xlab('Garden MAT') +
      ylab('Effect on Growth Increment') +
      geom_text_repel(label = rownames(df.posthoc), aes(x = mat, y = slope), box.padding = 0.5, min.segment.length = 1, size = 4) +
      theme(plot.title = element_text(size = 15),
            plot.subtitle = element_text(size = 10),
            legend.position = 'none')
    plot(p2)
    
    plots.summ[[n]] <- p2
    names(plots.summ)[n] <- trait
    
     # add legend
    leg_plot <- ggplot(df.posthoc, aes(x = mat, y = slope), na.rm  = TRUE) +
      geom_point(aes(color = mat))+
      scale_color_gradient2(high = "red2", mid = "grey", low = "blue2", midpoint = mean(range(sites.mat)), limits = range_mat, name = 'Garden MAT (°C)') +
      theme(legend.position = 'right')
    
    legend <- cowplot::get_legend(leg_plot)
    
  }
}
```

```
## Analysis of Deviance Table (Type II Wald chisquare tests)
## 
## Response: GrowthIncrement_2024
##                                                   Chisq Df Pr(>Chisq)    
## plasticity_rdpi_DOY_Stage2_2024                  1.8084  1    0.17870    
## garden_MAT_2024                                  4.4714  1    0.03447 *  
## Pt                                              31.8320  1  1.681e-08 ***
## plasticity_rdpi_DOY_Stage2_2024:garden_MAT_2024  2.8393  1    0.09199 .  
## ---
## Signif. codes:  0 '***' 0.001 '**' 0.01 '*' 0.05 '.' 0.1 ' ' 1
```

```
## boundary (singular) fit: see help('isSingular')
## boundary (singular) fit: see help('isSingular')
## boundary (singular) fit: see help('isSingular')
## boundary (singular) fit: see help('isSingular')
## boundary (singular) fit: see help('isSingular')
## boundary (singular) fit: see help('isSingular')
```

```
## `geom_smooth()` using formula = 'y ~ x'
```

```
## Warning: Removed 3 rows containing missing values or values outside the scale range
## (`geom_text_repel()`).
```

```
## Warning: Removed 3 rows containing missing values or values outside the scale range
## (`geom_point()`).
```

```
## Analysis of Deviance Table (Type II Wald chisquare tests)
## 
## Response: GrowthIncrement_2024
##                                                   Chisq Df Pr(>Chisq)    
## plasticity_rdpi_DOY_Stage3_2024                  4.6957  1    0.03024 *  
## garden_MAT_2024                                  4.4461  1    0.03498 *  
## Pt                                              37.9147  1  7.391e-10 ***
## plasticity_rdpi_DOY_Stage3_2024:garden_MAT_2024  4.3688  1    0.03660 *  
## ---
## Signif. codes:  0 '***' 0.001 '**' 0.01 '*' 0.05 '.' 0.1 ' ' 1
```

```
## boundary (singular) fit: see help('isSingular')
## boundary (singular) fit: see help('isSingular')
## boundary (singular) fit: see help('isSingular')
## boundary (singular) fit: see help('isSingular')
## boundary (singular) fit: see help('isSingular')
## boundary (singular) fit: see help('isSingular')
```

```
## `geom_smooth()` using formula = 'y ~ x'
```

```
## Warning: Removed 3 rows containing missing values or values outside the scale range
## (`geom_text_repel()`).
## Removed 3 rows containing missing values or values outside the scale range
## (`geom_point()`).
```

```
## Analysis of Deviance Table (Type II Wald chisquare tests)
## 
## Response: GrowthIncrement_2024
##                                                   Chisq Df Pr(>Chisq)    
## plasticity_rdpi_DOY_Stage6_2024                  0.0009  1    0.97659    
## garden_MAT_2024                                  4.4024  1    0.03589 *  
## Pt                                              31.7931  1  1.715e-08 ***
## plasticity_rdpi_DOY_Stage6_2024:garden_MAT_2024  0.0978  1    0.75447    
## ---
## Signif. codes:  0 '***' 0.001 '**' 0.01 '*' 0.05 '.' 0.1 ' ' 1
```

```
## boundary (singular) fit: see help('isSingular')
## boundary (singular) fit: see help('isSingular')
## boundary (singular) fit: see help('isSingular')
## boundary (singular) fit: see help('isSingular')
## boundary (singular) fit: see help('isSingular')
```

```
## `geom_smooth()` using formula = 'y ~ x'
```

```
## Warning: Removed 3 rows containing missing values or values outside the scale range
## (`geom_text_repel()`).
## Removed 3 rows containing missing values or values outside the scale range
## (`geom_point()`).
```

```
## boundary (singular) fit: see help('isSingular')
```

```
## Analysis of Deviance Table (Type II Wald chisquare tests)
## 
## Response: GrowthIncrement_2024
##                                                  Chisq Df Pr(>Chisq)   
## plasticity_rdpi_DOY_Stage7_2024                 0.4207  1    0.51661   
## garden_MAT_2024                                 6.9419  1    0.00842 **
## Pt                                              3.8122  1    0.05088 . 
## plasticity_rdpi_DOY_Stage7_2024:garden_MAT_2024 0.1770  1    0.67393   
## ---
## Signif. codes:  0 '***' 0.001 '**' 0.01 '*' 0.05 '.' 0.1 ' ' 1
```

```
## boundary (singular) fit: see help('isSingular')
```

```
## boundary (singular) fit: see help('isSingular')
## boundary (singular) fit: see help('isSingular')
## boundary (singular) fit: see help('isSingular')
```

```
## `geom_smooth()` using formula = 'y ~ x'
```

```
## Warning: Removed 3 rows containing missing values or values outside the scale range
## (`geom_text_repel()`).
## Removed 3 rows containing missing values or values outside the scale range
## (`geom_point()`).
```

```
## Analysis of Deviance Table (Type II Wald chisquare tests)
## 
## Response: GrowthIncrement_2024
##                                                        Chisq Df Pr(>Chisq)    
## plasticity_rdpi_DOY_last_budset_2024                  0.0082  1    0.92784    
## garden_MAT_2024                                       4.4339  1    0.03523 *  
## Pt                                                   28.7977  1  8.035e-08 ***
## plasticity_rdpi_DOY_last_budset_2024:garden_MAT_2024  0.2284  1    0.63270    
## ---
## Signif. codes:  0 '***' 0.001 '**' 0.01 '*' 0.05 '.' 0.1 ' ' 1
```

```
## boundary (singular) fit: see help('isSingular')
## boundary (singular) fit: see help('isSingular')
## boundary (singular) fit: see help('isSingular')
## boundary (singular) fit: see help('isSingular')
## boundary (singular) fit: see help('isSingular')
```

```
## `geom_smooth()` using formula = 'y ~ x'
```

```
## Warning: Removed 3 rows containing missing values or values outside the scale range
## (`geom_text_repel()`).
## Removed 3 rows containing missing values or values outside the scale range
## (`geom_point()`).
```

```
## Analysis of Deviance Table (Type II Wald chisquare tests)
## 
## Response: GrowthIncrement_2024
##                                                        Chisq Df Pr(>Chisq)    
## plasticity_rdpi_stage7_presence_2024                  1.2954  1  0.2550582    
## garden_MAT_2024                                       4.7517  1  0.0292693 *  
## Pt                                                   16.7084  1  4.359e-05 ***
## plasticity_rdpi_stage7_presence_2024:garden_MAT_2024 11.8461  1  0.0005778 ***
## ---
## Signif. codes:  0 '***' 0.001 '**' 0.01 '*' 0.05 '.' 0.1 ' ' 1
```

```
## boundary (singular) fit: see help('isSingular')
## boundary (singular) fit: see help('isSingular')
## boundary (singular) fit: see help('isSingular')
## boundary (singular) fit: see help('isSingular')
## boundary (singular) fit: see help('isSingular')
```

```
## `geom_smooth()` using formula = 'y ~ x'
```

```
## Warning: Removed 3 rows containing missing values or values outside the scale range
## (`geom_text_repel()`).
## Removed 3 rows containing missing values or values outside the scale range
## (`geom_point()`).
```

```
## Analysis of Deviance Table (Type II Wald chisquare tests)
## 
## Response: GrowthIncrement_2024
##                                                            Chisq Df Pr(>Chisq)
## plasticity_rdpi_growing_season_days_2024                  0.0135  1    0.90745
## garden_MAT_2024                                           2.9226  1    0.08734
## Pt                                                       30.4145  1  3.489e-08
## plasticity_rdpi_growing_season_days_2024:garden_MAT_2024  3.6634  1    0.05562
##                                                             
## plasticity_rdpi_growing_season_days_2024                    
## garden_MAT_2024                                          .  
## Pt                                                       ***
## plasticity_rdpi_growing_season_days_2024:garden_MAT_2024 .  
## ---
## Signif. codes:  0 '***' 0.001 '**' 0.01 '*' 0.05 '.' 0.1 ' ' 1
```

```
plots.summ <- plots.summ[lengths(plots.summ)>0]

# add legend to beginning of plot list
plots.summ[[length(plots.summ)+1]] <- legend

p3 <- grid.arrange(grobs = plots.summ, ncol = 4)
```

```
## `geom_smooth()` using formula = 'y ~ x'
```

```
## Warning: Removed 3 rows containing missing values or values outside the scale range
## (`geom_text_repel()`).
```

```
## `geom_smooth()` using formula = 'y ~ x'
```

```
## Warning: Removed 3 rows containing missing values or values outside the scale range
## (`geom_text_repel()`).
```

```
## `geom_smooth()` using formula = 'y ~ x'
```

```
## Warning: Removed 3 rows containing missing values or values outside the scale range
## (`geom_text_repel()`).
```

```
## `geom_smooth()` using formula = 'y ~ x'
```

```
## Warning: Removed 3 rows containing missing values or values outside the scale range
## (`geom_text_repel()`).
```

```
## `geom_smooth()` using formula = 'y ~ x'
```

```
## Warning: Removed 3 rows containing missing values or values outside the scale range
## (`geom_text_repel()`).
```

```
## `geom_smooth()` using formula = 'y ~ x'
```

```
## Warning: Removed 3 rows containing missing values or values outside the scale range
## (`geom_text_repel()`).
```

```
plot(p3)


#ggsave(file = 'results/fitness/fitnessEffects_summaryPlot_traitPlasticityxMAT_2024.png', p3, height = 8, width = 14)
#ggsave(file = 'results/fitness/fitnessEffects_summaryPlot_traitPlasticityxMAT_2024.pdf', p3, height = 8, width = 14)

# rename plots list for later use
plots.plast.mat.24 <- plots.summ
```

##### 3.2.2 Garden MAP

```
################################################
# with MAP

# reorder gardens by MAP
tmp <- aggregate(dat$garden_MAP_2024, by = list(dat$MiniCG_Site_2024), mean)
ord <- tmp$Group.1[order(tmp$x, decreasing = T)]
dat$MiniCG_Site_2024 <- factor(dat$MiniCG_Site_2024, levels = ord)

sites <- c("WI", "ID", "WYO", "EVERGREEN", "MSU", "OSU", "MORTON", "WSU", "PENN", "VA", "SU")

#sites <- levels(dat$MiniCG_Site_2024)
# remove UCM - height not measured
#sites <- sites[! sites == 'UCM']

sites.map <- dat$garden_MAP_2024[match(sites, dat$MiniCG_Site)]


# to save results
mods.map <- list()

# list to save summary plot for each trait with significant effects on fitness
plots.summ <- list()

for(n in 1:length(traits)){
  
  trait <- traits[n]
  
  mod <- lmer(paste0('GrowthIncrement_2024 ~', trait,  '* garden_MAP_2024 + Pt + (1 | MiniCG_Site/block) + (1 | Genotype)'),
              data = dat
  )
  
  print(car::Anova(mod))
  
  p1 <- plot_model(mod, vline.color = "black", show.values = T, type = 'std', 
                   title = paste('2024 Growth ~ ', trait, sep = ''))
  plot(p1)
  #plot_model(mod, type = 're')
  
  mods.map[[n]] <- mod
  names(mods.map)[n] <- trait
  
  #####################
  # posthoc tests
  # if either the climate or trait x climate are significant, run posthoc tests for each garden
  mod.info <- Anova(mod)
  # if any coefficients are significant, plot by garden
  # need to change indexes if model is changed!
  if(mod.info$`Pr(>Chisq)`[1] <= 0.05 | mod.info$`Pr(>Chisq)`[2] <= 0.05 | mod.info$`Pr(>Chisq)`[4] <=0.05){
    
# list to save plots for each site
    plots.sub <- list()
    
    # dataframe to save outputs from posthoc tests
    df.posthoc <- as.data.frame(matrix(ncol = 3, nrow = length(sites)))
    rownames(df.posthoc) <- sites
    colnames(df.posthoc) <- c('map', 'slope', 'pval')
    df.posthoc$map <- sites.map
    
    for(s in 1:length(sites)){
      
      site <- sites[s]
      dat.sub <- subset(dat, MiniCG_Site_2024 == site)
      # skip if site is missing data
      if(sum(!is.na(dat.sub[,trait])) > 10){
        mod.sub <- lmer(paste0('GrowthIncrement_2024 ~', trait, '+ Pt + (1 | block) + (1 | Genotype)'),
                        data = dat.sub)
        # get pval for trait
        mod.sub.info <- Anova(mod.sub)
        df.posthoc[site, 'pval'] <- mod.sub.info$`Pr(>Chisq)`[1]
        
        # get slopes from fixed effect of trait (effect on growth)
        df.posthoc[site, 'slope'] <- fixef(mod.sub)[2]
        
        # make title pretty
        maintitle <- paste(site, '\n', trait_names_df[trait, 'trait_names'], '\n', 'p = ', round(df.posthoc$pval[s], 3), sep = '')
        maintitle <- paste(strwrap(maintitle, width = 26), collapse = '\n')  
        
        # plot
        p <- ggplot(data = dat.sub, aes_string(y = 'GrowthIncrement_2024', x = trait, color = 'k2_tricho'), na.rm  = TRUE) +
          geom_point(na.rm = TRUE) +
          scale_color_gradient2(high = "darkolivegreen2", mid = "grey20", low = "dodgerblue2", midpoint = 0.5, name = 'P. trichocarpa\nancestry', guide = 'none') +
          ggtitle(maintitle) +
          geom_smooth(color = 'black', lty = ifelse(df.posthoc$pval[s] <= 0.05, 1,2), method = 'lm', formula = y ~ x, na.rm = TRUE) +
          theme(axis.text.x = element_text(angle = 90, vjust = 0.5, hjust=1, size = 12)) +
          theme(axis.title.x = element_blank())
        
        #plot(p)
        plots.sub[[s]] <- p
        
      }
    }

    # pvals are adjusted for summary plot
    df.posthoc$pvals.adj <- p.adjust(df.posthoc$pval, method = 'BH') 
    
    # plot panel with relationship for each garden
    # remove empty plots
    plots.sub <- plots.sub[lengths(plots.sub)>0]
    # plot in grid
    p1 <- grid.arrange(grobs = plots.sub, ncol = 4)
    #plot(p1)
    
    # setup text showing significance
    sig_text <- paste0('Plasticity ', pval_stars(mod.info[1, 'Pr(>Chisq)']), '\n',
                       'Climate ', pval_stars(mod.info[2, 'Pr(>Chisq)']), '\n',
                       'Plasticity x Climate ', pval_stars(mod.info[4, 'Pr(>Chisq)']))
    
    # wrap title
    maintitle <- trait_names_df[trait, 'trait_names']
    maintitle <- paste(strwrap(maintitle, width = 26), collapse = '\n')
    
    # plot slope of fitness relationship vs climate
    p2 <- ggplot(df.posthoc, aes(x = map, y = slope), na.rm  = TRUE) +
      geom_smooth(method = 'lm', col = 'black', na.rm = TRUE) +
      geom_hline(yintercept = 0, lty = 2) +
      geom_point(size = 5, stroke = 2, shape = ifelse(df.posthoc$pvals.adj <= 0.05, 16, 1), aes(col = map), na.rm = TRUE) +
      scale_color_gradient2(high = "steelblue2",  mid = 'grey', low = "sienna3", midpoint = mean(range(sites.map)), limits = range_map, name = 'Garden MAP (mm)') +
      ggtitle(maintitle,
              subtitle = sig_text) +
      xlab('Garden MAP') +
      ylab('Effect on Growth Increment') +
      geom_text_repel(label = rownames(df.posthoc), aes(x = map, y = slope), box.padding = 0.5, min.segment.length = 1, size = 4) +
        theme(plot.title = element_text(size = 15),
        plot.subtitle = element_text(size = 10),
        legend.position = 'none')
    plot(p2)
    
    plots.summ[[n]] <- p2
    names(plots.summ)[n] <- trait
    
     # add legend
    leg_plot <- ggplot(df.posthoc, aes(x = map, y = slope), na.rm  = TRUE) +
      geom_point(aes(color = map))+
      scale_color_gradient2(high = "steelblue2",  mid = 'grey', low = "sienna3", midpoint = mean(range(sites.map)), limits = range_map, name = 'Garden MAP (mm)') +
      theme(legend.position = 'right')
    
    legend <- cowplot::get_legend(leg_plot)
  }
}
```

```
## Warning: Some predictor variables are on very different scales: consider
## rescaling
```

```
## Analysis of Deviance Table (Type II Wald chisquare tests)
## 
## Response: GrowthIncrement_2024
##                                                   Chisq Df Pr(>Chisq)    
## plasticity_rdpi_DOY_Stage2_2024                  1.8705  1     0.1714    
## garden_MAP_2024                                  0.9975  1     0.3179    
## Pt                                              33.0875  1   8.81e-09 ***
## plasticity_rdpi_DOY_Stage2_2024:garden_MAP_2024  0.1067  1     0.7439    
## ---
## Signif. codes:  0 '***' 0.001 '**' 0.01 '*' 0.05 '.' 0.1 ' ' 1
```

```
## Warning: Some predictor variables are on very different scales: consider
## rescaling
```

```
## Analysis of Deviance Table (Type II Wald chisquare tests)
## 
## Response: GrowthIncrement_2024
##                                                   Chisq Df Pr(>Chisq)    
## plasticity_rdpi_DOY_Stage3_2024                  4.8509  1    0.02763 *  
## garden_MAP_2024                                  0.9727  1    0.32401    
## Pt                                              38.5745  1   5.27e-10 ***
## plasticity_rdpi_DOY_Stage3_2024:garden_MAP_2024  0.1649  1    0.68466    
## ---
## Signif. codes:  0 '***' 0.001 '**' 0.01 '*' 0.05 '.' 0.1 ' ' 1
```

```
## boundary (singular) fit: see help('isSingular')
## boundary (singular) fit: see help('isSingular')
## boundary (singular) fit: see help('isSingular')
## boundary (singular) fit: see help('isSingular')
## boundary (singular) fit: see help('isSingular')
## boundary (singular) fit: see help('isSingular')
```

```
## `geom_smooth()` using formula = 'y ~ x'
```

```
## Warning: Removed 3 rows containing missing values or values outside the scale range
## (`geom_text_repel()`).
```

```
## Warning: Removed 3 rows containing missing values or values outside the scale range
## (`geom_point()`).
```

```
## Warning: Some predictor variables are on very different scales: consider
## rescaling
```

```
## Analysis of Deviance Table (Type II Wald chisquare tests)
## 
## Response: GrowthIncrement_2024
##                                                   Chisq Df Pr(>Chisq)    
## plasticity_rdpi_DOY_Stage6_2024                  0.0012  1     0.9722    
## garden_MAP_2024                                  0.9828  1     0.3215    
## Pt                                              32.0612  1  1.494e-08 ***
## plasticity_rdpi_DOY_Stage6_2024:garden_MAP_2024  0.8863  1     0.3465    
## ---
## Signif. codes:  0 '***' 0.001 '**' 0.01 '*' 0.05 '.' 0.1 ' ' 1
```

```
## Warning: Some predictor variables are on very different scales: consider
## rescaling
```

```
## Analysis of Deviance Table (Type II Wald chisquare tests)
## 
## Response: GrowthIncrement_2024
##                                                  Chisq Df Pr(>Chisq)  
## plasticity_rdpi_DOY_Stage7_2024                 0.3896  1    0.53253  
## garden_MAP_2024                                 1.9881  1    0.15854  
## Pt                                              4.0986  1    0.04292 *
## plasticity_rdpi_DOY_Stage7_2024:garden_MAP_2024 0.5410  1    0.46203  
## ---
## Signif. codes:  0 '***' 0.001 '**' 0.01 '*' 0.05 '.' 0.1 ' ' 1
```

```
## Warning: Some predictor variables are on very different scales: consider
## rescaling
```

```
## Analysis of Deviance Table (Type II Wald chisquare tests)
## 
## Response: GrowthIncrement_2024
##                                                        Chisq Df Pr(>Chisq)    
## plasticity_rdpi_DOY_last_budset_2024                  0.0035  1     0.9532    
## garden_MAP_2024                                       0.9739  1     0.3237    
## Pt                                                   29.3453  1  6.056e-08 ***
## plasticity_rdpi_DOY_last_budset_2024:garden_MAP_2024  0.7913  1     0.3737    
## ---
## Signif. codes:  0 '***' 0.001 '**' 0.01 '*' 0.05 '.' 0.1 ' ' 1
```

```
## Warning: Some predictor variables are on very different scales: consider
## rescaling
```

```
## Analysis of Deviance Table (Type II Wald chisquare tests)
## 
## Response: GrowthIncrement_2024
##                                                        Chisq Df Pr(>Chisq)    
## plasticity_rdpi_stage7_presence_2024                  1.2880  1    0.25642    
## garden_MAP_2024                                       0.9981  1    0.31777    
## Pt                                                   17.3657  1  3.083e-05 ***
## plasticity_rdpi_stage7_presence_2024:garden_MAP_2024  6.6251  1    0.01005 *  
## ---
## Signif. codes:  0 '***' 0.001 '**' 0.01 '*' 0.05 '.' 0.1 ' ' 1
```

```
## boundary (singular) fit: see help('isSingular')
## boundary (singular) fit: see help('isSingular')
## boundary (singular) fit: see help('isSingular')
## boundary (singular) fit: see help('isSingular')
## boundary (singular) fit: see help('isSingular')
```

```
## `geom_smooth()` using formula = 'y ~ x'
```

```
## Warning: Removed 3 rows containing missing values or values outside the scale range
## (`geom_text_repel()`).
## Removed 3 rows containing missing values or values outside the scale range
## (`geom_point()`).
```

```
## Warning: Some predictor variables are on very different scales: consider
## rescaling
```

```
## Analysis of Deviance Table (Type II Wald chisquare tests)
## 
## Response: GrowthIncrement_2024
##                                                            Chisq Df Pr(>Chisq)
## plasticity_rdpi_growing_season_days_2024                  0.0045  1     0.9463
## garden_MAP_2024                                           0.9242  1     0.3364
## Pt                                                       31.7306  1  1.771e-08
## plasticity_rdpi_growing_season_days_2024:garden_MAP_2024  2.3634  1     0.1242
##                                                             
## plasticity_rdpi_growing_season_days_2024                    
## garden_MAP_2024                                             
## Pt                                                       ***
## plasticity_rdpi_growing_season_days_2024:garden_MAP_2024    
## ---
## Signif. codes:  0 '***' 0.001 '**' 0.01 '*' 0.05 '.' 0.1 ' ' 1
```

```
plots.summ <- plots.summ[lengths(plots.summ)>0]

# add legend to beginning of plot list

plots.summ[[length(plots.summ)+1]] <- legend

p3 <- grid.arrange(grobs = plots.summ, ncol = 3)
```

```
## `geom_smooth()` using formula = 'y ~ x'
```

```
## Warning: Removed 3 rows containing missing values or values outside the scale range
## (`geom_text_repel()`).
```

```
## `geom_smooth()` using formula = 'y ~ x'
```

```
## Warning: Removed 3 rows containing missing values or values outside the scale range
## (`geom_text_repel()`).
```

```
plot(p3)

#ggsave(file = 'results/fitness/fitnessEffects_summaryPlot_traitPlasticityxMAP_2024.png', p3, height = 6, width = 10)
#ggsave(file = 'results/fitness/fitnessEffects_summaryPlot_traitPlasticityxMAP_2024.pdf', p3, height = 6, width = 10)

# rename plots list for later use
plots.plast.map.24 <- plots.summ
```

##### 3.2.3 Garden TD

```
################################################
# with TD

# reorder gardens by TD
tmp <- aggregate(dat$garden_TD_2024, by = list(dat$MiniCG_Site_2024), mean)
ord <- tmp$Group.1[order(tmp$x, decreasing = T)]
dat$MiniCG_Site_2024 <- factor(dat$MiniCG_Site_2024, levels = ord)

sites <- c("WI", "ID", "WYO", "EVERGREEN", "MSU", "OSU", "MORTON", "WSU", "PENN", "VA", "SU")

#sites <- levels(dat$MiniCG_Site_2024)
# remove UCM - height not measured
#sites <- sites[! sites == 'UCM']

sites.td <- dat$garden_TD_2024[match(sites, dat$MiniCG_Site)]

# to save results
mods.td <- list()

# list to save summary plot for each trait with significant effects on fitness
plots.summ <- list()


for(n in 1:length(traits)){
  
  trait <- traits[n]
  
  mod <- lmer(paste0('GrowthIncrement_2024 ~', trait,  '* garden_TD_2024 + Pt + (1 | MiniCG_Site/block) + (1 | Genotype)'),
              data = dat
  )
  
  print(car::Anova(mod))
  
  p1 <- plot_model(mod, vline.color = "black", show.values = T, type = 'std', 
                   title = paste('2024 Growth ~ ', trait, sep = ''))
  plot(p1)
  #plot_model(mod, type = 're')
  
  mods.td[[n]] <- mod
  names(mods.td)[n] <- trait
  
  #####################
  # posthoc tests
  
  # if either the climate or trait x climate are significant, run posthoc tests for each garden
  mod.info <- Anova(mod)
  # if any coefficients are significant, plot by garden
  # need to change indexes if model is changed!
  if(mod.info$`Pr(>Chisq)`[1] <= 0.05 | mod.info$`Pr(>Chisq)`[2] <= 0.05 | mod.info$`Pr(>Chisq)`[4] <=0.05){
    
    
# list to save plots for each site
    plots.sub <- list()
    
    # dataframe to save outputs from posthoc tests
    df.posthoc <- as.data.frame(matrix(ncol = 3, nrow = length(sites)))
    rownames(df.posthoc) <- sites
    colnames(df.posthoc) <- c('td', 'slope', 'pval')
    df.posthoc$td <- sites.td
    
    for(s in 1:length(sites)){
      
      site <- sites[s]
      dat.sub <- subset(dat, MiniCG_Site_2024 == site)
      # skip if site is missing data
      if(sum(!is.na(dat.sub[,trait])) > 10){
        mod.sub <- lmer(paste0('GrowthIncrement_2024 ~', trait, '+ Pt + (1 | block) + (1 | Genotype)'),
                        data = dat.sub)
        # get pval for trait
        mod.sub.info <- Anova(mod.sub)
        df.posthoc[site, 'pval'] <- mod.sub.info$`Pr(>Chisq)`[1]
        
        # get slopes from fixed effect of trait (effect on growth)
        df.posthoc[site, 'slope'] <- fixef(mod.sub)[2]
        
        # make title pretty
        maintitle <- paste(site, '\n', trait_names_df[trait, 'trait_names'], '\n', 'p = ', round(df.posthoc$pval[s], 3), sep = '')
        maintitle <- paste(strwrap(maintitle, width = 26), collapse = '\n')  
        
        # plot
        p <- ggplot(data = dat.sub, aes_string(y = 'GrowthIncrement_2024', x = trait, color = 'k2_tricho'), na.rm  = TRUE) +
          geom_point(na.rm = TRUE) +
          scale_color_gradient2(high = "darkolivegreen2", mid = "grey20", low = "dodgerblue2", midpoint = 0.5, name = 'P. trichocarpa\nancestry', guide = 'none') +
          ggtitle(maintitle) +
          geom_smooth(color = 'black', lty = ifelse(df.posthoc$pval[s] <= 0.05, 1,2), method = 'lm', formula = y ~ x, na.rm = TRUE) +
          theme(axis.text.x = element_text(angle = 90, vjust = 0.5, hjust=1, size = 12)) +
          theme(axis.title.x = element_blank())
        
        #plot(p)
        plots.sub[[s]] <- p
        
      }
    }

    # pvals are adjusted for summary plot
    df.posthoc$pvals.adj <- p.adjust(df.posthoc$pval, method = 'BH') 
    
    # plot panel with relationship for each garden
    # remove empty plots
    plots.sub <- plots.sub[lengths(plots.sub)>0]
    # plot in grid
    p1 <- grid.arrange(grobs = plots.sub, ncol = 4)
    #plot(p1)
    
    # setup text showing significance
    sig_text <- paste0('Plasticity ', pval_stars(mod.info[1, 'Pr(>Chisq)']), '\n',
                       'Climate ', pval_stars(mod.info[2, 'Pr(>Chisq)']), '\n',
                       'Plasticity x Climate ', pval_stars(mod.info[4, 'Pr(>Chisq)']))
    
    # wrap title
    maintitle <- trait_names_df[trait, 'trait_names']
    maintitle <- paste(strwrap(maintitle, width = 26), collapse = '\n')
    
    
    # plot slope of fitness relationship vs climate
    p2 <- ggplot(df.posthoc, aes(x = td, y = slope), na.rm  = TRUE) +
            geom_smooth(method = 'lm', col = 'black', na.rm = TRUE) +
      geom_hline(yintercept = 0, lty = 2) +
      geom_point(size = 5, stroke = 2, shape = ifelse(df.posthoc$pvals.adj <= 0.05, 16, 1), aes(col = td), na.rm = TRUE) +
      scale_color_gradient2("orchid4",  mid = 'grey', low = "palegreen4", midpoint = mean(range(sites.td)), limits = range_td, name = 'Garden TD (°C)') +
      ggtitle(maintitle,
              subtitle = sig_text) +
      xlab('Garden TD') +
      ylab('Effect on Growth Increment') +
      geom_text_repel(label = rownames(df.posthoc), aes(x = td, y = slope), box.padding = 0.5, min.segment.length = 1, size = 4) +
        theme(plot.title = element_text(size = 15),
        plot.subtitle = element_text(size = 10),
        legend.position = 'none'
)
    
    plot(p2)
    
    plots.summ[[n]] <- p2
    names(plots.summ)[n] <- trait

    
     # add legend
    leg_plot <- ggplot(df.posthoc, aes(x = td, y = slope), na.rm  = TRUE) +
      geom_point(aes(color = td))+
      scale_color_gradient2("orchid4",  mid = 'grey', low = "palegreen4", midpoint = mean(range(sites.td)), limits = range_td, name = 'Garden TD (°C)') +
      theme(legend.position = 'right')
    
    legend <- cowplot::get_legend(leg_plot)
  }
}
```

```
## Analysis of Deviance Table (Type II Wald chisquare tests)
## 
## Response: GrowthIncrement_2024
##                                                  Chisq Df Pr(>Chisq)    
## plasticity_rdpi_DOY_Stage2_2024                 1.8527  1   0.173471    
## garden_TD_2024                                  0.0290  1   0.864812    
## Pt                                             33.7368  1   6.31e-09 ***
## plasticity_rdpi_DOY_Stage2_2024:garden_TD_2024  7.8332  1   0.005129 ** 
## ---
## Signif. codes:  0 '***' 0.001 '**' 0.01 '*' 0.05 '.' 0.1 ' ' 1
```

```
## boundary (singular) fit: see help('isSingular')
## boundary (singular) fit: see help('isSingular')
## boundary (singular) fit: see help('isSingular')
## boundary (singular) fit: see help('isSingular')
## boundary (singular) fit: see help('isSingular')
## boundary (singular) fit: see help('isSingular')
```

```
## `geom_smooth()` using formula = 'y ~ x'
```

```
## Warning: Removed 3 rows containing missing values or values outside the scale range
## (`geom_text_repel()`).
```

```
## Warning: Removed 3 rows containing missing values or values outside the scale range
## (`geom_point()`).
```

```
## Analysis of Deviance Table (Type II Wald chisquare tests)
## 
## Response: GrowthIncrement_2024
##                                                  Chisq Df Pr(>Chisq)    
## plasticity_rdpi_DOY_Stage3_2024                 4.9339  1    0.02633 *  
## garden_TD_2024                                  0.0189  1    0.89077    
## Pt                                             40.8578  1  1.637e-10 ***
## plasticity_rdpi_DOY_Stage3_2024:garden_TD_2024  5.9577  1    0.01465 *  
## ---
## Signif. codes:  0 '***' 0.001 '**' 0.01 '*' 0.05 '.' 0.1 ' ' 1
```

```
## boundary (singular) fit: see help('isSingular')
## boundary (singular) fit: see help('isSingular')
## boundary (singular) fit: see help('isSingular')
## boundary (singular) fit: see help('isSingular')
## boundary (singular) fit: see help('isSingular')
## boundary (singular) fit: see help('isSingular')
```

```
## `geom_smooth()` using formula = 'y ~ x'
```

```
## Warning: Removed 3 rows containing missing values or values outside the scale range
## (`geom_text_repel()`).
## Removed 3 rows containing missing values or values outside the scale range
## (`geom_point()`).
```

```
## Analysis of Deviance Table (Type II Wald chisquare tests)
## 
## Response: GrowthIncrement_2024
##                                                  Chisq Df Pr(>Chisq)    
## plasticity_rdpi_DOY_Stage6_2024                 0.0011  1     0.9732    
## garden_TD_2024                                  0.0284  1     0.8662    
## Pt                                             32.3959  1  1.257e-08 ***
## plasticity_rdpi_DOY_Stage6_2024:garden_TD_2024  0.0343  1     0.8531    
## ---
## Signif. codes:  0 '***' 0.001 '**' 0.01 '*' 0.05 '.' 0.1 ' ' 1
```

```
## Analysis of Deviance Table (Type II Wald chisquare tests)
## 
## Response: GrowthIncrement_2024
##                                                 Chisq Df Pr(>Chisq)  
## plasticity_rdpi_DOY_Stage7_2024                0.3359  1    0.56222  
## garden_TD_2024                                 0.2401  1    0.62412  
## Pt                                             4.0044  1    0.04538 *
## plasticity_rdpi_DOY_Stage7_2024:garden_TD_2024 0.8204  1    0.36506  
## ---
## Signif. codes:  0 '***' 0.001 '**' 0.01 '*' 0.05 '.' 0.1 ' ' 1
```

```
## Analysis of Deviance Table (Type II Wald chisquare tests)
## 
## Response: GrowthIncrement_2024
##                                                       Chisq Df Pr(>Chisq)    
## plasticity_rdpi_DOY_last_budset_2024                 0.0020  1     0.9645    
## garden_TD_2024                                       0.0285  1     0.8660    
## Pt                                                  29.9667  1  4.395e-08 ***
## plasticity_rdpi_DOY_last_budset_2024:garden_TD_2024  0.0196  1     0.8887    
## ---
## Signif. codes:  0 '***' 0.001 '**' 0.01 '*' 0.05 '.' 0.1 ' ' 1
```

```
## Analysis of Deviance Table (Type II Wald chisquare tests)
## 
## Response: GrowthIncrement_2024
##                                                       Chisq Df Pr(>Chisq)    
## plasticity_rdpi_stage7_presence_2024                 1.2814  1     0.2576    
## garden_TD_2024                                       0.0234  1     0.8785    
## Pt                                                  17.4688  1  2.921e-05 ***
## plasticity_rdpi_stage7_presence_2024:garden_TD_2024  0.0180  1     0.8931    
## ---
## Signif. codes:  0 '***' 0.001 '**' 0.01 '*' 0.05 '.' 0.1 ' ' 1
```

```
## Analysis of Deviance Table (Type II Wald chisquare tests)
## 
## Response: GrowthIncrement_2024
##                                                           Chisq Df Pr(>Chisq)
## plasticity_rdpi_growing_season_days_2024                 0.0018  1     0.9666
## garden_TD_2024                                           0.0002  1     0.9878
## Pt                                                      32.3202  1  1.307e-08
## plasticity_rdpi_growing_season_days_2024:garden_TD_2024  0.0182  1     0.8926
##                                                            
## plasticity_rdpi_growing_season_days_2024                   
## garden_TD_2024                                             
## Pt                                                      ***
## plasticity_rdpi_growing_season_days_2024:garden_TD_2024    
## ---
## Signif. codes:  0 '***' 0.001 '**' 0.01 '*' 0.05 '.' 0.1 ' ' 1
```

```
plots.summ <- plots.summ[lengths(plots.summ)>0]

# add legend to beginning of plot list
plots.summ[[length(plots.summ)+1]] <- legend

p3 <- grid.arrange(grobs = plots.summ, ncol = 3)
```

```
## `geom_smooth()` using formula = 'y ~ x'
```

```
## Warning: Removed 3 rows containing missing values or values outside the scale range
## (`geom_text_repel()`).
```

```
## `geom_smooth()` using formula = 'y ~ x'
```

```
## Warning: Removed 3 rows containing missing values or values outside the scale range
## (`geom_text_repel()`).
```

```
plot(p3)

#ggsave(file = 'results/fitness/fitnessEffects_summaryPlot_traitPlasticityxTD_2024.png', p3, height = 6, width = 10)
#ggsave(file = 'results/fitness/fitnessEffects_summaryPlot_traitPlasticityxTD_2024.pdf', p3, height = 6, width = 10)

# rename plots list for later use
plots.plast.td.24 <- plots.summ
```

#### 3.3 Plasticity plots for both years

```
# merge lists of plots created above

plots.plast.mat <- c(plots.plast.mat.23, plots.plast.mat.24)
plots.plast.map <- c(plots.plast.map.23, plots.plast.map.24)
plots.plast.td <- c(plots.plast.td.23, plots.plast.td.24)


# MAT
pMAT <- grid.arrange(grobs = plots.plast.mat, ncol = 4)
```

```
## `geom_smooth()` using formula = 'y ~ x'
## `geom_smooth()` using formula = 'y ~ x'
## `geom_smooth()` using formula = 'y ~ x'
## `geom_smooth()` using formula = 'y ~ x'
## `geom_smooth()` using formula = 'y ~ x'
## `geom_smooth()` using formula = 'y ~ x'
## `geom_smooth()` using formula = 'y ~ x'
## `geom_smooth()` using formula = 'y ~ x'
## `geom_smooth()` using formula = 'y ~ x'
## `geom_smooth()` using formula = 'y ~ x'
## `geom_smooth()` using formula = 'y ~ x'
## `geom_smooth()` using formula = 'y ~ x'
```

```
## Warning: Removed 3 rows containing missing values or values outside the scale range
## (`geom_text_repel()`).
```

```
## `geom_smooth()` using formula = 'y ~ x'
```

```
## Warning: Removed 3 rows containing missing values or values outside the scale range
## (`geom_text_repel()`).
```

```
## `geom_smooth()` using formula = 'y ~ x'
```

```
## Warning: Removed 3 rows containing missing values or values outside the scale range
## (`geom_text_repel()`).
```

```
## `geom_smooth()` using formula = 'y ~ x'
```

```
## Warning: Removed 3 rows containing missing values or values outside the scale range
## (`geom_text_repel()`).
```

```
## `geom_smooth()` using formula = 'y ~ x'
```

```
## Warning: Removed 3 rows containing missing values or values outside the scale range
## (`geom_text_repel()`).
```

```
## `geom_smooth()` using formula = 'y ~ x'
```

```
## Warning: Removed 3 rows containing missing values or values outside the scale range
## (`geom_text_repel()`).
```

```
## Warning: ggrepel: 2 unlabeled data points (too many overlaps). Consider increasing max.overlaps
## ggrepel: 2 unlabeled data points (too many overlaps). Consider increasing max.overlaps
## ggrepel: 2 unlabeled data points (too many overlaps). Consider increasing max.overlaps
## ggrepel: 2 unlabeled data points (too many overlaps). Consider increasing max.overlaps
```

```
plot(pMAT)
```

```
## Warning: ggrepel: 2 unlabeled data points (too many overlaps). Consider increasing max.overlaps
## ggrepel: 2 unlabeled data points (too many overlaps). Consider increasing max.overlaps
## ggrepel: 2 unlabeled data points (too many overlaps). Consider increasing max.overlaps
## ggrepel: 2 unlabeled data points (too many overlaps). Consider increasing max.overlaps
```

```
#ggsave(file = 'results/fitness/fitnessEffects_summaryPlot_traitPlasticityxMAT.png', pMAT, height = 16, width = 14)
#ggsave(file = 'results/fitness/fitnessEffects_summaryPlot_traitPlasticityxMAT.pdf', pMAT, height = 16, width = 14)

# MAP
pMAP <- grid.arrange(grobs = plots.plast.map, ncol = 3)
```

```
## `geom_smooth()` using formula = 'y ~ x'
## `geom_smooth()` using formula = 'y ~ x'
## `geom_smooth()` using formula = 'y ~ x'
## `geom_smooth()` using formula = 'y ~ x'
```

```
## Warning: Removed 3 rows containing missing values or values outside the scale range
## (`geom_text_repel()`).
```

```
## `geom_smooth()` using formula = 'y ~ x'
```

```
## Warning: Removed 3 rows containing missing values or values outside the scale range
## (`geom_text_repel()`).
```

```
plot(pMAP)
#ggsave(file = 'results/fitness/fitnessEffects_summaryPlot_traitPlasticityxMAP.png', pMAP, height = 12, width = 14)
#ggsave(file = 'results/fitness/fitnessEffects_summaryPlot_traitPlasticityxMAP.pdf', pMAP, height = 12, width = 14)

# TD
pTD <- grid.arrange(grobs = plots.plast.td, ncol = 3)
```

```
## `geom_smooth()` using formula = 'y ~ x'
## `geom_smooth()` using formula = 'y ~ x'
## `geom_smooth()` using formula = 'y ~ x'
```

```
## Warning: Removed 3 rows containing missing values or values outside the scale range
## (`geom_text_repel()`).
```

```
## `geom_smooth()` using formula = 'y ~ x'
```

```
## Warning: Removed 3 rows containing missing values or values outside the scale range
## (`geom_text_repel()`).
```

```
plot(pTD) 
#ggsave(file = 'results/fitness/fitnessEffects_summaryPlot_traitPlasticityxTD.png', pTD, height = 12, width = 14)
#ggsave(file = 'results/fitness/fitnessEffects_summaryPlot_traitPlasticityxTD.pdf', pTD, height = 12, width = 14)
```

```
# look at overlap in significant traits
g1 <- names(plots.trait.mat)
g2 <- names(plots.plast.mat)
g2 <- gsub('plasticity_rdpi_', '', g2 )
g2 <- gsub('_log', '', g2 )

overlap <- g1[g1 %in% g2]

overlap <- c("DOY_Stage6_2023", "licor_PhiPS2", "licor_Fs", 
"LMA_g_m2_2023", "DOY_last_budset_2024")

# subset of traits with significant interactions

panel <- grid.arrange(grobs = list(plots.trait.mat$leaf_thickness_avg_mm_2023,
                                   plots.trait.mat$DOY_Stage6_2023,
                                   plots.trait.mat$licor_PhiPS2, plots.trait.mat.24[[length(plots.trait.mat.24)]],
                                   plots.plast.mat$plasticity_rdpi_leaf_thickness_avg_mm_2023,
                                   plots.plast.mat.23$plasticity_rdpi_DOY_Stage6_2023 ,                                   plots.plast.mat$plasticity_rdpi_licor_PhiPS2), ncol = 4)
```

```
## `geom_smooth()` using formula = 'y ~ x'
```

```
## Warning: Removed 2 rows containing missing values or values outside the scale range
## (`geom_text_repel()`).
```

```
## `geom_smooth()` using formula = 'y ~ x'
```

```
## Warning: Removed 1 row containing missing values or values outside the scale range
## (`geom_text_repel()`).
```

```
## `geom_smooth()` using formula = 'y ~ x'
```

```
## Warning: Removed 3 rows containing missing values or values outside the scale range
## (`geom_text_repel()`).
```

```
## `geom_smooth()` using formula = 'y ~ x'
## `geom_smooth()` using formula = 'y ~ x'
## `geom_smooth()` using formula = 'y ~ x'
```

```
plot(panel)

#ggsave(file = 'results/fitness/fitnessEffectsPanel_MAT_TraitAndPlasticity.png', panel, height = 8, width = 14)
#ggsave(file = 'results/fitness/fitnessEffectsPanel_MAT_TraitAndPlasticity.pdf', panel, height = 8, width = 12)


grid.arrange(grobs = list(plots.trait.mat$LMA_g_m2_2023, plots.plast.mat$plasticity_rdpi_LMA_g_m2_2023), ncol= 2)
```

```
## `geom_smooth()` using formula = 'y ~ x'
```

```
## Warning: Removed 1 row containing missing values or values outside the scale range
## (`geom_text_repel()`).
```

```
## `geom_smooth()` using formula = 'y ~ x'
```

```
grid.arrange(grobs = list(plots.trait.mat$DOY_Stage6_2023, plots.plast.mat$plasticity_rdpi_DOY_Stage6_2023), ncol= 2)
```

```
## `geom_smooth()` using formula = 'y ~ x'
```

```
## Warning: Removed 1 row containing missing values or values outside the scale range
## (`geom_text_repel()`).
```

```
## `geom_smooth()` using formula = 'y ~ x'
```

```
##########################
# MAP
# look at overlap
g1 <- names(plots.trait.map)
g2 <- names(plots.plast.map)
g2 <- gsub('plasticity_rdpi_', '', g2 )
g2 <- gsub('_log', '', g2 )

overlap <- g1[g1 %in% g2]

dput(overlap)
```

```
## c("licor_gbw", "leaf_area_cm2_2023", "lower_stomata_density_mm2", 
## "")
```

```
#p <- grid.arrange(grobs = plots.trait.map[overlap], ncol = 4)

panel <- grid.arrange(grobs = list(plots.trait.map$licor_gbw, plots.trait.map$leaf_area_cm2_2023, plots.trait.map$lower_stomata_density_mm2,             plots.plast.map$plasticity_rdpi_licor_gbw, plots.plast.map$plasticity_rdpi_leaf_area_cm2_2023_log, plots.plast.map$plasticity_rdpi_lower_stomata_density_mm2),
                      ncol = 3)
```

```
## `geom_smooth()` using formula = 'y ~ x'
```

```
## Warning: Removed 3 rows containing missing values or values outside the scale range
## (`geom_text_repel()`).
```

```
## `geom_smooth()` using formula = 'y ~ x'
```

```
## Warning: Removed 1 row containing missing values or values outside the scale range
## (`geom_text_repel()`).
```

```
## `geom_smooth()` using formula = 'y ~ x'
```

```
## Warning: Removed 1 row containing missing values or values outside the scale range
## (`geom_text_repel()`).
```

```
## `geom_smooth()` using formula = 'y ~ x'
## `geom_smooth()` using formula = 'y ~ x'
## `geom_smooth()` using formula = 'y ~ x'
```

```
plot(panel)
#ggsave(file = 'results/fitness/fitnessEffectsPanel_MAP_TraitAndPlasticity.png', panel, height = 8, width = 12)

#ggsave(file = 'results/fitness/fitnessEffectsPanel_MAP_TraitAndPlasticity.pdf', panel, height = 8, width = 12)
```

### 4 Is there a tradeoff between plasticity and traits?

If traits and their plasticity have a significant relationship, there
could be a tradeoff between plasticity and expression of certain trait
values

```
# list of traits

traits <- c("DOY_Stage2_2023", "DOY_Stage3_2023", "DOY_Stage6_2023", "DOY_Stage7_2023",  "DOY_last_budset_2023", "growing_season_days_2023", "licor_gsw", "licor_gbw", "licor_ETR", "licor_PhiPS2", "licor_Fs", "licor_Fm.", "leaf_thickness_avg_mm_2023", "LMA_g_m2_2023", "leaf_mass_g_2023", "leaf_area_cm2_2023", "lower_stomata_pore_length_mean_um", "upper_stomata_presence", "upper_stomata_pore_length_mean_um",  "upper_stomata_density_mm2", "lower_stomata_density_mm2", "stomata_ratio", "DOY_Stage2_2024", "DOY_Stage3_2024", "DOY_Stage6_2024", "DOY_Stage7_2024", "DOY_last_budset_2024", "growing_season_days_2024")

plasts <- paste('plasticity_rdpi_', traits, sep = '')
dput(plasts)
```

```
## c("plasticity_rdpi_DOY_Stage2_2023", "plasticity_rdpi_DOY_Stage3_2023", 
## "plasticity_rdpi_DOY_Stage6_2023", "plasticity_rdpi_DOY_Stage7_2023", 
## "plasticity_rdpi_DOY_last_budset_2023", "plasticity_rdpi_growing_season_days_2023", 
## "plasticity_rdpi_licor_gsw", "plasticity_rdpi_licor_gbw", "plasticity_rdpi_licor_ETR", 
## "plasticity_rdpi_licor_PhiPS2", "plasticity_rdpi_licor_Fs", "plasticity_rdpi_licor_Fm.", 
## "plasticity_rdpi_leaf_thickness_avg_mm_2023", "plasticity_rdpi_LMA_g_m2_2023", 
## "plasticity_rdpi_leaf_mass_g_2023", "plasticity_rdpi_leaf_area_cm2_2023", 
## "plasticity_rdpi_lower_stomata_pore_length_mean_um", "plasticity_rdpi_upper_stomata_presence", 
## "plasticity_rdpi_upper_stomata_pore_length_mean_um", "plasticity_rdpi_upper_stomata_density_mm2", 
## "plasticity_rdpi_lower_stomata_density_mm2", "plasticity_rdpi_stomata_ratio", 
## "plasticity_rdpi_DOY_Stage2_2024", "plasticity_rdpi_DOY_Stage3_2024", 
## "plasticity_rdpi_DOY_Stage6_2024", "plasticity_rdpi_DOY_Stage7_2024", 
## "plasticity_rdpi_DOY_last_budset_2024", "plasticity_rdpi_growing_season_days_2024"
## )
```

```
sum(! plasts %in% colnames(dat))
```

```
## [1] 4
```

```
plasts[! plasts %in% colnames(dat)]
```

```
## [1] "plasticity_rdpi_leaf_mass_g_2023"         
## [2] "plasticity_rdpi_leaf_area_cm2_2023"       
## [3] "plasticity_rdpi_upper_stomata_density_mm2"
## [4] "plasticity_rdpi_stomata_ratio"
```

```
# manually edit names of plasticity columns that were log-transformed
dput(plasts)
```

```
## c("plasticity_rdpi_DOY_Stage2_2023", "plasticity_rdpi_DOY_Stage3_2023", 
## "plasticity_rdpi_DOY_Stage6_2023", "plasticity_rdpi_DOY_Stage7_2023", 
## "plasticity_rdpi_DOY_last_budset_2023", "plasticity_rdpi_growing_season_days_2023", 
## "plasticity_rdpi_licor_gsw", "plasticity_rdpi_licor_gbw", "plasticity_rdpi_licor_ETR", 
## "plasticity_rdpi_licor_PhiPS2", "plasticity_rdpi_licor_Fs", "plasticity_rdpi_licor_Fm.", 
## "plasticity_rdpi_leaf_thickness_avg_mm_2023", "plasticity_rdpi_LMA_g_m2_2023", 
## "plasticity_rdpi_leaf_mass_g_2023", "plasticity_rdpi_leaf_area_cm2_2023", 
## "plasticity_rdpi_lower_stomata_pore_length_mean_um", "plasticity_rdpi_upper_stomata_presence", 
## "plasticity_rdpi_upper_stomata_pore_length_mean_um", "plasticity_rdpi_upper_stomata_density_mm2", 
## "plasticity_rdpi_lower_stomata_density_mm2", "plasticity_rdpi_stomata_ratio", 
## "plasticity_rdpi_DOY_Stage2_2024", "plasticity_rdpi_DOY_Stage3_2024", 
## "plasticity_rdpi_DOY_Stage6_2024", "plasticity_rdpi_DOY_Stage7_2024", 
## "plasticity_rdpi_DOY_last_budset_2024", "plasticity_rdpi_growing_season_days_2024"
## )
```

```
plasts <- c("plasticity_rdpi_DOY_Stage2_2023", "plasticity_rdpi_DOY_Stage3_2023", 
"plasticity_rdpi_DOY_Stage6_2023", "plasticity_rdpi_DOY_Stage7_2023", 
 "plasticity_rdpi_DOY_last_budset_2023", 
"plasticity_rdpi_growing_season_days_2023", "plasticity_rdpi_licor_gsw", 
"plasticity_rdpi_licor_gbw", "plasticity_rdpi_licor_ETR", "plasticity_rdpi_licor_PhiPS2", 
"plasticity_rdpi_licor_Fs", "plasticity_rdpi_licor_Fm.", "plasticity_rdpi_leaf_thickness_avg_mm_2023", 
"plasticity_rdpi_LMA_g_m2_2023", "plasticity_rdpi_leaf_mass_g_2023_log", 
"plasticity_rdpi_leaf_area_cm2_2023_log", "plasticity_rdpi_lower_stomata_pore_length_mean_um", 
"plasticity_rdpi_upper_stomata_presence", "plasticity_rdpi_upper_stomata_pore_length_mean_um", 
"plasticity_rdpi_upper_stomata_density_mm2_log", "plasticity_rdpi_lower_stomata_density_mm2", 
"plasticity_rdpi_stomata_ratio_log", "plasticity_rdpi_DOY_Stage2_2024", 
"plasticity_rdpi_DOY_Stage3_2024", "plasticity_rdpi_DOY_Stage6_2024", 
"plasticity_rdpi_DOY_Stage7_2024",  
"plasticity_rdpi_DOY_last_budset_2024", "plasticity_rdpi_growing_season_days_2024"
)

cbind(traits, plasts)
```

```
##       traits                             
##  [1,] "DOY_Stage2_2023"                  
##  [2,] "DOY_Stage3_2023"                  
##  [3,] "DOY_Stage6_2023"                  
##  [4,] "DOY_Stage7_2023"                  
##  [5,] "DOY_last_budset_2023"             
##  [6,] "growing_season_days_2023"         
##  [7,] "licor_gsw"                        
##  [8,] "licor_gbw"                        
##  [9,] "licor_ETR"                        
## [10,] "licor_PhiPS2"                     
## [11,] "licor_Fs"                         
## [12,] "licor_Fm."                        
## [13,] "leaf_thickness_avg_mm_2023"       
## [14,] "LMA_g_m2_2023"                    
## [15,] "leaf_mass_g_2023"                 
## [16,] "leaf_area_cm2_2023"               
## [17,] "lower_stomata_pore_length_mean_um"
## [18,] "upper_stomata_presence"           
## [19,] "upper_stomata_pore_length_mean_um"
## [20,] "upper_stomata_density_mm2"        
## [21,] "lower_stomata_density_mm2"        
## [22,] "stomata_ratio"                    
## [23,] "DOY_Stage2_2024"                  
## [24,] "DOY_Stage3_2024"                  
## [25,] "DOY_Stage6_2024"                  
## [26,] "DOY_Stage7_2024"                  
## [27,] "DOY_last_budset_2024"             
## [28,] "growing_season_days_2024"         
##       plasts                                             
##  [1,] "plasticity_rdpi_DOY_Stage2_2023"                  
##  [2,] "plasticity_rdpi_DOY_Stage3_2023"                  
##  [3,] "plasticity_rdpi_DOY_Stage6_2023"                  
##  [4,] "plasticity_rdpi_DOY_Stage7_2023"                  
##  [5,] "plasticity_rdpi_DOY_last_budset_2023"             
##  [6,] "plasticity_rdpi_growing_season_days_2023"         
##  [7,] "plasticity_rdpi_licor_gsw"                        
##  [8,] "plasticity_rdpi_licor_gbw"                        
##  [9,] "plasticity_rdpi_licor_ETR"                        
## [10,] "plasticity_rdpi_licor_PhiPS2"                     
## [11,] "plasticity_rdpi_licor_Fs"                         
## [12,] "plasticity_rdpi_licor_Fm."                        
## [13,] "plasticity_rdpi_leaf_thickness_avg_mm_2023"       
## [14,] "plasticity_rdpi_LMA_g_m2_2023"                    
## [15,] "plasticity_rdpi_leaf_mass_g_2023_log"             
## [16,] "plasticity_rdpi_leaf_area_cm2_2023_log"           
## [17,] "plasticity_rdpi_lower_stomata_pore_length_mean_um"
## [18,] "plasticity_rdpi_upper_stomata_presence"           
## [19,] "plasticity_rdpi_upper_stomata_pore_length_mean_um"
## [20,] "plasticity_rdpi_upper_stomata_density_mm2_log"    
## [21,] "plasticity_rdpi_lower_stomata_density_mm2"        
## [22,] "plasticity_rdpi_stomata_ratio_log"                
## [23,] "plasticity_rdpi_DOY_Stage2_2024"                  
## [24,] "plasticity_rdpi_DOY_Stage3_2024"                  
## [25,] "plasticity_rdpi_DOY_Stage6_2024"                  
## [26,] "plasticity_rdpi_DOY_Stage7_2024"                  
## [27,] "plasticity_rdpi_DOY_last_budset_2024"             
## [28,] "plasticity_rdpi_growing_season_days_2024"
```

```
################################################
# Do trait values predict trait plasticity, suggesting trade-offs?
# models
################################################
# reorder gardens by MAT
tmp <- aggregate(dat$garden_MAT_2023, by = list(dat$MiniCG_Site_2023), mean)
ord <- tmp$Group.1[order(tmp$x, decreasing = T)]
dat$MiniCG_Site_2023 <- factor(dat$MiniCG_Site_2023, levels = ord)

sites <- levels(dat$MiniCG_Site_2023)
# remove UCM - height not measured
sites <- sites[! sites == 'UCM']

sites.mat <- dat$garden_MAT_2023[match(sites, dat$MiniCG_Site)]


# to save results
mods.mat <- list()

# list to save summary plot for each trait with significant effects on fitness
plots.summ <- list()

for(n in 1:length(traits)){
  
  trait <- traits[n]
  plast <- plasts[n]
  
  print(paste('Model for:', trait, 'and', plast))
  
  # mode;
  mod <- lmer(paste0(trait, '~', plast,  ' + (1 | MiniCG_Site/block) + (1 | Genotype)'),
              data = dat
  )
  
  print(car::Anova(mod))
  
  p1 <- plot_model(mod, vline.color = "black", show.values = T, type = 'std', 
                   title = paste('2023 Growth ~ ', trait, sep = ''))
  plot(p1)
  #plot_model(mod, type = 're')
  
  mods.mat[[n]] <- mod
  names(mods.mat)[n] <- trait
  
  
  #####################
  # posthoc tests
  # if trait significantly predicts plasticity in full model, test in each garden
  mod.info <- Anova(mod)
  # if any coefficients are significant, plot by garden
  # need to change indexes if model is changed!
  if(mod.info$`Pr(>Chisq)`[1] <= 0.05){
    
  # plot data for all sites
  p2 <- ggplot(data = dat, aes_string(y = plast, x = trait, color = 'k2_tricho'), na.rm  = TRUE) +
    geom_point(na.rm = TRUE) +
    scale_color_gradient2(high = "darkolivegreen2", mid = "grey20", low = "dodgerblue2", midpoint = 0.5, name = 'P. trichocarpa\nancestry', guide = 'none') +
    geom_smooth(color = 'black', lty=1, method = 'lm', formula = y ~ x, na.rm = TRUE) +
    xlab(trait_names_df[trait, "trait_names"]) +
    ylab( paste('Plasticity (RDPI) of', trait_names_df[trait, "trait_names"])) +
    theme(axis.text.x = element_text(angle = 90, vjust = 0.5, hjust=1, size = 12)) #+
  #facet_wrap(~ MiniCG_Site_2023, drop = T, scales = 'free')
  
  plot(p2)
  
   p3 <- ggplot(data = dat, aes_string(y = plast, x = trait, color = 'k2_tricho'), na.rm  = TRUE) +
    geom_point(na.rm = TRUE) +
    scale_color_gradient2(high = "darkolivegreen2", mid = "grey20", low = "dodgerblue2", midpoint = 0.5, name = 'P. trichocarpa\nancestry', guide = 'none') +
    geom_smooth(aes(group = MiniCG_Site_2023), lty=1, method = 'lm', formula = y ~ x, na.rm = TRUE) +
    xlab(trait_names_df[trait, "trait_names"]) +
    ylab( paste('Plasticity (RDPI) of', trait_names_df[trait, "trait_names"])) +
    theme(axis.text.x = element_text(angle = 90, vjust = 0.5, hjust=1, size = 12))
    
# list to save plots for each site
    plots.sub <- list()
    
    
    # dataframe to save outputs from posthoc tests
    df.posthoc <- as.data.frame(matrix(ncol = 4, nrow = length(sites)))
    rownames(df.posthoc) <- sites
    colnames(df.posthoc) <- c('mat', 'slope', 'cor', 'pval')
    df.posthoc$mat <- sites.mat
    
    for(s in 1:length(sites)){
      
      site <- sites[s]
      dat.sub <- subset(dat, MiniCG_Site_2023 == site)
      # skip if site is missing data
      if(sum(!is.na(dat.sub[,trait])) > 10){
        mod.sub <- lmer(paste0(trait, '~', plast,  ' + (1 | block) + (1 | Genotype)'),
              data = dat.sub
  )
        # get pval for trait
        mod.sub.info <- Anova(mod.sub)
        df.posthoc[site, 'pval'] <- mod.sub.info$`Pr(>Chisq)`[1]
        
        # get slopes from fixed effect of trait (effect of trait value on plasticity)
        df.posthoc[site, 'slope'] <- fixef(mod.sub)[2]
        
        # get standardized effect, equivalent to correlation
        eff.std <- effectsize::standardize_parameters(mod.sub)
        df.posthoc[site, 'cor'] <- eff.std[2, 'Std_Coefficient'] # 2nd row has coefficient for plasticity effect
        
        # plot
        p <- ggplot(data = dat.sub, aes_string(y = plast, x = trait, color = 'k2_tricho'), na.rm  = TRUE) +
          geom_point(na.rm = TRUE) +
          scale_color_gradient2(high = "darkolivegreen2", mid = "grey20", low = "dodgerblue2", midpoint = 0.5, name = 'P. trichocarpa\nancestry', guide = 'none') +
          ggtitle(paste(site, '\n', trait, '\n', 'p = ', round(df.posthoc$pval[s], 3), sep = '')) +
          geom_smooth(color = 'black', lty = ifelse(df.posthoc$pval[s] <= 0.05, 1,2), method = 'lm', formula = y ~x, na.rm = TRUE) +
          theme(axis.text.x = element_text(angle = 90, vjust = 0.5, hjust=1, size = 12)) +
          theme(axis.title.x = element_blank())
        
        #plot(p)
        plots.sub[[s]] <- p
        
      }
    }

    # pvals are adjusted for summary plot
    df.posthoc$pvals.adj <- p.adjust(df.posthoc$pval, method = 'BH') 
    
    # plot panel with relationship for each garden
    # remove empty plots
    plots.sub <- plots.sub[lengths(plots.sub)>0]
    # plot in grid
    p1 <- grid.arrange(grobs = plots.sub, ncol = 4)
    #plot(p1)
    
    # setup text showing significance
    sig_text <- paste0('Trait ', pval_stars(mod.info[1, 'Pr(>Chisq)']))
    
    # wrap title
    maintitle <- trait_names_df[trait, 'trait_names']
    maintitle <- paste(strwrap(maintitle, width = 26), collapse = '\n')
    
    # plot slope of fitness relationship vs climate
    p2 <- ggplot(df.posthoc, aes(x = mat, y = cor), na.rm  = TRUE) +
      geom_point(size = 5, stroke = 2, shape = ifelse(df.posthoc$pvals.adj <= 0.05, 16, 1), aes(col = mat), na.rm = TRUE) +
      scale_color_gradient2(high = "red2", mid = "grey", low = "blue2", midpoint = mean(range(sites.mat)), limits = range_mat, name = 'Garden MAT (°C)') +
      geom_smooth(method = 'lm', col = 'black', na.rm = TRUE) +
      geom_hline(yintercept = 0, lty = 2) +
      ggtitle(maintitle,
              subtitle = sig_text) +
      xlab('Garden MAT') +
      ylab('Correlation between Trait and Plasticity') +
      geom_text_repel(label = rownames(df.posthoc), aes(x = mat, y = cor), box.padding = 0.5, min.segment.length = 1, size = 4) +
        theme(plot.title = element_text(size = 15),
        plot.subtitle = element_text(size = 10),
        legend.position = 'none')
    plot(p2)
    
    plots.summ[[n]] <- p2
    names(plots.summ)[n] <- trait
    
    # add legend
    leg_plot <- ggplot(df.posthoc, aes(x = mat, y = slope), na.rm  = TRUE) +
      geom_point(aes(color = mat))+
      scale_color_gradient2(high = "red2", mid = "grey", low = "blue2", midpoint = mean(range(sites.mat)), limits = range_mat, name = 'Garden MAT (°C)') +
      theme(legend.position = 'right')
    
    legend <- cowplot::get_legend(leg_plot)
    
    
  }
}
```

```
## [1] "Model for: DOY_Stage2_2023 and plasticity_rdpi_DOY_Stage2_2023"
## Analysis of Deviance Table (Type II Wald chisquare tests)
## 
## Response: DOY_Stage2_2023
##                                  Chisq Df Pr(>Chisq)
## plasticity_rdpi_DOY_Stage2_2023 1.0063  1     0.3158
```

```
## [1] "Model for: DOY_Stage3_2023 and plasticity_rdpi_DOY_Stage3_2023"
## Analysis of Deviance Table (Type II Wald chisquare tests)
## 
## Response: DOY_Stage3_2023
##                                  Chisq Df Pr(>Chisq)
## plasticity_rdpi_DOY_Stage3_2023 0.3111  1      0.577
```

```
## [1] "Model for: DOY_Stage6_2023 and plasticity_rdpi_DOY_Stage6_2023"
## Analysis of Deviance Table (Type II Wald chisquare tests)
## 
## Response: DOY_Stage6_2023
##                                  Chisq Df Pr(>Chisq)    
## plasticity_rdpi_DOY_Stage6_2023 11.363  1  0.0007492 ***
## ---
## Signif. codes:  0 '***' 0.001 '**' 0.01 '*' 0.05 '.' 0.1 ' ' 1
```

```
## boundary (singular) fit: see help('isSingular')
## boundary (singular) fit: see help('isSingular')
## boundary (singular) fit: see help('isSingular')
## boundary (singular) fit: see help('isSingular')
## boundary (singular) fit: see help('isSingular')
## boundary (singular) fit: see help('isSingular')
## boundary (singular) fit: see help('isSingular')
## boundary (singular) fit: see help('isSingular')
## boundary (singular) fit: see help('isSingular')
## boundary (singular) fit: see help('isSingular')
## boundary (singular) fit: see help('isSingular')
## boundary (singular) fit: see help('isSingular')
## boundary (singular) fit: see help('isSingular')
## boundary (singular) fit: see help('isSingular')
```

```
## `geom_smooth()` using formula = 'y ~ x'
```

```
## Warning: Removed 1 row containing missing values or values outside the scale range
## (`geom_text_repel()`).
```

```
## Warning: Removed 1 row containing missing values or values outside the scale range
## (`geom_point()`).
```

```
## [1] "Model for: DOY_Stage7_2023 and plasticity_rdpi_DOY_Stage7_2023"
```

```
## boundary (singular) fit: see help('isSingular')
```

```
## Analysis of Deviance Table (Type II Wald chisquare tests)
## 
## Response: DOY_Stage7_2023
##                                 Chisq Df Pr(>Chisq)
## plasticity_rdpi_DOY_Stage7_2023 1.561  1     0.2115
```

```
## boundary (singular) fit: see help('isSingular')
```

```
## [1] "Model for: DOY_last_budset_2023 and plasticity_rdpi_DOY_last_budset_2023"
## Analysis of Deviance Table (Type II Wald chisquare tests)
## 
## Response: DOY_last_budset_2023
##                                       Chisq Df Pr(>Chisq)
## plasticity_rdpi_DOY_last_budset_2023 0.0098  1     0.9212
```

```
## [1] "Model for: growing_season_days_2023 and plasticity_rdpi_growing_season_days_2023"
## Analysis of Deviance Table (Type II Wald chisquare tests)
## 
## Response: growing_season_days_2023
##                                           Chisq Df Pr(>Chisq)
## plasticity_rdpi_growing_season_days_2023 2.1809  1     0.1397
```

```
## [1] "Model for: licor_gsw and plasticity_rdpi_licor_gsw"
## Analysis of Deviance Table (Type II Wald chisquare tests)
## 
## Response: licor_gsw
##                            Chisq Df Pr(>Chisq)
## plasticity_rdpi_licor_gsw 0.6928  1     0.4052
```

```
## [1] "Model for: licor_gbw and plasticity_rdpi_licor_gbw"
```

```
## Warning: Some predictor variables are on very different scales: consider
## rescaling
```

```
## Analysis of Deviance Table (Type II Wald chisquare tests)
## 
## Response: licor_gbw
##                            Chisq Df Pr(>Chisq)
## plasticity_rdpi_licor_gbw 0.4151  1     0.5194
```

```
## [1] "Model for: licor_ETR and plasticity_rdpi_licor_ETR"
## Analysis of Deviance Table (Type II Wald chisquare tests)
## 
## Response: licor_ETR
##                            Chisq Df Pr(>Chisq)    
## plasticity_rdpi_licor_ETR 11.226  1  0.0008067 ***
## ---
## Signif. codes:  0 '***' 0.001 '**' 0.01 '*' 0.05 '.' 0.1 ' ' 1
```

```
## Warning in checkConv(attr(opt, "derivs"), opt$par, ctrl = control$checkConv, :
## Model failed to converge with max|grad| = 0.00334794 (tol = 0.002, component 1)
```

```
## Warning in checkConv(attr(opt, "derivs"), opt$par, ctrl = control$checkConv, :
## Model failed to converge with max|grad| = 0.00334793 (tol = 0.002, component 1)
```

```
## boundary (singular) fit: see help('isSingular')
## boundary (singular) fit: see help('isSingular')
```

```
## `geom_smooth()` using formula = 'y ~ x'
```

```
## Warning: Removed 3 rows containing missing values or values outside the scale range
## (`geom_text_repel()`).
```

```
## Warning: Removed 3 rows containing missing values or values outside the scale range
## (`geom_point()`).
```

```
## [1] "Model for: licor_PhiPS2 and plasticity_rdpi_licor_PhiPS2"
```

```
## boundary (singular) fit: see help('isSingular')
```

```
## Analysis of Deviance Table (Type II Wald chisquare tests)
## 
## Response: licor_PhiPS2
##                               Chisq Df Pr(>Chisq)    
## plasticity_rdpi_licor_PhiPS2 30.024  1  4.267e-08 ***
## ---
## Signif. codes:  0 '***' 0.001 '**' 0.01 '*' 0.05 '.' 0.1 ' ' 1
```

```
## boundary (singular) fit: see help('isSingular')
```

```
## boundary (singular) fit: see help('isSingular')
## boundary (singular) fit: see help('isSingular')
## boundary (singular) fit: see help('isSingular')
## boundary (singular) fit: see help('isSingular')
## boundary (singular) fit: see help('isSingular')
## boundary (singular) fit: see help('isSingular')
## boundary (singular) fit: see help('isSingular')
## boundary (singular) fit: see help('isSingular')
## boundary (singular) fit: see help('isSingular')
## boundary (singular) fit: see help('isSingular')
```

```
## `geom_smooth()` using formula = 'y ~ x'
```

```
## Warning: Removed 3 rows containing missing values or values outside the scale range
## (`geom_text_repel()`).
## Removed 3 rows containing missing values or values outside the scale range
## (`geom_point()`).
```

```
## [1] "Model for: licor_Fs and plasticity_rdpi_licor_Fs"
```

```
## boundary (singular) fit: see help('isSingular')
```

```
## Analysis of Deviance Table (Type II Wald chisquare tests)
## 
## Response: licor_Fs
##                           Chisq Df Pr(>Chisq)
## plasticity_rdpi_licor_Fs 2.3595  1     0.1245
```

```
## boundary (singular) fit: see help('isSingular')
```

```
## [1] "Model for: licor_Fm. and plasticity_rdpi_licor_Fm."
```

```
## boundary (singular) fit: see help('isSingular')
```

```
## Analysis of Deviance Table (Type II Wald chisquare tests)
## 
## Response: licor_Fm.
##                            Chisq Df Pr(>Chisq)    
## plasticity_rdpi_licor_Fm. 17.783  1  2.476e-05 ***
## ---
## Signif. codes:  0 '***' 0.001 '**' 0.01 '*' 0.05 '.' 0.1 ' ' 1
```

```
## boundary (singular) fit: see help('isSingular')
```

```
## boundary (singular) fit: see help('isSingular')
## boundary (singular) fit: see help('isSingular')
## boundary (singular) fit: see help('isSingular')
## boundary (singular) fit: see help('isSingular')
## boundary (singular) fit: see help('isSingular')
## boundary (singular) fit: see help('isSingular')
## boundary (singular) fit: see help('isSingular')
## boundary (singular) fit: see help('isSingular')
```

```
## `geom_smooth()` using formula = 'y ~ x'
```

```
## Warning: Removed 3 rows containing missing values or values outside the scale range
## (`geom_text_repel()`).
## Removed 3 rows containing missing values or values outside the scale range
## (`geom_point()`).
```

```
## [1] "Model for: leaf_thickness_avg_mm_2023 and plasticity_rdpi_leaf_thickness_avg_mm_2023"
## Analysis of Deviance Table (Type II Wald chisquare tests)
## 
## Response: leaf_thickness_avg_mm_2023
##                                            Chisq Df Pr(>Chisq)
## plasticity_rdpi_leaf_thickness_avg_mm_2023 1.197  1     0.2739
```

```
## [1] "Model for: LMA_g_m2_2023 and plasticity_rdpi_LMA_g_m2_2023"
## Analysis of Deviance Table (Type II Wald chisquare tests)
## 
## Response: LMA_g_m2_2023
##                                Chisq Df Pr(>Chisq)   
## plasticity_rdpi_LMA_g_m2_2023 9.2645  1   0.002336 **
## ---
## Signif. codes:  0 '***' 0.001 '**' 0.01 '*' 0.05 '.' 0.1 ' ' 1
```

```
## boundary (singular) fit: see help('isSingular')
## boundary (singular) fit: see help('isSingular')
## boundary (singular) fit: see help('isSingular')
## boundary (singular) fit: see help('isSingular')
## boundary (singular) fit: see help('isSingular')
## boundary (singular) fit: see help('isSingular')
## boundary (singular) fit: see help('isSingular')
## boundary (singular) fit: see help('isSingular')
## boundary (singular) fit: see help('isSingular')
## boundary (singular) fit: see help('isSingular')
## boundary (singular) fit: see help('isSingular')
## boundary (singular) fit: see help('isSingular')
```

```
## `geom_smooth()` using formula = 'y ~ x'
```

```
## Warning: Removed 1 row containing missing values or values outside the scale range
## (`geom_text_repel()`).
```

```
## Warning: Removed 1 row containing missing values or values outside the scale range
## (`geom_point()`).
```

```
## [1] "Model for: leaf_mass_g_2023 and plasticity_rdpi_leaf_mass_g_2023_log"
## Analysis of Deviance Table (Type II Wald chisquare tests)
## 
## Response: leaf_mass_g_2023
##                                       Chisq Df Pr(>Chisq)   
## plasticity_rdpi_leaf_mass_g_2023_log 9.5504  1   0.001999 **
## ---
## Signif. codes:  0 '***' 0.001 '**' 0.01 '*' 0.05 '.' 0.1 ' ' 1
```

```
## boundary (singular) fit: see help('isSingular')
## boundary (singular) fit: see help('isSingular')
## boundary (singular) fit: see help('isSingular')
## boundary (singular) fit: see help('isSingular')
## boundary (singular) fit: see help('isSingular')
## boundary (singular) fit: see help('isSingular')
## boundary (singular) fit: see help('isSingular')
## boundary (singular) fit: see help('isSingular')
## boundary (singular) fit: see help('isSingular')
## boundary (singular) fit: see help('isSingular')
## boundary (singular) fit: see help('isSingular')
## boundary (singular) fit: see help('isSingular')
## boundary (singular) fit: see help('isSingular')
## boundary (singular) fit: see help('isSingular')
```

```
## `geom_smooth()` using formula = 'y ~ x'
```

```
## Warning: Removed 1 row containing missing values or values outside the scale range
## (`geom_text_repel()`).
## Removed 1 row containing missing values or values outside the scale range
## (`geom_point()`).
```

```
## [1] "Model for: leaf_area_cm2_2023 and plasticity_rdpi_leaf_area_cm2_2023_log"
```

```
## Warning in checkConv(attr(opt, "derivs"), opt$par, ctrl = control$checkConv, :
## Model failed to converge with max|grad| = 0.00294689 (tol = 0.002, component 1)
```

```
## Analysis of Deviance Table (Type II Wald chisquare tests)
## 
## Response: leaf_area_cm2_2023
##                                         Chisq Df Pr(>Chisq)
## plasticity_rdpi_leaf_area_cm2_2023_log 0.0163  1     0.8984
```

```
## Warning in checkConv(attr(opt, "derivs"), opt$par, ctrl = control$checkConv, :
## Model failed to converge with max|grad| = 0.00294689 (tol = 0.002, component 1)
```

```
## [1] "Model for: lower_stomata_pore_length_mean_um and plasticity_rdpi_lower_stomata_pore_length_mean_um"
## Analysis of Deviance Table (Type II Wald chisquare tests)
## 
## Response: lower_stomata_pore_length_mean_um
##                                                    Chisq Df Pr(>Chisq)
## plasticity_rdpi_lower_stomata_pore_length_mean_um 0.9888  1       0.32
```

```
## [1] "Model for: upper_stomata_presence and plasticity_rdpi_upper_stomata_presence"
## Analysis of Deviance Table (Type II Wald chisquare tests)
## 
## Response: upper_stomata_presence
##                                         Chisq Df Pr(>Chisq)    
## plasticity_rdpi_upper_stomata_presence 49.505  1  1.979e-12 ***
## ---
## Signif. codes:  0 '***' 0.001 '**' 0.01 '*' 0.05 '.' 0.1 ' ' 1
```

```
## boundary (singular) fit: see help('isSingular')
## boundary (singular) fit: see help('isSingular')
## boundary (singular) fit: see help('isSingular')
## boundary (singular) fit: see help('isSingular')
## boundary (singular) fit: see help('isSingular')
## boundary (singular) fit: see help('isSingular')
## boundary (singular) fit: see help('isSingular')
## boundary (singular) fit: see help('isSingular')
## boundary (singular) fit: see help('isSingular')
## boundary (singular) fit: see help('isSingular')
## boundary (singular) fit: see help('isSingular')
## boundary (singular) fit: see help('isSingular')
## boundary (singular) fit: see help('isSingular')
## boundary (singular) fit: see help('isSingular')
## boundary (singular) fit: see help('isSingular')
## boundary (singular) fit: see help('isSingular')
## boundary (singular) fit: see help('isSingular')
## boundary (singular) fit: see help('isSingular')
```

```
## `geom_smooth()` using formula = 'y ~ x'
```

```
## Warning: Removed 1 row containing missing values or values outside the scale range
## (`geom_text_repel()`).
## Removed 1 row containing missing values or values outside the scale range
## (`geom_point()`).
```

```
## [1] "Model for: upper_stomata_pore_length_mean_um and plasticity_rdpi_upper_stomata_pore_length_mean_um"
## Analysis of Deviance Table (Type II Wald chisquare tests)
## 
## Response: upper_stomata_pore_length_mean_um
##                                                    Chisq Df Pr(>Chisq)
## plasticity_rdpi_upper_stomata_pore_length_mean_um 1.5631  1     0.2112
```

```
## [1] "Model for: upper_stomata_density_mm2 and plasticity_rdpi_upper_stomata_density_mm2_log"
## Analysis of Deviance Table (Type II Wald chisquare tests)
## 
## Response: upper_stomata_density_mm2
##                                                Chisq Df Pr(>Chisq)
## plasticity_rdpi_upper_stomata_density_mm2_log 0.2951  1      0.587
```

```
## [1] "Model for: lower_stomata_density_mm2 and plasticity_rdpi_lower_stomata_density_mm2"
## Analysis of Deviance Table (Type II Wald chisquare tests)
## 
## Response: lower_stomata_density_mm2
##                                            Chisq Df Pr(>Chisq)
## plasticity_rdpi_lower_stomata_density_mm2 0.3974  1     0.5284
```

```
## [1] "Model for: stomata_ratio and plasticity_rdpi_stomata_ratio_log"
```

```
## boundary (singular) fit: see help('isSingular')
```

```
## Analysis of Deviance Table (Type II Wald chisquare tests)
## 
## Response: stomata_ratio
##                                    Chisq Df Pr(>Chisq)   
## plasticity_rdpi_stomata_ratio_log 7.7892  1   0.005256 **
## ---
## Signif. codes:  0 '***' 0.001 '**' 0.01 '*' 0.05 '.' 0.1 ' ' 1
```

```
## boundary (singular) fit: see help('isSingular')
```

```
## boundary (singular) fit: see help('isSingular')
## boundary (singular) fit: see help('isSingular')
## boundary (singular) fit: see help('isSingular')
## boundary (singular) fit: see help('isSingular')
## boundary (singular) fit: see help('isSingular')
## boundary (singular) fit: see help('isSingular')
## boundary (singular) fit: see help('isSingular')
## boundary (singular) fit: see help('isSingular')
## boundary (singular) fit: see help('isSingular')
## boundary (singular) fit: see help('isSingular')
## boundary (singular) fit: see help('isSingular')
## boundary (singular) fit: see help('isSingular')
## boundary (singular) fit: see help('isSingular')
## boundary (singular) fit: see help('isSingular')
```

```
## `geom_smooth()` using formula = 'y ~ x'
```

```
## Warning: Removed 1 row containing missing values or values outside the scale range
## (`geom_text_repel()`).
## Removed 1 row containing missing values or values outside the scale range
## (`geom_point()`).
```

```
## [1] "Model for: DOY_Stage2_2024 and plasticity_rdpi_DOY_Stage2_2024"
## Analysis of Deviance Table (Type II Wald chisquare tests)
## 
## Response: DOY_Stage2_2024
##                                 Chisq Df Pr(>Chisq)
## plasticity_rdpi_DOY_Stage2_2024 1.076  1     0.2996
```

```
## [1] "Model for: DOY_Stage3_2024 and plasticity_rdpi_DOY_Stage3_2024"
```

```
## boundary (singular) fit: see help('isSingular')
```

```
## Analysis of Deviance Table (Type II Wald chisquare tests)
## 
## Response: DOY_Stage3_2024
##                                  Chisq Df Pr(>Chisq)
## plasticity_rdpi_DOY_Stage3_2024 0.0754  1     0.7836
```

```
## boundary (singular) fit: see help('isSingular')
```

```
## [1] "Model for: DOY_Stage6_2024 and plasticity_rdpi_DOY_Stage6_2024"
## Analysis of Deviance Table (Type II Wald chisquare tests)
## 
## Response: DOY_Stage6_2024
##                                 Chisq Df Pr(>Chisq)
## plasticity_rdpi_DOY_Stage6_2024 0.392  1     0.5313
```

```
## [1] "Model for: DOY_Stage7_2024 and plasticity_rdpi_DOY_Stage7_2024"
```

```
## boundary (singular) fit: see help('isSingular')
```

```
## Analysis of Deviance Table (Type II Wald chisquare tests)
## 
## Response: DOY_Stage7_2024
##                                  Chisq Df Pr(>Chisq)
## plasticity_rdpi_DOY_Stage7_2024 2.4744  1     0.1157
```

```
## boundary (singular) fit: see help('isSingular')
```

```
## [1] "Model for: DOY_last_budset_2024 and plasticity_rdpi_DOY_last_budset_2024"
```

```
## boundary (singular) fit: see help('isSingular')
```

```
## Analysis of Deviance Table (Type II Wald chisquare tests)
## 
## Response: DOY_last_budset_2024
##                                       Chisq Df Pr(>Chisq)
## plasticity_rdpi_DOY_last_budset_2024 2.5143  1     0.1128
```

```
## boundary (singular) fit: see help('isSingular')
```

```
## [1] "Model for: growing_season_days_2024 and plasticity_rdpi_growing_season_days_2024"
```

```
## boundary (singular) fit: see help('isSingular')
```

```
## Analysis of Deviance Table (Type II Wald chisquare tests)
## 
## Response: growing_season_days_2024
##                                           Chisq Df Pr(>Chisq)   
## plasticity_rdpi_growing_season_days_2024 8.3645  1   0.003826 **
## ---
## Signif. codes:  0 '***' 0.001 '**' 0.01 '*' 0.05 '.' 0.1 ' ' 1
```

```
## boundary (singular) fit: see help('isSingular')
```

```
## boundary (singular) fit: see help('isSingular')
## boundary (singular) fit: see help('isSingular')
## boundary (singular) fit: see help('isSingular')
## boundary (singular) fit: see help('isSingular')
## boundary (singular) fit: see help('isSingular')
## boundary (singular) fit: see help('isSingular')
## boundary (singular) fit: see help('isSingular')
## boundary (singular) fit: see help('isSingular')
## boundary (singular) fit: see help('isSingular')
## boundary (singular) fit: see help('isSingular')
```

```
## `geom_smooth()` using formula = 'y ~ x'
```

```
## Warning: Removed 5 rows containing missing values or values outside the scale range
## (`geom_text_repel()`).
```

```
## Warning: Removed 5 rows containing missing values or values outside the scale range
## (`geom_point()`).
```

```
plots.summ <- plots.summ[lengths(plots.summ)>0]

    
# add legend to beginning of plot list

plots2 <- plots.summ
plots2[[length(plots.summ)+1]] <- legend

p3 <- grid.arrange(grobs = plots2, ncol = 4)
```

```
## `geom_smooth()` using formula = 'y ~ x'
```

```
## Warning: Removed 1 row containing missing values or values outside the scale range
## (`geom_text_repel()`).
```

```
## `geom_smooth()` using formula = 'y ~ x'
```

```
## Warning: Removed 3 rows containing missing values or values outside the scale range
## (`geom_text_repel()`).
```

```
## `geom_smooth()` using formula = 'y ~ x'
```

```
## Warning: Removed 3 rows containing missing values or values outside the scale range
## (`geom_text_repel()`).
```

```
## `geom_smooth()` using formula = 'y ~ x'
```

```
## Warning: Removed 3 rows containing missing values or values outside the scale range
## (`geom_text_repel()`).
```

```
## `geom_smooth()` using formula = 'y ~ x'
```

```
## Warning: Removed 1 row containing missing values or values outside the scale range
## (`geom_text_repel()`).
```

```
## `geom_smooth()` using formula = 'y ~ x'
```

```
## Warning: Removed 1 row containing missing values or values outside the scale range
## (`geom_text_repel()`).
```

```
## `geom_smooth()` using formula = 'y ~ x'
```

```
## Warning: Removed 1 row containing missing values or values outside the scale range
## (`geom_text_repel()`).
```

```
## `geom_smooth()` using formula = 'y ~ x'
```

```
## Warning: Removed 1 row containing missing values or values outside the scale range
## (`geom_text_repel()`).
```

```
## `geom_smooth()` using formula = 'y ~ x'
```

```
## Warning: Removed 5 rows containing missing values or values outside the scale range
## (`geom_text_repel()`).
```

```
plot(p3)

#ggsave(file = 'results/fitness/fitnessEffects_summaryPlot_traitPlasticityTradeoffMAT2023.png', p3, height = 12, width = 14)
#ggsave(file = 'results/fitness/fitnessEffects_summaryPlot_traitPlasticityTradeoffMAT_2023.pdf', p3, height = 12, width = 14)

# rename plots list for later use
plots.plast.mat.23 <- plots.summ
```
