## Supplementary material for "Phenotypic plasticity evolved for climate variability constrains performance under climate warming": Analysis Rmarkdown Files: variance_partitioning_GxE.html

GxE Variance Paritioning


### GxE Variance Paritioning

###### Alayna Mead

#### 2026-03-11

- 1 Setup
- 2 Statistics - GxE variance
  partitioning
- 3 Table with results
- 4 Nice plots

Determine the proportion of variance in a trait that is explained by
genotype (G), environment (E), genotype x environment (GxE), block
nested within garden, and the residuals.

### 1 Setup

```
library(ggplot2)
library(lme4)
```

```
## Loading required package: Matrix
```

```
library(dplyr)
```

```
## 
## Attaching package: 'dplyr'
```

```
## The following objects are masked from 'package:stats':
## 
##     filter, lag
```

```
## The following objects are masked from 'package:base':
## 
##     intersect, setdiff, setequal, union
```

```
library(tidyr)
```

```
## 
## Attaching package: 'tidyr'
```

```
## The following objects are masked from 'package:Matrix':
## 
##     expand, pack, unpack
```

```
library(viridis) # color palettes
```

```
## Loading required package: viridisLite
```

```
library(knitr) # kable() for tables

# load data
load('data/clean/mini_garden_phenotypic_and_climate_data_2021-2024.Rdata')

# set negative heights to NA
dat$GrowthIncrement_2023[dat$GrowthIncrement_2024 < 0] <- NA
dat$GrowthIncrement_2023[dat$GrowthIncrement_2023 < 0] <- NA
dat$GrowthIncrement_2022[dat$GrowthIncrement_2022 < 0] <- NA
dat$GrowthIncrement_2021[dat$GrowthIncrement_2021 < 0] <- NA

# remove NA genotypes and those without genetic info
dat <- dat[! is.na(dat$Genotype),]
dat <- dat[! is.na(dat$k2_tricho),]

# print session info, including package versions
sessionInfo()
```

```
## R version 4.5.2 (2025-10-31)
## Platform: x86_64-pc-linux-gnu
## Running under: Arch Linux
## 
## Matrix products: default
## BLAS:   /usr/lib/libblas.so.3.12.0 
## LAPACK: /usr/lib/liblapack.so.3.12.0  LAPACK version 3.12.0
## 
## locale:
##  [1] LC_CTYPE=en_US.UTF-8       LC_NUMERIC=C              
##  [3] LC_TIME=en_US.UTF-8        LC_COLLATE=en_US.UTF-8    
##  [5] LC_MONETARY=en_US.UTF-8    LC_MESSAGES=en_US.UTF-8   
##  [7] LC_PAPER=en_US.UTF-8       LC_NAME=C                 
##  [9] LC_ADDRESS=C               LC_TELEPHONE=C            
## [11] LC_MEASUREMENT=en_US.UTF-8 LC_IDENTIFICATION=C       
## 
## time zone: US/Eastern
## tzcode source: system (glibc)
## 
## attached base packages:
## [1] stats     graphics  grDevices datasets  utils     methods   base     
## 
## other attached packages:
## [1] knitr_1.50        viridis_0.6.5     viridisLite_0.4.2 tidyr_1.3.1      
## [5] dplyr_1.1.4       lme4_1.1-37       Matrix_1.7-4      ggplot2_3.5.2    
## 
## loaded via a namespace (and not attached):
##  [1] gtable_0.3.6      jsonlite_2.0.0    compiler_4.5.2    renv_0.17.3      
##  [5] Rcpp_1.0.14       tidyselect_1.2.1  gridExtra_2.3     jquerylib_0.1.4  
##  [9] splines_4.5.2     scales_1.3.0      boot_1.3-32       yaml_2.3.10      
## [13] fastmap_1.2.0     lattice_0.22-7    R6_2.6.1          generics_0.1.4   
## [17] rbibutils_2.3     MASS_7.3-65       tibble_3.2.1      nloptr_2.2.1     
## [21] munsell_0.5.1     minqa_1.2.8       bslib_0.9.0       pillar_1.10.2    
## [25] rlang_1.1.6       cachem_1.1.0      xfun_0.52         sass_0.4.10      
## [29] cli_3.6.5         withr_3.0.2       magrittr_2.0.3    Rdpack_2.6.4     
## [33] digest_0.6.37     grid_4.5.2        rstudioapi_0.17.1 lifecycle_1.0.4  
## [37] nlme_3.1-168      reformulas_0.4.0  vctrs_0.6.5       evaluate_1.0.3   
## [41] glue_1.8.0        colorspace_2.1-1  purrr_1.0.4       rmarkdown_2.29   
## [45] tools_4.5.2       pkgconfig_2.0.3   htmltools_0.5.8.1
```

### 2 Statistics - GxE variance partitioning

Get hte proportion of variance explained by factors modelled as
random effects using lmer (most traits) or glmer (traits with binary
measures).

```
# the list of traits to use

traits <- c("GrowthIncrement_2021", "GrowthIncrement_2022", "GrowthIncrement_2023", "GrowthIncrement_2024", # growth
            "DOY_Stage2_2021", "DOY_Stage3_2021", "DOY_Stage6_2021", "growing_season_days_2021", "Stage2_cGDD_2021", "Stage3_cGDD_2021", # 2021 phenology
            "DOY_Stage2_2022", "DOY_Stage3_2022", "DOY_Stage6_2022", "Stage2_cGDD_2022", "Stage3_cGDD_2022", "growing_season_days_2022", # 2022 phenology
            "DOY_Stage2_2023", "DOY_Stage3_2023", "Stage2_cGDD_2023", "Stage3_cGDD_2023", "DOY_Stage6_2023", "DOY_Stage7_2023", "stage7_presence_2023", "DOY_Stage8_2023", "DOY_last_budset_2023", "growing_season_days_2023", # 2023 phenology
            "DOY_Stage2_2024", "DOY_Stage3_2024", "Stage2_cGDD_2024", 
"Stage3_cGDD_2024", "DOY_Stage6_2024", "DOY_Stage7_2024",  "stage7_presence_2024", "DOY_Stage8_2024", "DOY_last_budset_2024", "growing_season_days_2024",
            "leaf_thickness_avg_mm_2023", "leaf_area_cm2_2023", "leaf_mass_g_2023", "LMA_g_m2_2023", # leaf morphology
"lower_stomata_pore_length_mean_um", "upper_stomata_pore_length_mean_um", "lower_stomata_density_mm2", "upper_stomata_density_mm2", "stomata_ratio", "upper_stomata_presence", "licor_gsw", # stomata 
             "licor_gbw", "licor_ETR", "licor_Fs", "licor_Fm.", "licor_PhiPS2" # photosynthesis
)

# presence/absence traits get modeled differently
traits_bin <- c("stage7_presence_2023", "stage7_presence_2024", "upper_stomata_presence")


# categorize traits for plotting
traits_group <- c(rep('Growth', 4), rep('Phenology', 32), rep('Leaf\nMorphology', 4), rep('Stomata', 7), rep('Photosynthesis', 5))
cbind(traits, traits_group)
```

```
##       traits                              traits_group      
##  [1,] "GrowthIncrement_2021"              "Growth"          
##  [2,] "GrowthIncrement_2022"              "Growth"          
##  [3,] "GrowthIncrement_2023"              "Growth"          
##  [4,] "GrowthIncrement_2024"              "Growth"          
##  [5,] "DOY_Stage2_2021"                   "Phenology"       
##  [6,] "DOY_Stage3_2021"                   "Phenology"       
##  [7,] "DOY_Stage6_2021"                   "Phenology"       
##  [8,] "growing_season_days_2021"          "Phenology"       
##  [9,] "Stage2_cGDD_2021"                  "Phenology"       
## [10,] "Stage3_cGDD_2021"                  "Phenology"       
## [11,] "DOY_Stage2_2022"                   "Phenology"       
## [12,] "DOY_Stage3_2022"                   "Phenology"       
## [13,] "DOY_Stage6_2022"                   "Phenology"       
## [14,] "Stage2_cGDD_2022"                  "Phenology"       
## [15,] "Stage3_cGDD_2022"                  "Phenology"       
## [16,] "growing_season_days_2022"          "Phenology"       
## [17,] "DOY_Stage2_2023"                   "Phenology"       
## [18,] "DOY_Stage3_2023"                   "Phenology"       
## [19,] "Stage2_cGDD_2023"                  "Phenology"       
## [20,] "Stage3_cGDD_2023"                  "Phenology"       
## [21,] "DOY_Stage6_2023"                   "Phenology"       
## [22,] "DOY_Stage7_2023"                   "Phenology"       
## [23,] "stage7_presence_2023"              "Phenology"       
## [24,] "DOY_Stage8_2023"                   "Phenology"       
## [25,] "DOY_last_budset_2023"              "Phenology"       
## [26,] "growing_season_days_2023"          "Phenology"       
## [27,] "DOY_Stage2_2024"                   "Phenology"       
## [28,] "DOY_Stage3_2024"                   "Phenology"       
## [29,] "Stage2_cGDD_2024"                  "Phenology"       
## [30,] "Stage3_cGDD_2024"                  "Phenology"       
## [31,] "DOY_Stage6_2024"                   "Phenology"       
## [32,] "DOY_Stage7_2024"                   "Phenology"       
## [33,] "stage7_presence_2024"              "Phenology"       
## [34,] "DOY_Stage8_2024"                   "Phenology"       
## [35,] "DOY_last_budset_2024"              "Phenology"       
## [36,] "growing_season_days_2024"          "Phenology"       
## [37,] "leaf_thickness_avg_mm_2023"        "Leaf\nMorphology"
## [38,] "leaf_area_cm2_2023"                "Leaf\nMorphology"
## [39,] "leaf_mass_g_2023"                  "Leaf\nMorphology"
## [40,] "LMA_g_m2_2023"                     "Leaf\nMorphology"
## [41,] "lower_stomata_pore_length_mean_um" "Stomata"         
## [42,] "upper_stomata_pore_length_mean_um" "Stomata"         
## [43,] "lower_stomata_density_mm2"         "Stomata"         
## [44,] "upper_stomata_density_mm2"         "Stomata"         
## [45,] "stomata_ratio"                     "Stomata"         
## [46,] "upper_stomata_presence"            "Stomata"         
## [47,] "licor_gsw"                         "Stomata"         
## [48,] "licor_gbw"                         "Photosynthesis"  
## [49,] "licor_ETR"                         "Photosynthesis"  
## [50,] "licor_Fs"                          "Photosynthesis"  
## [51,] "licor_Fm."                         "Photosynthesis"  
## [52,] "licor_PhiPS2"                      "Photosynthesis"
```

```
# set up dataframes to save results
df <- dat[,traits]
df$genotype <- as.character(dat$Genotype)
df$garden <- as.character(dat$MiniCG_Site)
df$block <- as.character(dat$block)

str(df)
```

```
## 'data.frame':    1510 obs. of  55 variables:
##  $ GrowthIncrement_2021             : num  30.8 37.7 30.7 34.7 25.6 19.6 24.1 24.1 20.1 29.1 ...
##  $ GrowthIncrement_2022             : num  NA 10.67 8.23 39.62 77.72 ...
##  $ GrowthIncrement_2023             : num  0 NA 22.6 NA NA ...
##  $ GrowthIncrement_2024             : num  6.1 -21.3 42.7 -6.1 -12.2 ...
##  $ DOY_Stage2_2021                  : num  92 88 88 88 85 85 95 92 88 88 ...
##  $ DOY_Stage3_2021                  : num  103 99 103 103 103 103 107 107 95 103 ...
##  $ DOY_Stage6_2021                  : num  200 264 200 264 283 266 190 283 190 200 ...
##  $ growing_season_days_2021         : num  108 176 112 176 198 181 95 191 102 112 ...
##  $ Stage2_cGDD_2021                 : num  488 462 462 462 441 ...
##  $ Stage3_cGDD_2021                 : num  554 534 554 554 554 ...
##  $ DOY_Stage2_2022                  : num  87 87 87 87 87 87 95 95 95 90 ...
##  $ DOY_Stage3_2022                  : num  95 95 95 98 95 95 98 98 98 98 ...
##  $ DOY_Stage6_2022                  : num  188 188 207 207 225 207 207 207 188 188 ...
##  $ Stage2_cGDD_2022                 : int  494 494 494 494 494 494 559 559 559 522 ...
##  $ Stage3_cGDD_2022                 : int  559 559 559 584 559 559 584 584 584 584 ...
##  $ growing_season_days_2022         : num  101 101 120 120 138 120 112 112 93 98 ...
##  $ DOY_Stage2_2023                  : num  97 101 93 101 93 93 101 101 93 97 ...
##  $ DOY_Stage3_2023                  : num  109 111 109 109 109 109 117 114 101 109 ...
##  $ Stage2_cGDD_2023                 : num  496 531 472 531 472 ...
##  $ Stage3_cGDD_2023                 : num  582 590 582 582 582 ...
##  $ DOY_Stage6_2023                  : num  181 163 181 163 162 NA 187 163 162 163 ...
##  $ DOY_Stage7_2023                  : num  NA 201 187 NA 194 194 NA NA NA NA ...
##  $ stage7_presence_2023             : num  0 1 1 0 1 1 0 0 0 0 ...
##  $ DOY_Stage8_2023                  : num  NA 215 201 NA 240 NA 225 NA NA NA ...
##  $ DOY_last_budset_2023             : num  181 215 201 163 240 NA 225 163 162 163 ...
##  $ growing_season_days_2023         : num  84 114 108 62 147 NA 124 62 69 66 ...
##  $ DOY_Stage2_2024                  : num  75 95 92 92 86 86 92 92 92 92 ...
##  $ DOY_Stage3_2024                  : num  92 99 95 95 92 95 99 95 95 95 ...
##  $ Stage2_cGDD_2024                 : num  418 614 584 584 531 ...
##  $ Stage3_cGDD_2024                 : num  584 643 614 614 584 ...
##  $ DOY_Stage6_2024                  : num  178 176 165 165 153 NA 165 178 165 165 ...
##  $ DOY_Stage7_2024                  : num  NA 193 178 NA 165 176 NA 193 NA NA ...
##  $ stage7_presence_2024             : num  0 1 1 0 1 1 0 1 0 0 ...
##  $ DOY_Stage8_2024                  : num  242 243 242 243 242 243 242 243 242 243 ...
##  $ DOY_last_budset_2024             : num  242 243 242 243 242 243 242 243 242 243 ...
##  $ growing_season_days_2024         : num  167 148 150 151 156 157 150 151 150 151 ...
##  $ leaf_thickness_avg_mm_2023       : num  0.212 0.25 0.224 0.234 0.232 0.314 0.302 0.226 0.198 0.212 ...
##  $ leaf_area_cm2_2023               : num  NA NA NA NA NA NA NA NA NA NA ...
##  $ leaf_mass_g_2023                 : num  NA NA NA NA NA NA NA NA NA NA ...
##  $ LMA_g_m2_2023                    : num  NA NA NA NA NA NA NA NA NA NA ...
##  $ lower_stomata_pore_length_mean_um: num  NA NA NA NA NA NA NA NA NA NA ...
##  $ upper_stomata_pore_length_mean_um: num  NA NA NA NA NA NA NA NA NA NA ...
##  $ lower_stomata_density_mm2        : num  NA NA NA NA NA NA NA NA NA NA ...
##  $ upper_stomata_density_mm2        : num  NA NA NA NA NA NA NA NA NA NA ...
##  $ stomata_ratio                    : num  NA NA NA NA NA NA NA NA NA NA ...
##  $ upper_stomata_presence           : num  NA NA NA NA NA NA NA NA NA NA ...
##  $ licor_gsw                        : num  NA NA NA NA NA NA NA NA NA NA ...
##  $ licor_gbw                        : num  NA NA NA NA NA NA NA NA NA NA ...
##  $ licor_ETR                        : num  NA NA NA NA NA NA NA NA NA NA ...
##  $ licor_Fs                         : num  NA NA NA NA NA NA NA NA NA NA ...
##  $ licor_Fm.                        : num  NA NA NA NA NA NA NA NA NA NA ...
##  $ licor_PhiPS2                     : num  NA NA NA NA NA NA NA NA NA NA ...
##  $ genotype                         : chr  "206" "206" "210" "210" ...
##  $ garden                           : chr  "EVERGREEN" "EVERGREEN" "EVERGREEN" "EVERGREEN" ...
##  $ block                            : chr  "1" "2" "1" "2" ...
```

```
# variance explained df
explained <- as.data.frame(matrix(nrow = length(traits), ncol = 5))
colnames(explained) <- c('Genotype x Environment', 'Genotype', 'Block', 'Environment',  'Residual')
rownames(explained) <- traits


# loop through and fit model to each trait

for(n in 1:length(traits)){
  
  trait <- traits[n]
  print(trait)
  
  # setup formula
  formula <- as.formula(paste(trait, "~ 1 + (1|genotype) + (1|garden/block) + (1|genotype:garden)"))
  
  # presence/absence traits have to be run separately
  # use a glmer with binomial link
  if(trait %in% traits_bin){
    
  # model with a binomial link function
   mod <- glmer(formula,
          data = df,
          family = binomial)
   
    vcs <- as.data.frame(VarCorr(mod))
    
    # calculate proportion variance explained
    # the sum and residuals are (pi^2)/3, see eg:
    # https://stats.stackexchange.com/questions/128750/residual-variance-for-glmer
    vcs_sum <- sum(vcs$vcov) + ((pi^2) / 3)
    explained[n,'Genotype x Environment'] <- (vcs[vcs$grp == 'genotype:garden', 'vcov']/vcs_sum)*100
    explained[n,'Genotype'] <- (vcs[vcs$grp == 'genotype', 'vcov']/vcs_sum)*100
    explained[n,'Environment'] <- (vcs[vcs$grp == 'garden', 'vcov']/vcs_sum)*100
    explained[n, 'Residual'] <- (((pi^2) / 3)/vcs_sum)*100
    explained[n,'Block'] <- (vcs[vcs$grp == 'block:garden', 'vcov']/vcs_sum)*100
    
  }else{
 
  # run the model
  mod <- lmer(formula, data = df)
  
  # save variances
  # need to make sure column names in explained match groups in vcs
  vcs <- as.data.frame(VarCorr(mod))
  explained[n,] <- vcs$vcov
    
  }
}
```

```
## [1] "GrowthIncrement_2021"
## [1] "GrowthIncrement_2022"
## [1] "GrowthIncrement_2023"
## [1] "GrowthIncrement_2024"
## [1] "DOY_Stage2_2021"
## [1] "DOY_Stage3_2021"
## [1] "DOY_Stage6_2021"
## [1] "growing_season_days_2021"
## [1] "Stage2_cGDD_2021"
## [1] "Stage3_cGDD_2021"
## [1] "DOY_Stage2_2022"
## [1] "DOY_Stage3_2022"
## [1] "DOY_Stage6_2022"
## [1] "Stage2_cGDD_2022"
```

```
## boundary (singular) fit: see help('isSingular')
```

```
## [1] "Stage3_cGDD_2022"
## [1] "growing_season_days_2022"
## [1] "DOY_Stage2_2023"
## [1] "DOY_Stage3_2023"
## [1] "Stage2_cGDD_2023"
## [1] "Stage3_cGDD_2023"
## [1] "DOY_Stage6_2023"
## [1] "DOY_Stage7_2023"
```

```
## boundary (singular) fit: see help('isSingular')
```

```
## [1] "stage7_presence_2023"
## [1] "DOY_Stage8_2023"
## [1] "DOY_last_budset_2023"
## [1] "growing_season_days_2023"
## [1] "DOY_Stage2_2024"
## [1] "DOY_Stage3_2024"
```

```
## boundary (singular) fit: see help('isSingular')
```

```
## [1] "Stage2_cGDD_2024"
## [1] "Stage3_cGDD_2024"
```

```
## boundary (singular) fit: see help('isSingular')
```

```
## [1] "DOY_Stage6_2024"
## [1] "DOY_Stage7_2024"
```

```
## boundary (singular) fit: see help('isSingular')
```

```
## [1] "stage7_presence_2024"
```

```
## boundary (singular) fit: see help('isSingular')
```

```
## [1] "DOY_Stage8_2024"
```

```
## boundary (singular) fit: see help('isSingular')
```

```
## [1] "DOY_last_budset_2024"
```

```
## boundary (singular) fit: see help('isSingular')
```

```
## [1] "growing_season_days_2024"
```

```
## boundary (singular) fit: see help('isSingular')
```

```
## [1] "leaf_thickness_avg_mm_2023"
## [1] "leaf_area_cm2_2023"
## [1] "leaf_mass_g_2023"
## [1] "LMA_g_m2_2023"
## [1] "lower_stomata_pore_length_mean_um"
## [1] "upper_stomata_pore_length_mean_um"
```

```
## boundary (singular) fit: see help('isSingular')
```

```
## [1] "lower_stomata_density_mm2"
## [1] "upper_stomata_density_mm2"
## [1] "stomata_ratio"
```

```
## boundary (singular) fit: see help('isSingular')
```

```
## [1] "upper_stomata_presence"
## [1] "licor_gsw"
## [1] "licor_gbw"
```

```
## boundary (singular) fit: see help('isSingular')
```

```
## [1] "licor_ETR"
## [1] "licor_Fs"
```

```
## boundary (singular) fit: see help('isSingular')
```

```
## [1] "licor_Fm."
```

```
## boundary (singular) fit: see help('isSingular')
```

```
## [1] "licor_PhiPS2"
```

```
# convert variance explained to a proportion of the total
explained.prop <- explained

for(n in 1:nrow(explained)){
  
  total <- sum(explained[n, c("Genotype x Environment", "Genotype", "Block", "Environment", "Residual")])
  
  prop <- explained[n, c("Genotype x Environment", "Genotype", "Block", "Environment", "Residual")]/total
  
  explained.prop[n, c("Genotype x Environment", "Genotype", "Block", "Environment", "Residual")] <- prop
  
}

sum(rownames(explained) != rownames(explained.prop))
```

```
## [1] 0
```

```
explained.prop$category <- explained$category

# plot as a stacked barplot

explained$trait <- row.names(explained)
explained$category <- traits_group
exp.long <- explained %>% mutate(trait = factor(trait)) %>% pivot_longer(cols = c('Genotype x Environment', 'Genotype', 'Environment', 'Residual', 'Block'))
exp.long$Effect <- factor(exp.long$name, levels = c('Residual', 'Block', 'Genotype x Environment', 'Genotype', 'Environment'))

ggplot(exp.long, aes(y = trait, x = value, fill = Effect)) +
  geom_bar(position = "fill", stat = "identity") +
  scale_fill_viridis_d(option = 'mako', direction = 1) +
  facet_grid(rows = vars(category), drop = T, scales = 'free_y', space = 'free_y') +
  xlab('Proportion Variance Explained') +
  ylab('Trait') +
  theme_set(theme_bw(base_size = 10))
```

```
#ggsave(file = 'results/plasticity/GxE_variance_explained_allTraits_allYears_withBlock.png', height = 10, width = 8)
```

### 3 Table with results

```
kable(explained, label = 'Raw variance explained')
```

|  | Genotype x Environment | Genotype | Block | Environment | Residual | trait | category |
| --- | --- | --- | --- | --- | --- | --- | --- |
| GrowthIncrement\_2021 | 72.7231532 | 133.1535890 | 12.3986760 | 2.821680e+02 | 3.099649e+02 | GrowthIncrement\_2021 | Growth |
| GrowthIncrement\_2022 | 68.4404928 | 152.8001931 | 2.3217447 | 4.050481e+02 | 4.133104e+02 | GrowthIncrement\_2022 | Growth |
| GrowthIncrement\_2023 | 199.4135752 | 150.2223443 | 16.4112062 | 5.502682e+02 | 6.432617e+02 | GrowthIncrement\_2023 | Growth |
| GrowthIncrement\_2024 | 42.0346670 | 73.5425513 | 54.4416537 | 4.981371e+01 | 7.281279e+02 | GrowthIncrement\_2024 | Growth |
| DOY\_Stage2\_2021 | 23.0389407 | 33.4698142 | 1.6950550 | 3.277383e+02 | 5.757598e+01 | DOY\_Stage2\_2021 | Phenology |
| DOY\_Stage3\_2021 | 14.4861927 | 28.2643584 | 0.6723750 | 2.904462e+02 | 9.863313e+01 | DOY\_Stage3\_2021 | Phenology |
| DOY\_Stage6\_2021 | 105.9931025 | 64.6307190 | 6.2226983 | 9.172884e+02 | 1.605088e+02 | DOY\_Stage6\_2021 | Phenology |
| growing\_season\_days\_2021 | 104.7715428 | 50.0284138 | 3.9130192 | 1.136927e+03 | 1.840510e+02 | growing\_season\_days\_2021 | Phenology |
| Stage2\_cGDD\_2021 | 3738.2273360 | 3114.9738463 | 402.8284968 | 6.629325e+04 | 8.428105e+03 | Stage2\_cGDD\_2021 | Phenology |
| Stage3\_cGDD\_2021 | 2637.3370296 | 3432.0223104 | 263.6367724 | 7.079791e+04 | 1.627887e+04 | Stage3\_cGDD\_2021 | Phenology |
| DOY\_Stage2\_2022 | 25.5100944 | 25.9630803 | 0.2327215 | 2.524839e+02 | 3.063128e+01 | DOY\_Stage2\_2022 | Phenology |
| DOY\_Stage3\_2022 | 23.5124271 | 24.5543607 | 0.6067540 | 2.349292e+02 | 3.946036e+01 | DOY\_Stage3\_2022 | Phenology |
| DOY\_Stage6\_2022 | 53.2108540 | 79.6164232 | 11.8110834 | 7.497629e+02 | 1.470707e+02 | DOY\_Stage6\_2022 | Phenology |
| Stage2\_cGDD\_2022 | 1643.2731131 | 2428.7083155 | 0.0000000 | 5.123039e+04 | 6.322939e+03 | Stage2\_cGDD\_2022 | Phenology |
| Stage3\_cGDD\_2022 | 2772.3257641 | 2538.5170975 | 259.8314340 | 6.034388e+04 | 7.213541e+03 | Stage3\_cGDD\_2022 | Phenology |
| growing\_season\_days\_2022 | 60.5706039 | 73.1736395 | 15.4179527 | 7.293230e+02 | 1.471073e+02 | growing\_season\_days\_2022 | Phenology |
| DOY\_Stage2\_2023 | 38.5440687 | 25.3083694 | 4.6841042 | 2.436103e+02 | 4.403450e+01 | DOY\_Stage2\_2023 | Phenology |
| DOY\_Stage3\_2023 | 32.5722633 | 27.1405676 | 4.4050664 | 2.073672e+02 | 5.458826e+01 | DOY\_Stage3\_2023 | Phenology |
| Stage2\_cGDD\_2023 | 5979.0497916 | 2561.5007946 | 860.9622946 | 4.859827e+04 | 7.806663e+03 | Stage2\_cGDD\_2023 | Phenology |
| Stage3\_cGDD\_2023 | 4794.2778093 | 3470.8154747 | 689.8954148 | 5.350062e+04 | 1.165397e+04 | Stage3\_cGDD\_2023 | Phenology |
| DOY\_Stage6\_2023 | 85.3760199 | 143.6529363 | 3.4311594 | 8.742704e+02 | 1.116987e+02 | DOY\_Stage6\_2023 | Phenology |
| DOY\_Stage7\_2023 | 357.5743820 | 0.0000000 | 0.0000000 | 4.424729e+02 | 1.684487e+02 | DOY\_Stage7\_2023 | Phenology |
| stage7\_presence\_2023 | 2.0967242 | 5.9019355 | 1.2546540 | 5.803732e+01 | 3.270937e+01 | stage7\_presence\_2023 | Phenology |
| DOY\_Stage8\_2023 | 171.7938852 | 19.5872272 | 0.0000106 | 3.044116e+02 | 7.317186e+02 | DOY\_Stage8\_2023 | Phenology |
| DOY\_last\_budset\_2023 | 129.1957628 | 103.1877538 | 11.5937981 | 6.499107e+02 | 6.083518e+02 | DOY\_last\_budset\_2023 | Phenology |
| growing\_season\_days\_2023 | 139.4628880 | 68.4786559 | 19.1222218 | 8.114974e+02 | 7.262632e+02 | growing\_season\_days\_2023 | Phenology |
| DOY\_Stage2\_2024 | 10.5731397 | 17.0936894 | 0.9336035 | 2.326590e+02 | 1.518456e+01 | DOY\_Stage2\_2024 | Phenology |
| DOY\_Stage3\_2024 | 0.3429855 | 15.2254321 | 0.0000000 | 1.937923e+02 | 6.081149e+01 | DOY\_Stage3\_2024 | Phenology |
| Stage2\_cGDD\_2024 | 671.9649426 | 1496.0924180 | 47.4208896 | 1.797899e+04 | 1.482448e+03 | Stage2\_cGDD\_2024 | Phenology |
| Stage3\_cGDD\_2024 | 0.0000000 | 1569.5125782 | 0.0000000 | 1.976480e+04 | 9.730394e+03 | Stage3\_cGDD\_2024 | Phenology |
| DOY\_Stage6\_2024 | 50.8841214 | 56.1175062 | 2.5952599 | 6.862421e+02 | 1.070112e+02 | DOY\_Stage6\_2024 | Phenology |
| DOY\_Stage7\_2024 | 76.4153891 | 0.0000000 | 0.0000000 | 7.307871e+02 | 1.563165e+02 | DOY\_Stage7\_2024 | Phenology |
| stage7\_presence\_2024 | 1.2917639 | 12.0924760 | 0.0000000 | 5.212448e+01 | 3.449128e+01 | stage7\_presence\_2024 | Phenology |
| DOY\_Stage8\_2024 | 152.1406709 | 52.8339072 | 0.0000000 | 4.361031e+02 | 3.315154e+02 | DOY\_Stage8\_2024 | Phenology |
| DOY\_last\_budset\_2024 | 120.3791166 | 93.4187833 | 0.0000000 | 6.140549e+02 | 4.127349e+02 | DOY\_last\_budset\_2024 | Phenology |
| growing\_season\_days\_2024 | 136.6342747 | 82.4409760 | 0.0000000 | 8.453748e+02 | 4.723982e+02 | growing\_season\_days\_2024 | Phenology |
| leaf\_thickness\_avg\_mm\_2023 | 0.0002633 | 0.0001173 | 0.0004433 | 1.754000e-04 | 1.501200e-03 | leaf\_thickness\_avg\_mm\_2023 | Leaf |
| Morphology |  |  |  |  |  |  |  |
| leaf\_area\_cm2\_2023 | 24.5471250 | 24.7246216 | 15.0301132 | 1.011113e+02 | 2.050319e+02 | leaf\_area\_cm2\_2023 | Leaf |
| Morphology |  |  |  |  |  |  |  |
| leaf\_mass\_g\_2023 | 0.0018880 | 0.0027980 | 0.0015441 | 1.028300e-02 | 2.466030e-02 | leaf\_mass\_g\_2023 | Leaf |
| Morphology |  |  |  |  |  |  |  |
| LMA\_g\_m2\_2023 | 0.0139404 | 0.0027978 | 0.0019167 | 8.444300e-03 | 2.792970e-02 | LMA\_g\_m2\_2023 | Leaf |
| Morphology |  |  |  |  |  |  |  |
| lower\_stomata\_pore\_length\_mean\_um | 1.0961793 | 2.4482165 | 0.0705351 | 4.161588e+00 | 7.904779e+00 | lower\_stomata\_pore\_length\_mean\_um | Stomata |
| upper\_stomata\_pore\_length\_mean\_um | 0.0000000 | 1.7908775 | 0.0458477 | 3.726627e+00 | 1.213422e+01 | upper\_stomata\_pore\_length\_mean\_um | Stomata |
| lower\_stomata\_density\_mm2 | 6.4723841 | 352.0045749 | 25.6206561 | 1.173995e+03 | 2.328184e+03 | lower\_stomata\_density\_mm2 | Stomata |
| upper\_stomata\_density\_mm2 | 10.6377613 | 149.2626629 | 5.1744161 | 5.507797e+01 | 4.239722e+02 | upper\_stomata\_density\_mm2 | Stomata |
| stomata\_ratio | 0.0025403 | 0.0074605 | 0.0000000 | 3.147500e-03 | 3.615330e-02 | stomata\_ratio | Stomata |
| upper\_stomata\_presence | 0.0000482 | 36.5851122 | 0.9191335 | 5.344188e+00 | 5.715152e+01 | upper\_stomata\_presence | Stomata |
| licor\_gsw | 0.0017004 | 0.0032747 | 0.0000947 | 2.712900e-03 | 1.031970e-02 | licor\_gsw | Stomata |
| licor\_gbw | 0.0000000 | 0.0000000 | 0.0000000 | 5.700000e-06 | 2.000000e-07 | licor\_gbw | Photosynthesis |
| licor\_ETR | 719.2760235 | 104.7323325 | 1101.0813764 | 2.401017e+03 | 2.570940e+03 | licor\_ETR | Photosynthesis |
| licor\_Fs | 110.8245471 | 0.0000000 | 74.2890521 | 1.506006e+02 | 6.904750e+02 | licor\_Fs | Photosynthesis |
| licor\_Fm. | 601.3806011 | 0.0000000 | 382.4396861 | 5.445138e+03 | 4.069906e+03 | licor\_Fm. | Photosynthesis |
| licor\_PhiPS2 | 0.0014394 | 0.0000426 | 0.0025714 | 1.819710e-02 | 1.245140e-02 | licor\_PhiPS2 | Photosynthesis |

```
kable(explained.prop, label = 'Proportion variance explained')
```

|  | Genotype x Environment | Genotype | Block | Environment | Residual |
| --- | --- | --- | --- | --- | --- |
| GrowthIncrement\_2021 | 0.0897364 | 0.1643043 | 0.0152993 | 0.3481801 | 0.3824799 |
| GrowthIncrement\_2022 | 0.0656868 | 0.1466524 | 0.0022283 | 0.3887513 | 0.3966811 |
| GrowthIncrement\_2023 | 0.1278639 | 0.0963225 | 0.0105229 | 0.3528317 | 0.4124591 |
| GrowthIncrement\_2024 | 0.0443422 | 0.0775798 | 0.0574303 | 0.0525483 | 0.7680994 |
| DOY\_Stage2\_2021 | 0.0519459 | 0.0754644 | 0.0038218 | 0.7389514 | 0.1298165 |
| DOY\_Stage3\_2021 | 0.0334939 | 0.0653508 | 0.0015546 | 0.6715484 | 0.2280523 |
| DOY\_Stage6\_2021 | 0.0844806 | 0.0515132 | 0.0049597 | 0.7311146 | 0.1279318 |
| growing\_season\_days\_2021 | 0.0708063 | 0.0338100 | 0.0026445 | 0.7683544 | 0.1243847 |
| Stage2\_cGDD\_2021 | 0.0456007 | 0.0379980 | 0.0049139 | 0.8086773 | 0.1028101 |
| Stage3\_cGDD\_2021 | 0.0282341 | 0.0367416 | 0.0028224 | 0.7579283 | 0.1742737 |
| DOY\_Stage2\_2022 | 0.0761902 | 0.0775431 | 0.0006951 | 0.7540861 | 0.0914855 |
| DOY\_Stage3\_2022 | 0.0727797 | 0.0760048 | 0.0018781 | 0.7271929 | 0.1221444 |
| DOY\_Stage6\_2022 | 0.0510920 | 0.0764461 | 0.0113408 | 0.7199070 | 0.1412142 |
| Stage2\_cGDD\_2022 | 0.0266656 | 0.0394109 | 0.0000000 | 0.8313206 | 0.1026030 |
| Stage3\_cGDD\_2022 | 0.0379105 | 0.0347133 | 0.0035531 | 0.8251805 | 0.0986425 |
| growing\_season\_days\_2022 | 0.0590591 | 0.0713477 | 0.0150332 | 0.7111235 | 0.1434364 |
| DOY\_Stage2\_2023 | 0.1082147 | 0.0710547 | 0.0131509 | 0.6839502 | 0.1236294 |
| DOY\_Stage3\_2023 | 0.0998924 | 0.0832345 | 0.0135094 | 0.6359526 | 0.1674110 |
| Stage2\_cGDD\_2023 | 0.0908581 | 0.0389248 | 0.0130833 | 0.7385032 | 0.1186307 |
| Stage3\_cGDD\_2023 | 0.0646917 | 0.0468336 | 0.0093091 | 0.7219124 | 0.1572532 |
| DOY\_Stage6\_2023 | 0.0700706 | 0.1179001 | 0.0028161 | 0.7175389 | 0.0916744 |
| DOY\_Stage7\_2023 | 0.3692058 | 0.0000000 | 0.0000000 | 0.4568660 | 0.1739282 |
| stage7\_presence\_2023 | 0.0209672 | 0.0590194 | 0.0125465 | 0.5803732 | 0.3270937 |
| DOY\_Stage8\_2023 | 0.1399530 | 0.0159569 | 0.0000000 | 0.2479909 | 0.5960993 |
| DOY\_last\_budset\_2023 | 0.0860021 | 0.0686893 | 0.0077177 | 0.4326278 | 0.4049632 |
| growing\_season\_days\_2023 | 0.0790237 | 0.0388020 | 0.0108352 | 0.4598176 | 0.4115215 |
| DOY\_Stage2\_2024 | 0.0382469 | 0.0618342 | 0.0033772 | 0.8416135 | 0.0549282 |
| DOY\_Stage3\_2024 | 0.0012695 | 0.0563545 | 0.0000000 | 0.7172918 | 0.2250842 |
| Stage2\_cGDD\_2024 | 0.0309991 | 0.0690178 | 0.0021876 | 0.8294072 | 0.0683883 |
| Stage3\_cGDD\_2024 | 0.0000000 | 0.0505240 | 0.0000000 | 0.6362462 | 0.3132299 |
| DOY\_Stage6\_2024 | 0.0563594 | 0.0621559 | 0.0028745 | 0.7600841 | 0.1185260 |
| DOY\_Stage7\_2024 | 0.0793086 | 0.0000000 | 0.0000000 | 0.7584563 | 0.1622350 |
| stage7\_presence\_2024 | 0.0129176 | 0.1209248 | 0.0000000 | 0.5212448 | 0.3449128 |
| DOY\_Stage8\_2024 | 0.1564279 | 0.0543227 | 0.0000000 | 0.4483922 | 0.3408572 |
| DOY\_last\_budset\_2024 | 0.0970339 | 0.0753020 | 0.0000000 | 0.4949710 | 0.3326930 |
| growing\_season\_days\_2024 | 0.0889055 | 0.0536429 | 0.0000000 | 0.5500704 | 0.3073812 |
| leaf\_thickness\_avg\_mm\_2023 | 0.1053042 | 0.0469009 | 0.1772887 | 0.0701593 | 0.6003469 |
| leaf\_area\_cm2\_2023 | 0.0662639 | 0.0667430 | 0.0405731 | 0.2729454 | 0.5534745 |
| leaf\_mass\_g\_2023 | 0.0458560 | 0.0679563 | 0.0375020 | 0.2497485 | 0.5989371 |
| LMA\_g\_m2\_2023 | 0.2533279 | 0.0508428 | 0.0348316 | 0.1534519 | 0.5075458 |
| lower\_stomata\_pore\_length\_mean\_um | 0.0699036 | 0.1561233 | 0.0044980 | 0.2653855 | 0.5040896 |
| upper\_stomata\_pore\_length\_mean\_um | 0.0000000 | 0.1011934 | 0.0025906 | 0.2105728 | 0.6856432 |
| lower\_stomata\_density\_mm2 | 0.0016654 | 0.0905763 | 0.0065926 | 0.3020874 | 0.5990783 |
| upper\_stomata\_density\_mm2 | 0.0165151 | 0.2317293 | 0.0080332 | 0.0855082 | 0.6582141 |
| stomata\_ratio | 0.0515259 | 0.1513237 | 0.0000000 | 0.0638418 | 0.7333086 |
| upper\_stomata\_presence | 0.0000005 | 0.3658511 | 0.0091913 | 0.0534419 | 0.5715152 |
| licor\_gsw | 0.0939339 | 0.1808974 | 0.0052299 | 0.1498625 | 0.5700763 |
| licor\_gbw | 0.0000000 | 0.0004510 | 0.0000802 | 0.9577733 | 0.0416955 |
| licor\_ETR | 0.1042875 | 0.0151851 | 0.1596453 | 0.3481225 | 0.3727595 |
| licor\_Fs | 0.1079962 | 0.0000000 | 0.0723931 | 0.1467572 | 0.6728535 |
| licor\_Fm. | 0.0572805 | 0.0000000 | 0.0364268 | 0.5186407 | 0.3876520 |
| licor\_PhiPS2 | 0.0414781 | 0.0012282 | 0.0740998 | 0.5243829 | 0.3588109 |

### 4 Nice plots

A nicer plot with trait categories and cleaned trait names

```
# simplify trait names, remove units
colnames(explained)
```

```
## [1] "Genotype x Environment" "Genotype"               "Block"                 
## [4] "Environment"            "Residual"               "trait"                 
## [7] "category"
```

```
explained$trait_names <- explained$trait
explained$trait_names <- gsub('Stage2_cGDD', 'Stage2_Bud_Flush_cGDD', explained$trait_names)
explained$trait_names <- gsub('DOY_Stage2', 'Stage2_Bud_Flush_DOY', explained$trait_names)
explained$trait_names <- gsub('DOY_Stage3', 'Stage3_Leaf_Emergence_DOY', explained$trait_names)
explained$trait_names <- gsub('DOY_Stage6', 'Stage6_First_Budset_DOY', explained$trait_names)
explained$trait_names <- gsub('DOY_Stage7', 'Stage7_Lammas_Growth_DOY', explained$trait_names)
explained$trait_names <- gsub('stage7_presence', 'Stage7_Lammas_Growth_presence', explained$trait_names)
explained$trait_names <- gsub('DOY_Stage8', 'Stage8_Budset_after_Lammas_DOY', explained$trait_names)
explained$trait_names <- gsub('DOY_last_budset', 'Stage8_Final_Budset_DOY', explained$trait_names)
explained$trait_names <- gsub('growing_season_days', 'Growing_Season_Days', explained$trait_names)
explained$trait_names <- gsub('licor_', '', explained$trait_names)
explained$trait_names <- gsub('Fm.', "Fm'", explained$trait_names)
explained$trait_names <- gsub('upper', "Adaxial", explained$trait_names)
explained$trait_names <- gsub('lower', "Abaxial", explained$trait_names)
explained$trait_names <- gsub('stomata_ratio', "Stomata_ratio", explained$trait_names)
explained$trait_names <- gsub('leaf', 'Leaf', explained$trait_names)

# remove units
explained$trait_names <- gsub('_cm2', '', explained$trait_names)
explained$trait_names <- gsub('_cm', '', explained$trait_names)
explained$trait_names <- gsub('_mm2', '', explained$trait_names)
explained$trait_names <- gsub('_m2', '', explained$trait_names)
explained$trait_names <- gsub('_mm', '', explained$trait_names)
explained$trait_names <- gsub('_um_', '', explained$trait_names)
explained$trait_names <- gsub('_g', '', explained$trait_names)


explained$trait_names <- gsub('_', ' ', explained$trait_names)

# reorder so they're alphabetical from top to bottom
explained$trait_names <- factor(explained$trait_names, levels = sort(explained$trait_names, decreasing = T))

# transform to long-form dataframe for ggplot
explained$trait <- row.names(explained)
explained$category <- traits_group
exp.long <- explained %>% mutate(trait = factor(trait)) %>% pivot_longer(cols = c('Genotype x Environment', 'Genotype', 'Environment', 'Residual', 'Block'))
exp.long$Effect <- factor(exp.long$name, levels = c('Residual', 'Block', 'Genotype x Environment', 'Genotype', 'Environment'))

# plot!
ggplot(exp.long, aes(y = trait_names, x = value, fill = Effect)) +
  geom_bar(position = "fill", stat = "identity") +
  scale_fill_viridis_d(option = 'mako', direction = 1) +
  facet_grid(rows = vars(category), drop = T, scales = 'free_y', space = 'free_y') +
  xlab('Proportion Variance Explained') +
  ylab('Trait') +
  theme_set(theme_bw(base_size = 10))
```

```
#ggsave(file = 'results/plasticity/GxE_variance_explained_allTraits_allYears.png', height = 10, width = 8)


#########################

# subset to only 2023 and 2024 measures
include <- c("GrowthIncrement_2023", "GrowthIncrement_2024","DOY_Stage2_2023", "DOY_Stage3_2023", "Stage2_cGDD_2023", "Stage3_cGDD_2023", "DOY_Stage6_2023", "DOY_Stage7_2023", "stage7_presence_2023", "DOY_Stage8_2023", "DOY_last_budset_2023", "growing_season_days_2023", "DOY_Stage2_2024", "DOY_Stage3_2024", "Stage2_cGDD_2024", "Stage3_cGDD_2024", "DOY_Stage6_2024", "DOY_Stage7_2024", "stage7_presence_2024", "DOY_Stage8_2024", "DOY_last_budset_2024", "growing_season_days_2024", "leaf_thickness_avg_mm_2023", "leaf_area_cm2_2023", "leaf_mass_g_2023", "LMA_g_m2_2023", "lower_stomata_pore_length_mean_um", "upper_stomata_pore_length_mean_um", "lower_stomata_density_mm2", "upper_stomata_density_mm2", "stomata_ratio", "upper_stomata_presence", "licor_gsw", "licor_gbw", "licor_ETR", "licor_Fs", "licor_Fm.", "licor_PhiPS2")
explained.sub <- explained[explained$trait %in% include,]


# reorder so they're alphabetical from top to bottom
# reversed because geom_bar plots them in reverse
explained.sub$trait_names <- factor(explained.sub$trait_names, levels = sort(explained.sub$trait_names, decreasing = F))

# transform to long-form dataframe for ggplot
exp.sub.long <- explained.sub %>% mutate(trait_names = factor(trait_names)) %>% pivot_longer(cols = c('Genotype x Environment', 'Genotype', 'Environment', 'Residual', 'Block'))
#exp.sub.long$Effect <- factor(exp.sub.long$name, levels =c('Residual', 'Block', 'Genotype x Environment', 'Genotype', 'Environment'))

exp.sub.long$Effect <- factor(exp.sub.long$name, levels =c("Genotype", "Environment", "Genotype x Environment", "Block", "Residual"))

exp.sub.long$category <- factor(exp.sub.long$category, levels = c('Growth', 'Phenology', 'Leaf\nMorphology', 'Stomata', 'Photosynthesis'))

# edit color palette
# I started with this palette: PNWColors::rev(pnw_palette('Starfish', 4))
# then edited colors to be a bit more distinguishable
cols <- c("#4760b8", "#015b58", "#380069", "#e69b99")

ggplot(exp.sub.long, aes(y = trait_names, x = value, fill = Effect)) +
  geom_bar(aes(fill = Effect), position = position_fill(reverse = TRUE), stat = "identity") +
  scale_fill_manual(values = c(cols, 'grey50')) +
  facet_grid(rows = vars(category), drop = T, scales = 'free_y', space = 'free_y') +
  xlab('Proportion Variance Explained') +
  ylab('Trait') +
  theme_set(theme_bw(base_size = 10))
```

```
#ggsave(file = 'results/plasticity/GxE_variance_explained_allTraits_2023-2024.png', height = 8, width = 8)

#ggsave(file = 'results/plasticity/GxE_variance_explained_allTraits_2023-2024.pdf', height = 8, width = 8)


##################
# Final plot
# subset to only 2023 and 2024 measures
# remove cGDD

include <- c("GrowthIncrement_2023", "GrowthIncrement_2024","DOY_Stage2_2023", "DOY_Stage3_2023","DOY_Stage6_2023", "DOY_Stage7_2023", "stage7_presence_2023", "DOY_last_budset_2023", "growing_season_days_2023", "DOY_Stage2_2024", "DOY_Stage3_2024", "DOY_Stage6_2024", "DOY_Stage7_2024", "stage7_presence_2024", "DOY_last_budset_2024", "growing_season_days_2024", "leaf_thickness_avg_mm_2023", "leaf_area_cm2_2023", "leaf_mass_g_2023", "LMA_g_m2_2023", "lower_stomata_pore_length_mean_um", "upper_stomata_pore_length_mean_um", "lower_stomata_density_mm2", "upper_stomata_density_mm2", "stomata_ratio", "upper_stomata_presence", "licor_gsw", "licor_gbw", "licor_ETR", "licor_Fs", "licor_Fm.", "licor_PhiPS2")
explained.sub <- explained[explained$trait %in% include,]

# reorder so they're alphabetical from top to bottom
explained.sub$trait_names <- factor(explained.sub$trait_names, levels = sort(explained.sub$trait_names, decreasing = T))

exp.sub.long <- explained.sub %>% mutate(trait_names = factor(trait_names)) %>% pivot_longer(cols = c('Genotype x Environment', 'Genotype', 'Environment', 'Residual', 'Block'))
exp.sub.long$Effect <- factor(exp.sub.long$name, levels =c('Residual', 'Block', 'Genotype x Environment', 'Genotype', 'Environment'))

# edit color palette
# I started with this palette: PNWColors::rev(pnw_palette('Starfish', 4))
# then edited colors to be a bit more distinguishable
cols <- c("#e69b99","#380069","#4760b8","#015b58")

ggplot(exp.sub.long, aes(y = trait_names, x = value, fill = Effect)) +
  geom_bar(position = "fill", stat = "identity") +
  scale_fill_manual(values = c('grey50', cols)) +
  facet_grid(rows = vars(category), drop = T, scales = 'free_y', space = 'free_y') +
  xlab('Proportion Variance Explained') +
  ylab('Trait') +
  theme_set(theme_bw(base_size = 10))
```

```
#ggsave(file = 'results/plasticity/GxE_variance_explained_withBlock_clean_2023-2024.png', height = 8, width = 8)
```
